# Luxar: Gaussian splatting for microscopy and scalable interactive web visualisation of multidimensional scientific data

**DOI:** 10.64898/2026.09.22.753676

**Authors:** Loïc A. Royer

## Abstract

Volumetric microscopy produces datasets too large to share as files and too rich to convey as still images, yet exploring them interactively means installing desktop software or building web infrastructure. Here we present Luxar, an open-source Python framework that compiles multidimensional data into compact archives that any modern browser renders from static file hosting. Fluorescence volumes are represented as mixtures of anisotropic Gaussians (Gaussian splats): the image becomes geometry that the GPU draws directly, with no voxel grid to rebuild. Splats share one scene with points, trajectories and surfaces. Blind-spot cross-validation sets the number of Gaussians without noise-free references. Across 17 volumes from 12 datasets and four modalities, this yields a median 99-fold (6–340-fold) size reduction at 26–67 dB peak signal-to-noise ratio (PSNR), in minutes per volume on one GPU. We demonstrate Luxar on a 500-timepoint embryogenesis recording, super-resolution localisations, singlecell atlases and lineage tracking, each explorable through a shareable link.

## Introduction

Modern scientific instruments generate multidimensional data at a pace that far outstrips our ability to share and explore it. A single light-sheet microscope run produces hundreds of gigabytes;^1, 2^ a single-cell atlas^3^ contains millions of cells embedded in high-dimensional spaces; a single embryo followed through development becomes a time-lapse of volumes that no still image can convey. Yet the dominant mode of dissemination remains downloading raw files or, more commonly, inspecting static projections that discard the very dimensionality that motivated the measurement.

Interactive exploration is no easier to share. Desktop tools such as napari,^4^ Fiji,^5^ Vaa3D,^6^ and BigDataViewer^7^ provide powerful visualisation and remain the natural home for analysis on the source voxels. BigDataViewer streams multi-resolution data from remote servers, and napari and Fiji can open remote OME-Zarr or N5 stores through plugins. Each requires a local installation, however, and remote access depends on specific formats and plugins; it is not the sharing path. Web-based viewers like Neuroglancer^8^ stream precomputed multiresolution voxel pyramids over HTTP and render them in the browser, with shared views of images, segmentations and annotations; they are organised around voxel slices and volume rendering of those pyramids, not around an explicit geometric representation that mixes data types, and a custom deployment is web-development work. Neither paradigm is designed for the full diversity of scientific data: volumetric microscopy, point-cloud embeddings, polyline trajectories, and higher-dimensional datasets all require different tools, formats, and expertise.

Here we present Luxar (Fig. 1; Supplementary Video 1), an open-source system that bridges the gap between data processing and analysis in Python and scalable interactive exploration in a web browser. Luxar supports four first-class geometry types (**points, lines, meshes**, and **Gaussian splats**, the last being how *n*D images are represented) within a hierarchical scene graph that handles arbitrary dimensions, physical units, and per-dimension transforms. Scientists work entirely in Python and NumPy: a handful of lines compiles data into a chunked Zarr^9^ archive laid out for lazy loading, indexed in space and along every non-displayed dimension and optionally carrying a level-of-detail grammar. From that archive a browser viewer (WebGL, or WebGPU) with HDR rendering and native *n*-dimensional navigation streams just the chunks, levels and timepoints the current view calls for (Fig. 1a,b). No knowledge of TypeScript, WebGL, or web development is required; users describe rich, large nD scenes in Python and can locally open the viewer in any modern browser, or share these scenes on the web.

**Figure 1.**
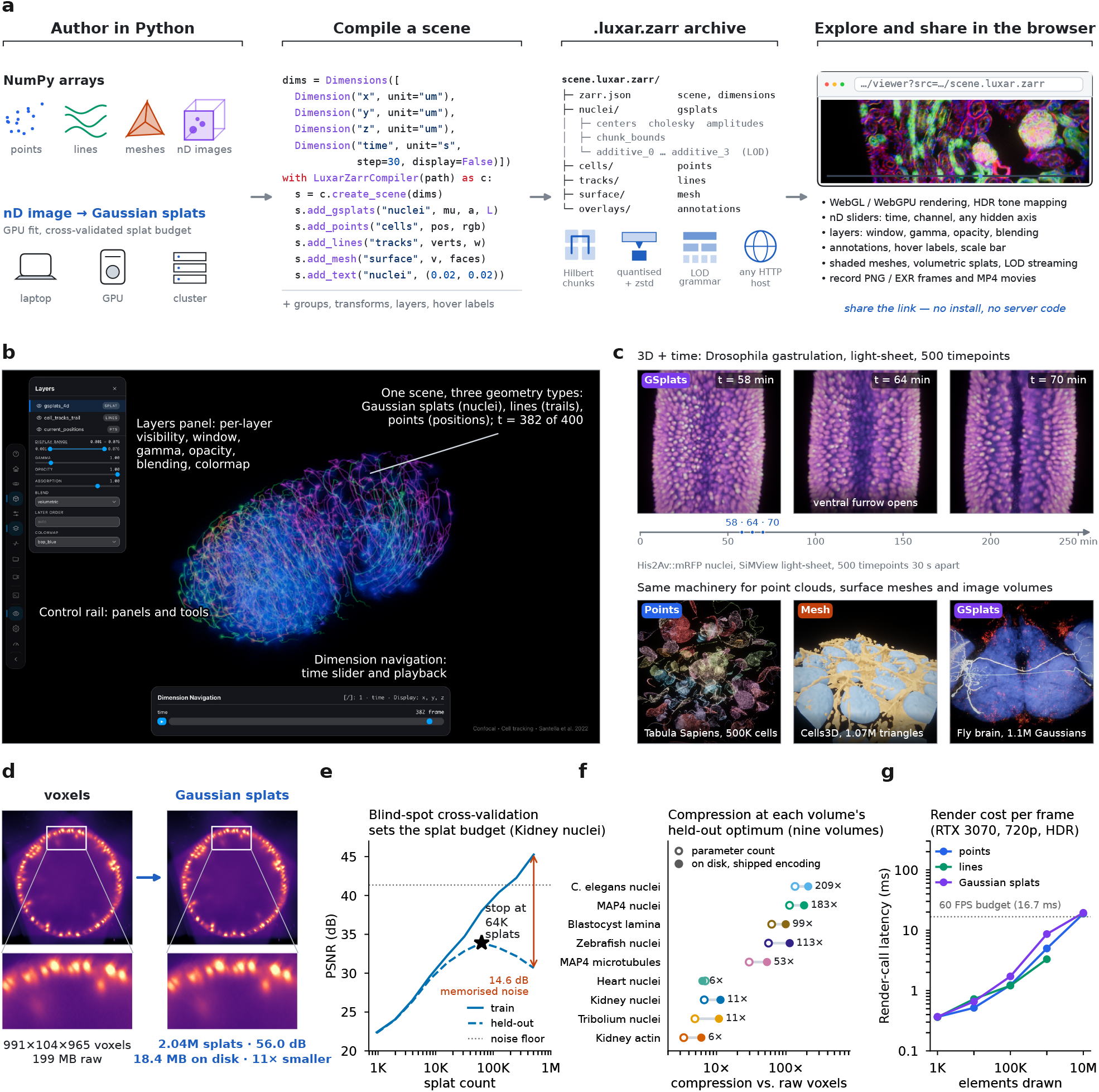
Luxar: Gaussian splats for microscopy volumes inside a Python-first, browser-rendered system for *n*D data. **a**, Authoring flow: NumPy arrays (points, lines, meshes, *n*D images) are described in a few lines of Python; *n*D images are first fitted to Gaussian splats (on a laptop, a GPU workstation or, tiled, a cluster), their splat budget set by blind-spot cross-validation (**e**). The scene graph is compiled once into a .`luxar.zarr` archive that any static HTTP host serves to the browser; sharing a scene is sharing a link. **b**, The viewer: a 4D *C. elegans* embryo^10^ at timepoint 382 of 400, with Gaussian splats (nuclei), fading-trail polylines and points in one scene (Supplementary Video 2). **c**, Top: gastrulation in a *Drosophila* embryo (His2Av::mRFP nuclei, SiMView light-sheet^11, 12^) at 58, 64 and 70 min (Supplementary Videos 1 and 3). Bottom: 500K Tabula Sapiens cells^13^ as points coloured by organ, isosurface meshes of the Cells3D volume, and 1.1 million Gaussian splats of a FlyLight MultiColor FlpOut *Drosophila* brain.^14^ **d**, One slice of the *Tribolium* light-sheet embryo^15^ (`uint16`, 199 MB as stored) and the same slice rendered from its fit at its operating point (2.04M splats after culling, 11 *×* smaller on disk, 56.0 dB PSNR); insets: the boxed region. **e**, On the Kidney nuclei volume, training PSNR keeps rising while held-out (blind-spot) PSNR peaks at 64K splats; beyond it the added splats memorise noise (train–held-out gap 14.6 dB at 512K). **f**, Compression at each volume’s held-out optimum for the nine volumes of the rate-distortion analysis (all 17: Supplementary Fig. 1), as a parameter count (open) and as bytes on disk with the shipped encoding (filled; Methods). **g**, Latency of a synchronised render-and-readback call versus elements drawn for points, lines and Gaussian splats (RTX 3070, 720p, HDR): within the 16.7 ms budget of 60 frames per second through 10^6^ elements of each. **b** and **c** are unretouched viewer frames; **d** shows data slices.

This Python-first, compile-then-view architecture enables applications across domains: protein embeddings and single-cell atlases as colour-coded point clouds with categorical dimension navigation, cell trajectories as polylines, segmented structures as shaded surface meshes, and *n*D microscopy images rendered as Gaussian splats (Fig. 1b,c). In this paper, we focus on the last of these, Gaussian splatting for microscopy volumes, which is our main novel contribution.

Fluorescence microscopy data is inherently sparse: signal is concentrated in structures (nuclei, membranes, filaments) while the vast majority of voxels are background. This sparsity has previously been exploited by content-adaptive representations such as the Adaptive Particle Representation,^16^ which adapts particle density to local image content for efficient storage and processing. Splatting, the rendering of a volume as a sum of projected Gaussian footprints, is a classical volume-rendering technique,^17, 18^ and radial-basis-function encodings of scalar fields have long been rendered interactively.^19^ Gaussian splatting^20, 21^ revived it as a *fitted* representation for real-time novel-view synthesis, an alternative to neural radiance fields:^22^ a scene is represented as a mixture of oriented, anisotropic Gaussian primitives optimised against the data, placing representational capacity only where signal exists. The resulting representation is continuous, compact and directly consumable by the GPU rasteriser: unlike voxel data, which is decoded chunk by chunk into grid textures at some resolution before it can be sampled, Gaussian splats are rasterised directly from their chunk-decoded parameters, with no intermediate voxel array. This makes them particularly well-suited for streaming, interactive visualisation in a web browser, where both bandwidth and client-side memory are constrained. We adapt nD Gaussian splatting for microscopy and validate the approach on 17 volumes from 12 datasets spanning spinning-disk, confocal, light-sheet and structured-illumination modalities. Using blind-spot cross-validation,^23, 24^ we show that the noise level sets an upper bound on useful model complexity, which provides a principled criterion for separating signal from noise. The Results describe the system, the fitting method and its accuracy across modalities, the cross-validated choice of splat count and a comparison with a video codec, fitting at scale by tiling and across whole recordings, the level-of-detail grammar for scenes that outgrow a single rendering budget, and the viewer.

## Results

### The Luxar system

Luxar is designed around a simple principle: scientists should be able to describe their data in Python and explore and share it in a browser, without requiring web development expertise. The core API exposes four geometry types, organised in a hierarchical scene graph with groups, transforms, and *n*-dimensional coordinate systems (Fig. 1a): `Points` (positions, colours, radii), `Lines` (vertices, widths, colours), `Mesh` (vertices, triangle faces, optional normals and texture), and `GSplats` (centres, amplitudes, Cholesky factors; the representation of *n*D images).

Data is compiled progressively to disk, so datasets far exceeding RAM can be processed on a laptop. Everything downstream is organised around one goal: load only what the current view requires, and load it as fast as possible. Elements are quantised, compressed and chunked along a space-filling curve led by the non-displayed dimensions, and every chunk carries its *n*D bounding box, so at each view or slice change the viewer fetches only the chunks that intersect the displayed frustum and the current slice. Stepping a time-lapse fetches the chunks of the new timepoint, whatever the recording’s length (Supplementary Document 14, Supplementary Videos 4 and 5). An optional level-of-detail grammar (below) adds the second axis of economy: additive ladders paint a coarse scene from the first chunk and sharpen it as data arrives, and substitutive levels and spatial tiles let a distant or off-screen object cost a fraction of its full size (Supplementary Videos 6 and 7). A client-side cache with prefetching hides network latency when the user orbits or scrubs a dimension (Methods). The compiled archive, a directory of chunks or a single zip file, is served by any static HTTP host. A browser viewer (WebGL, or WebGPU) with HDR tone mapping renders it, either run locally or as the hosted copy at https://luxarviewer.dev, which opens any archive from its URL (see Methods and Supplementary Document 10). Luxar scales from a handful of primitives to millions of points and splats while remaining interactive (measured below, under Viewer performance). Compilation has been exercised on 100-million-voxel light-sheet volumes (Table 1).

**Table 1.** Gaussian splat reconstruction quality across microscopy modalities. Every volume is reported at its cross-validation-optimal splat count (CV optimal: the target seed count; Splats, the count the fit retains after its conservative cull of *∼* 0.1%-amplitude splats, the count the archive stores): full-volume PSNR of the unmasked fit against the source volume, SSIM (in percent), on-disk compression ratio (CR: bytes of the source volume at its stored bit depth, Table 3, over the bytes of the stored, quantised and compressed splat archive) and the same archive as bits per source voxel and the optimisation time of one fit at that count (the Adam loop alone, excluding seeding and the post-fit cull) on a single NVIDIA RTX PRO 6000 (Blackwell) GPU, run alone on the card with the same protocol as the sweep (L1 loss, 20,000-iteration cap with early stopping, no background-floor suppression; on an 8 GiB RTX 3070, with the shipped floor default and a fixed 32K budget, the median end-to-end fit takes about 2.4 min, Supplementary Document 13). Best PSNR is the maximum over the full sweep (to 512K, or 2M for the extended light-sheet datasets); noise floor is the estimated PSNR ceiling (–: every MAD estimator returns exactly zero, so no ceiling is estimable; Supplementary Document 2). Volumes normalised to [0, 1]. Modality: SDC, spinning-disk confocal; LS, light-sheet; iSIM, instant structured-illumination. ^*^Signal-limited: held-out PSNR is still rising at the largest tested capacity (2M splats), so no held-out peak exists in the tested range; the count shown is the detector’s operating point, the smallest capacity within 0.3 dB of the held-out maximum (see Supplementary Document 2). ^†^Cull-limited: beyond this count the post-fit cull holds the retained splat count nearly constant (a growth of less than 1.5 *×* over at least two further doublings of the requested count), so the sweep past it no longer varies capacity and the small held-out decline there is not evidence of overfitting; the count shown is the held-out maximum.

| Dataset | Modality | Shape | CV optimal | Splats | PSNR | SSIM (%) | CR | bits/voxel | Optim. time (min) | Best PSNR | Noise floor |
| --- | --- | --- | --- | --- | --- | --- | --- | --- | --- | --- | --- |
| OpenCell-MAP4 nuclei | SDC | $51 \times 600 \times 600$ | 16K | 16,302 | 28.4 | 66.2 | $183\times$ | 0.09 | 0.4 | 30.6 | 32.6 |
| OpenCell-MAP4 microtubules | SDC | $51 \times 600 \times 600$ | 64K | 62,155 | 25.7 | 62.8 | $53\times$ | 0.30 | 0.6 | 28.0 | 30.4 |
| OpenCell-LMNB1 nuclei | SDC | $112 \times 600 \times 600$ | 32K | 31,961 | 25.6 | 50.8 | $218\times$ | 0.07 | 0.4 | 26.8 | 30.0 |
| OpenCell-LMNB1 nuclear lamina | SDC | $112 \times 600 \times 600$ | 64K | 62,021 | 34.6 | 90.7 | $118\times$ | 0.14 | 0.7 | 36.5 | 42.6 |
| Kidney nuclei | Confocal | $16 \times 512 \times 512$ | 64K | 63,042 | 38.1 | 98.1 | $11\times$ | 1.40 | 4.2 | 45.2 | 41.3 |
| Kidney actin | Confocal | $16 \times 512 \times 512$ | 128K | 124,277 | 39.1 | 98.8 | $6\times$ | 2.65 | 4.1 | 48.1 | 43.5 |
| Cells3D nuclei | Confocal | $60 \times 256 \times 256$ | 32K | 31,877 | 36.8 | 87.4 | $21\times$ | 0.77 | 0.4 | 44.8 | 37.6 |
| Cells3D membranes | Confocal | $60 \times 256 \times 256$ | 32K | 31,837 | 49.9 | 99.2 | $21\times$ | 0.77 | 0.5 | 58.1 | 53.5 |
| Blastocyst nuclear lamina | Confocal | $236 \times 275 \times 271$ | 32K | 28,203 | 39.2 | 93.0 | $99\times$ | 0.16 | 0.4 | 41.8 | 54.4 |
| C. elegans embryo nuclei | Confocal | $41 \times 512 \times 512$ | 8K | 7,948 | 39.3 | 88.9 | $209\times$ | 0.08 | 0.3 | 40.5 | 40.7 |
| Fly brain neurons | Confocal | $435 \times 478 \times 478$ | 64K | 60,723 | 44.9 | 95.9 | $313\times$ | 0.05 | 0.7 | 45.4 | 52.1 |
| Tribolium embryo nuclei | LS | $991 \times 104 \times 965$ | 2048K* | 2,039,735 | 56.0 | 99.8 | $11\times$ | 1.48 | 4.7 | 56.0 | 61.0 |
| Mouse heart nuclei | LS | $597 \times 174 \times 960$ | 2048K* | 1,452,517 | 47.7 | 99.2 | $6\times$ | 1.28 | 10.6 | 47.7 | – |
| Zebrafish embryo nuclei | LS | $407 \times 512 \times 512$ | 256K | 190,131 | 39.7 | 97.3 | $113\times$ | 0.14 | 5.5 | 43.2 | – |
| Drosophila embryo nuclei | LS | $108 \times 1352 \times 532$ | 256K | 228,920 | 34.8 | 88.7 | $65\times$ | 0.25 | 6.0 | 36.4 | 51.1 |
| Neuromast membranes | iSIM | $84 \times 580 \times 576$ | 128K† | 74,765 | 57.2 | 100.0 | $144\times$ | 0.22 | 2.3 | 57.5 | – |
| Neuromast nuclei | iSIM | $84 \times 580 \times 576$ | 64K† | 32,839 | 66.6 | 100.0 | $340\times$ | 0.09 | 2.1 | 68.0 | – |

For *n*-dimensional datasets, displayed spatial dimensions are navigated by orbiting and zooming; non-displayed dimensions (time, channel, experimental condition) are navigated via continuous sliders or categorical toggles (Supplementary Video 2). Points and Gaussian splats are treated as bounded *n*D extents whose visibility follows from intersection with the current slice; line segments are clipped against that slice and triangles are culled whole (see Methods).

### Fitting microscopy volumes to Gaussian splats

Luxar decomposes an nD microscopy volume *V* : ℝ^*D*^ → ℝ _*≥*0_ into a mixture of *K* anisotropic Gaussians:

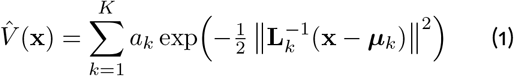

where ***µ***_*k*_ is the position, *a*_*k*_ *>* 0 the amplitude, and **L**_*k*_ the Cholesky factor of the covariance 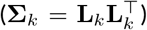, ensuring positive-definiteness by construction. Parameters are optimised with Adam;^25^ periodically, the least informative splats are relocated to regions of high reconstruction residual, a fixed-pool analogue of the pruning-and-seeding cycle used in standard 3D Gaussian splatting (3DGS) that keeps optimiser tensor shapes constant (see Methods). The reconstruction loss is L1 with regularisation on splat amplitudes and extents (see Methods). Initial positions are seeded automatically from the volume itself, from gradient-weighted samples of the volume’s edges complemented by a coarse uniform grid (see Methods; the whole pipeline, from raw stack to streamed scene, is shown in Supplementary Video 1). Fitting in a single pass reaches a higher training PSNR than a progressive residual-pass fitter at the same nominal budget on 15 of 17 volumes, at comparable cost (Supplementary Fig. 2; Supplementary Document 3).

### Benchmark volumes

We evaluated reconstruction across 17 volumes from 12 datasets; the Datasets section of Methods lists every volume with its source. The modalities are spinning-disk confocal (OpenCell-MAP4 nuclei and microtubules, OpenCell-LMNB1 nuclei and nuclear lamina^26^), confocal (Kidney nuclei and actin,^27^ Cells3D nuclei and membranes,^27^ Blastocyst nuclear lamina,^28^ *C. elegans* embryo nuclei,^29^ and Fly brain neurons from one raw FlyLight MultiColor FlpOut (MCFO) tile^30, 31^), light-sheet (*Tribolium* embryo nuclei,^15^ Mouse heart nuclei,^32^ Zebrafish embryo nuclei (*h2afva* histone line), and a raw SiMView frame of *Drosophila* embryo nuclei^11^) and instant structured-illumination microscopy (iSIM: Neuromast membranes and nuclei^33^). The volumes span 4 to 107 million voxels (Table 1; Fig. 2 shows four of them at increasing splat counts, Fig. 3 nine of them, Supplementary Document 2 all).

**Figure 2.**
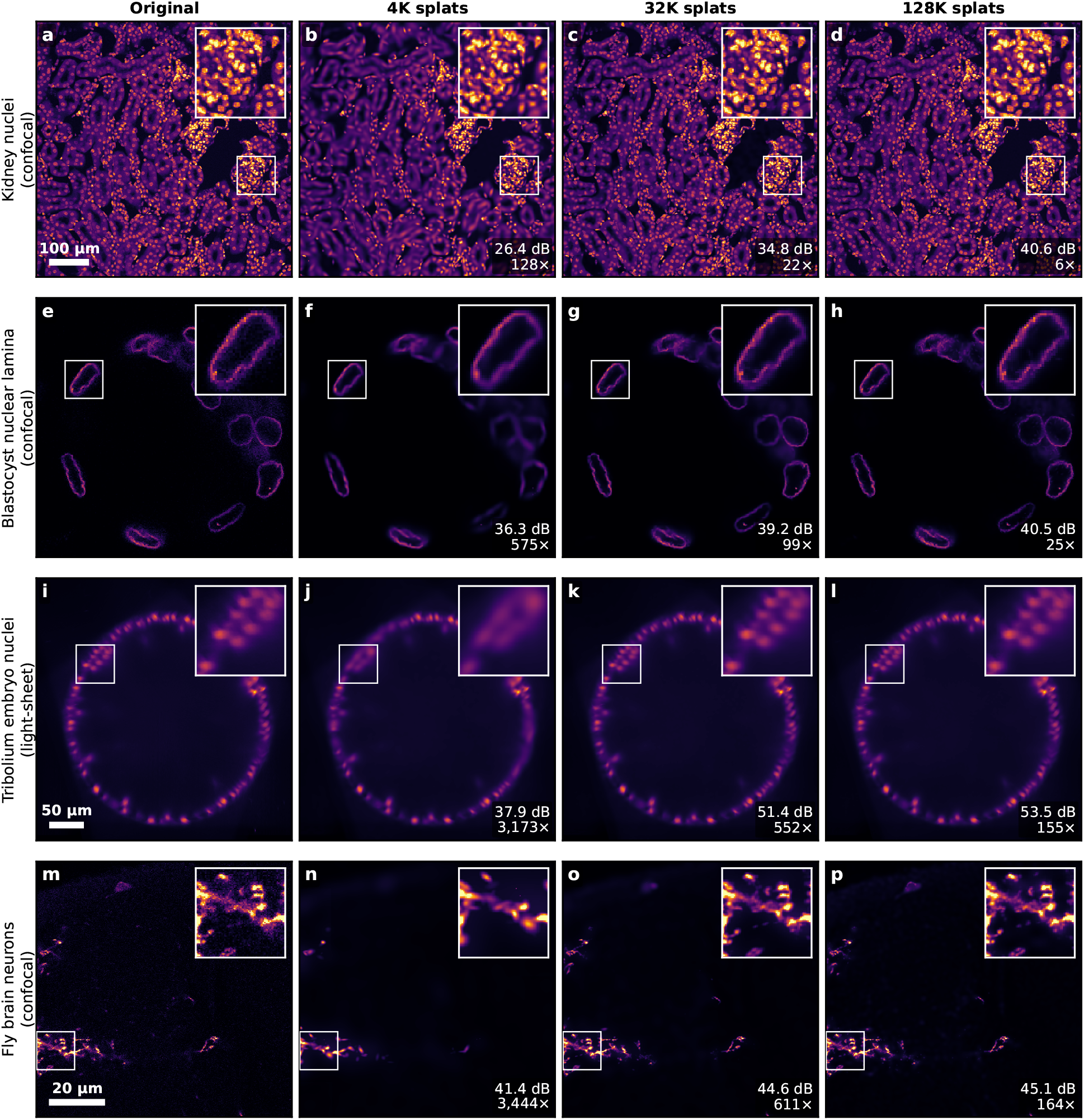
Visual quality of Gaussian splat reconstructions across modalities. Representative slices at increasing splat counts (left to right: ground truth, then increasing model capacity); the inset enlarges the boxed region of each panel. **a–d**, Mouse kidney^27^ (confocal, DAPI): 128K splats recover fine nuclear morphology at 6 *×* compression. **e–h**, Mouse blastocyst^28^ (confocal, LaminB1 immunostain): the nuclear envelopes are faithfully reconstructed as closed rings. **i–l**, *Tribolium* embryo^15^ (light-sheet): 4K splats already recover the embryo’s global morphology at 3,173 *×* compression 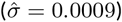. **m–p**, Fly brain neurons^30^ (confocal, one MCFO signal channel of a raw, unstitched FlyLight 63 *×* tile): sparse neurites against a near-empty, shot-noise-limited background; 4K splats already suppress the noise and recover the main tract at 3,444 *×* compression, and 32K recover the fine branches. This row is displayed with a window set to the 99.9th percentile of the original slice (the same window for all four panels) because the neurites are dim; the other rows use the full [0, 1] range. White annotations: PSNR and compression ratio. Compression ratio is the on-disk ratio: the bytes of the source volume at its stored bit depth (Methods, Datasets) divided by the summed size of the encoded chunk and metadata files of the fit’s stored splat archive, as the viewer reads them (Methods). The post-cull splat count may be slightly below the nominal seed count after the conservative pruning pass described in Methods. Scale bars are shown in the original panels where a voxel size is recorded (none is available for the Blastocyst volume); source dataset dimensions are listed in Methods.

**Figure 3.**
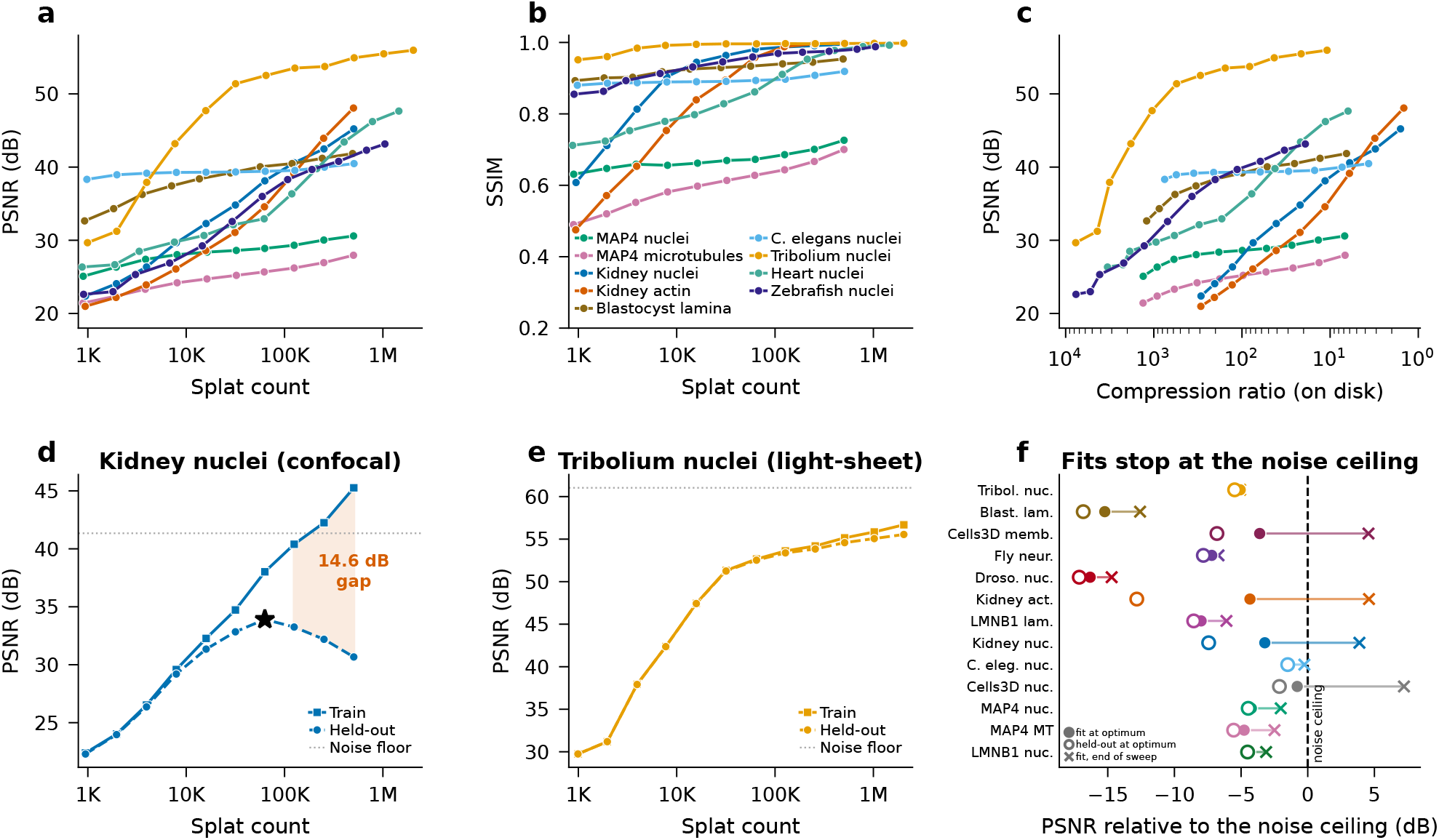
Quantitative analysis and noise-aware model selection. *Top row:* rate-distortion analysis across nine of the 17 benchmark volumes (all of them in Supplementary Fig. 3 and Supplementary Document 2). **a**, PSNR vs. splat count. All datasets show log-linear improvement. **b**, SSIM vs. splat count. **c**, PSNR vs. on-disk compression ratio, as in Table 1 (higher compression to the left). *Bottom row:* Blind-spot cross-validation. **d**, Kidney nuclei (confocal): held-out PSNR peaks at 64K splats (star) then declines, leaving a 14.6 dB overfitting gap at 512K. **e**, *Tribolium* embryo nuclei (light-sheet): no overfitting; train and held-out curves overlap. **f**, Every fit relative to its volume’s noise ceiling, for the thirteen volumes with an estimable one (volumes sorted by ceiling): PSNR of the fit at the operating point (filled) and of the held-out score there (open), and of the fit at the end of the sweep (cross), each minus the ceiling. At the operating point every fit sits below the ceiling (median 4.4 dB; held-out median 6.8 dB). Beyond it the fit crosses the ceiling on the Kidney and Cells3D volumes, which is memorised noise. *Tribolium* is signal-limited and its operating point is the 2M end of the sweep; Mouse heart nuclei, Zebrafish embryo nuclei and the two neuromast channels have no estimable ceiling (every median-absolute-deviation (MAD) estimator returns 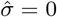, which bounds nothing) and are not shown.

### Quality scales log-linearly with splat count

All datasets exhibit a log-linear rate-distortion relationship (Fig. 1d, Fig. 3a). The *Tribolium* light-sheet volume (estimated noise level 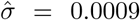 in normalised intensity units; Methods) reaches 51.4 dB PSNR at 32K splats (Fig. 2i–l), and keeps improving to 54.9 dB at 512K and 56.0 dB at 2M (Supplementary Fig. 3). The Kidney nuclei volume reaches 34.8 dB at 32K splats and 45.2 dB at 512K (Fig. 2a–d). Structural similarity (SSIM) follows a similar pattern (Fig. 3b). Convergence is rapid: at 2,000 iterations training PSNR is within 0.5 dB of its 20,000-iteration value on 77 of 85 volume–capacity cells and within 1 dB on 81. The largest shortfall is 1.50 dB (Neuromast nuclei at 16 K), and 67 cells are within 0.5 dB already at 1,000 iterations. Adding capacity also outperforms adding iterations at lower fitting cost: 16K splats × 500 iterations beat 4K splats × 20,000 iterations by an average of +3.03 dB across 17 volumes at 10 × fewer splat-iterations (the measured loop time was 11–52 × shorter on a shared GPU, an indicative range) (Supplementary Fig. 4, Supplementary Document 4). Fitting completes in minutes per volume on a single GPU at the 32K reference budget. Run to completion under the stopping rule (early stop at patience 500 or the iteration cap), a fit takes a median of about 2.4 min per volume at that fixed budget on an 8 GiB NVIDIA RTX 3070, a commodity consumer card, across all 17 benchmark volumes, none exceeding its memory at that count. twelve of the seventeen operating points of Table 1 (16,000, 32,000, 64,000 and 256,000 splats) lie on that card’s timed grid; the five at 8,000, 128,000 and 2,048,000 splats were not timed on it (Supplementary Document 13). On every volume the training PSNR keeps rising as splats are added. Whether the added splats encode structure or noise is the question we turn to next.

### Cross-validation identifies the boundary between signal and noise

The number of splats needed to represent a volume is primarily determined by its information content, the structural complexity of the biological signal. However, adding splats beyond what the signal requires causes the model to memorise noise. To detect this transition we used blind-spot cross-validation: 5% of the voxels are masked and replaced with donut-median neighbourhood values,^23, 24^ and the reconstruction is scored on the held-out voxels.

The held-out PSNR peaks and then declines for 10 of the 17 volumes swept; on two more, the iSIM neuromast channels, the maximum is interior but the post-fit cull holds the retained splat count nearly constant beyond it, so their declines to the end of the sweep, at most 0.19 dB, compare fits of the same size and are read as cull-limited, not as overfitting (six of the nine in Fig. 3; Fig. 1e, Fig. 3d, Supplementary Document 2). For the Kidney nuclei volume 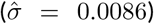, the train–held-out gap widens to 14.6 dB at 512K splats, and the training PSNR overshoots the volume’s noise ceiling by 4 dB, indicating that splats beyond the 64K optimum are fitting noise, not structure. Across volumes, the noise level constrains useful model capacity: the held-out peak provides a per-dataset stopping criterion, with structural complexity driving the absolute optimum (Supplementary Document 2). This provides a practical recipe: fit at increasing capacity, stop at the held-out peak.

### What the operating point buys

Compression ratios are quoted at each volume’s operating point, the splat count selected by cross-validation (for the two signal-limited light-sheet volumes, whose held-out curve is still rising at the end of the sweep, the 2M endpoint, below). At that point the on-disk compression ratio across the seventeen volumes ranges from 6 × to 340 × at 26–67 dB PSNR, with a median of 99.3 × (Table 1; Fig. 1f, Supplementary Fig. 1, Supplementary Document 7); over the nine volumes of Fig. 3c it spans 6–209 ×. The ratio is set by content and noise, not by modality: the sparse volumes compress most (Neuromast nuclei at 340 ×), while the dense light-sheet Tribolium embryo nuclei and Mouse heart nuclei volumes (11 × and 6 ×) and the two noisy confocal kidney channels compress least. On noisier confocal data a lower splat count yields higher-fidelity reconstructions than blindly adding more. These ratios are not bought by discarding the structures the data resolve. At the optimum, adding splats no longer improves the prediction of held-out voxels and starts storing noise, so the residual there is the noise itself plus whatever structure falls below the Gaussian scale. Across the thirteen volumes with a measurable noise floor, the unmasked fit at the optimum, scored against its noisy source, sits a median 4.4 dB below the noise ceiling, and for Cells3D nuclei and C. elegans embryo nuclei within 1.4 dB of it (Fig. 3f). That gap is an upper bound on how far the reconstruction sits from the noise level, because the ceiling is estimated from the background noise (Supplementary Document 2). It does not by itself say how much of the residual is noise and how much unrecovered signal, nor certify every dim or thin feature. The reconstructions are therefore also shown at the scale a reader can judge (Fig. 2, Fig. 4), and Supplementary Fig. 5 marks, automatically, where each fit at its optimum deviates most from its source. On five of six volumes the strongest automatically ranked deviation is a dim structure reproduced too faintly or dropped (the dimmest nucleus of a field, a faint membrane sheet), not an invented one; on the background-subtracted neuromast it is the dark gap between membranes filled in (Supplementary Document 2). The capacity beyond the optimum buys PSNR mostly by memorising noise (on Cells3D nuclei the full sweep ends 7.2 dB *above* the ceiling), which is exactly the part of a volume that cannot be compressed and that a voxel codec is forced to keep. At the operating point a fit alone on an NVIDIA RTX PRO 6000 takes 0.3– 6.0 min for the volumes with a finite optimum and up to 11 min for the two signal-limited light-sheet volumes at 2M splats (Table 1).

**Figure 4.**
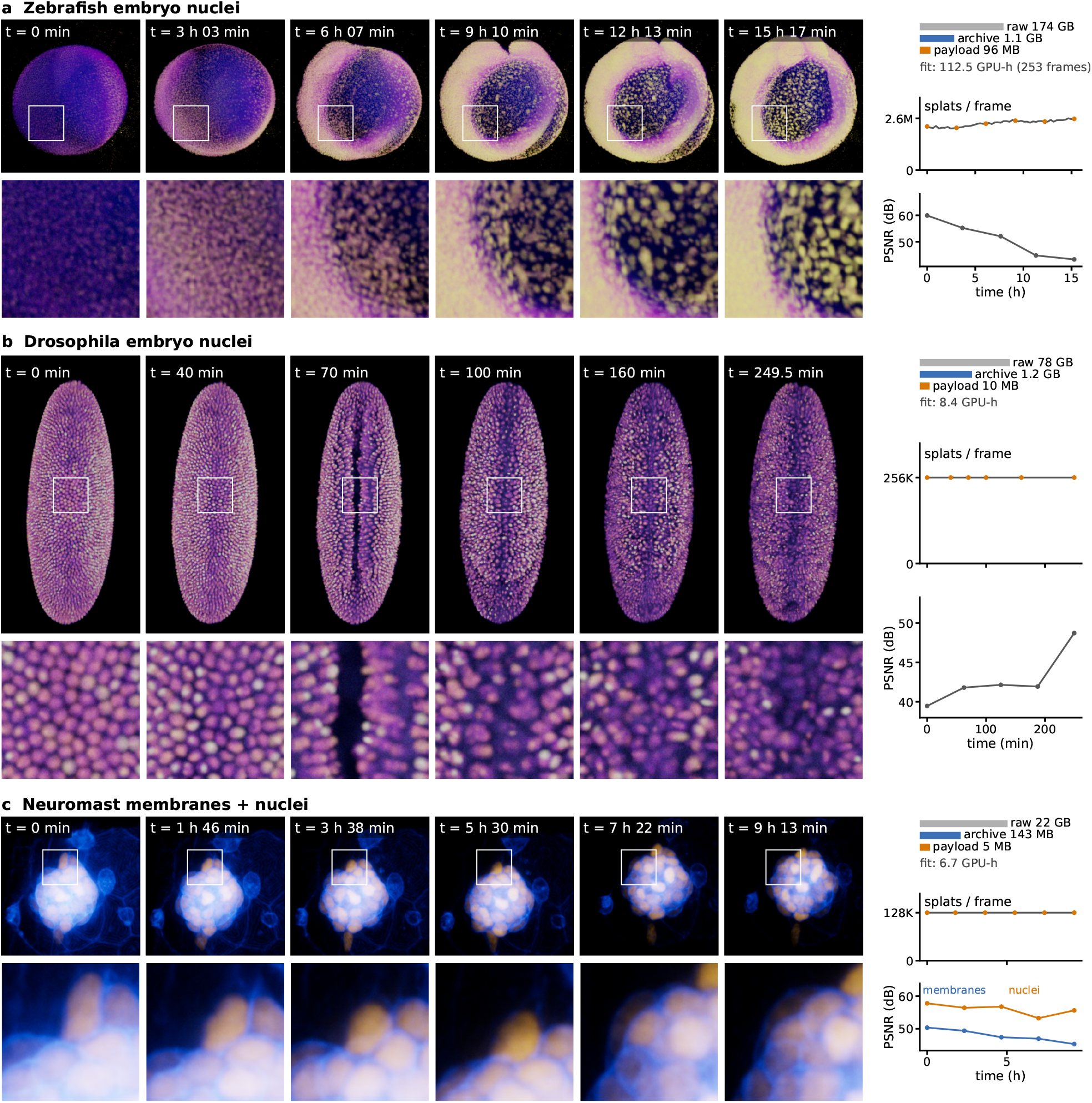
Gaussian-splat time-lapses. Three recordings fitted per timepoint, each shown at six timepoints (left to right) as viewer frames with the boxed region enlarged beneath: **a**, Zebrafish embryo nuclei (*h2afva* histone line; 51 of 253 timepoints, 220-s frames of which every fifth is kept, a median of 2.4M splats per frame); **b**, *Drosophila* embryo nuclei through gastrulation (SiMView light-sheet;^11^ 500 timepoints at 30 s, 256K per frame); **c**, the two-channel Neuromast recording (membranes and nuclei, iSIM; 100 timepoints at 5.6 min, 128K per frame over both channels). Frame labels give elapsed time from the first frame shown. Beside each row: the raw frames, the archive on disk and the decoded parameter payload of one timepoint’s splats (40 bytes per splat) on a logarithmic byte scale (raw-to-archive 157*×*, 67*×* and 157*×*), with the fitting cost of the recording, as elapsed hours of one RTX PRO 6000 GPU, beneath (Table 2). Then the splat count of every frame of the recording, the six shown frames marked, and the fidelity at five frames spread over the recording, as the PSNR of the shipped archive against the raw frame (both neuromast channels). Frames are captured with bloom off and every level-of-detail group at its finest level once loading had settled, through one window size per row placed on the drifting specimen, with the appearance each demo store carries and one camera per row (Supplementary Document 14). Supplementary Video 8 orbits the zebrafish embryo at one timepoint, Supplementary Video 3 plays the *Drosophila* and neuromast recordings in the viewer, and Supplementary Video 9 plays the whole *Drosophila* recording streamed from the demo site.

### When the held-out curve does not turn down

The lightsheet volumes are the exceptions: none of the four shows a held-out decline beyond the 0.06 dB by which the zebrafish curve dips from 1M to 2M splats (Fig. 3e and Supplementary Fig. 3). Part of this is expected from the theory. The blind-spot argument^23^ rests on the noise being independent across voxels given the signal, and any processing that mixes neighbouring voxels (multi-view fusion, deconvolution, denoising, resampling) correlates the noise, so the held-out curve need not turn down (Methods). The volumes fall into two regimes (Supplementary Fig. 6). The deconvolved Zebrafish embryo and the raw *Drosophila* frame reach a held-out plateau, both from 256K splats, as does the one confocal volume without a decline, the sparse Fly brain tile. *Tribolium* and Mouse heart nuclei are signal-limited: their train and held-out curves track each other and keep rising to 2M splats, for Mouse heart even with the iteration budget tripled, and *Tribolium*’s held-out PSNR is still 5.5 dB below its 61 dB noise ceiling at 2M (Methods; Supplementary Document 2). Wherever a noise ceiling is estimable the held-out curve, rising or flat, stays below it, so the same stopping criterion applies in every regime. The practical recommendation follows: calibrate the splat budget on unprocessed camera data whenever it is available, and where only processed data exists and a noise floor is estimable, stop before the training PSNR reaches it, since a fit above the floor is reproducing noise; the floor caps the budget from above and does not locate an optimum, because it says nothing about how close the fit is to the clean signal (Supplementary Document 2).

**Table 2.**
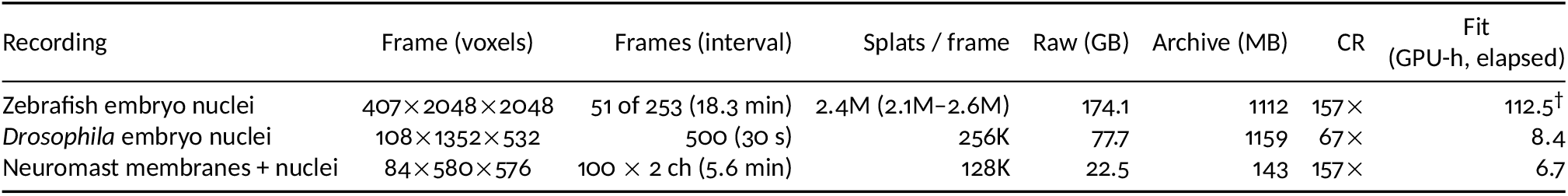
Gaussian-splat time-lapses: storage and fitting cost. The three recordings of Fig. 4, each fitted per timepoint: frame size, frames fitted and the interval between retained frames (the zebrafish keeps every fifth 220-s frame), splats per frame (median, with the 5th–95th percentile range where it varies), raw bytes of the frames the archive represents (the zebrafish store keeps every fifth of the 253 fitted frames) at the dtype each source is stored in (uint16 for the zebrafish and *Drosophila* stacks, float32 for the deconvolved neuromast) against the archive on disk and their ratio (CR, on the same stored-byte basis as Table 1), and the fitting cost on a single NVIDIA RTX PRO 6000 GPU. ^†^All 253 frames were fitted under content tiling with a configuration the archive does not record; the store keeps every fifth (Supplementary Document 14). Streaming cost and fidelity are in Supplementary Table 2; details and recipes are in Supplementary Document 14.

| Recording | Frame (voxels) | Frames (interval) | Splats / frame | Raw (GB) | Archive (MB) | CR | Fit (GPU-h, elapsed) |
| --- | --- | --- | --- | --- | --- | --- | --- |
| Zebrafish embryo nuclei | 407×2048×2048 | 51 of 253 (18.3 min) | 2.4M (2.1M–2.6M) | 174.1 | 1112 | 157× | 112.5<sup>†</sup> |
| *Drosophila* embryo nuclei | 108×1352×532 | 500 (30 s) | 256K | 77.7 | 1159 | 67× | 8.4 |
| Neuromast membranes + nuclei | 84×580×576 | 100 × 2 ch (5.6 min) | 128K | 22.5 | 143 | 157× | 6.7 |

### Against voxel codecs

Compared against H.265 video compression under the same blind-spot held-out scoring, which rewards predicting the held-out voxels over reproducing the noisy input, the two held-out rate-distortion curves cross. At matched bit rate, crediting H.265 with the larger of its best held-out result at any rate up to the bits per voxel of the Gaussian-splat cross-validation optimum and its interpolated value at that rate, Gaussian splats recover more signal on 15 of 17 volumes and tie within the 0.2 dB replicate band on 2 (median +0.98 dB). Read at the exact operating rate, the margin is +0.50 to +7.27 dB (median +1.32 dB) on 16 of 16. At H.265’s own, mostly lower, peak rate the codec stays ahead on 9 of the 12 volumes whose splat sweep reaches that rate. Compared at each method’s own peak, which occurs at a different rate for each, Gaussian splats attain the higher blind-spot optimum on 12 of 17 volumes, by up to +5.58 dB, with the remaining 5 inside the replicate band (the *Tribolium* volume at − 0.09 dB among them) (Supplementary Fig. 7, Supplementary Document 7).

### Tiled fitting for scalability

For volumes exceeding GPU memory, Luxar partitions the volume into overlapping tiles, each fitted independently, and concatenates the results. Raised-cosine taper windows form a partition of unity at tile boundaries: adjacent windows sum to exactly 1.0, so the windowed targets add back to the volume. Whether the fitted tiles do is an empirical question. In a controlled comparison against monolithic fits at the same splat budget (Supplementary Document 11), the tiled union’s seam-to-interior residual ratio stays within 1.01–1.13 of the seamless fit’s, at most 13% above it on the 81-tile grid (*T* = 128, *L* = 13) of the dense lightsheet volume, with an overlap of 5–20% of the tile size, on both a sparse confocal and a dense light-sheet volume (one fit per cell, no seed replicates). A hard cut without overlap raises it to 2.15–7.52, and overlapping tiles *without* the taper are worse still (8.0–30.8 dB below the monolithic fit). Tiling at a fixed splat budget and 5–20% overlap costs at most one decibel of PSNR (− 0.98 to +0.19 dB over 10 configurations of 4 to 100 tiles), the finest grids of a few hundred splats per tile paying the most; the budget is divided over the tiles in proportion to their intensity mass (Methods). Tiles are embarrassingly parallel, and the scheduling is built in: the batch fitter plans the tiling of a whole volume or recording once, then spreads the plan over the GPUs of one machine or submits it as Slurm array jobs on a cluster, and a fan-in merge assembles one scene that streams spatial tiles as parts (Methods). A 2048^3^ float32 volume that would exceed consumer GPU memory (e.g., NVIDIA RTX 3090, 24 GB) could thus be decomposed into overlapping 256^3^ tiles. The same batch system reads OME-Zarr^34^ 5D layouts (time, channel, Z, Y, X) directly, fitting every timepoint and channel as its own task (Supplementary Video 5). The partition-of-unity proof and the implementation conventions are detailed in Supplementary Document 11.

### Time-lapses at scale

The same machinery carries whole recordings as well as single volumes (Fig. 4, Table 2, Supplementary Videos 8, 3 and 9). Three time-lapses were fitted per timepoint with the batch fitter: a zebrafish embryo (*h2afva* histone line; 51 of its 253 timepoints kept, 2.4M splats per frame), a *Drosophila* embryo through gastrulation (SiMView light-sheet;^11^ 500 timepoints at 30 s, 256K splats per frame and 128,000,000 in all) and a two-channel lateral-line neuromast (iSIM; 100 timepoints, 128K splats per frame over both channels). Time is a hard ordering barrier in each archive: a timepoint’s splats are stored contiguously and no level-of-detail rung blends two frames, so one timepoint is resident at a time and a time step fetches only the chunks that hold the new frame. In the layout the demo site publishes, a step costs 1 request for the *Drosophila* scene and, for the zebrafish, one per-timepoint part (17.91–17.96 MB; Supplementary Table 2). A time-lapse is stepped far more often than it is opened, so the chunk layout is resolved in favour of the step (Methods). Fitting is the expensive step here too: the *Drosophila* recording took 8.4 hours of one RTX PRO 6000 (two fits in flight; the optimisation loop alone took a median 0.8 min per frame, seeding and I/O about as much again), the neuromast pair 6.7 hours, and the full 253-frame zebrafish recording, fitted as 253 frames of content-adaptive tiles, 112.5 hours. The archives are 157×, 67× and 157× smaller than the raw frames they were fitted from, and on 20 frames sampled across the three recordings the shipped archives reproduce the raw frames at 39.5–59.9 dB PSNR. That score is against each raw frame’s own range, not held-out and not on the normalised basis of Table 1 (Supplementary Table 2; Supplementary Document 14).

### A grammar of levels of detail

A fitted dataset is directly viewable, but scenes that outgrow a single rendering budget benefit from a level-of-detail (LOD) structure, which Luxar builds as a separate authoring step over any of its geometry types. Two families are available. *Additive* ladders order elements so that every prefix is a valid approximation of the whole, letting the viewer paint a coarse scene from the first chunk and sharpen it as more data arrives; *substitutive* levels instead merge elements into fewer representatives, so that a coarse level replaces a finer one as an object shrinks on screen. The two compose, and a single command selects among the resulting topologies (`flat, stream, levels, tiles, overview, adaptive`), which trade first-paint latency against peak detail; the spatial variants additionally let the viewer cull tiles outside the view frustum. At display time the viewer streams a ladder rung by rung and picks the coarsest substitutive level whose splats still project to at most 1.5 CSS pixels at the median. It never fetches a level or tile the view does not need (Supplementary Video 6; the six topologies built from one fit are compared side by side in Supplementary Video 7). A distinct *reveal* ordering, available on all four geometry types, instead sorts elements into concentric shells, so that a partial load reads as a growing object instead of a uniformly faint one. Together these form a small grammar for authoring large scenes. Recipe, ladder, level count, tile size and chunk layout are a handful of declarative choices whose consequences (bytes, requests, first paint, resident elements) are measurable, so a scene’s layout can be tuned to how it will be used. A time-lapse that is stepped far more often than it is opened wants different choices from a single volume that is zoomed into (Supplementary Document 14). Because the grammar is small, declarative and measurable, it is also tractable for AI coding agents: given the command line and its measurements, a coding agent can author a scene’s layout and optimise it against those numbers, which is how the largest scenes of the public gallery were laid out and tuned. LOD is opt-in, not a precondition for viewing: a compiled scene opens as is, and a ladder is added only when a scene outgrows a single rendering budget. The analytical foundations are developed in Supplementary Documents 1, 8, and 9: the closed-form merging error used to score substitutive merges, and the sub-modularity result guaranteeing that the greedy additive ordering captures at least 1−1*/e* of the optimal energy at every prefix.

### Viewer performance

On an NVIDIA RTX 3070 at 1280 × 720 the web viewer’s frame callback holds at least 59.3 frames per second (FPS) against its 60 FPS cap up to 10^6^ points, 10^6^ Gaussian splats, and 10^6^ line vertices (Supplementary Document 6). At *N* = 10^5^ a synchronised render-and-readback call with the default HDR pipeline completes in 1.21–1.72 ms, a tenth of the 16.7 ms frame budget (Fig. 1g); at *N* = 10^7^ it takes 18.8– 19.4 ms, 12.3–16.3% over the budget, while the callback rate still reads 60 FPS (Supplementary Document 6 gives two readings the records do not separate). Only large primitives are markedly more expensive. 32–64-pixel Gaussian splats take 41–157 ms per synchronised call, and on those cells the callback intervals during the rendering phase were 67–300 ms before the loop went idle (whether each is one long frame or two frames of the call’s length the records do not say), the regime in which the viewer’s adaptive resolution scaling trades pixel detail for frame rate (Methods). On real scenes, every one of the fifteen measured scenes holds 60 FPS, the heaviest, a 2.2M-splat time-lapse frame, using 5.6 ms of the 16.7 ms frame budget, and a deliberately overloaded 10^7^-splat scene still renders, at 18–44 FPS across repeat runs (Supplementary Table 1). GSplat scenes of 100K–300K splats load in 0.4–0.7 s and the million-splat scenes in 0.4–5.9 s (Supplementary Table 1); larger point-cloud datasets (millions of elements) render at full frame rate once loaded, though initial transfer time scales with dataset size even with progressive streaming (Supplementary Video 4). Full count, size, and viewport sweeps are reported in Supplementary Document 6.

### Layered, annotated, scriptable scenes

Beyond raw rendering, the Luxar viewer exposes a layer model (per-node visibility, opacity, absorption, display range, gamma, blending mode, and colormap with multi-select), text, image, video and HTML overlays with dimension-aware visibility, sound nodes for ambient beds and narration, and per-element hover annotations driven by GPU picking. These enable annotated atlases (e.g. Tabula Sapiens cells with on-hover labels, which can link out to a database entry), per-channel adjustment of multi-channel volumes, and dimension-conditional callouts that appear or disappear as the user navigates (Supplementary Video 10). Camera poses bound to positions along a hidden dimension turn that dimension into a guided tour (Supplementary Video 11). The same programmatic surface that embeds the viewer in a web page drives it remotely, so a scene can run as a narrated kiosk steered from a Python script or a touch panel. The viewer itself works by touch on phones and tablets (Supplementary Video 12). A per-dimension animation system with frame-data synchronisation, paired with a recording panel producing PNG/EXR sequences and WebM/MP4 videos in-browser through WebCodecs, turns nD scenes into shareable movies and figures with no scripting; 10-bit HDR is reached by exporting an EXR frame sequence and encoding it with the accompanying generated ffmpeg script. Exporting a scene to a self-contained folder or a native application, embedding the viewer in an ordinary web page and recording a movie from the panel are shown in Supplementary Video 13. Together these features make the viewer a complete environment for interactive exploration of nD scientific data, not merely a renderer (see Methods and Supplementary Document 10).

## Discussion

Fluorescence volumes can be represented as mixtures of anisotropic Gaussians whose size is set by the data themselves. Across 17 volumes from 12 datasets and four modalities, blind-spot cross-validation places the operating point where adding splats stops predicting heldout voxels, and there the fits store the volumes 6–340 ×smaller (median 99.3 ×) at 26–67 dB, in minutes of GPU time per volume, with the fitted geometry streaming to any browser from static file hosting. Luxar packages this inside a Python-first system, a choice that follows the convergence of bioimaging and scientific computing on Python,^4, 27^ from acquisition control^35^ and analysis pipelines^36^ to machine learning.^37^ It complements the desktop tools where analysis lives (napari,^4^ Fiji^5^) and the voxel-pyramid web viewers (Neuroglancer,^8^ Viv,^38^ Vitessce^39^) without replacing either: it represents images as fitted geometry, mixes them with points, lines and meshes in one scene, and moves only the bytes a view needs, so that data transfer stays practical over modest bandwidth (Supplementary Fig. 7 and Fig. 4).

Cross-validation does more than pick a number. The fixed-pool optimiser is part of why it can: standard 3D Gaussian splatting grows and prunes its Gaussian set during optimisation, whereas here the count is fixed and weak splats are relocated, so the splat count is an explicit quantity that cross-validation can set. The information content of a volume drives that count upward and its noise level caps it, and on the thirteen volumes with an estimable noise ceiling the fit at the operating point sits a median 4.4 dB below the ceiling (Fig. 3f). On the noisy confocal volumes the fit sits within a few decibels of the ceiling and further capacity crosses it, so their reconstruction quality is bounded by the data’s own noise, and no change to the fitter can raise it without a denoising prior. On the cleanest volumes, the Blastocyst lamina and the *Drosophila* frame, the held-out optimum instead lies more than 10 dB below the ceiling: what remains there is structure below the Gaussian scale or noise correlated by processing, not shot noise, and it marks where a finer primitive or a different loss could still gain. The comparison with H.265 reads the same way. Block-transform coding encodes coarse content cheaply and so leads at very low bit rates, while at their operating rate splats recover more held-out signal on 15 of 17 volumes and tie on the rest, with H.265 credited with the larger of its best result at any rate up to that budget and its interpolated value there, because they are fitted to predict the signal, not to reproduce the noisy input. The compressed form is also the renderable form: the splats are rasterised straight from their decoded parameters, with no voxel grid rebuilt on the client, which is what lets a compressed volume stream and render in a browser at all.

The same machinery serves any data that has coordinates. In the life sciences it already carries single-cell atlases^13^ navigated by organ, cell type or donor, super-resolution localisations by the million, 4D lineage tracking^10^ with trajectories over Gaussian-splat nuclei, protein-embedding landscapes, connectomes, tractography, clinical CT and histopathology (Fig. 1b,c; Supplementary Videos 14 and 15), and the three fitted timelapses of Fig. 4. The public gallery at https://demos.luxarviewer.dev holds more than 80 scenes, reaching from the Laniakea supercluster and the DESI and Gaia^40^ surveys to a NEXRAD thunderstorm as a 4D Gaussian-splat volume, ocean currents, and a photogrammetric Gaussian-splat capture of an insect imported from another tool (Supplementary Video 16). Nothing in the system knows what a nucleus or a galaxy is; it knows points, lines, meshes and Gaussians in *n* dimensions (Supplementary Video 17).

The most closely related prior work is the Adaptive Particle Representation (APR),^16^ which likewise exploits the sparsity of fluorescence images by placing primitives where signal exists. The two are complementary points in design space. APR is a content-adaptive substitute for the pixel grid, built for near-linear conversion and for running segmentation, tracking and filtering on the particles themselves; Gaussian splats are built for continuous, GPU-rasterised rendering, with anisotropic primitives that orient along structures and a representation a standard browser decodes. APR serves computation on the representation; splats serve sharing and interactive visualisation. In computer graphics, Gaussian splats have been fitted to tomographic volumes for X-ray novel-view synthesis and CT reconstruction;^41, 42^ those methods target radiodensity fields observed through projections, and here the fluorescence volume itself is the fitting target.

Luxar has limits, and they follow from its design. A Gaussian splat is an approximation: structures thinner than the local Gaussian scale, such as individual microtubules, blur, and on the cleanest volumes the fit plateaus well below the noise ceiling (above). The representation is built for looking, not measuring; quantification, segmentation and tracking belong on the source voxels. The pipeline is asymmetric: fitting is the expensive side and viewing the cheap one. A volume takes minutes of GPU time (Table 1), a 500-frame time-lapse about eight hours of one GPU (Supplementary Document 14), and the largest volumes still call for a cluster, whereas the result renders in milliseconds per frame on a laptop. The representation is made once and looked at many times, the right trade for sharing but not for a quick look at a single stack. Blind-spot cross-validation assumes noise that is independent across voxels, so on fused or deconvolved volumes a held-out peak is not guaranteed; where a noise floor is estimable it caps the budget from above, since a fit above it reproduces noise, without locating an optimum. The viewer’s performance was measured on one GPU and browser configuration and no other viewer was benchmarked, so those numbers are absolute costs, not a comparison. And every benchmark volume is fluorescence microscopy: the representation carries nothing specific to it, but its behaviour on other modalities is untested here.

The representation matters more than the tool. Once an image is a scene (Gaussians, points, lines and surfaces that stream at the resolution the view demands) and no longer an array, the boundary between visualising data and sharing it disappears: the object a scientist explores on a laptop is the same one a collaborator opens elsewhere or a reviewer inspects, in one coordinate system with the tracks, segmentations and annotations derived from it. Fitting can move closer to the microscope, so that a recording becomes explorable while it is still being acquired, and because the level-of-detail grammar is small and measurable, publishing an explorable scene can become as routine as exporting a figure. We expect the same machinery to serve electron and X-ray tomography, functional imaging, spatial omics, and the astronomical and geoscience surveys already in the gallery. The imaging data now being produced can only be shared and understood at its native scale through representations built for streaming, and the browser, which every scientist already has open, is where that understanding will happen. We hope Luxar makes *n*-dimensional scientific data as easy to share and explore as a web link.

## Methods

### Zebrafish embryo light-sheet imaging

Embryos of the tg(*h2afva:h2afva*-mCherry) line, whose every nucleus carries a fluorescent histone-H2A variant fusion, crossed with tg(*mezzo*:eGFP), were raised at 28.5^*°*^C under protocols approved by the UCSF institutional animal care and use committee. For imaging an embryo was dechorionated with forceps, embedded in 0.1% low-gelling-temperature agarose inside a fluorinatedethylene-propylene (FEP) tube held in an agarose-filled capillary holder, and kept in filtered embryo medium in the microscope chamber. It was imaged on OpenSiMView,^43^ an open-hardware simultaneous multi-view light-sheet microscope of the SiMView design:^11^ two opposing illumination arms sweep a light sheet through the specimen while two opposing water-dipping detection objectives (16 ×, 0.8 NA) image the illuminated plane onto sCMOS cameras, giving four views of every plane. The light sheets and detection planes are kept aligned by the adaptive optimisation of that design. The recording ran from gastrulation onward as a sequential two-channel time-lapse; the histone channel used here comprises 253 timepoints acquired every 220 s (15.4 h from first to last frame), each a stack of 407 × planes of 2048 2048 pixels at 0.406 *µ*m laterally and a 1.625 *µ*m axial step (four views per timepoint). The four views of each timepoint were registered, fused with intensity equalisation and deconvolved (ten iterations against the point-spread function of the 16×/0.8 NA objective) with our GPU-accelerated light-sheet processing library DEXP,^44^ yielding one 407×2048×2048 fused volume per timepoint. The Zebrafish embryo nuclei benchmark volume is the content-dense 407 × 512 × 512 sub-region of the last time-point of this series (Datasets, below), and the time-lapse of Supplementary Document 14 uses every fifth timepoint (51 of the 253) at full frame size.^45^

### Datasets

The 17 benchmark volumes come from 12 physical acquisitions (Table 3; co-registered channels of one acquisition share a dataset) and are registered in one place in the analysis code, from which every table, figure and count in this manuscript is generated. Except where noted, they are drawn from public repositories; a volume is named by its specimen and the structure the channel labels. Crops and slices: the Blastocyst volume is channel 0 of IDR idr0062 image 6001240, centre-cropped along its longest axis to at most 20 million voxels. The *C. elegans* volume is timepoint 100 of a Cell-Tracking-Challenge-style confocal recording of embryonic nuclei,^29, 46^ taken from Zenodo record 6460303 (`series mskcc_confocal_s1`).^10^

**Table 3.**
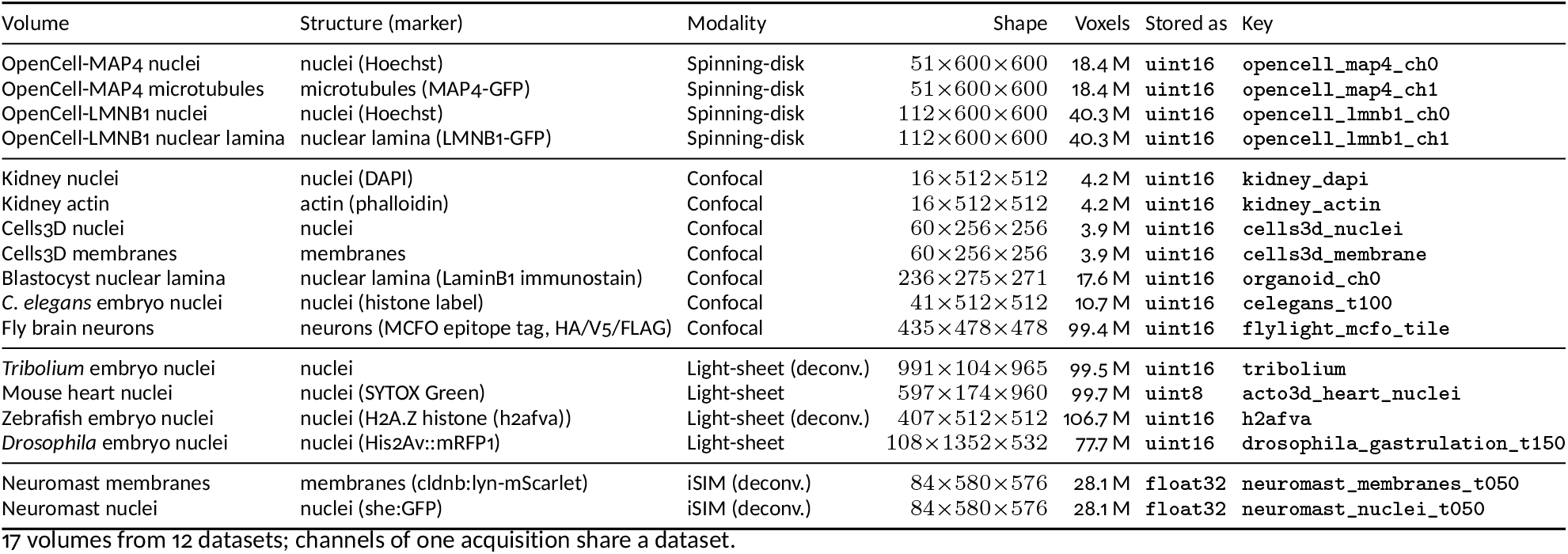
Benchmark volumes. Manuscript name (specimen and labelled structure), structure with its marker, imaging modality, voxel grid and count, the data type the source is stored as, and the key used in the analysis code; sources are given in the text above.

The *Tribolium* volume is the multi-view fused and deconvolved Cell Tracking Challenge recording,^15^ as re-hosted in Zenodo record 5270323 (the first timepoint, deposited as a supplemental file of the GIANI study^47^), centre-cropped along its longest axis to at most 100 million voxels. The Mouse heart volume, from the Acto3D project’s public sample data,^32^ is centre-cropped the same way to at most 100 million voxels. The Zebrafish embryo nuclei volume (*h2afva* histone line) is the content-dense 407 × 512 × 512 sub-region of the last timepoint of a multi-view fused, deconvolved time-lapse (Zebrafish embryo light-sheet imaging, above). The Fly brain neurons volume is one raw 16-bit FlyLight 63 × MCFO tile^30, 31^ (VT019012, slide 20140423_20_D5, right-dorsal image 2), of which we use the MCFO signal channel with the larger Otsu fore-ground fraction (channel 2) and the densest 478 × 478 lateral region over the full 435-plane depth (the same densest-region rule as the Zebrafish crop). The *Drosophila* embryo nuclei volume is frame 150 of the 500-timepoint SiMView recording,^11^ whose four views (two illumination arms, two detection cameras) were registered and fused into one volume per timepoint with the SiMView processing pipeline^48, 49^ without deconvolution (no crop). The Neuromast membranes and nuclei volumes are the two channels of zero-based timepoint 50 (label t51) of the 100-timepoint iSIM recording.^33^

### Normalisation and stored depth

The source volumes are stored as 16-bit integers, except the Mouse heart TIFF (8-bit) and the two neuromast channels, whose deconvolution output is float32 (quantising it back to 16 bit would itself be a lossy step, so its stored depth is the basis of its compression ratio). All volumes were normalised to [0, 1] before fitting by an affine map of their intensity range, with dataset-specific clipping of outlier voxels chosen when each loader was written. The Kidney, Cells3D, Blastocyst, *Tribolium* and Mouse heart volumes are not clipped (plain min–max). The four OpenCell channels are clipped to the 1st–99.5th percentiles, *C. elegans* to the 1st–99.999th, the sparse raw Fly brain tile to the 0th– 99.999th, and the Zebrafish volume (whose deconvolution leaves rare hot voxels) and the *Drosophila* frame to the 0th–99.9th. The two Neuromast channels first have their per-channel background floors subtracted (105.99 and 103.89 grey levels, pinned when the demo store was built, clipped at 0) and are then clipped to the 0th–99.9th percentiles. The percentile-clipped fraction is therefore at most 1.5% of voxels and 0.1% or less on the light-sheet and iSIM volumes (the neuromast floor subtraction additionally zeroes the sub-floor background). All single-volume PSNRs (Table 1, Fig. 3 and Supplementary Documents 2–5, 7 and 12) are referenced to these normalised volumes. The time-lapse fidelity scores of Fig. 4 and Supplementary Table 2 instead score each archived frame against its raw full-frame source (a constant background of 8 grey levels subtracted for the *Drosophila* recording, none for the others) over the source’s own intensity range, with a per-frame least-squares gain for the neuromast channels and an Otsu foreground mask for the foreground PSNR (Supplementary Document 14). They are therefore not directly comparable with the benchmark scores.

### Background floor

For the benchmarks reported here we applied no background-floor suppression (`--floor none`), so both the fits and the reported PSNR are referenced to the original volume including its background pedestal (consistent with the voxel-averaged MSE of the rate-distortion analysis, below). The shipped `gsplat fit` and `cal` tools instead default to automatic floor suppression (`--floor auto`), which subtracts a constant background pedestal before fitting. Because that pedestal is not structural signal, removing it concentrates representational capacity on real features and improves compression, but it lowers PSNR measured against the original (pedestal-bearing) volume and would require a background-relative reference to score on equal footing. We therefore report original-referenced numbers with floor suppression disabled, for comparability across datasets and against the voxel codecs; a background-relative evaluation is left to future work. The one exception is the commodity-GPU timing supplement (Supplementary Document 13), which measures wall-time, not quality, and was run with the shipped `--floor auto` default; its wall times describe that recipe (a different target changes the optimisation trajectory and where it stops), and its runs are not bit-comparable with the quality benchmarks.

### Gaussian splat model

Each splat *k* is parameterised by a centre ***µ***_*k*_ ∈ ℝ^*D*^, a positive amplitude *a*_*k*_ (via softplus activation), and a lower-triangular Cholesky factor **L**_*k*_ ∈ ℝ^*D*×*D*^. All results use the standard Gaussian falloff. The model prediction at voxel **x** is 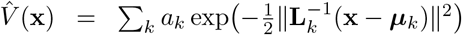 This un-bounded Gaussian is the analytical model used to score merges (Supplementary Documents 1 and 8). The fitter and the viewer evaluate each splat as the shifted, truncated 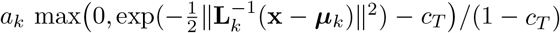 with *c*_*T*_ = exp(*T* ^2^/2) at *T* = 2.75 standard deviations, so that every splat has compact support and the sum is exactly zero far from any centre (Supplementary Document 1, §8.4). The Cholesky parameterisation stores *D*(*D*+1)/2 elements per splat (6 for 3D) and guarantees positive-definite covariance by construction; its diagonal is parameterised as *σ*_min_ +softplus(·) with 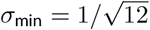 voxels, the standard deviation of a uniform voxel, so no splat is narrower than a voxel’s own spread. Amplitudes are made non-negative by a softplus activation and are upper-bounded by a configurable maximum *a*_max_ (by default the larger of 1 and the normalised peak, i.e. 1 for a volume normalised to [0, 1]). Fits are performed in voxel units and the physical voxel anisotropy is applied at display time. The smallest fitted splats are 0.50–1.38 voxels in every benchmark volume (0.50 voxels being the scale at which relocated splats are re-seeded, which sets the small end on four volumes; Supplementary Document 2). That is at or below the effective resolution of the data (resolution over splat FWHM 1.15–8.7, median 2.4; Supplementary Table 3, Supplementary Document 2), so the representation is not resolution-limited at its small end.

### Optimisation

We use standard PyTorch Adam^25^ (*β*_1_=0.9, *β*_2_=0.999, *ϵ*=10^*−*8^), with fused Adam on CUDA when available. Adaptive model capacity comes from a fixed-pool relocation mechanism (below), not from topology changes, which keeps optimiser tensor shapes constant and preserves the benefits of standard vectorised Adam. The base learning rate is 0.01 with gradient-dilution compensation (the residual passes of the progressive fitter use 0.03; Supplementary Document 3), and it follows a reduce-on-plateau schedule on the regularised training loss (patience 15 iterations, factor 0.9, threshold 10^*−*3^, floor 10^*−*8^). For *D* ≤ 3, the effective rate is scaled by *P*_*D*_*/P*_2_, where *P*_*D*_ = *D* + *D*(*D*+1)/2 is the number of position and Cholesky parameters per splat (5 for 2D, 9 for 3D). For *D* ≥ 4, an additional factor of *D*^0.8^ accounts for the larger spatial search (the exponent was chosen empirically to stabilise convergence in 4–8D). Amplitudes are given 3 × the dilution-compensated rate, as they converge faster than shapes, and centre positions 1.5 ×, since a mis-positioned Gaussian produces a large error whatever its shape or amplitude. Fits run to an iteration cap with early stopping on the regularised training loss; the benchmarks reported here use a 20,000-iteration cap and an early-stopping patience of 500 iterations (Rate-distortion analysis, below).

The loss function used for the single-pass benchmarks reported here (the loss ablation of Supplementary Document 5 and the residual passes of progressive fitting, Supplementary Document 3, are the exceptions) is:

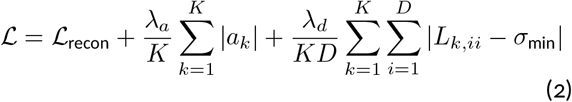

where ℒ_recon_ is the L1 reconstruction loss between the predicted and target volumes, and *λ*_*a*_ = 0.1 × lr and *λ*_*d*_ = 0.01 × lr (with lr = 0.01, yielding *λ*_*a*_ = 10^*−*3^ and *λ*_*d*_ = 10^*−*4^) regularise splat amplitude and extent. Both penalties are means over the *K* splats (and *D* diagonal entries) of the softplus-activated parameters (for the diagonal, the softplus term above the floor *σ*_min_), evaluated before the amplitude clamp *a*_max_, so their weight does not grow with *K* as the splat count is swept. The implementation also exposes an optional boundary-containment term *λ*_*b*_ℒ_boundary_ with 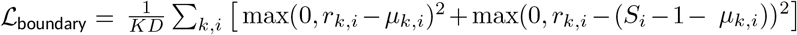 where 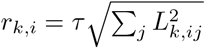 is the effective radius of splat *k* along dimension *i* at truncation threshold *τ* and *S*_*i*_ is the volume extent. This term is disabled (*λ*_*b*_ = 0) for all benchmarks reported in this paper. We adopt L1 over MSE based on a comparison of MSE, L1, and Poisson-deviance reconstruction losses across 17 volumes (Supplementary Document 5): on held-out PSNR, L1 beats MSE on 11 of them (by between +0.20 and +7.78 dB) and never trails it by more than 0.28 dB, and MSE is not the best training-PSNR loss for these regularised, early-stopped fits (best on only one volume).

### Dynamic operations

Every 50 iterations, splats are redistributed via fixed-pool relocation. A per-splat importance score *a*_*k*_ *П* _*i*_ *L*_*k,ii*_ (amplitude times approximate volume) identifies the least informative splats; the bottom 1% that sit at low residual (at most 64 per step, one per detected residual peak) are relocated to local maxima of the positive reconstruction residual 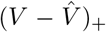, found with non-maximum suppression over the volume. Relocated splats are reinitialised isotropically at *σ* = 0.5 voxels and their Adam first- and second-moment estimates are zeroed; all other splats continue uninterrupted. A short cooldown (≥ 1 relocation step) prevents the same splat from being relocated again before it has had a chance to adapt. Every 1,000 iterations, splats are reordered along a Morton (Z-order) curve to improve GPU cache locality. After fitting completes, a single pass of conservative culling removes the splats accounting for the lowest fraction of cumulative amplitude (5% by default in the Python API; the shipped presets and the benchmarks reported here use the conservative 0.1%, i.e. 99.9% retention).

### Seed generation

Initial splat positions come from the volume itself. The default (auto) seeder, used for every single-pass fit reported here, draws 60% of the seeds from edges and 40% from a uniform grid. Sobel gradients are computed across all dimensions, and voxels whose normalised gradient magnitude exceeds 0.1 are sampled by gradient-weighted Poisson-disk sampling with a 2-voxel exclusion radius. The grid, whose spacing is set so that its 40% share of the seed budget tiles the volume (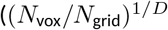, at least 2 voxels), guarantees coverage of smooth regions the gradient misses. Two further seeders are available and combinable: a scale-hierarchical decomposition seeder, and an intensity-weighted *peaks* seeder used on sparse residuals in progressive fitting (Supplementary Document 3). The seed target defaults to max 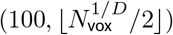 capped at 10,000 (the rate-distortion sweeps override this with explicit counts). The retained count falls below the requested seed count after the post-fit cull (Dynamic operations, above). Seeding runs on the GPU (CUDA, MPS) when one is available.

### Tiled fitting

For volumes exceeding GPU memory, Luxar partitions the volume into overlapping tiles on a regular grid with stride *T* − *L*, where *T* is the tile size and *L* is the overlap. Each tile is fitted independently with the same optimiser configuration. Before fitting, each tile’s voxel data is multiplied by a raised-cosine (half-Hann) taper 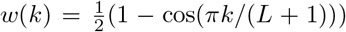 for *k* = 1, …, *L*, where *k* indexes voxels within the overlap region, applied along each overlap face, producing a partition-of-unity: adjacent windows sum to 1.0. The first and last tile on each grid axis retain unit weight on their outer face. The constraint *L* ≤ *T/*2 prevents triple overlap. The last candidate tile of an axis is folded into its predecessor when its unique coverage is thinner than the overlap, and an integer seed budget is divided over the tiles in proportion to their Hann-weighted voxel count times their mean windowed intensity above the floor raised to *α* = 0.44, with a quarter-share floor (Supplementary Document 11, Sec. 5.2). After fitting, the tile splat sets are unioned: by default one part per tile in a `kind=partition` tree, or a single flat leaf with `–flat`. The batch fitter (`luxar gsplat batch-fit`) drives this at scale: `run` executes a tile plan across the GPUs of one machine (one worker per GPU, or several per card sized from free memory) and `submit` submits the same plan as a Slurm array job, packing tiles per task and sizing tiles from a GPU profile. Per-tile outputs are written independently, so a run resumes by skipping finished tiles, and `validate detects` corrupt or stale tiles for re-fitting. `merge` streams the tiles into one partitioned scene without holding the volume in memory, optionally building a level-of-detail ladder per tile as it goes; its peak host memory is one tile’s splats over all timepoints (a recording fitted as one tile per frame becomes a single time-stacked leaf, so for it the whole archive). In the controlled tiling comparison of the Results (Supplementary Document 11; confocal blastocyst and light-sheet *Tribolium* embryo, tiles of 64 to 256 voxels, overlaps of 0–40%), the seam-to-interior residual ratio stays within 1.01–1.13 of the seamless fit’s at 5–20% overlap, at every tile size on both volumes (one fit per cell, no seed replicates), and the tiled union stays within −0.98 to +0.19 dB of the monolithic fit’s PSNR there. A hard cut without overlap raises the near-seam error itself up to 50-fold on the embryo, and overlapping tiles without the taper reach a seam-to-interior ratio of 3.91–5.10. The finest grids (81 and 90 tiles of 128 voxels, a median of 354– 434 seeds each) cost 0.78–0.98 dB (Supplementary Document 11, Sec. 7.5). OME-Zarr inputs are read directly, each timepoint and channel becoming its own task. We also evaluated progressive multi-pass fitting, where a nominal splat budget is distributed in equal shares across sequential passes, each fitting the residual left by the frozen earlier passes. At a common nominal 32K-seed budget and a common post-fit cull, single-pass fitting reaches a higher training PSNR than the progressive (residual-pass) fitter on 15 of 17 volumes (by 0.07–6.02 dB over 2-pass; 2-pass is ahead only on Mouse heart nuclei (+0.40 dB) and Zebrafish embryo nuclei (+0.85 dB)), while four or more passes trail on every volume; the smallest of those margins, Fly brain neurons (0.07 dB), is below the largest run-to-run spread measured in the replicates (0.093 dB), although on that volume itself the two conditions’ ranges do not overlap. On an otherwise idle GPU the two conditions take comparable time (1-pass/2-pass wall-time ratio 0.70–1.75 ×, median 1.07×), and single-pass also has the lower worst-case error on 12 of them (Supplementary Fig. 2; Supplementary Document 3).

### Rate-distortion analysis

For each benchmark volume (Table 3), we fitted Gaussian splat models at 10 splat counts geometrically spaced (doubling from 1K to 512K), extended to 1M and 2M splats on the four light-sheet volumes (*Tribolium*, Mouse heart, Zebrafish, *Drosophila*) under the same 20,000-iteration cap. All fits used conservative culling (99.9% amplitude retention), 20,000 maximum iterations, early stopping patience of 500 iterations, L1 loss, and enabled dynamic operations. PSNR was computed as 10 log_10_(1/MSE) on volumes normalised to [0, 1] (equivalent to the standard 10 log_10_(MAX^2^/MSE) with MAX = 1); MSE is averaged over all voxels including background. SSIM^50^ was computed with a window size of 11 and Gaussian weighting (*σ* = 1.5).

### Compression ratio

The compression ratio is defined on disk: CR = *s N*_voxels_*/B*_store_, where *s* is the number of bytes per voxel at which the source volume is stored (2 for the uint16 volumes, 1 for the 8-bit Mouse heart TIFF, 4 for the two deconvolved neuromast channels, which are float32 files; Table 3). The numerator *s N*_voxels_ is thus the uncompressed voxel payload of the benchmark volume at its stored sample depth. *B*_store_ is the summed size of the encoded chunk and metadata files of the fit’s `.gsplats.zarr` store as the default writer emits it and the viewer reads it (the outer ZIP container is not counted): `uint16` per-axis fixed-point centres, `uint8/uint16` log-encoded Cholesky factors and amplitudes on a bounded-linear `uint8/uint16` or geometriclog `uint16` grid chosen from the array’s dynamic range, Blosc-`zstd` chunks, plus the array and index metadata, fitting provenance excluded (Supplementary Document 12). *B*_store_ is read from the cached store of every sweep cell. Every PSNR in Table 1 and Fig. 3 is scored on the decoded store, not on the optimiser’s float32 parameters. The encoding costs a median of 0.003 dB, at most 0.15 dB at 32K splats and 0.72 dB over the whole sweep (Cells3D membranes at 512K), measured by rescoring every cached sweep cell (Supplementary Document 2); on fresh float32 fits the same encoding costs up to 0.679 dB (the *Tribolium* 2048K cell; at most 0.167 dB on the other reported cells; Supplementary Document 12).

Where a parameter count is more useful than bytes we also quote the encoding-agnostic ratio *N*_voxels_/(*K* × *F*_*D*_), with *K* the number of splats after culling and *F*_*D*_ = *D* + *D*(*D*+1)/2 + 1 float parameters per splat (10 for 3D); it is 3–164 × at the cross-validation optimum across the benchmark. The two ratios differ by the factor *sF*_*D*_*/b*, with *b* the measured bytes per splat of the shipped encoding (a median of 11.3 B over the 30 fresh-fit cells of Supplementary Document 12): 0.9–4.0 × (median 1.8 ×) across the benchmark, below 1 only for the 8-bit Mouse heart volume.

### Blind-spot cross-validation

We adapted the blind-spot cross-validation framework of Batson & Royer^23^ for Gaussian splat model selection. For each dataset and splat count, 5% of voxels were randomly selected (numpy `RandomState` with seed 42, shared across the GSplat and H.265 pipelines so all methods are scored on the same held-out voxels) and replaced with the median of their 26 neighbours in the 3 × 3 × 3 donut, the centre voxel excluded. In the canonical sweeps of Table 1 and Supplementary Document 2 that median runs over all 26 neighbours, other held-out voxels included (72–74% of held-out voxels have one); the codec baselines below use the same fill; the released code implements a stricter fill that also excludes other held-out voxels, so that the filled volume is a function of the unmasked voxels only. Re-sweeping four regime-spanning volumes with the stricter fill changed held-out PSNR by at most 0.25 dB and moved no optimum (Supplementary Document 2). Gaussian splats were fitted to this modified volume. The train PSNR was computed on unmasked voxels against the original volume; the held-out PSNR was computed on the masked voxels against the original (pre-replacement) values. The splat count at which the held-out PSNR peaks identifies the point beyond which additional splats memorise noise instead of capturing signal. This model-selection procedure is packaged as a single command, `luxar gsplat cal`, which sweeps splat count, reports the recommended *K*^*⋆*^, classifies the held-out curve (peak, plateau, or signal-limited), and estimates the ensemble noise-floor PSNR ceiling (Supplementary Video 5).

### Regimes of the held-out curve

The blind-spot guarantee holds only when a masked voxel’s noise is unpredictable from its neighbours, so that fitting it can only lower the held-out score. Any processing that mixes neighbouring voxels (multi-view fusion, deconvolution, denoising, resampling) correlates the noise and voids that guarantee; a model can then keep improving the held-out score by reproducing correlated noise, and the curve need not turn down. Of the light-sheet volumes whose held-out curve levels off, only the *Drosophila* frame is not deconvolved (its SiMView camera views are registered and fused, but no deconvolution spreads each voxel’s noise over its neighbours), and it is the one that plateaus cleanly; the Mouse heart volume, also undeconvolved, is instead signal-limited. For *Tribolium*,^15^ multi-view deconvolution could introduce such spatial correlations; its Laplacian and Haar noise estimates agree to within 5%, but that agreement measures a noise level and does not test independence between voxels, so we read the volume as signal-limited under the present model class and leave open whether correlated noise contributes to its rising curve. For Mouse heart nuclei^32^ the noise level is not estimable (every median-absolute-deviation (MAD) estimator returns exactly zero, which bounds nothing) and the model is signal-limited under L^1^ + bounded splat parameters, not noise-limited; the held-out curve keeps rising even when the sweep is extended to 2M splats and the iteration budget is tripled. *Tribolium* behaves the same way: its held-out PSNR is still rising at 2M splats (55.5 dB, 5.5 dB below its 61 dB noise ceiling), with the train and held-out PSNR of that blind-spot fit within 1.1 dB of each other. The deconvolved Zebrafish embryo instead reaches a finite optimum, a held-out plateau from 256K, so it is not signal-limited, and the *Drosophila* frame plateaus from 256K splats and is flat within 0.05 dB from 512K to 2M; the one confocal volume without a held-out decline, the sparse Fly brain tile, plateaus in the same way (Supplementary Document 2).

### Noise floor estimation

For each dataset, we estimated the noise standard deviation *σ* using three robust MAD-based estimators (in the style of Immerkær^51^ and Donoho & Johnstone^52^): (1) *Laplacian MAD* applies the standard *D*-dimensional discrete Laplacian (kernelnorm 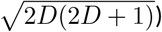) and scales the MAD of the response by 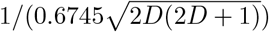. (2) *Haar finite-difference MAD* uses per-slice diagonal second differences *d* = *I*_*y,x*_ − *I*_*y,x*+1_ − *I*_*y*+1,*x*_ + *I*_*y*+1,*x*+1_, for which *σ* = median(|*d*|)/(0.6745 × 2). (3) *Background MAD* is taken on voxels below the 10th intensity percentile (when enough background voxels are present). The ensemble estimate is the median of whichever estimators are available for a given volume. The noise floor PSNR is 10 log_10_(1*/σ*^2^).

### Compression baseline comparison

We compared Gaussian splats against H.265 video compression (libx265 via FFmpeg, treating Z-stacks as video sequences) using the same blind-spot cross-validation framework^23^ applied identically to both methods. For each dataset the same seed-42 mask as above was filled with the median of all 26 donut neighbours, held-out donors included, as in the canonical sweeps; the released code excludes held-out donors, which moved held-out PSNR by at most 0.25 dB on the four re-swept volumes (Supplementary Document 2). Both GSplats and H.265 were then applied to this modified volume: GSplats were fitted to it (the optimiser sees donut-filled values at masked positions); H.265 encoded it at constant-rate-factor (CRF) values {0, 5, 10, 15, 20, 25, 30, 35, 40, 45, 51} with `libx265` preset `medium`, 16-bit gray (`gray16le`) input frames, and 12-bit gray (`gray12le`) internal pixel format. CRF = 0 enables x265’s lossless mode (`x265-params lossless`=1) on the 12-bit converted frames, so it is lossless for that input, not for the 16-bit source. On systems whose `libx265` build lacks 12-bit gray support, encoding falls back to 8-bit `yuv420p`; we verified this fallback was not engaged on the runs reported here.^*^ In both cases, the held-out PSNR was computed by comparing the output at masked positions against the *original* (pre-replacement) values, measuring each method’s ability to recover the true signal from surrounding context. Both methods are spatially aware (GSplats: global mixture model; H.265: block-based prediction and DCT), making the blind-spot framework applicable to both. The matched-rate comparison of the Results reads H.265 at the bits per voxel of the Gaussian-splat operating point, interpolated between adjacent CRF settings and never extrapolated. Under the envelope rule that credits H.265 with the larger of its best held-out result at any rate up to that bit budget and its interpolated value at that rate, Gaussian splats lead on 15 of 17 volumes and tie within the 0.2 dB replicate band of Supplementary Document 2 on 2 (median +0.98 dB; *Tribolium* at −0.09 dB; Supplementary Document 7).

### Scene graph and layers

A Luxar dataset is organised as a typed scene graph: a Scene root that owns the dimension definitions and writer, interior Group nodes that bundle children and may carry transforms, and leaf data nodes `Points, Lines, Mesh,` and `GSplats`. Each node persists a small set of attributes to its Zarr group: a 4 × 4 `transform`, a per-dimension `nd_transform`, rendering uniforms (`opacity, absorption, gamma, intensity, offset, blending_mode, colormap), visible, extend_to_all` (controls visibility along non-displayed dimensions), an optional `layer_order` that fixes the draw order between layers, and a layer flag. The root carries the dimension definitions and an optional structured dataset citation, so attribution travels with the data. Metadata is streamed to disk as nodes are added, so the in-memory scene graph never holds bulk data and datasets exceeding RAM compile naturally on a laptop. Nodes flagged `layer=True` populate the viewer’s Layers panel, where users adjust visibility, opacity, absorption (volumetric *κ*), display range, gamma, blending mode, and colormap per layer; display-range edits are converted on the fly to shader uniforms (intensity = 1/(max − min), offset = −min/(max − min)). Multi-select (shift/ctrl) propagates uniform changes to all selected layers, and groups can be exposed as composite layers, fanning controls to all data descendants.

### Spatial and nD transforms

Spatial 4 × 4 transforms compose hierarchically along the parent chain (the leaf’s transform is applied to coordinates first and the root’s last, *M*_world_ = *M*_root_ · · · *M*_leaf_) and are stored column-major in Zarr for Three.js compatibility, with transposition at the Python ? Zarr boundary. Non-displayed dimensions support per-node transforms of two kinds: continuous/discrete affine {scale, offset} and categorical {permutation}, with composition *s* = *sV*_*p*_*s*_*c*_, *o* = *s*_*p*_*o*_*c*_ + *o*_*p*_ for affines and *r*[*i*] = perm_*p*_[perm_*c*_[*i*]] for permutations. At query time the viewer never transforms point coordinates: the spatial index stores raw values, and the slice query is inverse-transformed per dimension instead of per point (constant work for an affine, one pass over the category list for a permutation). For an affine, raw_pos = (world_pos −offset)/scale and raw_tol = world_tol/ |scale|; for a permutation, inverse[perm[*i*]] = *i*; on a discrete axis the world value is rounded onto the declared grid and a value with no local preimage selects nothing, and a zero scale, accepted with a validation warning, leaves the query untransformed.

### Data encoding, indexing and lazy loading

Fitted splats are written to Zarr^9^ archives with semantic encoding (8 semantic types) under a width-aware Blosc-zstd level-9 policy: byte shuffle for multi-byte integer codes and no shuffle for single-byte codes and floating-point arrays (Supplementary Document 12). An archive is a directory of chunk files, or, when a single file is more convenient to move or host, the same store compiled directly into an uncompressed .`luxar.zarr.zip`, which the viewer reads in place: each chunk is one member of the archive and is fetched by HTTP range request without unpacking. Any static HTTP host that allows cross-origin reads (and range requests for the zipped form) can serve it. Meshes, whose faces are not independently meaningful, are chunked and compressed but loaded whole. Spatial indexing uses compound ordering: elements are sorted first by their discrete non-spatial coordinates (time, channel) and, within each such slice, along a space-filling curve over the spatial coordinates. Both Hilbert and Morton (Z-order) curves are supported; Hilbert provides superior spatial locality and is the default for serialisation, while Morton is used for in-memory reordering during fitting due to its lower computational cost. This ordering ensures that spatially proximate splats lie in adjacent chunks, so chunks selected by the slice-intersection query coalesce into a small number of HTTP range requests; tighter chunk extents also make the intersection test more selective. Chunk-level axis-aligned bounding boxes (AABBs) are precomputed and stored as metadata, and at load and on every view change a fast AABB test against each chunk’s precomputed nD extent (slice position ± tolerance per dimension) selects only the chunks overlapping the current view for fetching. Because the non-displayed dimensions lead the ordering, the rows of one value of a discrete axis such as time are contiguous, so a time step fetches the chunks of the new timepoint (plus the prefetched first chunks of its neighbours). A chunk that spans several timepoints brings its neighbours along, a level-of-detail ladder built per timepoint keeps the property rung by rung, and the prefetcher below adds the adjacent slice (Supplementary Document 14). In the layout the demo site publishes (Supplementary Table 2), whose 1 MB chunks hold many frames each, a time step costs 1 request for the *Drosophila* scene and 1 for the neuromast, and one time part (17.91–17.96 MB) in 65 requests for the zebrafish, whose 2.4M-splat frames are each their own part. That economy is bought at the opening: the *Drosophila* scene opens in 19.06 MB, against 5.27 MB at the compiler’s 64 KB chunks, which in turn make a step cost 266–268 requests. Where a scene carries a level-of-detail structure, the viewer streams additive ladders in prefix order, selects the coarsest substitutive level whose projected median splat footprint stays at or below 1.5 CSS pixels (the finest only when none does, with a 10% hysteresis against coarsening), and frustum-culls partition tiles, so a coarse or off-screen region is never fetched at full detail. Where no footprint is stamped, or the stamped dimensions are not the displayed ones, it falls back to the screen-area fraction the node’s bounding box projects to (Supplementary Document 10; the levels and ladders themselves are Supplementary Documents 8 and 9; Supplementary Video 6). Additive ladders are ordered by the greedy ordering of Supplementary Document 9 up to 5,000 splats and the *O*(*N* log *N*) self-energy ordering above that, for which no approximation guarantee is claimed (in the Supplementary Document 9 experiments greedy lowers the residual area under the loading curve by 0.2–2.6% relative to it). Fetched data passes through a multi-level cache: decoded per-slice geometry and decoded chunks in memory, compressed chunks in memory and in the browser’s origin-private file system, and HTTP range requests behind them. After an on-disk hit or a remote fetch a prefetcher requests the chunk on either side in the ordering, and an idle-time prefetcher warms the adjacent slice, hiding remote-fetch latency (of the order of 100 ms) when the user orbits or plays a slider (Supplementary Video 4).

### Web viewer architecture

The Luxar viewer is a TypeScript application built on Three.js,^53^ with two interchangeable rendering backends: WebGL (the default) and WebGPU, the latter driving the same materials through Three.js’ node-material system. The two were verified to render pixel-equivalent output. It runs both as a standalone served application and as an embeddable component (published as the @luxar/viewer npm package), exposed through a programmatic API (`LuxarApp`) with a scoped container and an event interface for integration into external web applications. Its interface (a control rail whose panels cover layers, rendering, dimension navigation, settings and recording, with every shortcut listed in a help overlay) is walked through in Supplementary Video 12. The same API is the viewer’s remote-control surface: camera flights (`flyTo`), rendering and per-layer setters, a full state snapshot and a stream of camera and dimension events are callable from the host page. `luxar serve --control` relays them over a WebSocket so that a Python script, an exhibit’s touch panel or an agent can drive a viewer on another display (`luxar.control.Viewer`). Scenes can also author *story waypoints*: a camera pose bound to a position along the hidden dimensions by the same visibility rule overlays use, so a hidden discrete “story” dimension carries a guided tour: the camera flies between poses, dimension-bound overlays can be timed to appear when the flight lands, and a controller need only step that dimension. The viewer is usable on phones and tablets: the canvas owns its touch gestures (one-finger orbit, pinch zoom, two-finger pan and twist-to-roll, with a touch vocabulary for fly mode as well), a tap picks and a long press opens the element’s menu. The layout adapts to a coarse pointer and to the device’s safe areas, and rendering budgets are clamped to mobile memory. Splat data is loaded progressively via HTTP range requests into a four-level cache: L2 (the browser’s persistent Origin Private File System, OPFS, a fixed 2 GB surviving browser restarts), L1 (compressed in-memory chunks, 100 MB ceiling), L0 (decompressed in-memory chunks, 200 MB ceiling), and above them an S-cache of fully decoded slice ladders that lets a revisited slice skip the chunk path entirely. The three in-memory tiers are heap-aware: L0 and L1 shrink from their configured ceilings when the heap is tight, and the S-cache grows with the heap from its 128 MB default up to 2 GB. Blosc/zstd chunk decompression runs in the Zarr client’s WebAssembly (WASM) codec; Luxar’s own WASM-accelerated *n*-dimensional compute kernels (effective radii, Mahalanobis distance) support up to 16 dimensions, falling back to a TypeScript implementation beyond that. For *n*-dimensional datasets, the viewer displays three spatial dimensions and navigates non-displayed dimensions via sliders (continuous) or toggles (categorical/boolean; Supplementary Video 2). Each geometry type resolves slice membership differently: point clouds intersected by the display hyperplane take an effective radius 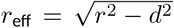, where *d* is the Euclidean distance to the hyperplane; Gaussian splats are attenuated by their marginal density along the hidden dimensions (a shifted Gaussian in the Mahalanobis distance from the splat centre to the slice). Line segments are clipped against the *n*D slab, with every attribute interpolated at the clip parameter, and a triangle is drawn only when all three of its vertices pass the slab membership test, a whole-triangle cull in place of *n*D polygon clipping.

#### Viewer performance measurement

The sweeps of Fig. 1g and Supplementary Document 6 run the web viewer in headless Chromium on the GPU via ANGLE/Vulkan (as for Supplementary Table 1) on an NVIDIA RTX 3070 at 1280 × 720, with the adaptive device-pixel-ratio (DPR) scaling pinned to 1.0. The device pixel ratio scales the render buffer along each axis, so the number of rendered pixels scales as its square; under load the viewer lowers it, rendering fewer pixels and upscaling the result, so that frame rate is held at the cost of sharpness, and pinning it to 1.0 reports the raw cost of rendering every pixel. The measurement window is 3 s at the reference nominal primitive size (4 px; lines 1.5 px), and the reported latency is that of a synchronised render-and-readback call, one render plus a readback with the pipeline drained. On the three heavy size cells the callback window is a rendering phase of callback intervals of 67–300 ms, 1.6–1.9 times the call, followed by idle cap ticks; whether each interval is one long frame or two frames of the call’s length the records do not say, and neither number there is a delivered frame rate. The deposited sweep does not record the harness settings it ran with, so the DPR pin and launch flags are asserted from the deposited harness, not evidenced by the sweep (Supplementary Document 6). At *N* = 10^5^ the call completes in 1.21 ms (lines), 1.23 ms (points) or 1.72 ms (Gaussian splats); at *N* = 10^7^ it takes 18.8 ms (points) and 19.4 ms (Gaussian splats). In production the adaptive DPR system steps the render scale down (to a floor of 0.5) whenever the measured rate falls below three quarters of the display’s refresh rate; the sweeps were run with it pinned, so the recovered rate is not measured.

### Geometry primitives and mesh shading

Points and Gaussian splats are drawn as camera-facing sprites carrying an analytic radial profile (Supplementary Video 17). Lines are drawn as *capsules*: a rotationally symmetric profile of the two-dimensional point-to-segment distance in screen space, whose joints are composed by a deficit rule so that a corner neither doubles its own brightness nor opens a gap. A per-node size policy sits above this choice: the capsule is used throughout except for line nodes whose effective segment load (the authored segment count scaled by the on-screen width its widest segment opens to at the opening framing) reaches two million, where the cheaper screen-space quad is built instead. The policy can be pinned to either primitive. The capsule is preferred over an exact volumetric primitive (the closed-form convolution of a segment with a three-dimensional Gaussian, integrated along the view ray) because it matched or beat it in a blinded visual comparison, including in the near-axial case, while the exact primitive costs 2.7–5× the screen-space quad per frame. The capsule and the quad are the two shipped primitives. The line primitive in force for the Supplementary Document 6 sweep is not recorded; the automatic policy selects the capsule below two million effective segments, which leaves only the 16-pixel size cell and the 10^6^-vertex cell uncertain, and the two primitives differ by 1.04–1.11× on an Apple GPU and about 1.5× on thin lines on a discrete NVIDIA GPU in the viewer’s own linematerial benchmark; Supplementary Document 6 quotes those figures and did not measure them. Meshes are the only shaded geometry type (the other three are purely emissive) and are lit by a light-free headlight anchored to the view, so a surface reads as solid without the scene having to define lights; stored vertex normals carry a companion attribute naming which three dimensions they describe. A mesh takes its base colour from exactly one source: per-vertex colours, a colormap over per-vertex scalars, or an image texture (optionally GPU-compressed as KTX2/Basis). A texture is paired with per-vertex coordinates in the same way normals are paired with their dimension names: either alone is refused, not merely warned about, since either would render something plausible and wrong. A mesh may instead opt into a physically based material (`material=“physical”`): roughness, metalness, clearcoat, iridescence and sheen give brushed and polished metals or a pearlescent shell. A transmission family (index of refraction, thickness, attenuation, dispersion) gives glass, which refracts the background and other meshes and, on request, the points, lines and splats behind it while the data in front of it stays crisp. Such a mesh is lit by a scene environment: a neutral room by default, an equirectangular HDR image, or a cube map captured from a probe inside the scene itself, so that a chrome sphere reflects the point cloud around it. The capture can be baked into the archive (`luxar env` bake) so a published scene never pays for it on a visit. Because a triangle’s extent is implied by its own vertices, a mesh contributes no size padding to the scene bounds, and because a face chunk is not independently meaningful, a mesh is loaded whole, not through the spatial index. Picking resolves to the vertex, not the face. Meshes support substitutive LOD levels and spatial partitioning, but they refuse two things by construction: volumetric blending, because a triangle has zero thickness and so presents no path length for absorption to attenuate; and an additive ladder over an arbitrary face order, because a prefix of such an order is a surface with holes, not a coarser surface. Only the spatially coherent reveal ordering yields a presentable prefix.

### Overlays, sound and hover annotations

The viewer carries a screen-space annotation layer atop the WebGL canvas, configured from a dedicated `overlays`/ group in the Zarr archive. Four overlay types are supported: `overlay_text` (rendered via `textContent` for XSS safety), `overlay_image` (viewport-relative sizing), `overlay_video` (a WebM or MP4 clip stored verbatim in the archive), and `overlay_html` (rendered after a client-side allowlist sanitisation that strips on* handlers and `javascript:` URLs). A video overlay’s container signature is checked at authoring, playability remaining the browser’s; it is played muted in a loop and paused whenever its visibility range does not match, so clips with disjoint visibility ranges cost one decode at a time. Transparency travels as a stacked alpha matte, colour above and alpha as a grey matte below, that the viewer recombines in a shader, since browsers drop a codec’s alpha plane inconsistently. Overlays support 9-point anchoring, opacity, `CSS mix-blend-mode`, fade transitions, and dimension-aware visibility via `visible_rang`e (perdimension exact values with tolerance, or [min, max] ranges). A scene can be heard as well as seen: a sound node stores an MP3 or AAC clip (an ambient bed, a positional source that follows a node, or a narration cued when a story waypoint is reached) and the viewer plays it through Web Audio, with the same hidden-dimension slab rule deciding audibility that decides which elements are drawn. A sound is a node of the scene graph but not a geometry, so it carries no appearance and never stretches the scene bounds. Hover annotations are driven by GPU picking: an RGBA32F render target at half the canvas resolution (capped at 1024 pixels on its longer side) encodes (R) node ID, (G, A) the low and high 16 bits of the element ID, and (B) element brightness. Fragment depth is 1 − brightness for points, lines, and Gaussian splats and meshes in the additive, max and luminous modes, so the brightest element at each pixel wins via depth testing, and the projected surface depth for Gaussian splats and meshes in the normal and opaque modes. A pick event dispatches a ray–AABB cull, then reads back a 5 × 5 neighbourhood around the cursor and applies brightness-weighted majority voting; the buffer is cached and re-rendered only when the view, geometry, appearance or viewport change and both cursor and scene have been still for 120 ms, making hover events on a static scene effectively free. Per-element metadata is stored in compressed-sparse-row layout: `label_offsets` (uint64) plus `label_bytes` (UTF-8) for text labels, and `image_label_offsets` plus `image_label_bytes` for pre-compressed JPEG/PNG/WebP blobs decoded to blob URLs with a 50 MiB LRU cache. When label arrays are present, default**_** hover_textand**_** hover_image overlays are auto-injected on scene finalisation; their templates substitute `{hover_key}`, `{hover_label}`, `{hover_image_label}`, `{hover_node}`, and `{hover_index}` on each pick (Supplementary Video 10). A picked element can also be made actionable: a node’s link and copy templates use the same placeholders, so a click opens the element’s URL (for instance a database entry for its label) and a right-click offers to copy its text or link.

### Post-processing pipeline

Rendering uses a 16-bit floating-point HDR pipeline. The scene is rendered into a HalfFloat target (optionally multisampled; MSAA, SSAA and FXAA are all off by default), followed by HDR-domain effects (mipmap-based bloom, chromatic lens distortion, physically based detector noise; off by default), a tone-map pass, an LDR-domain effect (vignette), and, when enabled, FXAA. The tone-map pass selects among LINEAR, REINHARD, CINEON, ACES filmic (the default), AgX, and NEUTRAL operators (None is an alias of LINEAR; both clamp to [0, 1]), with exposure (color × 2^*E*^), additive offset (clamped at zero), and gamma fused into the same shader to avoid an extra full-screen pass. Bloom and chromatic distortion are composited within this single fused fragment pass, with bloom mixed into the HDR sample before the per-channel chromatic sampling, instead of as separate library passes. Adaptive devicepixel-ratio scaling monitors a 1 s sliding window of frame timestamps and adjusts the render-target resolution using thresholds set relative to the display’s estimated achievable refresh rate (not fixed frame-rate constants), with a probe-and-verify guard that reverts a downscale if it does not actually improve throughput; it is locked during recording to prevent mid-capture resolution changes (Supplementary Document 10).

### Animation and recording

Each non-displayed dimension supports independent animation at FPS presets {0.5, 1, 2, 5, 10, 15, 30, 60, 120} or a custom rate, with three loop modes (once, loop, bounce); continuous dimensions are linearly interpolated over a configurable traverse time (10 s by default), while discrete dimensions step by their declared step size. Animation advance is gated by completion of asynchronous chunk loading so that every displayed frame has its data; actual FPS is measured per second and a warning is emitted when it drops substantially below target. A scene also carries its opening camera and appearance: an authored pose or a named lens preset, an optional turntable about any screen or scene axis with a slow dolly that breathes in and out, and the rendering settings a visitor first sees, so a published link opens on the view its author chose. The recording panel produces still images (PNG, WebP, JPEG, EXR), real-time video (WebM via `MediaRecorder`, VP8/VP9), offline video and turntable rotations (H.264, H.265, VP9, VP8 via mediabunny WebCodecs), and frame-sequence ZIPs (PNG/WebP/JPEG/EXR). A generated `ffmpeg` script accompanies each sequence: its active command encodes an SDR H.265 master, and its commented-out block, once a nominal peak luminance is chosen, produces 10-bit BT.2020/PQ HDR video from the exported EXR frames. HDR EXR captures sample the linear HalfFloat pipeline *before* tone-mapping (by default with bloom kept when it is enabled, lens distortion and detector noise switched off, and alpha set to 1; a raw-scene mode omits bloom too), preserving the full dynamic range. During a capture adaptive DPR and resizing are locked, and offline sequences run a keep-alive callback so the animation loop does not idle.

### Hardware and reproducibility

All fitting benchmarks were run on a single NVIDIA RTX PRO 6000 Blackwell (96 GB VRAM, sm_120) with PyTorch CUDA; viewer-performance measurements use a separate NVIDIA RTX 3070 (Supplementary Document 6). Fitting wall-time was additionally measured on that consumer RTX 3070 (8 GiB) across a five-count sweep whose fixed 32K reference budget is the basis of the “minutes per volume” figure; twelve of the seventeen Table 1 operating points lie on that sweep’s grid and the five at 8,000, 128,000 and 2,048,000 splats were not timed there (Supplementary Document 13). Each splat-count sweep cell uses its target seed count as the deterministic random seed for `torch.manual_seed` and `numpy.random.seed` (so the 8K-splat fit uses seed = 8000, the 16K fit uses seed = 16000, etc.); the blind-spot mask uses a separate fixed seed of 42 shared across pipelines (above). Edge-based seed generation and PyTorch CUDA operations can introduce small run-to-run variability; scripts that support repeated fits report replicate-level variation in their supplementary documents, and deterministic seeds are recorded where fixed. Viewer renderings (Fig. 1b,c) are frames captured from the shipped viewer in headless Chromium on a real GPU (ANGLE/Metal on an Apple M4 Max for the released figure; ANGLE/Vulkan on Linux), never from the software rasteriser, and are used as rendered apart from the camera pose, the time slice and, for the *Drosophila* frames of Fig. 1c, an absorption of 0.91. Every capture argument is recorded with the figure sources. The demo stores are generated by scripts in the sibling Luxar repository (https://github.com/royerlab/luxar); set LUXAR_DIR to override the default sibling path, or set LUXAR_SKIP_SCREENSHOTS=1 to skip capture and compose from the committed frames. The Luxar software used throughout, including the fitting, calibration, encoding, level-of-detail and export tools, is released at https://github.com/royerlab/luxar; the benchmark volumes and the fitted demo archives are the Zenodo records listed under Data and code availability.

## Supporting information

supp_doc_01_merging_error_analysis

supp_doc_02_splat_count_vs_quality

supp_doc_03_progressive_vs_single

supp_doc_04_convergence

supp_doc_05_loss_comparison

supp_doc_06_viewer_performance

supp_doc_07_compression_comparison

supp_doc_08_substitutive_lod

supp_doc_09_additive_lod

supp_doc_10_scene_graph_pipeline

supp_doc_11_tiled_fitting

supp_doc_12_codec_selection

supp_doc_13_consumer_gpu_timing

supp_doc_14_timelapse_splats

## Acknowledgements

We thank Kyle Harrington for discussions. We thank Merlin Lange for the zebrafish embryo light-sheet recording, and Adrian Jacobo and his laboratory at Biohub for the two-channel neuromast iSIM recording. The *Drosophila* embryogenesis time-lapse was acquired by Philipp J. Keller in the Keller lab at HHMI Janelia Research Campus, and we thank him for permission to redistribute a fitted representation of it. We are grateful to the teams who made the public datasets used here available: the OpenCell project, the FlyLight project at HHMI Janelia Research Campus, the Acto3D project, the Cell Tracking Challenge, the Image Data Resource, and the Allen Institute for Cell Science and the contributors of the scikit-image sample data. We thank the open-source communities behind Zarr, Three.js,^53^ PyTorch, NumPy and napari.^4^ Funding for this work was provided by Biohub. We thank the Biohub donors, Priscilla Chan and Mark Zuckerberg, for their generous support.

## Data and code availability

Luxar is open source under the BSD 3-Clause licence: https://github.com/royerlab/luxar. Installation: pip install luxar. The software is archived at Zenodo (concept DOI 10.5281/zenodo.22908222, resolving to the latest release).^54^ A gallery of more than 80 demonstration scenes is served interactively at https://demos.luxarviewer.dev, rebuilt from the same archives cited below. A hosted copy of the viewer is available to everyone at https://luxarviewer.dev: it opens any publicly reachable .`luxar.zarr` archive named in its URL (https://luxarviewer.dev/?src=<archiveURL>), so a compiled scene on any static host can be shared as a link with no server of one’s own. Nothing is uploaded; the browser fetches the archive directly from the host named in the link. Every gallery scene has a stable link of this kind, for example https://demos.luxarviewer.dev/d/gsplats_3d_cells3d_multichannel for the multichannel Cells3D volume.

The 17 benchmark volumes and their provenance are listed in Methods (Table 3). All but three are drawn from public repositories: the Zebrafish embryo nuclei (*h2afva*) recording was acquired by Merlin Lange, the Neuromast recording in the Jacobo laboratory, and the *Drosophila* SiMView recording^11^ in the Keller laboratory (Acknowledgements). The exact benchmark volumes as consumed by the analyses, sixteen of the seventeen (the Mouse heart nuclei volume is excluded because its source states no reuse terms), are deposited at Zenodo (DOI 10.5281/`zenodo.22682969`).^55^ That record as a whole is CC BY-SA 4.0 because it contains OpenCell material, on the same principle that splits the demo-data records into permissive and ShareAlike ones; each volume’s licence of origin is stated per volume in the record, and reuse of an individual volume may follow that more permissive licence.

The demo scenes shown in Fig. 1b,c and measured in Supplementary Table 1, along with the other bundled demo datasets, are archived as derived products in four Zenodo records; the FlyLight brain of Fig. 1c is instead rebuilt locally from the FISBe release.^14^ The records hold permissively licensed material (`DOI 10.5281/zenodo.21912279`),^33^ ShareAlike material whose derived products must themselves be relicensed CC BY-SA 4.0 (`DOI 10.5281/zenodo.21912281`),^56^ a 253-timepoint zebrafish histone-labelled time-lapse with a lighter 51timepoint fit (`DOI 10.5281/zenodo.21912283`),^57^ and a 500-timepoint *Drosophila* embryogenesis time-lapse (`DOI 10.5281/zenodo.22118694`).^12^ These are concept DOIs, which resolve to the latest version of each record: the demo products are refit as the fitter improves, and the current versions are the ones the demo site serves and Supplementary Document 14 measures. The benchmark-volume record is cited by concept DOI as well: the volumes it carries are byte-identical across its versions (their SHA-256 digests are recorded in the deposited manifest), so the concept DOI resolves to exactly the bytes analysed here while labels and descriptions can be corrected. These records hold *fitted* Gaussian-splat representations built for interactive visualisation: they are lossy, and are not a substitute for the source imagery in quantitative work. A few demo datasets are deliberately not redistributed, because their source licences do not permit redistributing even a derived product; those demos instead fetch the original data from its provider and refit locally. Every archived file is pinned by SHA-256 in the repository’s dataset manifest, so a download can be verified against the version cited here. The supplementary videos can be watched on the demo site (every mention in this manuscript links to the video’s page) and are deposited at Zenodo (`DOI 10.5281/zenodo.22825092`, a concept DOI resolving to the latest cut);^58^ the set is CC BY-SA 4.0 because two of the filmed datasets are ShareAlike, and the record lists each video’s licence of origin.

## AI use statement

Every line of Luxar’s code was designed, generated, debugged and tested with the help of Claude Code, to exacting engineering standards. The code base carries 33,497 automated tests (18,051 in Python, 206 of them exercising the CUDA kernels on GPU hosts, 15,323 in TypeScript, 118 in Rust and 5 in Go, plus 84 browser end-to-end specifications) at 91% line coverage, and the tests themselves are probed by mutation testing (Stryker). At 694,000 lines of Python, TypeScript, Rust, CUDA and Go, the project represents roughly 30 person-years of pre-AI software engineering at the author’s own rate of finished code, measured on his earlier projects; for a single author it would simply not have been possible without AI. All analyses, figures and tables were produced with the help of AI, and the text of this manuscript and its supplements was written with the help of AI, under the author’s direction and verification. The project, code, analyses and manuscript represent one year of the author’s work, and two years counting the ideation and reflection that preceded models capable of carrying it out. AI does more than make the mundane faster. It makes the impossible real.

## Author contributions

L.A.R. conceived the project, developed the software, performed the analyses, and wrote the manuscript.

## Competing interests

The author declares no competing interests.

## Supplementary Materials

### Supplementary Document 1 – Analytical merging error for Gaussian splats

Closed-form L^2^ integrated squared-error framework for merging two Gaussian splats, with geometric interpretation (relative merging error *ε* = sin^2^ *α* for the amplitude-optimised moment-matched template) and computational pseudocode.

### Supplementary Document 2 – Splat count vs. reconstruction quality

Systematic rate-distortion analysis across all 17 benchmark volumes with blind-spot cross-validation model selection, noise floor estimation, per-dataset quality curves, visual reconstructions, held-out validation, a cull-limited regime for the two iSIM neuromast channels whose realised splat count saturates beyond the operating point, a synthetic control on a known clean volume at two noise levels (the held-out and true-error maxima coincide), the effective-resolution analysis behind Supplementary Table 3, and the automatic residual-feature detection behind Supplementary Fig. 5.

### Supplementary Document 3 – Progressive vs. single-pass fitting

Comparison of single-pass and 2–8-pass residual fitting at a common nominal 32K-seed, 10K-iteration budget across 17 volumes (*n* = 1 per cell; retained counts 22,178–32,384). Single-pass wins on training PSNR for 15 of 17 by 0.07–6.02 dB over 2-pass (the smallest margin, Fly brain neurons (0.07 dB), is below the largest measured run-to-run spread of 0.093 dB); 2-pass wins on Mouse heart nuclei (+0.40 dB) and Zebrafish embryo nuclei (+0.85 dB); four and eight passes trail single-pass on every volume. On the six replicated volumes the 1-pass and 2-pass ranges are disjoint on six of the six. Wall time on an idle GPU is comparable (1-pass/2-pass ratio 0.70– 1.75×); single-pass has the lowest maximum absolute error on 12 of 17 and 2-pass the highest SSIM on 12.

### Supplementary Document 4 – Convergence behaviour of Gaussian-splat fitting

Convergence at 5 splat counts and 11 iteration checkpoints across 17 volumes, plus a blindspot sweep of 374 fits at each volume’s operating point and at a second count. 2,000 iterations leave training PSNR within 0.5 dB of its 20,000-iteration value on 77 of 85 volume–capacity cells and within 1 dB on 81 (largest shortfall 1.50 dB), and at each volume’s operating point held-out PSNR is within 0.2 dB of its peak by 2,000; under both metrics model capacity dominates optimisation budget.

### Supplementary Document 5 – Choice of reconstruction loss

Empirical and theoretical comparison of meansquared-error (MSE), L1, and Poisson-deviance reconstruction losses across 17 volumes, each fitted at its blindspot operating point plus a 5,000-splat margin (517,000 for the two signal-limited volumes) with *n* = 3 replications per cell. On held-out PSNR, L1 beats MSE on eleven volumes, by up to +7.78 dB (on the deconvolved iSIM Neuromast nuclei; +1.10 to +2.02 dB on the noisy confocal kidney channels), while MSE edges it by at most 0.28 dB on the other six. MSE is not the best training-PSNR loss for these stopped fits either: it has the highest training PSNR on only one volume (Poisson on nine, L1 on seven); Poisson deviance stops in the fewest iterations on eight volumes, with 7.6–13.3× fewer iterations than MSE on the three confocal volumes where MSE runs to the cap and Poisson does not. Explains why MSE is not the best training-PSNR loss for these regularised, early-stopped fits (the PSNR-optimality argument concerns the global minimiser) and states that the L1 adoption is an end-to-end result for one stopping rule and one L1-derived operating point.

### Supplementary Document 6 – Viewer rendering performance

Controlled count, size, and viewport sweeps for points, lines, and Gaussian splats (meshes are not covered by this sweep). Synthetic scenes vary *N* from 10^3^ to 10^7^ at fixed reference size, sweep object size from about one pixel nominal to thousands of pixels per element at fixed *N*, where the cost grows with covered area, and vary viewport from 720p to 4K. Reports the cadence-capped rAF callback rate and the synchronised render-and-readback latency (a 1-pixel gl.readPixels after each call; gl.finish and gl.fenceSync returned implausibly early in this harness and are not used), with three render-call variants (post_off, post_off_hdr, post_on) decomposing post-processing overhead into HDR-target cost vs. the fused post-processing pass (bloom and anti-aliasing off by default).

### Supplementary Document 7 – Gaussian splats vs. voxel codecs

Quantitative comparison of Gaussian splat compression against H.265 video coding and quantisation+zstd baselines, evaluated under blind-spot crossvalidation across all 17 volumes with a completed sweep, spanning spinning-disk, confocal, light-sheet and iSIM modalities. Includes a matched-rate comparison (H.265 read at the splat operating rate and the splats at the H.265 held-out peak rate, interpolated between measured sweep points and never extrapolated), a per-volume peak held-out PSNR table, and the very-low-rate regime against the voxel baselines: GSplats reach 0.002– 0.06 bits/voxel, at 21–38 dB against the original, 175– 3506× below the lowest rate the tested lossless blosc configurations reach on each volume at its stored dtype (47–9762× on the normalised float32 arrays) and 3.5– 407× elow the most aggressive quantisation+zstd setting, while the H.265 sweep’s lowest rates are comparable to the 1K-splat fits. Reports the compression ratio at the per-volume cross-validation optimum across the benchmark (range 6.0–340×, median 99.3×) and discusses the regimes where each method leads. Crediting H.265 with the larger of its best result at any rate up to the GSplat operating rate and its interpolated value there, GSplats recover more signal on 15 of 17 volumes and tie on 2 (median +0.98 dB); at the exact rate, on 16 of 16 (median +1.32 dB), while at H.265’s own, mostly lower, peak rate the codec leads on 9 of 12. Peak-to-peak, which compares at different rates, GSplats attain the higher blind-spot optimum on 12 of 17, by the most on the deconvolved iSIM neuromast channels, the Mouse heart and the noisy Kidney actin, and the two are within the 0.2 dB replicate band on5, the *Tribolium* volume among them (−0.09 dB), so no “H.265 wins on deconvolved light-sheet” rule holds.

### Supplementary Document 8 – Substitutive levels of detail for Gaussian splats

Cost-metric design space (*L*^2^, Cauchy–Schwarz, gKL, *W*_2_, MMD, *L*^*∞*^) for the substitutive-LOD problem, *K*-wise moment-matched merging with *L*^2^-optimal amplitude, three partitioning algorithms (greedy hierarchical à la Runnalls, cost-increment relocation, joint relaxation), the limits of spectral lower bounds (with an explicit counterexample), and connection to existing 3D Gaussian-splatting LOD methods. Numerical experiments on synthetic mixtures and on the full hosted DAPI nuclei fit: the summed per-bin error is, in the median, a faithful surrogate for the global error once a spatial partition separates the bins (single trials deviate by tens of percent at moderate separation); cost-increment relocation matches greedy merging within 5.5% at compression factors of four and above; on the full fit cost-aware merging beats mass-ranked culling, with and without an amplitude refit, while on sparse subsamples culling wins in the median.

### Supplementary Document 9 – Additive levels of detail for Gaussian splats

Order-optimisation framework for streaming progressive loading: closed-form quadratic-assignment AUC, monotone-submodular utility on the splat Gram matrix yielding a uniform (1 − 1*/e*) greedy guarantee, sparse-Gram + sparse-greedy implementation, and numerical experiments on four real Luxar gsplat datasets. Companion theory document for the additive ladders described in the main text.

### Supplementary Document 10 – Scene graph, transforms, overlays, and viewer pipeline

Reference for scene-graph node types and attributes, hierarchical 4 × 4 and per-dimension transform composition with the inverse-query algorithm, the layer model and how display-range/gamma/blending controls map to shader uniforms, the overlay system (text/image/video/HTML, dimension-aware visibility, anchoring), GPU picking encoding and compressed-sparse-row (CSR) encoded per-element labels and image-labels with auto-injected default hover overlays, the HDR post-processing pipeline (HDR pass ordering, tone mapping with fused exposure/offset/gamma, bloom/chromatic-distortion conflict resolution, adaptive device-pixel-ratio (DPR) scaling), and the per-dimension animation and recording system (loop modes, frame-data synchronisation, output formats including HDR EXR captured pre-tone-map). Chunk layout, encoding and the chunk cache are outside its scope.

### Supplementary Document 11 – Tiled fitting for large volumes

Analytical derivation of raised-cosine apodisation windows, proof of the partition-of-unity property at tile boundaries, linearity argument for splat-set union, tile-geometry conventions, and Slurm fan-in orchestration; then an empirical sweep against monolithic fits at equal splat budget on a confocal blastocyst and a light-sheet *Tribolium* embryo (tile sizes 64–256, overlaps 0–40%, with a no-taper control): seam-to-interior residual ratio 1.01– 1.13 at 5–20% overlap on both volumes, versus 2.15–7.52 with a hard cut and 3.91–5.10 with overlap but no taper (single fits, no seed replicates); at a fixed budget the tiled union stays within −0.98 to +0.19 dB of the monolithic fit in the moderate-overlap configurations, more overlap buys nothing and 40% costs 0.54–1.68 dB, and the finest grids (81–90 tiles of a few hundred splats each) pay the most; the CLI’s mass-weighted seed split and its quarter-share floor are tabulated per grid; per-tile GPU footprint (43–46 MB of allocated tensor memory per million voxels fitted at once, probed above one million voxels).

### Supplementary Document 12 – Splat parameter encoding: quantisation fidelity and codec selection, natively and in the browser

Quantisation fidelity of the shipped splat encoding (uint16 centres, certified uint8 Cholesky factors, amplitudes on a bounded-linear uint8/uint16 or geometric-log uint16 grid chosen from the array’s dynamic range) on fresh float32 fits of 17 benchmark volumes at 32K and at the cross-validation optimum (30 cells; for the two signal-limited volumes the 2048K endpoint). PSNR against the original changes by at most −0.679 dB (median −0.012; the largest loss is the *Tribolium* 2048K cell, the next largest among the reported cells −0.167 dB), the PSNR of the quantised render relative to the float32 render is 59–88 dB, and the store shrinks from 30–37 to 8.8–12.7 bytes/splat. Then a codec, filter, and level sweep (164 configurations per array) for the quantised splat-parameter arrays a Luxar store writes, benchmarked both natively and through the viewer’s in-browser WebAssembly decode path. Characterises the shipped width-aware per-dtype compressor policy (Blosc-zstd level 9; byte shuffle for multi-byte integer codes, no shuffle for single-byte codes and floating-point arrays; the float rule is not derived from this sweep), documents a level-dependent Blosc shuffle anomaly on 64 KiB chunks, and measures the four parameter arrays of the store as the writer emits it (its own atom-aligned chunks and the columnar delta prefilter its probe enables on the centres). The store costs 9.63 bytes/splat under the rule as shipped (10.09 bytes/splat for the same rule re-chunked to 64 KiB without the prefilter, 14.3% below the uniform Blosc-zstd level-3 bit-shuffle baseline) and 9.50 bytes/splat if the log-companded amplitude codes are left unshuffled (a further 1.44%; 15.6% below the baseline at 64 KiB); in the latter configuration every array decodes at 138–215 MB/s in the browser and 326–592 MB/s natively on the identical chunk set on the benchmark workstation.

### Supplementary Document 13 – Fitting wall-time on a commodity GPU

End-to-end Gaussian-splat fitting wall-time measured on a commodity NVIDIA GeForce RTX 3070 (8 GiB, 2020 consumer hardware) across 17 benchmark volumes and a splat-count sweep (*K* ∈ {4,000, …, 256,000}), under the same fitting configuration as the main rate-distortion analysis (early stopping at patience 500) except the shipped background-floor default. Supports the “minutes per volume” figure at the 32K reference budget on hardware widely available to individual laboratories (twelve of the seventeen operating points of Table 1 lie on the timed grid; the five at 8,000, 128,000 and 2,048,000 splats were not timed): at a fixed reference budget of *K* = 32,000 splats every timed volume fits in 39–379 s (median ≈ 2.4 min). Across the 85-cell sweep, 2 cells failed with a CUDA out-of-memory error, both at *K* = 256,000 on two volumes (Fly brain neurons and Neuromast nuclei) and recorded as such. Every other cell fits within 6.2 GiB, including the ∼ 100-megavoxel light-sheet volumes at *K* = 256,000; peak memory does not track voxel count (the two failures are a 28- and a 99-megavoxel volume). A final section costs the whole adoption path once on the same card and on public data (a 400-frame *C. elegans* confocal recording, Zenodo record 6460303): TIFF stacks to OME-Zarr, calibration, fitting and merging every frame, scene compilation, hosting re-chunking, export and a cold open from a laptop, with wall time, host memory and GPU memory per stage.

### Supplementary Document 14 – Gaussian-splat timelapses

The three shipped time-lapse archives, Zebrafish embryo nuclei (51 of 253 timepoints, 2.4M splats per frame), *Drosophila* embryo nuclei (500 timepoints, 256K per frame) and the two-channel Neuromast recording (100 timepoints, 128K per frame over both channels), as fitted per timepoint with time as a hard ordering barrier, so each timepoint’s splats are stored contiguously, no level-of-detail rung blends two timepoints and one time-point is resident at a time. Reports storage (raw frames to archive: 157×, 67× and 157×), the streaming cost of opening a frame and of a time step in the compiler’s and the published chunk layouts, and the fidelity of the shipped archives on 20 sampled frames, each scored against its raw frame (not held-out) (Fig. 4).

### Supplementary Videos

Every video is a screen capture of the shipped viewer or command-line tools, shown as rendered. Each opens with a title card; section cards or lower-third labels mark cuts between scenes, and data credits appear on screen either on those cards or in the scene’s own credit line; network conditions and time compression are stated on a card where they are part of the story. Only Video 11 carries sound. Each title below links to the video’s page on the demo site; all seventeen are deposited at Zenodo (https://zenodo.org/records/22825093; concept DOI 10.5281/zenodo.22825092; CC BY-SA 4.0) as 1080p HEVC masters; after each title, the file name links to that file and *watch* to the same cut streamed in the browser from https://demos.luxarviewer.dev/v/, which lists all seventeen with their provenance.

#### Supplementary Video 1 - A volume becomes a scene

SuppVideo01_volume_to_scene.mp4 A *Drosophila* embryo during gastrulation, one frame of the SiMView light-sheet recording of Fig. 4b: the raw stack is rendered in three dimensions in napari,^4^ turned and zoomed. Luxar gsplat fit then fits the whole 108 × 1352 × 532-voxel stack from 256K seeds, the operating point of Table 1, with the standard preset (5,000 iterations, 484 s on one consumer GPU, a GeForce RTX 3070), to 200,278 splats after the post-fit cull; the reconstruction is shown converging beneath the volume every 25 iterations, time-compressed, with a five-fold magnified inset of the same region of each and the PSNR curve drawn on the console. The fitted splats are scaled to physical microns, given a streaming ladder and built into a scene by about fifteen lines of Python, typed and run in a terminal. The scene opens in the browser, where the embryo turns about its long axis, is orbited and is zoomed into the nuclei of the ventral furrow.

#### Supplementary Video 2 - A 4D embryo and its lineage

SuppVideo02_celegans_lineage.mp4 The *C. elegans* embryo of Fig. 1b, nuclei as Gaussian splats fitted per time-point, lineage tracks as fading polylines and current positions as points, in one scene: time plays while the camera orbits, and as the slice moves along time the tracks are clipped and the splats fade in and out according to their extent along the hidden dimension.

#### Supplementary Video 3 - *Drosophila* gastrulation and the neuromast

SuppVideo03_drosophila_neuromast. mp4 The 500-frame *Drosophila* recording of Fig. 4b played through gastrulation while the camera turns about the embryo, and the two-channel neuromast recording of Fig. 4c played with the membrane and nucleus channels shown together and each alone through the Layers panel, then viewed close up from above the skin.

#### Supplementary Video 4 - Loading only what the view needs

SuppVideo04_lazy_loading.mp4 The 500-timepoint *Drosophila* time-lapse of Fig. 4b and the *Tribolium* embryo of the adaptive level-of-detail demo, each opened cold from the public demo site with the browser caches emptied and the data-loading monitor open, the *Drosophila* half with the browser’s network throttled to 30 Mbit/s and 40 ms latency, the *Tribolium* half over the unthrottled home connection. The fly embryo’s first time-point arrives through its additive ladder, coarsest rung first, sharpening into the full frame while the monitor counts the splats resident and the bytes loaded; each step along time then fetches the chunks that hold the new timepoint, the loaded total rising as new chunks are needed. The *Tribolium* fit is stored as seven substitutive levels coloured green (finest) to red (coarsest), and a coarse level paints first (18.5K splats, against 296.6K at the finest). Zooming in until the embryo fills the screen fetches the finer levels, the monitor’s visible-splat count rising to the full 296.6K; zooming back out returns to a coarser level and zooming in again to the finest, both from what is already resident.

#### Supplementary Video 5 - Fitting at scale: a crossvalidated splat count and a batch fit across timepoints

SuppVideo05_fitting_at_scale.mp4 Both runs on the GPU workstation (an RTX PRO 6000), consoles time-compressed. luxar gsplat cal sweeps nine splat counts from 1,000 to 2,048,000 on one frame of the *Drosophila* recording (108 × 1352 × 532 voxels), fitting each and scoring it on held-out blind-spot voxels; the sweep took 110 minutes. Held-out PSNR climbs by 13 dB to 512,000 splats and then flattens, gaining 0.1 dB over the next two doublings while the train–held-out gap keeps opening, so the report classifies the curve as a plateau and returns its onset, *K*^*⋆*^ = 512,000 (on the benchmark’s doubling grid the detector places the same frame’s onset one rung lower, at 256K, the Table 1 operating point); the first two pages of its PDF report are shown. Luxar gsplat batch-fit run then fits ten timepoints of the same recording, every fiftieth of the 500 (25 minutes apart), at 512,000 seeds each, two fits at a time on the same GPU, in 37 minutes of wall time, and merges them into one four-dimensional scene with a four-rung streaming ladder. Scaled to the recording’s voxel size (1.93 micrometres axially, 0.40625 laterally), the scene opens in the viewer, which steps through the ten timepoints with the data-loading monitor open and dollies into the gastrulating embryo.

#### Supplementary Video 6 - Near-unlimited scaling with adaptive level of detail

SuppVideo06_adaptive_lod.mp4 A hundred copies of an embryo laid along a line, each an adaptive level-of-detail group of seven levels, from 296,559 splats at the finest to 73 at the coarsest, and each coloured by the level it is currently drawn from, green for the finest through yellow and orange to red for the coarsest, with an on-screen legend giving every level’s splat count. The camera flies along the line and looks down it to the far end, where the embryos are dots: an embryo swaps to finer levels as it approaches and fills the screen and drops back to coarser ones as it recedes, so the detail drawn never exceeds what the view can resolve; the data-loading monitor is open throughout.

#### Supplementary Video 7 - The grammar of levels of detail: six topologies from one fit

SuppVideo07_lod_recipes. mp4 The *Tribolium* embryo fit of the recipe gallery laid out six times along a line, one per lod –recipe topology (flat, stream, levels, tiles, overview, adaptive), each column coloured by part and shaded by the level it is drawn from, opened from the public demo site with the scene’s own text overlays hidden. After a view of the whole line, the camera frames each level-of-detail column alone and dollies from far to near and back, pausing at each level. The levels column swaps from its coarse level (red) through its middle level to its finest (teal) as it grows to fill the screen, the tiles column loses parts to frustum culling as the camera closes in, the overview column bursts from one coarse cap into eight fine tiles, and in the adaptive column each tile settles on its own level, visible as differently shaded bands. The data-loading monitor, open during the column shots, counts the splats resident for everything in view.

#### Supplementary Video 8 - A large Gaussian-splat volume: one frame of the zebrafish recording

SuppVideo08_ zebrafish_embryo.mp4 The first frame of the 51-frame *h2afva* zebrafish recording of Fig. 4a, 2,217,045 Gaussian splats, held at that timepoint while the camera orbits the embryo in the viewer: a single large, complex microscopy volume rendered as splats.

#### Supplementary Video 9 - *Drosophila* gastrulation, the whole recording

SuppVideo09_drosophila_timelapse. mp4 The 500-frame *Drosophila* recording of Fig. 4b streamed from the public demo site, whose data is served from a content-delivery edge in San Jose, 7–12 network hops and 3–30 ms round trip from the recording machine, over a home Wi-Fi 6E link that measured 779 Mbit/s down and 9 ms to a Bay Area speed-test server. The camera turns, orbits and closes in on the ventral furrow while the first 80 minutes of development play slowly; the recording then plays from its first to its last frame (30 s intervals, 4 hours of development) in 33 s, from a dorsal view and, after 81 minutes, an oblique one.

#### Supplementary Video 10 - Annotated atlases: legends, hover labels and hover images

SuppVideo10_ annotated_atlases.mp4 The Tabula Sapiens atlas (about 500K cells from 24 tissues) orbited with its legend of tissues; the arXiv/bioRxiv/medRxiv embedding of scientific papers, where hovering a point names the paper through GPU picking; and the CytoSelf protein-localisation map, where hovering a protein shows the micrograph it was embedded from.

#### Supplementary Video 11 - A guided tour with overlays and narration

SuppVideo11_esm_stories_tour.mp4 The Swiss-Prot ESM C protein landscape of Supplementary Video 14 with a hidden *story* dimension whose steps are bound to camera poses: from the overview, five of its ten stories (hemoglobin, photosystem II, the prion protein, ATP synthase, insulin) are stepped through; at each step the camera flies to the family’s cluster, lights it up, and the dimension-aware overlays appear (an HTML panel of facts, a turntable video of a representative structure) while the scene’s own narration plays over its ambient bed, ducked under the voice as the scene’s audio configuration specifies. The picture is a screen capture of the viewer; the sound is the scene’s narration and bed mixed to the recorded timeline.

#### Supplementary Video 12 - The viewer interface

SuppVideo12_viewer_interface.mp4 A walk through the viewer’s controls on three scenes. On the *C. elegans* lineage scene of Fig. 1b: the help overlay listing every keyboard shortcut, the Dimension Navigation panel with the time slider dragged and played, the control rail and the panels it opens (Navigation with its camera modes and turntable, Rendering, Layers with a layer hidden and re-shown and its blending changed, the data monitor and the recording panel), and the three camera modes, orbit, fly and orthographic, with the turntable spinning the embryo. On the arXiv paper map, beside a browser window: hovering a point names the paper, the right-click menu copies its title or opens its link, and that link, a scholar search for the title, opens in the browser. On an iPad in Safari, the Cells3D volume driven by touch: one finger rotates, two fingers pan, a pinch zooms, a twist rolls the view, a double tap recentres, and a tap on the rail opens it.

#### Supplementary Video 13 - Sharing a scene: export, embed, record

SuppVideo13_sharing_surfaces.mp4 luxar export writes the *Drosophila* gastrulation scene to a self-contained folder and, with –native macos, to a double-clickable application; the application is launched from the Finder and opens the scene in its own window, where the embryo is framed and turned; the folder is opened from its own local server in a browser; the viewer, installed from npm as @luxar/viewer, is mounted into a canvas of an ordinary HTML page beside text; and the recording panel records a movie of the turning embryo in the browser, from the Record button to the saved file, which is then played back.

#### Supplementary Video 14 - Flying through the data

SuppVideo14_flythroughs.mp4 Free flights of the camera, in the viewer’s fly mode, inside four scenes: the Fly-Light MCFO brain of Fig. 1c; the ESM C protein landscape of 575,503 Swiss-Prot proteins coloured by taxon, entered from afar and threaded between its clusters; the human nuclear pore complex, approached head-on and flown through its central channel along the inner ring; and the Human Connectome Project (HCP) white-matter tractography atlas, flown along its tracts. Each flight turns the viewing direction as it moves.

#### Supplementary Video 15 - From molecules to atlases

SuppVideo15_tour_life_science.mp4 A tour of eight life-science scenes from the public gallery, one orbit each: a clinical CT coloured by 117 TotalSegmentator organ labels, the 87 tracts of the HCP tractography atlas, the Fly-Light MCFO neuron scene of a fly brain, the Zebrahub RNA-velocity UMAP with velocity streamlines, the FlyLight MCFO brain of Fig. 1c in its bounding box, the atoms of ATP synthase (PDB 5DN6) as points, the 4.9 million atoms of the human nuclear pore complex (an integrative model of all 808 chains) orbited and then entered through the pore, and the cryo-EM density of the PBCV-1 giant-virus capsid as Gaussian splats.

#### Supplementary Video 16 - Beyond biology

SuppVideo16_tour_beyond_biology.mp4 A tour of the public gallery, one orbit each: the Laniakea super-cluster with Cosmicflows-4 velocity streamlines, the DESI galaxy catalogue of about 9.75 million objects, the ocean currents of Earth, a macro-photogrammetry Gaussian-splat capture of a cluster fly (Dany Bittel, CC BY 4.0) imported from an INRIA-dialect PLY file, an evolving cumulus cloud played in time, a Mandelbulb fractal, and an animated particle collision.

#### Supplementary Video 17 - Four geometry types, one pipeline

SuppVideo17_geometry_types.mp4 The same rendering conventions on each primitive with real data: the atoms of ATP synthase (PDB 5DN6) as points, their sharpness parameter swept from soft Gaussian sprites to hard spheres and back along a hidden dimension of the scene; the chromosomes of a single cell’s genome (Dip-C) as polylines drawn as capsules, seen close up at their joints; the shaded isosurface meshes of the Cells3D volume lit by the view-anchored headlight; and the Gaussian splats of the *Drosophila* gastrulation frame switched between additive and volumetric blending while the embryo turns about its long axis.

**Supplementary Figure 1.**
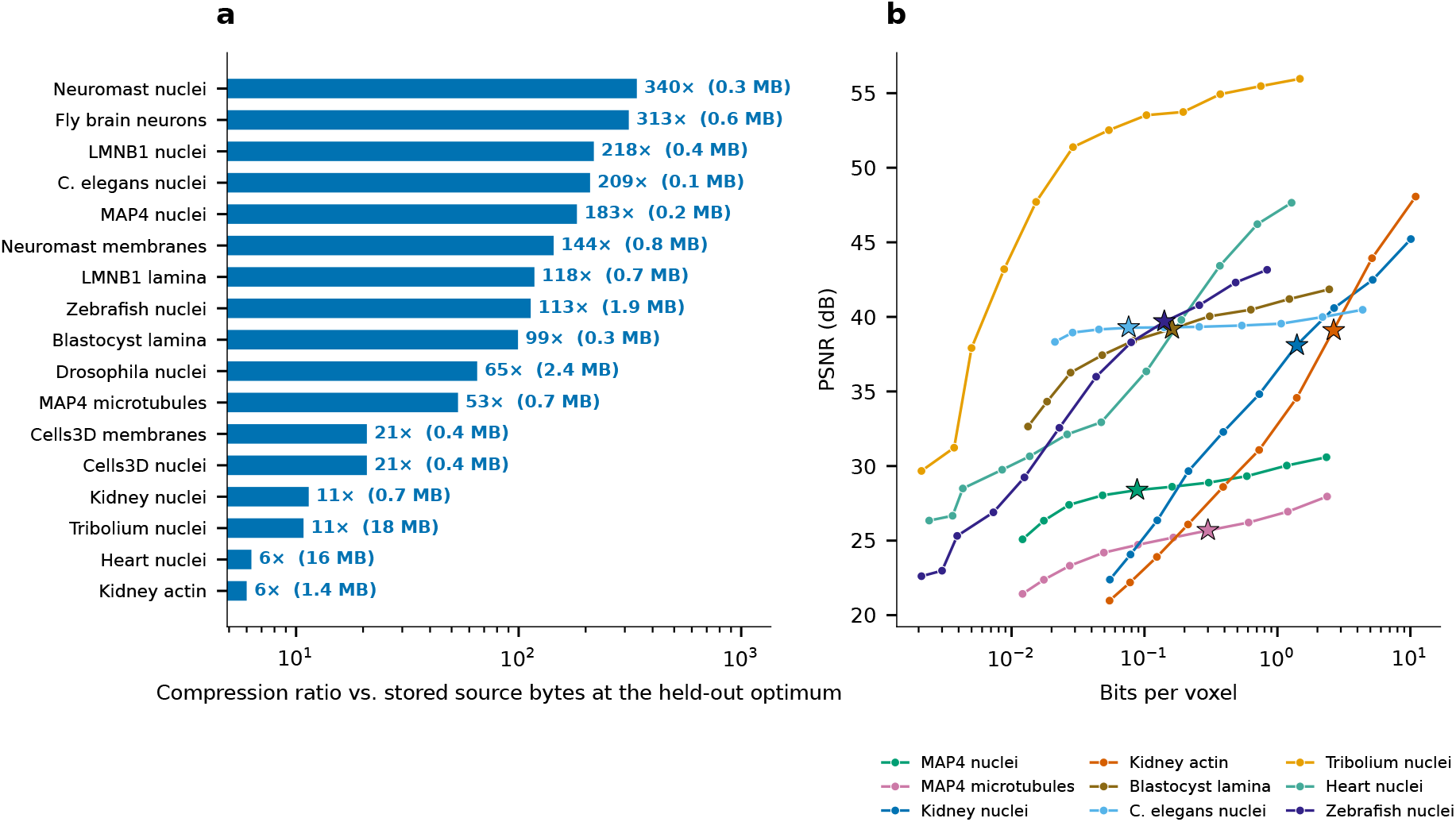
Compression at the per-dataset cross-validation optimum. **a**, Compression ratio of the fitted Gaussian-splat representation against the source volume at its stored bit depth at each volume’s held-out (cross-validation) optimum, for the 17 benchmark volumes with a completed sweep, ranked from the most to the least compressible; labels repeat the ratio and give the splat file size. **b**, Rate-distortion (PSNR vs. bits per voxel): the full curve of each of the nine volumes of Fig. 3, the operating point starred on the seven with a finite held-out optimum. Compression ratios range from 6.0× (Kidney actin, optimum at 128K splats) to 340*×* (Neuromast nuclei, optimum at 64K splats), with a median of 99.3*×* across the 17 volumes. 15 of 17 reach a finite held-out optimum; Mouse heart nuclei (6.3*×*) and *Tribolium* embryo nuclei (10.8*×*), both at 2M splats, are signal-limited: their held-out curves are still rising at the largest tested capacity, so the detector’s operating point is the endpoint of the sweep; the *Drosophila* embryo (65.1 *×* at 256K) plateaus, its held-out PSNR flat within 0.05 dB from 512K to 2M splats. The Mouse heart re-run with a tripled iteration cap, which confirms that it is signal-limited, is in Supplementary Document 2. See Supplementary Document 7.

**Supplementary Figure 2.**
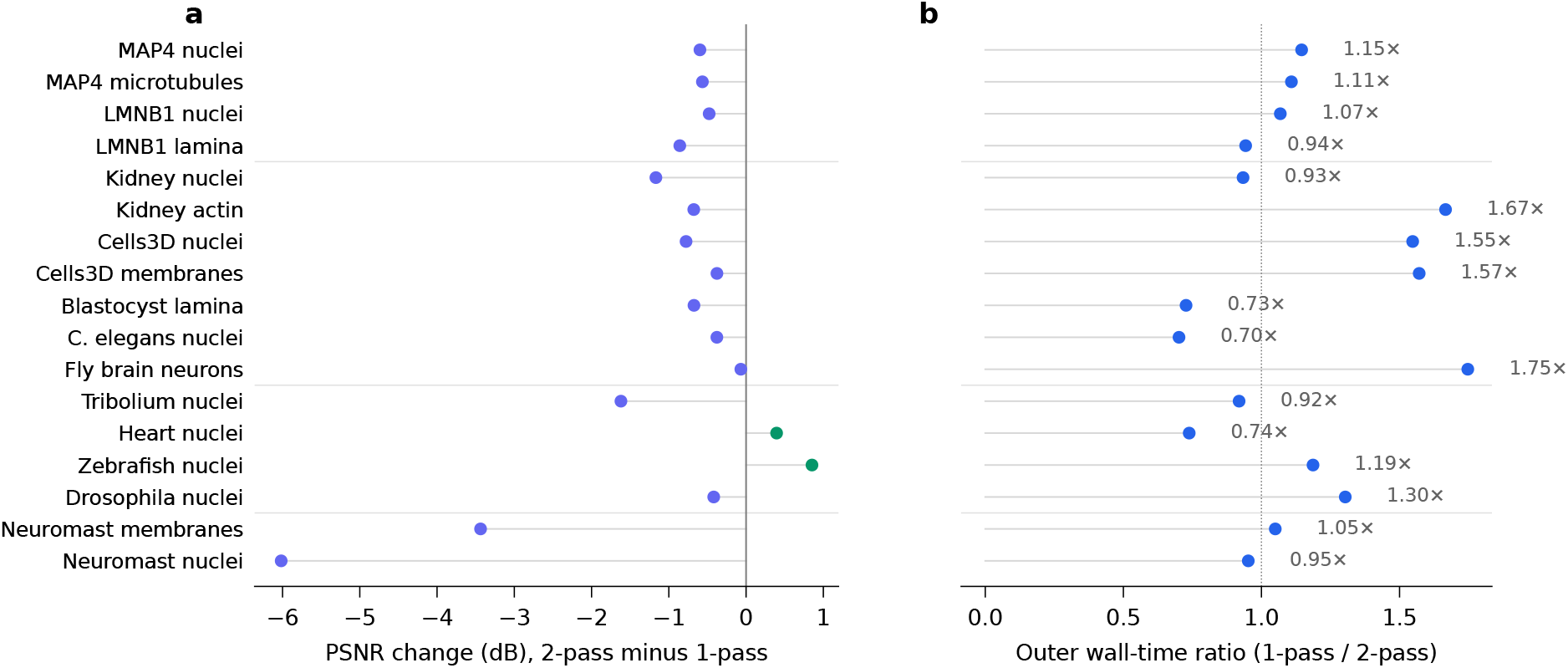
Progressive vs. single-pass fitting at a nominal 32K-seed budget. **a**, PSNR difference (2-pass minus 1-pass) across 17 benchmark volumes grouped by modality. Green markers: 2-pass progressive fitting achieves higher training PSNR; purple markers: single-pass is higher. Each cell is a single fit at a common nominal seed budget (retained post-cull counts 22,178–32,384). On the six volumes whose margin is below 0.5 dB, two further RNG seeds per condition put the run-to-run PSNR spread (range over three seeds; initialisation is deterministic in this fitter, so the spread measures GPU run-to-run nondeterminism and, in the 2-pass arm, the residual-seeder draw) at 0.002–0.093 dB (median 0.013 dB), and the 1-pass and 2-pass ranges are disjoint on six of the six under that spread (Supplementary Document 3). 2-pass is ahead only on Mouse heart nuclei (+0.40 dB) and Zebrafish embryo nuclei (+0.85 dB); the remaining fifteen volumes favour single-pass by 0.07–6.02 dB. **b**, Wall-time ratio of 1-pass over 2-pass (whole fitter call, CUDA-synchronised, sequential runs on an otherwise idle GPU; values above 1 mean 2-pass finished first). The ratio spans 0.70–1.75*×* (median 1.07*×*); 2-pass finishes first on 10 of 17 volumes. No mechanism is inferred from these single observations.

**Supplementary Figure 3.**
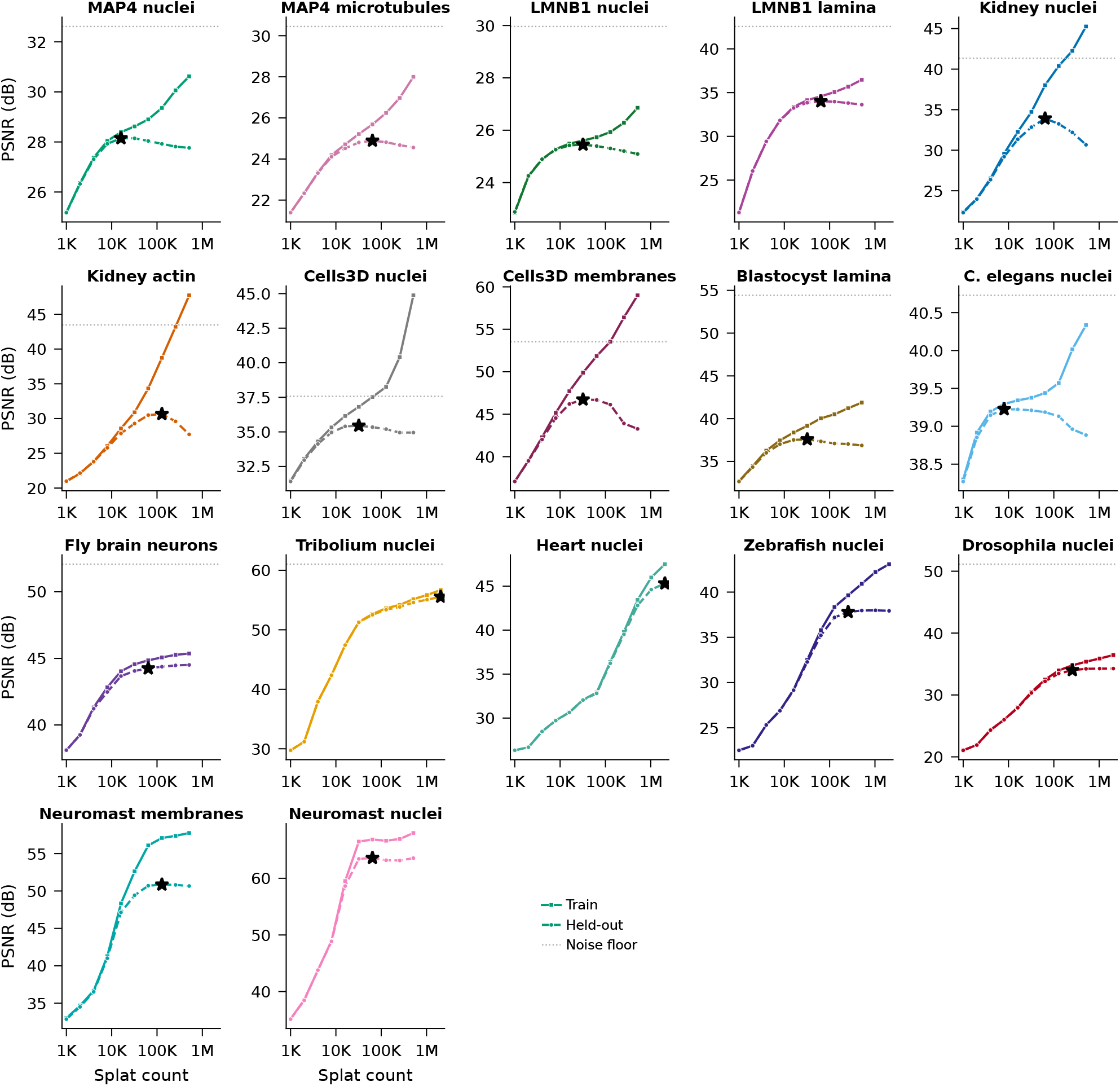
Blind-spot cross-validation for every benchmark volume. Train (solid) and held-out (dashed) PSNR vs. splat count for the seventeen volumes with a completed sweep, in the order of Table 3. Stars: the selected *N* ^*∗*^, the held-out peak where one exists, otherwise the plateau onset (Fly brain, Zebrafish, *Drosophila*) or the sweep endpoint for the signal-limited curves (*Tribolium*, Mouse heart). Dotted lines: noise floor. The x-axis is the requested splat count; the fitted count after culling is lower on sparse volumes (neuromast membranes 77,900 and nuclei 40,191 fitted at 512K requested; Supplementary Document 2). A held-out peak followed by a decline is seen on 10 volumes; on the two iSIM neuromast channels the maximum is interior but the post-fit cull holds the retained count nearly constant beyond it, so they are read as cull-limited; the seven volumes without a clear peak are Fly brain neurons, Tribolium embryo nuclei, Mouse heart nuclei, Zebrafish embryo nuclei, Drosophila embryo nuclei, Neuromast membranes, Neuromast nuclei. The per-volume regimes (deconvolution-induced noise correlations, estimator ceilings, plateaus) are discussed in Supplementary Document 2 and in Results.

**Supplementary Figure 4.**
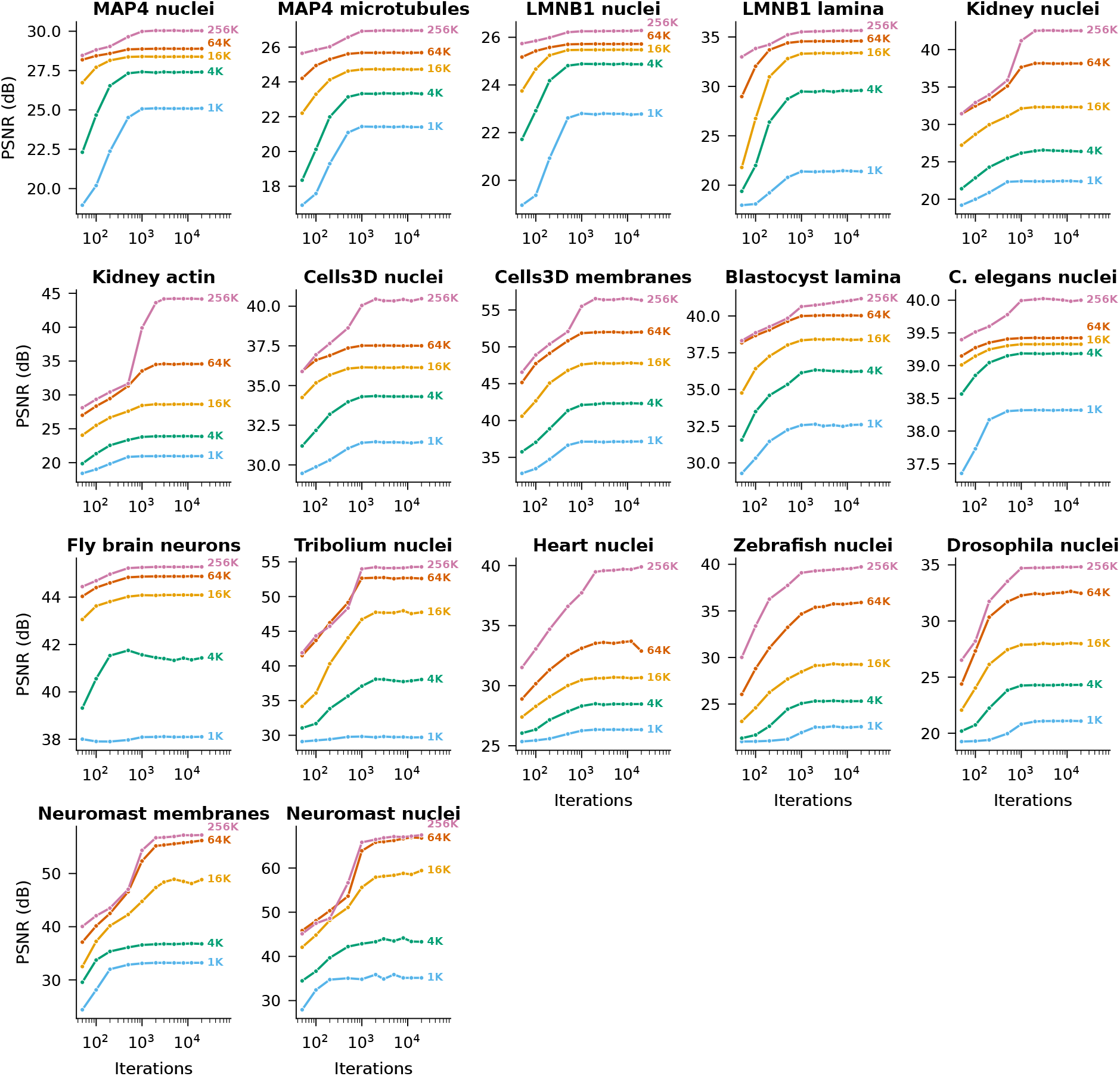
Convergence across volumes and splat counts. Training-mode PSNR vs. iteration for every volume with a convergence sweep (17 volumes, one panel each, in the order of Table 3); each curve is labelled by its splat count at its right end. Most quality is gained within 2,000 iterations; beyond this point, additional iterations yield diminishing returns in training PSNR regardless of splat count. Model capacity (splat count) has a substantially larger effect on reconstruction quality than optimisation duration, and 77 of the 85 cells are within 0.5 dB of their 20,000-iteration training PSNR by 2,000 iterations (81 within 1 dB; the largest shortfall is 1.50 dB, on Neuromast nuclei at 16 K). The held-out counterpart of this sweep, at each volume’s operating point, is in Supplementary Document 4, §3.5.

**Supplementary Table 1.** Viewer rendering performance on fifteen scenes. Measured with Playwright in headless Chromium on an NVIDIA RTX 3070 at 1280 *×* 720 with adaptive DPR pinned to 1.0 (Methods), 5 repeats per scene, on the demo datasets regenerated from the released code, in one session on an otherwise idle machine (1-min load average *≤* 2.9). Rows are bundled demo scenes, except where the label names a benchmark fit (a Gaussian-splat fit of a Table 3 volume converted to a scene; the heart 2M row is the 60,000-iteration re-run of Supplementary Fig. 1) or a time-lapse (one timepoint drawn). *Load*: a cold load into an empty browser cache, from first request to scene ready. *Elements*: primitives drawn at the scene’s opening view once it has settled, so streamed ladders and time-lapse frames are counted fully resident; a non-displayed dimension means only the current slice is drawn (storm opens on its 146 K-splat widefield fit, not on the 5.0 M single-molecule localisations that its second view value holds, each time-lapse row draws one timepoint, and forest one growth *×* season slice of a 1.5 M-element archive). *rAF FPS*: frames per second of wall-clock time on the viewer’s own requestAnimationFrame loop during the camera orbit, capped at the headless runtime’s begin-frame cadence of about 60 Hz (headless Chromium has no display). *Render ms*: mean wall time of one synchronised render-and-readback call (until a 1-pixel gl.readPixels returns), with the default post-processing pass enabled (Supplementary Document 6). Every measured scene holds 60 FPS, the heaviest (Zebrafish nuclei (time-lapse), 2.2M splats) costing 5.6 ms of the 16.7 ms budget. Because a frame rate pinned at its cap is not by itself evidence of headroom, the final row is a *negative control*: a deliberately overloaded synthetic scene of 10^7^ large splats, measured in a single static-camera run, which leaves the cap as it must; six further runs gave 18–44 FPS, and its splats are far larger on screen than the 4-pixel sprites of the Supplementary Document 6 sweep, whose 10^7^ cell costs 19.4 ms per synchronised call. Full count, size and viewport sweeps are in Supplementary Document 6.

| Dataset | Geometry | Elements | Load (s) | rAF FPS | Render (ms) | Heap (MB) |
| --- | --- | --- | --- | --- | --- | --- |
| Cells3D | GSplats | 99 921 | 0.5 | 60 | 1.55 | 39 |
| Neuromast membranes + nuclei (time-lapse) | GSplats | 110 614 | 0.6 | 60 | 1.37 | 50 |
| OpenCell MAP4 | GSplats | 119 087 | 0.5 | 60 | 1.22 | 46 |
| STORM microtubules | GSplats | 146 284 | 0.5 | 60 | 1.48 | 61 |
| Kidney (6D, 3 layers) | GSplats | 157 120 | 0.5 | 60 | 1.25 | 56 |
| Drosophila nuclei (time-lapse) | GSplats | 167 342 | 0.4 | 60 | 1.47 | 74 |
| L-system forest | mixed (all four) | 191 096 | 1.6 | 60 | 1.54 | 111 |
| Tribolium embryo | GSplats | 296 559 | 0.7 | 60 | 1.75 | 85 |
| Lorenz attractor | Points | 500 000 | 0.7 | 60 | 2.40 | 78 |
| Spiral galaxy | Points | 500 000 | 0.7 | 60 | 2.30 | 78 |
| Cells3D isosurfaces | Mesh | 1 067 632 | 0.9 | 60 | 1.06 | 88 |
| Fly brain neurons | GSplats | 1 139 140 | 5.9 | 60 | 4.63 | 336 |
| Heart nuclei, 2M fit | GSplats | 1 502 861 | 1.9 | 60 | 3.60 | 320 |
| Tribolium nuclei, 2M fit | GSplats | 2 039 735 | 2.4 | 60 | 4.55 | 417 |
| Zebrafish nuclei (time-lapse) | GSplats | 2 217 045 | 0.4 | 60 | 5.56 | 640 |
| *Negative control* | *GSplats (synthetic)* | *10 000 000* | *–* | *23.2* | *51.5* | *–* |

**Supplementary Table 2.** Gaussian-splat time-lapses: what a viewer streams and what it gets back. For the recordings of Table 2, in the chunk layout the demo site publishes: the decoded parameter payload of one timepoint (median over the six captured frames; float32 centres, amplitude and packed Cholesky factors, 40 B per splat; the renderer’s own GPU allocation is larger), the bytes and HTTP requests that open the scene at its first frame, the bytes and requests one time step costs, and the PSNR range of the shipped archive against the raw frame (its own range as *R*; not held-out, not the normalised basis of Table 1) over the five frames per recording of Fig. 4 (20 scores, both neuromast channels counted). Time is a hard ordering barrier in every archive, so a step fetches the chunks holding the new timepoint (for the *Drosophila* and neuromast nodes a 1 MB chunk already holds neighbouring frames, so most advances make only a zero-byte conditional re-read of the metadata document); the step values are the first two advances from the opening frame; the compiler-layout numbers are in Supplementary Document 14.

| Recording | Payload (MB / frame) | Opening (MB) | Opening (requests) | Step (MB) | Step (requests) | PSNR (dB) |
| --- | --- | --- | --- | --- | --- | --- |
| Zebrafish embryo nuclei | 96.4 | 40 | 480 | 17.9–18.0 | 65 | 43.3–59.9 |
| *Drosophila* embryo nuclei | 10.2 | 19 | 45 | 0.00 | 1 | 39.5–48.7 |
| Neuromast membranes + nuclei | 5.1 | 65 | 140 | 0.00 | 1 | 45.3–57.8 |

**Supplementary Table 3.** Effective resolution and splat scale of the 17 benchmark volumes. Resolution is estimated from the data by decorrelation analysis^59^ (median over slices; axial from a sector of the *xz*/*yz* spectra in physical frequency units), so it is the effective, noise-limited resolution of each volume as fitted, with a factor 1.1–1.3 estimator convention relative to a Gaussian FWHM and up to a factor 1.4–3 sensitivity on dim volumes (Supplementary Document 2). *ℓ*_10_ is the 10th-percentile axis-aligned splat standard deviation at the operating point *N* ^*∗*^, in voxels: 0.50–1.38 on every volume, against lateral resolutions of 2.2–24.4 pixels, so the smallest splats are at or below the data’s resolution on every volume: resolution over splat FWHM is 1.15–8.7 (median 2.4 *×*), above 2 on 13 of the 17 volumes. Kidney has 16 planes, too few for an axial estimate; ^‡^ marks curves with no held-out peak, whose *N* ^*∗*^ is the plateau onset or, for signal-limited curves, the sweep endpoint.

| Volume | Voxel ($z, y, x$), $\mu\text{m}$ | Lateral res. ($\mu\text{m}$) | Axial res. ($\mu\text{m}$) | $\ell_{10}$ lat. (vox) | $\ell_{10}$ ax. (vox) | $N^*$ |
| --- | --- | --- | --- | --- | --- | --- |
| OpenCell-MAP4 nuclei | 0.500, 0.200, 0.200 | 1.56 | 3.52 | 1.38 | 1.48 | 16K |
| OpenCell-MAP4 microtubules | 0.500, 0.200, 0.200 | 2.56 | 2.07 | 0.92 | 0.79 | 64K |
| OpenCell-LMNB1 nuclei | 0.200, 0.200, 0.200 | 2.21 | 4.88 | 1.19 | 1.35 | 32K |
| OpenCell-LMNB1 nuclear lamina | 0.200, 0.200, 0.200 | 1.09 | 1.31 | 0.89 | 0.98 | 64K |
| Kidney nuclei | 1.250, 1.240, 1.240 | 2.94 | – | 0.62 | 1.08 | 64K |
| Kidney actin | 1.250, 1.240, 1.240 | 3.35 | – | 0.55 | 1.12 | 128K |
| Cells3D nuclei | 0.290, 0.260, 0.260 | 1.09 | 1.72 | 0.71 | 0.90 | 32K |
| Cells3D membranes | 0.290, 0.260, 0.260 | 0.60 | 0.75 | 0.86 | 0.95 | 32K |
| Blastocyst nuclear lamina | – | 3.2 px | 9.2 z-vox | 0.66 | 1.00 | 32K |
| *C. elegans* embryo nuclei | 0.750, 0.150, 0.150 | 3.66 | 4.74 | 1.19 | 1.00 | 8K |
| Fly brain neurons | 0.380, 0.190, 0.190 | 0.91 | 1.77 | 0.91 | 0.95 | 64K<sup>‡</sup> |
| *Tribolium* embryo nuclei | 0.381, 0.381, 0.381 | 1.15 | 1.59 | 0.88 | 0.92 | 2M<sup>‡</sup> |
| Mouse heart nuclei | – | 2.2 px | 7.3 z-vox | 0.77 | 1.01 | 2M<sup>‡</sup> |
| Zebrafish embryo nuclei | 1.625, 0.406, 0.406 | 2.90 | 4.21 | 0.50 | 0.50 | 256K<sup>‡</sup> |
| *Drosophila* embryo nuclei | 1.930, 0.406, 0.406 | 3.72 | 4.80 | 0.50 | 0.49 | 256K<sup>‡</sup> |
| Neuromast membranes | 0.250, 0.108, 0.108 | 0.35 | 0.74 | 0.50 | 0.50 | 128K |
| Neuromast nuclei | 0.250, 0.108, 0.108 | 0.27 | 1.11 | 0.50 | 0.50 | 64K |

**Supplementary Figure 5.**
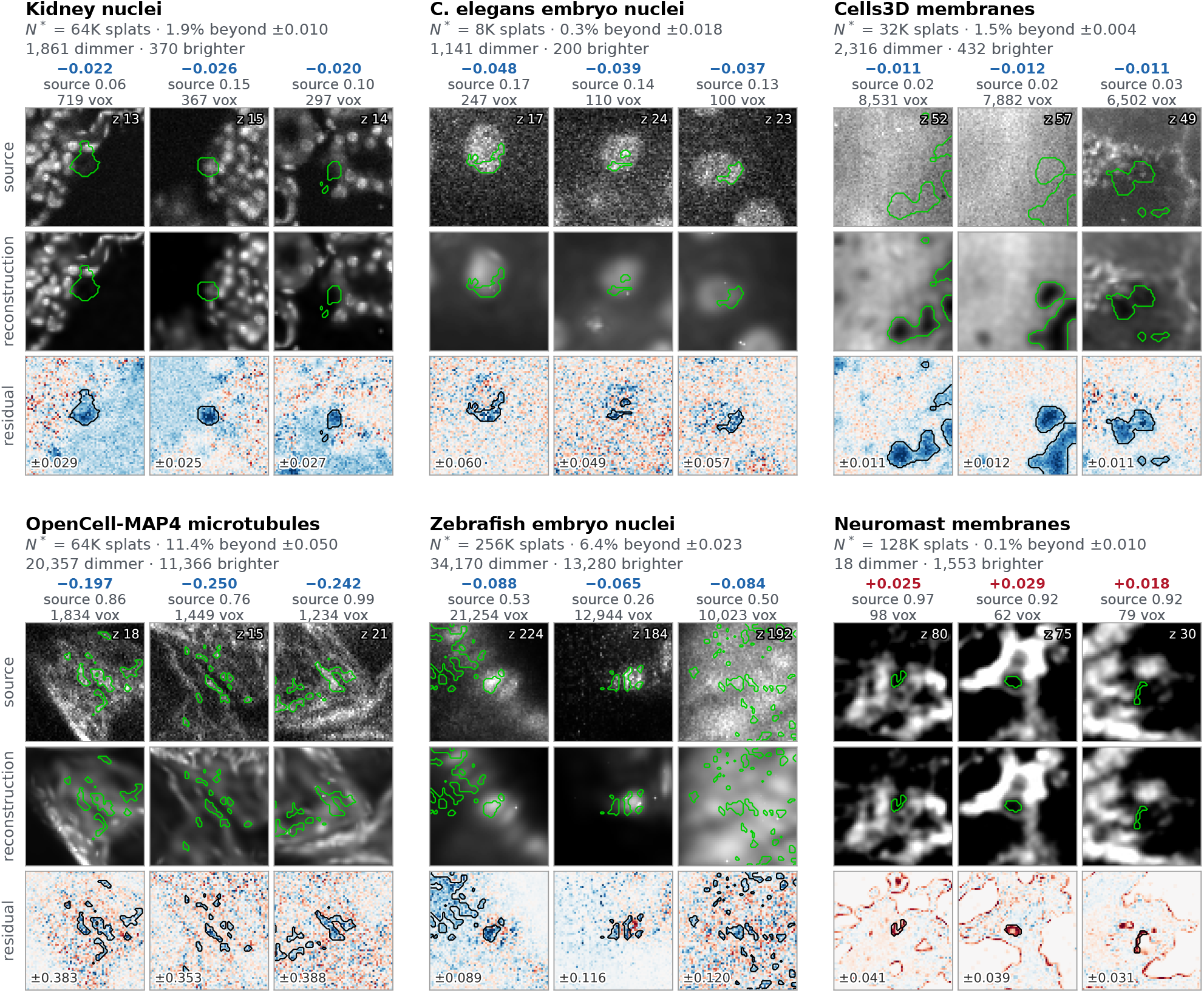
Where a fit at its held-out optimum deviates from its source. For six benchmark volumes spanning the regimes (confocal nuclei and membranes, spinning-disk microtubules, a deconvolved light-sheet embryo and a background-subtracted iSIM neuromast), the cached fit at the operating point *N* ^*∗*^ is rendered as the viewer decodes it and the normalised source is subtracted from it (residual = reconstruction *−* source). The signed residual is smoothed by one lateral voxel and thresholded at six robust standard deviations of its own noise, separately for dimmer (reconstruction below the source) and brighter voxels, and the connected components are ranked by the sum of |residual| over their voxels. Three components per volume are shown as source, reconstruction and signed residual: the strongest that pass a minimum-size filter and do not overlap an already chosen crop (a component of the opposite sign would take the third slot if one ranked among the twenty strongest; on these six volumes none did, so all three share the dominant sign), each shown through the slice where the smoothed residual peaks, with the component outlined and source and reconstruction on one display window per crop. Column titles give the signed peak residual (blue dimmer, red brighter), the local source maximum and the component size; the slice index is printed in the source crop and the residual display range in the residual crop. Block titles give *N* ^*∗*^, the share of Otsu-foreground voxels beyond the threshold (0.1–11.4%) and the component counts by sign. On 5 of the six volumes the strongest feature is an omission: a dim object among bright ones whose smoothed residual peak reaches 16–66% of the nearby source maximum; on the neuromast, whose background is exactly zero, it is a dark lumen between membranes partly filled. Statistics are in Supplementary Document 2.

**Supplementary Figure 6.**
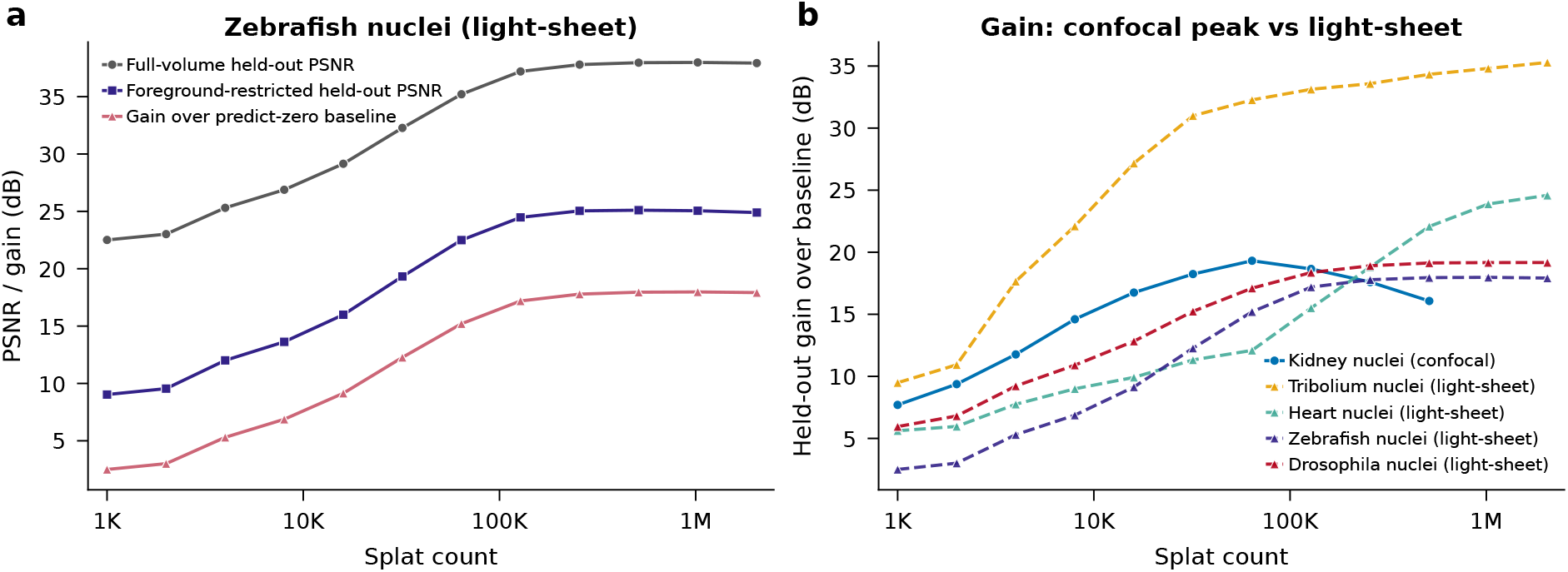
Held-out metrics on a sparse, deconvolved light-sheet volume. **a**, Three held-out measures for the zebrafish *h2afva* nuclei volume as a function of splat count: full-volume held-out PSNR, held-out gain over the predict-zero baseline, and held-out PSNR restricted to foreground voxels. On a volume that is mostly empty, the background inflates the full-volume held-out PSNR by about 13 dB relative to the foreground-only value, so the full-volume number overstates how well the structures themselves are recovered. **b**, Held-out gain curves for the confocal kidney DAPI volume, which peaks at 64K splats (the same optimum as its full-volume curve in Supplementary Fig. 3), and for the light-sheet volumes, whose gain is monotone or plateaus: on these processed, low-noise volumes there is no interior optimum, and the criterion selects the onset of the plateau for the Zebrafish and *Drosophila* volumes and the largest count swept for the signal-limited *Tribolium* and Mouse heart volumes (Supplementary Document 2). Curves and values are computed from the held-out sweep results (Supplementary Document 2).

**Supplementary Figure 7.**
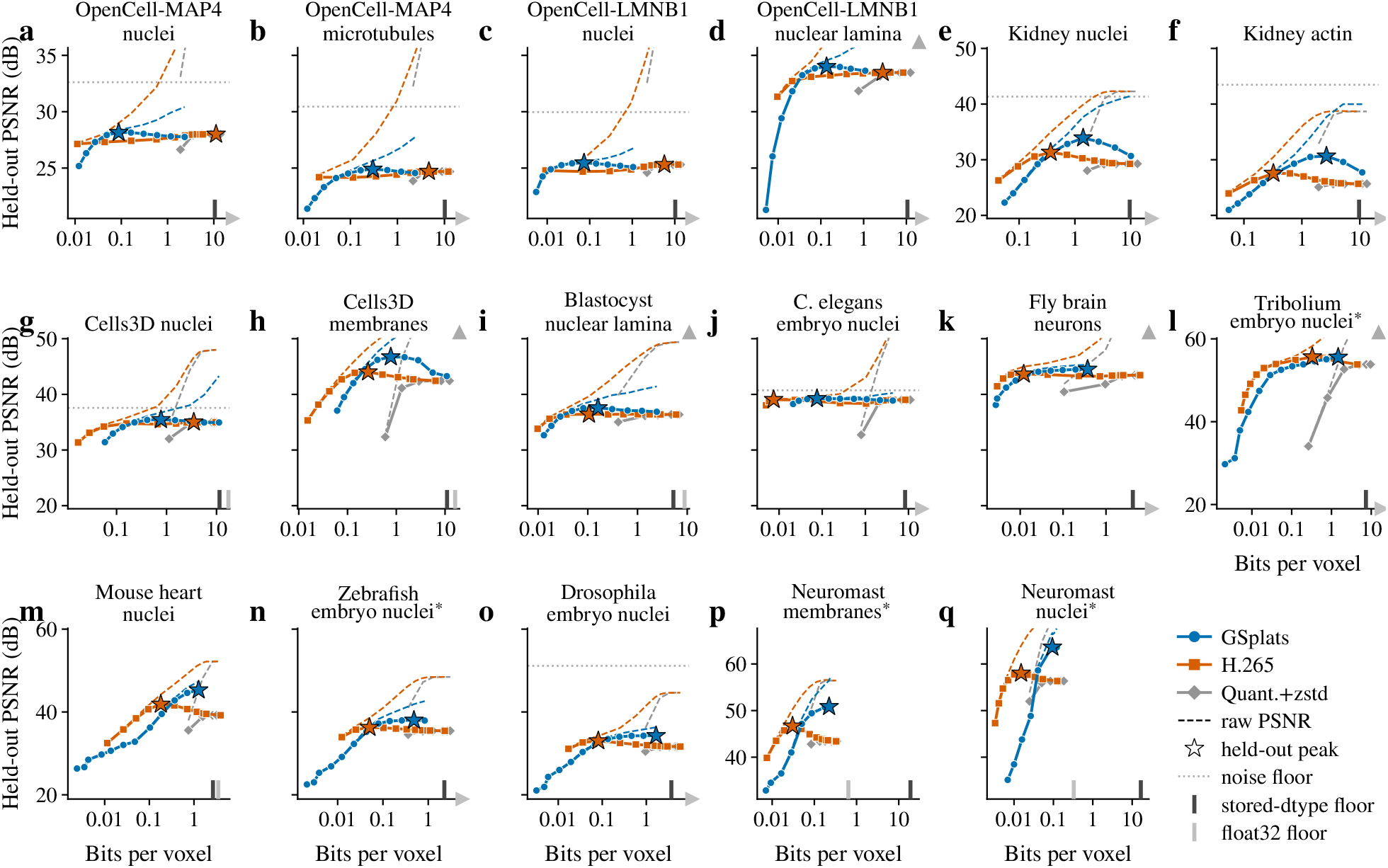
Cross-validation signal recovery: Gaussian splats vs. voxel codecs. Held-out PSNR (5% blind-spot mask) vs. bits per voxel for every benchmark volume with a completed sweep, one panel per volume in the order of Table 3 (Supplementary Document 7), spanning spinning-disk, confocal, light-sheet and iSIM modalities. Panel titles are the volume names of Table 3, an asterisk marking the deconvolved volumes; panels of one modality share a y range, and the first panel of each modality and of each row carries the y tick labels. Blue circles: GSplats. Orange squares: H.265 (Z-stack as video, constant rate factor 0–51). Grey diamonds: uniform quantisation (4–16 bit) + zstd, encoded from the same blind-spot-filled volume. Solid lines with markers: held-out PSNR. Thin dashed lines: raw PSNR against the noisy original (includes noise), which rises past the top of the panel at the highest rates. Stars: peak held-out PSNR for each method; on a curve still rising at its largest count the star sits at the endpoint. Dotted horizontal line: the noise-floor ceiling where it is estimable and lies within the shared y range; a grey triangle at the top right marks a ceiling above that range, and the volumes with no estimate carry neither. Ticks rising from the x axis: the lowest bit rate the tested lossless blosc configurations reach on that volume, on the source at its stored dtype (dark) and on the normalised float32 array (light; a light triangle at the right end marks a float32 floor beyond the plotted range); GSplats extend 175–3506*×* below the stored-dtype floor. Quantisation has no spatial model: at high precision its held-out PSNR approaches that of the mask-fill predictor however many bits it spends, and its raw PSNR beyond the noise floor is fidelity to detector noise, not to signal. **The two held-out curves cross**. H.265 leads at very low bit-rates, where its block-DCT structure efficiently encodes coarse content; at the bit rate of the GSplat operating point, with H.265 credited with the larger of its best result up to that budget and its interpolated value there, GSplats recover more held-out signal on 15 of 17 volumes and tie on 2; the stars, which compare each method’s peak at its own rate, favour GSplats on 12 of 17, with 5 inside the replicate band, the *Tribolium* embryo among them. Per-volume margins, the envelope rule and the comparison at H.265’s own peak rate are tabulated in Supplementary Document 7.

## Footnotes

* Exact encode invocation: `ffmpeg -y -f rawvideo -pix_fmt gray16le -s WxH -r 1 -i in.raw -c:v libx265 -preset medium -x265-params lossless={1 if crf==0 else 0} -crf N -pix_fmt gray12le out.mp4, decoded back with -f rawvideo` `-pix_fmt gray16le.`

## Notes

https://demos.luxarviewer.dev/

### Competing Interest Statement

The authors have declared no competing interest.

