## Supplementary material for "Luxar: Gaussian splatting for microscopy and scalable interactive web visualisation of multidimensional scientific data": supp_doc_01_merging_error_analysis

### Analytical Merging Error for Gaussian Splats: An $L^2$ Framework for the Moment-Matched Approximation

Supplementary Document 1 – Luxar

**Abstract**

Merging two Gaussian splats into one is the basic step of every level-of-detail reduction, and it loses information. We derive that loss in closed form. Within the $L^2$ (integrated squared error) framework the inner product of any two Gaussians has an analytical expression, and every term of the merging error reduces to such a product: the energy of the two-splat mixture, its overlap with the merged template, and the template's own energy. The merged splat takes its centre and covariance from moment matching and its amplitude from a one-line $L^2$ optimisation. The normalised error then equals $\sin^2\alpha$, where $\alpha$ is the angle in $L^2$ between the mixture and the template, so the score lies in $[0, 1]$ and is free of units. Two equal splats two standard deviations apart merge with a relative squared error of 1.2%. The inter-splat cosine, which needs one kernel evaluation and no template, is an amplitude-blind shortlist heuristic and no substitute for the score. Two caveats bound the result. The score is an upper bound on the error of the best single Gaussian, loose for well-separated pairs, because the moment-matched shape is $L^2$-optimal only for proportional splats. And the shipped level builder widens and rescales the representative it stores, so a stored level is a different object from the template analysed here, and its quality is measured directly on what is stored. Pseudocode for the score, the proxy and the merge is given.

#### Contents

**1 Introduction and motivation** 2

**2 The Gaussian $L^2$ inner product** 3

**3 The moment-matched approximation** 5

**4 The closed-form merging error** 7

**5 Cheaper proxy: the inter-splat cosine** 10

**6 Special cases** 11

**7 Algorithm** 13

**8 Optimality discussion** 13

**9 Summary** 18

### 1 Introduction and motivation

Gaussian splatting represents a scene as a collection of $N$ oriented Gaussians ("splats") [5]. Each splat is a smooth, bell-shaped density function in $D$-dimensional space, fully determined by three parameters: a *centre* $\boldsymbol{\mu}$ (its position), an *amplitude* $a$ (its peak height), and a *covariance matrix* $\boldsymbol{\Sigma}$ (which controls its shape, orientation, and width). For large scientific volumes, $N$ can reach millions, making simplification essential for interactive visualisation and efficient storage.

The most natural simplification primitive is *merging*: replacing two nearby or similar splats with a single splat that approximates their combined density as closely as possible (Fig. 1). Two questions follow: which pairs to merge first, and how much quality each merge costs. Both need a principled, closed-form measure of the **merging error**: the discrepancy between the two-splat mixture $\phi_1 + \phi_2$ and a single-splat approximation $g$ built by moment matching with an $L^2$-optimal amplitude. This document derives that measure, shows what it does and does not guarantee, and relates it to the merger Luxar ships.

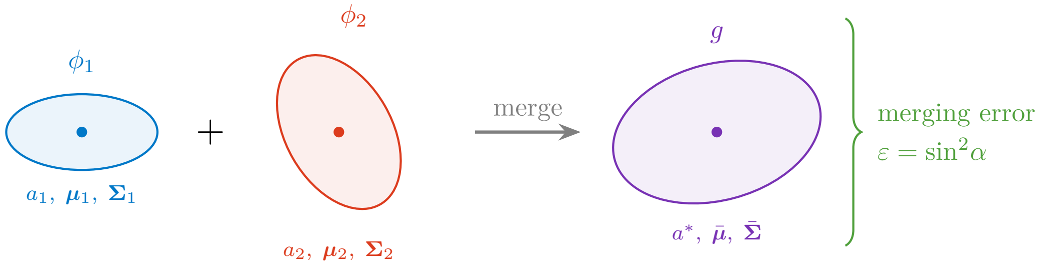


Figure 1: Schematic of the merging operation. Two oriented Gaussian splats $\phi_1$ and $\phi_2$ are replaced by a single splat $g$. The merging error $\varepsilon$ quantifies the fraction of the mixture's energy not captured by the moment-matched template (an upper bound on the error of the best single Gaussian).

**Choice of $L^2$.** To quantify how well a single Gaussian approximates a mixture, we need a *divergence*, a way to measure the "distance" between two functions. Several options exist (Table 1), but the $L^2$ (integrated squared error) framework is particularly convenient because *the integral of a product of two Gaussians has a closed-form expression.* The Cauchy–Schwarz divergence and the maximum mean discrepancy (MMD) with a Gaussian kernel are built from the same integrals, and Supplementary Document 8 uses them for the same reason. Every term in the error expansion reduces to such a product, yielding a fully analytical result. By contrast, the Kullback–Leibler divergence for a Gaussian mixture involves $\log(\phi_1 + \phi_2)$, which has no closed form [3]; the Hellinger distance requires $\sqrt{\phi_1 + \phi_2}$, equally intractable; and while the 2-Wasserstein distance between two individual Gaussians is elegant [1], it does not directly yield the error of approximating a *mixture*.

Table 1: Comparison of divergence measures for the merging problem. "Mixture error" refers to computing $d(\phi_1+\phi_2,\ g)$ analytically. KL, Hellinger and $W_2$ are defined between probability densities, so their rows assume that each function has been divided by its total mass; the $L^2$ row needs no such normalisation.

| **Divergence** | **Between two Gaussians** | **Mixture error** | **Optimal $g$** |
|---|---|---|---|
| $L^2$ (ISE) | Closed-form | **Closed-form** | Numerical† |
| KL | Closed-form | No closed-form | Moment-matched‡ |
| Hellinger | Closed-form | No closed-form | No closed-form |
| $W_2$ (Wasserstein) | Closed-form | No closed-form | Numerical§ |

† Amplitude is closed-form for a given shape; the $L^2$-optimal centre and covariance have no closed form (a local search from the moment-matched shape reaches a stationary point, not a certified global minimum).

‡ Minimises $\mathrm{KL}(f\|g)$ but the KL value itself has no closed form.

§ The Bures barycentre of the two components minimises $\sum_i w_i W_2^2(\phi_i, g)$, not $W_2^2(\phi_1+\phi_2, g)$; it is a surrogate, not the $W_2$-optimal Gaussian for the mixture.

**Outline.** Section 2 derives the Gaussian $L^2$ inner product. Section 3 constructs the moment-matched single-Gaussian approximation. Section 4 states the closed-form error and the $\sin^2\alpha$ mergeability score. Section 5 introduces a cheaper ranking proxy. Section 6 treats important special cases. Section 7 provides pseudocode. Section 8 discusses optimality and practical considerations.

##### Terms used in this document

**self-energy** The squared $L^2$ norm of one splat's Gaussian, $a_i^2\,\pi^{D/2}\,|\Sigma_i|^{1/2}$, the ordering weight of an additive ladder above the greedy size limit of 5,000 splats (below it a greedy residual ordering is used), the unit of the equal-energy breakpoints, and the quantity whose cumulative fraction stamps how much of a leaf a ladder prefix already carries.

**recipe** A named LOD topology built from a fitted splat set: `flat` (one bare leaf), `stream` (one stream-laddered leaf), `levels` (coarse-to-fine replacement levels), `tiles` (a BSP partition with one ladder per tile), `overview` (one coarse level above a `tiles` branch) and `adaptive` (`tiles` where every tile carries its own `levels`).

**Morton and Hilbert ordering** Two space-filling curves that sort points so that neighbours in space become neighbours in memory or on disk: Morton (Z-order) interleaves the bits of the quantised coordinates, Hilbert follows a continuous curve whose consecutive cells are always adjacent, so it has no long jumps between neighbouring runs. The compiler orders the stored elements along one of them; the fitter re-sorts its splats by Morton code periodically for GPU locality, and the substitutive LOD builder uses a Morton sort as its warm-start partition.

#### 2 The Gaussian $L^2$ inner product

##### 2.1 Setup and notation

We work in $\mathbb{R}^D$. A **Gaussian splat** is a function

$$\phi(\boldsymbol{x}) \;=\; a\,\exp\!\Big(-\tfrac{1}{2}\,(\boldsymbol{x}-\boldsymbol{\mu})^\top\boldsymbol{\Sigma}^{-1}(\boldsymbol{x}-\boldsymbol{\mu})\Big), \qquad a>0,\ \ \boldsymbol{\mu}\in\mathbb{R}^D,\ \ \boldsymbol{\Sigma}\succ 0 \tag{1}$$

where $\boldsymbol{\Sigma} \succ 0$ means $\boldsymbol{\Sigma}$ is symmetric positive definite (SPD; it defines a valid ellipsoidal shape). Where the parameters are written out we use $G(\boldsymbol{x}; \boldsymbol{\mu}, \boldsymbol{\Sigma})$ for the unit-amplitude kernel, so that $\phi(\boldsymbol{x}) = a\, G(\boldsymbol{x}; \boldsymbol{\mu}, \boldsymbol{\Sigma})$. The covariance is stored via its **Cholesky factor** $\boldsymbol{L}$, a lower-triangular matrix satisfying $\boldsymbol{\Sigma} = \boldsymbol{L}\boldsymbol{L}^\top$ (analogous to taking the "square root" of $\boldsymbol{\Sigma}$). We also write $\boldsymbol{P} = \boldsymbol{\Sigma}^{-1}$ for the **precision matrix** (the inverse covariance, which appears naturally in the Gaussian exponent). In the Luxar implementation, $\boldsymbol{L}$ is stored as its $D(D+1)/2$ lower-triangular elements, split on disk into a diagonal array and an off-diagonal array.

The $L^2$ **inner product** and norm used throughout are

$$\langle f, g \rangle_{L^2} = \int_{\mathbb{R}^D} f(\boldsymbol{x})\, g(\boldsymbol{x})\, d\boldsymbol{x}, \qquad \|f\|_{L^2}^2 = \langle f, f \rangle_{L^2}. \tag{2}$$

Every inner product and norm in this document is the $L^2$ one, so the subscript is dropped from here on. Since $a > 0$ and $\boldsymbol{\Sigma} \succ 0$, every splat has finite energy $\|\phi\|^2 < \infty$ and all inner products below are well-defined. Centre differences are written $\Delta\boldsymbol{\mu}_{ij} = \boldsymbol{\mu}_i - \boldsymbol{\mu}_j$; for the pair $(1, 2)$ the indices are dropped, $\Delta\boldsymbol{\mu} = \Delta\boldsymbol{\mu}_{12}$.

#### 2.2 Derivation of the kernel

The inner product $\langle \phi_i, \phi_j \rangle$ measures the *geometric overlap* between two Gaussians: it is large when they sit close together with similar shapes, and becomes small when they are far apart or their covariances differ strongly (for finite parameters it is strictly positive). The central result of this section is that this overlap has a simple closed-form expression. Its proof uses one matrix identity, stated first.

**Lemma 2.1.** *For symmetric positive definite* $\boldsymbol{\Sigma}_i, \boldsymbol{\Sigma}_j$*:*

$$\boldsymbol{\Sigma}_i^{-1}(\boldsymbol{\Sigma}_i^{-1} + \boldsymbol{\Sigma}_j^{-1})^{-1}\boldsymbol{\Sigma}_j^{-1} = (\boldsymbol{\Sigma}_i + \boldsymbol{\Sigma}_j)^{-1}. \tag{3}$$

*Proof.* Multiply both sides on the left by $\boldsymbol{\Sigma}_i$ and on the right by $\boldsymbol{\Sigma}_j$. The left side becomes $(\boldsymbol{\Sigma}_i^{-1} + \boldsymbol{\Sigma}_j^{-1})^{-1}$ and the right side becomes $\boldsymbol{\Sigma}_i(\boldsymbol{\Sigma}_i + \boldsymbol{\Sigma}_j)^{-1}\boldsymbol{\Sigma}_j$. Taking the inverse of both sides: $\boldsymbol{\Sigma}_i^{-1} + \boldsymbol{\Sigma}_j^{-1}$ versus $\boldsymbol{\Sigma}_j^{-1}(\boldsymbol{\Sigma}_i + \boldsymbol{\Sigma}_j)\boldsymbol{\Sigma}_i^{-1} = \boldsymbol{\Sigma}_j^{-1} + \boldsymbol{\Sigma}_i^{-1}$. These are equal since matrix addition commutes. □

**Theorem 2.2** (Gaussian $L^2$ inner product)**.** *For two splats* $\phi_i$*,* $\phi_j$ *as in* (1)*,*

$$\boxed{\langle \phi_i, \phi_j \rangle \;=\; a_i\, a_j\; (2\pi)^{D/2}\, \frac{\sqrt{|\boldsymbol{\Sigma}_i|\,|\boldsymbol{\Sigma}_j|}}{\sqrt{|\boldsymbol{\Sigma}_i + \boldsymbol{\Sigma}_j|}}\; \exp\!\Big(-\tfrac{1}{2}\, \Delta\boldsymbol{\mu}_{ij}^\top (\boldsymbol{\Sigma}_i + \boldsymbol{\Sigma}_j)^{-1}\, \Delta\boldsymbol{\mu}_{ij}\Big)} \tag{4}$$

*with* $\Delta\boldsymbol{\mu}_{ij} = \boldsymbol{\mu}_i - \boldsymbol{\mu}_j$ *as above.*

*Proof.* Write the product of the two Gaussian kernels:

$$\phi_i(\boldsymbol{x})\, \phi_j(\boldsymbol{x}) = a_i a_j \exp\!\big(-\tfrac{1}{2}(Q_i + Q_j)\big), \tag{5}$$

where $Q_k = (\boldsymbol{x} - \boldsymbol{\mu}_k)^\top \boldsymbol{\Sigma}_k^{-1}(\boldsymbol{x} - \boldsymbol{\mu}_k)$. The sum of quadratic forms is itself quadratic in $\boldsymbol{x}$:

$$Q_i + Q_j = \boldsymbol{x}^\top \underbrace{(\boldsymbol{\Sigma}_i^{-1} + \boldsymbol{\Sigma}_j^{-1})}_{\boldsymbol{P}_{ij}} \boldsymbol{x} - 2\,\boldsymbol{x}^\top \underbrace{(\boldsymbol{\Sigma}_i^{-1}\boldsymbol{\mu}_i + \boldsymbol{\Sigma}_j^{-1}\boldsymbol{\mu}_j)}_{\boldsymbol{\eta}} + \underbrace{\boldsymbol{\mu}_i^\top \boldsymbol{\Sigma}_i^{-1}\boldsymbol{\mu}_i + \boldsymbol{\mu}_j^\top \boldsymbol{\Sigma}_j^{-1}\boldsymbol{\mu}_j}_{c}. \tag{6}$$

Completing the square:

$$Q_i + Q_j = (\boldsymbol{x} - \boldsymbol{\mu}_{ij})^\top \boldsymbol{P}_{ij}\,(\boldsymbol{x} - \boldsymbol{\mu}_{ij}) + (c - \boldsymbol{\eta}^\top \boldsymbol{P}_{ij}^{-1}\,\boldsymbol{\eta}), \tag{7}$$

where $\boldsymbol{\mu}_{ij} = \boldsymbol{P}_{ij}^{-1}\boldsymbol{\eta}$ is the "merged mean" in precision space. Integrating the Gaussian part over $\mathbb{R}^D$:

$$\int_{\mathbb{R}^D} \exp\!\left(-\tfrac{1}{2}(\boldsymbol{x} - \boldsymbol{\mu}_{ij})^\top \boldsymbol{P}_{ij}\,(\boldsymbol{x} - \boldsymbol{\mu}_{ij})\right) d\boldsymbol{x} = (2\pi)^{D/2}\,|\boldsymbol{P}_{ij}^{-1}|^{1/2} = \frac{(2\pi)^{D/2}}{|\boldsymbol{P}_{ij}|^{1/2}}. \tag{8}$$

It remains to simplify the constant $c - \boldsymbol{\eta}^\top \boldsymbol{P}_{ij}^{-1}\boldsymbol{\eta}$. Expand $\boldsymbol{\eta}^\top \boldsymbol{P}_{ij}^{-1}\boldsymbol{\eta}$ with $\boldsymbol{\eta} = \boldsymbol{\Sigma}_i^{-1}\boldsymbol{\mu}_i + \boldsymbol{\Sigma}_j^{-1}\boldsymbol{\mu}_j$. The pure terms in $\boldsymbol{\mu}_i$ combine through $\boldsymbol{\Sigma}_i^{-1} - \boldsymbol{\Sigma}_i^{-1}\boldsymbol{P}_{ij}^{-1}\boldsymbol{\Sigma}_i^{-1} = \boldsymbol{\Sigma}_i^{-1}\boldsymbol{P}_{ij}^{-1}\boldsymbol{\Sigma}_j^{-1}$, which holds because $\boldsymbol{P}_{ij} - \boldsymbol{\Sigma}_i^{-1} = \boldsymbol{\Sigma}_j^{-1}$, and the pure terms in $\boldsymbol{\mu}_j$ combine through the $j$-counterpart of the same identity. Together with the cross terms they collect into a single quadratic form in $\Delta\boldsymbol{\mu}_{ij}$, which gives the first equality below. Lemma 2.1, whose right-hand side shows that the matrix is symmetric, gives the second:

$$c - \boldsymbol{\eta}^\top \boldsymbol{P}_{ij}^{-1}\boldsymbol{\eta} = \Delta\boldsymbol{\mu}_{ij}^\top\,\boldsymbol{\Sigma}_i^{-1}\,\boldsymbol{P}_{ij}^{-1}\,\boldsymbol{\Sigma}_j^{-1}\,\Delta\boldsymbol{\mu}_{ij} = \Delta\boldsymbol{\mu}_{ij}^\top\,(\boldsymbol{\Sigma}_i + \boldsymbol{\Sigma}_j)^{-1}\,\Delta\boldsymbol{\mu}_{ij}, \tag{9}$$

and the determinant (using the harmonic-mean identity $(\boldsymbol{\Sigma}_i^{-1}+\boldsymbol{\Sigma}_j^{-1})^{-1} = \boldsymbol{\Sigma}_i(\boldsymbol{\Sigma}_i+\boldsymbol{\Sigma}_j)^{-1}\boldsymbol{\Sigma}_j$, which follows from Lemma 2.1 by multiplying (3) on the left by $\boldsymbol{\Sigma}_i$ and on the right by $\boldsymbol{\Sigma}_j$):

$$|\boldsymbol{P}_{ij}^{-1}| = |(\boldsymbol{\Sigma}_i^{-1} + \boldsymbol{\Sigma}_j^{-1})^{-1}| = |\boldsymbol{\Sigma}_i(\boldsymbol{\Sigma}_i + \boldsymbol{\Sigma}_j)^{-1}\boldsymbol{\Sigma}_j| = \frac{|\boldsymbol{\Sigma}_i|\,|\boldsymbol{\Sigma}_j|}{|\boldsymbol{\Sigma}_i + \boldsymbol{\Sigma}_j|}. \tag{10}$$

Combining: the integral of (5) factors as $(2\pi)^{D/2}\,|\boldsymbol{P}_{ij}^{-1}|^{1/2}$ from (7) times $\exp(-\frac{1}{2}\cdot(9))$. Rewriting $|\boldsymbol{P}_{ij}^{-1}|$ via (10) yields (4). □

**Corollary 2.3** (Self inner product)**.** *Setting $i = j$ and using $|2\boldsymbol{\Sigma}| = 2^D\,|\boldsymbol{\Sigma}|$:*

$$\|\phi_i\|^2 = a_i^2\,\pi^{D/2}\,|\boldsymbol{\Sigma}_i|^{1/2}. \tag{11}$$

*Remark* 2.4 (In terms of Cholesky factors). Since $\boldsymbol{\Sigma}_i = \boldsymbol{L}_i\boldsymbol{L}_i^\top$, we have $|\boldsymbol{\Sigma}_i|^{1/2} = |\det \boldsymbol{L}_i| = \prod_{k=1}^{D}(\boldsymbol{L}_i)_{kk}$. To evaluate the kernel (4), form $\boldsymbol{S} = \boldsymbol{\Sigma}_1+\boldsymbol{\Sigma}_2 = \boldsymbol{L}_1\boldsymbol{L}_1^\top+\boldsymbol{L}_2\boldsymbol{L}_2^\top$, compute $\boldsymbol{R} = \mathrm{chol}(\boldsymbol{S})$, and then $|\boldsymbol{\Sigma}_1 + \boldsymbol{\Sigma}_2|^{1/2} = \prod_k \boldsymbol{R}_{kk}$. The Mahalanobis term $(\boldsymbol{\Sigma}_1+\boldsymbol{\Sigma}_2)^{-1}\Delta\boldsymbol{\mu}$ is obtained by forward–backward substitution with $\boldsymbol{R}$, avoiding explicit matrix inversion.

#### 3 The moment-matched approximation

With the closed-form kernel from Section 2, we construct the moment-matched single-Gaussian replacement for a pair of splats. The error of that replacement is the subject of Section 4.

Given two splats $\phi_1, \phi_2$, we seek a single splat $g(\boldsymbol{x}) = a\,\bar{G}(\boldsymbol{x})$ that approximates their sum $f = \phi_1 + \phi_2$, with the shape fixed by moment matching and the amplitude chosen optimally for that shape. Throughout, we write $G_i(\boldsymbol{x}) = G(\boldsymbol{x}; \boldsymbol{\mu}_i, \boldsymbol{\Sigma}_i)$ for the **unit-amplitude kernel** of splat $i$, so that $\phi_i = a_i\, G_i$. Similarly, $\bar{G}(\boldsymbol{x}) = G(\boldsymbol{x}; \bar{\boldsymbol{\mu}}, \bar{\boldsymbol{\Sigma}})$ denotes the unit-amplitude template for the merged splat.

##### 3.1 Motivation for moment matching

Any density function can be summarised by its *moments*: the total mass (zeroth moment), the mean position (first moment), and the spread or covariance (second moment). A single Gaussian is fully determined by its mean and covariance, so matching these moments is the most natural way to "compress" a mixture into one Gaussian.

This intuition has a rigorous justification. The Gaussian family forms an *exponential family* whose sufficient statistics are $\boldsymbol{x}$ and $\boldsymbol{x}\boldsymbol{x}^\top$ [2], and a classical result from information geometry guarantees that the minimiser of the Kullback–Leibler (KL) divergence from a target distribution to a single Gaussian matches the target's mean and covariance [2, 9].

Applied to our problem: we first normalise $f = \phi_1 + \phi_2$ to a probability distribution, find the Gaussian whose mean and covariance match (determining $\bar{\boldsymbol{\mu}}$ and $\bar{\boldsymbol{\Sigma}}$), and then set the amplitude separately, either by mass conservation ($\bar{a}$, Section 3.2) or by $L^2$ optimisation ($a^*$, Section 4.2). The moment-matched Gaussian is therefore the natural starting point for a single-component approximation, and its shape parameters have fully closed-form expressions. Whether it is also the $L^2$-optimal single Gaussian is a separate question, taken up in Section 8.

##### 3.2 Closed-form parameters

Define the **mass** of each splat and the mass-based weights:

$$m_i = a_i\,(2\pi)^{D/2}\,|\boldsymbol{\Sigma}_i|^{1/2} = \int_{\mathbb{R}^D} \phi_i(\boldsymbol{x})\,d\boldsymbol{x}, \qquad w_i = \frac{m_i}{m_1 + m_2}. \tag{12}$$

**Proposition 3.1** (Moment-matched parameters)**.** *The moment-matched single Gaussian $g$ has parameters:*

$$\bar{\boldsymbol{\mu}} = w_1\,\boldsymbol{\mu}_1 + w_2\,\boldsymbol{\mu}_2, \tag{13}$$

$$\bar{\boldsymbol{\Sigma}} = w_1\,\boldsymbol{\Sigma}_1 + w_2\,\boldsymbol{\Sigma}_2 + w_1 w_2\,\Delta\boldsymbol{\mu}\,\Delta\boldsymbol{\mu}^\top, \tag{14}$$

$$\bar{a} = \frac{m_1 + m_2}{(2\pi)^{D/2}\,|\bar{\boldsymbol{\Sigma}}|^{1/2}} = \frac{a_1|\boldsymbol{\Sigma}_1|^{1/2} + a_2|\boldsymbol{\Sigma}_2|^{1/2}}{|\bar{\boldsymbol{\Sigma}}|^{1/2}}, \tag{15}$$

*with $\Delta\boldsymbol{\mu} = \boldsymbol{\mu}_1 - \boldsymbol{\mu}_2$.*

*Proof.* Treat the mixture as an unnormalised density with total mass $m_1 + m_2$. Normalising: $p(\boldsymbol{x}) = w_1\,\mathcal{N}(\boldsymbol{x}; \boldsymbol{\mu}_1, \boldsymbol{\Sigma}_1) + w_2\,\mathcal{N}(\boldsymbol{x}; \boldsymbol{\mu}_2, \boldsymbol{\Sigma}_2)$. The mean is $\bar{\boldsymbol{\mu}} = w_1\boldsymbol{\mu}_1 + w_2\boldsymbol{\mu}_2$ by linearity. The covariance uses the law of total variance:

$$\bar{\boldsymbol{\Sigma}} = \sum_i w_i \, \boldsymbol{\Sigma}_i + \sum_i w_i (\boldsymbol{\mu}_i - \bar{\boldsymbol{\mu}})(\boldsymbol{\mu}_i - \bar{\boldsymbol{\mu}})^\top. \tag{16}$$

Substituting $\boldsymbol{\mu}_i - \bar{\boldsymbol{\mu}} = w_j \, (\boldsymbol{\mu}_i - \boldsymbol{\mu}_j)$ (where $j \neq i$) and $w_1 w_2^2 + w_2 w_1^2 = w_1 w_2$ yields (14). The amplitude (15) enforces mass conservation $\int g = m_1 + m_2$. □

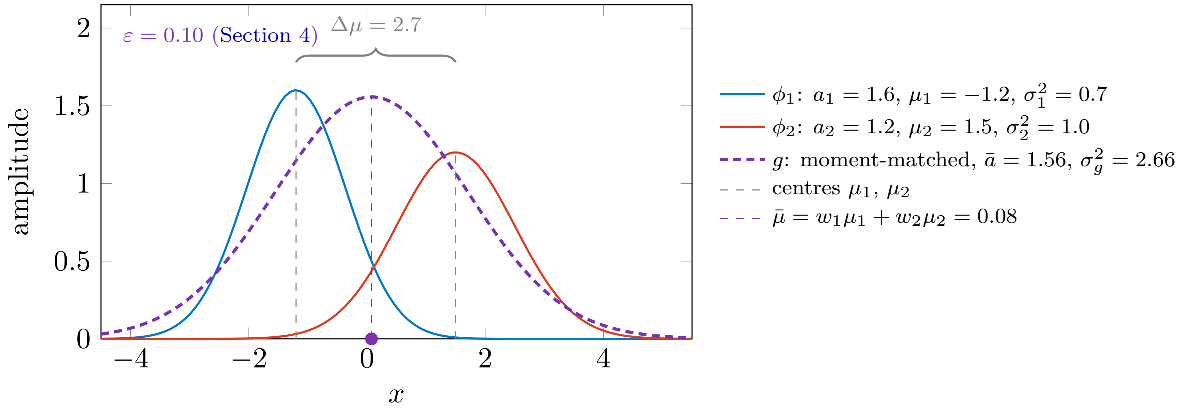


Figure 2: The moment-matched Gaussian $g$ (purple dashed) has its mean $\bar{\boldsymbol{\mu}}$ at the mass-weighted average of the two centres, and its covariance includes a rank-1 "spreading" term $w_1 w_2 \, \Delta\boldsymbol{\mu} \, \Delta\boldsymbol{\mu}^\top$ due to the separation: here $\sigma_g^2 = 2.66$ against the intrinsic $w_1\sigma_1^2 + w_2\sigma_2^2 = 0.84$. The amplitude shown is the mass-preserving $\bar{a}$ of (15); with the optimised amplitude of Section 4 the pair scores $\varepsilon = 0.10$.

*Remark* 3.2 (Geometric interpretation of $\bar{\boldsymbol{\Sigma}}$). The moment-matched covariance decomposes as $\bar{\boldsymbol{\Sigma}} = \bar{\boldsymbol{\Sigma}}_{\text{intra}} + \bar{\boldsymbol{\Sigma}}_{\text{inter}}$, where $\bar{\boldsymbol{\Sigma}}_{\text{intra}} = w_1 \boldsymbol{\Sigma}_1 + w_2 \boldsymbol{\Sigma}_2$ is the weighted average of the individual spreads (the "intrinsic" width), and $\bar{\boldsymbol{\Sigma}}_{\text{inter}} = w_1 w_2 \, \Delta\boldsymbol{\mu} \, \Delta\boldsymbol{\mu}^\top$ is a rank-1 term encoding the "extrinsic" spread from the separation of the two centres. When $\boldsymbol{\mu}_1 = \boldsymbol{\mu}_2$, the extrinsic term vanishes. Figure 2 shows the merge for a one-dimensional pair and Fig. 3 the two terms for an anisotropic pair in two dimensions.

### 4 The closed-form merging error

#### 4.1 Error decomposition

For *any* single-Gaussian approximation $g$, the $L^2$ error is:

$$E = \|\phi_1 + \phi_2 - g\|^2 = \underbrace{K_{11} + 2\,K_{12} + K_{22}}_{\|f\|^2} - 2\,K_{1g} - 2\,K_{2g} + K_{gg}, \tag{17}$$

where $K_{ij} = \langle \phi_i, \phi_j \rangle$ and all six terms use the kernel from Theorem 2.2. Figure 4 shows the residual whose squared integral is this error, for the mass-preserving amplitude of Proposition 3.1.

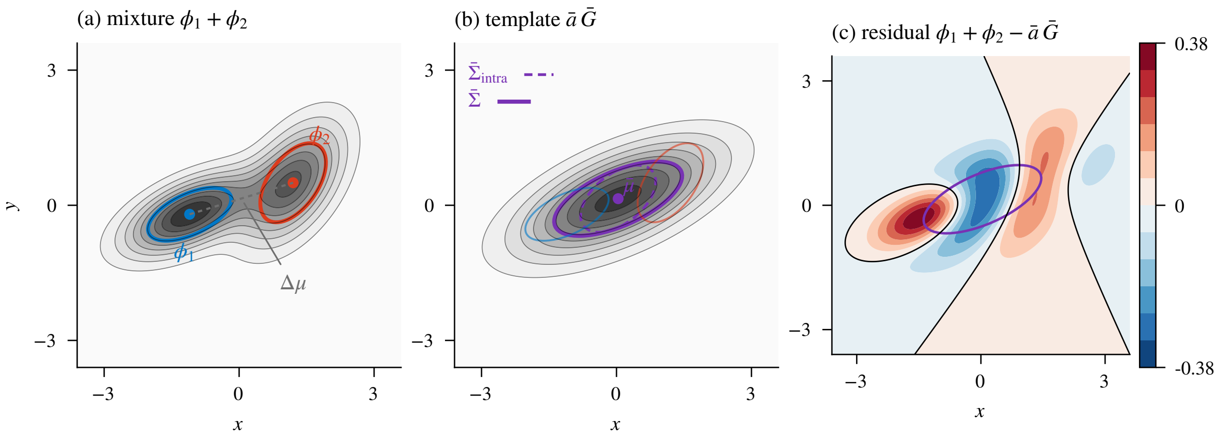


Figure 3: A moment-matched merge in two dimensions. (a) Two anisotropic splats, $a_1 = 1$ and $a_2 = 0.8$, with their $1\sigma$ ellipses and centres; grey levels show the mixture $\phi_1 + \phi_2$. (b) The template $\bar{a}\,\bar{G}$ of Proposition 3.1 on the same grey levels, with the $1\sigma$ ellipse of $\bar{\boldsymbol{\Sigma}}_{\text{intra}}$ (dashed) and of $\bar{\boldsymbol{\Sigma}}$ (solid): the rank-1 term $w_1 w_2\, \Delta\boldsymbol{\mu}\, \Delta\boldsymbol{\mu}^\top$ stretches the ellipse along $\Delta\boldsymbol{\mu}$ only, so the template is longer than either component in that direction and, relative to $\bar{\boldsymbol{\Sigma}}_{\text{intra}}$, no wider across it. (c) The residual $\phi_1 + \phi_2 - \bar{a}\,\bar{G}$ with its zero contour in black: the template is too low at the two centres and too high between them and far beyond them, the pattern of two modes that one Gaussian cannot resolve. Its squared integral is 10.5% of the mixture energy (10.4% with the optimised amplitude of Section 4).

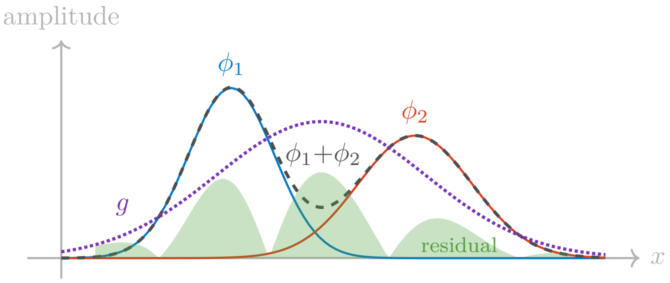


Figure 4: One-dimensional illustration. The mixture $\phi_1{+}\phi_2$ (dashed) cannot be exactly represented by a single Gaussian $g$ (dotted; here the moment-matched Gaussian of Proposition 3.1 with the mass-preserving amplitude $\bar{a}$). The shaded area (schematic) represents the absolute residual, whose squared values integrate to the error $E$ of (17) for that amplitude (the optimised amplitude of Section 4.2 lowers it to $E^*$).

#### 4.2 Amplitude optimisation

The mass-preserving amplitude $\bar{a}$ from (15) is not the $L^2$-optimal amplitude in general. We fix the moment-matched shape parameters $(\bar{\boldsymbol{\mu}}, \bar{\boldsymbol{\Sigma}})$, and with them the unit-amplitude template $\bar{G}$ of Section 3, and choose the amplitude that minimises the $L^2$ error. Since, for a generic amplitude $a$, $E = \|f\|^2 - 2a\,\langle f, \bar{G}\rangle + a^2\,\|\bar{G}\|^2$ is quadratic in $a$, the minimiser is

$$a^* = \frac{\langle f, \bar{G}\rangle}{\|\bar{G}\|^2} = \frac{a_1\langle G_1, \bar{G}\rangle + a_2\langle G_2, \bar{G}\rangle}{\pi^{D/2}\,|\bar{\boldsymbol{\Sigma}}|^{1/2}}, \tag{18}$$

and the corresponding minimum error is the classical **projection residual**:

$$\boxed{E^* = \|f\|^2 - \frac{|\langle f, \bar{G}\rangle|^2}{\|\bar{G}\|^2}} \tag{19}$$

This error is never worse than the mass-preserving one, and it is still fully closed-form.

##### 4.3 The mergeability score

The error $E^*$ has the units of the energy $\|f\|^2$ (amplitude squared times length$^D$), which depend on the amplitudes and dimension. To obtain a universal, dimensionless measure, we normalise by the mixture's total energy $\|f\|^2$. The result has a geometric interpretation (Fig. 5): think of the mixture $f$ and the template Gaussian $\bar{G}$ as vectors in the infinite-dimensional Hilbert space of square-integrable functions. The best amplitude $a^*$ is the scalar projection of $f$ onto $\bar{G}$, exactly as in ordinary Euclidean geometry, and the relative error is the sine squared of the angle between them:

$$\boxed{\varepsilon = \frac{E^*}{\|f\|^2} = 1 - \frac{|\langle f, \bar{G}\rangle|^2}{\|f\|^2\,\|\bar{G}\|^2} = 1 - \cos^2\alpha = \sin^2\alpha} \tag{20}$$

where $\alpha$ is the **angle between $f$ and $\bar{G}$ in $L^2$** (the Hilbert space of square-integrable functions).

- $\varepsilon = 0$ ($\alpha = 0$): the two splats are proportional ($\phi_1 \propto \phi_2$), and the merge is perfect.
- $\varepsilon \to 1$ ($\alpha \to \pi/2$): the mixture is nearly orthogonal to the template $\bar{G}$, so no amplitude can make $a\bar{G}$ resemble it, and the merge is lossy.

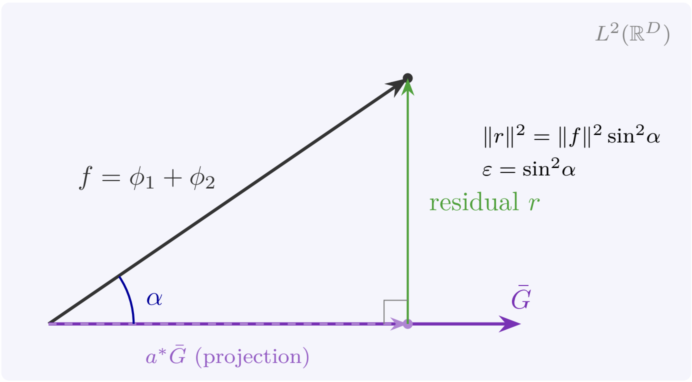


Figure 5: Geometric interpretation in $L^2$. The mixture $f = \phi_1 + \phi_2$ is a vector in the Hilbert space. Its projection onto the ray spanned by $\bar{G}$ (the unit-amplitude template) gives the optimal approximation $a^*\bar{G}$ (amplitude $a^*$). The residual $r = f - a^*\bar{G}$ is orthogonal to $\bar{G}$, and the normalised squared residual is $\varepsilon = \sin^2\alpha$.

##### 4.4 Expanding the score

For implementation, we expand the inner products in the mergeability score into explicit kernel evaluations. Writing out all terms in (20) using the kernel (4):

$$\varepsilon = 1 - \frac{\left(a_1 I_1 + a_2 I_2\right)^2}{\left(a_1^2 J_1 + 2\, a_1 a_2\, I_{12} + a_2^2 J_2\right) \;\cdot\; J_g} \tag{21}$$

with the four types of kernel evaluation listed in Table 2.

Table 2: The four types of kernel evaluation needed for the mergeability score ($J_i$ and $I_i$ each stand for two evaluations, $i = 1, 2$). All use the master formula (4).

| Symbol | Role | Expression |
|---|---|---|
| $J_i = \dfrac{\\|\phi_i\\|^2}{a_i^2}$ | Self-energy (per unit $a_i^2$) | $\pi^{D/2}\, \lvert\boldsymbol{\Sigma}_i\rvert^{1/2}$ |
| $J_g = \\|\bar{G}\\|^2$ | Template self-energy | $\pi^{D/2}\, \lvert\bar{\boldsymbol{\Sigma}}\rvert^{1/2}$ |
| $I_{12} = \langle G_1, G_2\rangle$ | Cross-kernel (input splats) | $(2\pi)^{D/2}\, \dfrac{\sqrt{\lvert\boldsymbol{\Sigma}_1\rvert\, \lvert\boldsymbol{\Sigma}_2\rvert}}{\sqrt{\lvert\boldsymbol{\Sigma}_1 + \boldsymbol{\Sigma}_2\rvert}}\; e^{-Q_{12}/2}$ |
| $I_i = \langle G_i, \bar{G}\rangle$ | Cross-kernel (splat vs. template) | $(2\pi)^{D/2}\, \dfrac{\sqrt{\lvert\boldsymbol{\Sigma}_i\rvert\, \lvert\bar{\boldsymbol{\Sigma}}\rvert}}{\sqrt{\lvert\boldsymbol{\Sigma}_i + \bar{\boldsymbol{\Sigma}}\rvert}}\; e^{-\bar{Q}_i/2}$ |

$Q_{12} = \Delta\boldsymbol{\mu}^\top(\boldsymbol{\Sigma}_1{+}\boldsymbol{\Sigma}_2)^{-1}\Delta\boldsymbol{\mu}$, $\quad \bar{Q}_i = (\bar{\boldsymbol{\mu}}{-}\boldsymbol{\mu}_i)^\top(\boldsymbol{\Sigma}_i{+}\bar{\boldsymbol{\Sigma}})^{-1}(\bar{\boldsymbol{\mu}}{-}\boldsymbol{\mu}_i)$.

#### 5 Cheaper proxy: the inter-splat cosine

Computing the full mergeability score requires four types of kernel evaluation (six per pair; four once the self-energies $J_i$, the *self-energy* of each splat per unit $a_i^2$, are precomputed) and forming the moment-matched covariance. When the goal is merely to *rank* splat pairs by mergeability (e.g., to select the best pair to merge next), a simpler proxy is a useful shortlist heuristic:

**Proposition 5.1** (Inter-splat $L^2$ cosine)**.** *The cosine similarity between two splats in $L^2$ is amplitude-independent:*

$$\boxed{\cos\theta = \frac{\langle\phi_1, \phi_2\rangle}{\|\phi_1\|\;\|\phi_2\|} = 2^{D/2}\, \frac{(\lvert\boldsymbol{\Sigma}_1\rvert\, \lvert\boldsymbol{\Sigma}_2\rvert)^{1/4}}{\lvert\boldsymbol{\Sigma}_1 + \boldsymbol{\Sigma}_2\rvert^{1/2}}\; \exp\!\left(-\tfrac{1}{2}\, Q_{12}\right)} \tag{22}$$

*where $Q_{12} = \Delta\boldsymbol{\mu}^\top(\boldsymbol{\Sigma}_1 + \boldsymbol{\Sigma}_2)^{-1}\Delta\boldsymbol{\mu}$.*

*Proof.* Direct computation from (4) and (11): the amplitudes $a_1, a_2$ cancel in the ratio, and $(2\pi)^{D/2}/(\pi^{D/2})^{1/2+1/2} = 2^{D/2}$. □

**Properties.**

- $\cos\theta = 1$ iff $\boldsymbol{\Sigma}_1 = \boldsymbol{\Sigma}_2$ and $\boldsymbol{\mu}_1 = \boldsymbol{\mu}_2$ (identical shape and location).
- $\cos\theta \to 0$ as $Q_{12} \to \infty$ (well-separated) or as the covariance shapes diverge.
- Requires only **one** kernel evaluation (plus trivial self-products).
- Correlated with $\varepsilon$ for pairs of comparable amplitude, but blind to amplitude ratios, whereas $\varepsilon$ depends on $a_1/a_2$ through $\|f\|^2$. Table 3 shows the failure mode: the proxy ranks a nearer

equal pair as more mergeable than a faint splat farther from a bright one, although the equal pair's $\varepsilon$ is a thousand times larger. Figure 6 traces $\varepsilon$ against separation and against width ratio at several amplitude ratios, with the proxy overlaid. Use the proxy to shortlist candidates and the full score to accept a merge. The proxy's ranking has not been validated against $\varepsilon$ over amplitude and covariance ranges.

Table 3: Two pairs of splats with equal covariances $\sigma^2\boldsymbol{I}$ and the two rankings they receive. The proxy prefers the first pair; the score prefers the second by three orders of magnitude.

| **Pair** | **Separation** | $\cos\theta$ | $\varepsilon$ |
|---|---|---|---|
| $a = (1, 1)$ | $d = 2\sigma$ | 0.37 | $1.2 \times 10^{-2}$ |
| $a = (1, 0.001)$ | $d = 3\sigma$ | 0.11 | $1.2 \times 10^{-5}$ |

*Remark* 5.2 (Relationship between $\theta$ and $\alpha$). The inter-splat angle $\theta$ (between $\phi_1$ and $\phi_2$) and the mergeability angle $\alpha$ (between the mixture $f$ and $\bar{G}$) are distinct quantities. They agree at one extreme: $\cos\theta = 1 \Leftrightarrow \varepsilon = 0$ (proportional splats). At the other, $\cos\theta \to 0$ does not force $\varepsilon \to 1$: two co-located Gaussians of comparable amplitude and very different widths have $\cos\theta \to 0$ while the moment-matched template absorbs the broad component and $\varepsilon \to 0$ (in $D = 3$ with $\sigma_2 = 10\sigma_1$: $\cos\theta = 0.09$ for any amplitudes, while $\varepsilon = 0.001$ at $a_1 = a_2$ but 0.85 at $a_1 = 100\, a_2$). Only for well-separated splats of fixed, comparable widths and weights does $\varepsilon$ approach 1. In between, the relationship is non-trivial because $\alpha$ depends on the moment-matched template $\bar{G}$, while $\theta$ is a direct pairwise measure. $\cos\theta$ remains a convenient shortlist heuristic.

### 6 Special cases

To build intuition for the general formulas, we examine three limiting cases where the expressions simplify and yield practical rules of thumb.

#### 6.1 Equal isotropic covariances

When $\boldsymbol{\Sigma}_1 = \boldsymbol{\Sigma}_2 = \sigma^2\boldsymbol{I}_D$, the kernel simplifies. Let $d = \|\Delta\boldsymbol{\mu}\|$:

$$K_{12} = a_1 a_2\, \pi^{D/2}\, \sigma^D\, e^{-d^2/(4\sigma^2)}, \tag{23}$$

$$\|f\|^2 = (a_1^2 + a_2^2)\, \pi^{D/2}\, \sigma^D + 2\, a_1 a_2\, \pi^{D/2}\, \sigma^D\, e^{-d^2/(4\sigma^2)}, \tag{24}$$

$$\cos\theta = e^{-d^2/(4\sigma^2)}. \tag{25}$$

The dimension-dependence vanishes because the $2^{D/2}$ prefactor in (22) exactly cancels the determinant ratio $|\sigma^2\boldsymbol{I}|^{1/2}/|2\sigma^2\boldsymbol{I}|^{1/2} = 2^{-D/2}$ for equal covariances. The inter-splat cosine thus depends only on the ratio $d/\sigma$ (Fig. 7), and so does the score: with $\boldsymbol{\Sigma}_1 = \boldsymbol{\Sigma}_2 = \sigma^2\boldsymbol{I}_D$ every direction orthogonal to $\Delta\boldsymbol{\mu}$ contributes the same factor to $\langle f, \bar{G}\rangle^2$ and to $\|f\|^2\, \|\bar{G}\|^2$, so $\varepsilon$ depends on $d/\sigma$ and $a_2/a_1$ alone and takes its one-dimensional value in every $D$. This offers a simple rule of thumb: at $d = 2\sigma$ (where $\cos\theta = e^{-1} \approx 0.37$) two equal splats merge with a relative squared error $\varepsilon = 1.2\%$; the acceptable tolerance is the user's, and the full score should decide.

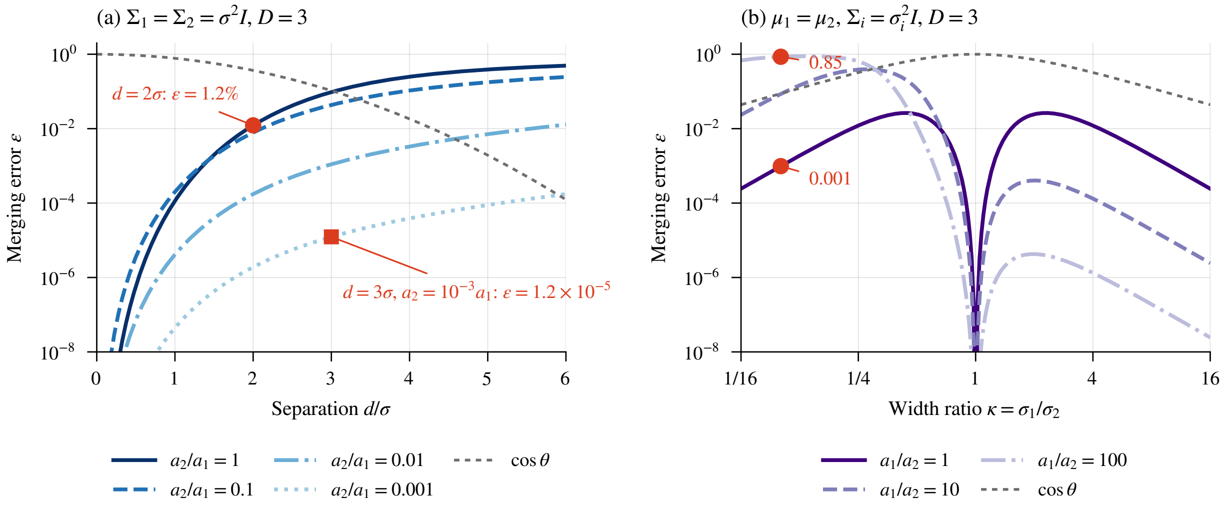


Figure 6: The merging error $\varepsilon$ of (20), evaluated from the kernel (4) with the moment-matched template of Proposition 3.1, in $D = 3$. (a) Equal isotropic covariances $\sigma^2 \boldsymbol{I}$ at separation $d$, for amplitude ratios $a_2/a_1$ from 1 to $10^{-3}$; the grey short-dashed curve is the proxy $\cos\theta = e^{-d^2/4\sigma^2}$ of (25). The two markers are the rows of Table 3: two equal splats at $d = 2\sigma$ merge with $\varepsilon = 1.2\%$, and a splat a thousand times fainter at $d = 3\sigma$ with $1.2 \times 10^{-5}$. The proxy is one curve for every amplitude ratio while $\varepsilon$ spans nearly four decades between them, which is the amplitude blindness of Proposition 5.1. (b) Co-located isotropic splats against the width ratio $\kappa = \sigma_1/\sigma_2$ for $a_1/a_2 \in \{1, 10, 100\}$; the grey short-dashed curve is $\cos\theta$ of (27). Every curve drops to zero at $\kappa = 1$ (proportional splats). The markers at $\kappa = 0.1$ are the values quoted in Remark 5.2: $\varepsilon = 0.001$ for equal amplitudes and 0.85 when the narrow splat is a hundred times brighter, at a single $\cos\theta = 0.09$.

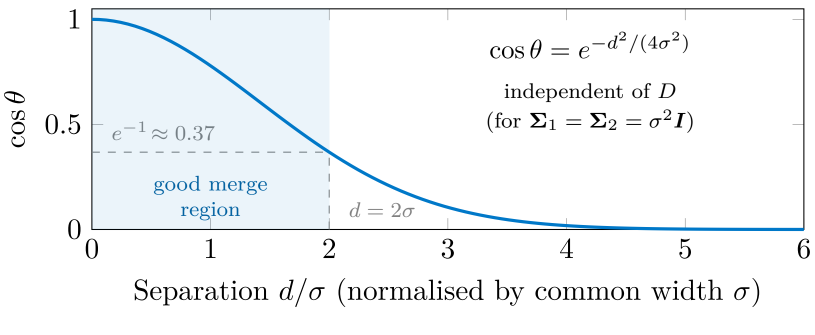


Figure 7: Inter-splat cosine similarity vs. normalised separation for two splats with equal isotropic covariances $\boldsymbol{\Sigma}_1 = \boldsymbol{\Sigma}_2 = \sigma^2 \boldsymbol{I}$. The result $\cos\theta = \exp(-d^2/4\sigma^2)$ is independent of the dimension $D$: the $2^{D/2}$ prefactor in (22) exactly cancels the determinant ratio for equal covariances. The shaded region ($d \lesssim 2\sigma$, $\cos\theta \gtrsim e^{-1}$) marks the rule of thumb of Section 6: for two equal splats the relative squared merging error there is about 1.2%.

##### 6.2 Co-located splats ($\boldsymbol{\mu}_1 = \boldsymbol{\mu}_2$)

When the centres coincide, $Q_{12} = 0$ and $\bar{\boldsymbol{\Sigma}}_{\text{inter}} = 0$, so:

$$\cos\theta = 2^{D/2} \frac{(|\boldsymbol{\Sigma}_1|\,|\boldsymbol{\Sigma}_2|)^{1/4}}{|\boldsymbol{\Sigma}_1 + \boldsymbol{\Sigma}_2|^{1/2}}. \tag{26}$$

This depends purely on the *shape* discrepancy. For isotropic splats with $\boldsymbol{\Sigma}_i = \sigma_i^2 \boldsymbol{I}$:

$$\cos\theta = \frac{2^{D/2}\,(\sigma_1\sigma_2)^{D/2}}{(\sigma_1^2 + \sigma_2^2)^{D/2}} = \left(\frac{2\sigma_1\sigma_2}{\sigma_1^2 + \sigma_2^2}\right)^{D/2} = \left(\frac{2}{\kappa + \kappa^{-1}}\right)^{D/2}, \tag{27}$$

where $\kappa = \sigma_1/\sigma_2$ is the width ratio (the letter $r$ is reserved for the residual of Fig. 5). This is the ratio of the geometric to the arithmetic mean of $\sigma_1^2$ and $\sigma_2^2$, raised to the $D/2$ power. It equals 1 when $\kappa = 1$ and decays rapidly as $\kappa$ departs from unity, especially in high dimensions (Fig. 8).

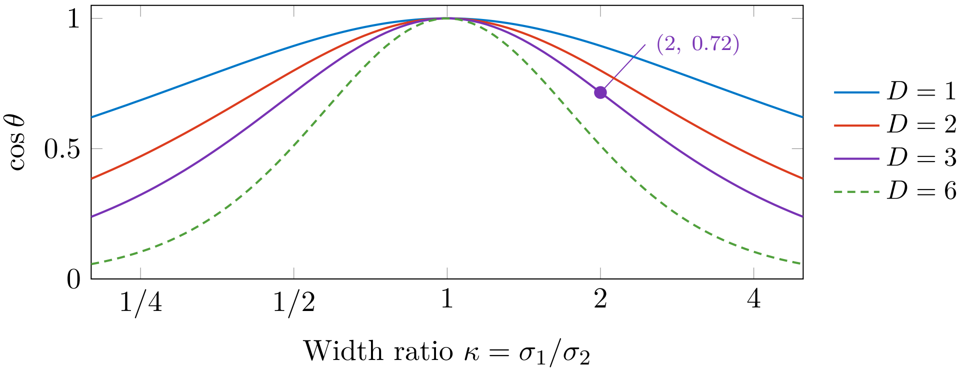


Figure 8: Inter-splat cosine for co-located isotropic splats as a function of the width ratio. Higher dimensions penalise shape differences more severely: in $D = 3$, a $2\times$ width ratio already drops the cosine to $\approx 0.72$.

##### 6.3 Proportional splats ($\phi_1 \propto \phi_2$)

When $\boldsymbol{\mu}_1 = \boldsymbol{\mu}_2$ and $\boldsymbol{\Sigma}_1 = \boldsymbol{\Sigma}_2$, the two splats are proportional: $\phi_1 + \phi_2 = (a_1 + a_2)\, G(\boldsymbol{x}; \boldsymbol{\mu}, \boldsymbol{\Sigma})$, which is already a single Gaussian. In this case $\cos\theta = 1$ and $\varepsilon = 0$: the merge is exact.

#### 7 Algorithm

Algorithm 1 computes the full mergeability score for a pair of splats. Algorithm 2 provides the cheaper cosine proxy for ranking. Algorithm 3 constructs the merged splat using either mass-preserving or $L^2$-optimal amplitude.

#### 8 Optimality discussion

##### 8.1 Optimality of the score

The mergeability score $\varepsilon$ uses the moment-matched $\bar{\boldsymbol{\mu}}, \bar{\boldsymbol{\Sigma}}$ with the $L^2$-optimised amplitude. The *globally* $L^2$-optimal Gaussian would also optimise $\bar{\boldsymbol{\mu}}$ and $\bar{\boldsymbol{\Sigma}}$. With the amplitude eliminated

**Algorithm 1:** Mergeability score for a pair of Gaussian splats

**Input:** Splats $(a_1, \boldsymbol{\mu}_1, \boldsymbol{L}_1)$ and $(a_2, \boldsymbol{\mu}_2, \boldsymbol{L}_2)$ in $\mathbb{R}^D$
**Output:** Mergeability score $\varepsilon \in [0, 1]$ (clamped at 0 against roundoff) and moment-matched merged parameters with $L^2$-optimal amplitude $(a^*, \bar{\boldsymbol{\mu}}, \bar{\boldsymbol{\Sigma}})$

`/* Step 1: Form covariances */`
$\boldsymbol{\Sigma}_i \leftarrow \boldsymbol{L}_i \boldsymbol{L}_i^\top$ for $i = 1, 2$;

`/* Step 2: Moment-matched parameters */`
$d_i \leftarrow \prod_k (\boldsymbol{L}_i)_{kk}$ ; `//` $|\boldsymbol{\Sigma}_i|^{1/2} = |\det \boldsymbol{L}_i|$
$m_i \leftarrow a_i \, (2\pi)^{D/2} \, d_i$ ; `// mass`
$w_i \leftarrow m_i \,/\, (m_1 + m_2)$;
$\bar{\boldsymbol{\mu}} \leftarrow w_1 \, \boldsymbol{\mu}_1 + w_2 \, \boldsymbol{\mu}_2$;
$\Delta\boldsymbol{\mu} \leftarrow \boldsymbol{\mu}_1 - \boldsymbol{\mu}_2$;
$\bar{\boldsymbol{\Sigma}} \leftarrow w_1 \, \boldsymbol{\Sigma}_1 + w_2 \, \boldsymbol{\Sigma}_2 + w_1 w_2 \, \Delta\boldsymbol{\mu} \, \Delta\boldsymbol{\mu}^\top$;
$\bar{\boldsymbol{L}} \leftarrow \mathrm{chol}(\bar{\boldsymbol{\Sigma}})$;
$d_g \leftarrow \prod_k \bar{\boldsymbol{L}}_{kk}$ ; `//` $|\bar{\boldsymbol{\Sigma}}|^{1/2}$

`/* Step 3: Self-energies */`
$J_i \leftarrow \pi^{D/2} \, d_i$ for $i = 1, 2$;
$J_g \leftarrow \pi^{D/2} \, d_g$;

`/* Step 4: Cross-kernels (input pair) */`
$\boldsymbol{S}_{12} \leftarrow \boldsymbol{\Sigma}_1 + \boldsymbol{\Sigma}_2$;
$\boldsymbol{R}_{12} \leftarrow \mathrm{chol}(\boldsymbol{S}_{12})$;
$\boldsymbol{v} \leftarrow$ solve $\boldsymbol{S}_{12} \, \boldsymbol{v} = \Delta\boldsymbol{\mu}$ ; `// via` $\boldsymbol{R}_{12}$
$Q_{12} \leftarrow \Delta\boldsymbol{\mu}^\top \boldsymbol{v}$;
$d_{12} \leftarrow \prod_k (\boldsymbol{R}_{12})_{kk}$;
$I_{12} \leftarrow (2\pi)^{D/2} \, (d_1 \, d_2) \,/\, d_{12} \;\cdot\; \exp(-Q_{12}/2)$;

`/* Step 5: Cross-kernels (splat vs. template) */`
**for** $i \in \{1, 2\}$ **do**
- $\boldsymbol{S}_i \leftarrow \boldsymbol{\Sigma}_i + \bar{\boldsymbol{\Sigma}}$;
- $\boldsymbol{R}_i \leftarrow \mathrm{chol}(\boldsymbol{S}_i)$;
- $\boldsymbol{z} \leftarrow$ solve $\boldsymbol{S}_i \, \boldsymbol{z} = \bar{\boldsymbol{\mu}} - \boldsymbol{\mu}_i$;
- $\bar{Q}_i \leftarrow (\bar{\boldsymbol{\mu}} - \boldsymbol{\mu}_i)^\top \boldsymbol{z}$;
- $I_i \leftarrow (2\pi)^{D/2} \, (d_i \, d_g) \,/\, \prod_k (\boldsymbol{R}_i)_{kk} \;\cdot\; \exp(-\bar{Q}_i/2)$;

`/* Step 6: Mergeability score */`
$\|f\|^2 \leftarrow a_1^2 \, J_1 + 2 \, a_1 a_2 \, I_{12} + a_2^2 \, J_2$;
$\langle f, \bar{G} \rangle \leftarrow a_1 \, I_1 + a_2 \, I_2$;
$\varepsilon \leftarrow \max\big(0,\; 1 - \langle f, \bar{G} \rangle^2 \,/\, (\|f\|^2 \cdot J_g)\big)$ ; `// clamp against roundoff`

`/*` Step 7: $L^2$`-optimal amplitude */`
$a^* \leftarrow \langle f, \bar{G} \rangle \,/\, J_g$;
**return** $\varepsilon$, $(a^*, \bar{\boldsymbol{\mu}}, \bar{\boldsymbol{\Sigma}})$;

through (18), the objective it minimises is the projection residual (19) seen as a function of the shape,

$$E^*(\bar{\boldsymbol{\mu}}, \bar{\boldsymbol{\Sigma}}) = \|f\|^2 - \frac{\langle f, \bar{G} \rangle^2}{\|\bar{G}\|^2}, \qquad \langle f, \bar{G} \rangle = \sum_i a_i \, \lambda_i, \quad \lambda_i = \langle G_i, \bar{G} \rangle, \tag{28}$$

**Algorithm 2:** Quick mergeability ranking via inter-splat cosine

**Input:** Splats $(a_1, \boldsymbol{\mu}_1, \boldsymbol{L}_1)$ and $(a_2, \boldsymbol{\mu}_2, \boldsymbol{L}_2)$
**Output:** Cosine similarity $\cos\theta \in [0, 1]$

$\boldsymbol{\Sigma}_i \leftarrow \boldsymbol{L}_i \boldsymbol{L}_i^\top$ for $i = 1, 2$;
$\boldsymbol{S} \leftarrow \boldsymbol{\Sigma}_1 + \boldsymbol{\Sigma}_2$;
$\boldsymbol{R} \leftarrow \mathrm{chol}(\boldsymbol{S})$;
$\Delta\boldsymbol{\mu} \leftarrow \boldsymbol{\mu}_1 - \boldsymbol{\mu}_2$;
$\boldsymbol{w} \leftarrow$ solve $\boldsymbol{S}\boldsymbol{w} = \Delta\boldsymbol{\mu}$;
$Q \leftarrow \Delta\boldsymbol{\mu}^\top \boldsymbol{w}$;
$\cos\theta \leftarrow 2^{D/2} \cdot \dfrac{\left(\prod_k (\boldsymbol{L}_1)_{kk}\right)^{1/2} \left(\prod_k (\boldsymbol{L}_2)_{kk}\right)^{1/2}}{\prod_k (\boldsymbol{R})_{kk}} \cdot e^{-Q/2}$;
**return** $\cos\theta$;

**Algorithm 3:** Construct the merged splat

**Input:** Splats $(a_i, \boldsymbol{\mu}_i, \boldsymbol{L}_i)$ for $i = 1, 2$; desired mode (`mass` or `L2`)
**Output:** Merged splat $(a_g, \bar{\boldsymbol{\mu}}, \bar{\boldsymbol{L}})$ with amplitude $a_g = \bar{a}$ (mass-preserving, (15)) or $a_g = a^*$ ($L^2$-optimal, (18))

Compute $\bar{\boldsymbol{\mu}}$, $\bar{\boldsymbol{\Sigma}}$, $\bar{\boldsymbol{L}} = \mathrm{chol}(\bar{\boldsymbol{\Sigma}})$ as in Algorithm 1 Steps 1–2;
**if** mode = `mass` **then**
  $m_i \leftarrow a_i\,(2\pi)^{D/2} \prod_k (\boldsymbol{L}_i)_{kk}$ for $i = 1, 2$;
  $a_g \leftarrow \bar{a} = (m_1 + m_2) \,/\, \left((2\pi)^{D/2} \prod_k \bar{\boldsymbol{L}}_{kk}\right)$ ; `// mass-preserving`
**else**
  Compute $I_1, I_2$ and $J_g$ as in Algorithm 1 Steps 3–5;
  $a_g \leftarrow a^* = (a_1 I_1 + a_2 I_2) \,/\, J_g$ ; `//` $L^2$`-optimal`
**return** $(a_g, \bar{\boldsymbol{\mu}}, \bar{\boldsymbol{L}})$;

where each $\lambda_i$ depends on $\bar{\boldsymbol{\mu}}$ and $\bar{\boldsymbol{\Sigma}}$ through the kernel (4). Differentiating (28) needs three further abbreviations: the summed covariances $\boldsymbol{S}_i = \boldsymbol{\Sigma}_i + \bar{\boldsymbol{\Sigma}}$ (as in Algorithm 1), the centre offsets $\boldsymbol{\delta}_i = \bar{\boldsymbol{\mu}} - \boldsymbol{\mu}_i$, and the normalised weights $\rho_i = a_i\lambda_i / \sum_j a_j\lambda_j$. A stationary shape then satisfies the mean condition

$$\sum_i a_i\,\lambda_i\,\boldsymbol{S}_i^{-1}\boldsymbol{\delta}_i = \boldsymbol{0} \tag{29}$$

together with the covariance condition

$$\sum_i \rho_i \left[\boldsymbol{S}_i^{-1} - \boldsymbol{S}_i^{-1}\boldsymbol{\delta}_i\boldsymbol{\delta}_i^\top \boldsymbol{S}_i^{-1}\right] = \tfrac{1}{2}\bar{\boldsymbol{\Sigma}}^{-1}. \tag{30}$$

The two conditions form a nonlinear system with no general closed-form solution. Thus:

- The score $\varepsilon$ is an **upper bound** on the true minimum relative $L^2$ error.
- For strongly overlapping pairs of similar covariance (the typical merge candidates) the moment-matched and $L^2$-optimal solutions are close; for well-separated pairs they are not. Take two Gaussians with $a_1 = a_2 = 1$ and $\boldsymbol{\Sigma}_1 = \boldsymbol{\Sigma}_2 = \boldsymbol{I}$ whose centres lie at $\pm 10$ along one axis, twenty standard deviations apart. Their moment-matched merge has $\varepsilon = 0.85$, while

keeping either component alone and discarding the other gives 0.5.

- For local refinement, Gauss–Newton iterations from the moment-matched solution can refine it to a stationary point; they do not certify a global minimum.
- The error formula (17) is exact for *any* choice of $g$, so one can always evaluate the true $L^2$ error once the optimal parameters are found numerically.

##### 8.2 Where moment matching is, and is not, $L^2$-optimal

1. **Proportional splats** ($\phi_1 \propto \phi_2$): the moment-matched template is the $L^2$-optimal Gaussian, and trivially so, since $\varepsilon = 0$ for both.
2. **Equal amplitudes and equal covariances** ($a_1 = a_2$, $\boldsymbol{\Sigma}_1 = \boldsymbol{\Sigma}_2$): the midpoint $(\boldsymbol{\mu}_1+\boldsymbol{\mu}_2)/2$ is a symmetry-enforced stationary point of (29) and the moment-matched mean, but not necessarily the global optimum (for well-separated pairs the optimum sits on one of the components). Even among midpoint-centred templates the optimal covariance differs from the moment-matched one: for unit Gaussians at $\pm h$ it is $h^2 + \sqrt{h^4+1}$ against the moment-matched $h^2 + 1$. Moment matching is therefore not $L^2$-optimal here, even in this symmetric case. Figure 9 draws both templates, and a single retained component, at $h = 0.5\sigma$, $\sigma$ and $2\sigma$.

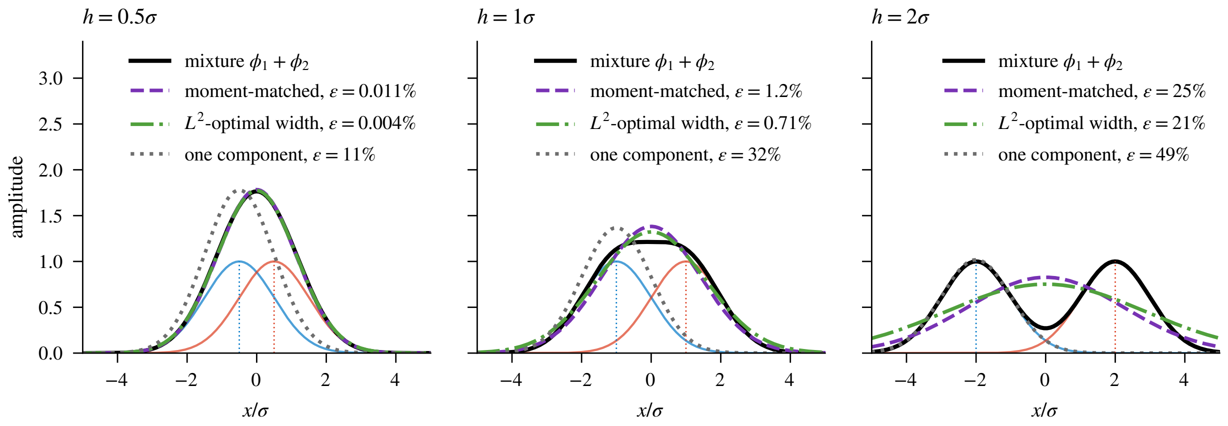


Figure 9: Three single-Gaussian templates for two unit splats at $\pm h$ ($\boldsymbol{\Sigma}_1 = \boldsymbol{\Sigma}_2 = \boldsymbol{I}$), one panel per separation, each with the $L^2$-optimal amplitude (18) for its shape and its score $\varepsilon$ of (20) in the legend. Dashed purple: the moment-matched template of Proposition 3.1, variance $h^2 + 1$. Dash-dotted green: the midpoint template with the $L^2$-optimal variance $h^2 + \sqrt{h^4+1}$ of case 2 above, wider than the moment-matched one at every separation and lower in error, by a factor of about 1.7 at $h = \sigma$ (0.71% against 1.2%). Dotted grey: one component kept as the shape, the alternative that wins for well-separated pairs; at these separations it is the worst of the three, and at $h = 2\sigma$ its 49% is close to the 0.5 of the twenty-sigma example above while the moment-matched score is still 25%.

Outside case 1 the moment-matched shape is at best a stationary point, so the score is a bound and not the minimum itself.

##### 8.3 Relationship to existing merging criteria

- **Runnalls' KL bound** [6]: an upper bound on the KL divergence cost of merging. It is $B_{12} = \frac{1}{2}\big[(w_1{+}w_2)\log|\bar{\boldsymbol{\Sigma}}| - w_1\log|\boldsymbol{\Sigma}_1| - w_2\log|\boldsymbol{\Sigma}_2|\big]$, with $w_i$ the components' probability masses and $\bar{\boldsymbol{\Sigma}}$ the moment-matched covariance (which contains the separation term), so the bound depends on amplitude ratios and separation alike; it bounds a KL cost, whereas our score is the exact fixed-template ISE.

- **ISE-based reduction**, the Williams–Maybeck criterion [10] and its descendants [4, 8]: uses $\|f-g\|^2$ directly, as we do. Our contribution is the explicit factorisation into the $\sin^2\!\alpha$ form and the amplitude-optimised closed-form expression.

- **Salmond's method** [7]: merges the pair with the smallest increase in within-component dispersion, $\mathrm{tr}\big(\boldsymbol{C}^{-1}\frac{w_i w_j}{w_i+w_j}(\boldsymbol{\mu}_i-\boldsymbol{\mu}_j)(\boldsymbol{\mu}_i-\boldsymbol{\mu}_j)^\top\big)$ with $\boldsymbol{C}$ the overall covariance of the mixture (a different matrix from the precision $\boldsymbol{P}$ of Section 2). This is a normalised separation criterion that vanishes for co-located pairs of unequal covariance; it is not the ISE increase, which is the Williams–Maybeck criterion above.

##### 8.4 Practical considerations

**Truncation.** The Luxar renderer uses a shifted truncated Gaussian $a\cdot s\cdot\max\!\big(0,\ e^{-Q/2}-c_T\big)$, where $Q=(\boldsymbol{x}-\boldsymbol{\mu})^\top\boldsymbol{\Sigma}^{-1}(\boldsymbol{x}-\boldsymbol{\mu})$ is the squared Mahalanobis distance (the quadratic form in the exponent of (1)), $s=1/(1{-}c_T)$ is a peak-preserving scale factor, and $c_T=e^{-T^2/2}\approx 0.023$ at the default truncation radius $T=2.75$ (shared by the Python and viewer constants). In 3D the mass beyond $T$ standard deviations is $\approx 5.6\%$ of the total and the squared-integral ($L^2$) energy beyond it $\approx 0.2\%$. The shift $c_T$ applied inside the support matters more than the tail: relative to the pure Gaussian, the shifted peak-preserving kernel carries 16.3% less mass and 7.4% less self-energy (a relative squared $L^2$ discrepancy of 0.8% per splat), so the pure-Gaussian analysis is a surrogate for the rendered quantity; its merge rankings have not been validated against the truncated renderer. Figure 10 draws both kernels and their difference.

**What the shipped merger computes.** Luxar's level builder, the `levels` *recipe* of Supplementary Document 8 (§8.2), uses the analysis of this document in one place and departs from it in another. Where it merges pairs greedily, which is its default for levels of at most 5,000 splats, it ranks candidate pairs by the unnormalised error $E^*$ of Algorithm 1. Above that size a warm start along a Morton curve (*Morton and Hilbert ordering*) followed by Lloyd reassignment replaces pairwise ranking altogether (Supplementary Document 8, §8.3).

The representative it stores, however, is not the template of Steps 2 and 7. By default the builder widens the inter-centre spread threefold, $\bar{\boldsymbol{\Sigma}}_{\text{out}}=\bar{\boldsymbol{\Sigma}}_{\text{intra}}+3\,\bar{\boldsymbol{\Sigma}}_{\text{inter}}$, so that neighbouring coarse splats sum to a flatter field; it then rescales the amplitude to preserve the mass the template had before the widening instead of re-optimising it, and finally applies a correction that pins the total mass of the whole level to that of its input. Only with the widening switched off and the level-wide correction disabled does the stored splat equal the template analysed here (the $L^2$-optimal amplitude is already the default).

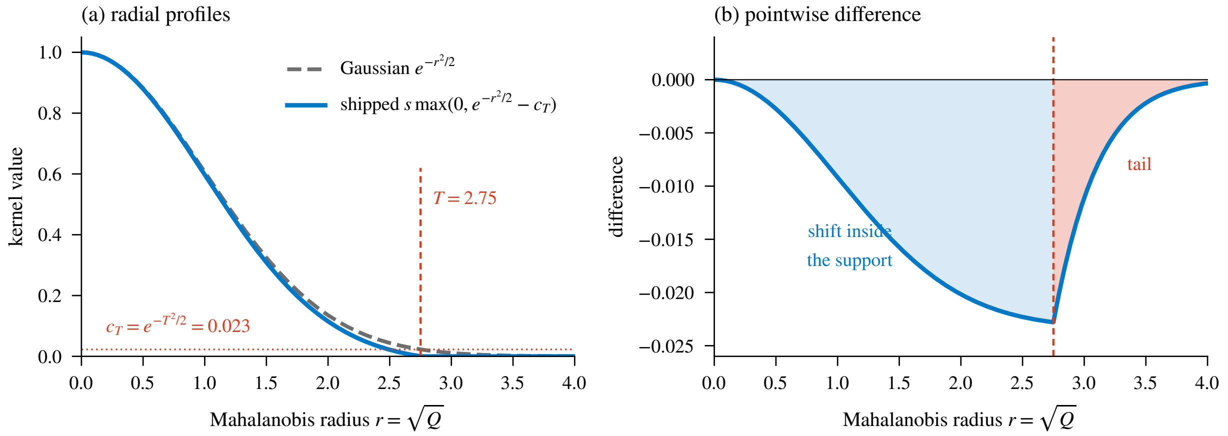


Figure 10: The kernel the renderer evaluates against the pure Gaussian of the analysis, as radial profiles in the Mahalanobis radius $r = \sqrt{Q}$. (a) The shipped kernel $s\ \max(0, e^{-r^2/2} - c_T)$ with $c_T = e^{-T^2/2} = 0.023$ at the default truncation radius $T = 2.75$ and $s = 1/(1 - c_T)$; it keeps the peak, sits below the Gaussian everywhere inside the support and is exactly zero beyond $T$. (b) Their pointwise difference. The shift inside the support (blue) is the larger of the two departures: in $D = 3$ the Gaussian mass beyond $T$ is 5.6% and its $L^2$ energy beyond $T$ is 0.2%, while the shipped kernel carries 16.3% less mass and 7.4% less self-energy than the Gaussian, a relative squared $L^2$ discrepancy $\|k - g\|^2/\|g\|^2$ of 0.8%.

Two consequences follow for how the numbers of this document should be read. First, the builder reports no per-pair $\varepsilon$: the unnormalised residual $E^*$ is used to rank, and neither it nor the template's $\varepsilon$ is written to the store. Second, the quality stamp the builder does write for each level is measured on the *stored* level, widened and mass-corrected, against the finest level of its group. The template's $\varepsilon$ derived here and that shipped stamp are therefore different quantities, and it is the stamp that describes what ships.

**Computational cost.** For $D = 3$: four Cholesky factorisations of $3 \times 3$ matrices ($\bar{\mathbf{\Sigma}}$, $\mathbf{\Sigma}_1 + \mathbf{\Sigma}_2$, $\mathbf{\Sigma}_1 + \bar{\mathbf{\Sigma}}$, $\mathbf{\Sigma}_2 + \bar{\mathbf{\Sigma}}$), three SPD solves and scalar arithmetic: a few hundred floating-point operations per pair, negligible compared to the rendering cost.

**Batch processing.** Before ranking candidate merges over a large splat set, precompute $J_i = \pi^{D/2}\,|\det \boldsymbol{L}_i|$ and $m_i$ for all splats. The per-pair cost then reduces to one kernel evaluation $I_{12}$ (for the cosine proxy) or four kernel evaluations (for the full score).

### 9 Summary

1. The $L^2$ integrated squared error is a divergence for which *all* terms in the Gaussian merging error have closed-form expressions.

2. The moment-matched Gaussian (KL-optimal) provides a closed-form approximation whose shape parameters are given by (13)–(14), with $L^2$-optimal amplitude from (18).

3. The normalised merging error is $\varepsilon = \sin^2\alpha$ (Eq. (20)), where $\alpha$ is the angle between the mixture and the template Gaussian in $L^2$.

4. For ranking pairs cheaply, the inter-splat cosine (Eq. (22)) requires only one kernel evaluation and is amplitude-independent.

5. For equal isotropic covariances, the cosine reduces to $\exp(-d^2/4\sigma^2)$, depending only on the normalised separation.

6. The score $\varepsilon$ is an upper bound on the relative $L^2$ error of the best single Gaussian, not that error itself. The moment-matched shape is $L^2$-optimal only for proportional splats (Section 8.2); in the symmetric equal-splat case its mean is a stationary point but its covariance is not optimal, and for well-separated pairs the bound is loose.

7. The representative the shipped level builder stores is not the template analysed here: by default it widens the inter-centre spread, rescales the amplitude to preserve mass and applies a level-wide mass correction (Section 8). The builder reports no per-pair $\varepsilon$, and the per-level quality stamp it writes is measured on the stored level against the finest level of its group, so the template's error derived here and the shipped stamp are different quantities, the latter measured on what ships.
