## Supplementary material for "Luxar: Gaussian splatting for microscopy and scalable interactive web visualisation of multidimensional scientific data": supp_doc_02_splat_count_vs_quality

### Splat count vs. reconstruction quality in Gaussian-splat fitting

Supplementary Document 2 – Luxar

**Abstract**

We ask how many Gaussian splats a microscopy volume needs and how to read the answer from noisy data alone. We sweep the splat count $N$ over nine doublings on seventeen benchmark volumes covering four modalities and a 27× range of voxel counts, and select $N$ by blind-spot cross-validation: a random 5% of voxels is hidden from the fit and the reconstruction is scored on them, so noise memorisation registers as a loss. An operating point $N^*$, a held-out peak or the onset of a plateau within 0.3 dB of the maximum, exists on 15/17 volumes, ten of them clear interior peaks. It spans more than an order of magnitude on the confocal and spinning-disk data ($N^* \approx$ 8K–128K), reaches 256K on the deconvolved zebrafish embryo, and follows signal density and noise level. On the two iSIM neuromast channels the post-fit cull holds the realised count nearly constant beyond the peak, so their small declines are not evidence of overfitting. On the Mouse heart and *Tribolium* light-sheet volumes the held-out score is still rising at 2M splats, consistent with a noise variance too small for the crossover into noise fitting to appear. On a synthetic volume with known clean signal the held-out maximum falls on the count where the true error is smallest. The criterion needs only the noisy data and a random mask, assuming zero-mean noise independent across voxels given the signal. A noise-floor estimate from median absolute deviations places each dataset in absolute terms beside its peak.

#### 1 Introduction

A Gaussian-splat fit has one free budget, the number of splats $N$. Too few and the reconstruction misses structure. Too many and, on data whose noise is not negligible, the splats begin to reproduce the noise of the particular acquisition. Choosing $N$ against the noisy source cannot tell the two apart: the training error keeps falling as $N$ grows whether the new splats capture signal or noise. A noise-floor ceiling does not settle it either. The ceiling says where agreement with the noisy source reaches the noise level, not how many splats the signal needs.

Two questions follow: how many splats a given volume needs, and how the answer can be read from the noisy data alone, without the clean volume a synthetic benchmark would supply. We answer both with *blind-spot cross-validation* [1]. A random 5% of the voxels is hidden from the fit and replaced by a neighbourhood median, and the fit is scored on those *held-out voxels* against their original values. A splat that captures signal improves the prediction at the hidden voxels; a splat that memorises noise cannot, because the noise there was never seen. The held-out score therefore peaks at the count where the signal is best recovered, and that peak, or the onset of a plateau, is the operating point $N^*$ we report. A noise-floor estimate places each volume's PSNR in absolute terms beside it.

The document sweeps $N$ over nine doublings on seventeen benchmark volumes, measures reconstruction PSNR, held-out PSNR and the noise floor, checks the criterion on a synthetic volume whose clean signal is known, and asks where the fits at $N^*$ go wrong. The detection rule and the noise-floor ensemble are the ones shipped in the `luxar gsplat cal` command, to which the Summary returns. §2 gives the protocol, §3 the results, §4 their interpretation, §5 the theory behind the criterion and §6 two diagnostics for sparse volumes.

#### Terms used in this document

**blind-spot cross-validation** A model-selection protocol in which a deterministic random 5% of voxels is hidden from the fit and the reconstruction is scored on those voxels afterwards, so that capacity spent on memorising noise shows up as a loss instead of a gain.

**held-out voxels** The masked voxels of a blind-spot split, whose original values the optimiser never sees and against which the held-out PSNR of each candidate splat count is scored.

**donut-median fill** The replacement of each masked voxel by the median of its $3^D - 1$ neighbours (the donut, 26 voxels in 3D); the shipped calibration draws the median over the unmasked neighbours only, and a document whose sweeps used all 26 neighbours, masked ones included, states that variant where it applies.

**PSNR** The peak signal-to-noise ratio, $10 \log_{10}(\text{peak}^2/\text{MSE})$ in decibels, where peak is the reference range of the values compared; unless a document states another range, the supplements take peak 1 on volumes normalised to $[0, 1]$, so that $\text{PSNR} = -10 \log_{10} \text{MSE}$.

**SSIM** The structural similarity index, the mean over local windows of a product of luminance, contrast and structure comparisons between two images, lying in $[-1, 1]$ with 1 meaning identical.

**cull (retention)** The removal of splats after a fit. The fit-time cull ranks splats by amplitude and keeps the top set accounting for a retention fraction of the total amplitude (0.95 by default); the contribution-based cull instead removes the splats whose contribution to the rendered field falls below an error budget or a fractional threshold, and is the baseline that removal-based level-of-detail methods use.

**relocation** The fixed-pool alternative to adding and removing splats during optimisation: the splat count is fixed at seeding, and the least informative splats are periodically moved to regions of high reconstruction residual, so the optimiser tensor shapes never change.

**plateau curve** A held-out PSNR curve of a splat-count sweep with a flat top and no clear interior peak, for which the smallest splat count within 0.3 dB of the maximum is reported as the operating point; the word is used in this sense only, not for a plateau of a training loss or a learning-rate schedule.

**signal-limited curve** A held-out PSNR curve whose maximum is the largest splat count tested and which is still climbing there (a total rise of at least 0.3 dB across the sweep, a trailing mean rise of at least 0.1 dB per step and a final step of at least 0.05 dB), so the peak lies beyond the sampled range.

***K**** The splat count recommended by a blind-spot sweep: the interior maximum of the held-out PSNR curve when there is a clear peak, the smallest count within 0.3 dB of the maximum on a plateau, and the largest count tested when the curve is still rising there (signal-limited). The shared operating-point table of these documents takes the knee instead, the smallest count within 0.3 dB of that maximum, which is the largest count itself on the two signal-limited benchmark volumes.

**$N^\dagger$** The population optimum of the splat count, the value minimising the expected held-out error over the distribution of masks, seeds and initialisations, as distinct from the operating point that one finite sweep detects.

**MAD** The median absolute deviation, a robust scale estimate that, divided by $0.6745 = 1/\Phi^{-1}(0.75)$, is a consistent estimator of the standard deviation of Gaussian noise.

**iSIM** Instant structured illumination microscopy, a super-resolution fluorescence technique that performs the structured-illumination reconstruction optically in a single exposure, giving roughly a twofold resolution gain at camera frame rates.

### 2 Methods

#### 2.1 Splat-count sweep

For each dataset we fit a single-pass Gaussian-splat model [5] at ten geometrically spaced splat counts $N \in \{1\text{K}, 2\text{K}, 4\text{K}, 8\text{K}, 16\text{K}, 32\text{K}, 64\text{K}, 128\text{K}, 256\text{K}, 512\text{K}\}$. Each fit uses identical hyperparameters: $L^1$ reconstruction loss, $n_{\text{iters}} = 20{,}000$, early-stop patience 500, learning rate 0.01, a conservative *cull (retention)* of 0.999 (post-fit removal of only the bottom 0.1% of cumulative splat amplitude), Adam optimiser with dynamic splat *relocation* enabled. Volumes are normalised to $[0, 1]$ before fitting by an affine map of each volume's intensity range with dataset-specific percentile clipping and, for the neuromast channels, background-floor subtraction, exactly as listed in the main Methods.

We record reconstruction *PSNR*, *SSIM* [6], MSE, fit time, and the realised splat count after culling. Throughout this document, on volumes normalised to $[0, 1]$ (peak 1),

$$\text{PSNR} = -10\log_{10}\text{MSE}, \qquad \text{MSE} = \frac{1}{|\Omega|}\sum_{j\in\Omega}(\hat{V}_j - Y_j)^2, \tag{1}$$

where $Y$ is the noisy volume over the voxel set $\Omega$ and $\hat{V}$ the rendered fit. The held-out PSNR of §2.2 restricts the sum to the masked voxels, and the noise-floor ceiling of §2.4 replaces the MSE by the noise variance. SSIM uses an $11^3$ Gaussian window of standard deviation 1.5 voxels with valid borders.

Across datasets the realised count stays within +2.5% and −16% of the requested $N$ on the confocal and spinning-disk data, with exceptions where redundant seeding in near-empty regions is culled post-fit. The count is fixed at seeding and relocation only moves splats: a seed pool within 5% of the request is accepted as it is, which is why 15 cells realise more splats than requested (by at most 2.5%), and the post-fit cull removes the rest. OpenCell-LMNB1 nuclear lamina loses 29% at 1K (and 3% at 512K), the sparse light-sheet volumes lose up to 29% (Mouse heart nuclei) and 49% (Zebrafish embryo nuclei), and the two background-subtracted *iSIM* neuromast channels lose 42–49% at $N^*$ and 85–93% at 512K. The tables below index cells by the requested $N$; the appendix curves plot the realised count on their x-axis, and the realised count is recorded per cell in the per-fit metrics tables deposited with the analysis.

The PSNR/SSIM of every rate-distortion cell is scored from the cell's cached store, written in the default encoding, as a viewer decodes it; relative to the optimiser's float32 parameters the encoding costs a median of 0.003 dB and at most 0.72 dB (Cells3D membranes at 512K). The held-out curves of §3.2 are scored on the fitter's parameters. Compression ratios are on-disk ratios: the bytes of the source volume at its stored dtype (`uint16` for most volumes, `uint8` for the Mouse heart, `float32` for the deconvolved neuromast channels) over the measured byte size of the cell's cached `.gsplats.zarr` store. The parameter-count ratio (voxels per float parameter) is recorded alongside it, and every ratio printed on a figure of this document is the on-disk one.

#### 2.2 Blind-spot cross-validation

To detect the splat count at which the model transitions from fitting signal to fitting noise we add a held-out evaluation following the blind-spot protocol of Batson and Royer [1]: 5% of voxels (seed 42) are masked and replaced by their 3×3×3 *donut-median fill* before fitting, so the model never sees the original values at masked positions. The donors define the donut: the centre is excluded and the donors are the 26 neighbours, other held-out voxels included (see below). Held-out PSNR is computed on the masked voxels against the *original* (pre-replacement) noisy intensities. The held-out fit re-uses the splat-count sweep above with otherwise identical hyperparameters. The theoretical justification for using this held-out PSNR as a model-selection criterion is given in §5. Fig. 1 draws the protocol.

**Which donors entered the fill.** The canonical sweeps of this document (and Table 1 of the main text) are scored with the donut median taken over all 26 neighbours, including other held-out voxels: 72–74% of held-out voxels have at least one such neighbour, and 1–60% of the fills change when those donors are excluded (Table 1). Sweeping four regime-spanning volumes again with the unmasked-donor fill changes held-out PSNR by at most 0.25 dB (mean −0.005 dB) and moves no optimum $N^*$ on those volumes (Table 2); the volumes are *C. elegans* embryo nuclei, Kidney nuclei, OpenCell-MAP4 microtubules and *Tribolium* embryo nuclei (to 512K of its 2M sweep). The leak the all-donor fill opens is this: a masked voxel's fill can carry the observed values of its held-out neighbours, which the model then sees during fitting, so at those voxels the score is not strictly blind. The share of held-out voxels with at least one held-out neighbour is what independent masking at 5% predicts, $1 - 0.95^{26} \approx 74\%$. The ablation covers four volumes to 512K; it is indicative for the other thirteen and for the 2M endpoints, not a proof that no ranking would move, and it does not include the volumes whose fills change most relative to their noise level. The unmasked-donor fill is the one Proposition 1 assumes and the released code implements; the tables of this document rest on the all-donor fill plus this ablation.

#### 2.3 Detection of $N^*$

We use a hybrid detection rule with three outcomes, plus one refinement applied afterwards from the realised splat counts. **(1) Peak**: if the held-out PSNR has a clear interior peak (the argmax is strictly interior and the mean of all pre-argmax values *and* the mean of all post-argmax values are both $\geq 0.1$ dB below the peak value), we report the argmax. **(2) Signal-limited**: if the argmax is the largest tested $N$, the curve rose by $\geq 0.3$ dB across the sweep, *and* it is still climbing at the top, the curve is a *signal-limited curve*. "Still climbing" means that the mean per-step rise over the trailing run is $\geq 0.1$ dB and the final step is $\geq 0.05$ dB, where the trailing run is the last four adjacent finite points of the sweep (three steps), or fewer when the sweep has gaps. The tail check prevents a flat-topped plateau whose noisy maximum merely happens to land on the last point from being mislabelled. **(3) Plateau**: otherwise the curve is a *plateau curve*, and we report the smallest $N$ within 0.3 dB of the maximum, the onset of diminishing returns.

The detector returns two counts. The *budget anchor* is the argmax for a peak, the plateau onset for a plateau, and the largest tested $N$ for a signal-limited curve; it is the count that `luxar`

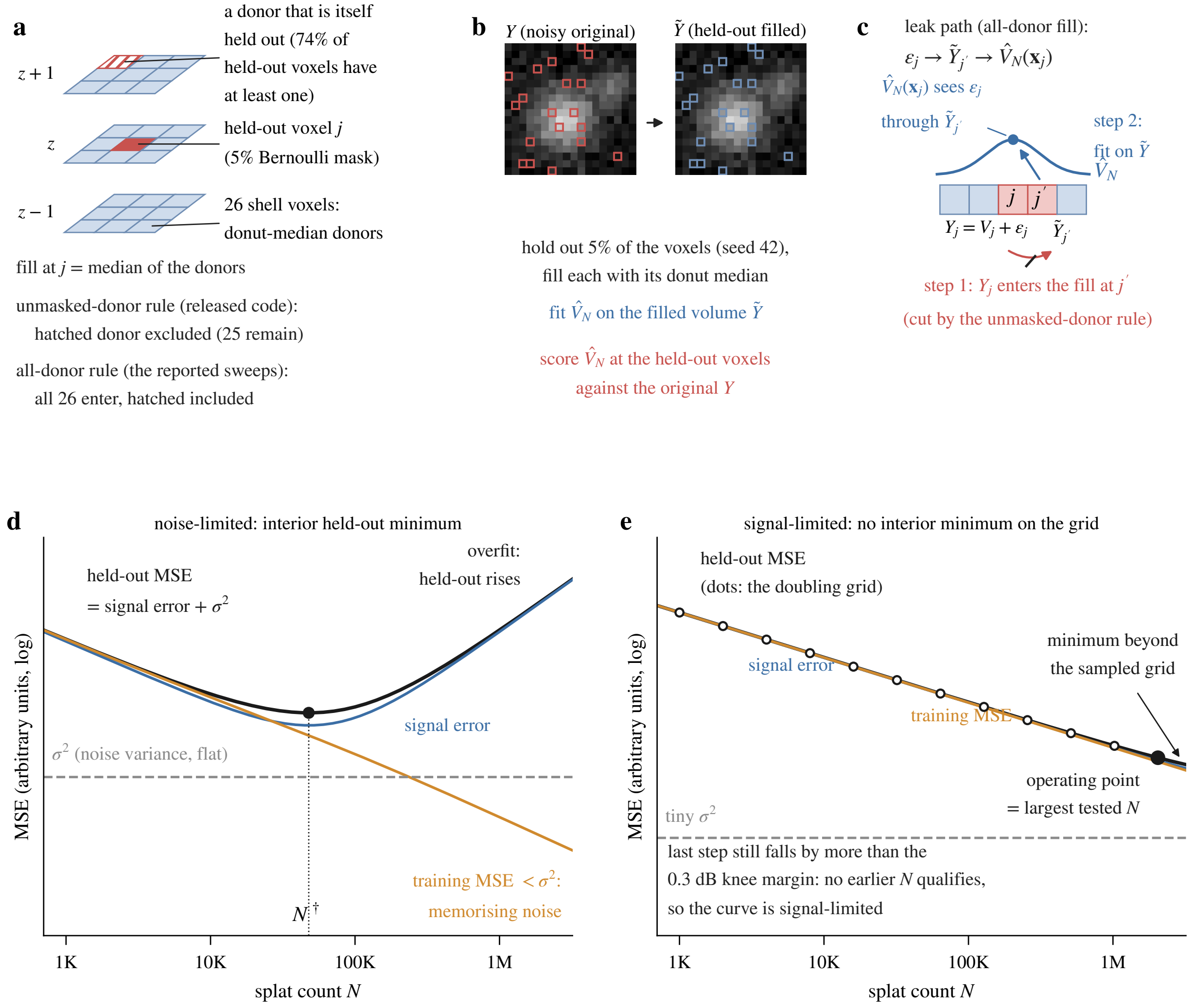


Figure 1: The blind-spot protocol. **(a)** A held-out voxel $j$ (5% Bernoulli mask) and its 26 shell voxels, whose median is the fill; one donor is itself held out, which the unmasked-donor rule of the released code excludes and the all-donor rule of the reported sweeps includes. **(b)** The shipped mask and fill on a patch: the fit sees the filled volume $\tilde{Y}$ and is scored at the held-out voxels against the original $Y$. **(c)** The leak of the all-donor fill: the observed value at $j$ enters the fill at a held-out neighbour $j'$, which the fit sees, so the prediction at $\boldsymbol{x}_j$ is no longer independent of $\epsilon_j$. **(d, e)** Idealised curves, not measured data: the held-out MSE is the signal error plus the flat noise variance $\sigma^2$ (Proposition 1), with an interior minimum when $\sigma^2$ is large (d) and none on the sampled grid when it is tiny (e), the signal-limited case in which the detector reports the endpoint.

`gsplat cal` prints as $K^*$. The *operating point* $N^*$ is the point of diminishing returns: the argmax for a clear peak, otherwise the smallest $N$ within 0.3 dB of the maximum. The two coincide except on a signal-limited curve, where $N^*$ may be a smaller count already within 0.3 dB of the top. Every $N^*$ in this document, and every "CV optimal" count in the main text, is the operating point. The rule is robust on all seventeen curves (§A). The two signal-limited light-sheet volumes (Mouse heart nuclei, *Tribolium* embryo nuclei) illustrate the branch: each sweep was extended to 2M splats and still showed no interior peak, and no smaller count lies within 0.3 dB of the 2M maximum, so anchor and operating point coincide at $N^* = $ 2M for both (Table 5, Type

Table 1: Held-out voxels with at least one held-out donut neighbour, share of fills that change under the unmasked-donor rule, mean absolute change of the fill (normalised intensity) and its RMS relative to the estimated noise level (– where every MAD estimator returns zero and no noise level is estimable).

| Volume | held-out voxels | with held-out neighbour (%) | fill changed (%) | mean $\lvert\Delta\rvert$ ($10^{-3}$) | RMS $\Delta/\hat{\sigma}$ |
|---|---|---|---|---|---|
| OpenCell-MAP4 nuclei | 916,483 | 72.9 | 59.8 | 1.38 | 0.15 |
| OpenCell-MAP4 microtubules | 916,483 | 72.9 | 59.2 | 2.03 | 0.18 |
| OpenCell-LMNB1 nuclei | 2,015,017 | 73.3 | 59.7 | 1.74 | 0.13 |
| OpenCell-LMNB1 nuclear lamina | 2,015,017 | 73.3 | 59.8 | 0.84 | 0.54 |
| Kidney nuclei | 209,624 | 71.6 | 57.0 | 1.76 | 0.56 |
| Kidney actin | 209,624 | 71.6 | 55.8 | 2.11 | 0.90 |
| Cells3D nuclei | 196,356 | 72.6 | 51.3 | 0.77 | 0.14 |
| Cells3D membranes | 196,356 | 72.6 | 58.9 | 0.32 | 0.49 |
| Blastocyst nuclear lamina | 877,917 | 73.3 | 22.8 | 0.29 | 0.85 |
| *C. elegans* embryo nuclei | 537,332 | 72.8 | 57.1 | 0.41 | 0.09 |
| Fly brain neurons | 4,969,182 | 73.4 | 26.2 | 0.16 | 0.22 |
| *Tribolium* embryo nuclei | 4,972,548 | 73.3 | 46.7 | 0.14 | 0.64 |
| Mouse heart nuclei | 4,985,912 | 73.4 | 14.3 | 0.51 | – |
| Zebrafish embryo nuclei | 5,334,977 | 73.5 | 13.0 | 0.41 | – |
| *Drosophila* embryo nuclei | 3,884,295 | 73.3 | 23.2 | 0.89 | 1.27 |
| Neuromast membranes | 1,401,763 | 73.2 | 1.2 | 0.14 | – |
| Neuromast nuclei | 1,401,763 | 73.2 | 0.7 | 0.04 | – |

Table 2: Fill ablation: held-out PSNR (dB) of the canonical sweep (all 26 donors) and of the same sweep with the unmasked-donor fill (other held-out voxels excluded from the median), per splat count $N$ and volume; $\Delta$ is the change.

| | *C. elegans* embryo nuclei | | | Kidney nuclei | | | OpenCell-MAP4 microtubules | | | *Tribolium* embryo nuclei | | |
|---|---|---|---|---|---|---|---|---|---|---|---|---|
| $N$ | all | unmasked | $\Delta$ | all | unmasked | $\Delta$ | all | unmasked | $\Delta$ | all | unmasked | $\Delta$ |
| 1K | 38.27 | 38.26 | −0.01 | 22.30 | 22.31 | +0.01 | 21.39 | 21.33 | −0.06 | 29.74 | 29.78 | +0.04 |
| 2K | 38.85 | 38.88 | +0.02 | 23.97 | 24.05 | +0.08 | 22.33 | 22.35 | +0.02 | 31.19 | 31.25 | +0.06 |
| 4K | 39.15 | 39.14 | −0.01 | 26.36 | 26.45 | +0.09 | 23.31 | 23.27 | −0.04 | 37.91 | 38.01 | +0.10 |
| 8K | 39.23 | 39.21 | −0.01 | 29.20 | 29.20 | 0.00 | 24.11 | 24.08 | −0.03 | 42.36 | 42.61 | +0.25 |
| 16K | 39.22 | 39.21 | −0.01 | 31.35 | 31.38 | +0.03 | 24.52 | 24.50 | −0.02 | 47.43 | 47.43 | 0.00 |
| 32K | 39.21 | 39.19 | −0.02 | 32.84 | 32.82 | −0.02 | 24.80 | 24.78 | −0.02 | 51.25 | 51.40 | +0.15 |
| 64K | 39.19 | 39.15 | −0.03 | 33.91 | 33.83 | −0.08 | 24.89 | 24.85 | −0.04 | 52.53 | 52.45 | −0.08 |
| 128K | 39.13 | 39.09 | −0.04 | 33.25 | 33.31 | +0.05 | 24.82 | 24.77 | −0.05 | 53.38 | 53.41 | +0.03 |
| 256K | 38.96 | 38.91 | −0.06 | 32.19 | 32.01 | −0.18 | 24.68 | 24.63 | −0.05 | 53.84 | 53.74 | −0.10 |
| 512K | 38.88 | 38.85 | −0.04 | 30.67 | 30.61 | −0.06 | 24.56 | 24.52 | −0.04 | 54.59 | 54.53 | −0.05 |

column "Signal-limited"). This three-way classification is exactly the rule implemented by the shipped `luxar gsplat cal` command (`find_k_star` in `luxar.gsplats.calibration`), which the analysis calls directly. Fig. 2 draws the rule as a decision tree.

**(4) Cull-limited** is a refinement applied on top of it from the realised (post-cull) counts, which the shipped command does not carry. When a peak or plateau is followed by requested counts at least 4× larger whose realised counts grow by less than 1.5×, the sweep beyond the operating point no longer varies capacity, the label becomes cull-limited, and $N^*$ is unchanged. The two such volumes are the iSIM neuromast channels (Neuromast membranes, Neuromast nuclei).

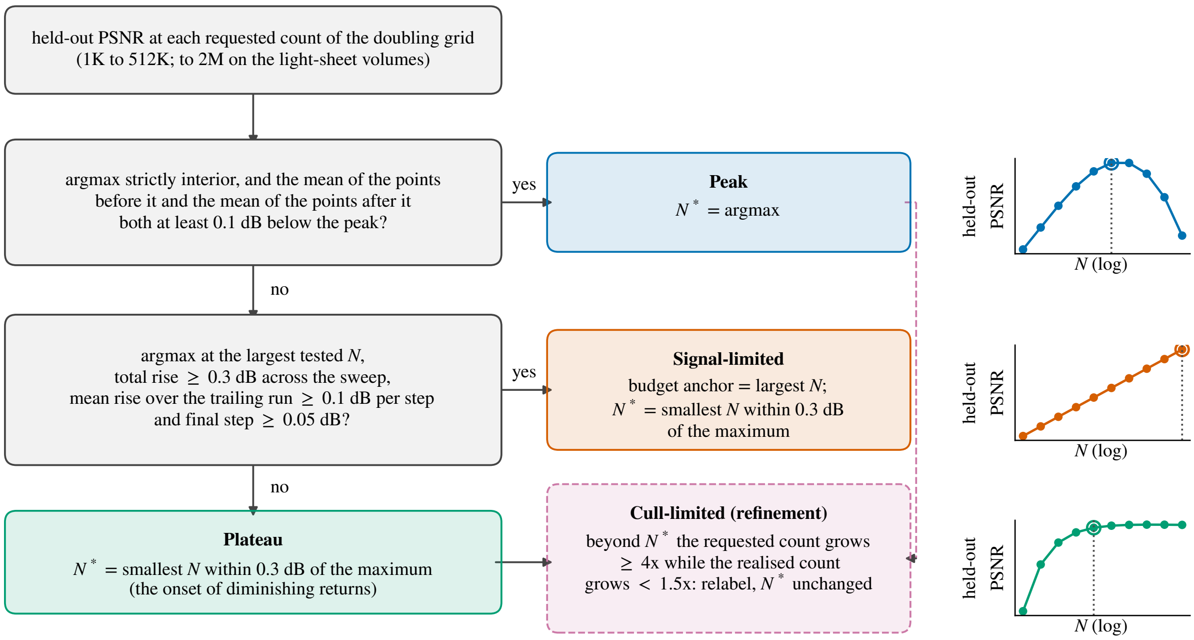


Figure 2: The detection rule as a decision tree. The two questions on the left are asked of the held-out PSNR curve in turn; each outcome on the right names the operating point it reports, and the dashed box is the cull-limited refinement applied afterwards from the realised counts. The constants are those of the shipped detector and of this document: 0.1 dB flank margin, 0.3 dB total rise and knee tolerance, 0.1 dB per step over the trailing run and 0.05 dB for the final step, and a realised count growing by less than 1.5× while the requested count grows by 4× or more. The three small curves are idealised sketches of each outcome, not measured data; the open circle and the dotted line mark the reported $N^*$.

#### 2.4 Noise-floor estimation

We estimate the per-dataset noise standard deviation $\hat{\sigma}$ using an ensemble of three robust estimators and set the PSNR ceiling for $[0, 1]$-normalised data to $\mathrm{PSNR}_{\max} = -20 \log_{10} \hat{\sigma}$. The ceiling is an upper bound: it is the PSNR a reconstruction equal to the clean signal would score against the noisy source if the noise were pixel-independent with this standard deviation everywhere. Robust estimators over a mostly dark volume measure the background noise level; under signal-dependent (Poisson-like) noise the level on the structures is higher and the true ceiling lower. A lower true ceiling leaves a smaller true gap, so a gap "below the ceiling" computed from the background estimate is an upper bound on the headroom, not a measure of it.

Fig. 3 sketches what each estimator filters.

**Laplacian MAD (adapted from Immerkær [4]).** Immerkær's estimator averages the absolute response of one 3×3 Laplacian-of-differences kernel. Here we take the median absolute deviation ($MAD$) of the $d$-dimensional discrete Laplacian $L = \nabla^2 Y$ of the noisy volume $Y$ instead, so that structure contributes to the tail and not to the scale:

$$\hat{\sigma}_{\mathrm{Lap}} \;=\; \mathrm{MAD}(L) \,/\, (0.6745 \sqrt{c_L}), \qquad c_L = 2d(2d{+}1),$$

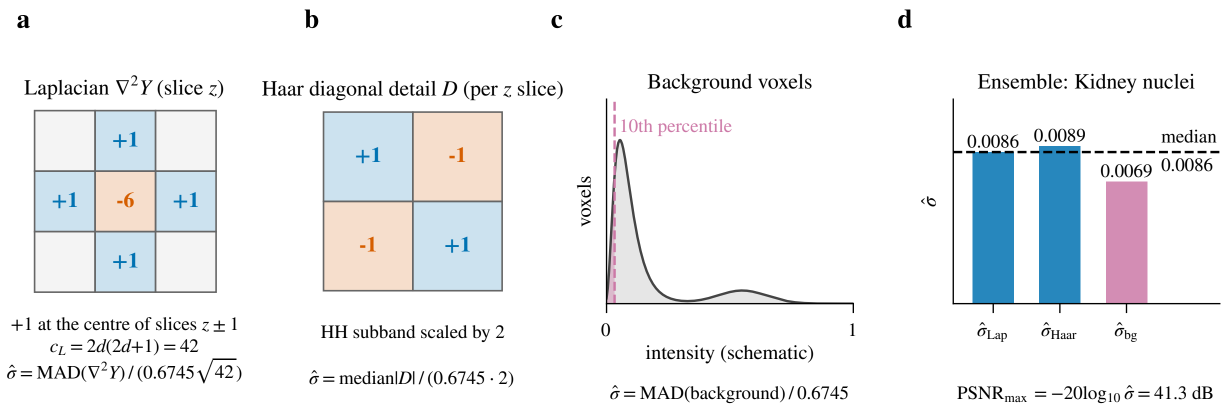


Figure 3: The three noise-floor estimators (panels a to c are schematic, not data). **(a)** Laplacian MAD: the 3D discrete Laplacian, centre weight $-6$ and $+1$ on the six face neighbours (the two out-of-slice ones sit at the centre of the slices above and below), whose squared weights sum to $c_L = 42$; it removes smooth signal and passes voxel-independent noise. **(b)** Haar MAD: the $2\times2$ diagonal-difference kernel applied within each $z$ slice, the HH subband scaled by 2. **(c)** Background MAD: the voxels below the 10th intensity percentile of a schematic histogram. **(d)** The ensemble is the median of the three, shown with the measured values for Kidney nuclei (Table 7), and the ceiling follows as $-20\log_{10}\hat{\sigma}$.

where $c_L$ is the sum of squared Laplacian kernel weights and $d$ is the spatial dimension.

**Haar finite-difference MAD [3].** For each $z$-slice compute the diagonal detail $D_{y,x} = Y_{y,x} - Y_{y,x+1} - Y_{y+1,x} + Y_{y+1,x+1}$ (algebraically the Haar HH subband, scaled by 2); then $\hat{\sigma}_{\text{Haar}} = \text{median}(|D|)/(0.6745 \cdot 2)$.

**Background-region MAD.** Restrict to voxels below the 10-th intensity percentile, then $\hat{\sigma}_{\text{bg}} = \text{MAD(background)}/0.6745$.

The ensemble estimate is the median $\hat{\sigma} = \text{median}(\hat{\sigma}_{\text{Lap}}, \hat{\sigma}_{\text{Haar}}, \hat{\sigma}_{\text{bg}})$; we discuss the rationale for the median in §5.3.[1]

#### 2.5 Effective resolution and splat scale

To relate the fitted splat sizes to what the data can resolve, we estimate each volume's effective resolution from the data itself with decorrelation analysis [2]: for a 2D slice with spectrum $F(\mathbf{k})$ and phase-only spectrum $F_n = F/|F|$, the decorrelation curve

$$d(r) = \Re \sum_{|\mathbf{k}|\le r} F\,\overline{F_n} \Big/ \sqrt{\sum |F|^2 \sum_{|\mathbf{k}|\le r} |F_n|^2}$$

peaks where the signal spectrum runs out of correlated content. The cutoff $k_c$ is the largest peak position over a family of Gaussian high-pass pre-filters, and the resolution is $2\,\Delta x/k_c$ ($k_c$ in units of the Nyquist frequency). Lateral resolution is the median over seven $xy$ slices, chosen

[1]The shipped implementation takes the median over whichever of these three estimators return a finite value for a given volume (an estimator is dropped only when it returns none, which for the background estimator requires an empty dark tail); all values in Table 7 use the three estimators above.

by total intensity among planes in the central half of the stack. Axial resolution uses $xz$ and $yz$ slices in physical frequency units ($k_z$ scaled by $\Delta z/\Delta x$) restricted to a $\pm 30°$ sector about the $z$ axis, so anisotropic data need no resampling, and stacks of fewer than 32 planes get no axial estimate. On synthetic point sources blurred with a Gaussian PSF and Gaussian noise (SNR 5–20), the estimator returns 1.06–1.26× the true FWHM (2.5–3.0 $s_{\mathrm{PSF}}$, with $s_{\mathrm{PSF}}$ the standard deviation of the PSF), coarser at low SNR, and the sector estimate recovers the axial FWHM of an anisotropically sampled volume within 14%. The per-slice interquartile range and the sensitivity to the peak rule (global vs. first local maximum) are reported with every estimate; on dim volumes they reach a factor 1.4–3, so these are effective, noise-limited resolutions with that uncertainty, not optical specifications.

Splat scale is read from the cached fit at each volume's operating point $N^*$. For every splat with covariance matrix $\Sigma$ we take the axis-aligned marginal width $\ell_i = \sqrt{\Sigma_{ii}}$ along axis $i$, in voxel units, and summarise it by its 10th percentile $\ell_{10}$ (the small end of the mixture) and its amplitude-weighted median $\ell^a_{50}$ (the scale that carries the intensity). The fitter's diagonal floor is $\sqrt{1/12} = 0.29$ voxels (the standard deviation of a uniform voxel), and the dynamic relocation step re-seeds splats at an isotropic $\ell = 0.5$ voxels, so both values are checked for in the distributions.

##### 2.6 Residual features

A held-out PSNR is one number per volume; it cannot say *which* structures a fit at $N^*$ gets wrong. To find them without choosing them by eye, we render the cached fit at the operating point exactly as a viewer decodes it, subtract the normalised source volume, and smooth the signed residual with a Gaussian of one lateral voxel (axial width scaled by the voxel anisotropy). The smoothed residual is thresholded at six robust standard deviations (median absolute deviation over the active voxels, where source or reconstruction is non-zero), separately for negative values (reconstruction dimmer than the source) and positive ones (brighter), and each side is grouped into 26-connected components. A component's extent score is the sum of $|r|$ over its voxels, so large *and* strong errors rank first; the three strongest per volume, at least three-quarters of a crop apart, are the "error features" of Fig. 5 and Table 9. The threshold is relative to each volume's own residual noise, so the flagged fraction of foreground voxels compares the volumes on one footing; the crops show the $xy$ slice through each component's strongest smoothed residual, with source and reconstruction on a shared display window.

##### 2.7 Datasets

Table 3 lists the 17 benchmark volumes (17 volumes from 12 datasets; a dataset is one physical acquisition, and co-registered channels of one acquisition share it). The seventeen volumes swept in this document span four microscopy modalities, at least six organisms (the Cells3D source names none), and a $\sim 27\times$ range of voxel counts. All are 3D float32 volumes normalised to $[0, 1]$ before fitting.

Table 3: The benchmark volumes (17 volumes from 12 datasets), grouped by modality. Channels of one acquisition (the two OpenCell-MAP4 channels, the two OpenCell-LMNB1 channels, the two kidney stains, the two Cells3D channels, the two neuromast channels) are co-registered: the same volume, two stains. Shape is $z \times y \times x$ in voxels.

| Volume | Modality | Shape | Voxels |
|---|---|---|---|
| OpenCell-MAP4 nuclei | Spinning-disk | 51×600×600 | 18.4 M |
| OpenCell-MAP4 microtubules | Spinning-disk | 51×600×600 | 18.4 M |
| OpenCell-LMNB1 nuclei | Spinning-disk | 112×600×600 | 40.3 M |
| OpenCell-LMNB1 nuclear lamina | Spinning-disk | 112×600×600 | 40.3 M |
| Kidney nuclei | Confocal | 16×512×512 | 4.2 M |
| Kidney actin | Confocal | 16×512×512 | 4.2 M |
| Cells3D nuclei | Confocal | 60×256×256 | 3.9 M |
| Cells3D membranes | Confocal | 60×256×256 | 3.9 M |
| Blastocyst nuclear lamina | Confocal | 236×275×271 | 17.6 M |
| *C. elegans* embryo nuclei | Confocal | 41×512×512 | 10.7 M |
| Fly brain neurons | Confocal | 435×478×478 | 99.4 M |
| *Tribolium* embryo nuclei | Light-sheet (deconv.) | 991×104×965 | 99.5 M |
| Mouse heart nuclei | Light-sheet | 597×174×960 | 99.7 M |
| Zebrafish embryo nuclei | Light-sheet (deconv.) | 407×512×512 | 106.7 M |
| *Drosophila* embryo nuclei | Light-sheet | 108×1352×532 | 77.7 M |
| Neuromast membranes | iSIM (deconv.) | 84×580×576 | 28.1 M |
| Neuromast nuclei | iSIM (deconv.) | 84×580×576 | 28.1 M |

##### 2.8 Compute

The canonical sweep, 17 volumes × 10 splat counts × 2 modes (rate-distortion + held-out blind-spot) = 340 fits plus the 1M and 2M extension cells on the light-sheet volumes swept further, ran on a single NVIDIA RTX PRO 6000 Blackwell (96 GB VRAM, sm_120). With early stopping (patience 500) most fits terminate well before the 20,000-iteration cap: in rate-distortion mode the median run is 2 500 iterations and 36 of 178 cells (the largest counts; 30 of them on the light-sheet and iSIM volumes, the other 6 on confocal ones) reach the cap; in held-out mode the median is 2 575 and again 36 of 178 reach it. No fit reached the configured `max_abs_error` threshold, so no rate-distortion cell is marked converged; termination is by loss patience or, for the 72 capped cells, by the iteration budget (§3.3 for the seed spread). The two further seeds of §3.3 add 400 fits and the cap-lifted heart run of Table 10 six more (three per mode). Per-fit wall-clock is not reported here (the figure-time panel in the per-dataset quality-curve figures plots fit time vs. $N$; the headline metrics tables in this document focus on quality). Seed deduplication contributes several minutes per fit at 256K and 512K seeds and dominates total wall-clock at the high-$N$ end of the sweep; it is an implementation cost of the seeding step, not a cost of the fit itself.

**Reproducibility.** The per-fit metrics tables of both sweep modes (one row per volume and splat count, with the realised count, the scoring domain and the iteration count of every cell), the seed-replicate and cap-lifted tables, the noise-floor table, the resolution and splat-scale table, the fill-ablation statistics and the residual-feature detections are deposited with the analysis; the detection rule of §2.3 and the noise-floor ensemble are the ones shipped in `luxar gsplat cal`.

Table 4: Reconstruction PSNR (dB) at five representative splat counts. Gain = PSNR(512K) − PSNR(1K); $PSNR_{max}$ is the noise-floor ceiling (Table 7). One fit per cell; on the ten confocal and spinning-disk datasets two further seeds differ from each other by at most 0.22 dB, while the canonical seed, scored from its decoded store, sits up to 0.85 dB from their mean (§3.3).

| Volume | 1K | 8K | 32K | 128K | 512K | Gain | $PSNR_{max}$ |
|---|---|---|---|---|---|---|---|
| OpenCell-MAP4 nuclei | 25.09 | 28.04 | 28.61 | 29.32 | 30.59 | +5.5 | 32.6 |
| OpenCell-MAP4 microtubules | 21.43 | 24.20 | 25.21 | 26.21 | 27.95 | +6.5 | 30.4 |
| OpenCell-LMNB1 nuclei | 22.81 | 25.28 | 25.59 | 25.90 | 26.84 | +4.0 | 30.0 |
| OpenCell-LMNB1 nuclear lamina | 21.45 | 31.83 | 34.14 | 35.05 | 36.46 | +15.0 | 42.6 |
| Kidney nuclei | 22.39 | 29.66 | 34.82 | 40.59 | 45.22 | +22.8 | 41.3 |
| Kidney actin | 20.98 | 26.08 | 31.08 | 39.14 | 48.07 | +27.1 | 43.5 |
| Cells3D nuclei | 31.41 | 35.34 | 36.78 | 38.21 | 44.79 | +13.4 | 37.6 |
| Cells3D membranes | 37.10 | 45.11 | 49.93 | 53.49 | 58.09 | +21.0 | 53.5 |
| Blastocyst nuclear lamina | 32.65 | 37.43 | 39.20 | 40.48 | 41.85 | +9.2 | 54.4 |
| *C. elegans* embryo nuclei | 38.32 | 39.27 | 39.33 | 39.55 | 40.48 | +2.2 | 40.7 |
| Fly brain neurons | 38.09 | 42.92 | 44.58 | 45.09 | 45.38 | +7.3 | 52.1 |
| *Tribolium* embryo nuclei | 29.67 | 43.20 | 51.38 | 53.52 | 54.93 | +25.3 | 61.0 |
| Mouse heart nuclei | 26.34 | 29.75 | 32.12 | 36.34 | 43.43 | +17.1 | – |
| Zebrafish embryo nuclei | 22.61 | 26.90 | 32.56 | 38.29 | 40.78 | +18.2 | – |
| *Drosophila* embryo nuclei | 21.07 | 25.96 | 30.52 | 33.92 | 35.37 | +14.3 | 51.1 |
| Neuromast membranes | 33.21 | 41.23 | 52.69 | 57.18 | 57.54 | +24.3 | – |
| Neuromast nuclei | 34.90 | 49.02 | 66.34 | 66.77 | 68.02 | +33.1 | – |

### 3 Results

#### 3.1 Rate-distortion: PSNR vs. splat count

Table 4 reports reconstruction PSNR (dB) at five representative splat counts (1, 8, 32, 128, and 512 thousand); the full ten-point sweep is in the per-dataset rate-distortion figures (§A). Total dynamic range over the sweep (Gain column) varies more than 15× across volumes: from +2.2 dB on *C. elegans* embryo nuclei (sparse signal already near the noise floor by 1K splats) to +25.3 dB on *Tribolium* embryo nuclei and +33.1 dB on Neuromast nuclei (dense nuclei and filaments with high representational demand, and, on the confocal volumes, ample noise to memorise).

The PSNR-vs-$N$ profile is qualitatively distinct across regimes. The light-sheet *Tribolium* embryo climbs steeply to 51.4 dB by 32K and then slowly (+3.5 dB over the next 16× in capacity, to 54.9 dB at 512K and 56.0 dB at 2M), still 5 dB short of its 61 dB noise ceiling. The sparse *C. elegans* embryo reaches 38.3 dB by 1K and barely moves through 512K (Gain +2.2 dB); the volume is mostly empty and the signal is captured early. Dense confocal tissue (Kidney nuclei, Kidney actin, Cells3D membranes) improves continuously through 512K without a knee, and four confocal volumes end *above* their estimated noise ceilings at 512K (Kidney nuclei +3.9, Kidney actin +4.6, Cells3D membranes +4.6, Cells3D nuclei +7.2 dB). A reconstruction PSNR above the noise floor is a direct indicator of noise memorisation: the model is reproducing the noisy voxels more faithfully than the noise-free signal could be, which the held-out criterion below confirms on exactly these datasets.

Table 5: Held-out PSNR (dB) summary: peak value, the operating point $N^*$ (the diminishing-returns knee: the interior argmax for a clear peak, else the smallest $N$ within 0.3 dB of the maximum), the drop from the peak to the largest tested $N$, and the train/held-out PSNR gap at 512K. Volumes are grouped by modality. Type is "Peak" when the held-out maximum is interior, stands at least 0.1 dB above the mean of the points on either side of it and the realised count keeps growing beyond it; "Signal-limited" when the maximum is the largest tested $N$ and the curve is still climbing there (§2.3); "Cull-limited" when the detector reads a peak or plateau but the realised count saturates beyond the operating point (a growth of less than 1.5× over at least 4× more requested splats), so that the drop is between fits of nearly the same size (it overrides "Peak"); and "Plateau" otherwise (the four light-sheet volumes were swept to 2M splats; all others to 512K). One fit per cell; replicate seeds move the held-out values by at most 0.41 dB and the operating point by at most one grid step (§3.3).

| Volume | Held-out peak (dB) | $N^*$ | Drop at largest $N$ (dB) | Train/held-out gap at 512K (dB) | Type |
|---|---|---|---|---|---|
| OpenCell-MAP4 nuclei | 28.15 | 16K | −0.39 | 2.86 | Peak |
| OpenCell-MAP4 microtubules | 24.89 | 64K | −0.33 | 3.44 | Peak |
| OpenCell-LMNB1 nuclei | 25.45 | 32K | −0.36 | 1.76 | Peak |
| OpenCell-LMNB1 nuclear lamina | 34.00 | 64K | −0.38 | 2.84 | Peak |
| Kidney nuclei | 33.91 | 64K | −3.24 | 14.59 | Peak |
| Kidney actin | 30.65 | 128K | −2.92 | 20.00 | Peak |
| Cells3D nuclei | 35.45 | 32K | −0.50 | 9.92 | Peak |
| Cells3D membranes | 46.73 | 32K | −3.46 | 15.73 | Peak |
| Blastocyst nuclear lamina | 37.58 | 32K | −0.71 | 5.01 | Peak |
| *C. elegans* embryo nuclei | 39.23 | 8K | −0.34 | 1.45 | Peak |
| Fly brain neurons | 44.51 | 64K | 0.00 | 0.86 | Plateau |
| *Tribolium* embryo nuclei | 55.55 | 2M | – | 0.56 | Signal-limited |
| Mouse heart nuclei | 45.30 | 2M | – | 0.64 | Signal-limited |
| Zebrafish embryo nuclei | 37.99 | 256K | −0.06 | 2.95 | Plateau |
| *Drosophila* embryo nuclei | 34.27 | 256K | 0.00 | 1.14 | Plateau |
| Neuromast membranes | 50.85 | 128K | −0.19 | 7.09 | Cull-limited |
| Neuromast nuclei | 63.60 | 64K | −0.03 | 4.43 | Cull-limited |

##### 3.2 Held-out PSNR (blind-spot cross-validation)

A finite operating point appears on fifteen of the seventeen volumes (Table 5). Of these, ten show a clear interior peak at $N^* = 8\text{K}$–$128\text{K}$: all four spinning-disk volumes and six of the seven confocal volumes, with a peak-to-largest-$N$ drop of 0.33–3.46 dB. The drop is largest on the kidney and Cells3D membranes volumes (−2.92 to −3.46 dB), where dense structure and moderate noise ($\sigma \approx 0.002$–$0.009$) combine to give the model substantial latitude to memorise noise once $N$ exceeds $N^*$.

The two deconvolved iSIM neuromast channels are cull-limited: their held-out maximum is interior (at 64K and 128K requested), but the post-fit cull holds the realised count nearly constant beyond it, at 32,715 to 40,191 splats for the nuclei and 74,825 to 77,900 for the membranes over the remaining doublings to 512K, so the later points are fits of nearly the same size. Their declines of 0.03 and 0.19 dB are below the largest held-out seed-to-seed difference measured on the replicated volumes (0.41 dB, §3.3; those replicates include no neuromast rows) and are not evidence of overfitting; the operating point is the held-out maximum.

A further 3 volumes plateau, i.e. their held-out curve stops within 0.3 dB of its maximum without a clear peak: the raw confocal Fly brain neurons tile from 64K, the fused, undeconvolved light-sheet *Drosophila* embryo from 256K (swept to 2M without turning over), and the deconvolved light-sheet Zebrafish embryo nuclei from 256K. The zebrafish held-out curve flattens from 256K (37.80 dB) to a 37.99 dB maximum at 1M and turns over by 2M, while its training PSNR keeps climbing: capacity spent on voxels the held-out metric does not reward.

The remaining two light-sheet volumes are signal-limited: on Mouse heart nuclei the held-out PSNR keeps rising through the largest tested capacity (extended to 2M splats, +0.70 dB over the last doubling), and on the *Tribolium* embryo it keeps rising through 2M as well (+0.48 dB over the last doubling), so neither reaches a finite optimum in its sampled range. That profile is consistent with a noise level low enough that the held-out signal-error term dominates the noise constant (§5) throughout the sweep.

The train/held-out gap at 512K is a parallel diagnostic: small on the two signal-limited datasets (0.56 and 0.64 dB) and on the two raw plateau volumes (0.86 and 1.14 dB), large (9.9–20.0 dB) on the kidney and Cells3D volumes where overfitting is most pronounced.

##### 3.3 Replicates

Every cell above is a single fit. To measure how much of a table entry is the seed, the ten confocal and spinning-disk datasets were re-fitted twice more in both modes with distinct RNG seeds (replicates 2 and 3; seed $= N \times 100 + \text{rep}$; 400 fits). Table 6 reports, at the five highlighted $N$, the spread between the two extra seeds and their mean's offset from the canonical seed, together with the operating point each replicate's held-out curve yields. Over the 100 (dataset, $N$) cells the seed-to-seed spread of the reconstruction PSNR has median 0.02 dB, 95th percentile 0.09 dB and maximum 0.22 dB (Kidney nuclei at 512K); the held-out PSNR behaves the same (median 0.01, maximum 0.20 dB). The canonical seed's reconstruction PSNR sits within 0.11 dB of the replicate mean on 94 of the 100 cells. The exceptions are Kidney actin (0.14 and 0.25 dB at 512K and 256K) and Cells3D membranes from 64K upward, where the offset reaches +0.85 dB at 512K (Table 6). Part of that offset is scoring domain, not seed variation: the canonical rate-distortion rows are scored from their decoded stores, while the replicate rows carry the optimiser's scores. In held-out PSNR the replicate-mean offset reaches 0.31 dB and a single replicate differs from the canonical seed by up to 0.41 dB (both Cells3D membranes at 128K). The detected $N^*$ agrees across all three seeds on nine volumes and moves by one doubling on the tenth (*C. elegans* embryo nuclei, 8K vs. 16K: its held-out curve is flat to within 0.02 dB between the two, so the knee is genuinely ambiguous there). Differences between table entries below $\sim$0.2 dB should therefore not be read as dataset effects on these ten volumes; the seven others (light-sheet, iSIM and the fly brain) have no replicates, and the two-seed spread is an observed range, not a confidence bound. Every difference this document interprets is larger.

##### 3.4 Noise floor

The Laplacian and Haar estimators agree within $\sim$15% on every dataset: the same noise variance is recovered by two qualitatively different high-pass filters, which is consistent with an estimated $\hat{\sigma}$ dominated by noise and not by residual signal structure (§5.3). The agreement is a one-voxel-

Table 6: Seed replicates on the ten confocal and spinning-disk datasets (rate-distortion PSNR, dB). Top: absolute difference between replicates 2 and 3, and the held-out operating point $N^*$ from each replicate's own blind-spot sweep. Bottom: mean of the two replicates minus the canonical run (replicate 1, the seed used for Tables 4–5).

| | \|rep 2 − rep 3\| (dB) | | | | | $N^*$ |
|---|---|---|---|---|---|---|
| Dataset | 1K | 8K | 32K | 128K | 512K | rep 1 / 2 / 3 |
| OpenCell-MAP4 nuclei | 0.02 | 0.01 | 0.01 | 0.01 | 0.01 | 16K / 16K / 16K |
| OpenCell-MAP4 microtubules | 0.02 | 0.01 | 0.01 | 0.00 | 0.01 | 64K / 64K / 64K |
| OpenCell-LMNB1 nuclei | 0.02 | 0.00 | 0.00 | 0.00 | 0.01 | 32K / 32K / 32K |
| OpenCell-LMNB1 nuclear lamina | 0.04 | 0.09 | 0.02 | 0.01 | 0.00 | 64K / 64K / 64K |
| Kidney nuclei | 0.01 | 0.09 | 0.06 | 0.03 | 0.22 | 64K / 64K / 64K |
| Kidney actin | 0.01 | 0.04 | 0.07 | 0.06 | 0.06 | 128K / 128K / 128K |
| Cells3D nuclei | 0.02 | 0.03 | 0.02 | 0.04 | 0.10 | 32K / 32K / 32K |
| Cells3D membranes | 0.01 | 0.05 | 0.03 | 0.00 | 0.07 | 32K / 32K / 32K |
| Blastocyst nuclear lamina | 0.21 | 0.02 | 0.02 | 0.01 | 0.05 | 32K / 32K / 32K |
| *C. elegans* embryo nuclei | 0.01 | 0.00 | 0.00 | 0.00 | 0.06 | 8K / 16K / 16K |

| | mean(rep 2, 3) − rep 1 (dB) | | | | |
|---|---|---|---|---|---|
| Dataset | 1K | 8K | 32K | 128K | 512K |
| OpenCell-MAP4 nuclei | -0.03 | +0.01 | +0.00 | +0.02 | +0.01 |
| OpenCell-MAP4 microtubules | -0.03 | +0.01 | +0.01 | -0.01 | +0.00 |
| OpenCell-LMNB1 nuclei | -0.04 | +0.01 | +0.00 | +0.00 | +0.00 |
| OpenCell-LMNB1 nuclear lamina | -0.05 | -0.01 | +0.01 | +0.01 | -0.04 |
| Kidney nuclei | +0.03 | +0.04 | +0.02 | +0.05 | +0.07 |
| Kidney actin | -0.02 | -0.01 | -0.02 | +0.02 | +0.14 |
| Cells3D nuclei | -0.03 | -0.00 | +0.01 | -0.00 | +0.04 |
| Cells3D membranes | +0.03 | +0.04 | +0.00 | +0.31 | +0.85 |
| Blastocyst nuclear lamina | -0.07 | +0.03 | -0.01 | +0.02 | -0.01 |
| *C. elegans* embryo nuclei | -0.00 | +0.01 | +0.03 | +0.01 | -0.01 |

scale check of the filtered residual and does not establish that the noise is independent between voxels: deconvolution correlates noise over the point-spread function, at scales both filters pass, so on a deconvolved volume the estimate is a level, not a test of independence. The background estimator consistently underestimates: at very dark voxels detector floors and pre-processing clamps compress the noise variance below the bulk value. The measurable noise floor spans 30–61 dB. From low to high, the order is: spinning-disk confocal (the OpenCell volumes, 30–43 dB), confocal (kidney, Cells3D nuclei, *C. elegans*, 37–44 dB), undeconvolved volumes (the multi-view fused *Drosophila* embryo nuclei 51 dB, the raw Fly brain neurons tile 52 dB), low-noise confocal (Blastocyst nuclear lamina, Cells3D membranes, 53–54 dB), and the deconvolved light-sheet and iSIM volumes (*Tribolium* 61 dB; the four others return zero MAD on every estimator, so their ceilings are not estimable). Low noise is where the held-out behaviour changes (Table 5): the three volumes without a held-out peak whose ceiling is estimable (signal-limited *Tribolium* and the two raw plateaus) have ceilings of 51 dB or higher, and the other two (signal-limited Mouse heart, the zebrafish plateau) return zero MAD. Every volume below 44 dB peaks. The two deconvolved neuromast channels also saturate the estimators; they are cull-limited (§3.2), and their small drops are not evidence of overfitting. Fig. 4 places every volume's operating point against its noise level.

Table 7: Per-dataset noise standard deviation from three robust estimators and the resulting theoretical PSNR ceiling $\text{PSNR}_{\max} = -20 \log_{10} \hat{\sigma}$. The ensemble $\hat{\sigma}$ is the median of the three estimators. The light-sheet Mouse heart nuclei and Zebrafish embryo nuclei volumes and the two deconvolved, floor-clipped iSIM neuromast channels return *exactly* 0.0 on every estimator: a zero MAD means that at least half of the filtered samples are zero and bounds nothing about the noise on the rest (§5.3); their ceilings are not estimable with these estimators and are marked '–'.

| Volume | $\hat{\sigma}_{\text{Lap}}$ | $\hat{\sigma}_{\text{Haar}}$ | $\hat{\sigma}_{\text{bg}}$ | $\hat{\sigma}$ | $\text{PSNR}_{\max}$ (dB) |
|---|---|---|---|---|---|
| OpenCell-MAP4 nuclei | 0.0234 | 0.0247 | 0.0039 | 0.0234 | 32.6 |
| OpenCell-MAP4 microtubules | 0.0300 | 0.0322 | 0.0050 | 0.0300 | 30.4 |
| OpenCell-LMNB1 nuclei | 0.0318 | 0.0338 | 0.0052 | 0.0318 | 30.0 |
| OpenCell-LMNB1 nuclear lamina | 0.0074 | 0.0079 | 0.0011 | 0.0074 | 42.6 |
| Kidney nuclei | 0.0086 | 0.0089 | 0.0069 | 0.0086 | 41.3 |
| Kidney actin | 0.0067 | 0.0075 | 0.0044 | 0.0067 | 43.5 |
| Cells3D nuclei | 0.0132 | 0.0134 | 0.0054 | 0.0132 | 37.6 |
| Cells3D membranes | 0.0022 | 0.0021 | 0.0020 | 0.0021 | 53.5 |
| Blastocyst nuclear lamina | 0.0019 | 0.0022 | 0.0000 | 0.0019 | 54.4 |
| *C. elegans* embryo nuclei | 0.0092 | 0.0093 | 0.0022 | 0.0092 | 40.7 |
| Fly brain neurons | 0.0025 | 0.0027 | 0.0000 | 0.0025 | 52.1 |
| *Tribolium* embryo nuclei | 0.0009 | 0.0008 | 0.0030 | 0.0009 | 61.0 |
| Mouse heart nuclei | 0.0000 | 0.0000 | 0.0000 | 0.0000 | – |
| Zebrafish embryo nuclei | 0.0000 | 0.0000 | 0.0000 | 0.0000 | – |
| *Drosophila* embryo nuclei | 0.0028 | 0.0030 | 0.0000 | 0.0028 | 51.1 |
| Neuromast membranes | 0.0000 | 0.0000 | 0.0000 | 0.0000 | – |
| Neuromast nuclei | 0.0000 | 0.0000 | 0.0000 | 0.0000 | – |

##### 3.5 Effective resolution and splat scale

Three facts follow from Table 8.

**The small end of the mixture.** The small end is set by the grid, not by the optics. The 10th-percentile splat width $\ell_{10}$ lies between 0.50 and 1.38 voxels on every volume, while the measured lateral resolution spans 2.2–24.4 pixels. Across the 17 volumes the two are only weakly related (Spearman $\rho = 0.45$, $p = 0.07$; log-log slope 0.23), and the resolution-to-$\ell_{10}$ ratio runs from 2.7 to 20 (median 5.7). The amplitude-weighted median $\ell_{50}^{a}$ (0.7–2.5 voxels) tracks resolution somewhat better ($\rho = 0.54$, $p = 0.02$) but still spans only a ∼3.5-fold range against the ∼11-fold range of the resolution in voxel units. In width-commensurate units (the decorrelation estimate is calibrated to a FWHM, so divide by the splat FWHM $2\sqrt{2 \ln 2}\, \ell_{10}$) the ratio is 1.15–8.7 (median 2.4), above 2 on 13 of the 17 volumes and close to 1 on Cells3D membranes and Mouse heart nuclei (1.15 and 1.21). The smallest fitted splats are therefore at or below the resolution of the data on every volume, well below it on most: the mixture spends its finest elements on voxel-scale intensity structure that the data sample but do not resolve, which is also why the fits are done in voxel units and the physical anisotropy is applied at display time. This is a statement about the scales the fit uses, not about the fidelity of any feature: a small $\ell_{10}$ shows the representation has no scale floor above the data, and says nothing about which features survive at $N^*$ (that is what the held-out curves and the residual maps measure).

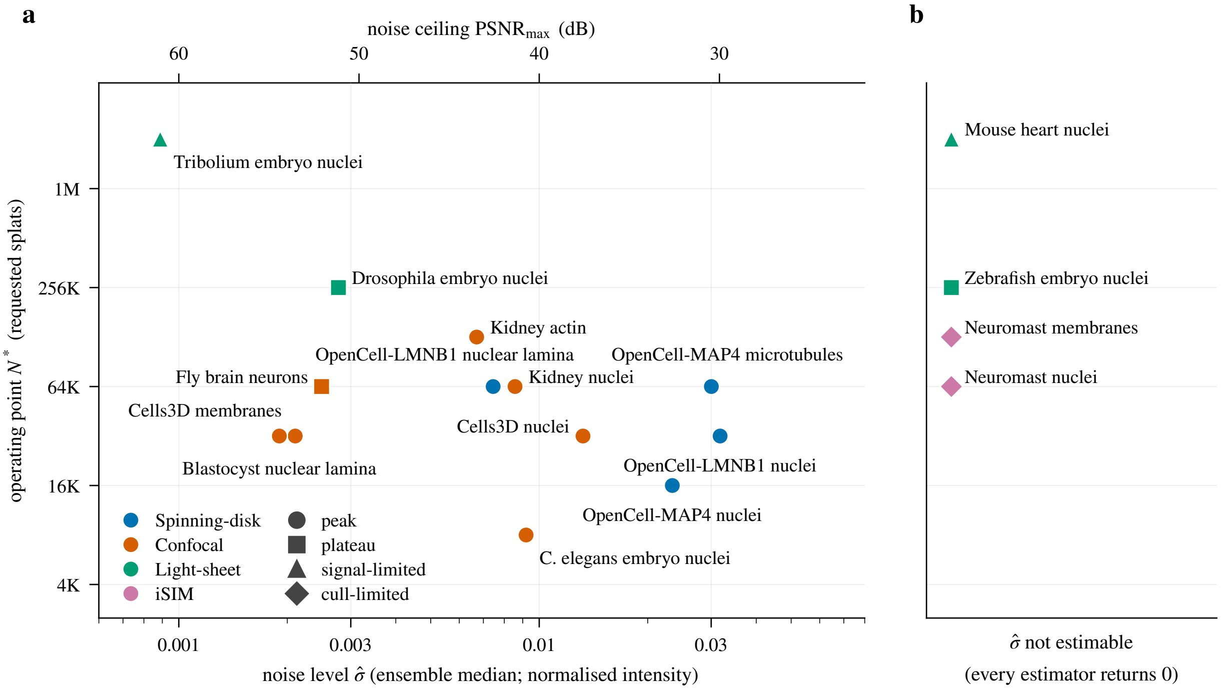


Figure 4: Operating point against noise level for the 17 volumes (Tables 5 and 7). **(a)** $N^*$ (requested splats) against the ensemble estimate $\hat{\sigma}$, with the noise ceiling on the top axis; marker colour is the imaging modality and marker shape the curve type. **(b)** The four volumes whose MAD estimators all return zero have no estimable $\hat{\sigma}$ and sit in the side strip at their $N^*$. Every volume with a ceiling below 44 dB shows a held-out peak, the plateaus and the signal-limited curve lie at the low-noise end, and the largest operating points belong to the light-sheet volumes.

**Resolution elements and $N^*$.** Over the 12 volumes with an interior held-out maximum (ten peaks, two cull-limited), $N^*$ correlates with the number of resolution cells (voxels per lateral$^2$×axial resolution) at $\rho = 0.65$ ($p = 0.02$, log-log slope 0.34) and with the foreground resolution cells (Otsu mask) at $\rho = 0.48$ ($p = 0.11$). A rank correlation over 12 volumes from 7 acquisitions is exploratory: paired channels of one acquisition are not independent samples, so the nominal $p$-values are optimistic, the kidney channels' unmeasured axial resolution is set equal to the lateral one, and a rank statistic says nothing about how much of the variance in $N^*$ it explains; the content-tiling calibration uses the local-maxima feature count instead ($N^* \propto n_{\text{features}}^{0.44}$; `luxar gsplat cal --fit-exponent`). We have not compared the two predictors on the same volumes here, so no claim is made that one is better than the other.

**Relocated splats.** Relocated splats keep their initialisation scale. In four volumes (Zebrafish embryo nuclei, *Drosophila* embryo nuclei, Neuromast membranes, Neuromast nuclei) between 11% and 44% of all splats at $N^*$ have a smallest lateral width within 0.01 voxel of $\ell = 0.5$, the scale at which the dynamic relocation step re-seeds the weakest splats every 50 iterations. The statistic tests that lateral width only (not isotropy, amplitude or relocation history) and is consistent with re-seeded splats keeping their initialisation scale on these long, high-$N$ fits. The held-out optimum is measured with them in place, so $N^*$ is unaffected, but the smallest-splat statistic of these four fits is a fitter constant, not a data property, and the same quality may be reachable with fewer,

Table 8: Effective resolution (top) and splat scale (bottom) for the 17 volumes. Lateral resolution by decorrelation analysis (median over slices, interquartile range in brackets) in pixels and, where the voxel size is recorded, in $\mu$m; axial resolution from the sectorial estimate in $z$-voxels and $\mu$m (†: fewer than 32 planes, no estimate). $\ell_{10}$: 10th-percentile axis-aligned splat width $\ell$ at $N^*$ (§2.5); $\ell^a_{50}$: amplitude-weighted median; "$\ell$ at 0.5": share of splats whose lateral $\ell$ is within 0.01 voxel of the relocation initialisation. ‡: no interior held-out maximum; $N^*$ is the plateau onset or the sweep endpoint (the cull-limited channels, interior maximum, carry no mark).

| Volume | Voxel ($z$, $y$, $x$), $\mu$m | Lateral resolution px [IQR] | Lateral resolution $\mu$m | Axial resolution $z$-vox ($\mu$m) |
|---|---|---|---|---|
| OpenCell-MAP4 nuclei | 0.500, 0.200, 0.200 | 7.8 [7.5–11.0] | 1.56 | 7.0 (3.52) |
| OpenCell-MAP4 microtubules | 0.500, 0.200, 0.200 | 12.8 [10.7–17.7] | 2.56 | 4.1 (2.07) |
| OpenCell-LMNB1 nuclei | 0.200, 0.200, 0.200 | 11.0 [10.7–12.8] | 2.21 | 24.4 (4.88) |
| OpenCell-LMNB1 nuclear lamina | 0.200, 0.200, 0.200 | 5.4 [4.3–5.9] | 1.09 | 6.6 (1.31) |
| Kidney nuclei | 1.250, 1.240, 1.240 | 2.4 [2.3–2.4] | 2.94 | –† |
| Kidney actin | 1.250, 1.240, 1.240 | 2.7 [2.6–2.7] | 3.35 | –† |
| Cells3D nuclei | 0.290, 0.260, 0.260 | 4.2 [4.0–4.3] | 1.09 | 5.9 (1.72) |
| Cells3D membranes | 0.290, 0.260, 0.260 | 2.3 [2.3–2.8] | 0.60 | 2.6 (0.75) |
| Blastocyst nuclear lamina | – | 3.2 [3.0–3.5] | – | 9.2 |
| *C. elegans* embryo nuclei | 0.750, 0.150, 0.150 | 24.4 [24.4–24.4] | 3.66 | 6.3 (4.74) |
| Fly brain neurons | 0.380, 0.190, 0.190 | 4.8 [4.8–4.8] | 0.91 | 4.7 (1.77) |
| *Tribolium* embryo nuclei | 0.381, 0.381, 0.381 | 3.0 [2.6–5.9] | 1.15 | 4.2 (1.59) |
| Mouse heart nuclei | – | 2.2 [2.2–2.2] | – | 7.3 |
| Zebrafish embryo nuclei | 1.625, 0.406, 0.406 | 7.1 [6.6–8.7] | 2.90 | 2.6 (4.21) |
| *Drosophila* embryo nuclei | 1.930, 0.406, 0.406 | 9.2 [9.2–9.4] | 3.72 | 2.5 (4.80) |
| Neuromast membranes | 0.250, 0.108, 0.108 | 3.2 [3.1–3.4] | 0.35 | 2.9 (0.74) |
| Neuromast nuclei | 0.250, 0.108, 0.108 | 2.5 [2.5–2.6] | 0.27 | 4.4 (1.11) |

| Volume | $\ell_{10}$ lateral vox | $\ell^a_{50}$ lateral vox | $\ell_{10}$ axial vox | $\ell$ at 0.5 % | $N^*$ |
|---|---|---|---|---|---|
| OpenCell-MAP4 nuclei | 1.38 | 1.93 | 1.48 | 1 | 16K |
| OpenCell-MAP4 microtubules | 0.92 | 1.31 | 0.79 | 0 | 64K |
| OpenCell-LMNB1 nuclei | 1.19 | 1.84 | 1.35 | 1 | 32K |
| OpenCell-LMNB1 nuclear lamina | 0.89 | 1.54 | 0.98 | 3 | 64K |
| Kidney nuclei | 0.62 | 0.90 | 1.08 | 1 | 64K |
| Kidney actin | 0.55 | 0.71 | 1.12 | 2 | 128K |
| Cells3D nuclei | 0.71 | 1.00 | 0.90 | 1 | 32K |
| Cells3D membranes | 0.86 | 1.25 | 0.95 | 1 | 32K |
| Blastocyst nuclear lamina | 0.66 | 0.85 | 1.00 | 2 | 32K |
| *C. elegans* embryo nuclei | 1.19 | 2.08 | 1.00 | 4 | 8K |
| Fly brain neurons | 0.91 | 1.00 | 0.95 | 1 | 64K‡ |
| *Tribolium* embryo nuclei | 0.88 | 2.50 | 0.92 | 0 | 2M‡ |
| Mouse heart nuclei | 0.77 | 1.17 | 1.01 | 2 | 2M‡ |
| Zebrafish embryo nuclei | 0.50 | 1.30 | 0.50 | 13 | 256K‡ |
| *Drosophila* embryo nuclei | 0.50 | 1.71 | 0.49 | 11 | 256K‡ |
| Neuromast membranes | 0.50 | 1.29 | 0.50 | 32 | 128K |
| Neuromast nuclei | 0.50 | 1.39 | 0.50 | 44 | 64K |

resized splats.

##### 3.6 Where the fit at $N^*$ goes wrong

Fig. 5 shows three residual features per volume for six volumes spanning the regimes (the strongest components that pass the minimum-size and non-overlap filters of §2.6; the third slot would go to the strongest component of the opposite sign if one ranked among the twenty strongest, which on these six volumes none did, so the three features of a volume share one sign), and Table 9 the detection statistics. Three regularities hold.

**The strongest deviations.** The strongest deviations are omissions. On 5 of the 6 volumes the strongest feature is a place where the reconstruction is *dimmer* than the source, and the dimmer components outnumber the brighter ones by 1.8–5.7× everywhere except on the background-subtracted neuromast. The features are the faint end of each volume's structures: a dim nucleus between bright neighbours (Kidney nuclei; residual peak 37% of the nearby source maximum), dim nuclei (*C. elegans*, 29%), faint membrane sheets at 2% of the display range (Cells3D membranes, 66%), and dense filament bundles on the noisiest volume (OpenCell-MAP4 microtubules, 23%; the only volume where more than a tenth of the foreground is flagged, 11.4%). The percentages are the smoothed residual peak divided by the unsmoothed source maximum in its neighbourhood: a contrast statistic that ranks features, not the fraction of an object's peak intensity that is lost (the point-wise loss at the same voxels is larger, 25–88%).

**The neuromast gaps.** On the neuromast the strongest brighter components are the gaps. On the background-subtracted neuromast membranes the strongest brighter components are the small dark lumina between membranes, where the smoothed residual reaches 0.025 on a 0–1 range (the reconstruction itself reaches 0.033 where the source is exactly zero) beside membranes at 0.97. Gaussians cannot hold an exact zero next to a bright wall, so on a volume whose background is exactly zero the strongest errors are spill-overs (brighter components exist on every volume, Table 9, but rank below the omissions elsewhere). Only 0.1% of its foreground is flagged.

**The flagged fraction.** The flagged fraction, not the PSNR, separates the regimes. Between 0.1% and 11.4% of the foreground voxels lie beyond the threshold of six robust standard deviations. The sparse confocal volumes sit at the low end (a few dim objects), the deconvolved zebrafish volume at 6.4% (bright nuclear clusters under-fitted by 16% at 256K splats; the capacity, not the noise, is what limits it, consistent with its plateau held-out curve), and the MAP4 volume highest. These are the features a user should look for in a reconstruction: the dimmest objects of a field, thin faint sheets, and the darkest gaps of dense structures; bright, isolated objects did not appear among the strongest deviations on these six volumes.

##### 3.7 Cross-dataset summary

The three diagnostics (rate-distortion, held-out, noise-floor) for each dataset are reproduced in the per-dataset figure pages of §A: (i) rate-distortion curves (PSNR / SSIM / fit time / cull-percent vs. $N$) with the noise-floor line overlaid. (ii) Reconstruction slice montages at the lowest count, 16K, the detected $N^*$ and the largest count, annotated with the measured on-disk compression ratio. (iii) Blind-spot train- and held-out-PSNR curves with the noise constant overlaid (the held-out peak is the detected $N^*$). Three features are visible at a glance across datasets.

Table 9: Residual-feature detection at the operating point. Threshold: six robust standard deviations of the smoothed residual, in normalised intensity units; "fg %": share of Otsu-foreground voxels beyond it; component counts by sign; the strongest feature's sign, peak smoothed residual, local source maximum and voxel count.

| | | | flagged | | strongest feature | | | |
|---|---|---|---|---|---|---|---|---|
| Volume | $N^*$ | threshold | fg % | dimmer / brighter | sign | peak $\lvert r\rvert$ | source | voxels |
| Kidney nuclei | 64K | 0.010 | 1.9 | 1861 / 370 | dimmer | 0.022 | 0.06 | 719 |
| Cells3D membranes | 32K | 0.004 | 1.5 | 2316 / 432 | dimmer | 0.011 | 0.02 | 8 531 |
| *C. elegans* embryo nuclei | 8K | 0.018 | 0.3 | 1141 / 200 | dimmer | 0.048 | 0.17 | 247 |
| OpenCell-MAP4 microtubules | 64K | 0.050 | 11.4 | 20357 / 11366 | dimmer | 0.197 | 0.86 | 1 834 |
| Neuromast membranes | 128K | 0.010 | 0.1 | 18 / 1553 | brighter | 0.025 | 0.97 | 98 |
| Zebrafish embryo nuclei | 256K | 0.023 | 6.4 | 34170 / 13280 | dimmer | 0.088 | 0.53 | 21 254 |

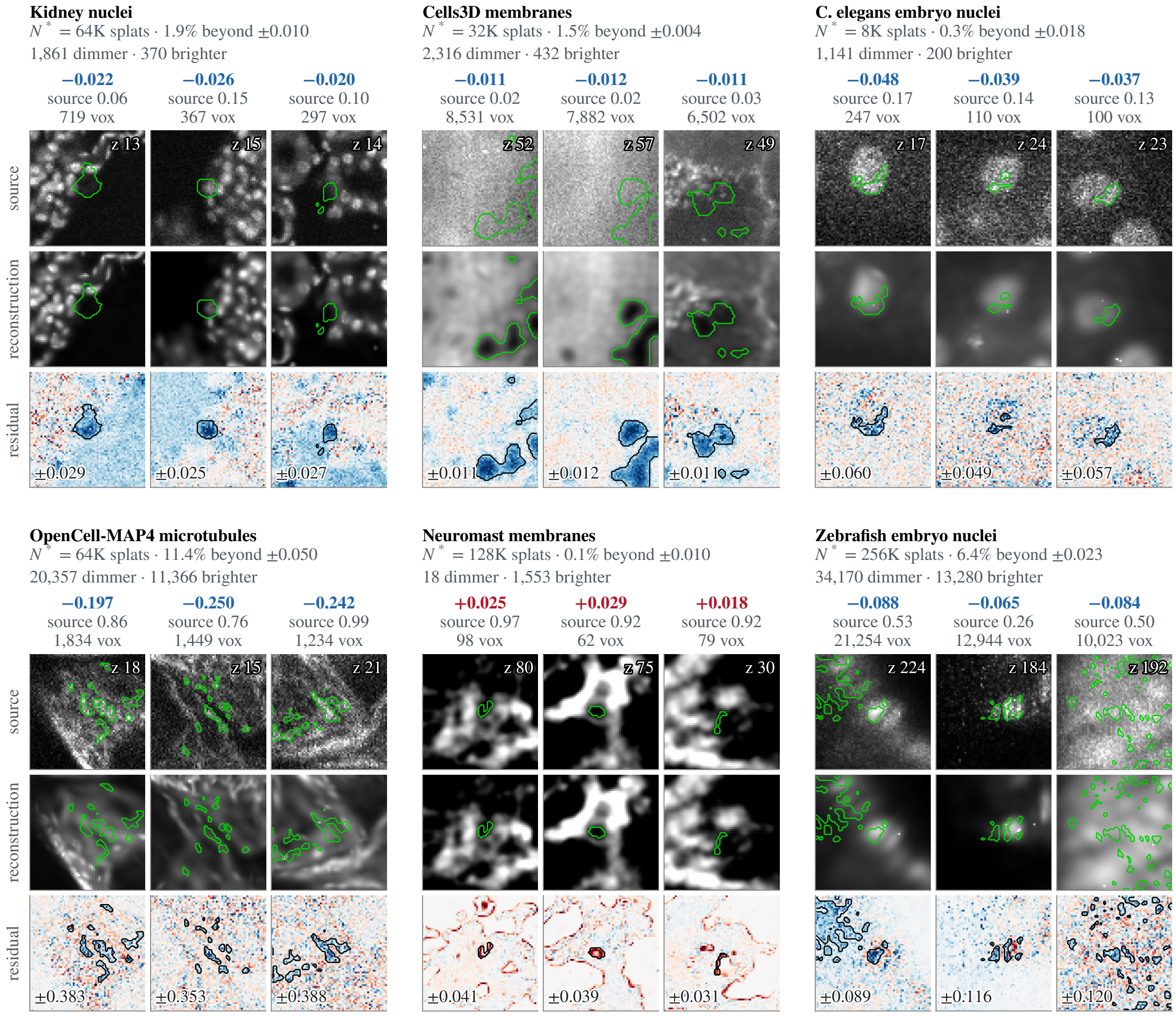


Figure 5: The three strongest residual features of six fits at $N^*$ (also Supplementary Fig. 5 of the main text). Per volume: source, reconstruction and signed residual (blue: reconstruction dimmer, red: brighter) through the slice where the smoothed residual peaks, with the detected component outlined; source and reconstruction share one display window per crop. Column titles give the sign, the peak smoothed residual, the local source maximum and the component size; the slice index is printed in the source crop and the residual display range in the residual crop.

(a) Every confocal and spinning-disk dataset shows a divergence between train and held-out PSNR above $N^*$ (the gap-at-512K column of Table 5), confirming noise memorisation.

(b) No light-sheet volume shows a held-out decline beyond the 0.06 dB by which the zebrafish curve dips from 1M to 2M. The *Tribolium* embryo and Mouse heart nuclei have train and held-out PSNR tracking each other (gap $\leq$ 0.64 dB through 512K, opening to 1.1 and 2.2 dB at the 2M extension point). On the zebrafish embryo the held-out curve plateaus as the training curve keeps rising (2.95 dB gap at 512K, 5.2 dB at 2M): capacity spent on voxels the held-out metric does not reward, without the decline that would mark noise memorisation.

(c) Reconstruction PSNR at 512K crosses the noise-floor ceiling on four confocal volumes (Kidney nuclei +3.9, Kidney actin +4.6, Cells3D membranes +4.6, Cells3D nuclei +7.2 dB above ceiling) and stays below it everywhere else. These are not the four with the largest held-out drops: Blastocyst's 0.7 dB drop exceeds Cells3D nuclei's 0.5 dB. The gap to ceiling ranges from 0.25 dB (*C. elegans*, near-saturated) to 6 dB on the *Tribolium* embryo, 13 dB on Blastocyst nuclear lamina and 16 dB on the *Drosophila* embryo, and is not estimable on the four volumes whose MAD estimators return zero.

#### 4 Interpretation

**Overfitting on 10 of 17 volumes.** Splats overfit microscopy data on 10 of the 17 volumes. With conservative culling (cull-retention 0.999) the model's full capacity is preserved post-fit, and held-out PSNR peaks and declines on every confocal and spinning-disk dataset tested except the fly brain, which plateaus (Table 5). On the two iSIM neuromast channels the same cull removes most of the pool above the operating point, so the sweep no longer varies capacity there and the question stays open for those two volumes.

With cull-retention 0.99 the post-fit cull removes a large fraction of the splats and the held-out drop is small or absent; that configuration is not part of this document, so we state it as the reason for the present setting, not as a measured ablation, and whether the removed splats were the noise-fitting ones or merely the faintest was not tested. The present setting (0.999) is the regime in which the question *"what splat count is right?"* is answered by capacity, not by post-hoc filtering.

**The spread of $N^*$.** $N^*$ varies by more than an order of magnitude. Across the 10 overfit-exhibiting volumes, it ranges from 8K to 128K (Table 5); the largest $N^*$ (Kidney actin, 128K) is 16× the smallest (*C. elegans* embryo nuclei, 8K). Two consistent determinants: (i) signal density: sparse signals saturate at small $N^*$ (the *C. elegans* embryo is mostly empty); dense filaments require more capacity (Kidney actin); (ii) noise level: higher $\sigma$ leaves a smaller window between under- and over-fitting before noise memorisation begins (OpenCell-MAP4 nuclei, $\sigma = 0.023$, peaks at 16K, is flat to within 0.01 dB at 32K and declines from 64K on). Volume size is confounded with both in the present sample: the three largest tested volumes (*Tribolium* embryo nuclei, Mouse heart nuclei, Zebrafish embryo nuclei) are also three of the four light-sheet (low-noise) volumes, none of which overfits at any tested $N$.

**The light-sheet exception.** The three light-sheet volumes with the lowest measured noise (*Tribolium* embryo nuclei, ceiling 61 dB; Mouse heart nuclei and Zebrafish embryo nuclei, zero MAD, ceiling not estimable) have noise levels low enough that the held-out signal-error term dominates the noise constant $\sigma^2$ in the decomposition (§5) across most of the tested capacity range. Two of them show no finite optimum: on Mouse heart nuclei the held-out PSNR is still rising at the largest tested capacity (+0.70 dB from 1M to 2M splats) and on the *Tribolium* embryo it is still rising at 2M (+0.48 dB from 1M), so both are classified signal-limited (Table 5, drop column blank). The zebrafish embryo *does* turn over: its held-out PSNR is within 0.3 dB of its maximum from 256K, peaks at 1M and dips −0.06 dB by 2M.

Two non-exclusive readings of the light-sheet behaviour follow. First: when $\sigma^2$ is small the signal-error term dominates the held-out MSE across the tested capacity, so any crossover is pushed to very high $N$ or below the rounding floor of our peak detector (0.1 dB; §2.3). Second: on the *Tribolium* embryo the training PSNR of the blind-spot fit still sits 4 dB below the 61 dB noise-floor ceiling at 2M (56.7 dB), and its train and held-out curves are within 1.1 dB of each other, so the model has not yet exhausted the recoverable signal: it is signal-limited (capacity-limited under $L^1$ + bounded splat parameters), not noise-limited, so far as the independence assumption of §5 survives deconvolution (§5.3). The two readings are consistent with each other: when $\sigma^2$ is very small, the held-out signal-error term *is* the same quantity that the rate-distortion curve measures.

The heart's rise has a possible confound: every fit at $N \geq 64$K, including the extended 1M and 2M points, hits the 20,000-iteration cap (the recorded iteration count is 20,000 for each), so the still-rising held-out curve could be under-convergence at the largest capacities instead of signal that only more splats can capture. A cap-lifted run tests this (Table 10: a 60,000-iteration cap, otherwise the identical recipe, against the 512K, 1M and 2M rows of the canonical sweep). Tripling the budget changes the train PSNR by at most 0.23 dB and the held-out PSNR by −0.19 to +0.05 dB, and no fit converges under either cap. Under-convergence is therefore not what makes the held-out curve rise: at 60,000 iterations the held-out PSNR still climbs, 42.85 → 44.54 → 45.11 dB, and the last doubling adds +0.57 dB against +0.70 dB at the cap. The signal-limited classification stands, with the operating point at 2M (the smallest $N$ within 0.3 dB of the maximum). The train/held-out gap grows with $N$ under both budgets, from 0.64 dB at 512K to 2.2 dB at 2M at the cap and from 0.81 to 2.4 dB with the cap tripled, as the noise-memorisation term of §5 allows. The opening gap is consistent with a held-out peak beyond 2M splats (the volume is 99.7 M voxels, one splat per 50 voxels at 2M), but a rising held-out curve with an opening train gap is equally consistent with no finite peak, so we claim only that the heart is signal-limited *within the tested range*.

**Two complementary stopping criteria.** The noise floor and the held-out peak provide complementary stopping criteria. The noise floor gives a per-dataset *reference level* ($\text{PSNR}_{\text{max}}$, the PSNR of the clean signal against its own noisy measurement), independent of model class: a reconstruction scored against the noisy source cannot exceed it without fitting noise, though the level says nothing about how close a reconstruction is to the clean signal. The held-out peak gives the per-dataset *optimum* $N^*$ given a fixed model class (here, $L^1$-fitted Gaussian splats) and a fixed

Table 10: Iteration-cap sensitivity on Mouse heart nuclei (blind-spot sweep, 5% mask): train and held-out PSNR (dB) and their gap at the 20,000-iteration cap used throughout this document and with the cap tripled. The 20,000-iteration rows are the canonical sweep of Table 5; the 60,000-iteration rows are the same sweep with the cap tripled. No fit converges under either cap. Wall time in seconds on an RTX PRO 6000; the 60,000-iteration fits share the card with other jobs, so their times are upper bounds. Single fit per cell.

| $N$ | iterations | train | held-out | gap | time (s) |
|---|---|---|---|---|---|
| 512K | 20,000 | 43.44 | 42.80 | 0.64 | 376 |
| 1M | 20,000 | 45.98 | 44.61 | 1.37 | 440 |
| 2M | 20,000 | 47.50 | 45.30 | 2.19 | 632 |
| 512K | 60,000 | 43.66 | 42.85 | 0.81 | 2701 |
| 1M | 60,000 | 46.15 | 44.54 | 1.61 | 2953 |
| 2M | 60,000 | 47.54 | 45.11 | 2.43 | 3464 |

splat parameterisation. They answer different questions: "Where does agreement with the noisy source reach the noise level?" vs. "How many splats *should* we use?". Read together (Tables 4–7), they place each dataset in absolute terms: confocal and spinning-disk reconstructions sit 0.8–8.0 dB below their floors at $N^*$ (with the exception of Blastocyst nuclear lamina, 15 dB below: its noise floor is unusually high for confocal at 54 dB and the model is far from saturating the recoverable signal), so their agreement with the noisy source is not yet at the noise level. The four dense confocal volumes (Kidney nuclei, Kidney actin, Cells3D nuclei, Cells3D membranes) cross their estimated floors by 4–7 dB at 512K, a direct signal of noise memorisation. The *Tribolium* embryo sits 5 dB below its floor at 2M and the two other light-sheet datasets have no estimable floor (their MAD estimators return zero, so their headroom is not measurable).

### 5 Theoretical context

#### 5.1 Held-out loss decomposition

Let $V : \Omega \to \mathbb{R}$ be the (unknown) clean volume and $Y_j = V_j + \epsilon_j$ the noisy observation, with $\epsilon_j$ independent across $j$, zero-mean, and variance $\sigma^2$. Let $M \subset \Omega$ be the blind-spot mask and $\tilde{Y}$ the fit-time volume in which $Y_M$ has been replaced by donut medians of the *unmasked* voxels of the 3×3×3 neighbourhood, so $\tilde{Y}$ is a function of the unmasked voxels only. Let $\hat{V}_N = \hat{V}_N(\tilde{Y})$ be the splat-fitted volume at splat count $N$, and $\hat{V}_N(\boldsymbol{x}_j)$ its value at the position $\boldsymbol{x}_j$ of voxel $j$.

**Proposition 1** (held-out MSE decomposition)**.** *For $j \in M$, $\hat{V}_N(\boldsymbol{x}_j)$ depends on $\tilde{Y}$ (and hence on $V$ and on $\epsilon_{\Omega\setminus M}$) but* not *on $\epsilon_j$ (since $Y_j$ was replaced before fitting). Then:*

$$\mathbb{E}[\mathrm{MSE}_{\textit{held-out}}(N)] = \underbrace{\frac{1}{|M|}\sum_{j\in M}\mathbb{E}[(\hat{V}_N(\boldsymbol{x}_j) - V_j)^2]}_{\textit{signal error at } N} + \sigma^2. \tag{2}$$

*Proof.* Expand $(\hat{V}_N(\boldsymbol{x}_j) - Y_j)^2 = (\hat{V}_N(\boldsymbol{x}_j) - V_j - \epsilon_j)^2$. The cross-term has expectation zero because $\hat{V}_N(\boldsymbol{x}_j)$ and $\epsilon_j$ are independent (the former is a function only of $\tilde{Y}$, which omits $Y_j$; the latter is the noise at $j$) and $\mathbb{E}[\epsilon_j] = 0$. The squared-noise term has expectation $\sigma^2$. Sum over $j \in M$ and divide by $|M|$. □

**Corollary 1.** *Since $\sigma^2$ is independent of $N$,*

$$N^\dagger \;=\; \arg\min_N \mathbb{E}[\mathrm{MSE}_{\textit{held-out}}(N)] \;=\; \arg\min_N \frac{1}{|M|} \sum_{j \in M} \mathbb{E}[(\hat{V}_N(\boldsymbol{x}_j) - V_j)^2].$$

*The held-out MSE-minimising splat count is the splat count that minimises expected signal-reconstruction error at the masked voxels.*

We write $N^\dagger$ for this expected-value optimum to keep it apart from the reported operating point $N^*$ of §2.3, which is read from one noisy realisation of the curve on a sparse grid and, for a plateau, is deliberately the onset within 0.3 dB of the maximum and not the argmax. The two coincide in expectation for a clear peak sampled at its argmax and differ by construction on a plateau.

This is the special case of the self-supervised loss decomposition of Batson and Royer [1] for an explicitly $\mathcal{J}$-invariant estimator $\hat{V}_N$ (an estimator is $\mathcal{J}$-invariant for a set of voxels $\mathcal{J}$ when its prediction at every voxel of $\mathcal{J}$ does not depend on the observed values in $\mathcal{J}$; here $\mathcal{J} = M$). Gaussian splatting is not intrinsically $\mathcal{J}$-invariant (a single splat covers many voxels), but the donut-median replacement step *enforces* $\mathcal{J}$-invariance with respect to $M$ by construction: $Y_j$ is replaced before fitting, so $\hat{V}_N(\boldsymbol{x}_j)$ has no functional dependence on $\epsilon_j$ (or on $Y_j$ at all). The constant $\sigma^2$ is independent of $N$, so the held-out PSNR peak detects the $N^*$ that minimises signal error.

##### 5.2 Synthetic control: does the held-out maximum find the true optimum?

On real data the clean volume is unknown, so the argmax of the held-out curve can only be compared with itself. A synthetic volume makes the comparison direct. We build a clean volume of 48×128×128 voxels from 2500 anisotropic Gaussian blobs of varied width, amplitude and in-plane orientation over a smooth dim background, add pixel-independent Gaussian noise of standard deviation $\sigma = 0.02$ (a light-sheet-like level, ceiling 34.0 dB) or $\sigma = 0.08$ (a confocal-like level, ceiling 21.9 dB), and run the pipeline of §2.2 unchanged: the same 5% mask, the donut-median fill, the shipped fitter at 9 requested counts from 250 to 64K with 2,000 iterations, and the same renderer. Both fills are run, the unmasked-donor fill that Proposition 1 assumes and the all-donor fill the canonical sweeps used. For every fit we record the held-out PSNR against the noisy source (the criterion) and the true PSNR against the clean volume, which is unavailable on real data.

The result (Fig. 6) is the one Proposition 1 predicts. In all 4 arms the held-out maximum and the true-PSNR maximum fall on the same requested count (4 of 4): 4K at $\sigma = 0.02$ and 2K at $\sigma = 0.08$, so the optimum moves down by one doubling when the noise quadruples, and the detector of §2.3 reads every arm as a peak and returns that count, at no cost in true PSNR. The two fills agree to within the seed spread at every count, so the leak of §2.2 does not move the optimum on this volume either. The noise level shows up as predicted: at $\sigma = 0.08$ the held-out PSNR spans 1.04 dB across the sweep while the true PSNR spans 3.67 dB, and beyond

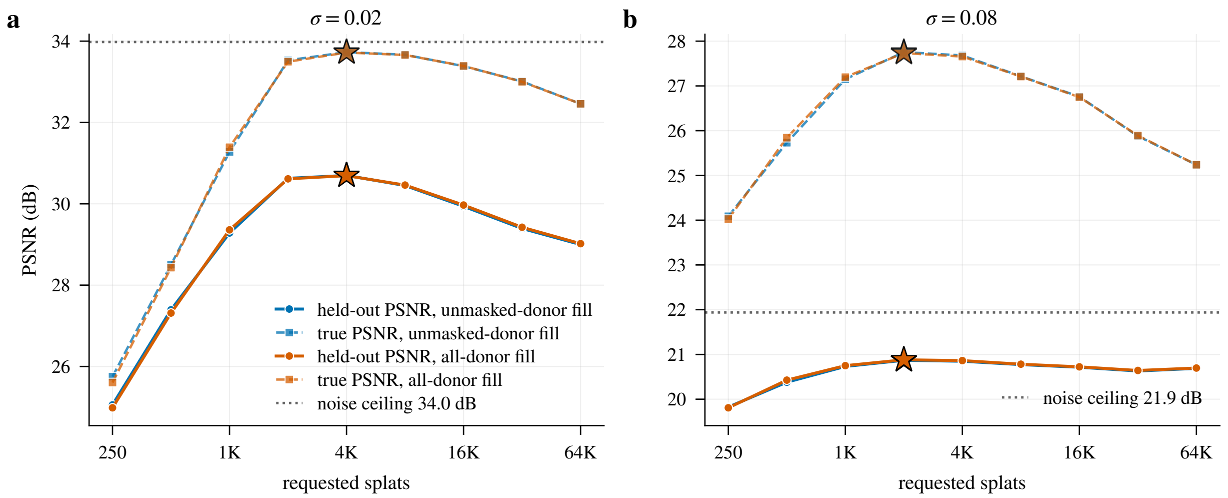


Figure 6: Synthetic control. Held-out PSNR against the noisy source (circles, solid) and true PSNR against the clean volume (squares, dashed) versus requested splat count, for the unmasked-donor fill (blue) and the all-donor fill (vermilion), at **(a)** $\sigma = 0.02$ and **(b)** $\sigma = 0.08$; stars mark the maximum of each curve and the dotted line the noise ceiling $-20\log_{10}\sigma$. The held-out curve sits at the noise constant plus the signal error (Proposition 1), so its range is compressed at high noise while its maximum stays at the count where the true error is smallest.

the optimum the held-out curve falls by 0.18 dB where the true PSNR falls by 2.52 dB, because the noise constant $\sigma^2$ dominates the held-out MSE and compresses the signal-error term; the maximum is nonetheless in the right place, and the training PSNR, which keeps rising with the count, would not find it. At $\sigma = 0.02$ the two curves have comparable ranges (5.64 and 7.98 dB) and the held-out curve sits below the ceiling by the signal-error term at its maximum. This is one volume, one noise model and one mask; it validates the criterion under the assumptions of the proposition, not under the correlated noise of §5.4.

#### 5.3 Noise-floor estimators

The Laplacian [4] and Haar [3] estimators are both high-pass: they apply a local linear filter that annihilates low-order polynomial structure in the clean signal $V$ but passes pixel-independent noise. Under the observation model $Y = V + \epsilon$ with independent $\epsilon$, the filter output's robust scale (MAD, rescaled to $\sigma$ via $\Phi^{-1}(0.75) \approx 0.6745$) is proportional to $\sigma$ with a known kernel-dependent constant. The two filters differ in their sensitivity profile (Laplacian: isotropic, 3D high-pass; Haar: directional, slice-by-slice 2D), and their agreement within $\sim$15% on every dataset (Table 7) is consistent with a filtered residual dominated by noise and not by unsuppressed signal; it does not test whether that noise is independent between neighbouring voxels. The background-region estimator is sensitivity-limited: detector or background-subtraction floors at the darkest voxels compress the noise variance below the bulk value, so $\hat{\sigma}_{\mathrm{bg}}$ systematically underestimates and is excluded from the headline ceiling by the median (which is robust to one outlier estimator on the low side). On the *Tribolium* embryo the background estimator instead *over*estimates (0.0030 vs. Laplacian/Haar at 0.0009) because the dark-voxel population includes structured background artefacts that the high-pass filters suppress; the median correctly defers to the high-pass estimators.

On Mouse heart nuclei (and likewise on Zebrafish embryo nuclei) all three estimators return *exactly* 0.0 (Table 7). A MAD of zero means that at least half of the filtered samples are exactly zero (here because the quietest regions are flat at the source quantisation step), and it bounds nothing: sparse noise of any variance on the other half leaves the MAD at zero. We therefore report no ceiling for these four volumes ('–' in the tables) and treat their noise level as unmeasured, not as below $10^{-3}$.

##### 5.4 Limitations of the held-out criterion

Three assumptions limit the strict applicability of Proposition 1.

1. **Pixel-independent noise.** The cross-term in the proof vanishes because $\mathbb{E}[\epsilon_j] = 0$ *and* because $\hat{V}_N(\boldsymbol{x}_j) \perp \epsilon_j$ (independence). If preprocessing (percentile clipping, histogram stretching, deconvolution) introduces spatial correlations into the residual, $\hat{V}_N(\boldsymbol{x}_j)$ becomes a (weak) function of $\epsilon_j$ via the neighbouring unmasked voxels, and the cross-term is no longer exactly zero. This flattens the held-out peak and may explain the relatively shallow drop on OpenCell-LMNB1 nuclear lamina ($-0.38$ dB), where the acquired data has been clip-percentile-normalised.

2. **The donut-median fill biases the prediction towards the local mean.** With the unmasked-donor fill, $\tilde{Y}$ at a masked position is a function of the unmasked voxels only, so its correlation with the signal $V_j$ is exactly what self-supervision relies on and is not a violation of $\mathcal{J}$-invariance. What the fill does introduce is a modelling bias: for very fine-grained signal it pulls the fitted $\hat{V}_N$ at masked positions towards the local mean instead of the true value. That changes the signal-error term of Eq. (2) but not the noise term. (The canonical sweeps' all-donor fill, which also used other held-out voxels as donors, is the separate concern of §2.2.)

3. **Discrete sampling of $N$.** The sweep evaluates ten geometrically spaced splat counts; the true $N^*$ lies between sampled points. A finer sweep around each detected peak (e.g., five extra counts in $[N^*/2, 2N^*]$) would tighten the empirical optimum and could move a "Plateau"-type volume into the "Peak" category.

#### 6 Regime-robust held-out metrics for sparse, deconvolved volumes

On large, sparse volumes the full-volume held-out PSNR is *background-dominated*: because most voxels are empty, a trivial predict-zero reconstruction already scores tens of dB, so the absolute held-out PSNR reflects empty space as much as signal fidelity. This is most acute on the deconvolved light-sheet Zebrafish embryo nuclei volume, whose MAD estimators all return zero (no estimable ceiling) and whose held-out PSNR flattens by its 256K operating point (37.80 dB; it edges up only to a 37.99 dB maximum at 1M splats and then turns over by 2M: a plateau, Table 5).

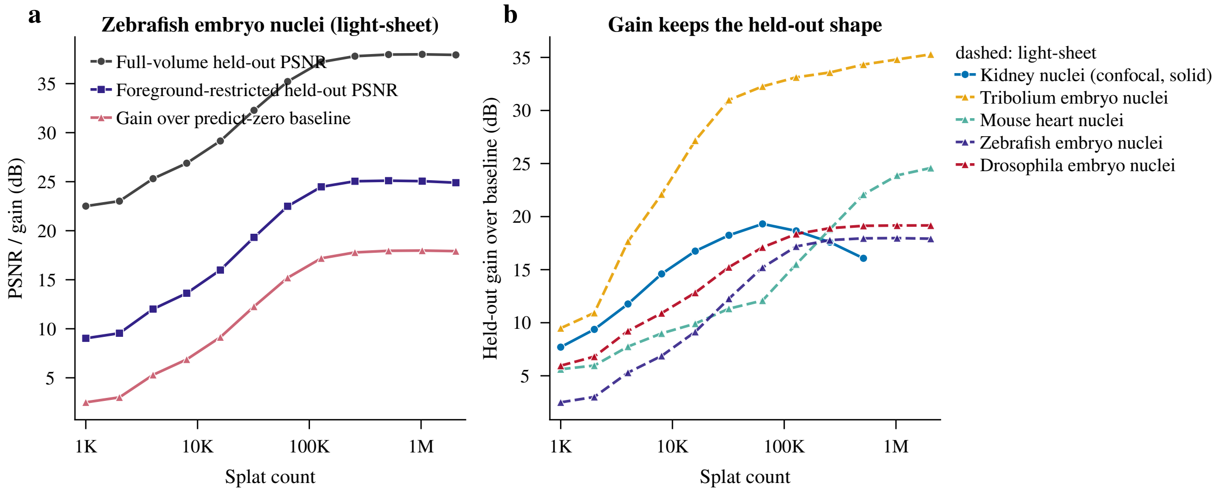


Figure 7: Regime-robust held-out diagnostics. **(a)** On the deconvolved light-sheet Zebrafish embryo nuclei volume, the full-volume held-out PSNR (37.80 dB at 256K) is inflated by correctly-reconstructed background; the gain over the predict-zero baseline (17.79 dB) and the foreground-restricted held-out PSNR (25.04 dB) expose the true signal fidelity. **(b)** The gain curve is the held-out PSNR curve shifted by a per-volume constant, so it keeps the held-out shape: it peaks at the same 64K on Kidney nuclei (solid) and keeps rising or levels off on the four light-sheet volumes (dashed); only its absolute level changes from volume to volume.

We therefore compute two complementary held-out diagnostics on the same blind-spot sweep. The *gain over the predict-zero baseline*, $10\log_{10}(\mathrm{MSE}_0/\mathrm{MSE}_{\mathrm{held}})$ where $\mathrm{MSE}_0$ is the error of an all-zeros reconstruction at the masked voxels, subtracts the free background credit; and the *foreground-restricted held-out PSNR* evaluates only masked voxels that lie above an Otsu foreground threshold. On the zebrafish embryo these reveal that the 37.80 dB figure overstates signal fidelity by $\sim$13 dB: the gain over baseline is only 17.79 dB and the foreground-restricted PSNR only 25.04 dB (Fig. 7a). The gain differs from the held-out PSNR by a per-volume constant ($10\log_{10}\mathrm{MSE}_0$), so it selects exactly the same $N^*$ and curve type on every volume (Fig. 7b shows the identical peak on Kidney nuclei): it changes how a score reads across volumes, not the operating point. The foreground-restricted PSNR is a different loss and can move the optimum, at the price of depending on an intensity-defined mask. Both are exposed by the shipped calibration command (`luxar gsplat cal --k-star-metric gain` or `psnr_foreground`) as reporting diagnostics for large sparse or deconvolved volumes.

### 7 Summary

The findings of this document, in the order the sections establish them. Blind-spot cross-validation identifies an operating point $N^*$ on 15 of the 17 volumes (Table 5): ten clear interior peaks, three plateaus and the two cull-limited iSIM neuromast channels, whose realised count stops growing beyond the peak so that their small declines are not evidence of overfitting. $N^*$ spans 8K–128K on the confocal and spinning-disk data, reaches 256K on the deconvolved zebrafish embryo, and follows signal density and noise level. The two signal-limited light-sheet volumes (Mouse heart nuclei, *Tribolium* embryo nuclei) are still rising at 2M splats, and on the heart a tripled iteration budget leaves the curve rising while the train/held-out gap opens (Table 10), so the label holds within the tested range. Seed replicates move $N^*$ by at most one doubling on the ten replicated volumes (§3.3). The synthetic control (§5.2) puts the held-out maximum on the same count as the true-PSNR maximum in all 4 arms, at two noise levels and under both fills. The smallest fitted splats sit at or below the data's resolution on every volume (§3.5), and on 5 of the 6 volumes examined the strongest errors at $N^*$ are omissions of the faintest structures (§3.6).

The held-out criterion (Corollary 1) requires only the noisy data and a random mask, and assumes no noise distribution beyond zero-mean noise that is independent across voxels given the signal (§5; its limits are listed in §5.4). The noise-floor analysis answers the other question, where the noise level of the source lies, through two high-pass MAD estimators that agree within $\sim$15%; on the four volumes whose estimators return zero MAD it returns no estimate. The ceiling is an upper bound (§2.4), so a gap below it only bounds the headroom.

The same framework is packaged as the `luxar gsplat cal` command: it sweeps splat count under the donut-median blind-spot protocol, applies the three-way peak/plateau/signal-limited detection rule of §2.3, estimates the noise-floor PSNR ceiling from the same MAD ensemble, and emits its recommended count, $K^*$ in the command's own notation (§2.3), plus the curve type. It differs from the benchmark sweeps in two ways: it defaults to background-floor suppression (`--floor auto`; the sweeps used `none`) and uses the unmasked-donor fill of §2.2. The same calibration also underlies content-adaptive fitting: a feature-density budget (`gsplat fit --tiling content`) transfers the calibrated splat count to sub-regions of a large volume via the relation $N^* \propto n_{\text{features}}^{\alpha}$, where the exponent $\alpha$ can be measured directly with `gsplat cal --fit-exponent` (by calibrating $N^*$ at several spatial scales and regressing $\log N^*$ on $\log n_{\text{features}}$). The shipped default $\alpha = 0.44$ is an engineering heuristic; the exponent is dataset-dependent and a robust per-dataset calibration is left for future work.

The principal limitations are scope and replication. The seventeen volumes, all 3D microscopy, leave open whether the same patterns hold under non-pixel-independent noise (after deconvolution, denoising, or aggressive preprocessing), under different optimisers or regularisers, or for 4D / time-series data. The doubling grid locates $N^*$ to the grid's resolution (one doubling) for the peaked curves and not at all for the two rising ones; seed replicates (§3.3) exist for the ten confocal and spinning-disk volumes, where they move $N^*$ by at most one doubling, while the seven others (light-sheet, iSIM and the fly brain) are single fits. A finer sweep around each detected peak, seed replicates on those seven volumes and replication with distinct masks would tighten the empirical optimum.

### A Per-dataset figures

For each dataset, three panels are reproduced: **(a)** rate-distortion curves (PSNR, SSIM, fit time, and culling percentage vs. splat count), with the estimated noise floor (Table 7) overlaid on the PSNR panel. **(b)** Reconstruction slice montages at the lowest, 16K, the detected-$N^*$, and the largest splat count. **(c)** Blind-spot train- and held-out-PSNR curves, the noise variance overlaid on the MSE panel; the held-out peak is $N^*$.

#### A.1 OpenCell-MAP4 nuclei: Spinning-disk

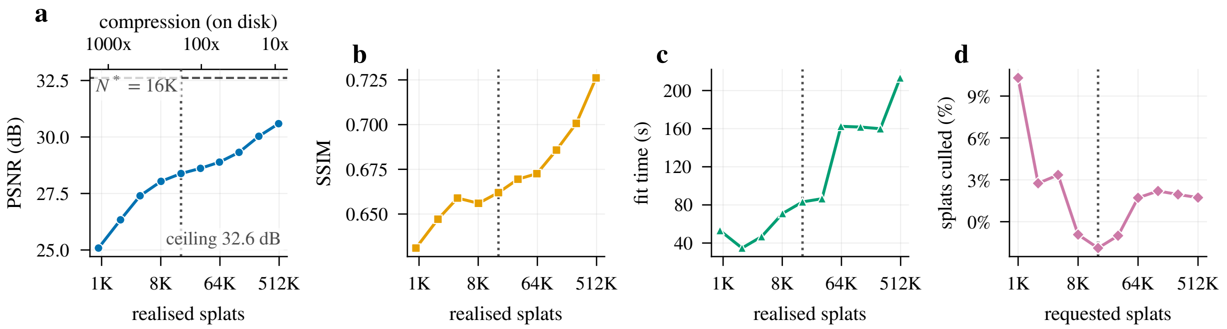


Figure 8: Rate-distortion curves, OpenCell-MAP4 nuclei. (a) PSNR rises from 25.1 dB at 1K to 30.6 dB at 512K against a noise ceiling of 32.6 dB; (b) SSIM reaches 0.66 at $N^* = 16\text{K}$ (dotted line in every panel); (c) fit time is wall-clock including seed deduplication (several minutes at 256K and 512K) and early stopping; (d) the realised count runs from 1.9% above the request (a seed pool within 5% of it is accepted as it is) to 10.3% below it.

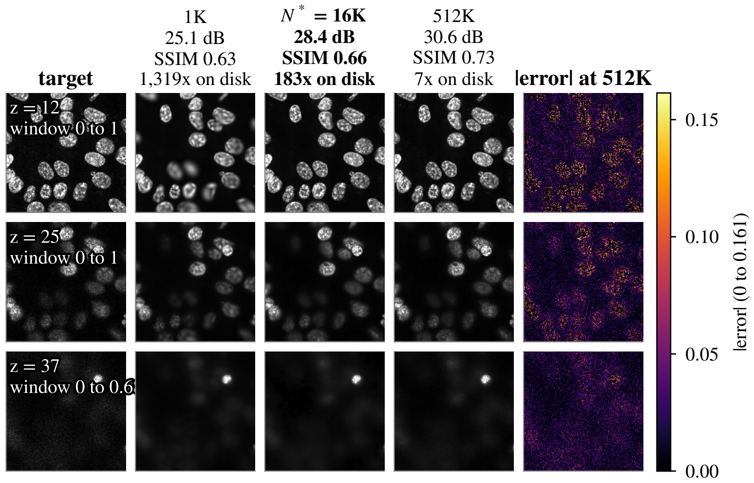


Figure 9: Reconstruction slices, OpenCell-MAP4 nuclei. Target and reconstructions of z planes 12, 25, 37, at 1K, $N^* = 16\text{K}$ and 512K (column titles: PSNR, SSIM, on-disk compression), and the absolute error at 512K on a 0 to 0.161 range; each plane's index and display window are printed on its target.

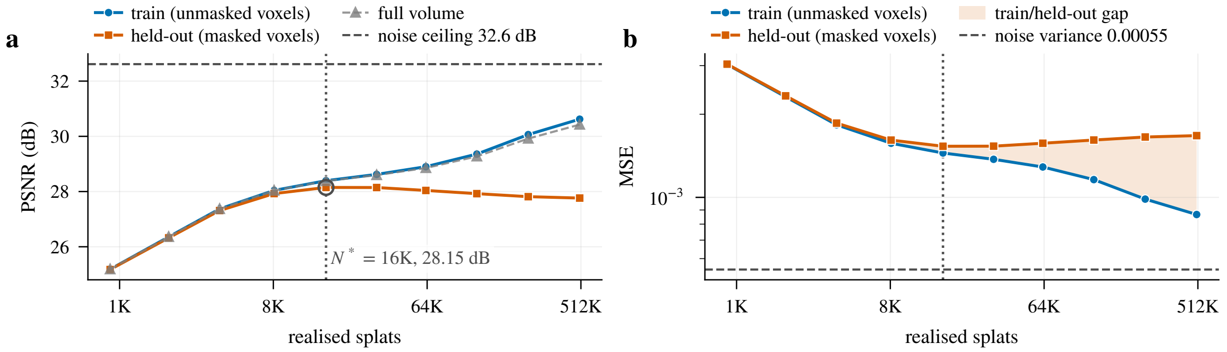


Figure 10: Blind-spot cross-validation, OpenCell-MAP4 nuclei. (a) The held-out PSNR peaks at $N^* = 16\text{K}$ (28.15 dB) and falls by 0.39 dB to 512K while the train PSNR keeps rising; (b) the train/held-out gap at 512K is 2.86 dB.

#### A.2 OpenCell-MAP4 microtubules: Spinning-disk

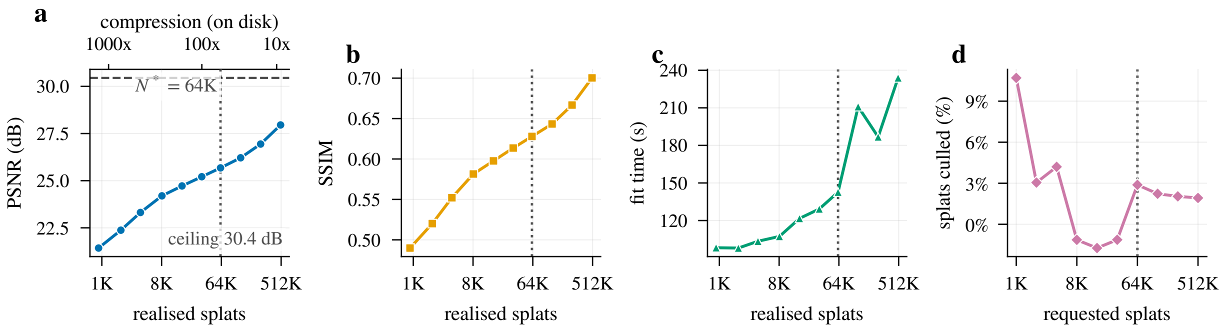


Figure 11: Rate-distortion curves, OpenCell-MAP4 microtubules. (a) PSNR rises from 21.4 dB at 1K to 28.0 dB at 512K against a noise ceiling of 30.4 dB; (b) SSIM reaches 0.63 at $N^* = 64K$ (dotted line in every panel); (c) fit time is wall-clock including seed deduplication (several minutes at 256K and 512K) and early stopping; (d) the realised count runs from 1.7% above the request (a seed pool within 5% of it is accepted as it is) to 10.7% below it.

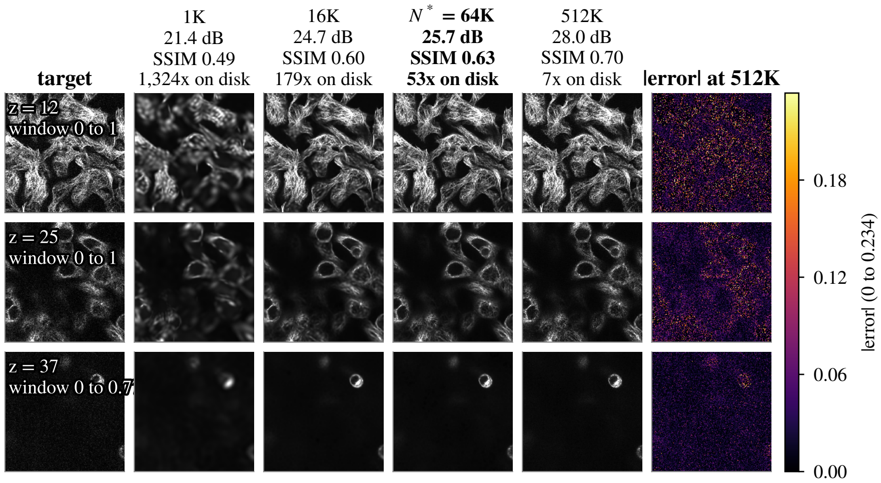


Figure 12: Reconstruction slices, OpenCell-MAP4 microtubules. Target and reconstructions of z planes 12, 25, 37, at 1K, 16K, $N^* = 64K$ and 512K (column titles: PSNR, SSIM, on-disk compression), and the absolute error at 512K on a 0 to 0.234 range; each plane's index and display window are printed on its target.

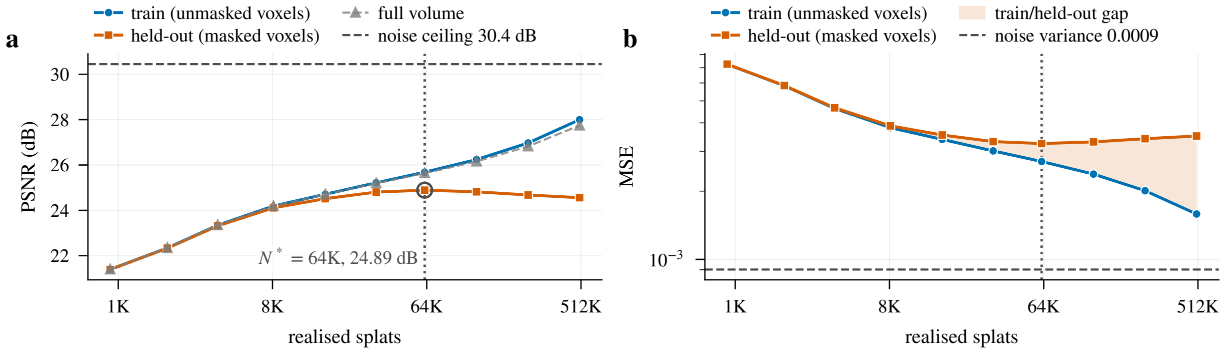


Figure 13: Blind-spot cross-validation, OpenCell-MAP4 microtubules. (a) The held-out PSNR peaks at $N^* = 64K$ (24.89 dB) and falls by 0.33 dB to 512K while the train PSNR keeps rising; (b) the train/held-out gap at 512K is 3.44 dB.

#### A.3 OpenCell-LMNB1 nuclei: Spinning-disk

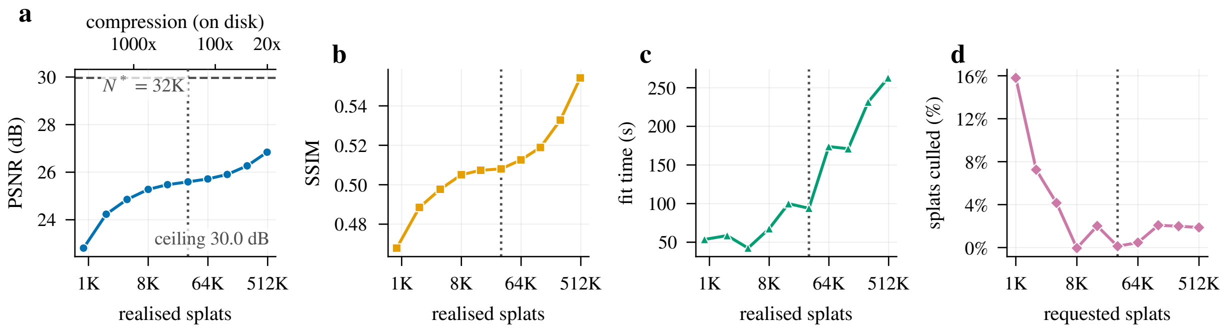


Figure 14: Rate-distortion curves, OpenCell-LMNB1 nuclei. (a) PSNR rises from 22.8 dB at 1K to 26.8 dB at 512K against a noise ceiling of 30.0 dB; (b) SSIM reaches 0.51 at $N^* = 32K$ (dotted line in every panel); (c) fit time is wall-clock including seed deduplication (several minutes at 256K and 512K) and early stopping; (d) the post-fit cull removes 0.0 to 15.8% of the requested splats.

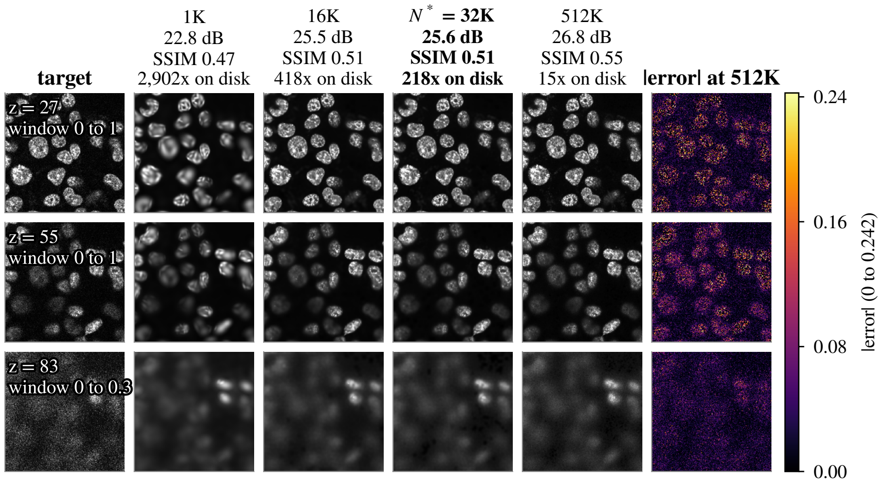


Figure 15: Reconstruction slices, OpenCell-LMNB1 nuclei. Target and reconstructions of z planes 27, 55, 83, at 1K, 16K, $N^* = 32K$ and 512K (column titles: PSNR, SSIM, on-disk compression), and the absolute error at 512K on a 0 to 0.242 range; each plane's index and display window are printed on its target.

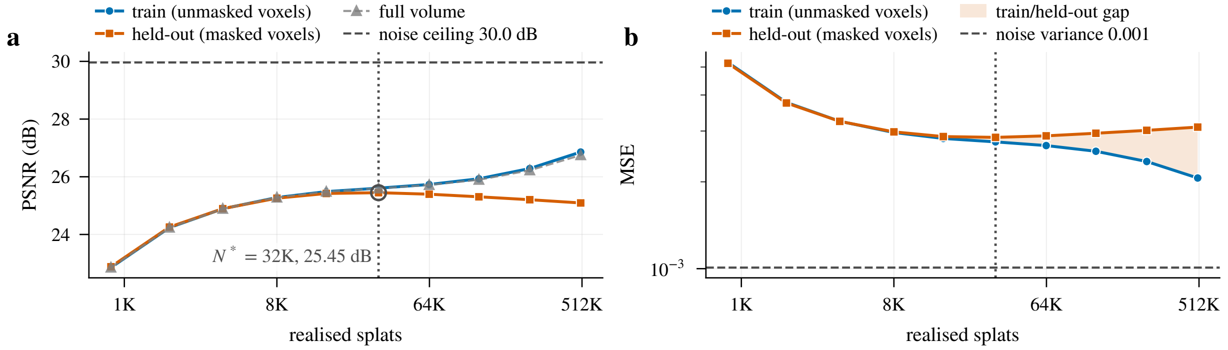


Figure 16: Blind-spot cross-validation, OpenCell-LMNB1 nuclei. (a) The held-out PSNR peaks at $N^* = 32K$ (25.45 dB) and falls by 0.36 dB to 512K while the train PSNR keeps rising; (b) the train/held-out gap at 512K is 1.76 dB.

#### A.4 OpenCell-LMNB1 nuclear lamina: Spinning-disk

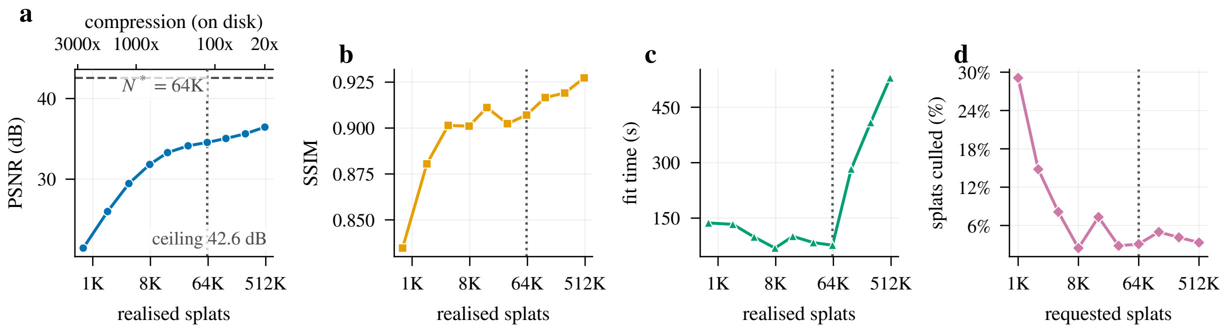


Figure 17: Rate-distortion curves, OpenCell-LMNB1 nuclear lamina. (a) PSNR rises from 21.5 dB at 1K to 36.5 dB at 512K against a noise ceiling of 42.6 dB; (b) SSIM reaches 0.91 at $N^* =$ 64K (dotted line in every panel); (c) fit time is wall-clock including seed deduplication (several minutes at 256K and 512K) and early stopping; (d) the post-fit cull removes 2.5 to 29.1% of the requested splats.

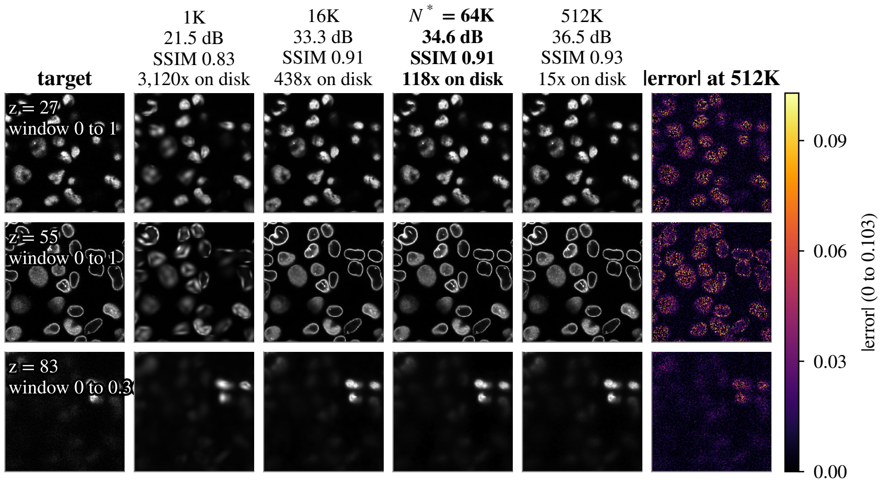


Figure 18: Reconstruction slices, OpenCell-LMNB1 nuclear lamina. Target and reconstructions of z planes 27, 55, 83, at 1K, 16K, $N^* =$ 64K and 512K (column titles: PSNR, SSIM, on-disk compression), and the absolute error at 512K on a 0 to 0.103 range; each plane's index and display window are printed on its target.

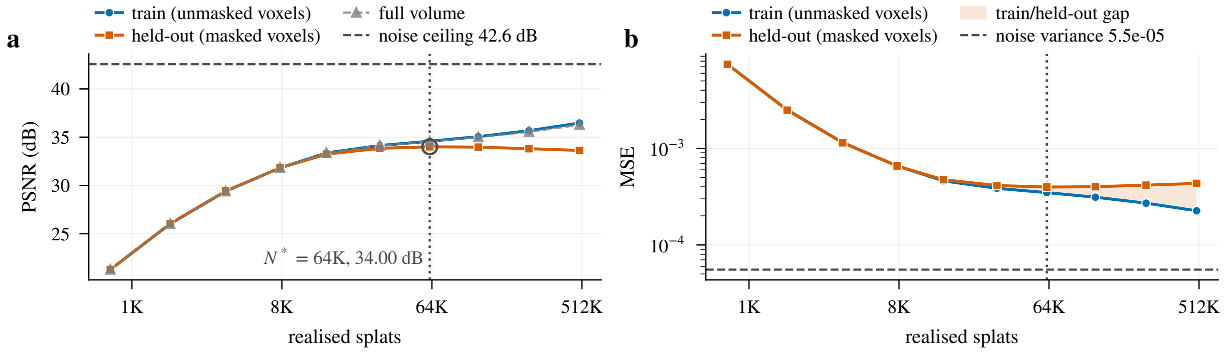


Figure 19: Blind-spot cross-validation, OpenCell-LMNB1 nuclear lamina. (a) The held-out PSNR peaks at $N^* =$ 64K (34.00 dB) and falls by 0.38 dB to 512K while the train PSNR keeps rising; (b) the train/held-out gap at 512K is 2.84 dB.

#### A.5 Kidney nuclei: Confocal

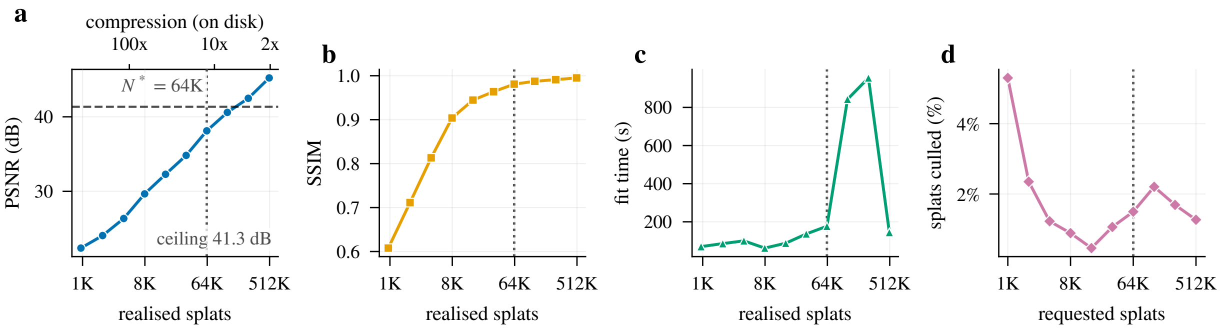


Figure 20: Rate-distortion curves, Kidney nuclei. (a) PSNR rises from 22.4 dB at 1K to 45.2 dB at 512K against a noise ceiling of 41.3 dB; (b) SSIM reaches 0.98 at $N^* = 64\text{K}$ (dotted line in every panel); (c) fit time is wall-clock including seed deduplication (several minutes at 256K and 512K) and early stopping; (d) the post-fit cull removes 0.5 to 5.3% of the requested splats.

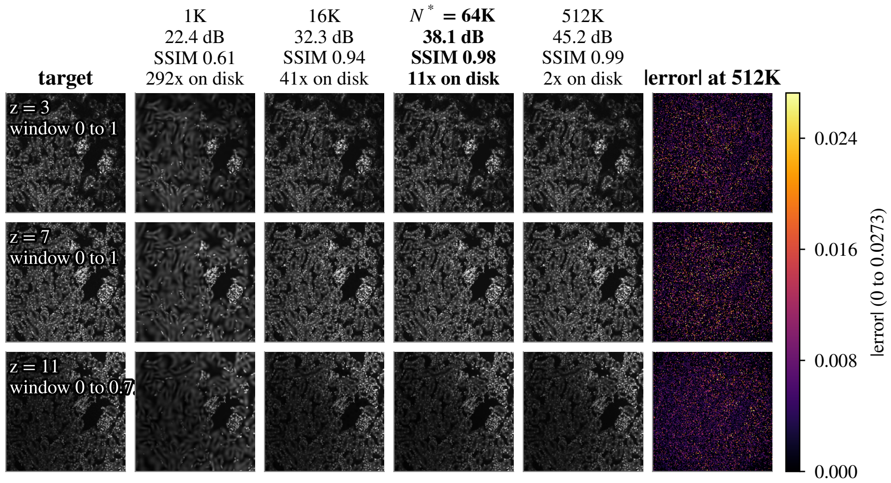


Figure 21: Reconstruction slices, Kidney nuclei. Target and reconstructions of z planes 3, 7, 11, at 1K, 16K, $N^* = 64\text{K}$ and 512K (column titles: PSNR, SSIM, on-disk compression), and the absolute error at 512K on a 0 to 0.0273 range; each plane's index and display window are printed on its target.

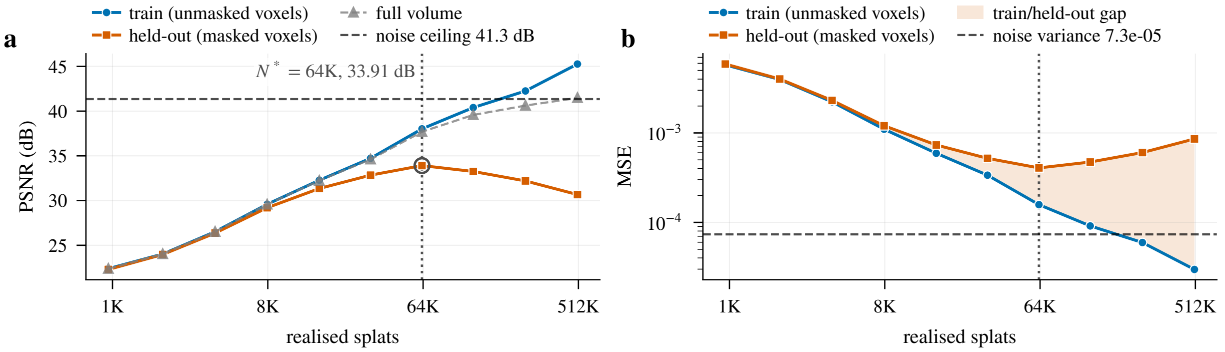


Figure 22: Blind-spot cross-validation, Kidney nuclei. (a) The held-out PSNR peaks at $N^* = 64\text{K}$ (33.91 dB) and falls by 3.24 dB to 512K while the train PSNR keeps rising; (b) the train/held-out gap at 512K is 14.59 dB.

#### A.6 Kidney actin: Confocal

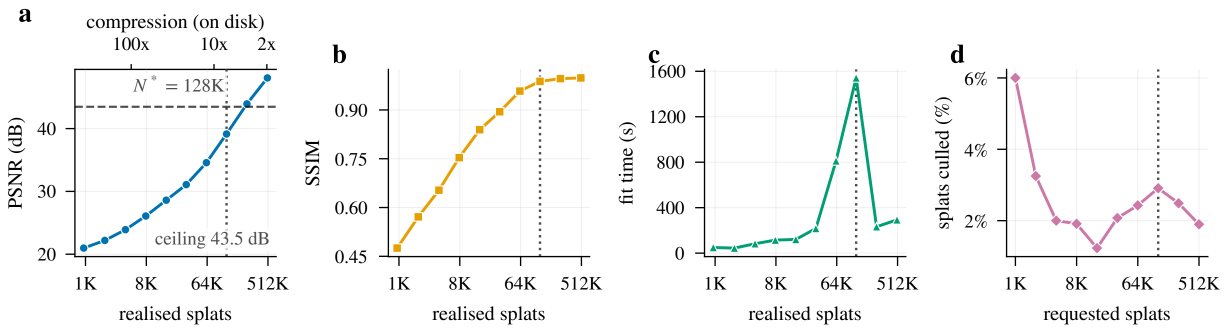


Figure 23: Rate-distortion curves, Kidney actin. (a) PSNR rises from 21.0 dB at 1K to 48.1 dB at 512K against a noise ceiling of 43.5 dB; (b) SSIM reaches 0.99 at $N^* = 128K$ (dotted line in every panel); (c) fit time is wall-clock including seed deduplication (several minutes at 256K and 512K) and early stopping; (d) the post-fit cull removes 1.2 to 6.0% of the requested splats.

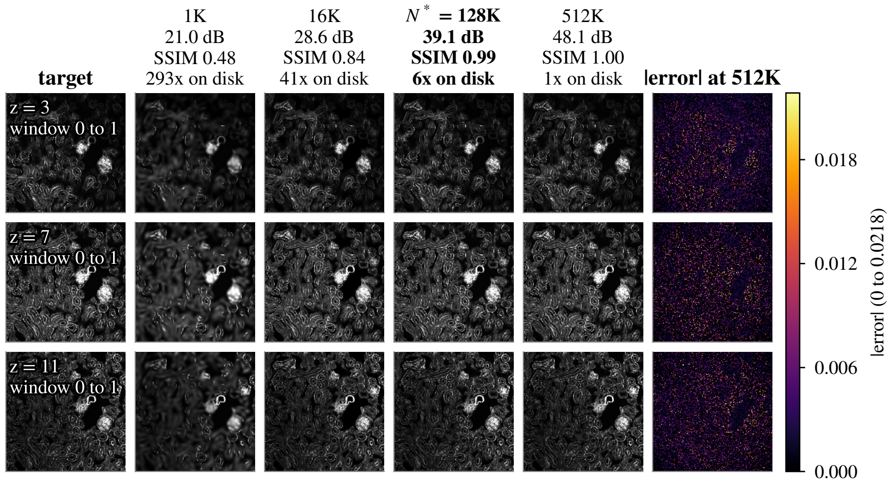


Figure 24: Reconstruction slices, Kidney actin. Target and reconstructions of z planes 3, 7, 11, at 1K, 16K, $N^* = 128K$ and 512K (column titles: PSNR, SSIM, on-disk compression), and the absolute error at 512K on a 0 to 0.0218 range; each plane's index and display window are printed on its target.

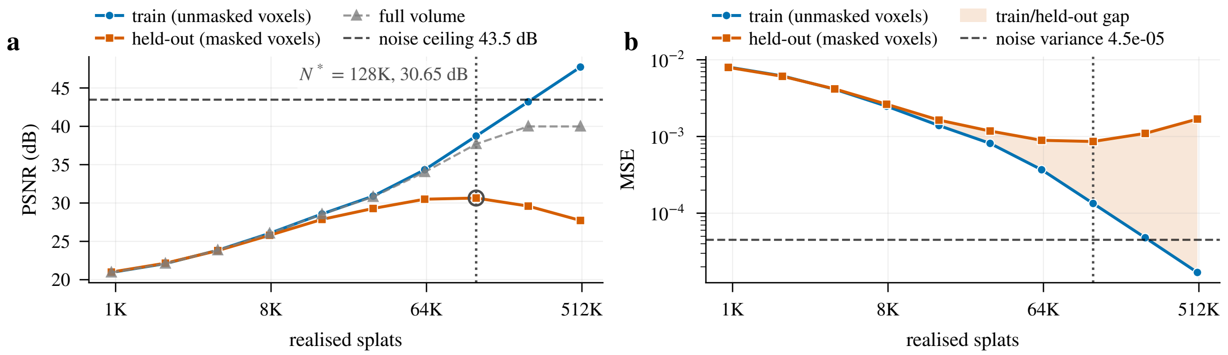


Figure 25: Blind-spot cross-validation, Kidney actin. (a) The held-out PSNR peaks at $N^* = 128K$ (30.65 dB) and falls by 2.92 dB to 512K while the train PSNR keeps rising; (b) the train/held-out gap at 512K is 20.00 dB.

#### A.7 Cells3D nuclei: Confocal

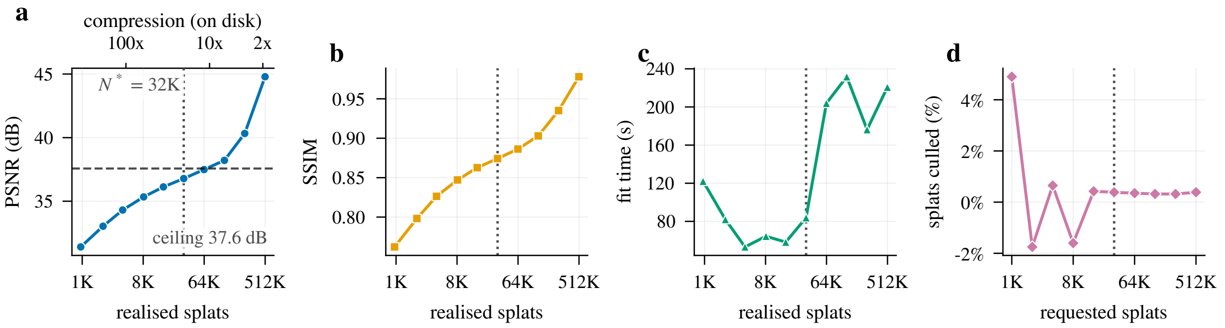


Figure 26: Rate-distortion curves, Cells3D nuclei. (a) PSNR rises from 31.4 dB at 1K to 44.8 dB at 512K against a noise ceiling of 37.6 dB; (b) SSIM reaches 0.87 at $N^* = 32$K (dotted line in every panel); (c) fit time is wall-clock including seed deduplication (several minutes at 256K and 512K) and early stopping; (d) the realised count runs from 1.8% above the request (a seed pool within 5% of it is accepted as it is) to 4.9% below it.

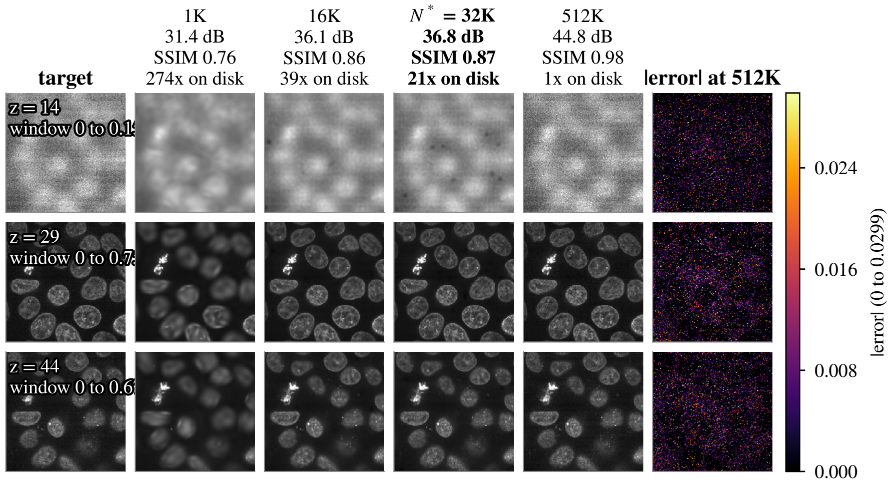


Figure 27: Reconstruction slices, Cells3D nuclei. Target and reconstructions of z planes 14, 29, 44, at 1K, 16K, $N^* = 32$K and 512K (column titles: PSNR, SSIM, on-disk compression), and the absolute error at 512K on a 0 to 0.0299 range; each plane's index and display window are printed on its target.

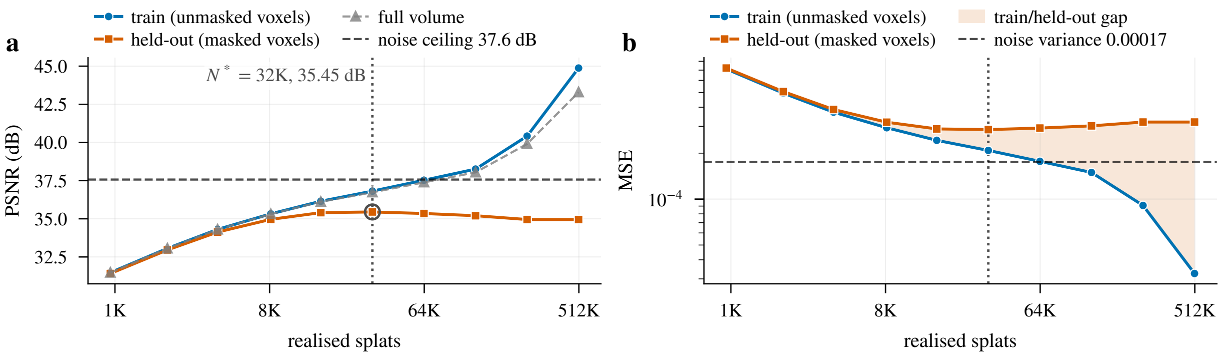


Figure 28: Blind-spot cross-validation, Cells3D nuclei. (a) The held-out PSNR peaks at $N^* = 32$K (35.45 dB) and falls by 0.50 dB to 512K while the train PSNR keeps rising; (b) the train/held-out gap at 512K is 9.92 dB.

#### A.8 Cells3D membranes: Confocal

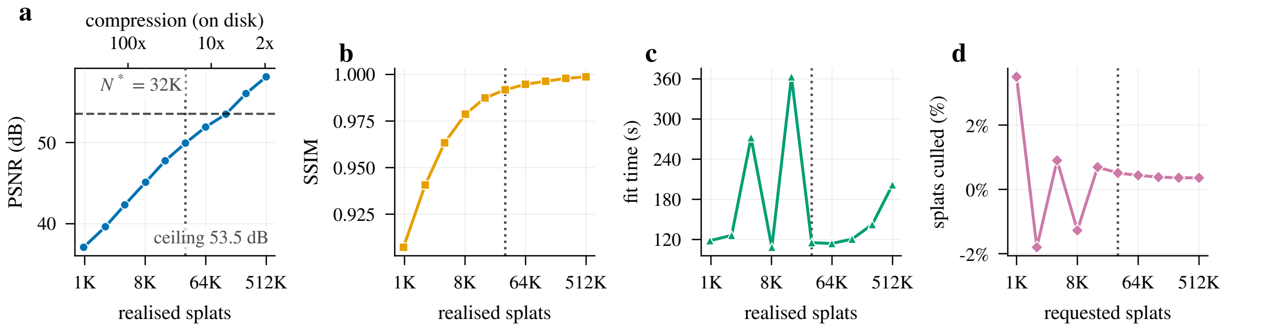


Figure 29: Rate-distortion curves, Cells3D membranes. (a) PSNR rises from 37.1 dB at 1K to 58.1 dB at 512K against a noise ceiling of 53.5 dB; (b) SSIM reaches 0.99 at $N^* = 32K$ (dotted line in every panel); (c) fit time is wall-clock including seed deduplication (several minutes at 256K and 512K) and early stopping; (d) the realised count runs from 1.8% above the request (a seed pool within 5% of it is accepted as it is) to 3.5% below it.

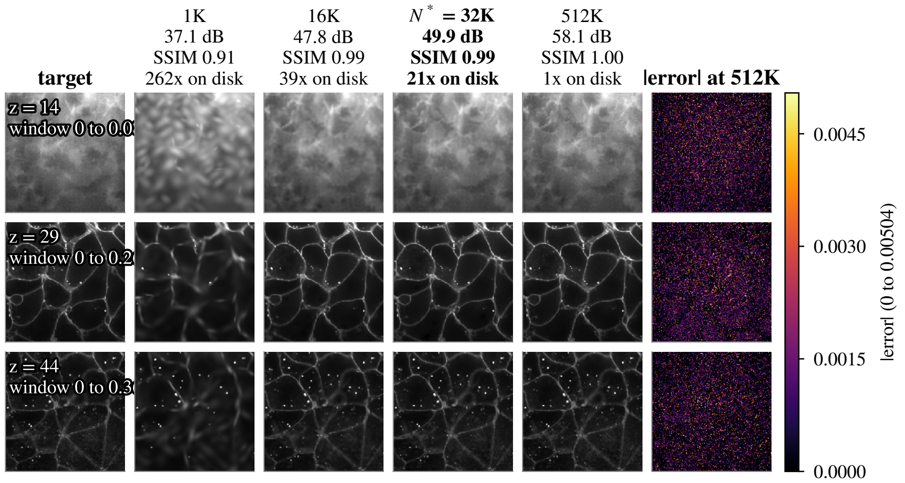


Figure 30: Reconstruction slices, Cells3D membranes. Target and reconstructions of z planes 14, 29, 44, at 1K, 16K, $N^* = 32K$ and 512K (column titles: PSNR, SSIM, on-disk compression), and the absolute error at 512K on a 0 to 0.00504 range; each plane's index and display window are printed on its target.

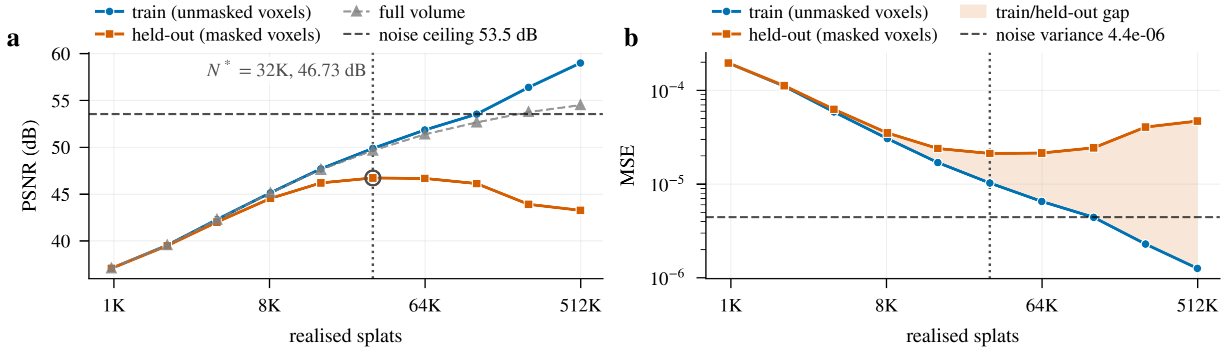


Figure 31: Blind-spot cross-validation, Cells3D membranes. (a) The held-out PSNR peaks at $N^* = 32K$ (46.73 dB) and falls by 3.46 dB to 512K while the train PSNR keeps rising; (b) the train/held-out gap at 512K is 15.73 dB.

#### A.9 Blastocyst nuclear lamina: Confocal

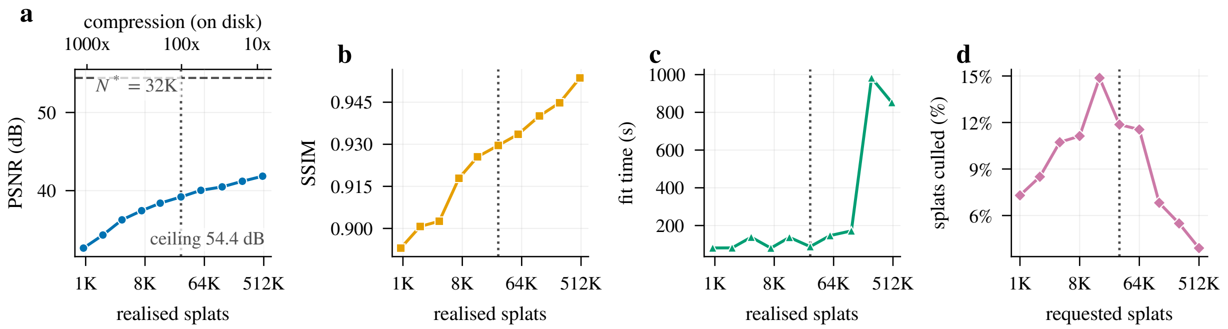


Figure 32: Rate-distortion curves, Blastocyst nuclear lamina. (a) PSNR rises from 32.6 dB at 1K to 41.8 dB at 512K against a noise ceiling of 54.4 dB; (b) SSIM reaches 0.93 at $N^* = 32K$ (dotted line in every panel); (c) fit time is wall-clock including seed deduplication (several minutes at 256K and 512K) and early stopping; (d) the post-fit cull removes 3.9 to 14.9% of the requested splats.

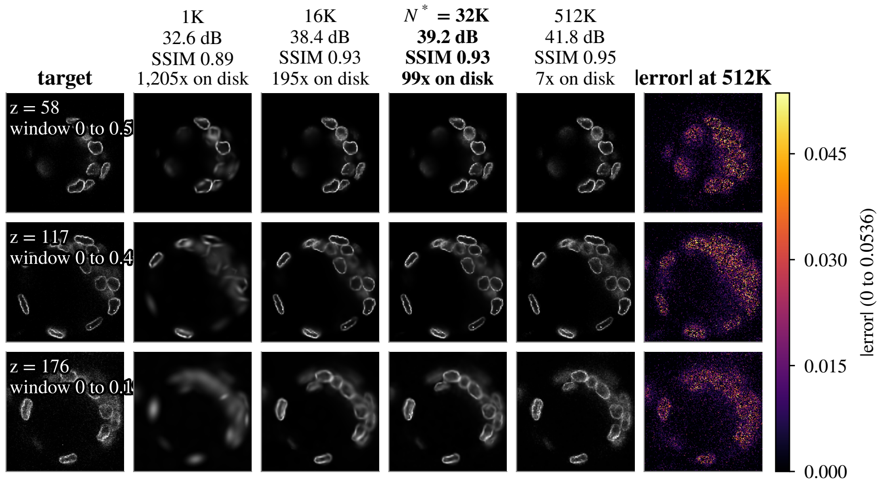


Figure 33: Reconstruction slices, Blastocyst nuclear lamina. Target and reconstructions of z planes 58, 117, 176, at 1K, 16K, $N^* = 32K$ and 512K (column titles: PSNR, SSIM, on-disk compression), and the absolute error at 512K on a 0 to 0.0536 range; each plane's index and display window are printed on its target.

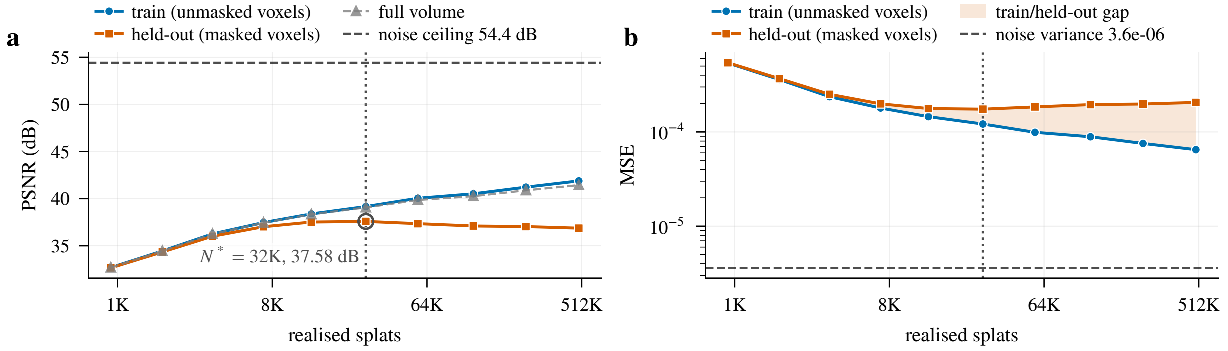


Figure 34: Blind-spot cross-validation, Blastocyst nuclear lamina. (a) The held-out PSNR peaks at $N^* = 32K$ (37.58 dB) and falls by 0.71 dB to 512K while the train PSNR keeps rising; (b) the train/held-out gap at 512K is 5.01 dB.

#### A.10 *C. elegans* embryo nuclei: Confocal

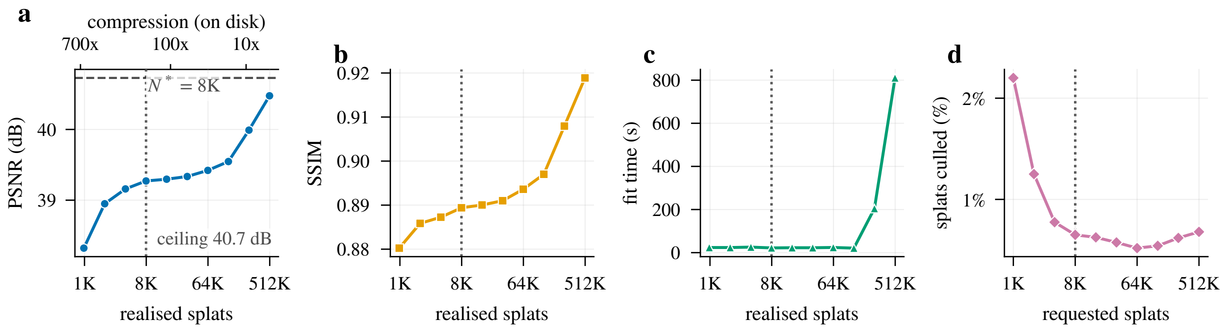


Figure 35: Rate-distortion curves, *C. elegans* embryo nuclei. (a) PSNR rises from 38.3 dB at 1K to 40.5 dB at 512K against a noise ceiling of 40.7 dB; (b) SSIM reaches 0.89 at $N^* = 8\text{K}$ (dotted line in every panel); (c) fit time is wall-clock including seed deduplication (several minutes at 256K and 512K) and early stopping; (d) the post-fit cull removes 0.5 to 2.2% of the requested splats.

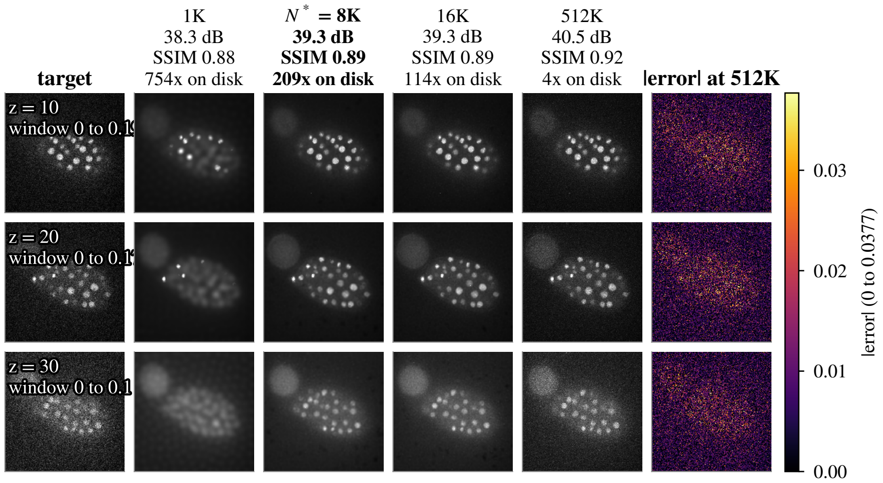


Figure 36: Reconstruction slices, *C. elegans* embryo nuclei. Target and reconstructions of z planes 10, 20, 30, at 1K, $N^* = 8\text{K}$, 16K and 512K (column titles: PSNR, SSIM, on-disk compression), and the absolute error at 512K on a 0 to 0.0377 range; each plane's index and display window are printed on its target.

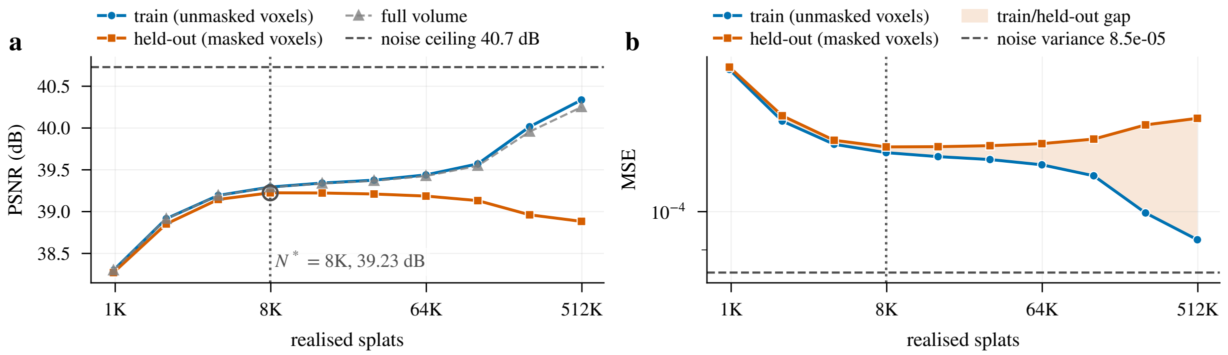


Figure 37: Blind-spot cross-validation, *C. elegans* embryo nuclei. (a) The held-out PSNR peaks at $N^* = 8\text{K}$ (39.23 dB) and falls by 0.34 dB to 512K while the train PSNR keeps rising; (b) the train/held-out gap at 512K is 1.45 dB.

#### A.11 Fly brain neurons: Confocal

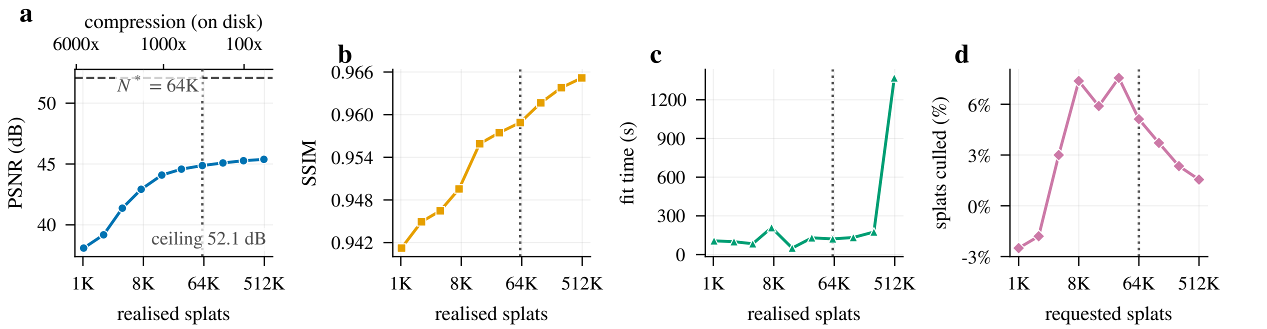


Figure 38: Rate-distortion curves, Fly brain neurons. (a) PSNR rises from 38.1 dB at 1K to 45.4 dB at 512K against a noise ceiling of 52.1 dB; (b) SSIM reaches 0.96 at $N^* = 64$K (dotted line in every panel); (c) fit time is wall-clock including seed deduplication (several minutes at 256K and 512K) and early stopping; (d) the realised count runs from 2.5% above the request (a seed pool within 5% of it is accepted as it is) to 7.6% below it.

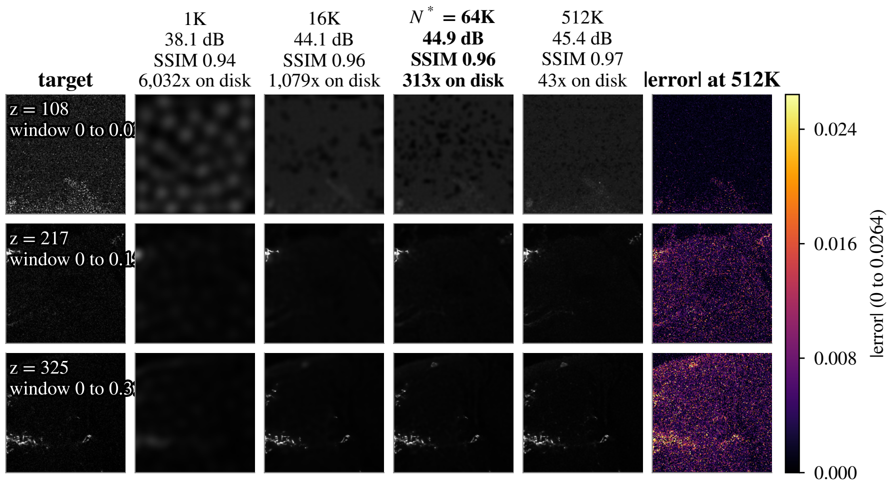


Figure 39: Reconstruction slices, Fly brain neurons. Target and reconstructions of z planes 108, 217, 325, at 1K, 16K, $N^* = 64$K and 512K (column titles: PSNR, SSIM, on-disk compression), and the absolute error at 512K on a 0 to 0.0264 range; each plane's index and display window are printed on its target.

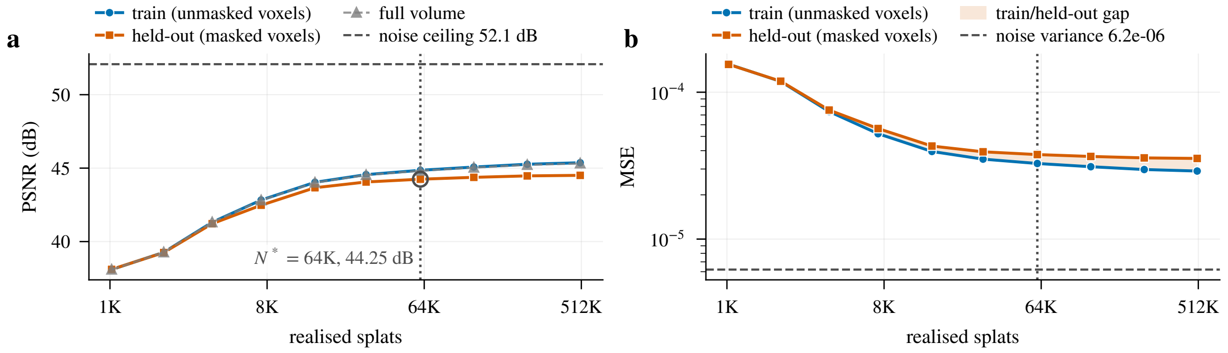


Figure 40: Blind-spot cross-validation, Fly brain neurons. (a) The held-out PSNR is within 0.3 dB of its 44.51 dB maximum from $N^* = 64$K (plateau onset) and changes by +0.00 dB between the maximum and 512K; (b) the train/held-out gap at 512K is 0.86 dB.

#### A.12 *Tribolium* embryo nuclei: Light-sheet (deconv.)

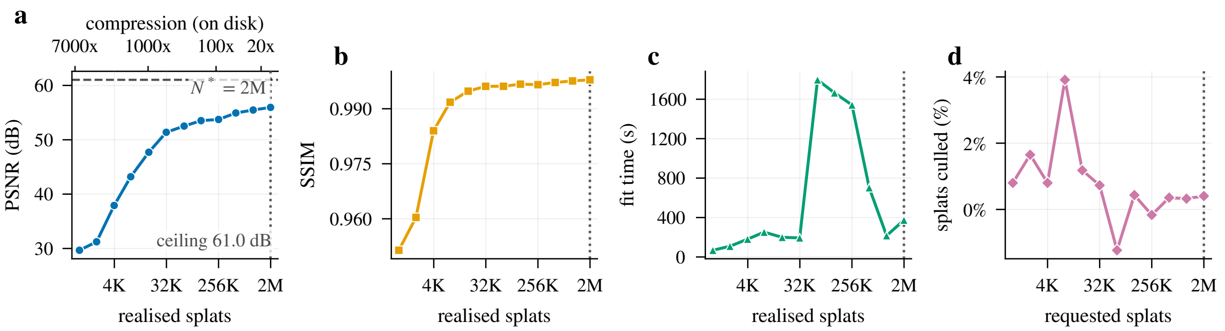


Figure 41: Rate-distortion curves, *Tribolium* embryo nuclei. (a) PSNR rises from 29.7 dB at 1K to 56.0 dB at 2M against a noise ceiling of 61.0 dB; (b) SSIM reaches 1.00 at $N^* = 2\text{M}$ (dotted line in every panel); (c) fit time is wall-clock including seed deduplication (several minutes at 256K and 512K) and early stopping; (d) the realised count runs from 1.2% above the request (a seed pool within 5% of it is accepted as it is) to 3.9% below it.

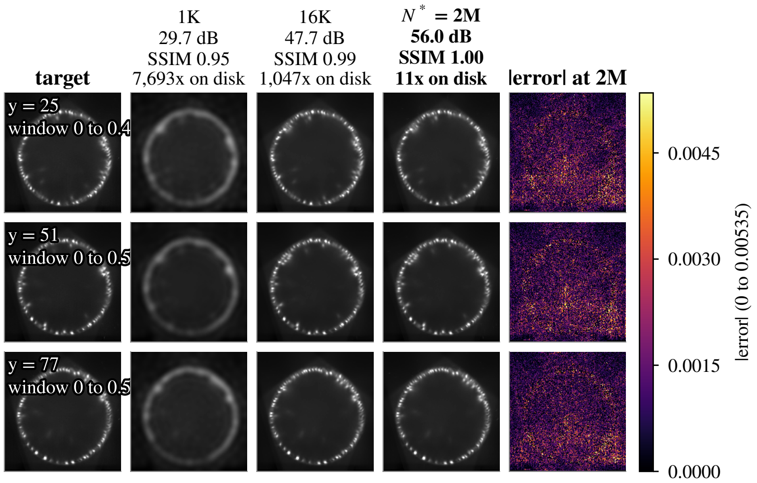


Figure 42: Reconstruction slices, *Tribolium* embryo nuclei. Target and reconstructions of y planes 25, 51, 77, cut along $y$, the volume's shortest axis, at 1K, 16K and $N^* = 2\text{M}$ (column titles: PSNR, SSIM, on-disk compression), and the absolute error at 2M on a 0 to 0.00535 range; each plane's index and display window are printed on its target.

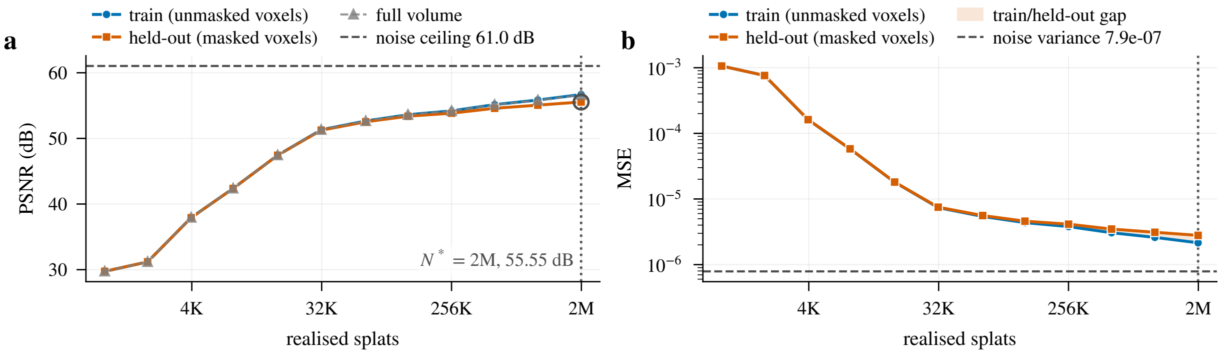


Figure 43: Blind-spot cross-validation, *Tribolium* embryo nuclei. (a) The held-out PSNR is still rising at the largest tested count, 2M (55.55 dB, +0.48 dB over the last doubling), so the operating point is the endpoint $N^* = 2\text{M}$; (b) the train/held-out gap at 512K is 0.56 dB.

#### A.13 Mouse heart nuclei: Light-sheet

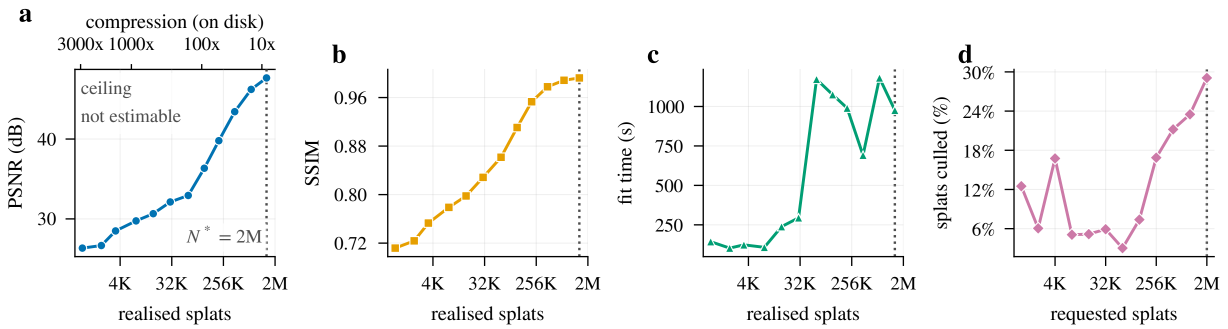


Figure 44: Rate-distortion curves, Mouse heart nuclei. (a) PSNR rises from 26.3 dB at 1K to 47.7 dB at 2M with no estimable noise ceiling; (b) SSIM reaches 0.99 at $N^* = 2M$ (dotted line in every panel); (c) fit time is wall-clock including seed deduplication (several minutes at 256K and 512K) and early stopping; (d) the post-fit cull removes 3.0 to 29.1% of the requested splats.

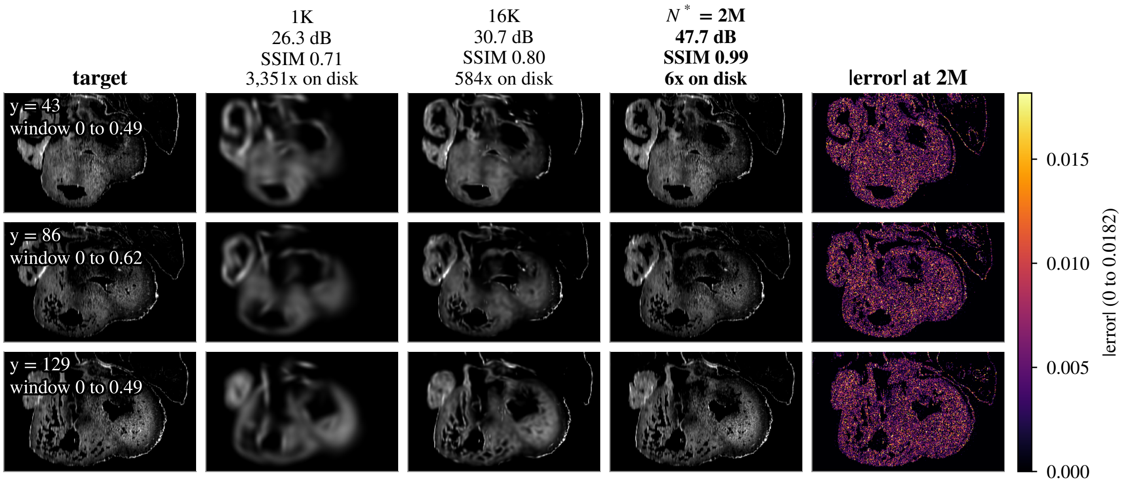


Figure 45: Reconstruction slices, Mouse heart nuclei. Target and reconstructions of y planes 43, 86, 129, cut along $y$, the volume's shortest axis, at 1K, 16K and $N^* = 2M$ (column titles: PSNR, SSIM, on-disk compression), and the absolute error at 2M on a 0 to 0.0182 range; each plane's index and display window are printed on its target.

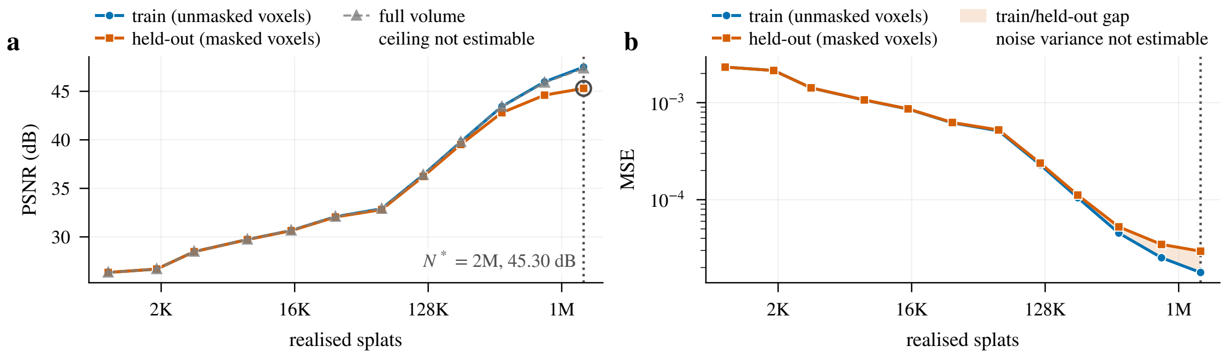


Figure 46: Blind-spot cross-validation, Mouse heart nuclei. (a) The held-out PSNR is still rising at the largest tested count, 2M (45.30 dB, +0.70 dB over the last doubling), so the operating point is the endpoint $N^* = 2M$; (b) the train/held-out gap at 512K is 0.64 dB.

#### A.14 Zebrafish embryo nuclei: Light-sheet (deconv.)

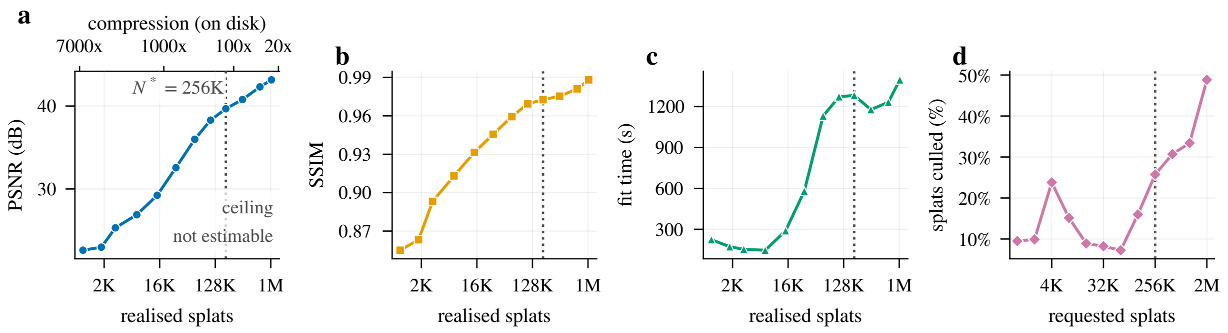


Figure 47: Rate-distortion curves, Zebrafish embryo nuclei. (a) PSNR rises from 22.6 dB at 1K to 43.2 dB at 2M with no estimable noise ceiling; (b) SSIM reaches 0.97 at $N^* = 256K$ (dotted line in every panel); (c) fit time is wall-clock including seed deduplication (several minutes at 256K and 512K) and early stopping; (d) the post-fit cull removes 7.3 to 48.8% of the requested splats.

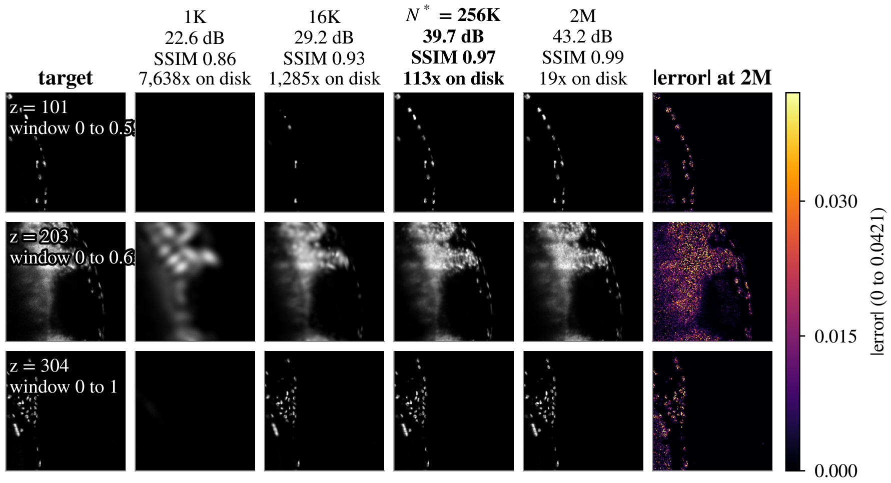


Figure 48: Reconstruction slices, Zebrafish embryo nuclei. Target and reconstructions of z planes 101, 203, 304, at 1K, 16K, $N^* = 256K$ and 2M (column titles: PSNR, SSIM, on-disk compression), and the absolute error at 2M on a 0 to 0.0421 range; each plane's index and display window are printed on its target.

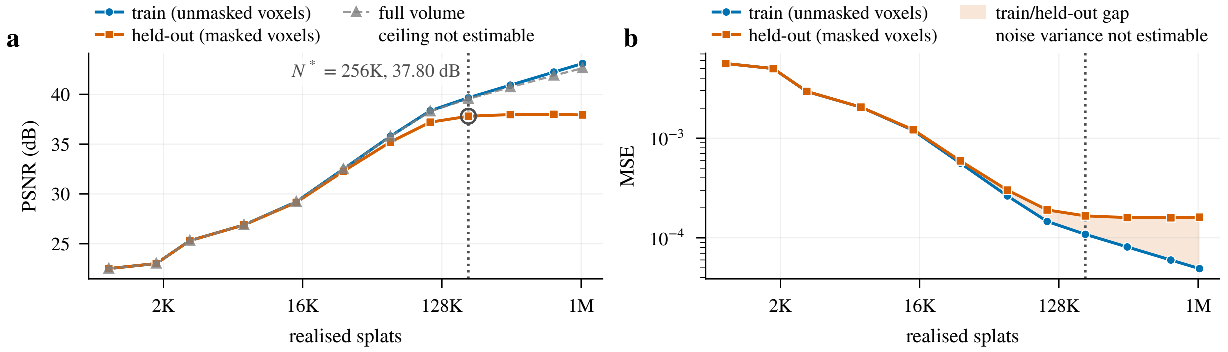


Figure 49: Blind-spot cross-validation, Zebrafish embryo nuclei. (a) The held-out PSNR is within 0.3 dB of its 37.99 dB maximum from $N^* = 256K$ (plateau onset) and changes by -0.06 dB between the maximum and 2M; (b) the train/held-out gap at 512K is 2.95 dB.

#### A.15 *Drosophila* embryo nuclei: Light-sheet

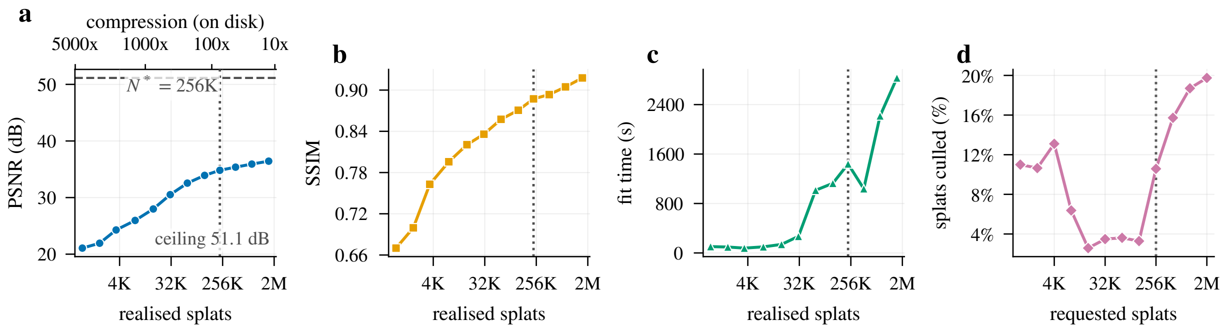


Figure 50: Rate-distortion curves, *Drosophila* embryo nuclei. (a) PSNR rises from 21.1 dB at 1K to 36.4 dB at 2M against a noise ceiling of 51.1 dB; (b) SSIM reaches 0.89 at $N^* = 256K$ (dotted line in every panel); (c) fit time is wall-clock including seed deduplication (several minutes at 256K and 512K) and early stopping; (d) the post-fit cull removes 2.6 to 19.7% of the requested splats.

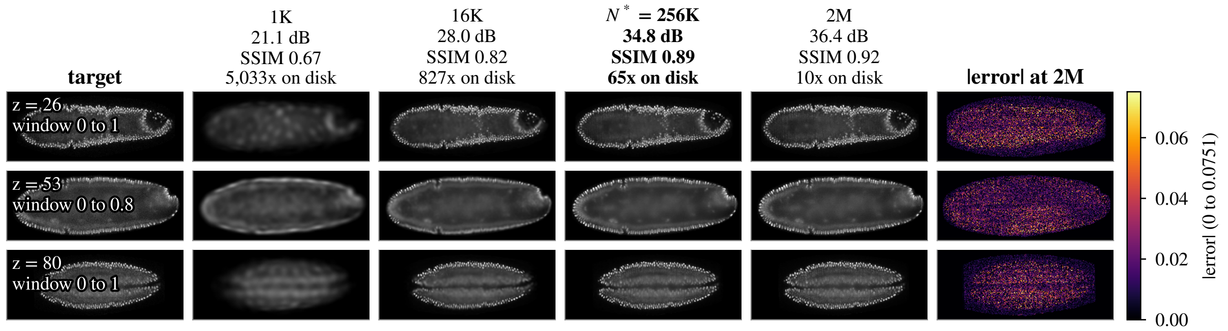


Figure 51: Reconstruction slices, *Drosophila* embryo nuclei. Target and reconstructions of z planes 26, 53, 80 (shown transposed), at 1K, 16K, $N^* = 256K$ and 2M (column titles: PSNR, SSIM, on-disk compression), and the absolute error at 2M on a 0 to 0.0751 range; each plane's index and display window are printed on its target.

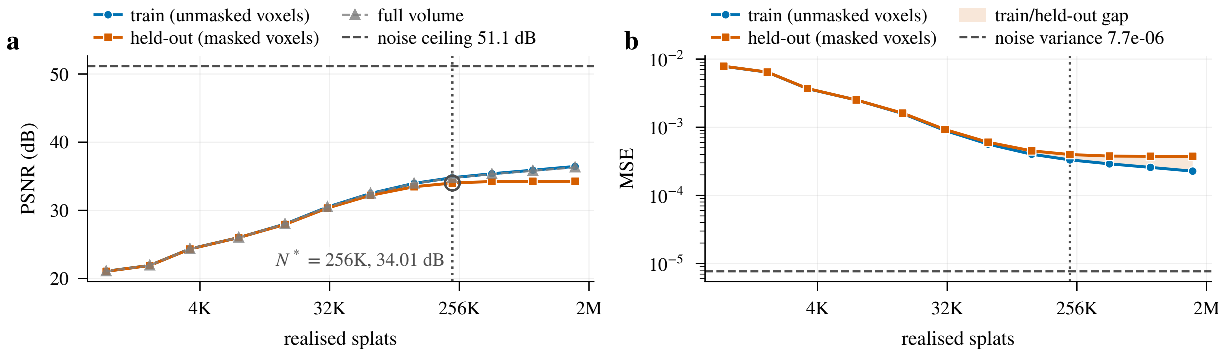


Figure 52: Blind-spot cross-validation, *Drosophila* embryo nuclei. (a) The held-out PSNR is within 0.3 dB of its 34.27 dB maximum from $N^* = 256K$ (plateau onset) and changes by +0.00 dB between the maximum and 2M; (b) the train/held-out gap at 512K is 1.14 dB.

#### A.16 Neuromast membranes: iSIM (deconv.)

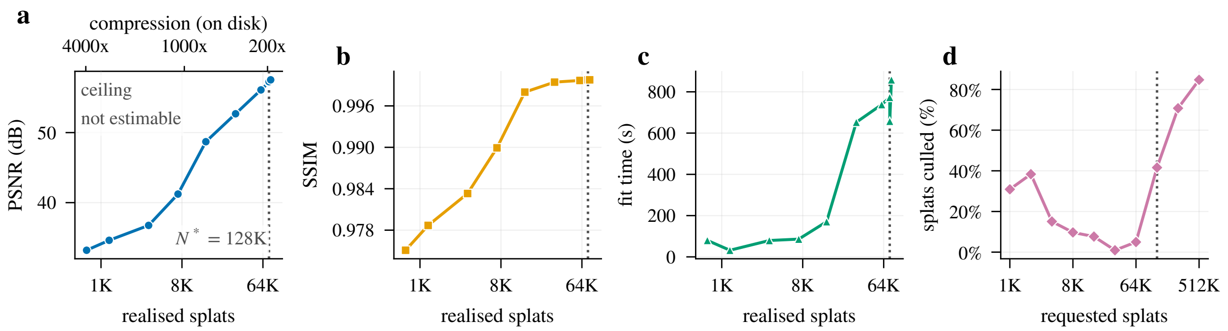


Figure 53: Rate-distortion curves, Neuromast membranes. (a) PSNR rises from 33.2 dB at 1K to 57.5 dB at 512K with no estimable noise ceiling; (b) SSIM reaches 1.00 at $N^* = 128\text{K}$ (dotted line in every panel); (c) fit time is wall-clock including seed deduplication (several minutes at 256K and 512K) and early stopping; (d) the post-fit cull removes 0.9 to 84.8% of the requested splats.

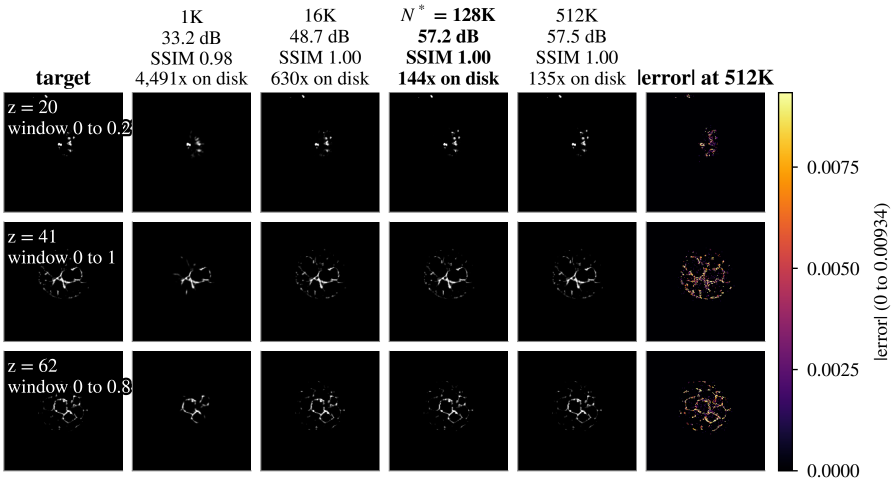


Figure 54: Reconstruction slices, Neuromast membranes. Target and reconstructions of z planes 20, 41, 62, at 1K, 16K, $N^* = 128\text{K}$ and 512K (column titles: PSNR, SSIM, on-disk compression), and the absolute error at 512K on a 0 to 0.00934 range; each plane's index and display window are printed on its target.

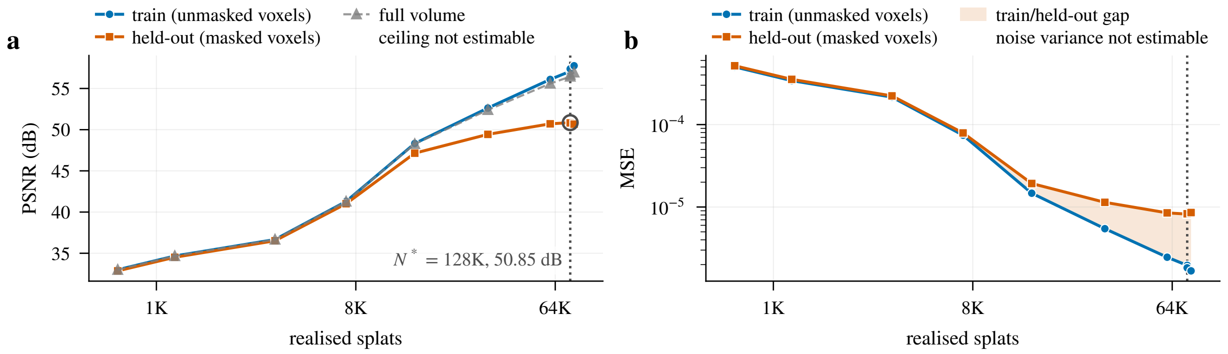


Figure 55: Blind-spot cross-validation, Neuromast membranes. (a) The held-out maximum is at $N^* = 128\text{K}$ (50.85 dB) and the decline beyond it, 0.19 dB, is between fits whose realised count stays between 74,793 and 77,900 splats (cull-limited); (b) the train/held-out gap at 512K is 7.09 dB.

#### A.17 Neuromast nuclei: iSIM (deconv.)

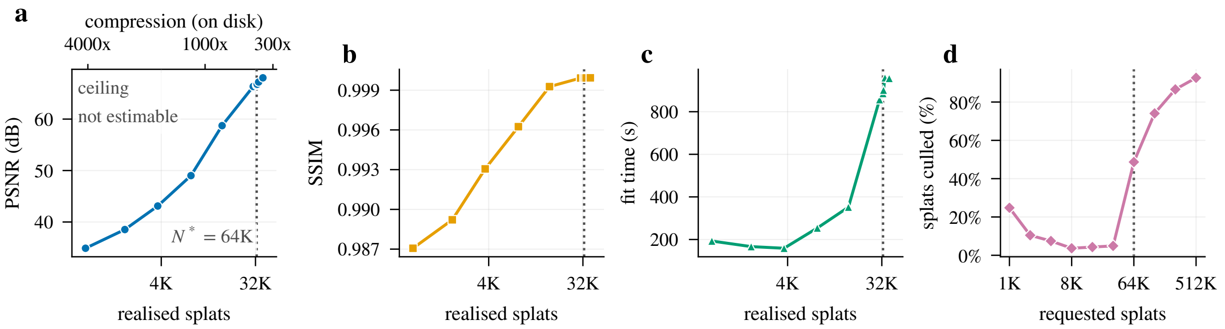


Figure 56: Rate-distortion curves, Neuromast nuclei. (a) PSNR rises from 34.9 dB at 1K to 68.0 dB at 512K with no estimable noise ceiling; (b) SSIM reaches 1.00 at $N^*$ = 64K (dotted line in every panel); (c) fit time is wall-clock including seed deduplication (several minutes at 256K and 512K) and early stopping; (d) the post-fit cull removes 3.7 to 92.6% of the requested splats.

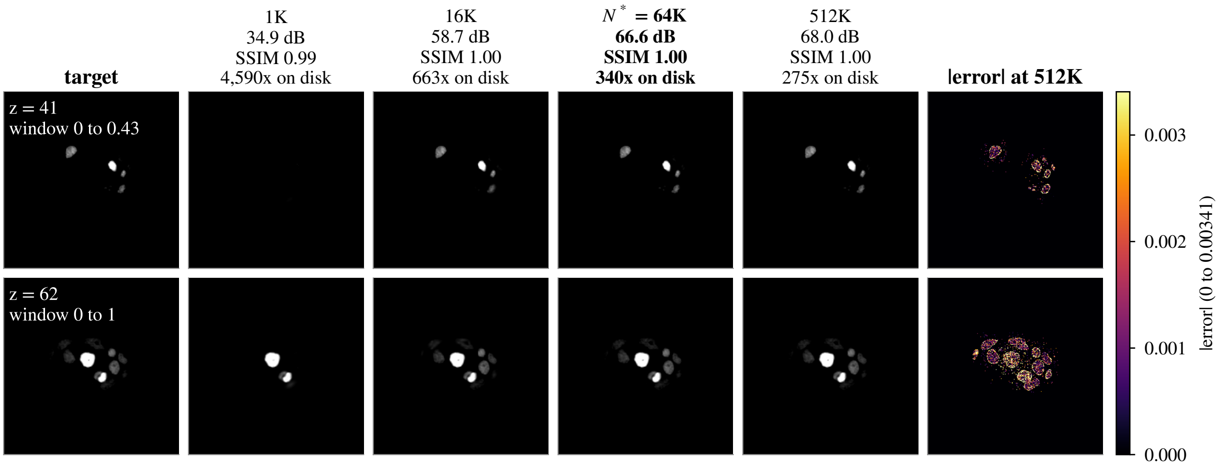


Figure 57: Reconstruction slices, Neuromast nuclei. Target and reconstructions of z planes 41, 62, at 1K, 16K, $N^*$ = 64K and 512K (column titles: PSNR, SSIM, on-disk compression), and the absolute error at 512K on a 0 to 0.00341 range; each plane's index and display window are printed on its target.

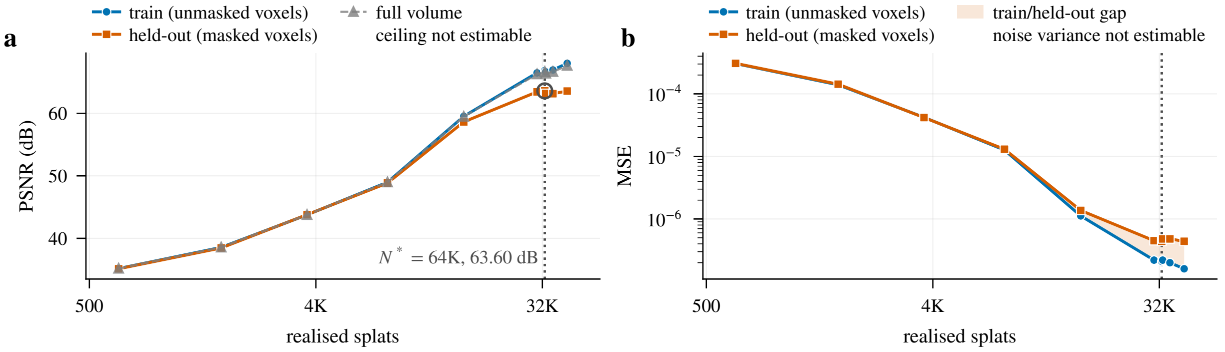


Figure 58: Blind-spot cross-validation, Neuromast nuclei. (a) The held-out maximum is at $N^*$ = 64K (63.60 dB) and the decline beyond it, 0.03 dB, is between fits whose realised count stays between 32,715 and 40,191 splats (cull-limited); (b) the train/held-out gap at 512K is 4.43 dB.
