## Supplementary material for "Luxar: Gaussian splatting for microscopy and scalable interactive web visualisation of multidimensional scientific data": supp_doc_03_progressive_vs_single

### Progressive vs. single-pass Gaussian-splat fitting at a nominal fixed budget

Supplementary Document 3 – Luxar

**Abstract**

Progressive residual fitting splits a Gaussian-splat budget into passes, each fitting new splats to what the earlier passes left unexplained, as an alternative to one joint fit. We test whether it helps at a nominal budget of 32,000 seeds and 10,000 iterations on 17 microscopy volumes, comparing one joint pass with 2, 4 and 8 residual passes, one fit per volume and pass count. The single pass reaches the higher training PSNR on 15 of the 17 volumes. The 2-pass minus 1-pass margin, the convention of every table here, runs on those volumes from $-6.02$ to $-0.07$ dB (median $-0.67$ dB); 2-pass leads only on Mouse heart nuclei (+0.40 dB) and Zebrafish embryo nuclei (+0.85 dB). Four and eight passes trail the single pass on every volume, by up to 13.7 and 19.8 dB. The single pass also has the lowest maximum absolute error on 12 of 17 volumes, while 2-pass has the higher SSIM on 12 of 17 by small margins. Wall time on an idle GPU is comparable (1-pass/2-pass ratio 0.70× to 1.75×, median 1.07×). On the 6 volumes whose margin is below 0.5 dB, 2 further RNG seeds per condition give within-condition PSNR spreads of 0.002 to 0.093 dB, and the 1-pass and 2-pass ranges are disjoint on 6 of the 6. Initialisation is deterministic in this fitter, so that spread samples GPU nondeterminism and the residual-seeder draw only. For the volumes and budget tested, a single joint fit is the recommended default.

#### 1 Introduction

A Gaussian-splat fit approximates a volume $V$ by a sum of anisotropic Gaussians (splats) whose centres, amplitudes and covariances are optimised jointly against a training loss. Progressive residual fitting spends the same budget in stages. A first pass fits a fraction of the splats to $V$; every later pass fits new splats to the positive residual that the accumulated reconstruction leaves behind, while the earlier splats stay frozen. The case for it is that a residual pass places capacity exactly where the reconstruction is still poor, that each pass optimises a smaller problem, and that seeding from the residual reaches faint structure a whole-volume seeder skips. The case against is that a staged fit can never revise its earlier passes and gives each stage fewer splats and fewer iterations. Which effect dominates at a fixed budget is an empirical question.

This document answers it for the shipped fitter with three of its defaults overridden (§ 2.1) and every stopping rule disabled (§ 2.3): one joint pass against 2, 4 and 8 residual passes at a common nominal budget on seventeen microscopy volumes, scored by training *PSNR*, *SSIM*, maximum absolute error and wall time. The progressive fitter also changes the loss, the seeding and the learning rate on its residual passes, so the comparison is between two complete procedures at a common budget and does not isolate the effect of staging alone.

##### Terms used in this document

**PSNR** The peak signal-to-noise ratio, $10\log_{10}(\text{peak}^2/\text{MSE})$ in decibels, where peak is the reference range of the values compared; unless a document states another range, the supplements take peak 1 on volumes normalised to $[0, 1]$, so that $\text{PSNR} = -10\log_{10}\text{MSE}$.

**SSIM** The structural similarity index, the mean over local windows of a product of luminance, contrast and structure comparisons between two images, lying in $[-1, 1]$ with 1 meaning identical.

**cull (retention)** The removal of splats after a fit. The fit-time cull ranks splats by amplitude and keeps the top set accounting for a retention fraction of the total amplitude (0.95 by default); the contribution-based cull instead removes the splats whose contribution to the rendered field falls below an error budget or a fractional threshold, and is the baseline that removal-based level-of-detail methods use.

**relocation** The fixed-pool alternative to adding and removing splats during optimisation: the splat count is fixed at seeding, and the least informative splats are periodically moved to regions of high reconstruction residual, so the optimiser tensor shapes never change.

**leaf** A node that carries splat arrays directly (centres, amplitudes, Cholesky factors) together with an additive ladder of at least one sub-LOD, as opposed to a group node that only holds children.

**iSIM** Instant structured illumination microscopy, a super-resolution fluorescence technique that performs the structured-illumination reconstruction optically in a single exposure, giving roughly a twofold resolution gain at camera frame rates.

### 2 Methods

#### 2.1 Single-pass and progressive fitting as run

The nominal budget is $K = 32{,}000$ seeds (the initial splat positions from which optimisation starts) and $I = 10{,}000$ optimisation iterations. A cell is one volume and one pass count $N \in \{1, 2, 4, 8\}$; each cell holds one fit.

- **Single-pass** ($N = 1$): one call to `fit_gaussian_splats` with $K$ requested seeds and $I$ iterations on the full volume $V$. Seeds come from the `auto` seeder, a deduplicated mixture of edge-based (60% of the budget) and grid (40%) seeds; the seeder skips its subsampling step when the candidate count lies within 5% above the request, so a single-pass fit may start from slightly more than $K$ seeds. The optimiser is Adam with a reduce-on-plateau learning-rate schedule (the learning rate is multiplied by 0.9 after 15 iterations without loss improvement) and fixed-pool *relocation* (every 50 iterations the least informative splats are moved to residual peaks; the pool size never changes). The fitter tracks the lowest training loss it visits and returns *that* state (column `best_iteration` in Table 3), not the state at iteration $I$. A post-fit cumulative-amplitude *cull (retention)* at retention 0.999 then removes the splats that together carry the last 0.1% of the total amplitude.
- **Progressive** ($N \in \{2, 4, 8\}$): one call to the progressive fitter requesting $K$ seeds in total, $K/N$ seeds and $I/N$ iterations per pass and at most $N$ passes, with every shared keyword forwarded to each pass (function `fit_progressive_gaussian_splats`; arguments `max_splats`, `max_splats_per_pass`, `iters_per_pass`, `max_passes`). Pass 0 fits $K/N$ seeds to $V$ for $I/N$ iterations with the single-pass seeding, loss and learning rate. Pass $n > 0$ fits a further $K/N$ new splats to the clipped residual $\max(V - \hat{V}^{(n-1)}, 0)$, where $\hat{V}^{(n-1)}$ is the

rendered reconstruction of all splats accumulated through pass $n-1$; the previously fitted splats are frozen and only the new splats receive gradient updates. Each residual pass is normalised to $[0,1]$ by its own range before fitting (the clipped residual's minimum is 0, so by its own maximum) and its amplitudes are rescaled back on return, so the loss, the regularisers, the learning rate and the tolerances act in residual-relative units that shrink from pass to pass. Each pass is itself a `fit_gaussian_splats` call and returns its own lowest-loss state. Per-pass culling is disabled; the same cumulative cull at retention 0.999 is applied once to the accumulated set, and the result is flattened into a single *leaf*. The four pass counts therefore request $(K/N,\ I/N)$ = (32,000, 10,000), (16,000, 5,000), (8,000, 2,500) and (4,000, 1,250) per pass.

Figure 1 sketches the two procedures, and Figure 2 shows, on one slice of two volumes, the target, a saved reconstruction, the clipped residual a later pass is fitted to and an intensity-weighted seed draw on it.

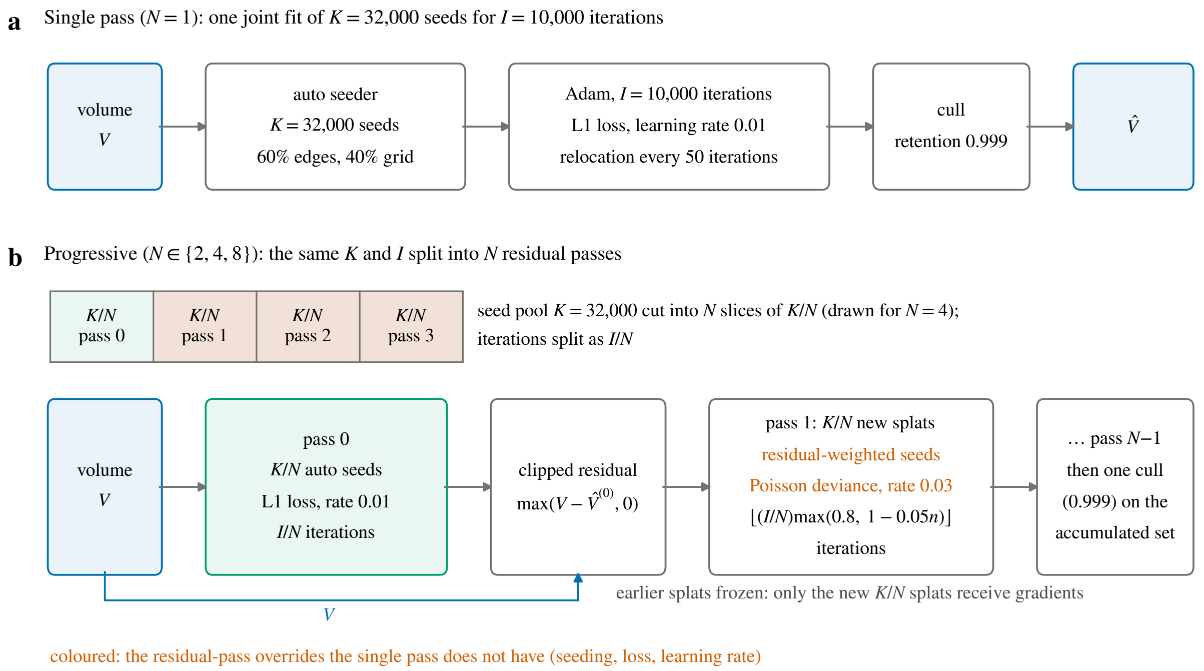


Figure 1: Schematic of the two procedures (no data plotted). (a) The single pass fits $K = 32{,}000$ auto-seeded splats to $V$ in one Adam run of $I = 10{,}000$ iterations under the L1 loss at learning rate 0.01, then culls at retention 0.999. (b) The progressive fitter cuts the seed pool into $N$ slices of $K/N$ and the iterations into $I/N$. Pass 0 fits its slice to $V$ with the single-pass seeding, loss and learning rate. Every later pass fits $K/N$ new splats to the clipped residual $\max(V - \hat{V}^{(n-1)}, 0)$ of everything fitted so far, whose splats stay frozen; the three settings in colour (intensity-weighted residual seeding, Poisson-deviance loss, learning rate 0.03) are the residual-pass overrides the single pass does not have, and the iteration count decays by 5% per pass down to 80% of $I/N$. One cull at retention 0.999 then acts on the accumulated set. Table 1 lists every setting.

Residual passes ($n > 0$) apply overrides that the progressive fitter hard-codes and that the single-pass baseline does not have:

- **Loss**: Poisson deviance instead of L1. This is an empirical choice of a non-negative divergence

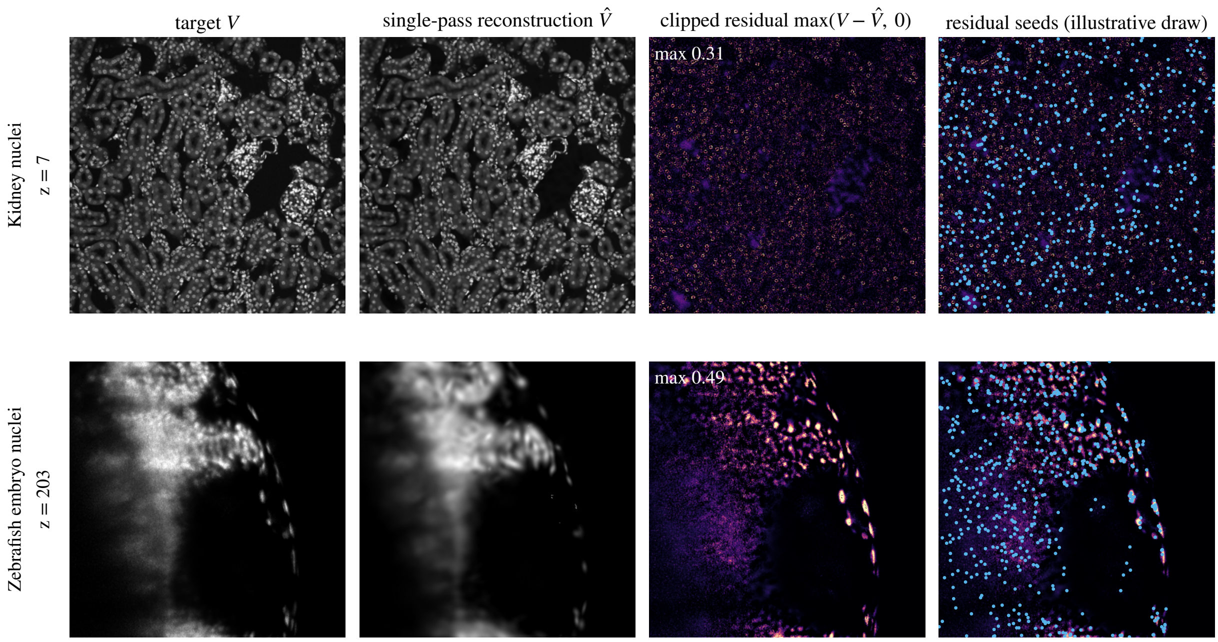


Figure 2: What a residual pass sees, on one saved slice of two volumes: Kidney nuclei, which the single pass won, and Zebrafish embryo nuclei, which 2-pass won. Columns: the target $V$; the saved single-pass reconstruction $\hat{V}$ ($K = 32{,}000$ seeds, $I = 10{,}000$ iterations); the clipped residual $\max(V - \hat{V}, 0)$, the quantity a residual pass is fitted to, each panel stretched to its own 99.5th percentile with its maximum printed; and the same residual with an intensity-weighted draw of seed positions made for this figure on this slice with the residual seeder's rule (multinomial sampling without replacement, probability proportional to residual intensity). The residual that pass 1 of the 2-pass fit saw is the residual of that fit's own pass 0 ($K/2$ seeds, $I/2$ iterations), which was not saved, so the third and fourth columns apply the residual operation to a saved reconstruction and the seed draw is illustrative, not the fitter's (which samples the whole 3-D residual). On the dense confocal volume the residual is a speckle spread over the tissue and the seeds land almost uniformly; on the light-sheet embryo the residual concentrates on the bright nuclei at the periphery, and so do the seeds.

for a sparse, clipped residual; the clipped residual is not Poisson-distributed (clipping a zero-mean fluctuation at zero changes its variance-to-mean ratio), so the loss is used as a heuristic, not as a likelihood.

- **Seeding**: intensity-weighted sampling of the positive residual voxels (`seed_from_peaks`: multinomial sampling without replacement with probability proportional to residual intensity) instead of the auto seeder's edge/grid mixture.
- **Learning rate**: 0.03 instead of 0.01.
- **Iteration count**: $\max\{500,\ \lfloor (I/N)\ \max(0.8,\ 1 - 0.05\,n) \rfloor\}$, where $\lfloor\cdot\rfloor$ rounds down; the schedule is a 5% decay per pass down to 80% of $I/N$, and the lower bound of 500 does not bind at this budget. The totals are 9,750 ($N = 2$), 9,250 ($N = 4$) and 8,624 ($N = 8$) iterations, i.e. 2.5–13.8% below the single pass's 10,000 (column `iters_actual_total`).

The residual seeder has one capacity limit. On a residual with more than $2^{24} - 1$ ($\approx 16.8\,$M) positive voxels, the category limit of `torch.multinomial`, the sampling support is first truncated to its $2^{24} - 1$ brightest voxels. Of the seventeen volumes, twelve have more voxels than that in total (Table 2); the positive-voxel count of each residual is not recorded, so whether the truncation fired is unknown.

Table 1 lists the setting of every stage. Two of the progressive fitter's own defaults are *not* in force here because the runner passes the single-pass values through. The asymmetric over-prediction penalty (a factor multiplying the loss wherever the reconstruction exceeds the target) is 1.0, that is symmetric, on every pass; the fitter's default is 10 on residual passes, capped at 3 on pass 0. Fixed-pool relocation is *enabled* on every pass, where the fitter's default is off. Conversely, the progressive fitter removes the eccentricity bound (a cap on the ratio of a splat's largest to smallest Cholesky diagonal entry) on every pass; the runner removes it for the single pass as well, so the two conditions share it. Background-floor subtraction inside the fitter is off (`floor="none"`) in both conditions, so PSNR is measured against the volume as the shared registry loader normalises it; for two volumes that loader itself subtracts a background floor before normalising (§ 2.2), so their target is background-relative.

This is a specified Python-API configuration, not the shipped CLI. The single-pass API defaults are 1,000 iterations, loss patience 300, absolute-error stop 0.01, eccentricity bound `max_eccentricity` $= 10$ (enforced as $\max_j \ell_{jj} / \min_j \ell_{jj} \leq \sqrt{10}$, with $\ell_{jj}$ the diagonal entries of a splat's Cholesky factor), cull retention 0.95 and `floor="auto"`. The progressive API defaults are asymmetric penalty 10, relocation off, cull retention 0.98 and an inter-pass PSNR patience of 0.5 dB, the minimum PSNR gain a pass must deliver for the next pass to run.

Table 1: Condition-by-stage configuration as executed. Every setting in the first column reaches both fitters; the third column's loss, seeding and learning rate are the progressive fitter's residual-pass overrides. The RNG seed (the integer that initialises the torch, CUDA and NumPy random number generators) is the pass count $N$ itself $(1, 2, 4, 8)$, one run per cell. The replicate fits of § 2.6 use $N + 1000\,r$ for replicate $r \in \{1, 2\}$.

| Setting | Single pass | Progressive pass 0 | Progressive passes $n \geq 1$ |
|---|---|---|---|
| Target | $V$ | $V$ | $\max(V - \hat{V}^{(n-1)}, 0)$ |
| Seeds requested | $K$ | $K/N$ | $K/N$ |
| Seeding | auto (edges + grid) | auto (edges + grid) | intensity-weighted residual voxels |
| Loss | L1 | L1 | Poisson deviance |
| Asymmetric penalty | 1.0 (symmetric) | 1.0 | 1.0 |
| Learning rate | 0.01 | 0.01 | 0.03 |
| Iterations | $I$ | $I/N$ | $\lfloor (I/N) \max(0.8, 1 - 0.05n) \rfloor$ |
| Relocation (fixed pool) | on | on | on |
| Eccentricity bound | none | none | none |
| Background floor | none | none | none |
| Loss patience | off | off | off |
| Absolute-error stop | $10^{-9}$ (unreachable) | $10^{-9}$ | $10^{-9}$ |
| Inter-pass PSNR patience | n/a | $-\infty$ | $-\infty$ |
| Earlier splats | n/a | n/a | frozen |
| State returned | lowest-loss state | lowest-loss state | lowest-loss state |
| Culling | post-fit cumulative, 0.999 | none | once, cumulative 0.999, on the accumulated set |

Differences between conditions therefore reflect a combination of (i) *how* the budget is spent (jointly on the full volume vs. in $N$ residual-targeted stages) and (ii) the residual-pass overrides above. The comparison is between two compound fitting procedures at a common nominal seed budget, not an isolated staging ablation.

##### 2.2 Datasets

The seventeen benchmark volumes (17 volumes from 12 datasets; Table 2) span four microscopy modalities (spinning-disk confocal, confocal, light-sheet, *iSIM*) and a $\sim 27\times$ range of voxel counts (from 3.9 M to $\sim$107 M). The sweep covers all seventeen.

Table 2: The seventeen benchmark volumes (17 volumes from 12 datasets), in registry order with modality groups separated by rules. Volumes are normalised to $[0, 1]$ before fitting; the clipping range is per volume and recorded in the shared dataset registry, matching the loaders shared with Supplementary Document 2. For two volumes that loader also subtracts a background floor before normalising, clipping at 0 (Neuromast membranes (106.0) and Neuromast nuclei (103.9); raw intensity units), so their fits and metrics are against a background-relative volume.

| Volume | Modality | Shape | Voxels |
|---|---|---|---|
| OpenCell-MAP4 nuclei | Spinning-disk | 51×600×600 | 18.4 M |
| OpenCell-MAP4 microtubules | Spinning-disk | 51×600×600 | 18.4 M |
| OpenCell-LMNB1 nuclei | Spinning-disk | 112×600×600 | 40.3 M |
| OpenCell-LMNB1 nuclear lamina | Spinning-disk | 112×600×600 | 40.3 M |
| Kidney nuclei | Confocal | 16×512×512 | 4.2 M |
| Kidney actin | Confocal | 16×512×512 | 4.2 M |
| Cells3D nuclei | Confocal | 60×256×256 | 3.9 M |
| Cells3D membranes | Confocal | 60×256×256 | 3.9 M |
| Blastocyst nuclear lamina | Confocal | 236×275×271 | 17.6 M |
| *C. elegans* embryo nuclei | Confocal | 41×512×512 | 10.7 M |
| Fly brain neurons | Confocal | 435×478×478 | 99.4 M |
| *Tribolium* embryo nuclei | Light-sheet (deconv.) | 991×104×965 | 99.5 M |
| Mouse heart nuclei | Light-sheet | 597×174×960 | 99.7 M |
| Zebrafish embryo nuclei | Light-sheet (deconv.) | 407×512×512 | 106.7 M |
| *Drosophila* embryo nuclei | Light-sheet | 108×1352×532 | 77.7 M |
| Neuromast membranes | iSIM (deconv.) | 84×580×576 | 28.1 M |
| Neuromast nuclei | iSIM (deconv.) | 84×580×576 | 28.1 M |

##### 2.3 Stopping rules and realised budgets

No stopping rule fired in any of the 68 fits. The loss patience is disabled in both conditions (`early_stop_patience=None`, forwarded to every progressive pass), the absolute-error convergence stop is set to an unreachable $10^{-9}$ of the normalised range (its default of 0.01 would otherwise be active), and the inter-pass PSNR patience is $-\infty$, so every requested pass runs. The progressive fitter's residual-negligible stop (residual maximum below $10^{-6}$) stayed armed but never triggered: all 51 progressive rows record `n_passes_actual` equal to the requested pass count and the stop reason `max_splats` (the loop ends because the seed budget is exhausted); the single-pass rows record `iteration_budget` with 10,000 iterations executed.

The number of retained splats is not fixed (Table 3). It varies with the seeder (the 5% tolerance above) and with the post-fit cull, which retains a fraction of the total *amplitude*, not of the count: across all 68 cells the retained count runs from 22,178 (Neuromast nuclei, 2-pass; 69% of the nominal budget) to 32,384 (OpenCell-MAP4 microtubules, 1-pass). The nominal 32,000 is exceeded by two single-pass cells (OpenCell-MAP4 nuclei 1-pass, 32,320 and OpenCell-MAP4 microtubules 1-pass, 32,384). Per condition the medians are 31,390 (1-pass), 30,762 (2-pass), 30,440 (4-pass) and 30,220 (8-pass). The two neuromast channels retain the fewest splats in the progressive conditions. The culled splats are the ones that together carried the last 0.1% of the amplitude, so a smaller retained count means the fit left more of its pool as near-zero contributors: on Neuromast nuclei the 2-pass fit left 31% of the nominal pool below the cull. The count gap therefore measures capacity the progressive arm failed to use, not capacity withheld from it, and every quality comparison below is between fits of a common *nominal* seed budget; the counts are tabulated so the reader can see how much of the pool each condition used.

The single-pass fitter returns the lowest-loss state it visited. Its `best_iteration` lies between 1,000 and 10,000, below half the budget on 13 of 17 volumes and at the final iteration on two (Neuromast membranes and Neuromast nuclei). The training loss includes the fitter's L1 penalties on amplitudes and Cholesky diagonals (Supplementary Document 4) and is perturbed by every relocation step, so the lowest-loss iteration is not a convergence statement about PSNR; it records which state the reported single-pass metrics describe. The progressive fitter applies the same best-state restoration inside every pass; its per-pass best iterations are not in the results table.

Table 3: Retained (post-cull) splats per condition (nominal request of 32,000 seeds in every column) and the iteration at which the single pass reached its lowest training loss (the state it returns). Executed iterations are identical across volumes: 10,000 (1-pass), 9,750 (2-pass), 9,250 (4-pass) and 8,624 (8-pass).

| Volume | 1-pass | 2-pass | 4-pass | 8-pass | Best iter. (1-pass) |
|---|---|---|---|---|---|
| OpenCell-MAP4 nuclei | 32,320 | 31,478 | 31,303 | 31,178 | 1,100 |
| OpenCell-MAP4 microtubules | 32,384 | 31,420 | 31,239 | 31,146 | 1,350 |
| OpenCell-LMNB1 nuclei | 31,965 | 31,507 | 31,302 | 31,179 | 1,000 |
| OpenCell-LMNB1 nuclear lamina | 31,111 | 30,567 | 30,440 | 30,101 | 1,400 |
| Kidney nuclei | 31,668 | 30,728 | 30,418 | 30,220 | 2,050 |
| Kidney actin | 31,390 | 30,607 | 30,446 | 30,702 | 2,300 |
| Cells3D nuclei | 31,878 | 31,531 | 31,403 | 31,322 | 1,450 |
| Cells3D membranes | 31,840 | 30,908 | 30,482 | 30,108 | 2,000 |
| Blastocyst nuclear lamina | 28,458 | 29,318 | 30,263 | 30,417 | 1,550 |
| *C. elegans* embryo nuclei | 31,815 | 31,808 | 31,727 | 31,604 | 1,050 |
| Fly brain neurons | 29,605 | 30,839 | 28,364 | 30,047 | 1,000 |
| *Tribolium* embryo nuclei | 31,773 | 30,834 | 30,168 | 29,916 | 2,249 |
| Mouse heart nuclei | 30,199 | 30,762 | 30,787 | 30,447 | 5,584 |
| Zebrafish embryo nuclei | 29,352 | 29,987 | 29,858 | 29,459 | 7,350 |
| *Drosophila* embryo nuclei | 30,935 | 30,517 | 30,153 | 29,999 | 4,476 |
| Neuromast membranes | 30,426 | 24,941 | 24,730 | 23,165 | 10,000 |
| Neuromast nuclei | 26,006 | 22,178 | 24,173 | 23,183 | 10,000 |

##### 2.4 Reconstruction quality metrics and timers

Each fit's reconstruction $\hat{V}$ is rendered back to volume at the original shape and compared to the target volume $V$ on the full-volume grid, with $V_i$ and $\hat{V}_i$ the values at voxel $i$:

- **Training PSNR (dB)**: on the normalised volume,

$$\text{PSNR} = 10 \log_{10} \frac{\text{MAX}^2}{\text{MSE}},$$

  with MAX the target's range (= 1) and MSE the mean of $(\hat{V}_i - V_i)^2$ over the full grid.

- **Training SSIM**: 3-D structural similarity [1] with an $11^3$ Gaussian window ($\sigma = 1.5$), constants $C_1 = (0.01L)^2$, $C_2 = (0.03L)^2$ with $L = 1$ (the normalised range) and population moments. The local values are averaged over the valid interior, with five voxels excluded at every boundary; on the 16-slice kidney volumes only six axial positions therefore contribute. The implementation is identical across all conditions.

- **Maximum absolute error**: $\max_i |\hat{V}_i - V_i|$ on the normalised volume, a worst-case fidelity diagnostic that complements the average-case PSNR and SSIM; the rendered reconstruction is not clipped to $[0, 1]$, so this error can exceed 1.

- **Outer wall time (s)** (`fit_wall_s`): the runner's timer around the whole fitter call, CUDA-synchronised on both sides. Seeding, optimisation, per-pass residual renders and the final cull are inside. In the progressive arm only, so is the fitter's own scoring: a global-PSNR computation after every pass and a foreground-PSNR score at the end (PSNR with the error averaged over the target's foreground voxels only), neither of which the single-pass call performs. Loading and the runner's metric render and scoring are outside. This is the one boundary the two conditions share and is the timer used for every speed statement in this document.

- **Fitter-internal time (s)** (`fit_time_s`): the fitter's own `stats["time_seconds"]`, whose scope differs by condition. For the single pass it is the Adam loop only, with seeding, preprocessing and the cull excluded. For the progressive fitter it runs from normalisation through the last pass's global-PSNR render and the fitter's final foreground-PSNR score, so per-pass seeding, residual renders and per-pass PSNR scoring are included and the final cull is excluded. It is tabulated for completeness (Table 7) and not used for cross-condition ratios in the prose. On this sweep the single pass's outer time exceeds its internal time by 7–15 s (seeding and cull), while the two progressive timers agree to within 0.03 s.

All "fidelity" statements are fidelity to the training input: these scores do not separate fitted signal from fitted acquisition noise.

##### 2.5 Compute and provenance

Both conditions are direct calls into the Luxar Python API (`luxar.gsplats`); the resolved configuration of every call is recorded in the run manifest that accompanies the results. All 68 fits ran sequentially on GPU 0 of one NVIDIA RTX PRO 6000 Blackwell Workstation Edition (96 GB, compute capability 12.0) with no other fitting job on the card, under Python 3.12.3, PyTorch 2.14.0+cu130 (CUDA 13.0), luxar 2026.06.05 at commit `5ec6c9852` (all recorded in the run manifest). The sweep's row timestamps (each row is written after its fit has been scored) span 2.9 h, opening at the end of the first fit (OpenCell-MAP4 nuclei 1-pass, 97 s of outer wall time, which the span therefore excludes). The outer wall times of all 68 fits sum to 2.79 h (internal timers 2.74 h), those of the 67 fits inside the span to 2.76 h; the remainder of the span is volume loading, metric rendering and scoring. Individual fits took 78–456 s. The per-volume results table carries, per row, the retained splat count, the executed pass and iteration counts, the stop reason, the RNG seed and both timers.

##### 2.6 Run-to-run replicates

The six volumes whose 1-pass vs. 2-pass training-PSNR margin is below 0.5 dB (OpenCell-LMNB1 nuclei, Cells3D membranes, *C. elegans* embryo nuclei, Fly brain neurons, Mouse heart nuclei and *Drosophila* embryo nuclei) are fitted two more times per condition, for the 1-pass and 2-pass conditions only. The replicates use the same budget, settings and software as the sweep (the same luxar commit, PyTorch 2.14.0+cu130, GPU 0 of the same NVIDIA RTX PRO 6000 Blackwell Workstation Edition), record their kernel path (`cuda_extension`; the sweep manifest does not record it) and use the RNG seed $N + 1000\,r$ for replicate $r \in \{1, 2\}$. The replicate fits have their own per-volume results tables and their own run manifest. With the sweep's own row as replicate 0, every such cell has three seeds (12 cells, 24 additional fits; row timestamps spanning 1.2 h).

Initialisation is deterministic: the initial `auto` seeding draws only from generators with fixed seeds and the relocation step's residual sample comes from a fixed private generator, so the seed changes only the progressive arm's residual seeder (an intensity-weighted `torch.multinomial` draw on the global stream). The remaining variation, all of the 1-pass arm's included, comes from sources the seed does not fix, among them the order of the CUDA kernels' atomic accumulation. The spread is therefore a run-to-run spread of the whole procedure, not a seed sensitivity alone, and is expected to understate the spread of a procedure whose initialisation also varied. No such replicate exists; the seed-varied 2-pass arm's spread is smaller than the 1-pass arm's on five of the six replicated volumes and larger on Mouse heart nuclei (by 0.001 dB)(Table 6).

The replicate fits share the GPU with other jobs, so their wall times are recorded in the results tables but not reported. The within-condition spread of a cell is the range (maximum minus minimum) of its training PSNR over the seeds; two conditions are called disjoint on a volume when their ranges do not overlap. The same range is taken over SSIM. The 4- and 8-pass conditions and the other volumes are not replicated.

### 3 Results

#### 3.1 Training PSNR, SSIM and maximum absolute error

Tables 4 and 5 list the three fidelity metrics of every fit; Table 6 and Figure 3 give the run-to-run replicates of the six volumes whose 1-pass vs. 2-pass margin is below 0.5 dB.

Table 4: Training PSNR (dB) and training SSIM (valid-interior average, § 2.4), full-volume fit, one fit per cell; the columns of each group are the pass counts $N$. Bold marks the best condition per volume and metric: the highest PSNR, the highest SSIM. *PSNR.* Single-pass wins on 15/17 volumes, by 0.07 dB (Fly brain neurons) to 6.02 dB (Neuromast nuclei) over 2-pass, median 0.67 dB; 2-pass wins on Mouse heart nuclei (+0.40 dB) and Zebrafish embryo nuclei (+0.85 dB). PSNR declines strictly from 2 to 4 to 8 passes on all 17 volumes; 4-pass trails 1-pass by 1.7–13.7 dB and 8-pass by 4.0–19.8 dB (worst: Neuromast nuclei, then Neuromast membranes at 11.5 dB). The 1-pass vs. 2-pass margin is below 0.5 dB in either direction on six volumes (OpenCell-LMNB1 nuclei (−0.48 dB), Cells3D membranes (−0.38 dB), *C. elegans* embryo nuclei (−0.38 dB), Fly brain neurons (−0.07 dB), Mouse heart nuclei (+0.40 dB) and *Drosophila* embryo nuclei (−0.42 dB)). On these the 1-pass and 2-pass conditions carry two further seeds each (Table 6), and the two PSNR ranges over the three seeds are disjoint on six of the six. That spread varies GPU nondeterminism and, in the 2-pass arm, the residual-seeder draw, not the initialisation (§ 2.6). Of the fifteen single-pass wins, one is below the largest measured spread of 0.093 dB: Fly brain neurons (0.07 dB); 1-pass spread 0.007 dB, 2-pass spread 0.006 dB, ranges disjoint. Retained splat counts in Table 3. *SSIM.* 2-pass has the highest SSIM on 12/17 volumes and 1-pass on 5; the SSIM-best condition differs from the PSNR-best one on ten volumes (SSIM-best condition, with its SSIM margin over 1-pass): 2-pass on OpenCell-MAP4 nuclei (+0.0054), OpenCell-MAP4 microtubules (+0.0032), OpenCell-LMNB1 nuclei (+0.0048), OpenCell-LMNB1 nuclear lamina (+0.0164), Kidney nuclei (+0.0011), Kidney actin (+0.0350), Cells3D membranes (+0.0009), Blastocyst nuclear lamina (+0.0172), Fly brain neurons (+0.0089) and *Drosophila* embryo nuclei (+0.0204). On every such volume the SSIM-best condition has the lower PSNR. Several of these margins are in the third or fourth decimal; three of them (OpenCell-MAP4 microtubules (+0.0032), Kidney nuclei (+0.0011) and Cells3D membranes (+0.0009)) are below the largest measured SSIM spread. On the six replicated volumes (§ 2.6) the run-to-run SSIM spread over three seeds is at most 0.0040 (*Drosophila* embryo nuclei, 1-pass; median 0.0001). The 1-pass and 2-pass SSIM ranges are disjoint on six of them; the SSIM margins of the other volumes are single observations.

| | Training PSNR (dB) | | | | Training SSIM | | | |
|---|---|---|---|---|---|---|---|---|
| Volume | $N$=1 | 2 | 4 | 8 | $N$=1 | 2 | 4 | 8 |
| OpenCell-MAP4 nuclei | **28.62** | 28.03 | 26.21 | 23.98 | 0.6696 | **0.6750** | 0.6402 | 0.5925 |
| OpenCell-MAP4 microtubules | **25.28** | 24.72 | 22.77 | 20.39 | 0.6167 | **0.6199** | 0.5831 | 0.5296 |
| OpenCell-LMNB1 nuclei | **25.60** | 25.12 | 23.40 | 21.29 | 0.5085 | **0.5133** | 0.4832 | 0.4423 |
| OpenCell-LMNB1 nuclear lamina | **34.22** | 33.37 | 31.28 | 28.74 | 0.9065 | **0.9229** | 0.8887 | 0.8352 |
| Kidney nuclei | **34.96** | 33.80 | 30.05 | 26.32 | 0.9640 | **0.9651** | 0.9440 | 0.9074 |
| Kidney actin | **31.72** | 31.05 | 27.36 | 23.47 | 0.9086 | **0.9436** | 0.9071 | 0.8353 |
| Cells3D nuclei | **36.85** | 36.07 | 34.08 | 31.75 | **0.8750** | 0.8731 | 0.8609 | 0.8433 |
| Cells3D membranes | **50.18** | 49.80 | 46.73 | 43.15 | 0.9921 | **0.9930** | 0.9891 | 0.9809 |
| Blastocyst nuclear lamina | **39.35** | 38.67 | 36.52 | 34.00 | 0.9310 | **0.9482** | 0.9183 | 0.8533 |
| *C. elegans* embryo nuclei | **39.36** | 38.99 | 37.32 | 35.35 | **0.8920** | 0.8853 | 0.8495 | 0.8009 |
| Fly brain neurons | **44.60** | 44.53 | 42.29 | 39.02 | 0.9575 | **0.9663** | 0.9244 | 0.8581 |
| *Tribolium* embryo nuclei | **51.38** | 49.76 | 45.80 | 41.77 | **0.9961** | 0.9959 | 0.9946 | 0.9921 |
| Mouse heart nuclei | 32.41 | **32.81** | 30.53 | 27.63 | 0.8358 | **0.8583** | 0.8216 | 0.7633 |
| Zebrafish embryo nuclei | 32.97 | **33.82** | 31.25 | 27.52 | 0.9485 | **0.9694** | 0.9585 | 0.9304 |
| *Drosophila* embryo nuclei | **31.07** | 30.65 | 27.79 | 24.00 | 0.8462 | **0.8667** | 0.8477 | 0.8094 |
| Neuromast membranes | **53.62** | 50.18 | 45.67 | 42.14 | **0.9994** | 0.9987 | 0.9972 | 0.9946 |
| Neuromast nuclei | **66.54** | 60.52 | 52.87 | 46.79 | **0.9999** | 0.9995 | 0.9981 | 0.9961 |

Table 5: Maximum absolute error on the normalised volume, one fit per cell; columns are the pass counts $N$; lower is better, and bold marks the lowest error per volume. 1-pass has the smallest worst-case error on 12/17 volumes (OpenCell-MAP4 nuclei, OpenCell-LMNB1 nuclear lamina, Kidney nuclei, Kidney actin, Cells3D nuclei, Cells3D membranes, Fly brain neurons, *Tribolium* embryo nuclei, Mouse heart nuclei, Zebrafish embryo nuclei, *Drosophila* embryo nuclei and Neuromast membranes) and 2-pass on 5 (OpenCell-MAP4 microtubules (0.694 vs. 0.704), OpenCell-LMNB1 nuclei (0.682 vs. 0.711), Blastocyst nuclear lamina (0.299 vs. 0.330), *C. elegans* embryo nuclei (0.160 vs. 0.171) and Neuromast nuclei (0.088 vs. 0.099)). 4- and 8-pass never hold the minimum.

| | Maximum absolute error | | | |
|---|---|---|---|---|
| Volume | $N$=1 | 2 | 4 | 8 |
| OpenCell-MAP4 nuclei | **0.541** | 0.559 | 0.588 | 0.658 |
| OpenCell-MAP4 microtubules | 0.704 | **0.694** | 0.753 | 0.939 |
| OpenCell-LMNB1 nuclei | 0.711 | **0.682** | 0.766 | 0.840 |
| OpenCell-LMNB1 nuclear lamina | **0.379** | 0.669 | 0.740 | 1.036 |
| Kidney nuclei | **0.322** | 0.467 | 0.590 | 0.817 |
| Kidney actin | **0.457** | 0.638 | 0.753 | 0.885 |
| Cells3D nuclei | **0.156** | 0.169 | 0.191 | 0.218 |
| Cells3D membranes | **0.104** | 0.106 | 0.143 | 0.213 |
| Blastocyst nuclear lamina | 0.330 | **0.299** | 0.383 | 0.506 |
| *C. elegans* embryo nuclei | 0.171 | **0.160** | 0.210 | 0.211 |
| Fly brain neurons | **0.357** | 0.506 | 0.614 | 0.717 |
| *Tribolium* embryo nuclei | **0.113** | 0.181 | 0.324 | 0.238 |
| Mouse heart nuclei | **0.570** | 0.647 | 0.779 | 0.785 |
| Zebrafish embryo nuclei | **0.903** | 0.933 | 1.251 | 1.189 |
| *Drosophila* embryo nuclei | **0.714** | 0.854 | 1.001 | 1.221 |
| Neuromast membranes | **0.233** | 0.278 | 0.737 | 0.430 |
| Neuromast nuclei | 0.099 | **0.088** | 0.225 | 0.515 |

Table 6: Run-to-run replicates (§ 2.6): training PSNR over three RNG seeds per cell (initialisation identical across the seeds, § 2.6) on the six volumes with a sub-0.5 dB margin. Spread is the per-cell range; the margin is 2-pass minus 1-pass over the seed means; the last column says whether the two ranges overlap. Per-cell spreads are 0.002–0.093 dB (median 0.013 dB; largest Cells3D membranes 1-pass); the largest spread per volume is OpenCell-LMNB1 nuclei (0.003 dB), Cells3D membranes (0.093 dB), *C. elegans* embryo nuclei (0.003 dB), Fly brain neurons (0.007 dB), Mouse heart nuclei (0.055 dB) and *Drosophila* embryo nuclei (0.075 dB). The 1-pass and 2-pass ranges are disjoint on six of the six volumes (winner by mean, margin of the means: OpenCell-LMNB1 nuclei (1-pass, −0.48 dB), Cells3D membranes (1-pass, −0.41 dB), *C. elegans* embryo nuclei (1-pass, −0.38 dB), Fly brain neurons (1-pass, −0.06 dB), Mouse heart nuclei (2-pass, +0.43 dB) and *Drosophila* embryo nuclei (1-pass, −0.37 dB)). Figure 3 plots every seed.

| Volume | 1-pass PSNR (dB) | | 2-pass PSNR (dB) | | Margin | Ranges |
|---|---|---|---|---|---|---|
| | min–max | spread | min–max | spread | (2-pass − 1-pass) | overlap |
| OpenCell-LMNB1 nuclei | 25.60–25.60 | 0.003 | 25.12–25.12 | 0.002 | −0.48 | no |
| Cells3D membranes | 50.17–50.26 | 0.093 | 49.78–49.80 | 0.019 | −0.41 | no |
| *C. elegans* embryo nuclei | 39.36–39.37 | 0.003 | 38.98–38.99 | 0.003 | −0.38 | no |
| Fly brain neurons | 44.59–44.60 | 0.007 | 44.53–44.54 | 0.006 | −0.06 | no |
| Mouse heart nuclei | 32.38–32.43 | 0.054 | 32.81–32.86 | 0.055 | +0.43 | no |
| *Drosophila* embryo nuclei | 31.00–31.07 | 0.075 | 30.65–30.68 | 0.026 | −0.37 | no |

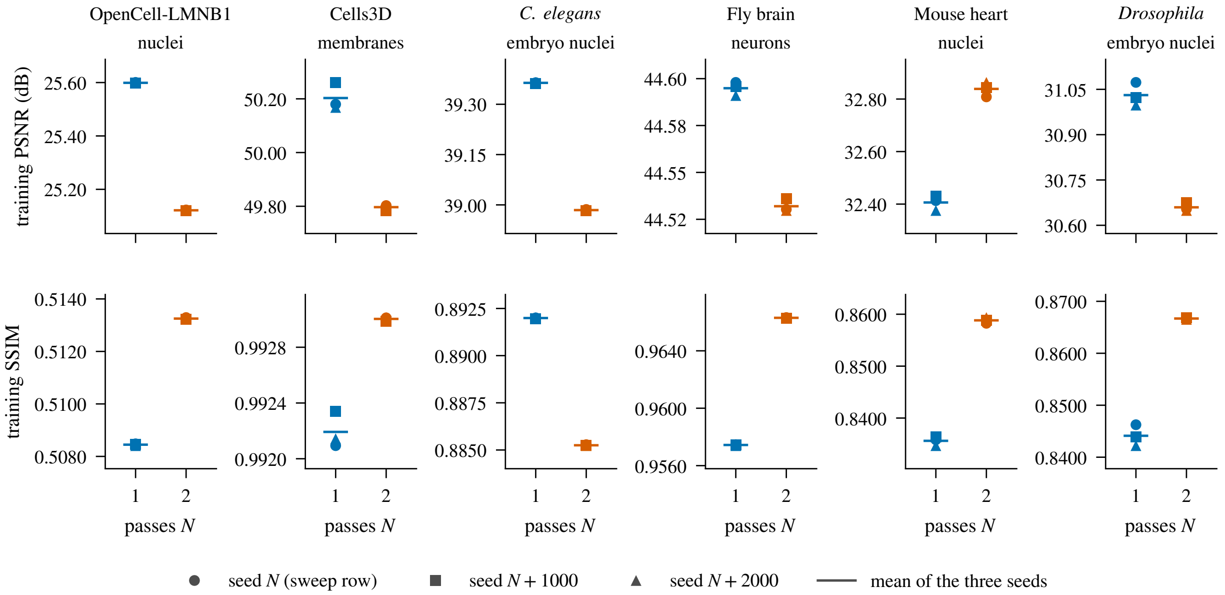


Figure 3: Run-to-run replicates of Table 6, every seed shown. One column per replicated volume, 1-pass (blue) and 2-pass (vermilion) side by side, one marker per RNG seed (circle: the sweep row, seed $N$; square: seed $N + 1000$; triangle: seed $N + 2000$) and a short line at the mean of the three; top row training PSNR, bottom row training SSIM. Each panel has its own vertical range, so the within-condition spread (thousandths to tenths of a decibel) is read against the between-condition margin of the same volume; the 1-pass and 2-pass ranges are disjoint on six of the six volumes for PSNR and on six for SSIM. The seed varies only the 2-pass arm's residual seeder; the 1-pass arm's spread is GPU nondeterminism alone (§ 2.6).

##### 3.2 Fit time

Table 7 lists both timers of every fit.

##### 3.3 Cross-dataset summary

Figure 4 restates the 2-pass minus 1-pass margin of Tables 4, 5 and 7 per volume on one diverging colour scale per metric, and Figure 5 follows every volume across the four pass counts. Three patterns are visible at a glance. First, the PSNR column is red (1-pass ahead) on fifteen volumes and blue on two (Mouse heart nuclei (+0.40 dB) and Zebrafish embryo nuclei (+0.85 dB)), and every volume's line in Figure 5a falls from 2 to 4 to 8 passes. Second, the SSIM column is blue on 12 of 17 volumes, by margins in the second to fourth decimal. Third, the maximum-error column is red on 12 volumes, while the wall-time column has no dominant colour (2-pass finished first on 10 of 17).

#### 4 Interpretation

**Single-pass wins on PSNR for 15/17 volumes.** Splitting the nominal budget into $N$ residual stages reduces both the number of splats optimised jointly in any stage ($K/N$) and the iterations each stage receives ($I/N$, less the decay), while the earlier stages are frozen. On fifteen volumes

Table 7: Outer wall time of the whole fitter call (`fit_wall_s`, common boundary) and the fitter-internal timer (`fit_time_s`), in seconds, per pass count $N$; each $t_1/t_2$ column is the 1-pass value divided by the 2-pass value of the same timer, so values above 1 mean 2-pass finished first. Bold marks the fastest condition per volume and timer. *Outer wall time.* Measured sequentially on an otherwise idle GPU. 2-pass finishes first on 10/17 volumes and 1-pass on 7; the ratio runs from 0.70× (*C. elegans* embryo nuclei) to 1.75× (Fly brain neurons), median 1.07×. The fastest condition per volume is 1-pass on 2, 2-pass on 3, 4-pass on 7 and 8-pass on 5 volumes; 8-pass finishes before 1-pass on 15 of 17. These are single observations per cell in a fixed condition order (1, 2, 4, 8) and no mechanism is inferred from them. *Fitter-internal timer.* The scope differs by condition (§ 2.4): the single-pass value is the Adam loop alone, the progressive value includes per-pass seeding, residual renders and the fitter's own PSNR scoring. The 1-pass/2-pass ratio runs from 0.63× to 1.69× (median 1.01×; 2-pass lower on 9/17), consistent with the outer-timer columns once the single pass's excluded seeding and cull time (7–15 s) is accounted for. Reported for completeness; the prose uses the outer timer.

| | Outer wall time (s) | | | | | Fitter-internal time (s) | | | | |
|---|---|---|---|---|---|---|---|---|---|---|
| Volume | $N$=1 | 2 | 4 | 8 | $t_1/t_2$ | $N$=1 | 2 | 4 | 8 | $t_1/t_2$ |
| OpenCell-MAP4 nuclei | 96.8 | **84.5** | 91.9 | 90.9 | 1.15× | 86.2 | **84.5** | 91.9 | 90.9 | 1.02× |
| OpenCell-MAP4 microtubules | 90.3 | **81.5** | 83.0 | 82.5 | 1.11× | 82.2 | **81.5** | 83.0 | 82.5 | 1.01× |
| OpenCell-LMNB1 nuclei | 125.1 | 117.1 | **96.2** | 98.3 | 1.07× | 117.9 | 117.1 | **96.1** | 98.3 | 1.01× |
| OpenCell-LMNB1 nuclear lamina | **122.0** | 129.4 | 148.7 | 183.1 | 0.94× | **114.0** | 129.4 | 148.7 | 183.1 | 0.88× |
| Kidney nuclei | 120.2 | 128.7 | **84.1** | 110.3 | 0.93× | 109.0 | 128.7 | **84.1** | 110.3 | 0.85× |
| Kidney actin | 142.7 | 85.6 | **79.4** | 112.0 | 1.67× | 133.4 | 85.6 | **79.4** | 111.9 | 1.56× |
| Cells3D nuclei | 146.8 | 94.9 | **93.5** | 129.1 | 1.55× | 138.1 | 94.9 | **93.5** | 129.1 | 1.46× |
| Cells3D membranes | 146.4 | **93.2** | 111.5 | 124.5 | 1.57× | 135.4 | **93.2** | 111.5 | 124.4 | 1.45× |
| Blastocyst nuclear lamina | 130.6 | 179.6 | 127.5 | **118.8** | 0.73× | 118.8 | 179.6 | 127.4 | **118.8** | 0.66× |
| *C. elegans* embryo nuclei | 98.7 | 140.6 | 131.4 | **79.1** | 0.70× | 88.6 | 140.6 | 131.4 | **79.1** | 0.63× |
| Fly brain neurons | 453.4 | 259.5 | 444.1 | **247.6** | 1.75× | 438.2 | 259.5 | 444.1 | **247.6** | 1.69× |
| *Tribolium* embryo nuclei | 419.3 | 456.3 | 351.7 | **256.3** | 0.92× | 409.2 | 456.3 | 351.7 | **256.2** | 0.90× |
| Mouse heart nuclei | 128.1 | 173.5 | **110.3** | 127.0 | 0.74× | 119.2 | 173.5 | **110.3** | 127.0 | 0.69× |
| Zebrafish embryo nuclei | 191.8 | 161.6 | **145.0** | 151.4 | 1.19× | 180.3 | 161.6 | **145.0** | 151.4 | 1.12× |
| *Drosophila* embryo nuclei | 174.6 | 133.9 | **120.8** | 136.6 | 1.30× | 159.1 | 133.9 | **120.8** | 136.6 | 1.19× |
| Neuromast membranes | 87.1 | 82.9 | 83.4 | **78.3** | 1.05× | **77.7** | 82.9 | 83.4 | 78.3 | 0.94× |
| Neuromast nuclei | **95.7** | 100.5 | 208.7 | 129.3 | 0.95× | **83.5** | 100.5 | 208.7 | 129.3 | 0.83× |

the single joint fit scores higher, by 0.07–6.02 dB over 2-pass (Table 4); the gap widens with $N$ on every volume, to a median of 2.8 dB at 4 passes and 5.5 dB at 8. The largest losses are on the two deconvolved iSIM neuromast channels (Neuromast nuclei −19.8 dB, Neuromast membranes −11.5 dB at 8 passes), which are also the volumes whose progressive fits retain the fewest splats after the cull (Table 3). Since the cull removes only near-zero contributors, the smaller count is a symptom of the staged fit leaving part of its pool unused, not a separate cause to be netted out.

**2-pass wins on two of the four light-sheet volumes; the margins are small.** The exceptions are Mouse heart nuclei (+0.40 dB) and Zebrafish embryo nuclei (+0.85 dB). A residual pass can only help where the first pass leaves unfit signal that intensity-weighted seeding then reaches; that this happened on these two volumes and not on the other two light-sheet volumes (*Tribolium* embryo nuclei (−1.62 dB) and *Drosophila* embryo nuclei (−0.42 dB), 2-pass minus 1-pass) is an observation, and we have no coverage or residual-content measurement that explains it. A 2-of-4 record does not make light-sheet data a domain where 2-pass is expected to win. Both

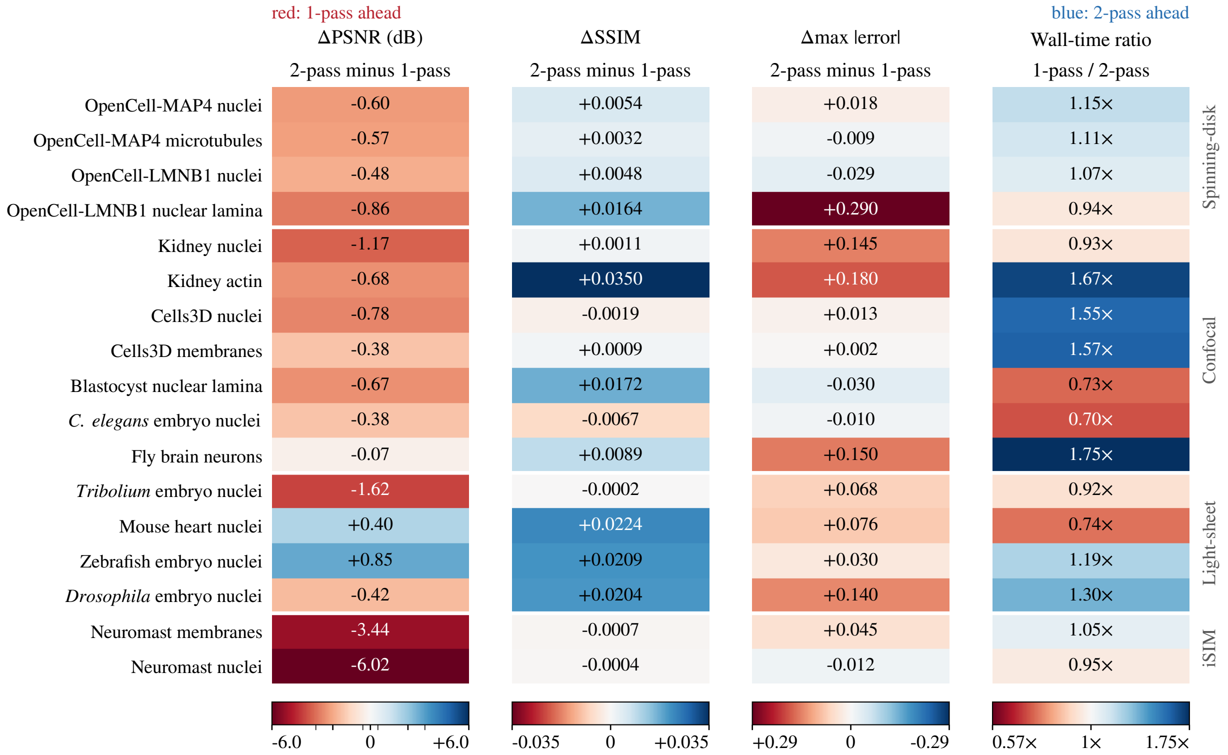


Figure 4: Cross-dataset summary as 2-pass minus 1-pass margins (one fit per cell). Rows are the seventeen volumes in registry order, grouped by modality as in Table 2; each column is one metric of Tables 4, 5 and 7, coloured on a diverging scale symmetric about zero whose limits are the largest magnitude in that column, so no cell is clipped. In every column blue means 2-pass ahead and red 1-pass ahead: the PSNR and SSIM columns show 2-pass minus 1-pass; the maximum-error column shows 2-pass minus 1-pass with the colour sign reversed (a lower error is better); the wall-time column shows 1-pass divided by 2-pass (above 1×, 2-pass finished first), coloured by its logarithm so 2× and 0.5× are equally saturated. The PSNR colour scale is linear within ±0.5 dB and logarithmic beyond, so the sub-decibel margins are visible beside the 6.02 dB one. Every cell prints its value.

gains are below 1 dB; the smaller holds across three seeds (which vary GPU nondeterminism and the residual draw only, § 2.6), the larger is a single fit.

**No volume favoured 4 or 8 passes at this budget.** PSNR declines strictly from 2 to 4 to 8 passes on all 17 volumes, and neither 4- nor 8-pass holds the minimum maximum error on any volume. Whether the per-pass iteration count (2,500 and then fewer at $N = 4$; 1,250 and then fewer at $N = 8$), the per-pass seed count, the frozen earlier stages or the residual-pass overrides drive the decline is not identified here; a larger budget could move the crossover. The statement is confined to this sweep and this budget.

**Fit time is comparable on an idle GPU.** With both conditions timed at the same boundary and no other job on the card, the 1-pass/2-pass outer-wall ratio is 0.70–1.75× (median 1.07×): 2-pass finishes first on 10 volumes and 1-pass on 7 (Table 7). The fastest condition varies by volume, with 4-pass fastest on 7 of 17. We do not attach a mechanism to these ratios: each is

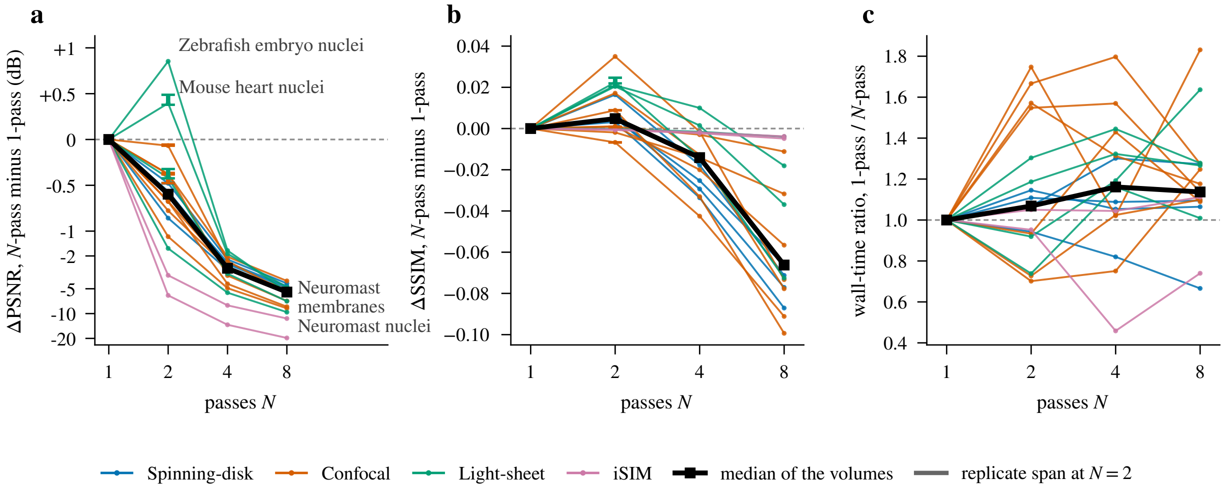


Figure 5: Effect size against the pass count, one thin line per volume coloured by imaging modality, the median over the seventeen volumes in bold. (a) Training PSNR of the $N$-pass fit minus the 1-pass fit; the axis is linear within $\pm 1$ dB and logarithmic beyond, so the sub-decibel 2-pass margins and the 19.8 dB 8-pass decline are both legible; the two volumes 2-pass won and the two largest 8-pass declines are named. (b) Training SSIM of the $N$-pass fit minus the 1-pass fit. (c) Outer wall time of the 1-pass fit divided by the $N$-pass fit; above the dashed line the $N$-pass fit finished first. In (a) and (b), at $N = 2$ the six replicated volumes carry a capped bar spanning the smallest and the largest 2-pass minus 1-pass margin over every pair of their three seeds per condition (§ 2.6); a bar that crosses zero is a volume whose 1-pass and 2-pass ranges overlap in Table 6, and most bars are shorter than the marker. Every value is a single fit; the sweep ran the conditions in a fixed order on an idle GPU.

one observation, the conditions differ in more than the pass count (seeding, loss, learning rate, iteration totals), and the sweep ran in a fixed order without interleaving or repetition.

**SSIM and PSNR pick different winners on ten volumes.** 2-pass has the highest SSIM on 12/17 volumes while 1-pass has the highest PSNR on 15/17 (Table 4). SSIM depends on local means, variances and covariances, so a fit can trade a little global MSE for local structure; a spatial analysis of the residual that would show *how* the two conditions differ has not been done, and several of the SSIM margins are in the third or fourth decimal (Table 4).

**Maximum absolute error favours 1-pass.** 1-pass has the lowest worst-case error on 12/17 volumes and 2-pass on 5, by narrow margins (Table 5). Together with the PSNR result, the single pass is the better choice on both average and worst-case fidelity on most volumes in this sweep.

### 5 Conclusion

Across seventeen microscopy volumes and four pass counts at a nominal 32,000-seed, 10,000-iteration budget, single-pass fitting has the higher training PSNR on 15/17 volumes (0.07–6.02 dB over 2-pass), the lower maximum error on 12/17, and takes comparable time (1-pass/2-pass outer-wall ratio 0.70–1.75×, median 1.07×). 2-pass has the higher PSNR on Mouse heart nuclei

(+0.40 dB) and Zebrafish embryo nuclei (+0.85 dB). Four and eight passes score below one pass on every volume (1.7–13.7 and 4.0–19.8 dB).

For the volumes and budget tested, a single joint fit is the default choice. 2-pass has the higher SSIM on 12/17 volumes, but only the six replicated volumes carry a measured SSIM spread: there the 1-pass and 2-pass SSIM ranges are disjoint on all six and 2-pass leads on five (OpenCell-LMNB1 nuclei, Cells3D membranes, Fly brain neurons, Mouse heart nuclei and *Drosophila* embryo nuclei). The other SSIM margins are single observations, three of them (OpenCell-MAP4 microtubules (+0.0032), Kidney nuclei (+0.0011) and Cells3D membranes (+0.0009)) below the largest measured SSIM spread of 0.0040. 2-pass also won PSNR on the two light-sheet volumes listed above (it lost on the other two light-sheet volumes, *Tribolium* embryo nuclei (−1.62 dB) and *Drosophila* embryo nuclei (−0.42 dB)), with the caveat that those margins are below 1 dB and only the sub-0.5 dB one of them is replicated (§ 2.6). We make no recommendation beyond this budget.

*Replication.* Each of the 68 cells of the sweep is a single fit with the pass count as its RNG seed. The within-condition spread is measured only for the 1-pass and 2-pass conditions on the six volumes whose margin is below 0.5 dB, at three seeds per cell (§ 2.6). There the per-cell PSNR range is 0.002–0.093 dB (median 0.013 dB) and the two conditions' PSNR ranges are disjoint on six of the six. That spread does not sample initialisation, which is deterministic, so it samples only the GPU nondeterminism (both arms) and the residual-seeder draw (2-pass). Of the single-pass wins, one is below the largest spread it did measure: Fly brain neurons (0.07 dB). The 0.5 dB figure is the cutoff that selected the replicated volumes, about 5.4 times the largest measured spread; the other volumes' 1-pass vs. 2-pass margins are at least 0.5 dB but are not replicated, and neither are the 4- and 8-pass conditions. The 1-pass and 2-pass SSIM ranges are disjoint on six of the replicated volumes (Table 4). The third- and fourth-decimal SSIM margins elsewhere and the timing ratios near 1 remain single observations (the replicate fits share the GPU, so their wall times are not compared), and no threshold from another experiment is borrowed.

*Retained capacity.* The retained splat counts differ by condition and volume (Table 3). Since the cull removes near-zero contributors, that difference records how much of the common nominal pool each condition used, not a mismatch in the pool offered.

*Design.* The progressive condition changes loss, seeding and learning rate on residual passes as well as the staging, so it evaluates the progressive fitter as a procedure and does not isolate the effect of splitting the budget. Both conditions return their lowest-loss state, not the state at the final iteration. The budget of 32,000 seeds and 10,000 iterations is one operating point; at a larger per-pass budget the ranking of $N$ may change. The interaction with a more aggressive cull (retention well below 0.999, which removes more of the retained capacity after the fit) is outside the present scope.
