## Supplementary material for "Luxar: Gaussian splatting for microscopy and scalable interactive web visualisation of multidimensional scientific data": supp_doc_04_convergence

### Convergence behaviour of Gaussian-splat fitting

Supplementary Document 4 – Luxar


**Abstract**

A Gaussian-splat fit has two budgets, the splat count and the number of optimiser iterations; this document asks which one sets reconstruction quality and how many iterations a fit needs. We fit 17 microscopy volumes at five splat counts (1 K to 256 K) and eleven requested Adam budgets (50 to 20,000 iterations), every cell an independent fit (935 fits), and repeat the budget axis in blind-spot mode at two counts per volume (374 further fits scored on held-out voxels). Training PSNR saturates early: at 2,000 requested iterations it is within 0.5 dB of the 20,000-iteration value on 77 of 85 cells, and capping the budget at 5,000 to 8,000 iterations changes any cell by at most +1.08 dB. Held-out PSNR at the calibrated splat count is within 0.2 dB of its peak at 2,000 on every volume; past the peak, training PSNR keeps rising while the held-out score stays flat or, on two iSIM channels, falls. Capacity dominates iterations: 4× the splats fitted for 40× fewer iterations beats the smaller, longer fit on every volume, by a mean of +3.03 dB, at 10× fewer splat-iterations. Convergence speed shows no detectable association with volume size ($r \approx +0.09$); the ten slow cells fall on six volumes, and on Kidney actin and the two neuromast channels, where they fall, the held-out sweep shows late training gains reaching no held-out voxel. We recommend setting the iteration budget once, at 2,000, and spending further effort on the splat count chosen by blind-spot calibration.


#### 1 Introduction

Fitting Gaussian splats to a volume is an optimisation with two budgets. The splat count $K$ fixes the capacity of the model, and the iteration count fixes how long Adam refines it. A user who wants a better reconstruction can raise either, and the two are not interchangeable in cost: every extra iteration renders the whole model, so its price scales with $K$. The first question of this document is which budget dominates quality, and the second is how many iterations a fit needs before more buy nothing that a caller would notice.

Answering the second question needs a convergence reference. An optimiser has no natural end point on this objective, so we take the training PSNR of the longest fit in the sweep as the reference and report how far shorter budgets fall short of it. That reference is a late-budget value, not a proven asymptote, and it is measured on the fitted voxels, so it rewards noise memorisation as much as signal recovery. A second sweep in *blind-spot cross-validation* mode scores the same budgets on *held-out voxels* the fit never saw, which separates the two.

**Notation.** $V$ is the target volume, normalised to $[0, 1]$, and $\hat{V}$ the rendered reconstruction of a fitted model. *PSNR* is $10 \log_{10}(1/\mathrm{MSE})$ in decibels with MSE the mean squared error between $\hat{V}$ and $V$ over the scored voxels. $K$ is the requested splat count and $n$ the requested iteration budget; a *cell* is one (volume, $K$) pair, a *checkpoint* one (volume, $K$, $n$) fit. $P_{20\text{K}}$ is the training PSNR of a cell's 20,000-iteration fit, the convergence reference. $n^*$ is the first checkpoint at which a cell's training PSNR reaches 99% of $P_{20\text{K}}$ in dB, and $\bar{n}^*$ its mean over the five splat counts of a volume. In the blind-spot sweep the same fits carry two crossings: $n^*_{\text{tr}}$ applies the $n^*$ criterion to their unmasked-voxel PSNR and $n^*_{\text{ho}}$ is the first checkpoint at or above 99% of the held-out peak. $K^*$ is the splat count that blind-spot calibration recommends for a volume (Supplementary Document 2), called the operating point below.

#### Terms used in this document

**PSNR** The peak signal-to-noise ratio, $10\log_{10}(\text{peak}^2/\text{MSE})$ in decibels, where peak is the reference range of the values compared; unless a document states another range, the supplements take peak 1 on volumes normalised to $[0, 1]$, so that $\text{PSNR} = -10\log_{10}\text{MSE}$.

**SSIM** The structural similarity index, the mean over local windows of a product of luminance, contrast and structure comparisons between two images, lying in $[-1, 1]$ with 1 meaning identical.

**blind-spot cross-validation** A model-selection protocol in which a deterministic random 5% of voxels is hidden from the fit and the reconstruction is scored on those voxels afterwards, so that capacity spent on memorising noise shows up as a loss instead of a gain.

**held-out voxels** The masked voxels of a blind-spot split, whose original values the optimiser never sees and against which the held-out PSNR of each candidate splat count is scored.

**donut-median fill** The replacement of each masked voxel by the median of its $3^D - 1$ neighbours (the donut, 26 voxels in 3D); the shipped calibration draws the median over the unmasked neighbours only, and a document whose sweeps used all 26 neighbours, masked ones included, states that variant where it applies.

**$K^*$** The splat count recommended by a blind-spot sweep: the interior maximum of the held-out PSNR curve when there is a clear peak, the smallest count within 0.3 dB of the maximum on a plateau, and the largest count tested when the curve is still rising there (signal-limited). The shared operating-point table of these documents takes the knee instead, the smallest count within 0.3 dB of that maximum, which is the largest count itself on the two signal-limited benchmark volumes.

**signal-limited curve** A held-out PSNR curve whose maximum is the largest splat count tested and which is still climbing there (a total rise of at least 0.3 dB across the sweep, a trailing mean rise of at least 0.1 dB per step and a final step of at least 0.05 dB), so the peak lies beyond the sampled range.

**relocation** The fixed-pool alternative to adding and removing splats during optimisation: the splat count is fixed at seeding, and the least informative splats are periodically moved to regions of high reconstruction residual, so the optimiser tensor shapes never change.

**cull (retention)** The removal of splats after a fit. The fit-time cull ranks splats by amplitude and keeps the top set accounting for a retention fraction of the total amplitude (0.95 by default); the contribution-based cull instead removes the splats whose contribution to the rendered field falls below an error budget or a fractional threshold, and is the baseline that removal-based level-of-detail methods use.

**Morton and Hilbert ordering** Two space-filling curves that sort points so that neighbours in space become neighbours in memory or on disk: Morton (Z-order) interleaves the bits of the quantised coordinates, Hilbert follows a continuous curve whose consecutive cells are always adjacent, so it has no long jumps between neighbouring runs. The compiler orders the stored elements along one of them; the fitter re-sorts its splats by Morton code periodically for GPU locality, and the substitutive LOD builder uses a Morton sort as its warm-start partition.

**iSIM** Instant structured illumination microscopy, a super-resolution fluorescence technique that performs the structured-illumination reconstruction optically in a single exposure, giving roughly a twofold resolution gain at camera frame rates.

### 2 Methods

#### 2.1 Experimental design

We measure how reconstruction quality evolves during optimisation as a joint function of model capacity (number of Gaussian splats) and optimisation budget (number of Adam iterations); Figure 1 lays the design out. For each dataset we fit at five splat counts $K \in \{1\,\mathrm{K}, 4\,\mathrm{K}, 16\,\mathrm{K}, 64\,\mathrm{K}, 256\,\mathrm{K}\}$ and for each splat count we record metrics at eleven iteration checkpoints

$$n \in \{50, 100, 200, 500, 1000, 2000, 3000, 5000, 8000, 12000, 20000\},$$

giving $5 \times 11 = 55$ fits per volume and 935 fits total across the seventeen volumes with results (Table 1). Each cell is a separate fit from scratch (no warm-starting; the RNG seed is $K + n$, so no two cells at the same splat count share an initial state, and cells at different $K$ differ in size), with $n$ the *requested* budget. A curve in this document is therefore a budget-response curve of independently fitted models, not the trajectory of one optimisation run. The reference for every convergence statement below is $P_{20\mathrm{K}}$, the training PSNR of the 20,000-iteration cell of the same (volume, $K$): a late-budget value, not a demonstrated asymptote, and a single checkpoint, so every cell reaches it at its last budget. Shortfalls against it are reported in absolute dB. A second sweep repeats the eleven budgets in blind-spot (held-out) mode at two splat counts per volume (§ 2.5): $17 \times 2 \times 11 = 374$ further fits that score each checkpoint on masked voxels the fit never saw.

#### 2.2 Datasets

The seventeen benchmark volumes (17 volumes from 12 datasets; Table 1) span confocal, spinning-disk, light-sheet and iSIM modalities and a 27× range of voxel counts. They are the same volumes used for the splat count vs. quality study (Supplementary Document 2), so the convergence analysis is directly comparable to those rate-distortion results. The results below cover the seventeen volumes for which the sweep has run.

#### 2.3 Fitting protocol

Each splat is parameterised by a centre, an amplitude and a lower-triangular Cholesky factor $L$ of its covariance, $\Sigma = LL^{\mathsf{T}}$, whose diagonal entries $L_{ii}$ ($i = 1, 2, 3$) set its extent along each axis. All cells share one set of caller settings: base learning rate 0.01, L1 reconstruction loss (symmetric), dynamic splat *relocation* enabled, a post-fit *cull (retention)* of 0.999, automatic seed placement, and `floor=none` (no background-floor subtraction, so every score is against the original normalised volume). Everything else is the fitter's default in the reference software (luxar at commit `5ec6c9852`), which resolves as follows. The optimiser is Adam [1] ($\beta = (0.9, 0.999)$, $\epsilon = 10^{-8}$, fused on CUDA) with per-group learning rates. The base rate is scaled by the gradient-dilution factor 9/5 to 0.018 for the Cholesky factors; the factor is the ratio of parameters per splat in 3D (three centre coordinates and six Cholesky entries) to 2D (two and three), and compensates for the

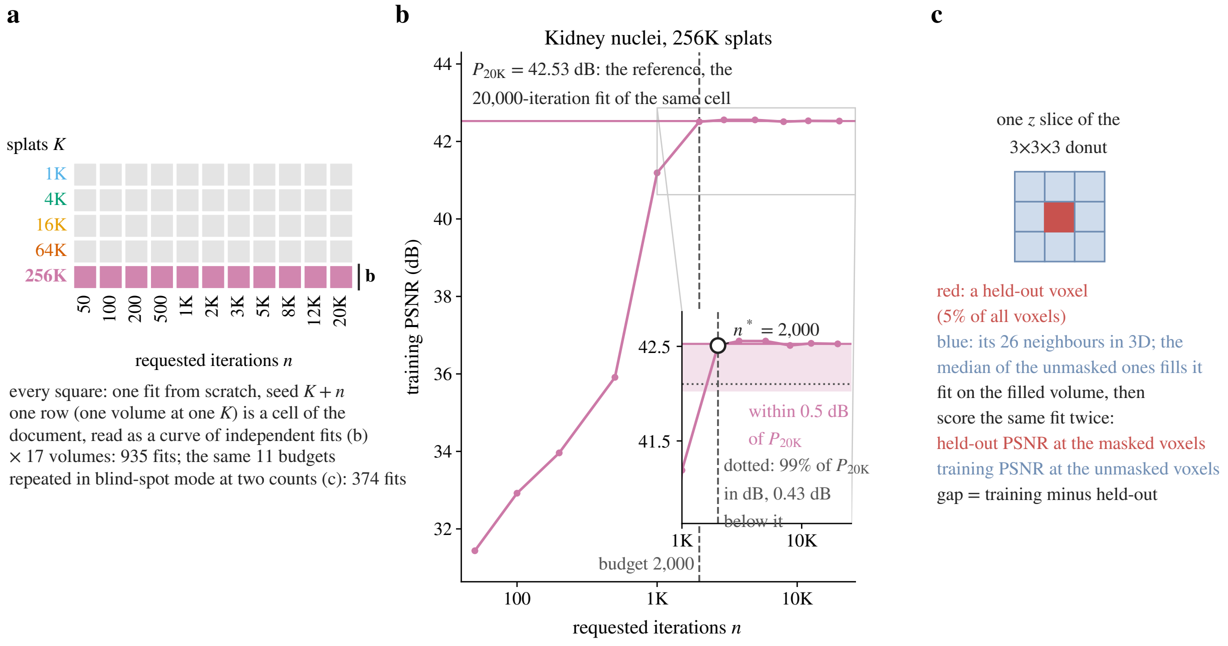


Figure 1: Experimental design. (a) The grid of one volume: five splat counts $K$ by eleven requested budgets $n$, every square an independent fit from scratch; one row (one volume at one $K$) is a cell of this document, read as a budget-response curve, and the grid is repeated for every volume (schematic). (b) The curve of one such row from the results, Kidney nuclei at 256 K splats: the reference $P_{20\text{K}}$ is the 20,000-iteration fit of the same row, the shaded band reaches 0.5 dB below it, the dotted line is 99% of $P_{20\text{K}}$ in dB, the hollow marker is the first checkpoint at or above that line ($n^*$) and the dashed line is the recommended budget of 2,000 iterations. (c) The blind-spot repeat (schematic): a held-out voxel is replaced by the median of its unmasked 3×3×3 neighbours, the fit sees the filled volume, and the same fit is scored on the masked voxels (held-out PSNR) and on the unmasked ones (training PSNR).

gradient of one splat being spread over more parameters. The centres take 1.5× that rate (0.027) and the amplitudes 3× (0.054). A `ReduceLROnPlateau` schedule [2] acts on the regularised training loss (patience 15 iterations, factor 0.9, relative threshold $10^{-3}$, minimum rate $10^{-8}$). The loss adds L1 regularisers $\lambda_a = 0.1\,\mathrm{lr} = 10^{-3}$ on the amplitudes and $\lambda_d = 0.01\,\mathrm{lr} = 10^{-4}$ on the Cholesky diagonals, each a mean over splats of the softplus-activated parameter ($\mathrm{softplus}(x) = \log(1 + e^x)$, the map that keeps amplitudes and diagonals positive). Eccentricity is bounded per splat at $\max_i L_{ii} / \min_i L_{ii} \le \sqrt{10}$ (the fitter's `max_eccentricity` = 10). Relocation moves the 1% least important splats (at most 64 per step, the fitter default `max_relocations_per_step` = 64, which the runner does not override) every 50 iterations (§ 5.2); splats are re-sorted along a Morton curve (*Morton and Hilbert ordering*) every 1,000 iterations. One fit per cell; no replicates.

**Stopping rules and the returned state.** The loss-patience early stop is disabled in both sweeps (`early_stop_patience=None`). The fitter has a second stop, on the absolute error (terminate when $\max|\hat{V} - V| < 0.01$, checked every 25 iterations, or every iteration for budgets below 100). The 935 training rows leave `max_abs_error` at the fitter default (0.01); the held-out rows pin it to `1e-9`. The smallest recorded maximum absolute error on the Supplementary Document 2 fits of the same volumes is 0.020, so the default threshold is not reached, but the training rows

| Volume | Modality | Shape | Voxels |
|---|---|---|---|
| OpenCell-MAP4 nuclei | Spinning-disk | 51×600×600 | 18.4 M |
| OpenCell-MAP4 microtubules | Spinning-disk | 51×600×600 | 18.4 M |
| OpenCell-LMNB1 nuclei | Spinning-disk | 112×600×600 | 40.3 M |
| OpenCell-LMNB1 nuclear lamina | Spinning-disk | 112×600×600 | 40.3 M |
| Kidney nuclei | Confocal | 16×512×512 | 4.2 M |
| Kidney actin | Confocal | 16×512×512 | 4.2 M |
| Cells3D nuclei | Confocal | 60×256×256 | 3.9 M |
| Cells3D membranes | Confocal | 60×256×256 | 3.9 M |
| Blastocyst nuclear lamina | Confocal | 236×275×271 | 17.6 M |
| *C. elegans* embryo nuclei | Confocal | 41×512×512 | 10.7 M |
| Fly brain neurons | Confocal | 435×478×478 | 99.4 M |
| *Tribolium* embryo nuclei | Light-sheet (deconv.) | 991×104×965 | 99.5 M |
| Mouse heart nuclei | Light-sheet | 597×174×960 | 99.7 M |
| Zebrafish embryo nuclei | Light-sheet (deconv.) | 407×512×512 | 106.7 M |
| *Drosophila* embryo nuclei | Light-sheet | 108×1352×532 | 77.7 M |
| Neuromast membranes | iSIM (deconv.) | 84×580×576 | 28.1 M |
| Neuromast nuclei | iSIM (deconv.) | 84×580×576 | 28.1 M |

Table 1: The seventeen benchmark volumes (17 volumes from 12 datasets), in registry order with modality groups separated by rules. Volumes are normalised to $[0, 1]$ before fitting; the per-volume clipping range is recorded in the shared dataset registry, matching the loaders in the splat-count-vs-quality study (Supplementary Document 2). Requested splat counts are reported throughout; the retained count $K_{\text{final}}$ differs from $K$ through seeding and the post-fit cull (§ 2.3).

record neither the executed iteration count nor a stopping flag, so this is not verified per row, and they carry no per-run software manifest. Every training-mode table and macro reads one per-volume metrics table written by the training sweep. The 374 held-out rows set the threshold to an unreachable $10^{-9}$ (`max_abs_error=1e-9`) and record the executed count, the iteration of the returned state and the stop flag: all 374 executed exactly the requested count and none stopped early. Their run manifest pins the same luxar commit, PyTorch `2.14.0+cu130`, CUDA `13.0` and the GPU.

In both sweeps the fitter returns the parameters of the iteration with the lowest regularised training loss, not the live final iterate, so a checkpoint's score is the score of that returned state. The held-out rows show what this means: the best-loss iteration equals the requested count on 162 of the 374 rows and lies earlier on 212. At the 20,000-iteration checkpoint the loss minimum sits within 100 iterations of the end on nine of the 34 cells (OpenCell-LMNB1 nuclear lamina at 64 $K$, Kidney nuclei at 64 $K$, Kidney actin at 128 $K$, *Tribolium* embryo nuclei at 256 $K$, Mouse heart nuclei at 256 $K$, Zebrafish embryo nuclei at 256 $K$, *Drosophila* embryo nuclei at 256 $K$, Neuromast membranes at 128 $K$ and Neuromast nuclei at 64 $K$, the high-capacity cells whose training PSNR keeps rising) and between iterations 750 and 3,497 on the other 25, every 4 $K$-splat cell among them. On those cells the regularised loss never improves after the first few thousand iterations, and the flat tail of the curve from there to 20,000 is the score of that early state, re-found by each independently seeded fit, not of a still-moving iterate. The training rows come from the same fitter, so the same reading applies to their plateaus; their best-loss iterations were not recorded. A budget-cap statement in this document is therefore a statement about the state the fitter returns at that cap, which is what a caller receives.

**Scoring.** After each fit the float32 model (not a decoded store) is rendered to the target grid and four metrics are recorded against the target: PSNR $= 10\log_{10}(1/\text{MSE})$ on the $[0,1]$-normalised volume, SSIM [3] with an $11^3$ Gaussian window ($\sigma = 1.5$, valid convolution), MSE, and relative $\text{L}^2$ error $\|\hat{V} - V\|_2/\|V\|_2$. `fit_time_s` is the fitter's internal optimisation-loop time (the Adam loop only; seeding, best-state rescoring, culling, rendering and scoring are excluded); every time axis and speed-up in this document uses it.

**Requested vs. retained counts.** The retained splat count $K_{\text{final}}$ differs from the requested seed count $K$: the automatic seeder may place a different number of seeds than requested (retained counts above $K$ occur), and the post-fit cull removes the splats carrying the lowest 0.1% of cumulative amplitude (relocation itself keeps the pool size fixed). Over the 935 rows $K_{\text{final}}/K$ ranges from 8% to 104% (median 96%; 412 rows below 95%), the smallest being Neuromast nuclei at $K = 256\,K$, $n = 50$ with 21,088 splats retained. The training rows record only the retained count, so seeding and cull are not separated per row. This document labels every cell by its requested count.

#### 2.4 Compute

All 935 fits ran on a single NVIDIA RTX PRO 6000 Blackwell (96 GB VRAM, compute capability 12.0), as two concurrent lanes sharing the card on a workstation that was simultaneously running unrelated CI jobs (1-minute load average 12–36 on a 32-thread host; the sweep ran at `nice` 15). Per-fit optimisation-loop time ranges from 0.44 s (Cells3D nuclei at $K = 256\,K$, $n = 50$) to 2,703 s (Fly brain neurons at $K = 16\,K$, $n =$ 20,000, a 99 M-voxel volume rendered at every step; the same volume's 256 K fit at that budget took 2,039 s, so individual times do not order by splat count); the per-fit loop times sum to 44.2 h, i.e. about 22 h of wall-clock across the two lanes (concurrent fits overlap, so the sum is not exclusive GPU service time; the sweep's own elapsed time was not logged). The sweep shared the GPU and ran under CPU contention, so the time axis of the figures below is indicative of relative, not absolute, cost; the iteration axis is the one that carries the convergence claims. The 374 held-out fits ran on the same card, again as two lanes sharing the GPU, with per-fit loop times of 0.85–1,207 s summing to 12.7 h.

#### 2.5 Metrics: training PSNR and the held-out checkpoint sweep

All quality metrics of the $5 \times 11$ sweep are computed in *training mode*: the model is fitted to the full volume and the reconstruction is scored against the same volume. Training-mode PSNR conflates signal recovery with noise memorisation: at high splat counts on noisy volumes the late-iteration "improvement" includes the model fitting voxel-level noise that predicts nothing outside the fitted voxels. Whether, and when, late training gains stop reaching unseen voxels is a question about the iteration axis that only a held-out measurement along that axis can answer, so the sweep was repeated in a blind-spot mode, which this document reports.

**Held-out protocol.** For each volume a fixed 5% of voxels is drawn as the held-out set (`RandomState(42)`, the mask of Supplementary Document 2's blind-spot fits; 0.20–5.33 M voxels per volume). Every held-out voxel is replaced by a *donut-median fill*: the median of its *unmasked* 3×3×3 neighbours with the centre excluded, so the fitted volume is independent of every held-out value. The fit runs on that volume with the settings of § 2.3, with the absolute-error stop set unreachable. The reconstruction is then scored against the *original* volume on two disjoint voxel sets of the same fit: *held-out PSNR* on the masked voxels (what the fit predicts where it saw a fill value) and *training PSNR* on the unmasked voxels (what it reproduces where it saw the data); the *gap* is their difference.

Two splat counts are swept per volume, each over the same eleven budgets as independent fits (seed $K + n$, no replicates): a $4\,K$-splat control, and the volume's operating point $K^*$ from Supplementary Document 2 ($8\,K$–$256\,K$ here) capped at the largest training-sweep count, $256\,K$. The cap binds on *Tribolium* embryo nuclei and Mouse heart nuclei, whose held-out curves are still rising at the largest count Supplementary Document 2 tested (*signal-limited curve*), so their $256\,K$ cells are below their operating point. That gives $17 \times 2 = 34$ held-out cells and 374 fits. A held-out "peak" below is the highest of a cell's eleven held-out scores, and $n^*_{\mathrm{ho}}$ the first checkpoint at or above 99% of it in dB; $n^*_{\mathrm{tr}}$ is the training criterion $n^*$ (§ 3.2) applied to the same fits' unmasked-voxel PSNR. With one fit per checkpoint, differences of a few hundredths of a dB between neighbouring budgets are within the seed-to-seed scatter visible on the flat parts of the curves (§ 3.5) and are not read as trends.

**What the two sweeps can and cannot say.** The "within 0.5 dB of $P_{20\mathrm{K}}$ at 2,000 iterations on 77 of 85 cells" statement (Table 3) and the "$n^* \leq 2{,}000$ on 75 cells" count (Table 4) are training-PSNR statements; the held-out sweep says what happens to unseen voxels over the same budgets. $n^*$ is the first sampled budget at which an independently fitted model reaches 99% of the same cell's 20,000-iteration training PSNR in dB. It is neither a held-out convergence time nor the fitter's stopping rule (§ 6).

In the held-out sweep the 99%-of-peak crossing $n^*_{\mathrm{ho}}$ is at or before $n^*_{\mathrm{tr}}$ on every one of the 34 cells (§ 3.5), so on these volumes the training criterion does not understate the budget the held-out score needs. What it does not capture is the other direction. On Kidney actin at $128\,K$ the held-out PSNR peaks at 30.62 dB after 1,000 iterations and is 30.52 dB at 20,000, while the training PSNR of the same fits climbs to 38.64 dB. That gap of +8.12 dB reaches no held-out voxel and is therefore read as reproduction of the fitted voxels' noise; the sweep does not separate that from the donut fill's own error at the masked voxels. How much of a late *training* rise is signal is therefore read off the held-out curve of the same cell in § 3.5, per volume. The sweep does not decompose it further, and with one fit per cell it bounds the seed-to-seed scatter without measuring it.

$P_{20\mathrm{K}}$ *values* and the capacity-dominates *magnitudes* of the $5 \times 11$ sweep are training-mode numbers: the 20,000-iteration PSNR table (Table 2) and the capacity-vs-iterations deltas (Table 5) are scored on the fitted volume. The held-out sweep tests the same capacity-vs-iterations question at its two counts (operating point at 500 iterations against $4\,K$ splats at 20,000; § 3.5). The ranking holds there under held-out PSNR, by +0.07 to +11.91 dB (mean +5.40 dB), at a different

| Volume | 1 K | 4 K | 16 K | 64 K | 256 K |
|---|---|---|---|---|---|
| OpenCell-MAP4 nuclei | 25.10 | 27.41 | 28.38 | 28.89 | 30.03 |
| OpenCell-MAP4 microtubules | 21.40 | 23.31 | 24.72 | 25.68 | 26.95 |
| OpenCell-LMNB1 nuclei | 22.78 | 24.87 | 25.48 | 25.72 | 26.28 |
| OpenCell-LMNB1 nuclear lamina | 21.39 | 29.59 | 33.39 | 34.60 | 35.66 |
| Kidney nuclei | 22.38 | 26.39 | 32.31 | 38.14 | 42.53 |
| Kidney actin | 20.97 | 23.88 | 28.62 | 34.57 | 44.16 |
| Cells3D nuclei | 31.44 | 34.30 | 36.13 | 37.51 | 40.47 |
| Cells3D membranes | 37.16 | 42.31 | 47.75 | 52.00 | 56.33 |
| Blastocyst nuclear lamina | 32.60 | 36.23 | 38.39 | 40.02 | 41.18 |
| *C. elegans* embryo nuclei | 38.32 | 39.18 | 39.33 | 39.42 | 40.00 |
| Fly brain neurons | 38.10 | 41.43 | 44.09 | 44.88 | 45.28 |
| *Tribolium* embryo nuclei | 29.71 | 38.04 | 47.73 | 52.60 | 54.28 |
| Mouse heart nuclei | 26.34 | 28.47 | 30.67 | 32.89 | 39.89 |
| Zebrafish embryo nuclei | 22.56 | 25.31 | 29.24 | 35.90 | 39.73 |
| *Drosophila* embryo nuclei | 21.08 | 24.31 | 27.98 | 32.47 | 34.83 |
| Neuromast membranes | 33.21 | 36.77 | 48.85 | 56.21 | 57.24 |
| Neuromast nuclei | 35.14 | 43.31 | 59.43 | 66.80 | 67.47 |

Table 2: Training-mode PSNR (dB) at the 20,000-iteration checkpoint, $P_{20K}$, the reference for every convergence statement in this document. These are full-volume fit-vs-target PSNRs and are inflated by noise memorisation at high $K$ on noisy datasets; see § 2.5. The highest training PSNR on each row is at $K = 256$ K on every volume.

high count (the operating point, not 16 K), so those margins are not comparable to Table 5. For the principled signal-recovery splat count on each dataset, refer to the operating points of Supplementary Document 2.

### 3 Results

#### 3.1 Training PSNR at the 20,000-iteration checkpoint

Table 2 reports $P_{20K}$, the training-mode PSNR of the 20,000-iteration cell, for each (dataset, $K$); the curves have flattened well before that budget on most cells (Figure 3), and this single checkpoint is the reference for every convergence statement in the document. These are training-mode values, not signal-recovery values; see § 2.5 for the distinction, § 3.5 for the held-out PSNR along the budget axis at the operating point, and Supplementary Document 2 for the held-out PSNR across splat counts. $P_{20K}$ rises monotonically with $K$ on all 17 volumes. The marginal value of capacity is volume-dependent: Neuromast nuclei gains +32.3 dB and *Tribolium* embryo nuclei +24.6 dB from 1 K to 256 K splats, while *C. elegans* embryo nuclei gains only +1.7 dB and OpenCell-LMNB1 nuclei +3.5 dB over the same range. Volumes where structure is sparse relative to volume (the *C. elegans* embryo, the OpenCell nuclear channels) saturate at low $K$; dense or fine-textured volumes (Kidney actin, *Tribolium* embryo nuclei, Mouse heart nuclei, Zebrafish embryo nuclei) continue to gain. On the noisy confocal volumes, however, training PSNR at the operating point runs +4.14 (Kidney nuclei, 64 $K$) and +8.12 dB (Kidney actin, 128 $K$) above the held-out PSNR of the same fits at 20,000 iterations (§ 3.5), so much of the high-$K$ gain there is noise memorisation.

| Volume | largest $P_{20K} - \text{PSNR}(n)$ over $K$ (dB), with that $K$ $n = 1{,}000$ | $n = 2{,}000$ | $n = 5{,}000$ |
|---|---|---|---|
| OpenCell-MAP4 nuclei | 0.05 (256 K) | 0.04 (4 K) | 0.02 (4 K) |
| OpenCell-MAP4 microtubules | 0.04 (256 K) | 0.01 (256 K) | 0.00 (256 K) |
| OpenCell-LMNB1 nuclei | 0.03 (256 K) | 0.03 (256 K) | 0.02 (256 K) |
| OpenCell-LMNB1 nuclear lamina | 0.14 (256 K) | 0.14 (4 K) | 0.13 (4 K) |
| Kidney nuclei | 1.34 (256 K) | 0.02 (256 K) | 0.01 (64 K) |
| Kidney actin | 4.26 (256 K) | 0.55 (256 K) | 0.06 (64 K) |
| Cells3D nuclei | 0.43 (256 K) | 0.03 (256 K) | 0.14 (256 K) |
| Cells3D membranes | 0.83 (256 K) | 0.09 (4 K) | 0.02 (1 K) |
| Blastocyst nuclear lamina | 0.54 (256 K) | 0.44 (256 K) | 0.27 (256 K) |
| *C. elegans* embryo nuclei | 0.01 (256 K) | 0.00 (16 K) | 0.00 (1 K) |
| Fly brain neurons | 0.02 (256 K) | 0.02 (16 K) | 0.10 (4 K) |
| *Tribolium* embryo nuclei | 1.03 (16 K) | 0.04 (256 K) | 0.17 (4 K) |
| Mouse heart nuclei | 2.16 (256 K) | 0.42 (256 K) | 0.28 (256 K) |
| Zebrafish embryo nuclei | 1.24 (64 K) | 0.51 (64 K) | 0.30 (256 K) |
| *Drosophila* embryo nuclei | 0.28 (1 K) | 0.09 (256 K) | 0.05 (256 K) |
| Neuromast membranes | 4.09 (16 K) | 1.48 (16 K) | 0.62 (64 K) |
| Neuromast nuclei | 3.81 (16 K) | 1.50 (16 K) | 1.08 (16 K) |

Table 3: Shortfall of a shorter budget against the 20,000-iteration reference: for each volume the largest $P_{20K} - \text{PSNR}(n)$ over the five splat counts at $n = 1{,}000$, 2,000 and 5,000, with the splat count at which it occurs. Cells are tinted by the shortfall (white at or below zero, full tint at 2 dB or more). A negative value means the shorter budget scored above the 20,000-iteration cell. Over all 85 cells the shortfall at 2,000 is within 0.5 dB on 77 and within 1 dB on 81.

##### 3.2 Convergence speed

Table 3 reports, per volume, the largest shortfall of a shorter budget against the 20,000-iteration reference, $P_{20K} - \text{PSNR}(n)$ over the five splat counts, at $n = 1{,}000$, 2,000 and 5,000. At 2,000 iterations the shortfall is at most 0.5 dB on 77 of the 85 cells and at most 1 dB on 81 (median 0.00 dB; largest 1.50 dB, Neuromast nuclei at 16 $K$; on 38 cells the 2,000-iteration fit scores above the 20,000 one). At 1,000 iterations 67 cells are within 0.5 dB and 73 within 1 dB (largest 4.26 dB, Kidney actin at 256 $K$); at 5,000, 82 and 84 (largest 1.08 dB, Neuromast nuclei at 16 $K$).

Table 4 reports the same data as a crossing count: the smallest checkpoint $n^*$ at which $\text{PSNR}(n^*) \geq 0.99\, P_{20K}(K)$. The threshold is 99% of the *decibel value*, a chosen tolerance and not a fraction of recovered signal: at $P_{20K} = 60$ dB it admits a 0.6 dB shortfall, i.e. about 15% more MSE than the reference (about 7% at 30 dB), which is why the absolute shortfalls above are the primary statement. $n^*$ is a first crossing by an independently fitted model at a sampled budget: an upper bound on the crossing budget, resolved to the checkpoint grid, not an optimiser stopping time, and by construction every cell crosses at 20,000 at the latest. On 75 of 85 cells, $n^* \leq 2{,}000$; on 63 of 85 cells, $n^* \leq 1{,}000$. First crossing is not the whole story on a noisy volume, where training PSNR can wobble between independent fits: 75 of 85 cells stay at or above 99% of $P_{20K}$ at *every* checkpoint from 2,000 iterations onward (82 from 5,000 onward). The ten cells past the bound, Kidney actin at 256 K, Blastocyst nuclear lamina at 256 K, Mouse heart nuclei at 256 K, Zebrafish embryo nuclei at 64 K, Zebrafish embryo nuclei at 256 K, Neuromast membranes at 16 K, Neuromast membranes at 64 K, Neuromast nuclei at 16 K, Neuromast nuclei at 64 K

| Volume | 1 K | 4 K | 16 K | 64 K | 256 K |
|---|---|---|---|---|---|
| OpenCell-MAP4 nuclei | 1,000 | 500 | 200 | 500 | 1,000 |
| OpenCell-MAP4 microtubules | 1,000 | 500 | 500 | 500 | 1,000 |
| OpenCell-LMNB1 nuclei | 500 | 500 | 200 | 200 | 500 |
| OpenCell-LMNB1 nuclear lamina | 1,000 | 1,000 | 1,000 | 500 | 1,000 |
| Kidney nuclei | 500 | 1,000 | 1,000 | 2,000 | 2,000 |
| Kidney actin | 500 | 1,000 | 1,000 | 2,000 | 3,000 |
| Cells3D nuclei | 1,000 | 500 | 500 | 500 | 2,000 |
| Cells3D membranes | 1,000 | 1,000 | 1,000 | 1,000 | 2,000 |
| Blastocyst nuclear lamina | 1,000 | 1,000 | 500 | 500 | 3,000 |
| *C. elegans* embryo nuclei | 200 | 100 | 50 | 50 | 500 |
| Fly brain neurons | 50 | 200 | 200 | 200 | 200 |
| *Tribolium* embryo nuclei | 200 | 2,000 | 2,000 | 1,000 | 1,000 |
| Mouse heart nuclei | 1,000 | 1,000 | 1,000 | 1,000 | 3,000 |
| Zebrafish embryo nuclei | 2,000 | 1,000 | 2,000 | 5,000 | 3,000 |
| *Drosophila* embryo nuclei | 2,000 | 1,000 | 1,000 | 1,000 | 1,000 |
| Neuromast membranes | 1,000 | 1,000 | 3,000 | 8,000 | 2,000 |
| Neuromast nuclei | 500 | 2,000 | 20,000 | 5,000 | 3,000 |

Table 4: Iterations to reach 99% of $P_{20\mathrm{K}}$: the smallest checkpoint $n^*$ such that $\mathrm{PSNR}(n^*) \geq 0.99\,P_{20\mathrm{K}}$, the PSNR of the same cell's 20,000-iteration fit (99% of the dB value, a chosen tolerance: a 0.6 dB shortfall, $\approx$ 15% more MSE, at 60 dB). 75 of 85 cells first cross by iteration 2,000 and the remaining ten at six at 3,000, two at 5,000, one at 8,000 and one at 20,000; 63 of 85 cross by iteration 1,000; 75 of 85 remain at or above the threshold at every checkpoint from 2,000 onward. Table 3 gives the shortfalls in dB.

and Neuromast nuclei at 256 K, cross at six at 3,000, two at 5,000, one at 8,000 and one at 20,000; the one at 20,000 (Neuromast nuclei at 16 K) reaches 99% of its own 20,000-iteration value at no earlier checkpoint, so for that cell the crossing is the reference itself. The next tier, twelve cells at 2,000 iterations distributed across Kidney nuclei, Kidney actin, Cells3D nuclei, Cells3D membranes, *Tribolium* embryo nuclei, Zebrafish embryo nuclei, *Drosophila* embryo nuclei, Neuromast membranes and Neuromast nuclei, mixes modalities (five confocal, five light-sheet and two iSIM cells) and capacities (six of them at 64 K or 256 K). The fastest cells are *C. elegans* embryo nuclei at $K \in \{16\,K, 64\,K\}$ and Fly brain neurons at $K \in \{1\,K\}$, which already exceed 99% of $P_{20\mathrm{K}}$ at iteration 50 (the first checkpoint).

##### 3.3 Capacity dominates iterations

A direct comparison demonstrates that adding capacity is more effective than adding optimisation budget. Table 5 and Figure 2 contrast two cells: 4 K splats fitted for 20,000 iterations, versus 16 K splats fitted for only 500 iterations. The latter has 4× the model capacity but 40× fewer optimisation steps. The two are *not* cost-matched: the 16 K-splat, 500-iteration fit performs 10× fewer splat-iterations (splats times iterations, the cost figure to quote), and its optimisation-loop time was 11–52× shorter (median 36×; time columns in the table), a ratio measured as two concurrent lanes on the GPU of § 2.4 shared with unrelated jobs, so it is indicative only. On every volume in the sweep, the 16 K-splat, 500-iteration fit nevertheless achieves *higher* training PSNR than the 4 K-splat, 20,000-iteration one (by +0.12 to +7.77 dB, mean +3.03 dB), so the larger,

| Volume | 4 K @ 20,000 iters PSNR (dB) | time (s) | 16 K @ 500 iters PSNR (dB) | time (s) | Δ (dB) | speed-up |
|---|---|---|---|---|---|---|
| OpenCell-MAP4 nuclei | 27.41 | 237 | 28.37 | 21 | +0.96 | 11× |
| OpenCell-MAP4 microtubules | 23.31 | 596 | 24.62 | 14 | +1.31 | 43× |
| OpenCell-LMNB1 nuclei | 24.87 | 423 | 25.46 | 12 | +0.59 | 36× |
| OpenCell-LMNB1 nuclear lamina | 29.59 | 632 | 32.81 | 17 | +3.22 | 36× |
| Kidney nuclei | 26.39 | 438 | 31.09 | 11 | +4.70 | 38× |
| Kidney actin | 23.88 | 276 | 27.60 | 7 | +3.72 | 38× |
| Cells3D nuclei | 34.30 | 421 | 36.07 | 12 | +1.77 | 36× |
| Cells3D membranes | 42.31 | 240 | 46.79 | 18 | +4.49 | 13× |
| Blastocyst nuclear lamina | 36.23 | 647 | 38.03 | 14 | +1.79 | 46× |
| *C. elegans* embryo nuclei | 39.18 | 433 | 39.30 | 12 | +0.12 | 38× |
| Fly brain neurons | 41.43 | 2,359 | 44.02 | 76 | +2.59 | 31× |
| *Tribolium* embryo nuclei | 38.04 | 732 | 44.06 | 20 | +6.02 | 36× |
| Mouse heart nuclei | 28.47 | 520 | 30.01 | 15 | +1.54 | 35× |
| Zebrafish embryo nuclei | 25.31 | 326 | 27.68 | 9 | +2.37 | 35× |
| *Drosophila* embryo nuclei | 24.31 | 2,155 | 27.44 | 41 | +3.13 | 52× |
| Neuromast membranes | 36.77 | 1,118 | 42.29 | 32 | +5.52 | 35× |
| Neuromast nuclei | 43.31 | 581 | 51.08 | 16 | +7.77 | 37× |

Table 5: Adding capacity beats adding iterations. Training-mode PSNR (dB) and optimisation-loop time (s; § 2.3) of the single fit for two cells: 4 K splats trained for 20,000 iterations versus 16 K splats trained for only 500 iterations; speed-up is the ratio of the two times. The 16 K-splat, 500-iteration cell wins on every volume, with gains of +0.12 dB (*C. elegans* embryo nuclei) to +7.77 dB (Neuromast nuclei); mean gain +3.03 dB, at 10× fewer splat-iterations; its loop time was 11–52× shorter (median 36×) on the shared GPU of § 2.4, an indicative range. These are training-mode scores; the held-out sweep repeats the comparison at its own two counts (operating point at 500 iterations vs. 4 $K$ splats at 20,000) and the larger, shorter fit wins in held-out PSNR on every volume there too (Table 6, § 3.5).

shorter fit is both better and cheaper. This is a dominance result for these two configurations; no matched-time frontier or cost-scaling law across $K$ is computed here.

##### 3.4 Cross-dataset summary

Figures 3 and 4 summarise the entire sweep in two complementary views. The 6×3 grid of Figure 3 (17 panels used) shows PSNR-vs-iterations small-multiples for every volume, one panel per volume, five coloured curves per panel (one per splat count), with the 2,000-iteration budget and each curve's 0.5 dB shortfall band drawn as guides. Figure 4 plots PSNR against optimisation-loop time for all 935 fits as small multiples by splat count: one panel per count, one curve per volume (85 (volume, $K$) groups, each connected in time order) and the cross-volume median in bold. Every fit is drawn (dominated points are not filtered out), and PSNRs of different target volumes are not interchangeable objective values, so the panels are not a Pareto frontier.

Three patterns are visible at a glance: (i) the curves in the small-multiple panels flatten by $n \approx$ 1,000–2,000 on most volumes (the neuromast channels keep rising slowly: up to +1.50 dB from 2,000 to 20,000 iterations on Neuromast nuclei at 16 $K$). (ii) Within each panel the five curves are *stacked vertically* by splat count instead of fanning out along iterations, visualising the same finding as Table 5. (iii) In Figure 4 every panel is a set of horizontal "shelves" (each splat

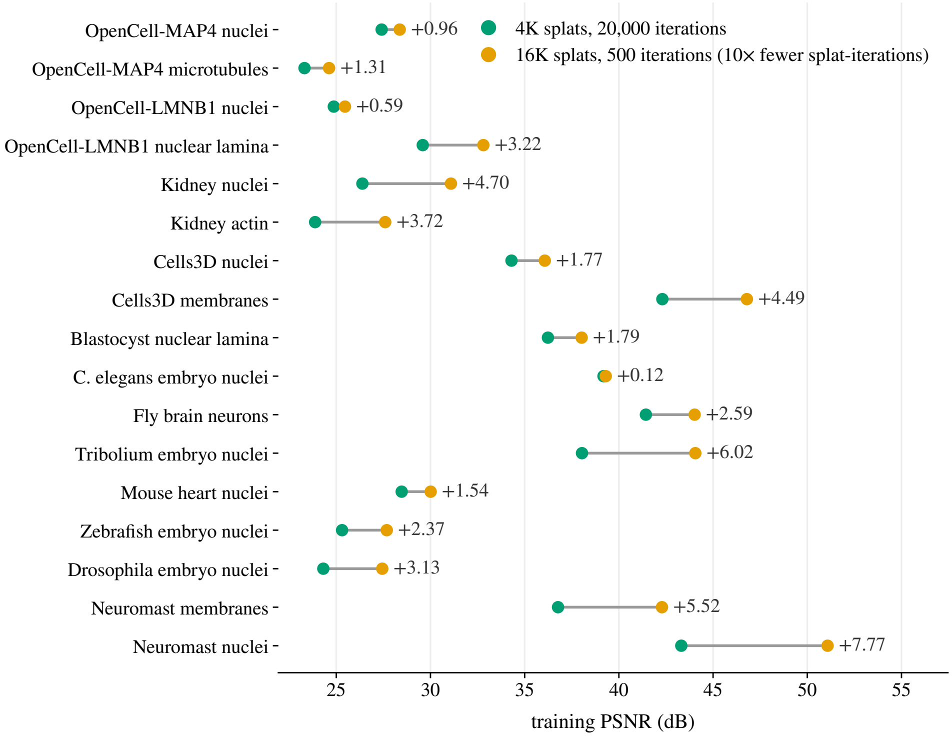


Figure 2: Capacity against iterations, per volume: training PSNR of the 4 K-splat fit at 20,000 iterations and of the 16 K-splat fit at 500 iterations, joined per volume, with the gain of the larger, shorter fit written at its marker. The larger, shorter fit performs 10× fewer splat-iterations and scores higher on every row. Each pair is the corresponding row of Table 5.

count defines a roughly time-independent PSNR ceiling per volume), and the shelves, read across panels, are spaced widely in PSNR by $K$ but move little in time within a $K$.

##### 3.5 Held-out convergence at the operating point

Tables 6 and 7 and Figure 6 report the blind-spot sweep of § 2.5: held-out PSNR (masked voxels) and training PSNR (unmasked voxels) of the same fits at eleven requested budgets, at the Supplementary Document 2 operating point (capped at 256 $K$) and at the 4 $K$-splat control.

**At the operating point.** Held-out PSNR peaks at a requested budget between 500 and 20,000: by 1,000 iterations on 5 of 17 volumes, by 2,000 on 8, by 5,000 on 12; two cells, *Tribolium* embryo nuclei at 256 $K$ and Zebrafish embryo nuclei at 256 $K$, are still rising at 20,000, both light-sheet volumes at the sweep's largest count. The 99%-of-peak crossing $n^*_{\text{ho}}$ is never later than the training crossing $n^*_{\text{tr}}$ of the same fits (17 of 17 cells; strictly earlier on 10), and is at most 2,000.

One caveat governs everything that follows. The sweep has no replicate seeds, so the peak *iteration* on a cell whose curve is flat over a decade of budgets is located only to within that flat range, and single-fit differences of a few hundredths of a dB are not read as trends. A decline is read as real only where it repeats: the five consecutive declines across independently seeded fits on the neuromast channels below are more than seed scatter. With that caveat, what happens after the peak separates three regimes (Figure 5 sketches them; Figure 6 has the measured curves).

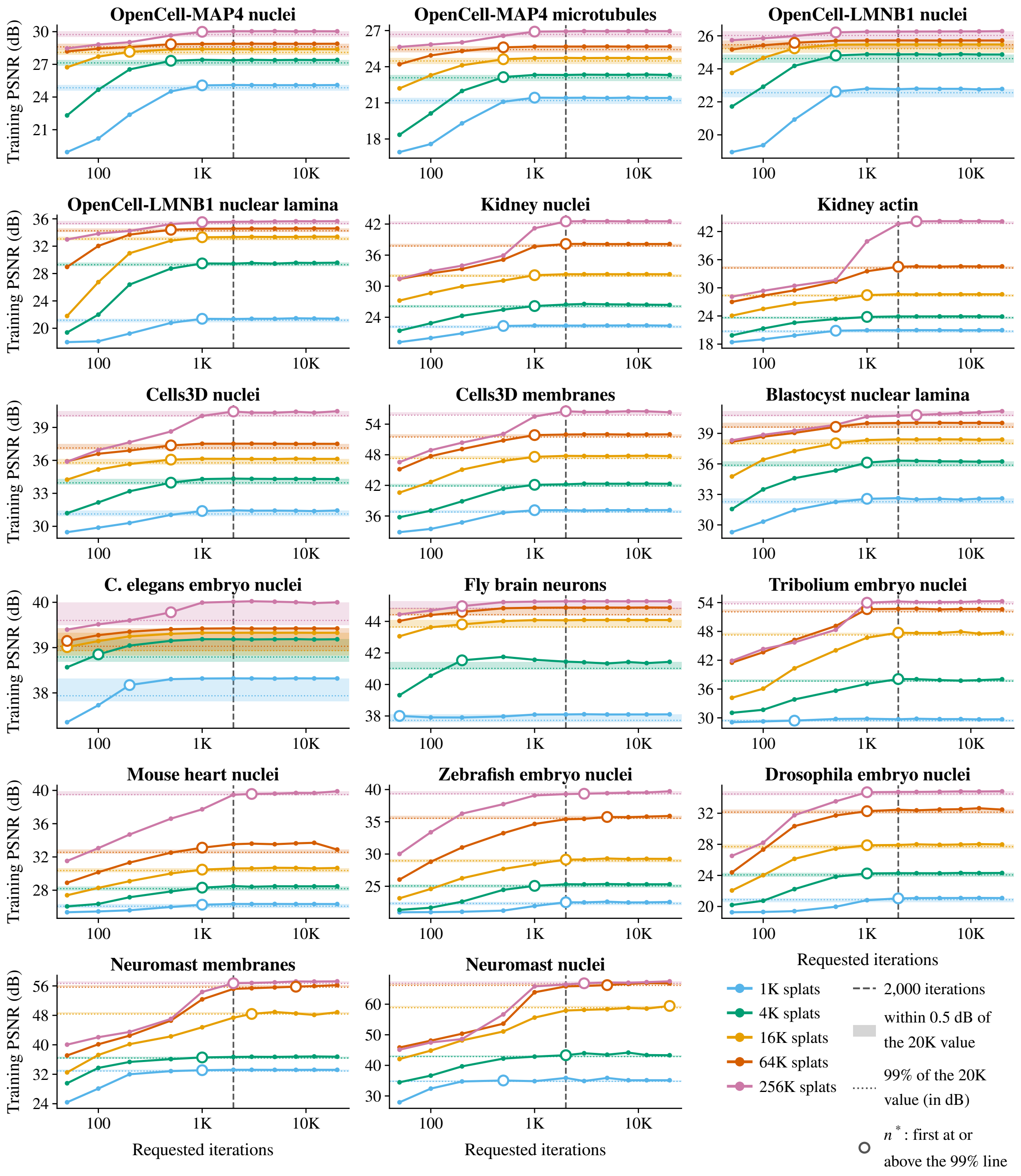


Figure 3: Cross-dataset convergence summary: 6×3 grid (17 panels) of training PSNR vs. requested iterations, one panel per volume (named as in Table 1), five splat-count colours per panel. The dashed line marks the 2,000-iteration budget; the band in each curve's colour spans from 0.5 dB below that curve's 20,000-iteration value up to it, so a curve inside its band at the dashed line is a cell that Table 3 counts as within 0.5 dB at 2,000. Dotted, in each curve's colour: 99% of that curve's 20,000-iteration value in dB; hollow marker: its first crossing $n^*$ (Table 4). Training PSNR reaches 99% of its 20,000-iteration value by $n \approx$ 1,000–2,000 on most cells (the ten slower cells, Kidney actin at 256 K, Blastocyst nuclear lamina at 256 K, Mouse heart nuclei at 256 K, Zebrafish embryo nuclei at 64 K, Zebrafish embryo nuclei at 256 K, Neuromast membranes at 16 K, Neuromast membranes at 64 K, Neuromast nuclei at 16 K, Neuromast nuclei at 64 K and Neuromast nuclei at 256 K, later).

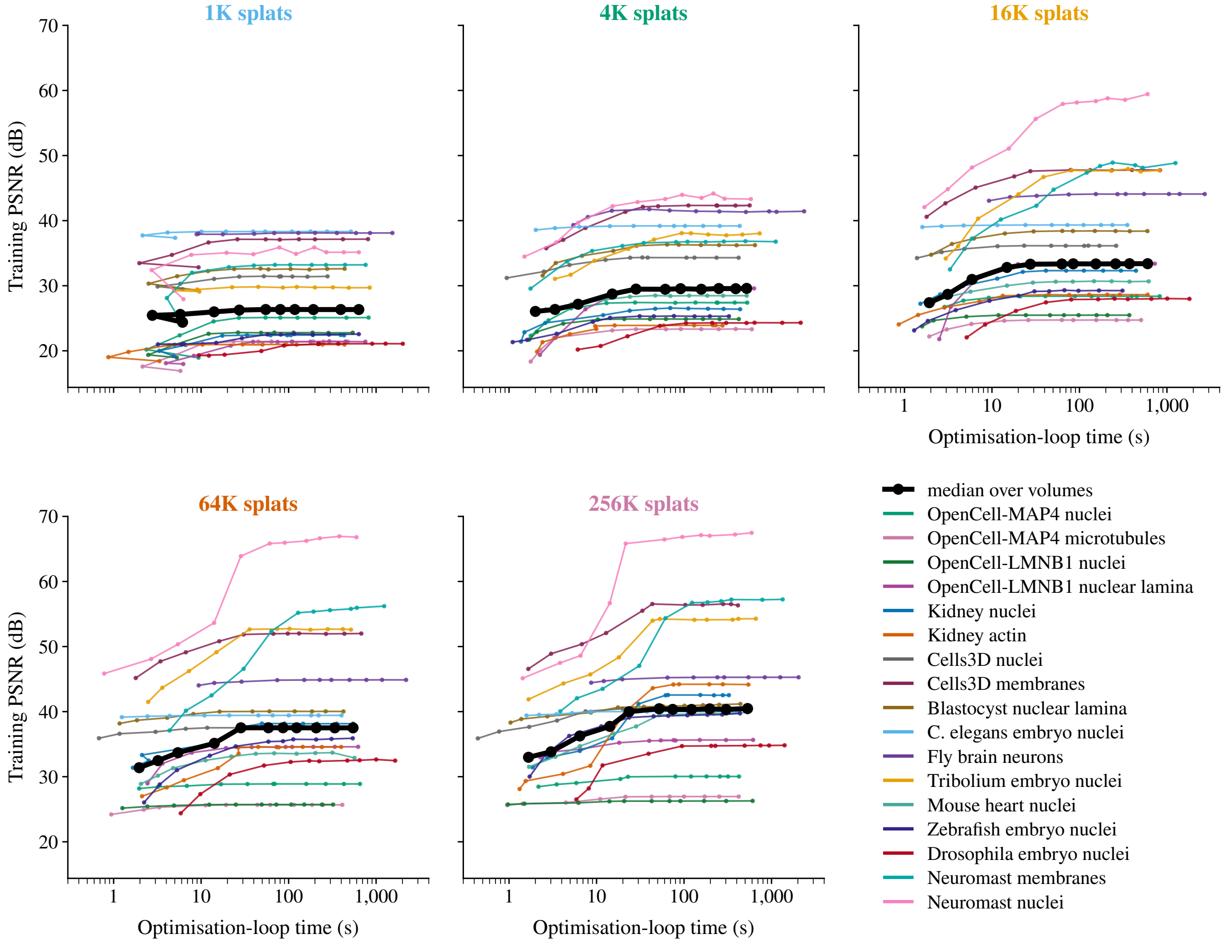


Figure 4: Training PSNR vs. optimisation-loop time for all 935 fits (17 volumes × 5 splat counts × 11 iter checkpoints; every fit drawn, no non-dominance filtering), as small multiples by splat count: one panel per count, one thin curve per volume (colour by volume, the eleven checkpoints as markers connected in time order) and, in bold black, the cross-volume median (median time and median PSNR at each requested budget). The panels share both axes. Within a panel the curves flatten early in loop time; across panels the level rises with capacity, not with further optimisation. Time was measured on a shared, contended GPU (§ 2.4) and is indicative only.

*Flat held-out, flat training* (the four OpenCell channels, *C. elegans*, Fly brain, and *Tribolium*, Mouse heart and *Drosophila* at 256 $K$). Both curves reach their level within the first few thousand iterations and stay there. The held-out score at 20,000 is within 0.1 dB of its peak and the train/held-out gap at 20,000 is below 1 dB on all of them, a few tenths of a dB on the two signal-limited light-sheet cells (+0.37 on *Tribolium*, +0.28 on Mouse heart); 5 of the 17 operating-point cells have a gap $\leq 0.5$ dB.

*Flat held-out, rising training* (Kidney nuclei 64 $K$, Kidney actin 128 $K$, Cells3D, Blastocyst, Zebrafish 256 $K$). Held-out PSNR reaches its peak by 5,000 iterations on all of them but Zebrafish and then changes by about 0.1 dB or less to 20,000; the largest such change, Kidney actin's, is 0.104 dB to three decimals, 0.10 dB as rounded in Table 6 (peak 30.62 at 1,000, 30.52 at 20,000). Training PSNR meanwhile keeps rising, to a gap of +4.14 (Kidney nuclei) and +8.12 dB (Kidney actin) at 20,000; the gap exceeds 1 dB on 8 of the 17 operating-point cells (Kidney nuclei, Kidney

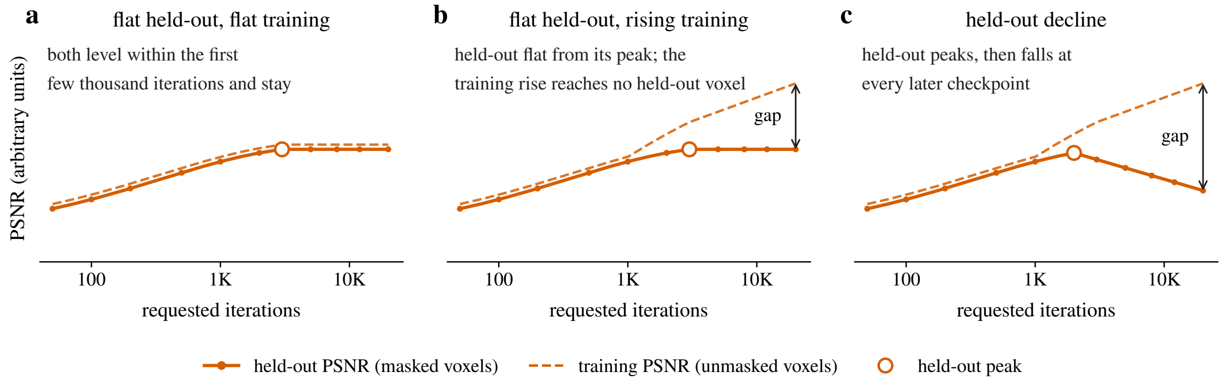


Figure 5: The three held-out regimes (schematic; idealised curves, not data). Held-out PSNR (solid) and training PSNR (dashed) of the same blind-spot fits against requested iterations, hollow marker on the held-out peak. (a) Flat held-out, flat training: both reach their level within the first few thousand iterations and stay there. (b) Flat held-out, rising training: the held-out score is flat from its peak while training keeps rising, so a gap opens that reaches no held-out voxel. (c) Held-out decline: the held-out score peaks and then falls at every later checkpoint while training rises. Figure 6 shows the measured curves.

actin, Cells3D nuclei, Cells3D membranes, Blastocyst nuclear lamina, Zebrafish embryo nuclei, Neuromast membranes and Neuromast nuclei). On these cells every training gain after the held-out peak is reproduction of the fitted voxels only. Zebrafish is the border case: its held-out score is still creeping up at 20,000 (the peak) while its training PSNR rises faster, so its gap (+1.89 dB) opens without a held-out decline.

*Held-out decline* (the two iSIM neuromast channels). Both peak at 2,000 requested iterations (63.52 dB at 64 $K$; 51.74 dB at 128 $K$) and then fall at each of the five later checkpoints, to 62.31 and 50.33 dB at 20,000 (losses of 1.21 and 1.42 dB), while training PSNR rises to 66.89 and 57.06 dB. These are the only two cells whose held-out drop from peak to 20,000 exceeds 0.5 dB; the median drop over the 17 operating-point cells is 0.02 dB and 14 of them stay within 0.1 dB. On the neuromasts the late training rise that makes them the slowest cells of the training sweep (§ 3.2) is harmful on held-out voxels, not just unproductive: the 5,000-iteration state is already 0.77 and 0.83 dB below the 2,000-iteration one in held-out PSNR.

**At 4 $K$ splats.** Held-out and training PSNR track each other: the gap at 20,000 is at most 0.33 dB (Cells3D membranes), the held-out score at 20,000 is within 0.1 dB of its peak on 13 of 17 volumes. The largest drop from peak is 0.47 dB on Fly brain neurons, whose 4 $K$ curve peaks at 500 iterations and whose *training* PSNR falls by a similar amount (0.42 dB), a seed-to-seed wobble of a fit that has nothing left to learn at this capacity, not overfitting. The held-out peak lands at 5,000 iterations or later on 14 of 17 volumes (5 at 20,000), i.e. the flat tail wanders by hundredths of a dB and the argmax falls wherever the wobble does. Under-capacity fits do not overfit within 20,000 iterations on any of these volumes.

**Budget checkpoints under held-out PSNR.** At the operating point the 2,000-iteration state is within 0.2 dB of the held-out peak on all 17 volumes (largest shortfall 0.16 dB, Mouse

| Volume | $K$ | 99% crossing | | held-out | held-out PSNR (dB) | | | training PSNR (dB) | |
|---|---|---|---|---|---|---|---|---|---|
| | | training | held-out | peak at | peak | at 20,000 | change | at 20,000 | gap |
| OpenCell-MAP4 nuclei | 16 K | 200 | 200 | 3,000 | 28.13 | 28.12 | −0.01 | 28.39 | +0.26 |
| OpenCell-MAP4 microtubules | 64 K | 500 | 200 | 1,000 | 24.87 | 24.86 | −0.01 | 25.68 | +0.82 |
| OpenCell-LMNB1 nuclei | 32 K | 200 | 200 | 3,000 | 25.43 | 25.43 | +0.00 | 25.60 | +0.17 |
| OpenCell-LMNB1 nuclear lamina | 64 K | 500 | 500 | 12,000 | 33.98 | 33.97 | −0.01 | 34.60 | +0.63 |
| Kidney nuclei | 64 K | 2,000 | 1,000 | 2,000 | 33.86 | 33.81 | −0.06 | 37.95 | +4.14 |
| Kidney actin | 128 K | 2,000 | 1,000 | 1,000 | 30.62 | 30.52 | −0.10 | 38.64 | +8.12 |
| Cells3D nuclei | 32 K | 500 | 100 | 500 | 35.45 | 35.43 | −0.02 | 36.77 | +1.34 |
| Cells3D membranes | 32 K | 1,000 | 500 | 5,000 | 46.74 | 46.66 | −0.09 | 49.86 | +3.20 |
| Blastocyst nuclear lamina | 32 K | 500 | 200 | 1,000 | 37.58 | 37.55 | −0.03 | 39.19 | +1.63 |
| *C. elegans* embryo nuclei | 8 K | 100 | 100 | 8,000 | 39.22 | 39.22 | +0.00 | 39.29 | +0.07 |
| Fly brain neurons | 64 K | 200 | 100 | 5,000 | 44.24 | 44.23 | −0.01 | 44.85 | +0.61 |
| *Tribolium* embryo nuclei | 256 K | 1,000 | 1,000 | 20,000 | 53.87 | 53.87 | +0.00 | 54.24 | +0.37 |
| Mouse heart nuclei | 256 K | 2,000 | 2,000 | 8,000 | 39.45 | 39.39 | −0.05 | 39.67 | +0.28 |
| Zebrafish embryo nuclei | 256 K | 3,000 | 1,000 | 20,000 | 37.77 | 37.77 | +0.00 | 39.66 | +1.89 |
| *Drosophila* embryo nuclei | 256 K | 1,000 | 1,000 | 1,000 | 34.11 | 34.01 | −0.09 | 34.83 | +0.81 |
| Neuromast membranes | 128 K | 5,000 | 1,000 | 2,000 | 51.74 | 50.33 | −1.42 | 57.06 | +6.73 |
| Neuromast nuclei | 64 K | 5,000 | 1,000 | 2,000 | 63.52 | 62.31 | −1.21 | 66.89 | +4.58 |

Table 6: Held-out checkpoint sweep at the operating point ($K$ = the Supplementary Document 2 blind-spot operating point, capped at 256 $K$). 99% crossing: the first checkpoint at $\geq$ 99% of the 20,000-iteration training (unmasked-voxel) PSNR of these fits ($n^*_{\mathrm{tr}}$) and the first checkpoint at $\geq$ 99% of the held-out peak in dB ($n^*_{\mathrm{ho}}$); held-out peak at: the checkpoint of the highest held-out PSNR. Held-out PSNR: the peak, the value at 20,000 and their difference (change). Training PSNR: the value at 20,000 and the gap, training minus held-out at 20,000, same fit. One fit per checkpoint.

heart nuclei); at 4 $K$ splats on 15 of 17 (largest 0.54 dB, Fly brain neurons, the 500-iteration peak above). At the 5,000 and 8,000 checkpoints the held-out score differs from the 20,000 one by at most 0.59 and 0.31 dB (both Neuromast membranes, where the shorter budget is the better one) at the operating point and, at the 5,000 checkpoint, by at most 0.11 dB at 4 $K$; the 5,000 state is at or above the 20,000 state on 11 of the 17 operating-point cells. At 500 iterations the held-out shortfall from the peak is a mean 0.7 dB at 4 $K$ splats (at most 2.3, *Tribolium* embryo nuclei) and 1.5 dB at the operating point (at most 8.6, Neuromast nuclei), with 6 and 9 of 17 volumes within 0.5 dB.

**Capacity vs. iterations, held-out.** The operating-point fit at 500 iterations beats the 4 $K$-splat fit at 20,000 in held-out PSNR on every volume, by +0.07 (*C. elegans* embryo nuclei, whose operating point is only 2× the control) to +11.91 dB (Neuromast nuclei), mean +5.40 dB. The capacity ratio varies from 2× to 64× across volumes and the cells are not cost-matched, so this is the held-out counterpart of the dominance result of § 3.3, not a scaling law.

##### 3.6 Per-volume SSIM and time curves

Figures 7 and 8 display the remaining budget-response curves of the training sweep for every volume, one panel per volume as in Figure 3: training SSIM vs. requested iterations (log $x$; the $y$ range is per panel and spans at least 0.2, so a flat curve is not stretched into apparent noise), and

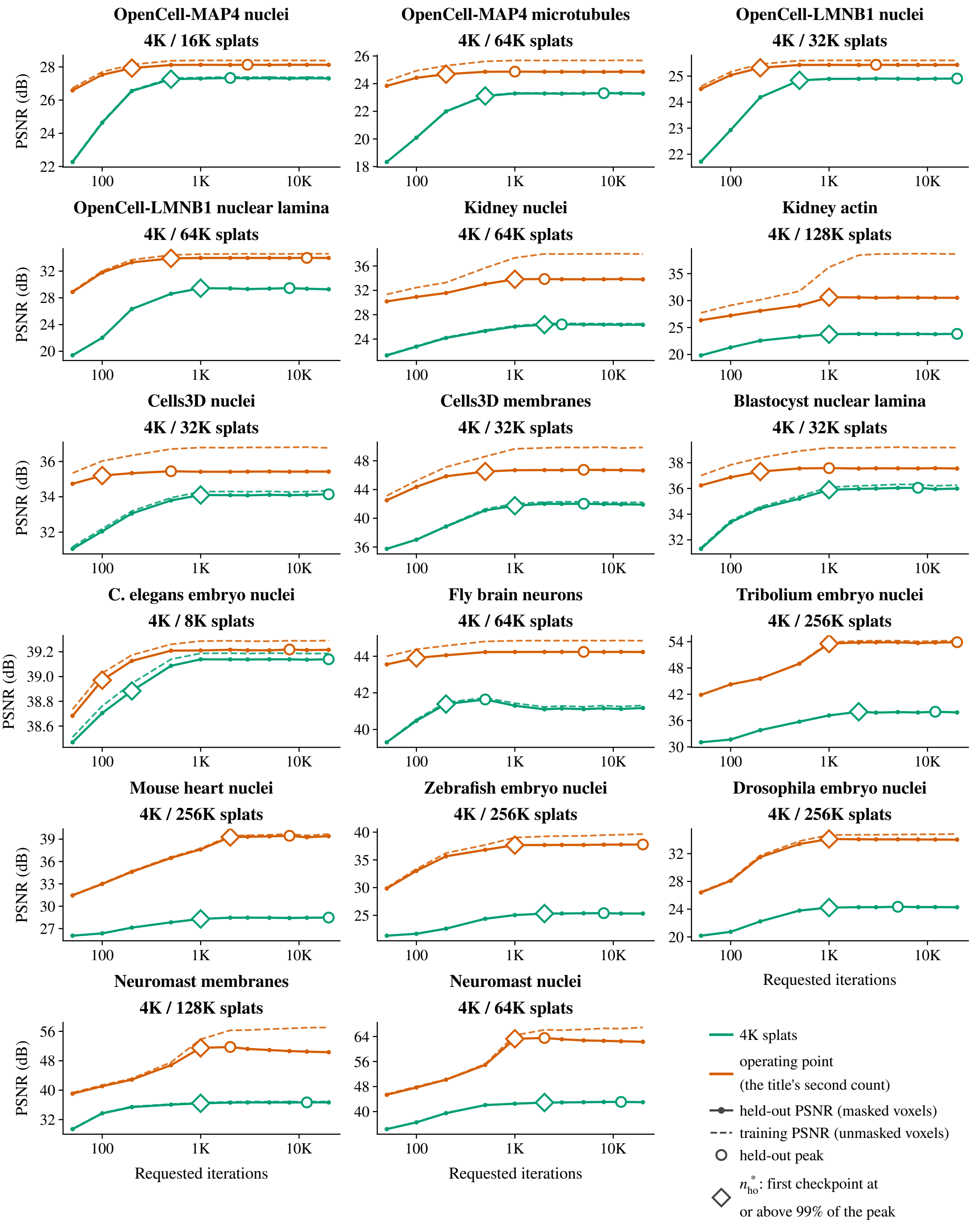


Figure 6: Held-out checkpoint sweep, one panel per volume (title: the volume and its two splat counts, the $4\,K$ control first). Solid: held-out PSNR at the masked voxels; dashed: training PSNR at the unmasked voxels of the same blind-spot fits; hollow circle: held-out peak; hollow diamond: $n^*_{\mathrm{ho}}$, the first checkpoint at or above 99% of it in dB (Tables 6 and 7). Green: $4\,K$ splats; orange: the operating point. Where the two curves of a colour separate, later training gains reach no held-out voxel; on the two neuromast channels the held-out curve falls after its 2,000-iteration peak.

| Volume | 99% crossing | | held-out | held-out PSNR (dB) | | | training PSNR (dB) | |
|---|---|---|---|---|---|---|---|---|
| | training | held-out | peak at | peak | at 20,000 | change | at 20,000 | gap |
| OpenCell-MAP4 nuclei | 500 | 500 | 2,000 | 27.33 | 27.31 | −0.03 | 27.38 | +0.07 |
| OpenCell-MAP4 microtubules | 500 | 500 | 8,000 | 23.31 | 23.28 | −0.04 | 23.31 | +0.03 |
| OpenCell-LMNB1 nuclei | 500 | 500 | 20,000 | 24.91 | 24.91 | +0.00 | 24.91 | +0.00 |
| OpenCell-LMNB1 nuclear lamina | 1,000 | 1,000 | 8,000 | 29.46 | 29.28 | −0.19 | 29.26 | −0.02 |
| Kidney nuclei | 2,000 | 2,000 | 3,000 | 26.41 | 26.34 | −0.06 | 26.51 | +0.17 |
| Kidney actin | 1,000 | 1,000 | 20,000 | 23.82 | 23.82 | +0.00 | 23.88 | +0.05 |
| Cells3D nuclei | 1,000 | 1,000 | 20,000 | 34.14 | 34.14 | +0.00 | 34.34 | +0.20 |
| Cells3D membranes | 1,000 | 1,000 | 5,000 | 42.02 | 41.91 | −0.11 | 42.23 | +0.33 |
| Blastocyst nuclear lamina | 1,000 | 1,000 | 8,000 | 36.05 | 35.99 | −0.06 | 36.26 | +0.27 |
| *C. elegans* embryo nuclei | 200 | 200 | 20,000 | 39.14 | 39.14 | +0.00 | 39.19 | +0.05 |
| Fly brain neurons | 200 | 200 | 500 | 41.64 | 41.17 | −0.47 | 41.31 | +0.14 |
| *Tribolium* embryo nuclei | 2,000 | 2,000 | 12,000 | 38.01 | 37.86 | −0.15 | 37.88 | +0.01 |
| Mouse heart nuclei | 1,000 | 1,000 | 20,000 | 28.48 | 28.48 | +0.00 | 28.48 | +0.00 |
| Zebrafish embryo nuclei | 2,000 | 2,000 | 8,000 | 25.40 | 25.33 | −0.07 | 25.34 | +0.01 |
| *Drosophila* embryo nuclei | 1,000 | 1,000 | 5,000 | 24.35 | 24.28 | −0.07 | 24.30 | +0.02 |
| Neuromast membranes | 1,000 | 1,000 | 12,000 | 36.68 | 36.68 | +0.00 | 36.89 | +0.21 |
| Neuromast nuclei | 2,000 | 2,000 | 12,000 | 43.09 | 43.00 | −0.09 | 43.01 | +0.01 |

Table 7: Held-out checkpoint sweep at the $4\,K$-splat control; columns as in Table 6.

training PSNR vs. optimisation-loop time (log $x$, shared across panels; a hollow circle sits on the 2,000-iteration checkpoint of each curve). Both grids also mark each curve's $n^*$ (Table 4) with a hollow marker, so the crossing can be read in SSIM and in seconds.

### 4 Cross-dataset complexity correlation

A natural question is whether convergence speed correlates with dataset complexity ("complex datasets need more iterations to converge"). Voxel count is not a usable predictor: averaging $n^*$ (iterations to 99% of $P_{20\text{K}}$) across the five splat counts and correlating against $\log_{10}$(voxels) gives Pearson $r \approx +0.09$ ($R^2 \approx 1\%$, $p = 0.729$, 95% CI $[-0.41, +0.55]$) on the 17 volumes. No association is detected in this sample, which is not the same as showing that none exists: the interval admits moderate correlations of either sign. $P_{20\text{K}}$ at $K = 256\,\text{K}$, which one might read as a measure of how much the fit can still extract (or memorise) from the volume, correlates moderately ($r \approx +0.70$, $R^2 \approx 49\%$, $p = 0.00168$, CI $[+0.33, +0.88]$, $n = 17$). That nominal test treats the 17 channels as independent, but they come from 12 acquisitions (five of them contribute two channels each). Averaging paired channels within their acquisition gives $r \approx +0.61$ ($p = 0.0359$, 12 acquisitions). The estimate leans on the slowest volume: without Neuromast nuclei it is $r \approx +0.49$ ($p = 0.0555$), and without both neuromast channels $r \approx +0.28$ ($p = 0.315$, 15 volumes). The response also depends on $P_{20\text{K}}$ through the relative-dB threshold itself. We therefore read that association as exploratory and specific to this sample: it points at the training metric, not at data size, and neither covariate measures structural complexity directly.

A linear regression

$$\hat{n}^*(D) = a + b \cdot \log_{10}(\text{voxels}(D))$$

fits to $\hat{n}^* \approx -336 + 242\,\log_{10}(\text{voxels})$, i.e. a $10\times$ increase in voxel count predicts on the order of

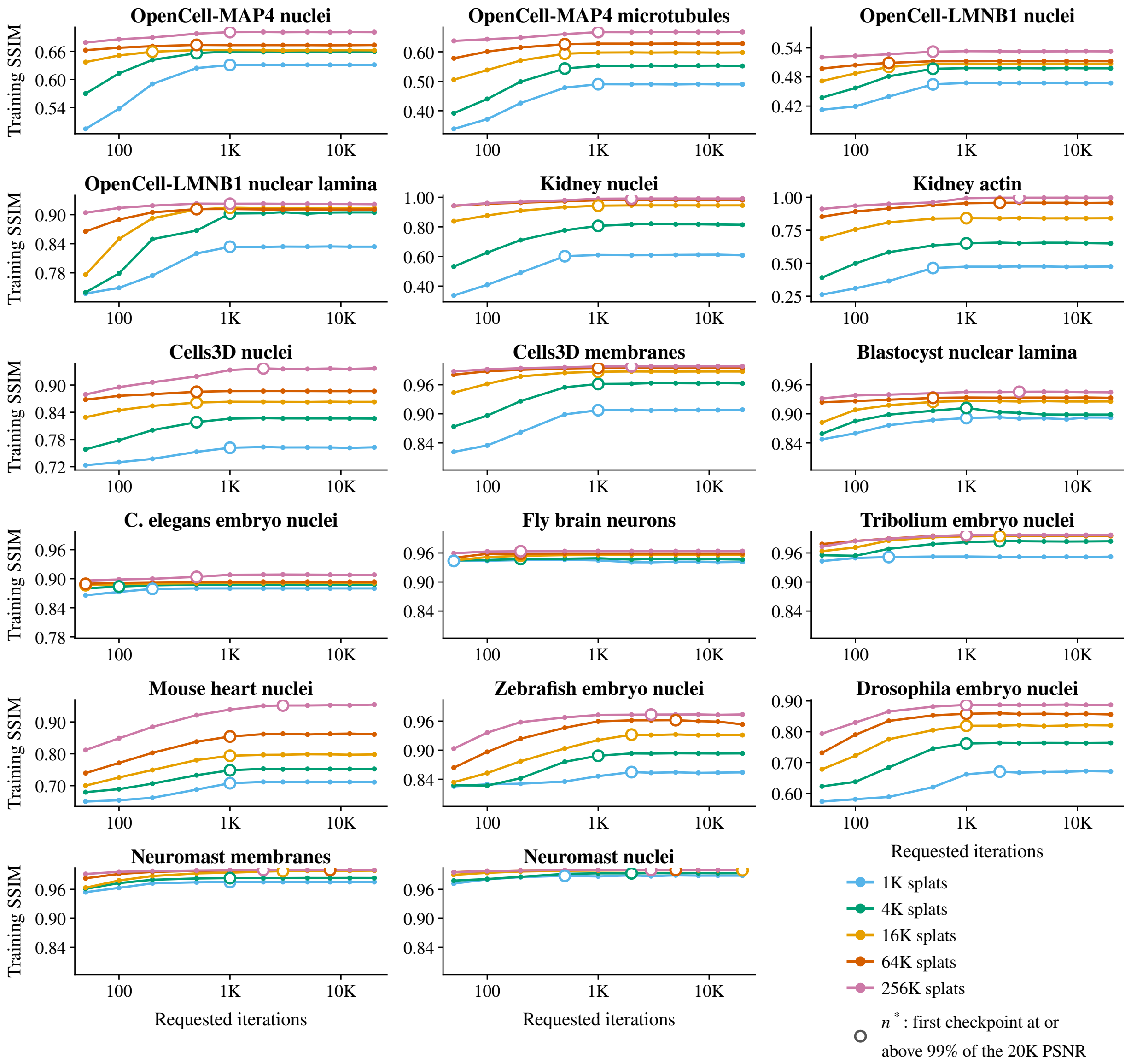


Figure 7: Training SSIM vs. requested iterations, one panel per volume, five splat-count colours as in Figure 3; hollow marker: the curve's $n^*$ (Table 4). The $y$ range is per panel and spans at least 0.2 SSIM. SSIM stacks by splat count and flattens within the first few thousand iterations, as PSNR does in Figure 3.

$\sim$242 *additional* iterations, less than the spacing between adjacent checkpoints anywhere above $n = 500$, and not distinguishable from zero on 17 volumes. Across the 3.9 M–106.7 M voxel range covered here ($\Delta \log_{10} \approx 1.4$) the regression predicts a span of $\sim$347 iterations.

The endpoints make the point directly. Ranked by mean $n^*$ across the five splat counts, the slowest volumes are Neuromast nuclei (28.1 M voxels, $\bar{n}^* = 6{,}100$), then Neuromast membranes (28.1 M voxels, $\bar{n}^* = 3{,}000$), then Zebrafish embryo nuclei (106.7 M voxels, $\bar{n}^* = 2{,}600$), then Kidney actin (4.2 M voxels, $\bar{n}^* = 1{,}500$), then Mouse heart nuclei (99.7 M voxels, $\bar{n}^* = 1{,}400$); the fastest is Fly brain neurons (99.4 M, $\bar{n}^* = 170$) and the second fastest *C. elegans* embryo nuclei (10.7 M, $\bar{n}^* = 180$). Small and large volumes sit at both ends. What separates the slow cells from the fast ones is not how many voxels there are but what the fit is doing late in

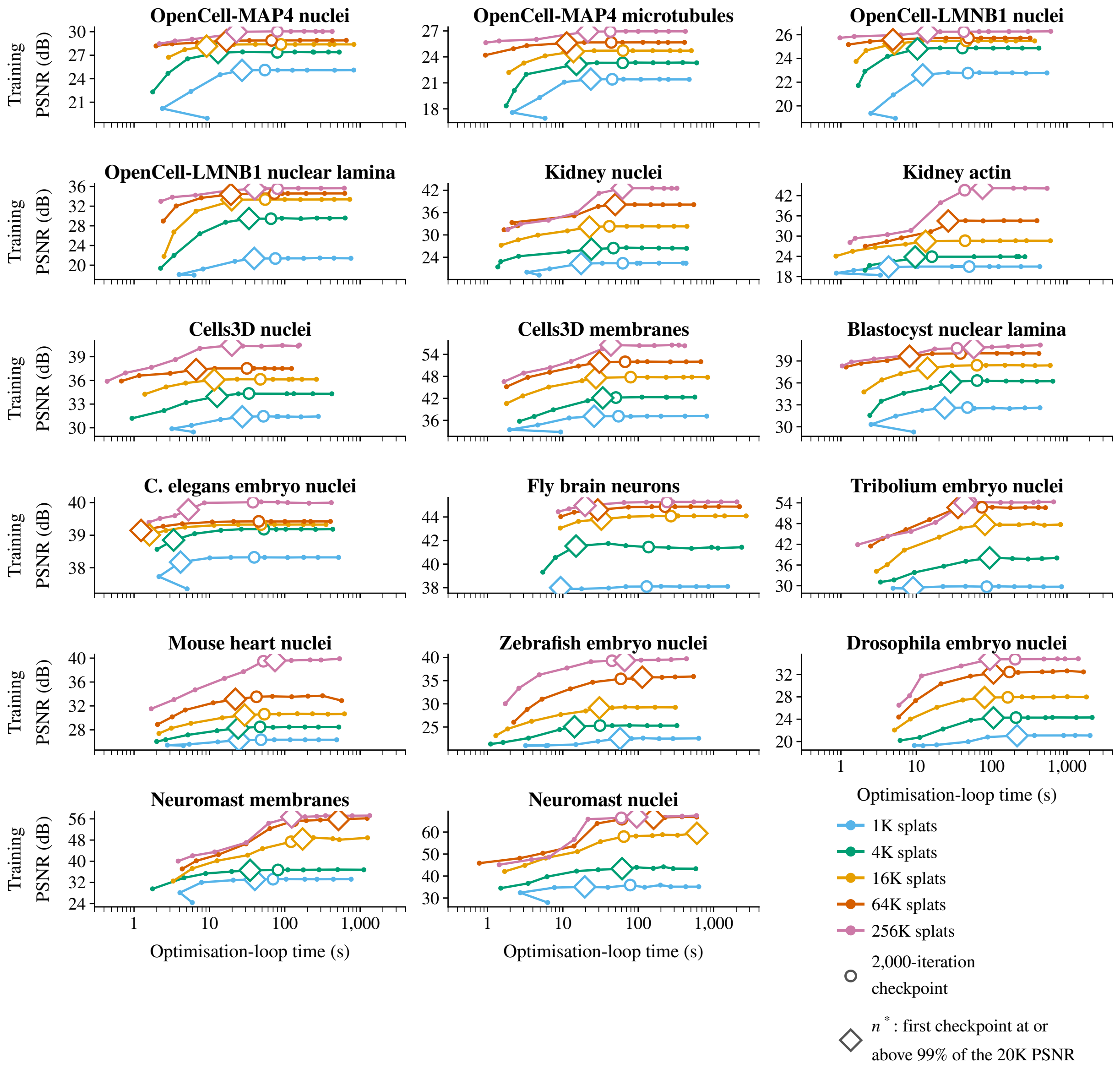


Figure 8: Training PSNR vs. optimisation-loop time (log $x$, shared across panels), one panel per volume, five splat-count colours as in Figure 3; hollow circle: the 2,000-iteration checkpoint; hollow diamond: $n^*$ (Table 4). Loop time grows with the budget and the curves flatten in time as they do in iterations; across splat counts it does not order monotonically and the 1 K fits are not the fastest (the 256 K fit took longer than the 1 K fit of the same volume and budget on 71 of 187 pairs, median ratio 0.90). On every volume the 50-iteration fit of the 1 K curve took longer than the 100-iteration one, a timing wobble at the sub-second end of the axis. Time was measured on a shared, contended GPU (§ 2.4) and is indicative only.

optimisation. On Kidney actin the cell past the bound is the 256 $K$ fit. The held-out sweep at the two kidney stacks' operating points shows training PSNR +4.14 (64 $K$, Kidney nuclei) and +8.12 dB (128 $K$, Kidney actin) above held-out PSNR at 20,000 iterations, with the held-out score flat from 1,000–2,000 iterations on (§ 3.5); they are memorising noise. On *C. elegans* embryo nuclei and the OpenCell nuclear channels the dominant Gaussian placements are fixed within the first few hundred iterations and there is little left to fit. The sweep does not include a structure-density or noise-level covariate, so this reading is directional. The practical reading for this benchmark is that dataset size alone does not predict $n$, and that the budget below is set from this sweep as a whole (2,000 iterations, within 0.5 dB of $P_{20K}$ on 77 of 85 cells and at most 1.50 dB short on any), not a size-scaled schedule; it has not been validated on volumes outside the benchmark.

### 5 Empirical saturation and two hypotheses

#### 5.1 What the curves show

This document makes no appeal to optimiser convergence theory. The objective is a non-smooth L1 reconstruction loss with L1 regularisers, the fit interleaves discrete relocation steps with the Adam updates, and the learning rate follows an adaptive plateau schedule (§ 2.3). Generic convergence rates for Adam-type methods hold only under conditions this problem does not meet as stated, and a first-order stationarity rate would in any case say nothing about iterations to a PSNR plateau. The finding is empirical: at 2,000 iterations the training PSNR is within 0.5 dB of $P_{20K}$ on 77 of 85 cells (99% of it in dB is crossed by then on 75), and the training gain from 5,000 to 20,000 iterations exceeds 0.5 dB on only three cells (Neuromast membranes and Neuromast nuclei).

Two hypotheses are consistent with these observations; this sweep isolates neither:

1. **Capacity-limited saturation.** With $K$ Gaussian splats of bounded amplitude and eccentricity the attainable reconstruction error is bounded below by the basis, and once a fit is near that bound further optimisation buys little. The curves are consistent with this (most rise to a per-$K$ plateau, and the plateau rises with $K$ on every volume), but consistency is not identification. The bound need not be positive (a target made of $K$ admissible splats is exactly representable); saturation at a plateau could equally reflect a local optimum or the plateau scheduler's learning-rate decay; and the neuromast channels keep rising slowly through 20,000 iterations.
2. **Relocation shortens the exploration phase.** Without dynamic operations, an Adam-only fit must move splats to their final positions through gradient descent on position parameters, a slow process for splats initialised far from signal. Dynamic relocation explicitly moves under-utilised splats onto high-residual regions early in optimisation (§ 5.2). The large PSNR jump from iteration 50 to 200–500 visible in every panel of Figure 3 occurs with relocation enabled; no relocation-off control with matched initial states was run, so the share of that jump attributable to relocation, as opposed to ordinary early descent, is not measured.

#### 5.2 Dynamic operations and the early jump

The fitting protocol enables dynamic splat relocation: every 50 iterations throughout the fit, the 1% least important splats (by amplitude × approximate volume, at most 64 per step) are eligible for relocation onto high-residual positions. Residual and under-utilisation gates can leave an opportunity unused, so this is a schedule of opportunities, not a guarantee that 64 splats move. This is a discrete operation that re-initialises a small subset of splats; under continuous gradient descent alone, the same correction would require many descent steps to overcome the amplitude-shrinkage induced by the L1 amplitude regulariser.

An observation consistent with this mechanism is the *disproportionate* PSNR gain over the first $\sim$500 iterations relative to the next $\sim$19,500. On Kidney nuclei at $K = 64$ K, for example, PSNR rises by +3.75 dB over iterations 50–500, +3.03 dB over iterations 500–2,000 and −0.02 dB over the remaining 18,000 iterations (each value from a separately fitted cell). Relocation is active throughout, so the profile does not by itself separate relocation from early gradient descent.

Supplementary Document 3 compares progressive against single-pass fitting with relocation enabled in both arms; it is not a relocation-on/off ablation, and no such ablation is reported here.

#### 6 Practical implications

**Iteration budget.** Setting $n = 2{,}000$ leaves the training PSNR within 0.5 dB of its 20,000-iteration value on 77 of the 85 cells in this sweep and within 1 dB on 81 (Table 3; the 99% crossing count is 75). The benchmark recipe of Supplementary Document 2 (the `n2s` and `ultra` CLI presets: a 20,000-iteration cap with loss-patience 500) is generous by that measure; the fitting API's own default is 1,000 iterations with patience 300, and the `standard` preset 5,000 with patience 300.

Production stopping is a different rule from the retrospective PSNR crossing used here: the fitter stops when the regularised training loss has not improved for `early_stop_patience` iterations (or at the cap), then returns its best-loss state. The two need not agree, and on the Supplementary Document 2 fits of these volumes at the same five counts they do not. The recorded stopping iteration is a median 3.15× the cell's $n^*$, 21 of 85 fits run past 5,000 iterations and 13 reach the 20,000 cap.

Lowering the cap to 5,000–8,000 iterations is therefore a speed/quality trade-off, not a free saving. Against the 20,000 checkpoint it changes the training PSNR of a cell by at most +1.08 dB (Neuromast nuclei at 16 $K$, 5,000 vs. 20,000; +0.64 dB from 8,000). In the other direction the 5,000 and 8,000 states exceed the 20,000 one by up to 0.74 and 0.86 dB (Neuromast nuclei at 1 $K$ and Neuromast nuclei at 4 $K$). The training PSNR rises by more than 0.5 dB on three cells (all on Neuromast membranes and Neuromast nuclei), and by a median $|\Delta|$ of 0.02 dB. Under held-out PSNR the same cap costs at most 0.20 dB on the cells the blind-spot sweep covers (*Tribolium* embryo nuclei at 8,000): at the operating point the 5,000- and 8,000-iteration states are within 0.59 and 0.31 dB of the 20,000 one on every volume and at or above it on 11 of 17. The three cells whose training PSNR gains more than 0.5 dB from 5,000 to 20,000 are all on Neuromast membranes and Neuromast nuclei, and on both neuromast channels the held-out score *falls* over that range (§ 3.5). The cap is a trade-off in training PSNR only.

**Splat budget.** Capacity dominates iterations in the tested comparison (Table 5). Within that regime (4× the splats for 40× fewer iterations) adding capacity beat adding iterations on every volume, so as a heuristic for a new dataset prefer increasing $K$ over increasing $n$; no matched-time frontier was computed, so this is not an allocation rule. Averaged over the two doublings from 1 K to 4 K, each doubling of $K$ buys +1.9 dB of training PSNR across this sweep, and still +1.3 dB per doubling over the two doublings from 64 K to 256 K. At the high end on noisy confocal data that gain is largely noise memorisation (§ 2.5), so the held-out optimum $K^*$ of Supplementary Document 2, not this table, should set $K$. Past 2,000 iterations, going to 20,000 raises training PSNR by more than 0.5 dB on eight of 85 cells (at most +1.50 dB, Neuromast nuclei at 16 $K$).

**Speed/quality recommendations.**

- For draft previews: $K \in [4\,\text{K}, 16\,\text{K}]$, $n = 500$. In this sweep the 4 K-splat fit at 500 iterations reaches 76% of the volume's highest 20,000-iteration training PSNR in dB on average (range 53–98%; 4 of 17 volumes at or above 90%) and sits a mean 0.6 dB (at most 2.4 dB) below its own 4 K 20,000-iteration value. The 16 K-splat fit reaches 84% (range 62–98%; 7 of 17 at or above 90%), a mean 1.5 dB (at most 8.4 dB, Neuromast nuclei) below its 16 K 20,000-iteration value. A percentage of dB is a chosen tolerance, not a fraction of recovered signal. In held-out PSNR the 500-iteration state at 4 $K$ splats is a mean 0.7 dB (at most 2.3) below that cell's held-out peak, and at the operating point 1.5 dB (at most 8.6, Neuromast nuclei): a preview at a high count is further from its own ceiling than one at a low count.
- For publication-quality fits: pick $K$ at the per-dataset blind-spot operating point of Supplementary Document 2 (8 $K$–128 $K$ on the confocal and spinning-disk volumes, 64 $K$–128 $K$ on the iSIM channels, Zebrafish embryo nuclei (256 $K$) and *Drosophila* embryo nuclei (256 $K$); *Tribolium* embryo nuclei and Mouse heart nuclei are signal-limited and continue to benefit up to the largest count tested there, 2048 $K$), with $n = 2{,}000$. In the blind-spot sweep at the operating point the 2,000-iteration state is within 0.2 dB of the held-out peak on all 17 volumes (Table 6). The training sweep puts two operating points in its slow set (Zebrafish embryo nuclei at 256 $K$ ($n^* = 3{,}000$) and Neuromast nuclei at 64 $K$ ($n^* = 5{,}000$)), but that late training gain is not a held-out gain: on Neuromast nuclei the held-out score at 5,000 is 0.77 dB *below* its 2,000-iteration value, and on Zebrafish the 5,000-iteration held-out state differs from the 2,000 one by +0.02 dB, so the training criterion's longer budget is not recommended there. Counts above 256 K were not swept in either mode, so the budget for the signal-limited volumes at their operating point is untested; at 256 K their held-out score is still rising at 20,000 (*Tribolium*) or peaks at 8,000 (Mouse heart), and the 5,000-iteration state differs from the 2,000 one by +0.02 (*Tribolium*) and +0.08 dB (Mouse heart).
- Beyond $n = 2{,}000$, the training-PSNR gain to 20,000 exceeds 0.5 dB on eight of the 85 cells tested; on those cells the remaining budget is a small-gain trade-off in training PSNR, and on the rest it buys nothing measurable. In held-out PSNR the budget past 2,000 buys at most 0.16 dB on any of the 17 operating-point cells (and 0.20 dB on one 4 K cell, Neuromast nuclei) and costs 1.21–1.42 dB on the two neuromast channels.

### 7 Conclusion

The two budgets of a Gaussian-splat fit are not symmetric. Capacity sets the level a fit reaches and iterations only decide how quickly it gets there: in every panel of Figure 3 the five curves stack by splat count and flatten within the first few thousand iterations, and a larger model fitted briefly beats a smaller one fitted forty times longer on every volume (§ 3.3). The held-out sweep confirms the ranking on unseen voxels at its own two counts, so the effect is not an artefact of noise memorisation.

The iteration budget can be set once. At 2,000 requested iterations the training PSNR is within 0.5 dB of the 20,000-iteration reference on 77 of 85 cells (Table 3), and the held-out score at the operating point is within 0.2 dB of its peak on every volume (Table 6). What a longer fit buys after that is training PSNR on the fitted voxels: the held-out curve is flat on most volumes and falls on the two iSIM neuromast channels, where the late training rise is noise memorisation that a caller would be better off without. The 20,000-iteration cap of the benchmark recipe is therefore generous, and a cap of 5,000 to 8,000 iterations costs at most 0.20 dB of held-out PSNR on the cells swept here (*Tribolium* embryo nuclei at 8,000).

Dataset size does not predict the budget. Convergence speed shows no detectable association with voxel count on this sample (§ 4), and the ten cells that converge late are one on Kidney actin, one on Blastocyst nuclear lamina, one on Mouse heart nuclei, two on Zebrafish embryo nuclei, two on Neuromast membranes and three on Neuromast nuclei. A size-scaled schedule would therefore be tuning on the wrong covariate; the budget is set from the sweep's worst case instead. The dataset features that plausibly matter, structure density and noise level, are not quantified here and are left as follow-up.

Three recommendations follow. Choose $K$ by blind-spot calibration (Supplementary Document 2), since capacity is the axis that moves quality and training PSNR alone cannot tell signal from memorised noise at high $K$. Fit for 2,000 iterations at that count; for a draft preview, 500 iterations at 4 K splats already reach 76% of a volume's best 20,000-iteration training PSNR in dB on average (§ 6). Treat the regularity of the curves as consistent with a capacity-limited regime in which relocation handles early exploration and Adam's remaining work is local refinement, while remembering that this sweep isolates neither hypothesis (§ 5.1).
