## Supplementary material for "Luxar: Gaussian splatting for microscopy and scalable interactive web visualisation of multidimensional scientific data": supp_doc_05_loss_comparison

### Choice of reconstruction loss in Gaussian-splat fitting

Supplementary Document 5 – Luxar


**Abstract**

L1 is the default reconstruction loss of the Luxar splat fitter, although mean-squared error (MSE) is the loss that maximises PSNR at a global optimum and Poisson deviance is the likelihood for photon-counted data. We test the three losses end to end on 17 microscopy volumes, each fitted at its blind-spot operating point with three seeded replicates per loss and mode, scoring training PSNR and SSIM, held-out (blind-spot) PSNR, iterations and wall-clock time. On held-out PSNR, the model-selection metric of the main paper, the L1 loss beats the MSE loss on 11 of 17 volumes, by +0.20 to +7.78 dB, with the largest gaps on the deconvolved iSIM volumes, the Tribolium and Mouse heart light-sheet volumes and the noisy or dense confocal volumes, and trails it by at most 0.28 dB on the rest, where all three losses lie within 0.3 dB of one another. MSE is not the best training-PSNR loss for these regularised, early-stopped fits either: it leads on 1 volume, against 9 for Poisson deviance and 7 for L1. Poisson deviance tracks MSE on held-out PSNR to within ±0.1 dB on 7 volumes; its distinguishing property is early termination on confocal data where MSE runs to the iteration cap. The PSNR-optimality argument for MSE does not reach these stopped iterates of a regularised non-convex objective. We recommend keeping L1 as the default and Poisson deviance when time to termination on confocal data matters more than the last few tenths of a decibel.


#### 1 Introduction

A Gaussian-splat fit minimises a reconstruction loss between the rendered model and the target volume, and the fitter's default loss is L1. That choice needs justifying. Mean-squared error is the natural loss for a PSNR benchmark: PSNR is a monotone function of the mean squared error, so the model of lowest mean squared error in any class has the highest training PSNR that class allows. Poisson deviance is the likelihood loss for photon-counted data, which fluorescence microscopy is at the sensor. Either could be expected to beat L1 on its own terms, and the fitter offers all three. This document measures what each loss delivers when it is used as the fitter uses it, inside a regularised objective, under early stopping and followed by a cull.

Two scores are needed because they answer different questions. Training PSNR scores the reconstruction on the voxels the fit saw, so a fit that memorises noise scores well. Held-out PSNR scores it on the *held-out voxels* that the *blind-spot cross-validation* protocol hides from the fit, so memorised noise counts against it; it is the model-selection metric of the main paper and the score this document ranks the losses by. A third question, how long each loss takes to terminate, matters because the fitter stops on a plateau of its own objective, and the losses reach that plateau at very different rates.

**Notation.** $V$ is the target volume with $N$ voxels, normalised to $[0, 1]$, and $\hat{V}$ the rendered reconstruction; $V_i$ and $\hat{V}_i$ are their values at voxel $i$. The model holds $K$ splats in $D = 3$ spatial dimensions; splat $k$ has parameters $\boldsymbol{\theta}_k$, namely a centre, an amplitude $a_k$ and a lower-triangular Cholesky factor $L_k$ of its covariance $\Sigma_k = L_k L_k^\top$, whose diagonal entries $L_{k,ii}$ set its extent along each axis. $K^*$ is the splat count that blind-spot calibration recommends for a volume (Supplementary Document 2), called the operating point below. MSE, L1 and Poisson name the three losses wherever they are compared, in the prose, the table heads and the figures; the loss built on the squared residual is written $\mathcal{L}_{\text{MSE}}$ where its formula matters. The statistic, the mean of $(\hat{V}_i - V_i)^2$ over the scored voxels, is called the mean squared error in words and written MSE only inside a formula. *PSNR* on the unit-range volume is

$$\text{PSNR} = 10 \log_{10}(1/\text{MSE}), \tag{1}$$

with MSE the statistic over the scored voxels; every PSNR in this document, training or held-out, is this quantity.

#### Terms used in this document

**PSNR** The peak signal-to-noise ratio, $10 \log_{10}(\text{peak}^2/\text{MSE})$ in decibels, where peak is the reference range of the values compared; unless a document states another range, the supplements take peak 1 on volumes normalised to $[0, 1]$, so that $\text{PSNR} = -10 \log_{10} \text{MSE}$.

**SSIM** The structural similarity index, the mean over local windows of a product of luminance, contrast and structure comparisons between two images, lying in $[-1, 1]$ with 1 meaning identical.

**blind-spot cross-validation** A model-selection protocol in which a deterministic random 5% of voxels is hidden from the fit and the reconstruction is scored on those voxels afterwards, so that capacity spent on memorising noise shows up as a loss instead of a gain.

**held-out voxels** The masked voxels of a blind-spot split, whose original values the optimiser never sees and against which the held-out PSNR of each candidate splat count is scored.

**donut-median fill** The replacement of each masked voxel by the median of its $3^D - 1$ neighbours (the donut, 26 voxels in 3D); the shipped calibration draws the median over the unmasked neighbours only, and a document whose sweeps used all 26 neighbours, masked ones included, states that variant where it applies.

***K**** The splat count recommended by a blind-spot sweep: the interior maximum of the held-out PSNR curve when there is a clear peak, the smallest count within 0.3 dB of the maximum on a plateau, and the largest count tested when the curve is still rising there (signal-limited). The shared operating-point table of these documents takes the knee instead, the smallest count within 0.3 dB of that maximum, which is the largest count itself on the two signal-limited benchmark volumes.

**signal-limited curve** A held-out PSNR curve whose maximum is the largest splat count tested and which is still climbing there (a total rise of at least 0.3 dB across the sweep, a trailing mean rise of at least 0.1 dB per step and a final step of at least 0.05 dB), so the peak lies beyond the sampled range.

**relocation** The fixed-pool alternative to adding and removing splats during optimisation: the splat count is fixed at seeding, and the least informative splats are periodically moved to regions of high reconstruction residual, so the optimiser tensor shapes never change.

**cull (retention)** The removal of splats after a fit. The fit-time cull ranks splats by amplitude and keeps the top set accounting for a retention fraction of the total amplitude (0.95 by default); the contribution-based cull instead removes the splats whose contribution to the rendered field falls below an error budget or a fractional threshold, and is the baseline that removal-based level-of-detail methods use.

**iSIM** Instant structured illumination microscopy, a super-resolution fluorescence technique that performs the structured-illumination reconstruction optically in a single exposure, giving roughly a twofold resolution gain at camera frame rates.

### 2 Methods

#### 2.1 Reconstruction losses

We compare three reconstruction losses for Gaussian-splat fitting of microscopy volumes:

$$\mathcal{L}_{\mathrm{MSE}} = \tfrac{1}{N}\sum_i (\hat{V}_i - V_i)^2, \tag{2}$$

$$\mathcal{L}_{\mathrm{L1}} = \tfrac{1}{N}\sum_i |\hat{V}_i - V_i|, \tag{3}$$

$$\mathcal{L}_{\mathrm{Poi}} = \tfrac{1}{N}\sum_i \big[\hat{V}_i - V_i + V_i \log(V_i/\hat{V}_i)\big], \tag{4}$$

where $V$ is the target volume, $\hat{V}$ is the model prediction, and the sum runs over all $N$ voxels. $\mathcal{L}_{\mathrm{Poi}}$ is the unit (half) Poisson deviance [6, 7]; the $V_i \log V_i/\hat{V}_i$ term is taken to be 0 when $V_i = 0$. The implementation evaluates the deviance with the conventional factor of two and three clamps (the target at $V_i \geq 0$, the prediction at $\hat{V}_i \geq \varepsilon$ and the ratio inside the logarithm at $V_i/\hat{V}_i \geq \varepsilon$, with $\varepsilon = 10^{-8}$), so the reconstruction term actually minimised is $2\mathcal{L}_{\mathrm{Poi}}$ on the clamped quantities. All three losses are evaluated in their symmetric form (`asymmetric_penalty`$= 1$: the factor that would multiply the loss wherever the prediction exceeds the target is one).

The full training objective is

$$\mathcal{L} = \mathcal{L}_{\mathrm{recon}} + \lambda_a \frac{1}{K}\sum_{k=1}^{K} |a_k| + \lambda_d \frac{1}{KD}\sum_{k=1}^{K}\sum_{i=1}^{D} |L_{k,ii} - \sigma_{\min}|, \tag{5}$$

where $\mathcal{L}_{\mathrm{recon}}$ is $\mathcal{L}_{\mathrm{MSE}}$, $\mathcal{L}_{\mathrm{L1}}$ or $2\mathcal{L}_{\mathrm{Poi}}$, $a_k$ is the softplus-activated amplitude of splat $k$ (evaluated before the amplitude clamp) and $L_{k,ii} - \sigma_{\min}$ the softplus term of its Cholesky diagonal above the per-axis floor $\sigma_{\min}$ (the diagonal is parametrised as $\sigma_{\min}$ plus a softplus, so the penalty acts on the part above the floor, as in the main Methods), $K$ is the splat count and $D = 3$ the number of spatial dimensions, so the second regulariser averages over the $KD$ diagonal entries. The coefficients are the fitter's defaults at learning rate $\mathrm{lr} = 0.01$, $\lambda_a = 0.1\,\mathrm{lr} = 10^{-3}$ and $\lambda_d = 0.01\,\mathrm{lr} = 10^{-4}$, and are the same for all three losses. Because the reconstruction terms differ in scale and gradient structure (Section 4.1) while the regularisers do not, the three configurations are not equally regularised. Doubling the Poisson term, for instance, halves the regularisers' weight relative to it and changes the first-order conditions of the objective, and Adam's per-parameter normalisation does not undo this. The results below therefore compare three particular regularised optimiser configurations, not the reconstruction losses in isolation; an ablation with the regularisers switched off or retuned per loss was not run.

Table 1: Volumes of the loss comparison, in registry order (the fixed order of the benchmark's dataset registry, with modality groups separated by rules). CV optimum is the operating point $K^*$ of the canonical sweep of Supplementary Document 2; for a signal-limited volume the column shows the largest tested count, at which the curve is still rising. Splats used is $K^*$ plus the 5,000 margin, or the 512,000 endpoint of the canonical sweep plus the margin when signal-limited.

| Volume | Modality | Shape | Voxels | CV optimum | Splats used |
|---|---|---|---|---|---|
| OpenCell-MAP4 nuclei | Spinning-disk | 51×600×600 | 18.4 M | 16,000 | 21,000 |
| OpenCell-MAP4 microtubules | Spinning-disk | 51×600×600 | 18.4 M | 64,000 | 69,000 |
| OpenCell-LMNB1 nuclei | Spinning-disk | 112×600×600 | 40.3 M | 32,000 | 37,000 |
| OpenCell-LMNB1 nuclear lamina | Spinning-disk | 112×600×600 | 40.3 M | 64,000 | 69,000 |
| Kidney nuclei | Confocal | 16×512×512 | 4.2 M | 64,000 | 69,000 |
| Kidney actin | Confocal | 16×512×512 | 4.2 M | 128,000 | 133,000 |
| Cells3D nuclei | Confocal | 60×256×256 | 3.9 M | 32,000 | 37,000 |
| Cells3D membranes | Confocal | 60×256×256 | 3.9 M | 32,000 | 37,000 |
| Blastocyst nuclear lamina | Confocal | 236×275×271 | 17.6 M | 32,000 | 37,000 |
| *C. elegans* embryo nuclei | Confocal | 41×512×512 | 10.7 M | 8,000 | 13,000 |
| Fly brain neurons | Confocal | 435×478×478 | 99.4 M | 64,000 | 69,000 |
| *Tribolium* embryo nuclei | Light-sheet (deconv.) | 991×104×965 | 99.5 M | ≥2,048,000 (rising) | 517,000 |
| Mouse heart nuclei | Light-sheet | 597×174×960 | 99.7 M | ≥2,048,000 (rising) | 517,000 |
| Zebrafish embryo nuclei | Light-sheet (deconv.) | 407×512×512 | 106.7 M | 256,000 | 261,000 |
| *Drosophila* embryo nuclei | Light-sheet | 108×1352×532 | 77.7 M | 256,000 | 261,000 |
| Neuromast membranes | iSIM (deconv.) | 84×580×576 | 28.1 M | 128,000 | 133,000 |
| Neuromast nuclei | iSIM (deconv.) | 84×580×576 | 28.1 M | 64,000 | 69,000 |

##### 2.2 Datasets and splat counts

The loss comparison runs on every benchmark volume that has a canonical held-out sweep, the ten-count blind-spot sweep from 1,000 to 512,000 splats of Supplementary Document 2 (Table 1: all 17 volumes from 12 datasets of the benchmark). At each volume the splat count is fixed at the operating point $K^*$ of that sweep plus a 5,000-splat margin. The operating point comes from L1 fits and the margin is a convention: it gives every loss the same pool, but does not by itself establish that MSE or Poisson fitting has sufficient capacity at that count (Section 4.3). A volume whose held-out curve is still rising at the largest tested count has no interior optimum inside the sweep. Such a *signal-limited curve* volume is fitted at the endpoint of the canonical sweep plus the same margin, not at the 2,048,000 endpoint of the extended sweep to which the two light-sheet volumes were additionally taken (Table 1); that operating point is a convention, not a measured optimum. The tables below cover these 17 volumes.

##### 2.3 Fitting protocol

Each (loss, dataset) pair is fitted in two *modes*: *training mode* (full-volume fit, which scores training PSNR/SSIM and times convergence) and *held-out mode* (blind-spot fit, defined below, which scores held-out PSNR). A *cell* is one (loss, mode) pair on one volume. Volumes are normalised to $[0, 1]$ before fitting by the benchmark's per-dataset affine map, with the dataset-specific percentile clipping and, for the two neuromast channels, background subtraction described in the main Methods.

The 102 cells (3 losses × 17 volumes × 2 modes) share one resolved configuration. The iteration cap is 30,000, with early stopping after 1,000 iterations without improvement of the training objective (5); the fitter returns its best-state snapshot, the parameters one optimiser step after the lowest training objective seen. The optimiser is Adam [5] at learning rate 0.01 with the fitter's plateau scheduler, which multiplies the learning rate by 0.9 after 15 iterations without loss improvement. Dynamic splat *relocation* is enabled, seeding is automatic (`seed_method=auto`), and there is no background-floor subtraction (`floor=none`, whereas the shipped CLI default is `auto`). The regularisers are $\lambda_a = 10^{-3}$ and $\lambda_d = 10^{-4}$. A final *cull (retention)* at 0.999 amplitude retention follows every fit, after which the returned model holds 0.500–0.997× the requested splat count (no model is returned uncut). The retained fraction differs by loss on the same volume, by up to 0.09 of the requested count (training mode: Blastocyst nuclear lamina MSE 0.94, L1 0.85, Poisson 0.90; Neuromast nuclei MSE 0.56, L1 0.50, Poisson 0.55), so the scored models are matched in the requested pool, not in final splat count.

Each cell is fitted three times. Replicate $r$ seeds torch, CUDA and NumPy with $r$ before fitting (`torch.manual_seed`, `torch.cuda.manual_seed_all`, `np.random.seed`), so the three replicates differ only in seed-dependent initialisation and relocation (and any GPU non-determinism), not in specimen, mask or configuration. The spread they measure is seed variability, not sampling uncertainty over specimens or masks. This gives 306 fits in total; cells in Tables 2–6 report mean ± sample std over the three replicates. PSNR is Eq. (1) with MSE taken over the scored voxels. *SSIM* [8] is computed in 3-D with an 11-voxel Gaussian window ($\sigma = 1.5$), constants $C_1 = 0.01^2$ and $C_2 = 0.03^2$ for unit data range, population moments and valid (no-padding) convolution, averaged over the volume. The result files record, per fit, the requested and retained splat counts, the replicate index, wall-clock time, iteration count and timestamp; they carry no software-commit or library-version fingerprint.

The iteration cap is intentionally generous; 184 of the 306 runs terminate through early stopping and the remaining 122 hit the cap. Reaching the cap is almost always a property of the (volume, loss) pair, not of the seed. In training mode MSE runs to the cap in every replicate on eight volumes (Kidney nuclei, Kidney actin, Cells3D nuclei, Mouse heart nuclei, Zebrafish embryo nuclei, Drosophila embryo nuclei, Neuromast membranes and Neuromast nuclei), L1 on seven (Kidney actin, Blastocyst nuclear lamina, Mouse heart nuclei, Zebrafish embryo nuclei, Drosophila embryo nuclei, Neuromast membranes and Neuromast nuclei) and Poisson on five (Mouse heart nuclei, Zebrafish embryo nuclei, Drosophila embryo nuclei, Neuromast membranes and Neuromast nuclei). On those five light-sheet and *iSIM* volumes all three losses run to the cap in both modes, and Poisson never reaches it on a confocal or spinning-disk volume. Only three training-mode cells reach the cap in some replicates but not others: Kidney nuclei L1 (2 of 3 reps), Cells3D membranes MSE (1 of 3 reps) and Tribolium embryo nuclei L1 (1 of 3 reps). These mixed cells produce the large iteration-count standard deviations in Table 5. Termination at the cap or by patience is not evidence of stationarity: no fit anywhere reaches the error-based convergence criterion ($\max_i |\hat{V}_i - V_i| < 0.01$ on the normalised volume) in any replicate of any cell.

##### 2.4 Blind-spot evaluation

For the held-out mode, 5% of voxels (seed 42) are masked with their $3 \times 3 \times 3$ *donut-median fill* value before fitting. Held-out PSNR is computed at the masked positions against the *original* (pre-replacement) intensities. This is the Noise2Self blind-spot self-supervision protocol of Batson and Royer [1]; the masking matches the cross-validation experiments of Supplementary Document 2 exactly (the fit budget differs: 30,000 iterations and patience 1,000 here against 20,000 and 500 there) and isolates signal recovery from noise memorisation. The fits reported here filled every masked voxel with the median of all 26 shell voxels, other held-out voxels included (the all-donor fill), as Supplementary Document 2 states for its canonical sweeps; the released code excludes held-out donors from the fill. Figure 1 sketches the protocol.

##### 2.5 Compute

All 306 fits (102 cells × 3 reps) ran on a single NVIDIA RTX PRO 6000 Blackwell (96 GB VRAM, compute capability 12.0), in concurrent lanes that shared the card with other fitting jobs. The fit durations sum to 55.7 aggregate fitting hours (lanes overlapped, so this is not exclusive device time), dominated by the cells that run to the 30,000-iteration cap; the longest fits are the Drosophila embryo nuclei under MSE at 261,000 splats (45.3 min) and the Tribolium embryo nuclei under L1 at 517,000 splats (41.4 min). The competing load was neither controlled nor recorded, so the wall-clock figures below are observations under these schedules: inflated relative to an idle card, comparable only loosely within this document, and not measurements of the intrinsic per-iteration cost of a loss (Section 3.5).

#### 3 Results

##### 3.1 Training PSNR

MSE is not the best training-PSNR loss for these fits, although it is the loss whose global minimiser would be (Section 4.3). It has the highest mean training PSNR on one volume of seventeen (OpenCell-MAP4 microtubules), Poisson deviance on nine and L1 on seven (Cells3D membranes, Tribolium embryo nuclei, Mouse heart nuclei, Zebrafish embryo nuclei, Drosophila embryo nuclei, Neuromast membranes and Neuromast nuclei); a rounded tie counts for no loss (0 here). These are the headline win counts of the document. The L1−MSE difference is positive on fourteen volumes, by +0.14 to +10.21 dB with the largest gap on the deconvolved iSIM Neuromast nuclei, and negative by 0.06–0.25 dB on three spinning-disk channels (OpenCell-MAP4 nuclei, OpenCell-MAP4 microtubules and OpenCell-LMNB1 nuclei), where all three losses land within 0.3 dB of one another (Table 2). The between-replicate standard deviation is at most 0.24 dB on every PSNR cell of this document.

##### 3.2 Training SSIM

On structural similarity the three losses are close on most volumes at the splat counts used (Table 3). Poisson deviance has the highest SSIM on nine volumes and L1 on seven, with 1

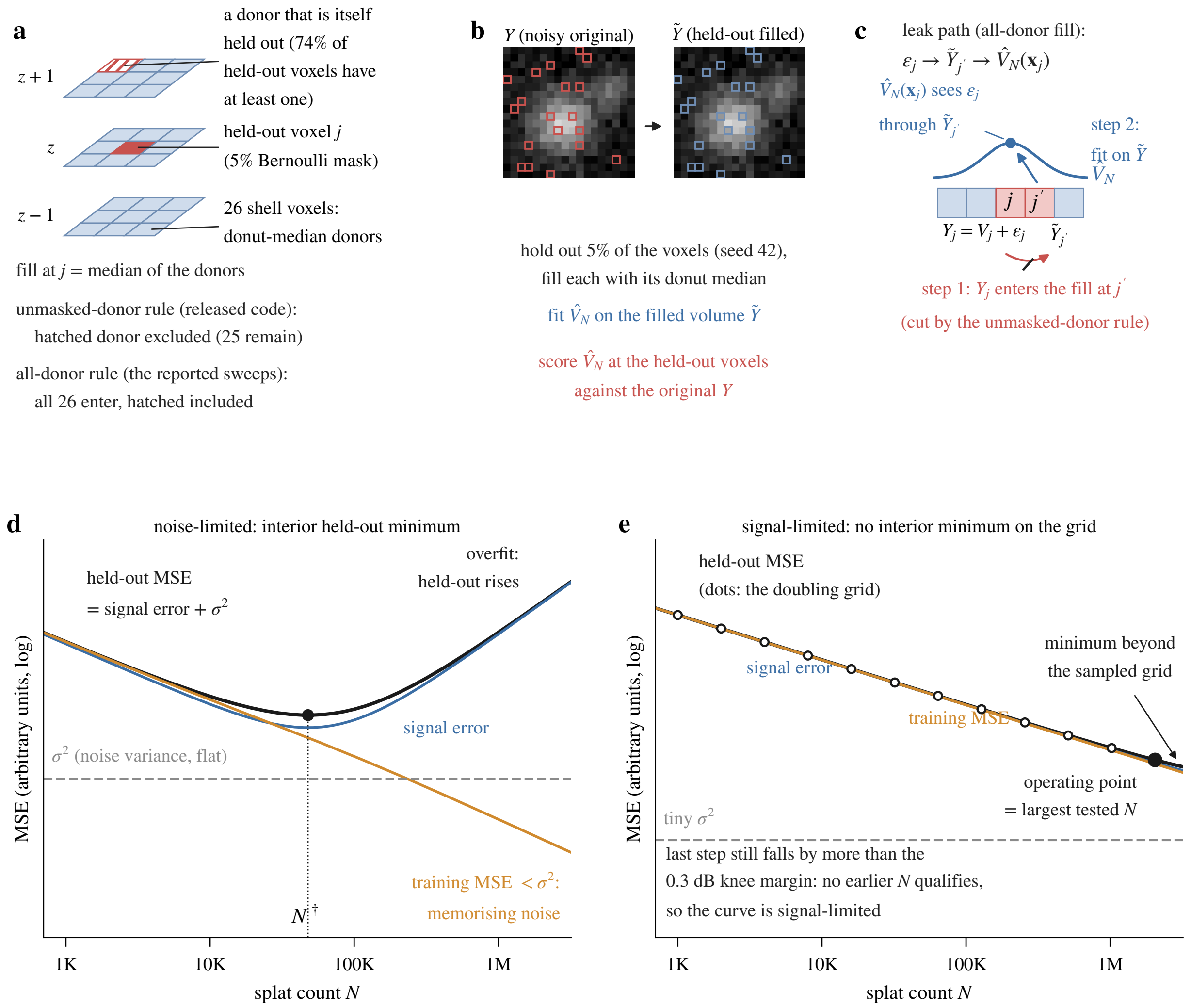


Figure 1: Schematic of the blind-spot protocol (shared with Supplementary Document 2). (a) A $3 \times 3 \times 3$ voxel cube as three exploded slices: the centre voxel is held out and its 26 shell voxels donate the donut median. (b) A small patch run through the masking and the donut fill: the fit sees the filled volume and is scored at the held-out voxels against the original values. (c) The neighbour leak between two adjacent held-out voxels, which the fill rule of the released code avoids; the fits of this document used the all-donor fill (Section 2.4). (d, e) The held-out decomposition against splat count, where the schematic's $N$ is this document's $K$ and its $N^\dagger$ is $K^*$: signal error, the flat noise variance $\sigma^2$, their sum with its interior minimum, and the training error falling below $\sigma^2$; in (e) a small $\sigma^2$ leaves no interior minimum on the sampled grid, the signal-limited case. Panels (d) and (e) are idealised curves, not data from any volume; the sweeps themselves are in Supplementary Document 2.

Table 2: Training PSNR (dB), full-volume fit, mean ± sample std over three seeded replicates. In this and the following tables bold marks the loss whose rounded mean is best at the printed precision; a rounded tie leaves every tied loss bold. A replicate spread that rounds to zero at the printed precision is omitted, so a cell showing the mean alone has three replicates agreeing to that precision. The column heads MSE, L1 and Poisson name the losses; the L1 − MSE column is the difference of the two printed means and Best names the bold loss (tie when more than one is bold); Tables 3 to 6 share this layout.

| Volume | MSE | L1 | Poisson | L1 − MSE (dB) | Best |
|---|---|---|---|---|---|
| OpenCell-MAP4 nuclei | 28.54 | 28.48 | **28.58** | −0.06 | Poi |
| OpenCell-MAP4 microtubules | **25.97** | 25.72 | 25.95 | −0.25 | MSE |
| OpenCell-LMNB1 nuclei | 25.69 | 25.63 | **25.72** | −0.06 | Poi |
| OpenCell-LMNB1 nuclear lamina | 34.48 | 34.62 ± 0.01 | **34.65 ± 0.01** | +0.14 | Poi |
| Kidney nuclei | 37.89 ± 0.04 | 38.45 ± 0.01 | **38.71 ± 0.04** | +0.56 | Poi |
| Kidney actin | 38.31 ± 0.01 | 39.41 ± 0.03 | **40.55 ± 0.02** | +1.10 | Poi |
| Cells3D nuclei | 36.73 ± 0.01 | 36.92 ± 0.01 | **36.99 ± 0.01** | +0.19 | Poi |
| Cells3D membranes | 47.27 ± 0.02 | **50.25 ± 0.02** | 50.08 ± 0.03 | +2.98 | L1 |
| Blastocyst nuclear lamina | 38.94 ± 0.01 | 39.38 ± 0.01 | **39.59** | +0.44 | Poi |
| C. elegans embryo nuclei | 38.86 ± 0.04 | 39.31 | **39.39** | +0.45 | Poi |
| Fly brain neurons | 42.62 ± 0.04 | 44.90 ± 0.01 | **45.20** | +2.28 | Poi |
| Tribolium embryo nuclei | 50.10 ± 0.05 | **55.13 ± 0.08** | 52.78 ± 0.13 | +5.03 | L1 |
| Mouse heart nuclei | 39.57 ± 0.02 | **43.48 ± 0.11** | 41.75 ± 0.03 | +3.91 | L1 |
| Zebrafish embryo nuclei | 39.35 ± 0.03 | **39.89 ± 0.05** | 39.53 ± 0.03 | +0.54 | L1 |
| Drosophila embryo nuclei | 34.48 | **34.88** | 34.71 ± 0.01 | +0.40 | L1 |
| Neuromast membranes | 52.39 ± 0.02 | **57.38 ± 0.07** | 53.92 ± 0.13 | +4.99 | L1 |
| Neuromast nuclei | 56.67 ± 0.01 | **66.88 ± 0.12** | 63.73 ± 0.24 | +10.21 | L1 |

rounded tie (on Neuromast nuclei). The inter-loss range of the means is $\leq 0.02$ on fifteen of the seventeen volumes; the exceptions are Fly brain neurons (0.126) and Mouse heart nuclei (0.033). The largest within-volume range (0.126, Fly brain neurons) is about 3.9× smaller than the across-volume range of the means ($\approx 0.49$, set by image content), and the between-replicate std is $\leq 0.0034$ on every cell. No equivalence margin was specified in advance, so the small ranges describe closeness, not demonstrated equivalence.

##### 3.3 Held-out PSNR (blind-spot cross-validation)

Held-out PSNR is the score this document ranks the losses by, and it splits by volume (Table 4). The L1 loss exceeds the MSE loss on eleven volumes (Kidney nuclei, Kidney actin, Cells3D membranes, C. elegans embryo nuclei, Fly brain neurons, Tribolium embryo nuclei, Mouse heart nuclei, Zebrafish embryo nuclei, Drosophila embryo nuclei, Neuromast membranes and Neuromast nuclei), by +0.20 to +7.78 dB; MSE is ahead of L1 by 0.04–0.28 dB on the other six (OpenCell-MAP4 nuclei, OpenCell-MAP4 microtubules, OpenCell-LMNB1 nuclei, OpenCell-LMNB1 nuclear lamina, Cells3D nuclei and Blastocyst nuclear lamina). A two-sided sign test on these 17 paired differences gives $p = 0.33$, not significant at $n = 17$; the count of wins alone does not establish the direction, and the case for L1 rests on the sizes of the gaps. Those sizes are very unequal. On seven volumes (OpenCell-MAP4 nuclei, OpenCell-MAP4 microtubules, OpenCell-LMNB1 nuclei, OpenCell-LMNB1 nuclear lamina, Cells3D nuclei, Blastocyst nuclear lamina and Zebrafish embryo nuclei) all three losses lie within 0.3 dB of one another, and every MSE win falls in this tight set. On the remaining ten L1 leads on eight and Poisson on two (C. elegans embryo nuclei

Table 3: Training SSIM, mean ± std over three replicates; bold marks the best rounded mean, ties both bold; a spread below 0.00005 is omitted. The L1 − MSE column is the difference of the printed means.

| Volume | MSE | L1 | Poisson | L1 − MSE | Best |
|---|---|---|---|---|---|
| OpenCell-MAP4 nuclei | 0.6776 | 0.6699 ± 0.0001 | **0.6780 ± 0.0001** | −0.0077 | Poi |
| OpenCell-MAP4 microtubules | 0.6350 ± 0.0001 | 0.6289 ± 0.0001 | **0.6367 ± 0.0001** | −0.0061 | Poi |
| OpenCell-LMNB1 nuclei | 0.5159 ± 0.0002 | 0.5121 ± 0.0001 | **0.5172 ± 0.0001** | −0.0038 | Poi |
| OpenCell-LMNB1 nuclear lamina | 0.9219 ± 0.0003 | 0.9174 ± 0.0003 | **0.9263 ± 0.0002** | −0.0045 | Poi |
| Kidney nuclei | 0.9792 ± 0.0002 | **0.9819** | 0.9816 ± 0.0001 | +0.0027 | L1 |
| Kidney actin | 0.9835 ± 0.0002 | 0.9888 | **0.9896** | +0.0053 | Poi |
| Cells3D nuclei | 0.8736 ± 0.0001 | 0.8765 ± 0.0001 | **0.8771 ± 0.0002** | +0.0029 | Poi |
| Cells3D membranes | 0.9872 | **0.9923 ± 0.0001** | 0.9920 | +0.0051 | L1 |
| Blastocyst nuclear lamina | 0.9419 ± 0.0007 | 0.9303 ± 0.0002 | **0.9493** | −0.0116 | Poi |
| C. elegans embryo nuclei | 0.8795 ± 0.0012 | 0.8907 | **0.8932** | +0.0112 | Poi |
| Fly brain neurons | 0.8443 ± 0.0034 | 0.9591 | **0.9704** | +0.1148 | Poi |
| Tribolium embryo nuclei | 0.9903 ± 0.0001 | **0.9972 ± 0.0001** | 0.9946 ± 0.0002 | +0.0069 | L1 |
| Mouse heart nuclei | 0.9456 ± 0.0002 | **0.9781 ± 0.0007** | 0.9696 ± 0.0002 | +0.0325 | L1 |
| Zebrafish embryo nuclei | 0.9671 ± 0.0008 | **0.9734 ± 0.0006** | 0.9719 ± 0.0003 | +0.0063 | L1 |
| Drosophila embryo nuclei | 0.8768 ± 0.0001 | **0.8880 ± 0.0001** | 0.8845 ± 0.0002 | +0.0112 | L1 |
| Neuromast membranes | 0.9980 | **0.9998** | 0.9997 | +0.0018 | L1 |
| Neuromast nuclei | 0.9990 | **0.9999** | **0.9999** | +0.0009 | tie |

and Fly brain neurons, by +0.03 and +0.27 dB over L1), MSE on none. The large L1 margins are on the noisy or dense confocal volumes (Kidney nuclei +1.10, Kidney actin +2.02, Cells3D membranes +2.34 dB over MSE), on the Tribolium and Mouse heart light-sheet volumes (+4.79 and +3.97 dB) and on both iSIM volumes (+2.36 dB on the Neuromast membranes, +7.78 dB on the Neuromast nuclei); on the other two light-sheet volumes L1 leads by +0.20 dB (Zebrafish embryo nuclei, inside the tight set) and +0.40 dB (Drosophila embryo nuclei).

Poisson deviance is within ±0.1 dB of MSE on seven volumes and ranges from −0.44 dB (Kidney actin) to +4.86 dB (Neuromast nuclei) relative to it. Every between-replicate std in Table 4 is $\leq$ 0.10 dB. Each entry scores the model the fitter returns, the best-state snapshot after its loss-dependent stop and the final cull, not the best held-out score seen during fitting (Section 5).

##### 3.4 Convergence speed

The iteration cap of 30,000 is intentionally generous. Whether a fit reaches it is almost always a property of the (volume, loss) pair, not of the seed (Section 2.3): in training mode MSE runs to the cap in every replicate on eight volumes, L1 on seven and Poisson on five, and Poisson never reaches it on a confocal or spinning-disk volume (Table 5). Poisson stops in the fewest iterations on eight of seventeen volumes. On the five light-sheet and iSIM volumes all three losses run to the cap in both modes, so a mean of 30,000 in the table is a capped observation, not an early stop, and those rows are exact three-way ties. The cells with very large standard deviations are the three mixed cells that cap in some replicates only (Kidney nuclei L1 (2 of 3 reps), Cells3D membranes MSE (1 of 3 reps) and Tribolium embryo nuclei L1 (1 of 3 reps)); the remaining spread (for instance OpenCell-MAP4 microtubules under L1, mean 4,100) is seed-dependent early stopping well below the cap.

Table 4: Held-out PSNR (dB) at blind-spot-masked voxels, mean ± std over three replicates; bold marks the best rounded mean, ties both bold (1 rounded tie, on OpenCell-MAP4 nuclei).

| Volume | MSE | L1 | Poisson | L1 − MSE (dB) | Best |
|---|---|---|---|---|---|
| OpenCell-MAP4 nuclei | **28.26** | 28.16 | **28.26** | −0.10 | tie |
| OpenCell-MAP4 microtubules | **25.01 ± 0.01** | 24.89 | 24.98 | −0.12 | MSE |
| OpenCell-LMNB1 nuclei | **25.54** | 25.45 | 25.53 | −0.09 | MSE |
| OpenCell-LMNB1 nuclear lamina | 34.05 | 34.01 | **34.08** | −0.04 | Poi |
| Kidney nuclei | 32.81 ± 0.01 | **33.91 ± 0.03** | 32.89 ± 0.02 | +1.10 | L1 |
| Kidney actin | 28.51 ± 0.03 | **30.53 ± 0.02** | 28.07 ± 0.05 | +2.02 | L1 |
| Cells3D nuclei | **35.60 ± 0.01** | 35.45 | 35.56 | −0.15 | MSE |
| Cells3D membranes | 44.53 ± 0.01 | **46.87 ± 0.05** | 45.08 ± 0.01 | +2.34 | L1 |
| Blastocyst nuclear lamina | **37.85 ± 0.01** | 37.57 ± 0.01 | 37.77 | −0.28 | MSE |
| C. elegans embryo nuclei | 38.70 ± 0.03 | 39.23 | **39.26** | +0.53 | Poi |
| Fly brain neurons | 42.32 ± 0.06 | 44.26 | **44.53** | +1.94 | Poi |
| Tribolium embryo nuclei | 49.83 ± 0.06 | **54.62** | 52.49 ± 0.03 | +4.79 | L1 |
| Mouse heart nuclei | 38.87 ± 0.03 | **42.84 ± 0.07** | 41.06 ± 0.06 | +3.97 | L1 |
| Zebrafish embryo nuclei | 37.63 ± 0.03 | **37.83 ± 0.01** | 37.74 ± 0.02 | +0.20 | L1 |
| Drosophila embryo nuclei | 33.63 ± 0.01 | **34.03 ± 0.02** | 33.77 ± 0.01 | +0.40 | L1 |
| Neuromast membranes | 48.44 ± 0.07 | **50.80 ± 0.02** | 48.76 ± 0.04 | +2.36 | L1 |
| Neuromast nuclei | 55.82 ± 0.02 | **63.60 ± 0.10** | 60.68 ± 0.03 | +7.78 | L1 |

Table 5: Iterations to termination, training mode (cap 30,000; patience 1,000), mean ± std over three replicates; bold marks the fewest at the printed precision, ties all bold, and Best names that loss. A cell showing 30,000 alone is three replicates at the cap.

| Volume | MSE | L1 | Poisson | L1 − MSE | Best |
|---|---|---|---|---|---|
| OpenCell-MAP4 nuclei | 2,150 | **1,933 ± 58** | 2,000 | −217 | L1 |
| OpenCell-MAP4 microtubules | 2,800 ± 50 | 4,100 ± 1,472 | **2,283 ± 29** | +1,300 | Poi |
| OpenCell-LMNB1 nuclei | 2,217 ± 29 | **2,117 ± 29** | 2,133 ± 29 | −100 | L1 |
| OpenCell-LMNB1 nuclear lamina | 2,733 ± 29 | 4,600 ± 100 | **2,300** | +1,867 | Poi |
| Kidney nuclei | 30,000 | 24,517 ± 9,497 | **2,867 ± 29** | −5,483 | Poi |
| Kidney actin | 30,000 | 30,000 | **3,933 ± 126** | +0 | Poi |
| Cells3D nuclei | 30,000 | 2,850 ± 350 | **2,250** | −27,150 | Poi |
| Cells3D membranes | 13,075 ± 14,660 | **2,800 ± 50** | 2,967 ± 29 | −10,275 | L1 |
| Blastocyst nuclear lamina | 5,833 ± 58 | 30,000 | **2,783 ± 29** | +24,167 | Poi |
| C. elegans embryo nuclei | 4,917 ± 161 | 1,917 ± 29 | **1,817 ± 29** | −3,000 | Poi |
| Fly brain neurons | 10,067 ± 225 | **2,133 ± 29** | 2,167 ± 58 | −7,934 | L1 |
| Tribolium embryo nuclei | 3,867 ± 142 | 18,225 ± 12,181 | **3,767 ± 38** | +14,358 | Poi |
| Mouse heart nuclei | **30,000** | **30,000** | **30,000** | +0 | tie |
| Zebrafish embryo nuclei | **30,000** | **30,000** | **30,000** | +0 | tie |
| Drosophila embryo nuclei | **30,000** | **30,000** | **30,000** | +0 | tie |
| Neuromast membranes | **30,000** | **30,000** | **30,000** | +0 | tie |
| Neuromast nuclei | **30,000** | **30,000** | **30,000** | +0 | tie |

Table 6: Wall-clock fit time (seconds), training mode, mean ± std over three replicates on a shared GPU; bold marks the fastest at the printed precision, ties both bold (1 rounded tie, on Mouse heart nuclei), and Best names that loss.

| Volume | MSE | L1 | Poisson | L1 − MSE (s) | Best |
|---|---|---|---|---|---|
| OpenCell-MAP4 nuclei | 106.5 ± 1.2 | **92.4 ± 2.1** | 100.8 ± 3.0 | −14.1 | L1 |
| OpenCell-MAP4 microtubules | 97.7 ± 1.9 | 134.8 ± 48.1 | **84.8 ± 1.6** | +37.1 | Poi |
| OpenCell-LMNB1 nuclei | **98.2 ± 4.6** | 104.5 ± 5.1 | 99.0 ± 31.1 | +6.3 | MSE |
| OpenCell-LMNB1 nuclear lamina | 122.0 ± 3.5 | 259.3 ± 32.3 | **113.4 ± 5.6** | +137.3 | Poi |
| Kidney nuclei | 1,209.9 ± 225.8 | 1,090.3 ± 472.0 | **143.6 ± 7.6** | −119.6 | Poi |
| Kidney actin | 1,252.5 ± 104.4 | 782.4 ± 111.9 | **124.6 ± 8.2** | −470.1 | Poi |
| Cells3D nuclei | 744.0 ± 162.7 | 60.5 ± 9.0 | **50.7 ± 2.8** | −683.5 | Poi |
| Cells3D membranes | 297.6 ± 321.7 | **64.8 ± 2.1** | 71.8 ± 3.6 | −232.8 | L1 |
| Blastocyst nuclear lamina | 274.0 ± 13.4 | 1,196.3 ± 84.9 | **113.4 ± 15.9** | +922.3 | Poi |
| C. elegans embryo nuclei | 145.2 ± 6.6 | 82.9 ± 1.0 | **72.4 ± 8.2** | −62.3 | Poi |
| Fly brain neurons | 601.6 ± 43.6 | **164.9 ± 13.5** | 250.9 ± 30.0 | −436.7 | L1 |
| Tribolium embryo nuclei | 513.9 ± 53.6 | 2,485.2 ± 1,748.7 | **432.5 ± 44.0** | +1,971.3 | Poi |
| Mouse heart nuclei | 1,865.5 ± 480.9 | **1,610.5 ± 71.1** | **1,610.5 ± 545.7** | −255.0 | tie |
| Zebrafish embryo nuclei | **1,222.3 ± 90.1** | 1,248.2 ± 77.8 | 1,234.6 ± 106.8 | +25.9 | MSE |
| Drosophila embryo nuclei | 2,715.5 ± 171.9 | 1,550.8 ± 64.9 | **1,533.0 ± 69.0** | −1,164.7 | Poi |
| Neuromast membranes | **1,329.9 ± 288.1** | 1,769.1 ± 49.0 | 1,437.5 ± 55.1 | +439.2 | MSE |
| Neuromast nuclei | 945.1 ± 91.6 | 900.5 ± 37.7 | **848.2 ± 51.3** | −44.6 | Poi |

##### 3.5 Wall-clock fit time

Wall-clock time was measured on a GPU shared with other, uncontrolled jobs (Section 2.5), so the ratios here are observations under that load, not intrinsic costs. Mean time per iteration differs across losses by up to 1.94× within a volume (Fly brain neurons), so wall-clock only roughly tracks iteration count. Poisson is fastest on ten of seventeen volumes (Table 6). On the three volumes where MSE runs to the cap and Poisson stops early (Kidney nuclei, Kidney actin and Cells3D nuclei) MSE's mean time is 8.4 to 14.7 times Poisson's, the largest ratio on Cells3D nuclei; on the two where L1 caps and Poisson does not (Kidney actin and Blastocyst nuclear lamina) L1's mean time is 6.3 to 10.5 times Poisson's. Where all three losses run to the cap (the five light-sheet and iSIM volumes) Poisson has no iteration advantage and is not always the fastest column; on the Zebrafish embryo nuclei all three losses take ≈ 20 min.

##### 3.6 Cross-dataset summary

Figure 2 restates Tables 2, 4 and 5 as paired differences against MSE, L1 − MSE and Poisson − MSE, one row per volume in registry order, with the replicate spread of each difference as an error bar. Three patterns are visible at a glance. (i) On training PSNR, both differences are positive on fourteen of seventeen rows, so MSE is the lowest of the three losses there, and both are negative on one (OpenCell-MAP4 microtubules, where L1 is 0.25 dB below MSE, small but reproducible across the replicates). (ii) On held-out PSNR, the L1 − MSE difference is large and positive on the noisy or dense confocal rows, on the Tribolium and Mouse heart rows and on both iSIM rows, where the error bars are far from the zero line; Poisson edges L1 on the *C. elegans* embryo and the Fly brain neurons, and MSE is ahead only on rows where all three losses sit within 0.3 dB of one another (Section 3.3). (iii) On iterations to termination the Poisson − MSE difference is large and negative on the confocal rows where MSE runs to the cap and Poisson does not, and the wide error bars belong to the mixed cells that cap in some replicates only; on the rows where all three losses cap, both iteration differences are zero. The SSIM differences of Table 3 are not drawn: they are below 0.02 on all but 2 volumes.

### 4 Theoretical context

#### 4.1 Gradient structure

Let $r_i = \hat{V}_i - V_i$ be the signed residual at voxel $i$. The per-voxel gradients of the three losses with respect to $\hat{V}_i$ are

$$\frac{\partial \mathcal{L}_{\mathrm{MSE}}}{\partial \hat{V}_i} = \tfrac{2}{N}\, r_i, \tag{6}$$

$$\frac{\partial \mathcal{L}_{\mathrm{L1}}}{\partial \hat{V}_i} = \tfrac{1}{N}\, \mathrm{sgn}(r_i), \tag{7}$$

$$\frac{\partial \mathcal{L}_{\mathrm{Poi}}}{\partial \hat{V}_i} = \tfrac{1}{N}\left(1 - \tfrac{V_i}{\hat{V}_i}\right) = \tfrac{1}{N}\, \frac{r_i}{\hat{V}_i}. \tag{8}$$

These gradients pull back to the parameters $\boldsymbol{\theta}_k$ of splat $k$ via $\partial \hat{V}_i / \partial \boldsymbol{\theta}_k$, which is identical for all three losses. The loss only enters through the weight assigned to each voxel's residual.

**Proposition 1** (voxel-weighting profile)**.** *Each loss weights voxel $i$'s contribution to the parameter gradient by a voxel-specific factor $w_i$, in the sense that $\partial \mathcal{L} / \partial \hat{V}_i = w_i / N$, with*

$$w_i^{\mathrm{MSE}} = 2\, r_i, \qquad w_i^{\mathrm{L1}} = \mathrm{sgn}(r_i), \qquad w_i^{\mathrm{Poi}} = r_i / \hat{V}_i.$$

*The common factor $1/N$ is dropped from the $w_i$ and restored wherever an absolute per-voxel contribution is quoted below.*

Figure 3 draws the three profiles. Three consequences follow:

**MSE.** $w_i^{\mathrm{MSE}}$ grows linearly with residual magnitude. A bright-outlier voxel with residual 100 produces a gradient contribution $100\times$ larger than a background voxel with residual 1. This pours gradient budget into the largest residuals, which is PSNR-optimal at the global minimiser of the objective but outlier-seeking in finite-iteration optimisation.

**L1.** $w_i^{\mathrm{L1}}$ is bounded: every voxel contributes $\pm 1/N$ regardless of residual magnitude (bounded influence in the sense of Huber [3]). In the limit, L1 converges to a per-voxel median, not a mean.

**Poisson.** $w_i^{\mathrm{Poi}} = r_i / \hat{V}_i$ divides the residual by the prediction. Relative to MSE the per-voxel gradient is $w_i^{\mathrm{Poi}} / w_i^{\mathrm{MSE}} = 1/(2\hat{V}_i)$ (or $1/\hat{V}_i$ for the doubled deviance the implementation minimises): a ratio set by the prediction, not by the residual, so it does not shrink as $\hat{V}_i \to V_i$. On a $[0,1]$-normalised volume the unit-deviance gradient exceeds MSE's wherever $\hat{V}_i < 0.5$ (on background and on most dim structure) and falls below it only on the brightest voxels. Unlike L1's weight, $w_i^{\mathrm{Poi}}$ is neither bounded nor residual-independent: it is linear in $r_i$ and unbounded as $\hat{V}_i \to 0$ with

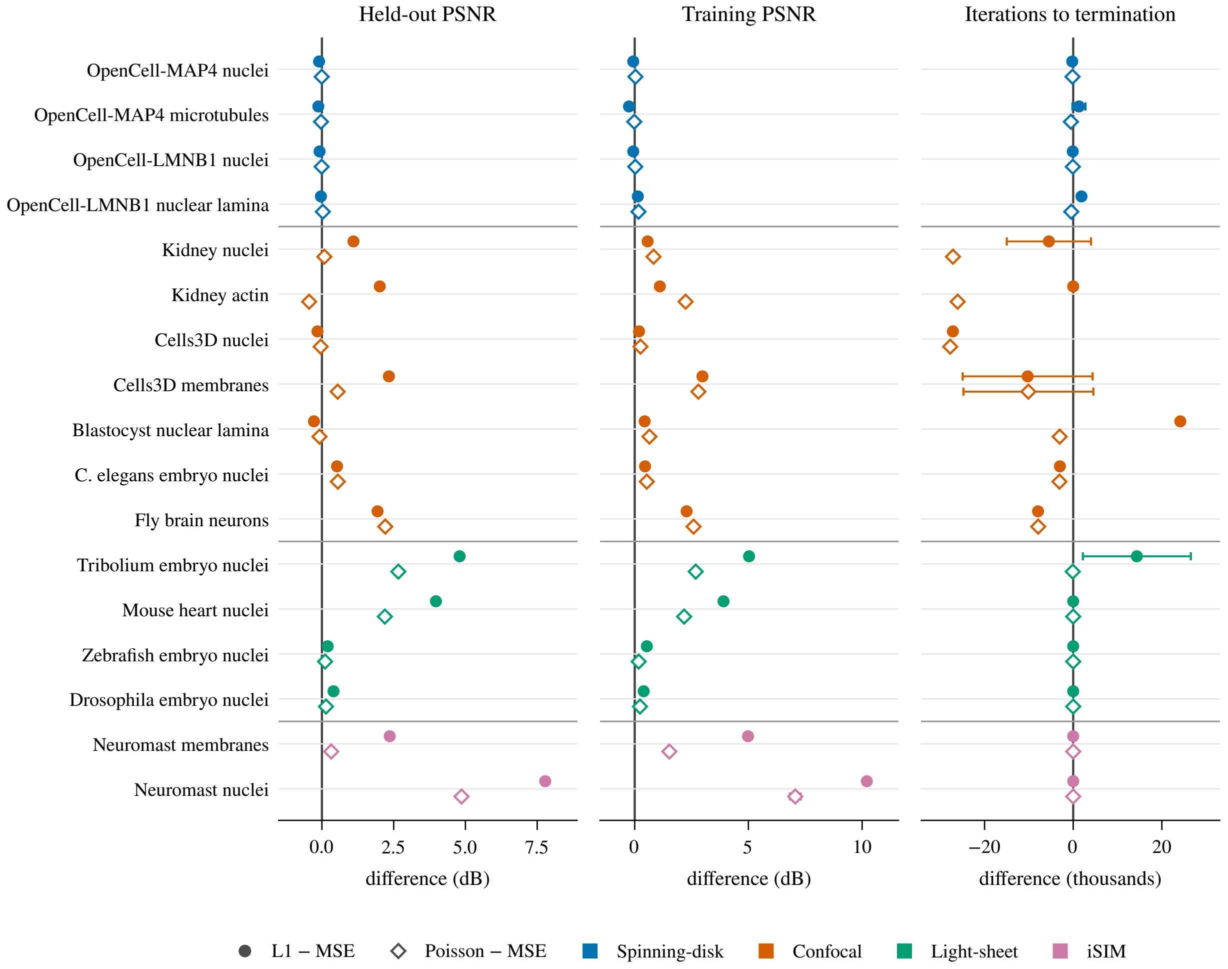


Figure 2: Paired differences against MSE of held-out PSNR (masked-volume fits), training PSNR and iterations to termination (full-volume fits): L1 − MSE (filled circles) and Poisson − MSE (open diamonds) of the replicate means, one row per volume in registry order, coloured by imaging modality with thin rules between modality groups. A point right of the zero line means the loss scored higher (or ran longer) than MSE on that volume. Error bars are the replicate spread of the difference, $\sqrt{s_a^2 + s_b^2}$ for the two sample standard deviations over the three seeded replicates; they measure seed variability only. The iteration panel is in thousands of iterations; a difference of exactly zero with no bar is a row on which both losses ran to the 30,000 cap in every replicate.

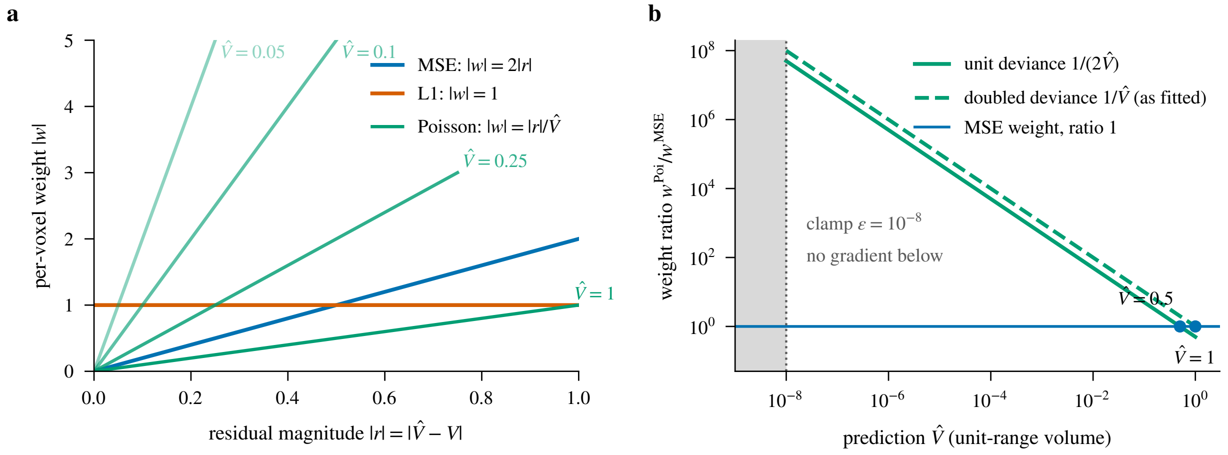


Figure 3: Per-voxel weight profiles of Proposition 1 (schematic: closed-form curves, no data). (a) Weight magnitude $|w|$ against the residual magnitude $|r| = |\hat{V} - V|$ on the unit-range volume: MSE grows linearly ($2|r|$), L1 is flat at 1, and Poisson is a fan of lines of slope $1/\hat{V}$, each drawn over the residuals feasible at that prediction ($|r| \le \max(\hat{V}, 1 - \hat{V})$ because $V \in [0, 1]$). The line for $\hat{V} = 0.5$ is omitted: it coincides with the MSE line. (b) The Poisson-to-MSE weight ratio against the prediction on log axes, $1/(2\hat{V})$ for the unit deviance and $1/\hat{V}$ for the doubled deviance the implementation minimises; the ratio crosses 1 at $\hat{V} = 0.5$ and $\hat{V} = 1$ respectively and grows without bound as $\hat{V} \to 0$ until the clamp $\hat{V} \ge \varepsilon = 10^{-8}$, below which the derivative is zero.

$V_i > 0$. A low prediction under positive signal, not a dark target, is the singular case (for $V_i = 0$ the derivative is exactly $1/N$); the implementation's clamp $\hat{V}_i \leftarrow \max(\hat{V}_i, \varepsilon)$ zeroes the derivative below $\varepsilon$ and caps it at $1 - V_i/\varepsilon$ just above. This is the weighting implied by the Poisson likelihood for photon-counted data; the normalised, clipped and in several cases deconvolved intensities fitted here are not calibrated counts (Section 4.4).

#### 4.2 Behaviour under noise

Consider a voxel with true signal $\mu \ge 0$ and noisy observed value $V = \mu + \eta$, where $\eta$ is zero-mean and independent across voxels. We ask which constant prediction $\hat{V}$ minimises the expected loss, where the expectation is taken over the noise distribution of $\eta$:

**Proposition 2** (noise-distribution minimiser)**.** *Let $V$ have finite second moment. The minimisers of $\mathbb{E}_\eta[\mathcal{L}(\hat{V}, V)]$ for the three losses are*

$$\hat{V}^*_{\text{MSE}} = \mathbb{E}[V] = \mu, \quad \hat{V}^*_{\text{L1}} = \text{median}(V), \quad \hat{V}^*_{\text{Poi}} = \mathbb{E}[V] = \mu,$$

*where the Poisson statement additionally requires $V \ge 0$ almost surely and $\hat{V} > 0$ (the deviance is defined only there), and the median is any median when it is not unique. When $\mu = 0$ the Poisson case degenerates: $V = 0$ almost surely, the expected deviance reduces to $\hat{V}$, and the infimum is approached as $\hat{V} \to 0^+$ on the boundary of the domain, so no minimiser exists inside it.*

MSE and Poisson are unbiased for the mean; L1 targets the median. For symmetric noise these coincide. Under the Poisson-Gaussian sensor model of fluorescence microscopy [2] (a photon-count term plus additive Gaussian read noise) the noise is skewed but not heavy-tailed in the statistical sense. Skewness alone fixes no direction: the median of a Poisson or Poisson-Gaussian variate can lie on either side of its mean, so no systematic under-prediction of bright pixels by L1 follows from this proposition. What does follow is the bounded influence of Section 4.1: a single noisy bright voxel contributes a per-voxel gradient of magnitude at most $1/N$ to the L1 objective whatever its residual, whereas its contribution to the MSE objective grows in proportion to the residual. This is a per-voxel, infinite-data argument and does not directly carry over to multi-voxel fitting with shared splat parameters; it is a possible robustness mechanism, not a prediction of the magnitude or even the sign of a gap on any particular volume.

At convergence under Gaussian noise with a well-specified model, estimator efficiency gives the expected sign: the L1 minimiser is the sample median, whose asymptotic relative efficiency against the mean under Gaussian noise is $2/\pi$ [3, 4], so an L1 fit pays a variance penalty that MSE does not. The argument gives the sign of the gap and not its magnitude: the efficiency ratio is a property of the estimator, and turning it into a PSNR gap would need the ratio of fitted parameters to voxels, which this document does not derive. The fits here are early-stopped iterates, and the six volumes on which the MSE loss edges L1 by 0.04–0.28 dB (OpenCell-MAP4 nuclei, OpenCell-MAP4 microtubules, OpenCell-LMNB1 nuclei, OpenCell-LMNB1 nuclear lamina, Cells3D nuclei and Blastocyst nuclear lamina) include some of the noisiest volumes in the benchmark while the two noisy kidney channels fall on the L1 side, so the argument is offered as context, not as an explanation of which volumes fall on which side.

For finite-budget fitting it suggests, without proving, that:

- MSE, whose per-voxel weight grows with the residual, can spend more of its gradient budget on the noisiest bright voxels than L1 at the same splat pool, which would widen the train/held-out gap.
- Poisson's weighting is set by the prediction, not the residual (Section 4.1): for the unit deviance it exceeds MSE's on dark and dim voxels and falls below it only where $\hat{V}_i > 0.5$ on a unit-range volume (the implemented doubled deviance exceeds MSE's weight wherever $\hat{V}_i < 1$). It therefore behaves neither like MSE nor like L1, and its held-out behaviour has to be measured, not predicted from the single-voxel argument.

The two kidney rows of Table 4 are consistent with the bounded-influence mechanism in direction (L1 > MSE by +1.10 and +2.02 dB); the analysis does not predict the magnitude of the gap, nor does it explain why Poisson lands next to MSE on Kidney nuclei (+0.08 dB) and below it on Kidney actin (−0.44 dB).

##### 4.3 Why the PSNR-optimality argument for MSE does not reach these fits

PSNR (Eq. (1)) is a strictly decreasing function of the mean squared error, whatever the model class. Minimising $\mathcal{L}_{\text{MSE}}$ therefore minimises the exact quantity PSNR measures, and the corollary is that the global minimiser of the unregularised MSE objective within any model class, however restricted, has the highest training PSNR attainable in that class. Finite model capacity does not weaken the corollary. Provided the infimum of the mean squared error over the class is attained (an assumption, since a class of $K$ Gaussian splats is not closed: a splat can collapse or drift away), the class has a member of lowest mean squared error, and that member beats the class's L1-optimal member on training PSNR (or ties it).

The corollary is a statement about global minimisers of the unregularised objective, and the returned models here are neither:

1. **Stopped iterates, not global minimisers.** The fits are finite-iteration Adam runs on a non-convex objective that terminate by patience or at the cap; a local critical point would not suffice, and the runs were not tested for local optimality either (the residual tolerance is a distortion target, not a stationarity test). All 306 fits (102 cells × 3 reps) terminate either via early stop (no improvement of the training objective for 1,000 iterations) or via the 30,000-iteration cap, and none reach the error-based convergence criterion ($\max_i |\hat{V}_i - V_i| < 0.01$ on the normalised volume) in any replicate of any cell.

2. **Auxiliary regularisation.** The full objective (5) adds L1 regularisation on amplitudes and Cholesky diagonals to the reconstruction term, which moves the joint optimum away from the raw MSE minimum. The coefficients are identical across losses, but the reconstruction terms differ in scale and gradient structure, so the effective regularisation differs (Section 2.1).

Given that the returned models are stopped iterates, a candidate mechanism for the observed ranking is an optimisation effect at a finite budget. MSE's residual-magnitude weighting steers the gradient budget of a finite-iteration, finite-pool optimiser toward the voxels with the largest residuals (bright outliers, sharp edges), while L1's bounded weighting spreads it more evenly across the volume; the mixtures the two losses settle into differ accordingly. This is a hypothesis about optimisation trajectories, distinct from the approximation capacity of the class, and the 5,000-splat margin above the L1-selected operating point (Section 2.2) says nothing about it. The training-PSNR ranking (Table 2) is consistent with the hypothesis: MSE ranks lowest of the three losses on fourteen of seventeen volumes and highest on one, with L1 ahead by up to +10.21 dB and Poisson by up to +7.06 dB (both on Neuromast nuclei). The experiments here do not isolate the mechanism; a clean ablation would compare against MSE fits with bright-pixel residuals masked or down-weighted, and against fits with the regularisers switched off.

The Cells3D membranes result is consistent with this optimisation hypothesis: L1 leads MSE by +2.98 dB on training and +2.34 dB on held-out PSNR on a volume whose estimated noise ceiling, 53.5 dB per Supplementary Document 2, lies above every fit. It does not require an appeal to noise statistics, although it does not exclude them; MSE on clean data still spends gradient budget on the largest residuals first.

##### 4.4 Poisson-deviance practical concerns

Two reasons are commonly given for why Poisson deviance might under-perform:

1. **Volume normalisation.** The pipeline normalises input volumes to $[0, 1]$ before fitting. Poisson deviance is homogeneous of degree one under a pure rescaling, $D(cV, c\hat{V}) = c\, D(V, \hat{V})$ for $c > 0$, so a multiplicative change of units rescales the objective and its minimiser together and leaves the ranking of predictions unchanged (the ratio $V_i/\hat{V}_i$ is itself scale-free).

That identity covers only the multiplicative part of the preprocessing. The benchmark's normalisation is an affine map with percentile clipping on most volumes and a background subtraction on the neuromast channels (main Methods), under which $(V_i - b)/(\hat{V}_i - b) \neq V_i/\hat{V}_i$; several volumes are deconvolved; and none of the normalised intensities are calibrated photon counts. Poisson deviance is therefore used here as a non-negative-intensity discrepancy with a prediction-weighted gradient, not as a likelihood. Two fixed quantities do not rescale with the data, the clamp $\varepsilon = 10^{-8}$ and the regulariser coefficients of Eq. (5), so the normalised scale does matter through them.

2. **Dark-voxel amplification.** The $1/\hat{V}_i$ weighting gives a voxel whose prediction is far below a positive target a very large gradient (Section 4.1), which early in optimisation, before $\hat{V}$ has learned the background, could pull splats toward under-predicted regions instead of onto the signal.

Empirically, neither effect prevented any Poisson fit from completing, and the clamp keeps the objective finite; but the clamp bounds the gradient only just above $\varepsilon$, and Adam normalises the aggregate gradient of each parameter after the voxel contributions have been summed, so neither mechanism limits the influence of an individual voxel. Across $n = 3$ replications, Poisson uses fewer training-mode iterations than MSE on twelve of seventeen volumes (mean ratio 7.6–13.3× on the three volumes where MSE runs to the cap and Poisson does not; Table 5) and is the highest-mean training PSNR on nine of seventeen (Table 2; L1 is highest on Cells3D membranes, Tribolium embryo nuclei, Mouse heart nuclei, Zebrafish embryo nuclei, Drosophila embryo nuclei, Neuromast membranes and Neuromast nuclei, MSE on OpenCell-MAP4 microtubules).

### 5 Interpretation

**Training PSNR.** MSE is not the best training-PSNR loss for the regularised, early-stopped fits measured here (Section 3.1), although the global minimiser of the unregularised MSE objective would be: since PSNR is a decreasing function of the mean squared error, minimising $\mathcal{L}_{\text{MSE}}$ maximises PSNR at that minimiser within any model class. Its one win, on OpenCell-MAP4 microtubules, is by 0.02 dB over Poisson, and the L1−MSE gap runs from −0.25 dB on that spinning-disk channel to +10.21 dB on Neuromast nuclei (Table 2). The PSNR-optimality argument is a statement about the global minimiser of the unregularised objective (a critical point is not sufficient for a non-convex mixture fit), and finite-iteration Adam under early stopping on a regularised objective does not deliver it. It delivers it least of all for MSE, the loss that most often runs to the iteration cap without reaching the residual tolerance (Table 5; that tolerance is a distortion target, not a stationarity test). Section 4.3 unpacks the assumptions.

**Held-out PSNR.** The held-out ranking of Section 3.3 has a clear structure. L1's wins over MSE are decisive on the noisy or dense confocal volumes, on the Tribolium and Mouse heart light-sheet volumes and on both iSIM volumes, and on the *C. elegans* embryo and the Fly brain neurons Poisson is marginally ahead of both. The volumes on which the MSE loss edges L1 are exactly the tight ones, where all three losses sit within 0.3 dB of one another. These tight volumes are not a converged regime: on Cells3D nuclei MSE runs to the cap in every training replicate and on the blastocyst L1 does (Table 5), and a run that stops by patience has reached a plateau of its training objective, not a stationary point.

Two candidate mechanisms could contribute to the large L1 margins. The first is *noise-robustness*: L1's bounded per-voxel influence limits how much any one noisy bright voxel can move the shared parameters, while MSE's residual-proportional weighting lets bright-pixel noise be memorised (Section 4.2). The second is a *finite-budget optimisation* effect: MSE's residual-magnitude weighting steers a finite-iteration, finite-pool optimiser toward the largest residuals at the expense of the rest of the volume (Section 4.3). The two kidney gaps are plausibly dominated by the first: they are noisy volumes whose canonical sweep in Supplementary Document 2 shows their training PSNR overshooting the noise ceiling (the PSNR at which a reconstruction's residual equals the estimated noise alone) at large $K$. For Cells3D membranes that ceiling (53.5 dB) lies above every fit (training PSNR 47.27–50.25 dB), so approximation error plausibly contributes to the residual there. The ceiling is itself an estimate, and a scale comparison of this kind neither decomposes the residual into noise and structure nor says why two losses differ. The present experiments do not separate the two mechanisms and do not rule out partial overlap.

**What the held-out numbers score.** Every entry of Table 4 is the held-out PSNR of the model the fitter returns: its best-state snapshot, the parameters one optimiser step after the lowest training objective (5) seen before the loss-dependent stop (patience or cap), after the final 0.999-retention cull. The ranking therefore combines the reconstruction loss with its stopping behaviour and post-processing, and holds for these end-to-end configurations, not for the loss in isolation or at matched convergence. The held-out PSNR was also tracked every 250 iterations during each masked-volume fit; the tracked maximum exceeds the returned endpoint by more than 0.5 dB in 7 of the 51 (volume, loss) cells, by up to 1.69 dB (Neuromast membranes, L1). Figure 4 places each replicate's tracked maximum and returned endpoint against the iteration at which they occurred. Those maxima are taken on the same mask that scores them and on intermediate, un-culled predictions, so they are optimistic and are not reported as an alternative endpoint; they do show that part of a gap in Table 4 can be a stopping or post-processing effect, not a property of the loss. Separating the two would need checkpoint trajectories scored on an independent mask, which this study does not include.

**Poisson deviance.** Poisson deviance is the best training-PSNR loss on nine and the best training-SSIM loss on nine of seventeen volumes (Tables 2, 3). On held-out PSNR it sits next to MSE (Section 3.3), above it on eleven volumes by up to +4.86 dB (Neuromast nuclei). Relative to L1 it is ahead on eight volumes, by +0.03 to +0.27 dB: on the six of them inside the 0.3 dB band by +0.07 to +0.20 dB, and outside that band only on the C. elegans embryo nuclei and Fly brain neurons. It trails L1 by up to 2.92 dB (Neuromast nuclei; −2.46 dB on Kidney actin), so it captures only part of L1's advantage where that advantage is large.

Its distinguishing property is early termination on the confocal volumes (Section 3.4). The wall-clock consequence is largest on the noisy kidney volumes: Poisson reaches a training PSNR of 38.71 and 40.55 dB in 2.4 and 2.1 min, where MSE needs 20.2 and 20.9 min for 37.89 and

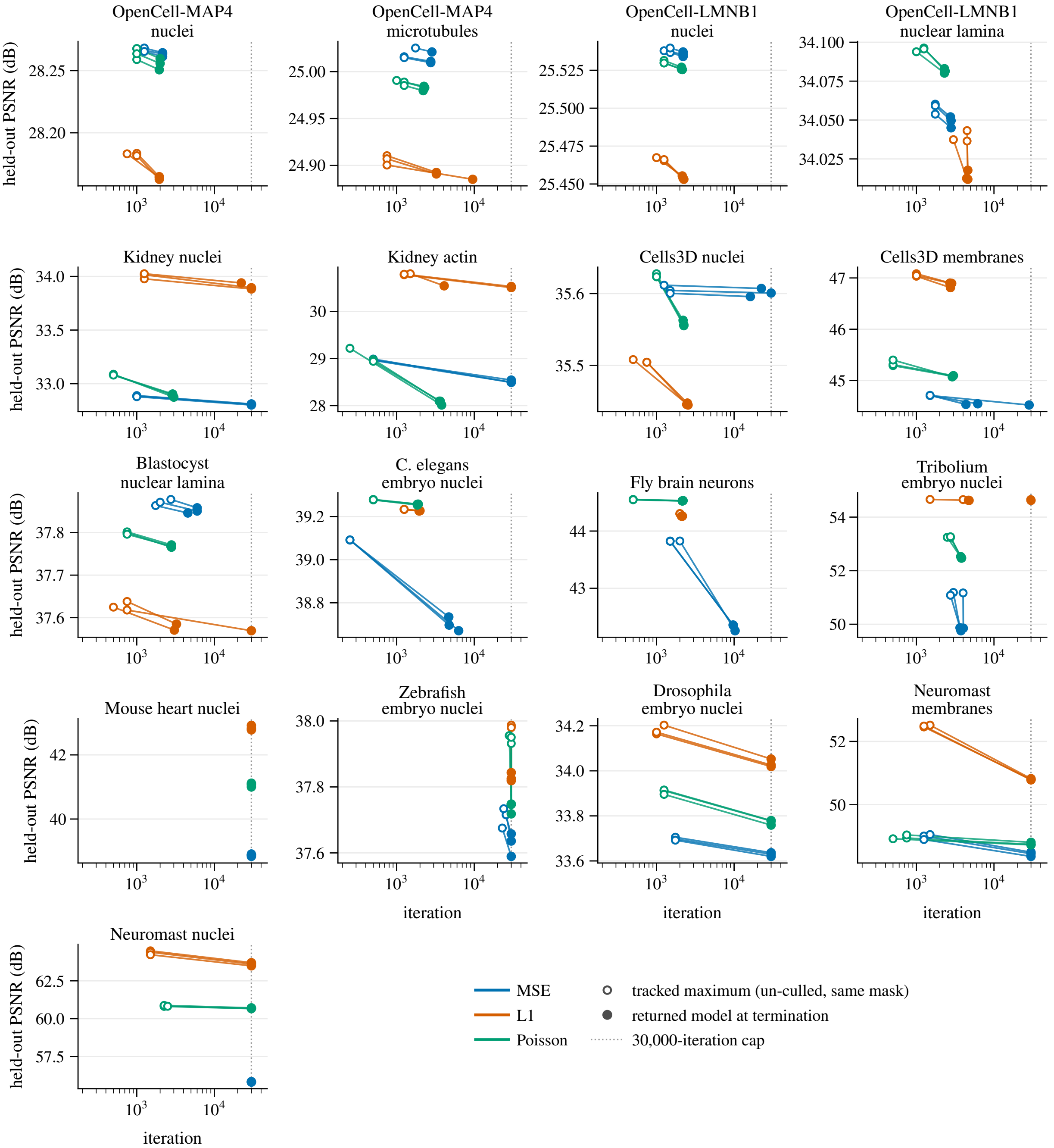


Figure 4: Held-out PSNR against iteration in the masked-volume fits, one panel per volume in registry order. For every replicate of every loss an open marker is the tracked maximum of the held-out PSNR (sampled every 250 iterations, scored on the same mask, on the un-culled intermediate model) and a filled marker is the model the fitter returns at termination (best-state snapshot after the loss-dependent stop and the 0.999-retention cull, the value tabulated in Table 4); the line between them is a connector, not the trajectory, which was not stored. The dotted line is the 30,000-iteration cap. Where the maximum sits far to the left of the endpoint, the fit kept running long after its held-out score peaked; where the two markers coincide, the returned model scores the same as the tracked best to the resolution of the plot.

38.31 dB (Table 6), all on a shared GPU. Where every loss runs to the cap, Poisson has no iteration advantage. The numerical concerns about dark-voxel amplification discussed in Section 4.4 did not prevent any fit from completing, but neither the $\hat{V}_i \geq \varepsilon$ clamp nor Adam bounds the influence of an individual voxel (Section 4.1).

**Structural similarity and appearance.** The SSIM exceptions of Section 3.2 are real: on the Fly brain neurons MSE reaches 0.844 against 0.959 (L1) and 0.970 (Poisson), a range of 0.126 that is many replicate standard deviations wide, and on the Mouse heart nuclei the range is 0.033. Figure 5 shows, for two volumes on which L1 leads MSE on held-out PSNR, Neuromast nuclei (the largest gap) and the noisy confocal Kidney actin, the middle slice of the training-mode fit (replicate 1) and the signed difference between each reconstruction and the target: the three reconstructions are visually similar at the inspection scale, and the difference images are where the losses separate. Being training fits they do not show the visual cost of the held-out differences, and the figure does not cover the Fly brain, where the SSIM difference is largest.

### 6 Conclusion

The loss a splat fitter minimises is inseparable from how it stops. MSE is the PSNR-optimal loss only at the global minimiser of an unregularised objective, and the fitter never delivers that object: it returns a stopped iterate of a regularised, non-convex objective, and it is MSE that most often runs to the iteration cap without settling. Judged on what the fitter actually returns, MSE is the best training-PSNR loss on one volume and the best held-out loss on none of the volumes where the losses differ by more than 0.3 dB (Section 3). What matters is which mixture each loss settles into, not which target residual it would minimise at a global optimum.

On held-out PSNR, the metric this study selects models by, L1 is the safe default. Its wins over the MSE loss are large where the data are noisy or dense, on the deconvolved iSIM volumes and on the Tribolium and Mouse heart light-sheet volumes, and its losses are confined to the tight volumes, where they never exceed 0.28 dB. The win count alone is not significant at $n = 17$ (Section 3.3); the recommendation rests on the asymmetry of the gap sizes. Two mechanisms are consistent with the large gaps, L1's bounded per-voxel influence under noise (Section 4.2) and a finite-budget optimisation effect of MSE's residual weighting (Section 4.3); the experiments do not separate them, and every held-out entry scores an endpoint that mixes the loss with its stopping rule and post-processing (Section 5). The inter-loss SSIM range is $\leq 0.02$ on fifteen of the seventeen volumes and wider on Fly brain neurons (0.126) and Mouse heart nuclei (0.033) (Section 3.2); the slice montages are visually similar at the inspection scale used, and the difference images (Figure 5) locate the residual that separates the losses at the boundaries and interiors of the bright structures.

Poisson deviance is the alternative when time to termination on confocal data matters. It tracks MSE on held-out PSNR, gives up part of L1's advantage where that advantage is large, and stops early where MSE and L1 run to the cap, which makes it the fastest loss on ten of seventeen volumes on this shared GPU (Section 3.5). Where all three losses run to the cap it offers no speed advantage.

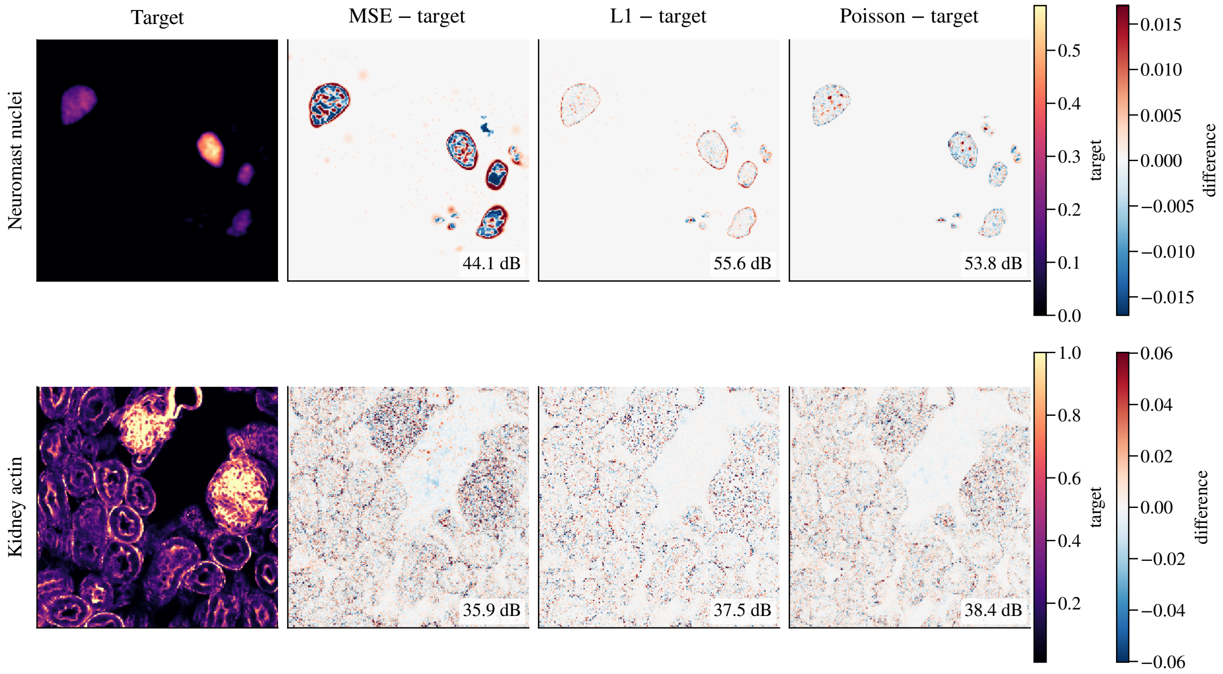


Figure 5: Difference images of the training-mode fits (replicate 1) on the two headline volumes. Each row shows the target of the middle slice along the most-square axis and, on one diverging scale per row (symmetric at the 99.5th percentile of the absolute difference over the three panels), the signed difference reconstruction − target for the MSE, L1 and Poisson fits; red is over-prediction, blue under-prediction. Top: Neuromast nuclei (69,000 splats), cropped to the region of the slice that holds signal. Bottom: Kidney actin (133,000 splats), cropped to the 256-voxel window of highest brightness and edge content. On the Neuromast nuclei the MSE fit leaves a rim of over-prediction around every nucleus and speckle inside it that L1 and Poisson largely remove; on the Kidney actin the MSE residual is densest inside the bright structure, where L1 and Poisson leave a lighter, less structured residual. The label in each difference panel is the PSNR of that 2-D slice, computed from the displayed arrays and referenced to the target's dynamic range: a per-slice diagnostic, not the volume-wide score of the tables (Table 4 gives L1 − MSE of +7.78 and +2.02 dB on held-out PSNR for these volumes).

These conclusions hold for one end-to-end configuration: one stopping rule (patience 1,000 under a 30,000-iteration cap, best-state snapshot, 0.999-retention cull), one L1-derived operating point ($K^*$ plus 5,000 splats, a convention at the sweep endpoint for the two signal-limited light-sheet volumes) and fixed regularisers. They say nothing about the losses at matched convergence or at per-loss operating points, about 4-D data or synthetic noise, or about intrinsic per-iteration cost, which the shared GPU did not allow us to measure. The three seeded replicates bound seed variability, small for PSNR ($\leq 0.24$ dB) and SSIM ($\leq 0.0034$) and large only for the mixed cap cells, and say nothing about variability across specimens or masks. A per-loss operating point, a regulariser ablation and timing on an idle device are the natural follow-ups.
