## Supplementary material for "Luxar: Gaussian splatting for microscopy and scalable interactive web visualisation of multidimensional scientific data": supp_doc_06_viewer_performance

### Viewer rendering performance: count, size, and viewport sweeps

Supplementary Document 6 – Luxar


**Abstract**

A browser viewer feels interactive when it draws a new frame at every display refresh, so the practical question is how large a scene fits inside one 16.7 ms refresh interval. We measured the Luxar web viewer on an NVIDIA RTX 3070 in headless Chromium across three sweeps: element count from $10^3$ to $10^7$ for points, lines and Gaussian splats, nominal on-screen size from about one pixel to thousands of pixels per element, and viewport from 720p to 4K. Every cell records the wall time of one synchronised render-and-readback call, which scales with the scene, and the callback rate of the browser's animation loop, capped at the refresh cadence. Up to $10^6$ points, lines or Gaussian splats the callback rate (cell medians) holds at least 59.3 FPS, and one synchronised call takes 1.21 to 1.72 ms at $10^5$ elements, well inside the budget. At $10^7$ points or splats the call exceeds the budget by 12.3 to 16.3%. Cost is set by covered pixels: above a nominal 16 px per element the latency grows close to quadratically with size, and on the largest elements the callback intervals of the rendering phase stretched to 300 ms; whether each is one long frame or two frames of the call's length the record does not say. The HDR render target adds a coverage-dependent cost, while the fused post-processing pass costs at most a millisecond at 720p. A scene over budget is therefore rescued by reducing the pixels its elements cover, not by disabling post-processing.


#### 1 Introduction

The Luxar viewer renders four first-class geometries (points, lines, meshes and Gaussian splats) in a WebGL pipeline that targets 60 frames per second. A browser presents at most one frame per tick of its begin-frame cadence, so "interactive" has a concrete meaning: the viewer's frame must fit inside one refresh interval, 16.7 ms at 60 Hz, and the frame rate a user can observe never exceeds that cadence. Two questions follow: up to what scene size the viewer keeps every refresh, and how the cost of one frame grows with element count, element size and viewport once it does not.

The two quantities measured here answer those questions from opposite sides. The callback rate of a *rAF* loop is what a page can observe: it saturates at the cadence and moves only once frames overrun it, so it says whether the cap was kept and little else. The latency of one synchronised render-and-readback call has no cap: it is the wall time from issuing one render until a one-pixel readback returns, and it scales with the scene. The call serialises work that a running loop would overlap, so the two numbers are not the same quantity and neither is a delivered frame time; §2.1 defines both and §4.1 says how to read them together.

The sweeps cover points, lines and Gaussian splats. Meshes are omitted because the synthetic scene generator emits none and a mesh has no per-element size, so it does not fit the size sweep's calibrated pixel grid. Throughout, a *cell* is one combination of geometry, element count, nominal size and viewport in the measurement plan; each cell is measured under three render-call variants and three repeats.

##### Terms used in this document

**rAF** `requestAnimationFrame`, the browser callback that fires once per begin-frame tick (the display refresh when there is one); a page's rAF rate is capped at that cadence and counts callbacks, not frames the viewer drew, so on its own it is not a frame rate.

**HDR and LDR** High versus low dynamic range: HDR colours are unbounded floating-point intensities (supplied as float32; the AUTO encoding stores them as per-channel logarithmic `uint16` and decodes back to float32, PRECISION keeps float32), rendered into a half-float target, while LDR colours are clamped to $[0, 1]$ as a display expects; tone mapping converts the first into the second.

**DPR** The device pixel ratio, the number of physical pixels per CSS pixel; the viewer renders at DPR 1.0 unless high DPR is explicitly allowed, and an adaptive controller may walk the render resolution below the live window value when the frame rate drops.

**anti-aliasing (MSAA, SSAA, FXAA, SMAA)** Techniques that soften the stair-stepping of rasterised edges: SSAA shades several samples per pixel and averages them, MSAA shades once per pixel and keeps per-sample coverage, while FXAA and SMAA are post-process filters that smooth edges detected in the finished image.

#### 2 Methods

##### 2.1 The two metrics

**rAF rate.** The harness runs a `requestAnimationFrame` loop beside the viewer's own animation loop for a 3 s window and records the timestamp of every callback. The rate is the reciprocal of the mean interval $\Delta t$ between successive callbacks (frames per elapsed second). It is capped at the runtime's *begin-frame cadence*, the periodic tick at which headless Chromium schedules a frame and delivers animation callbacks, about 60 Hz in this runtime (headless Chromium has no display to synchronise with). Callbacks are delivered in the same per-tick batch as the viewer's frame, so when a viewer frame overruns the tick the whole batch slips and the harness sees a longer interval. The loop does not count the viewer's render calls, and it keeps ticking at the cadence when the viewer's loop goes idle: the viewer's animation controller pauses after inactivity, and nothing in the measurement kept it awake (§2.6).

The reciprocal of the mean interval is the right estimator. The mean of per-callback instantaneous rates, $\overline{1/\Delta t}$, is biased high under jitter by Jensen's inequality: one long interval contributes a single small term while the short intervals around it contribute many large ones. The two coincide only when every interval is equal, that is on cells pinned at the cap. The runner also stores the median and 95th-percentile interval and the raw per-callback intervals of every repeat, which §3.2 uses to describe the windows of the heavy cells. The rAF loop always observes the viewer's default pipeline; the variant label on an rAF value names the latency measurement it was paired with, not a change of rendering path.

**Synchronised render-and-readback call.** The latency is the wall-clock time from just before a single render call until a one-pixel `gl.readPixels` issued right after it returns, in a tight loop; the mean over the loop is the per-repeat value. The readback forces the driver to flush all queued commands and complete the preceding rendering to the framebuffer it reads. Because the clock stops when the read returns, the number includes CPU submission, driver and compositor work as well as GPU execution. It is a synchronous end-to-end render cost, not an isolated GPU-timer (`EXT_disjoint_timer_query`) measurement, and we do not call it GPU time. In this harness `gl.finish()` and `gl.fenceSync` with `gl.clientWaitSync` returned sub-microsecond timings on default-backbuffer renders that take tens of milliseconds under `readPixels`, so neither is used as the synchronisation point; the cause was not isolated (§2.6).

The call measures one render plus one readback. It stops on the readback, so it serialises work that a running loop pipelines, and the ratio of the 16.7 ms budget to the call compares two durations and predicts no frame rate. On the heavy size cells the raw rAF intervals of the rendering phase were 1.6 to 1.9 times the synchronised call (§3.2); whether each is one long frame or two frames of the call's length the rows do not say, so the call is not a bound on a frame time in either direction. It is the number that scales with the scene, and we use it for that.

**Aggregation.** Every cell is measured three times, and the central value is the median across the three repeats, not the mean. A repeat can carry a degenerate rAF reading, and the deposited rows hold more than one. On gsplats at 64 px (`post_off_hdr`) repeat 0 holds 1 interval (0.02 s) and reads 59.88 FPS against siblings at 28.62/28.62. On points at 64 px (`post_off_hdr`) repeat 0 holds 72 intervals over 1.20 s and reads 60.00 against 35.47/36.46. On gsplats at 32 px (`post_off_hdr`) repeat 0 has a 2.33 s window and reads 50.57 against 43.76/44.42. Table 1 lists the window length and rate of every repeat of the 14 cell-variant groups with a repeat below 59.9 FPS. The median rejects one such repeat out of three; the mean does not. The error bars in Figs. 4–7 are the sample standard deviation of the three per-repeat means, plotted about the median.

We avoid the framing "renders per second" for the latency metric. The inverse of the mean latency is the throughput a tight loop would achieve, but it is not the rate at which a user sees new frames, which the cadence caps at about 60.

#### 2.2 Synthetic scenes

To separate element count from element size we generate three families of synthetic Luxar scenes (Table 2). All cells place their elements in $[-0.5, 0.5]^3$ and the viewer auto-frames the bounding box. The count $N$ is the number of points, the number of Gaussian splats, or, for lines, the number of vertices of a single polyline, which has $N - 1$ segments. The polyline is a random walk whose step standard deviation is $1/\sqrt{N}$, rescaled to fit the box, so its segment length shrinks with $N$ along the count axis while its extent stays fixed.

The size axis is nominal. The viewer's default vertical field of view is 47° and it computes its own framing distance; the size calibration instead assumes a 60° field of view and a framing distance of 1.5 world units for the default 1280×720 viewport, a framing assumption that gives about 416 pixels per world unit at the focus plane. A nominal target of 4 pixels then corresponds to a world radius of about 0.0096. The projected footprint was not recorded, so the pixel sizes below are planning values. The nominal size also names a different quantity per geometry: the visible disc radius for points, the Gaussian standard deviation $\sigma$ for gsplats (visible extent about

Table 1: Per-repeat rAF window (number of intervals, summed duration in seconds) and callback rate for every cell-variant group in which any repeat reads below 59.9 FPS (14 groups). A short window with a reading at the cap beside sub-cap siblings is a degenerate repeat, which the median rejects.

| Cell | Variant | rAF window per repeat: intervals / s / FPS | | |
|---|---|---|---|---|
| | | repeat 0 | repeat 1 | repeat 2 |
| points $10^6$ | `post_off` | 181 / 3.02 / 60.00 | 173 / 3.02 / 57.35 | 179 / 3.02 / 59.34 |
| points $10^6$ | `post_on` | 181 / 3.02 / 60.00 | 178 / 3.02 / 59.01 | 181 / 3.02 / 60.00 |
| points $10^7$ | `post_off_hdr` | 181 / 3.03 / 59.67 | 181 / 3.02 / 60.00 | 177 / 2.98 / 59.33 |
| gsplats $10^7$ | `post_off_hdr` | 179 / 3.02 / 59.34 | 181 / 3.02 / 60.00 | 181 / 3.02 / 60.00 |
| gsplats $10^7$ | `post_on` | 182 / 3.03 / 60.00 | 181 / 3.02 / 60.00 | 171 / 3.02 / 56.69 |
| points 64 px | `post_off` | 107 / 3.02 / 35.47 | 109 / 3.02 / 36.13 | 107 / 3.02 / 35.47 |
| points 64 px | `post_off_hdr` | 72 / 1.20 / 60.00 | 107 / 3.02 / 35.47 | 110 / 3.02 / 36.46 |
| points 64 px | `post_on` | 106 / 3.02 / 35.14 | 107 / 3.02 / 35.47 | 110 / 3.02 / 36.47 |
| gsplats 32 px | `post_off` | 132 / 3.02 / 43.76 | 132 / 3.02 / 43.76 | 133 / 3.02 / 44.09 |
| gsplats 32 px | `post_off_hdr` | 118 / 2.33 / 50.57 | 132 / 3.02 / 43.76 | 134 / 3.02 / 44.42 |
| gsplats 32 px | `post_on` | 134 / 3.02 / 44.42 | 134 / 3.02 / 44.42 | 133 / 3.02 / 44.09 |
| gsplats 64 px | `post_off` | 83 / 2.88 / 28.79 | 83 / 2.90 / 28.62 | 83 / 2.90 / 28.62 |
| gsplats 64 px | `post_off_hdr` | 1 / 0.02 / 59.88 | 83 / 2.90 / 28.62 | 83 / 2.90 / 28.62 |
| gsplats 64 px | `post_on` | 82 / 2.88 / 28.44 | 83 / 2.90 / 28.62 | 82 / 2.88 / 28.44 |

Table 2: The three sweeps. Lines stop at $N = 10^6$ in sweep A ($10^7$ line vertices is heavy on disk and the line shader is slower), so sweep A is 5 levels × 2 geometries plus 4 levels × 1 geometry, 14 cells. Sweep C reuses the $10^5$-point and $10^5$-gsplat cells of sweep A at three further viewports under `post_on` only, hence "+6" plan entries and no new cells. Unique cells: 29; plan size with three render-call variants and three repeats: $29 \times 3 \times 3 + 6 \times 1 \times 3 = 279$ measurements.

| Sweep | Geometry | N levels | Size levels | Cells |
|---|---|---|---|---|
| A. Count | points / lines / gsplats | 5 ($10^3$–$10^7$) | fixed reference | 14 |
| B. Size | points / lines / gsplats | fixed ($10^5$) | 5 (px-calibrated) | 15 |
| C. Viewport | points / gsplats | fixed ($10^5$) | fixed reference | + 6 |

$3\sigma$) and the stroke half-width for lines. The same nominal value is therefore not the same screen footprint across the three curves. Figure 1 draws the calibration camera and the reference glyphs at pixel scale.

For sweep B the nominal sizes are 1, 4, 16, 32 and 64 px for points and gsplats, and 0.5, 2, 4, 8 and 16 px for lines. The endpoints bracket a footprint of about one pixel per element and one of thousands of pixels per element. Because the footprints are nominal, the per-fragment regime is inferred from the measured curves (§3.2), not from the parameter value.

##### 2.3 Render-call variants

The latency of every count and size cell is measured under three render-call variants, which separate the cost of the render target's dynamic range (*HDR and LDR*) from the cost of the post-processing pass. The additional viewports of sweep C use `post_on` only. Figure 2 draws the three paths and the two differences taken between them.

- `post_off`: the scene rendered into the default RGBA8 backbuffer, four bytes per pixel. Pure scene cost in LDR.

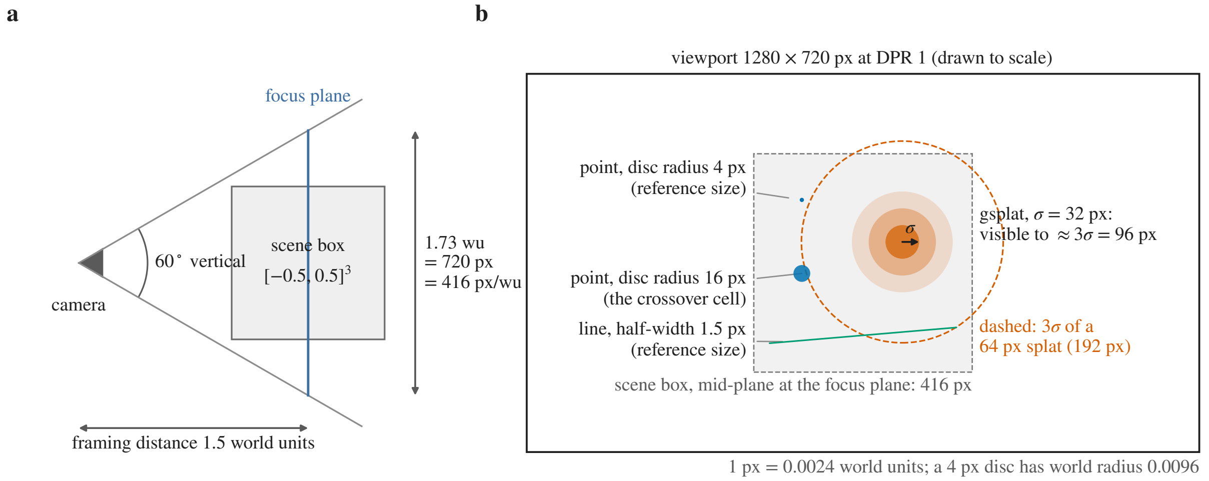


Figure 1: The nominal-size calibration (schematic). (a) The planning camera: a 60° vertical field of view at a framing distance of 1.5 world units puts 1.73 world units across the 720 rows of the viewport, about 416 px per world unit at the focus plane. The viewer's own default field of view is 47° and it computes its own framing distance, so these are planning values and the projected footprint was not recorded. (b) The 1280×720 viewport drawn to scale with the scene box's mid-plane cross-section at the focus plane (416 px; the front face, at 1.0 world units, would project at 624 px and the back face at 312 px) and the reference glyphs at true pixel scale: the 4 px disc and the 1.5 px half-width line of the count sweep, the 16 px disc at the crossover of the size sweep, and a $\sigma = 32$ px splat visible to about $3\sigma$; the dashed circle is the $3\sigma$ extent of a 64 px splat. The same nominal value covers a different area per geometry.

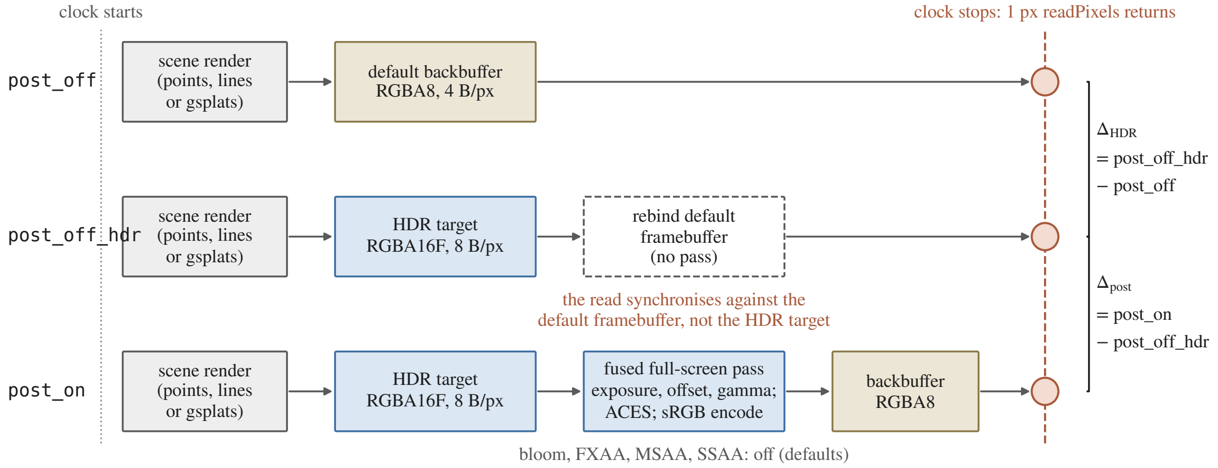


Figure 2: The three render-call variants (schematic). Each variant renders the scene once, and the clock runs from just before the render call until a one-pixel `readPixels` returns (the dashed column). `post_off` draws into the default RGBA8 backbuffer. `post_off_hdr` draws into the RGBA16F target and rebinds the default framebuffer with no pass, so its read synchronises against the default framebuffer and not the target it drew. `post_on` draws into the RGBA16F target and runs the single fused pass into the backbuffer, with bloom and anti-aliasing off at their defaults. The brackets are the two differences the overhead is decomposed into.

- `post_off_hdr`: the same scene render, but into the pipeline's HDR target, an RGBA16F texture (four half-float channels, eight bytes per pixel), with no post-processing pass. The harness binds the HDR target, renders the scene, and rebinds the default framebuffer. This isolates the cost of writing the scene into an HDR float target relative to LDR, with the synchronisation caveat of §2.6.
- `post_on`: the viewer's post-processing pipeline as configured for the scene: the scene is rendered into the RGBA16F target, then a single fused full-screen pass applies exposure, offset and gamma, tone mapping (the ACES filmic curve, Academy Color Encoding System, by default) and sRGB encoding into the backbuffer. Bloom (a luminance-threshold pass plus a mipmap pyramid whose output the fused pass mixes in) and FXAA (a separate final pass) are optional stages that the pipeline runs only when enabled, as are MSAA and SSAA; the synthetic scenes carry no viewer configuration and the harness sets none, so all four are at their default, off. `post_on` therefore measures the default pipeline, HDR target plus the fused pass, without bloom or *anti-aliasing (MSAA, SSAA, FXAA, SMAA).*

The two differences attribute the post-processing overhead:

- $\Delta_{\text{HDR}} =$ `post_off_hdr` $-$ `post_off` is the measured cost of the RGBA16F render target relative to RGBA8. An RGBA16F pixel is 8 bytes against 4, so this term is expected to track per-fragment bandwidth and to be geometry-independent at fixed coverage; it is an empirical timing difference between two runs, not an isolated stage cost.
- $\Delta_{\text{post}} =$ `post_on` $-$ `post_off_hdr` is the cost of the fused full-screen pass: one full-screen fragment pass for each rendered image, whose cost depends on the viewport size and not on the scene content when bloom is off, as here.

##### 2.4 Hardware, runtime and harness

All measurements were collected on an NVIDIA RTX 3070 through Playwright driving headless Chromium (`chrome-headless-shell`) on the GPU, the runtime of the main paper's Supplementary Table 1. Chromium was launched with Vulkan through ANGLE, Chromium's graphics layer that translates WebGL to a native API (`--use-angle=vulkan`, `--enable-features=Vulkan`). GPU rasterisation, WebGL and WebGL2 were enabled, the GPU blocklist ignored and precise memory information enabled. Background timer throttling and renderer backgrounding were disabled; the two stop Chromium from throttling timers and animation frames in a non-foreground renderer, so they bear on the rAF cadence. GPU acceleration, as opposed to SwiftShader (Chromium's CPU software renderer, the fallback when no GPU is usable), is confirmed per cell from the `WEBGL_debug_renderer_info` extension. Every deposited row records the renderer string verbatim: `ANGLE (NVIDIA, Vulkan 1.4.312 (NVIDIA NVIDIA GeForce RTX 3070 (0x00002484)), NVIDIA)`.

Each row records that renderer string, the viewport and, for the $10^6$ and $10^7$ cells, the settled drawn-element count and the cache budget; the run's metadata records the browser mode (headless) and the run timestamp once. The deposit does not record the DPR pin, the launch flags, the harness revision, the viewer commit, the Chromium, Playwright or NVIDIA driver versions, the operating-system version, the CPU and memory, the machine load, the resolved rendering settings (bloom, anti-aliasing and the line primitive are taken to be at their defaults, §2.3 and §4.5) or the camera projection. The run is therefore pinned by GPU, backend and timestamp, and the settings above are asserted by the harness, not evidenced by the rows.

**Adaptive DPR is disabled.** The viewer ships with an adaptive device-pixel-ratio (*DPR*) controller that lowers the render scale when the measured frame rate drops. Evaluated every 500 ms, it steps the render scale down by 10% whenever the rAF rate falls below 0.75 of the estimated refresh cap (45 FPS at 60 Hz), one step per probe window, down to a floor of DPR = 0.5 ($640 \times 360$ at this viewport), and steps back up once the rate has stayed above 0.9 of the cap for 3 s. Each step is probed for 1.5 s and reverted, with the pre-step scale learned as a floor, if it does not raise the rate by at least 5%. Left on, it would conflate raw rendering cost at the stated viewport with a dynamically adapted resolution, so the benchmark disables it and pins the DPR to 1.0 immediately after load and before any measurement. Every number here is the cost at the stated viewport's pixel count ($1280 \times 720$ for the count and size sweeps).

**Camera.** The camera is static during measurement. Both measurement loops write a camera position on every iteration and then update the orbit controls, but the controls re-apply their stored pose on update and require an explicit re-initialisation after an external position write, which the harness does not perform. The render call and the readback execute on every iteration regardless (three.js issues the full draw whether or not the camera moved), so the latency is the cost of drawing the scene from the auto-framed view.

**Settling and residency.** For the $10^6$ and $10^7$ count cells the harness settles before measuring: it polls the viewer's drawn-element count every 0.5 s and proceeds only once the count has been unchanged, with no load pass running, for 8 s. The settled count, the per-geometry counts and the dropped-element count are recorded in every such row, so a cell that is not fully *resident*, one whose elements were not all loaded, uploaded and drawn, is visible in the data. The viewer's *cache budget*, the total in-memory allowance for its data caches, is set to 2048 MB through the `?cacheBudgetMB=` parameter, the value the native launcher passes. Under it the settled $10^7$ cells draw all $10^7$ elements with none dropped, and a probe of the same cells at the default budget also records $10^7$ drawn and none dropped. The $10^6$ line cell settles at 999,999 drawn segments, one polyline of $10^6$ vertices having $N - 1$ segments.

**Isolation.** Each cell runs in its own fresh browser context, the browser process is recycled every four cells, and the viewer's readiness predicate is polled every 500 ms instead of at animation-frame rate. All three keep the viewer's initialisation from being disturbed on heavy cells: state accumulated in the GPU process across several sequential cells otherwise breaks the next cell's initialisation, and a predicate polled at 60 Hz competes with the viewer for main-thread time. Variants and viewports within a cell are exercised on the same loaded page (one navigation per cell), because each navigation discards the WebGL context. Each cell is served with `luxar serve <cell.zarr> --viewer`.

**Plan and window.** Figure 3 lays out the sequence for one cell. The full plan is 279 measurements (Table 2). The rAF window's deadline is armed before 30 warm-up callbacks and

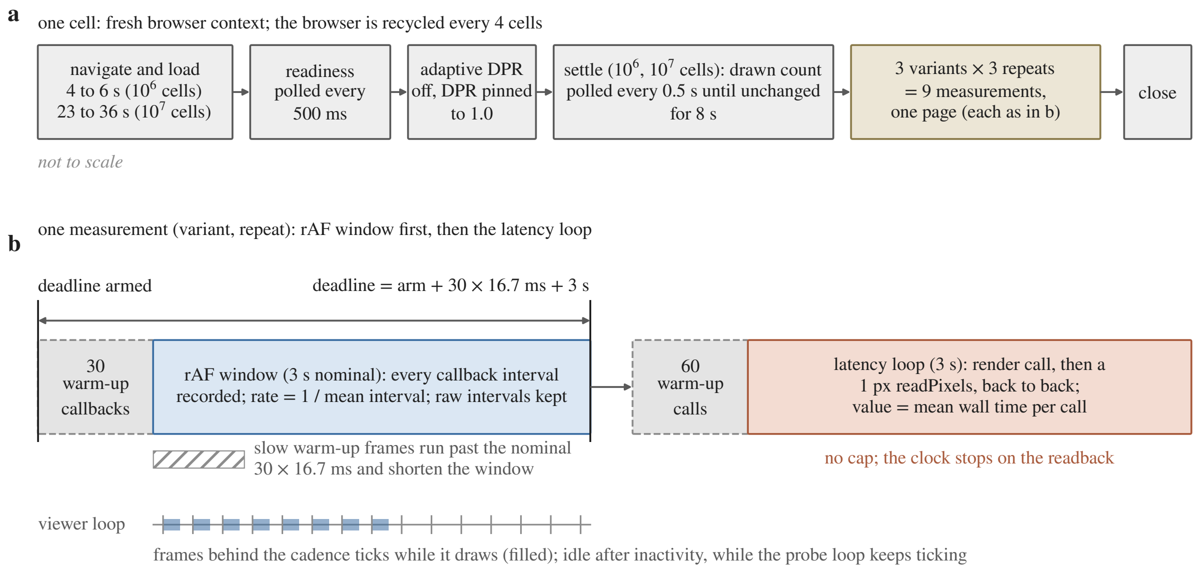


Figure 3: The measurement sequence of one cell (schematic, not to scale). (a) A fresh browser context loads the cell; the harness polls the viewer's readiness every 500 ms, pins the DPR to 1.0 and, on the $10^6$ and $10^7$ cells, waits for the drawn-element count to hold for 8 s; the nine measurements (three variants, three repeats) then run on the same page. (b) One measurement: the rAF window (30 warm-up callbacks, then a nominal 3 s of recorded callback intervals) followed by the latency loop (60 warm-up synchronised calls, then 3 s of render-and-readback calls). The window's deadline is armed before the warm-up and extended by a nominal $30 \times 16.7$ ms, so slow warm-up frames shorten the window. The viewer's own loop draws frames behind the cadence ticks while it is active and goes idle after inactivity, while the probe loop ticks on either way.

extended by a nominal $30 \times 16.7$ ms, not by the warm-up's actual duration, so slow initial frames shorten the measured window; per-repeat sample counts range from 1 to 182 intervals (summed duration 0.017 to 3.03 s). The latency loop warms up with 60 synchronised calls and then measures for a full 3 s. Per-cell load times are 4–6 s for the $10^6$ cells and 23–36 s for the $10^7$ cells, each followed by the 8 s settle; the total wall-clock time of the sweep was not logged.

#### 2.5 Reproducibility

The synthetic scene generator, the benchmark harness, the plotting step and the build of this document are deposited with the analysis; they are not part of the Luxar software tree. The generator writes the 29 cells, each with a random seed derived from the SHA-256 of its name (the leading 32 bits reduced modulo $2^{31} - 1$) and recorded in a manifest beside the cells; the manifest embedded in the deposited sweep data carries no seeds. The harness runs headless Chromium on ANGLE/Vulkan with the explicit 2048 MB cache budget and three repeats; headed Chromium on the default backend is not the reported configuration. The deposited harness differs from the one that produced the deposited sweep in four respects: it arms the rAF window's deadline at the first tick after the warm-up, it calls the viewer's `renderOnce` keep-alive on every tick, it re-initialises the orbit controls after each camera write, and it settles the drawn-element count on every cell. In the deposited data the rows at $N \geq 10^6$ come from a settled pass over those cells with that cache budget (one filtered run supplies both the $10^6$ and the $10^7$ rows); rows of those cells from other passes and the default-budget probe of the $10^7$ cells are retained beside them and are not used here. The plotting step writes the figures and the per-cell tables; the tables include all three render-call variants, report medians across the three repeats and are the source of truth for any follow-up analysis.

##### 2.6 Limitations of the measurement

Four properties of the instrument bear on how the numbers should be read.

**An idle viewer loop reads as 60 FPS.** The rAF loop ticks at the cadence whether or not the viewer draws a frame behind each tick, and the viewer's animation controller pauses after inactivity. Nothing in the measurement kept the viewer's loop awake, so on a light cell a 60 FPS reading does not by itself show that frames were being drawn. On the heavy size cells the raw intervals show a rendering phase of long intervals, preceded by lead cap ticks and followed by idle cap ticks (§3.2, Fig. 6), so the rate on those cells averages rendering with idle ticks and is not comparable to the light cells' 60 FPS.

**Measurement window.** Because the rAF deadline is armed before the warm-up (§2.4), slow initial frames shorten the window, and a shortened window can hold too few intervals to characterise the cell. In three groups of Table 1 (gsplats at 64 px, points at 64 px and gsplats at 32 px, all `post_off_hdr`) a shortened repeat reads well above its two siblings, and the median across repeats rejects each of them.

**Variant scope.** Only the latency loop switches render paths. The rAF loop observes the viewer's own loop, which runs its default pipeline for every variant label (§2.3); the three rAF values of a cell are three repeats of one measurement.

**Synchronisation endpoint.** In `post_off_hdr` the default framebuffer is rebound before the `readPixels`, so the read synchronises against the default framebuffer, not the HDR target that was just drawn. The three variants share a synchronisation point but not the framebuffer it reads; $\Delta_{\mathrm{HDR}}$ is reported as an empirical timing difference for that reason, and it is slightly negative on the two $10^7$ cells (§3.4).

#### 3 Results

##### 3.1 Count scaling at fixed reference size

Figure 4 shows the count sweep at the reference nominal size (4 px for points and gsplats, 1.5 px for lines) under the default HDR pipeline (`post_on`) at 1280×720. At $N = 10^5$ the synchronised render-and-readback call takes 1.21 ms for lines, 1.23 ms for points and 1.72 ms for gsplats; the 16.7 ms budget is 9.7 to 13.9 times the call across the row. At $N = 10^6$ the call is still inside the budget: 3.31 ms (lines), 4.98 ms (points) and 8.64 ms (gsplats), the budget being 1.9 to 5.0 times the call. At $N = 10^7$ points cost 18.8 ms and gsplats 19.4 ms, 12.3% and 16.3% over the budget. Lines were measured through $10^6$ vertices. All $10^6$ and $10^7$ rows carry their settled drawn-element count, equal to $N$ for points and gsplats and 999,999 segments for the $10^6$ line cell, with nothing dropped.

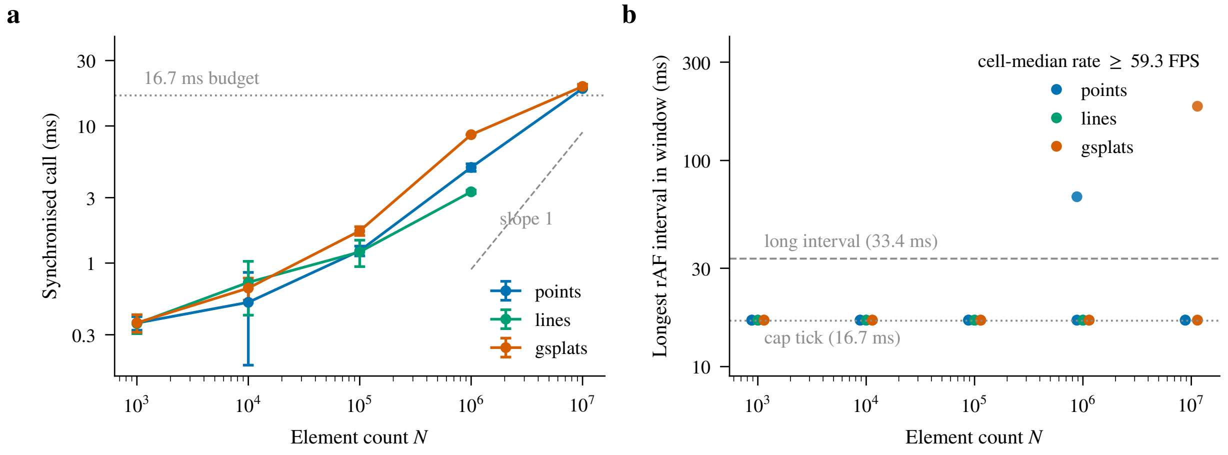


Figure 4: Count scaling at the reference nominal size (4 px for points and gsplats, 1.5 px for lines), 1280×720 viewport, default HDR pipeline (`post_on`). (a) Latency of the synchronised render-and-readback call against $N$ (log-log; medians of three repeats, bars: sample SD across the repeats), with the 16.7 ms budget (dotted) and a slope-1 guide (dashed). (b) The longest interval in each repeat's rAF window (three repeats per cell, log scale). Every window of the count sweep is cap ticks of 16.7 ms (dotted) apart from isolated single long intervals; the dashed line is the 33.4 ms long-interval threshold. The callback-rate medians all lie at the cap (at least 59.3 FPS); the panel shows the intervals themselves because a rate does not say how its window was made up.

Every cell median of the rAF rate in the count sweep lies at or above 59.3 FPS, the $10^7$ cells included. Two readings fit the 60 FPS median at $10^7$. An overrun of 12.3 to 16.3% may not move a callback rate that is batched per begin-frame tick; or the viewer's own loop may have been idle during the window, in which case the callback loop ticked with no frame behind it (§2.6). The rows do not decide between them. One gsplat repeat at $10^7$ reads 56.69 FPS (Table 1); its window is 170 cap ticks and one interval of 183 ms, the same single-outlier shape that 3 of the 27 repeats at $10^6$ show (one interval of 50 to 150 ms among cap ticks), so it is not evidence of a sustained overrun.

The latency curves of Fig. 4a are empirical fits, not shader exponents. Log-log least squares over the measured cells gives slopes of 0.31 (lines, $10^3$–$10^6$), 0.44 (points) and 0.46 (gsplats, $10^3$–$10^7$), with $R^2$ of 0.98, 0.96 and 0.98 and local decade slopes between 0.15 and 0.70. An affine model $a + bN$ fits the same cells with $R^2 = 0.97$, 0.98 and 0.89 (lines, points, gsplats). The two models describe the data about equally well, so a sublinear log slope is also consistent with a fixed per-call overhead plus strictly linear per-element work; the curves do not by themselves identify which GPU stage sets the cost. For lines the segment length also shrinks as $1/\sqrt{N}$ along the count axis (§2.2).

The full per-cell numbers, including the `post_off` and `post_off_hdr` variants, are in the per-cell tables deposited with the analysis. The representative-cell numbers from real Luxar datasets in the main paper's Supplementary Table 1 are consistent with the 1–2 ms synchronised call reported here at $10^5$ elements.

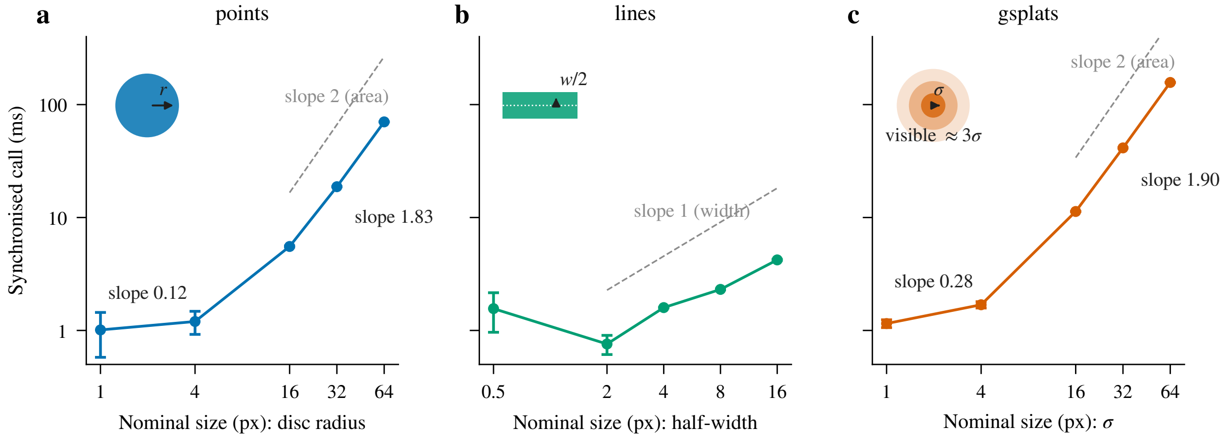


Figure 5: Size scaling at $N = 10^5$ across nominal screen-space sizes $\{1, 4, 16, 32, 64\}$ px (lines: $\{0.5, 2, 4, 8, 16\}$), `post_on`; medians of three repeats, bars: sample SD across the repeats; log-log axes shared across (a) points, (b) lines and (c) gsplats. The inset in each panel draws what the nominal size measures: the disc radius $r$ for points, the stroke half-width $w/2$ for lines and $\sigma$ for gsplats (visible extent about $3\sigma$), so the three x-axes are not the same footprint (Fig. 1). The printed slopes are the log-log endpoint slopes between the cells named in the text; the dashed guides are a cost proportional to covered area (slope 2) and to width (slope 1).

#### 3.2 Size scaling at fixed count

For points and gsplats the small-size end of Fig. 5 is nearly flat (log-log slopes of 0.12 and 0.28 from 1 to 4 px), while from 16 to 64 px the slopes are 1.83 and 1.90, close to the 2 expected if the cost scaled with the area each element covers. A flat curve is what a per-element (vertex or instance) cost produces; an upward-sloping one, a cost that grows with the pixels each element covers. The crossover, between 4 and 16 px nominal, is where a $10^5$-element scene stops being priced per element and starts being priced per pixel. Lines are not flat: their latency rises from 0.76 ms at 2 px to 4.21 ms at 16 px (5.6×), more slowly than points and gsplats over their range, and the 0.5 px cell (1.56 ms) is slower than the 2 px cell, which the data do not explain. The 16 px endpoint carries a primitive caveat (§4.5): it sits at the viewer's automatic capsule-to-quad switch while the 0.5–8 px cells are capsule cells, so the primitive-safe span of the curve is 2–8 px (3.0×, 0.76 to 2.31 ms, for a 4× width increase).

On three cells the cell-median rAF rate left the cap: gsplats at 32 px (44.4 FPS; synchronised call 41.4 ms), gsplats at 64 px (28.4 FPS; 157 ms) and points at 64 px (35.5 FPS; 70.3 ms). Neither number on these cells is a delivered frame rate, and the raw intervals say why. Each window holds a rendering phase, preceded by lead cap ticks and followed by idle cap ticks. In the rendering phase, intervals longer than 33.4 ms (two cap ticks) alternate with cap ticks, almost always a single one. Gsplats at 64 px show 6 long intervals of median 300 ms (maximum 300) over 1.7 s. Points at 64 px show 13 of median 117 ms (maximum 183) over 1.6 s, and gsplats at 32 px 16 of median 67 ms over 1.3 s (medians over the three repeats, long intervals pooled). The rendering phase is preceded by 1, 4 and 17 lead cap ticks and followed by 70, 78 and 86 trailing cap ticks with no long interval among them, the pattern an idle viewer loop produces (§2.6). The callback intervals during the rendering phase were therefore 300, 117 and 67 ms, 1.6 to 1.9 times the synchronised

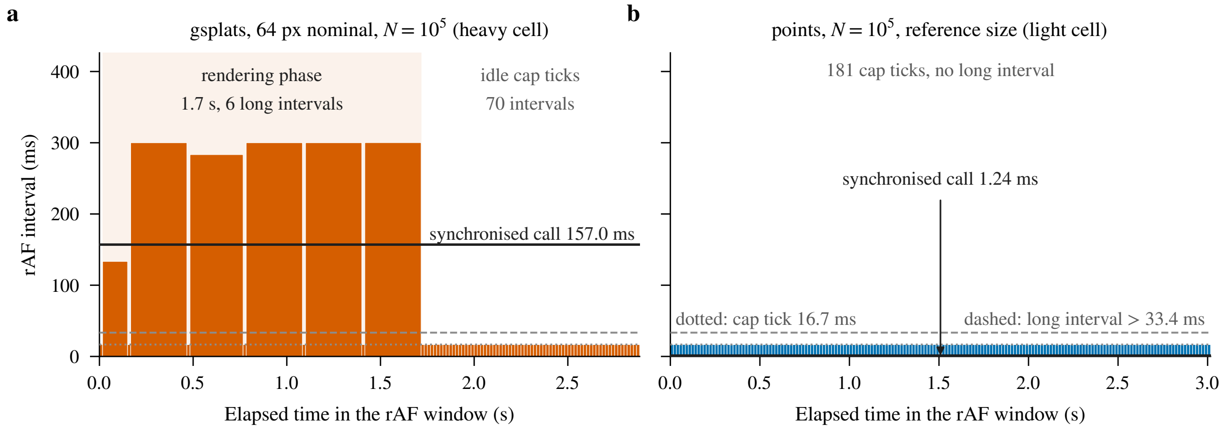


Figure 6: The rAF window as measured, interval by interval, for (a) the gsplats 64 px cell and (b) the points $10^5$ cell at the reference size (`post_on`; the repeat with the median callback rate of its three; each bar spans its interval in time and its height is the interval). The dotted line is the 16.7 ms cap tick, the dashed line the 33.4 ms long-interval threshold and the solid line the synchronised call of the same repeat. The heavy cell's window is a rendering phase of long intervals alternating with single cap ticks, then idle cap ticks with no long interval among them; its callback rate averages rendering with idle ticks and is not a frame rate, and the synchronised call is shorter than the long intervals of the rendering phase. The light cell's window is cap ticks throughout.

call on the same cell; whether each is one long frame or two frames of the call's length the rows do not say, since one long interval plus its cap tick comes close to two synchronised calls on each cell. The reported rate averages that phase with the idle ticks around it. The rate on these cells is not comparable to the 60 FPS of the light cells, and the synchronised call is not a bound on the frame time in either direction. Figure 6 shows one heavy window beside a light one.

##### 3.3 Viewport scaling

At $N = 10^5$ under `post_on` the synchronised call grows with the viewport's pixel count but less than proportionally: the log-log slopes from 720p to 4K are 0.51 for points and 0.63 for gsplats (Fig. 7). Both the scene render and the full-screen post-processing pass scale with the viewport, so the slope does not separate the two; the decomposition of the next section does that at 720p.

##### 3.4 Post-processing overhead, decomposed

Figure 8 splits the total post-processing overhead into its two signed components at three $N$ levels per geometry. The HDR-target term $\Delta_{\mathrm{HDR}}$ grows with scene coverage: a few hundredths of a millisecond on the sparse cells, +1.37 ms at $10^6$ gsplats, and +15.9 and +25.5 ms on the 64 px points and gsplats of the size sweep (Fig. 8b). At $N = 10^7$ it is −0.19 ms (points) and −0.16 ms (gsplats), within the run-to-run noise of two independent medians. The fused-pass term $\Delta_{\mathrm{post}}$ is 0.05–0.60 ms on the count cells of Fig. 8a and −0.10 to +1.00 ms across every paired cell of both sweeps, with no dependence on scene content; bloom, the one content-dependent stage, is off by default and was off here.

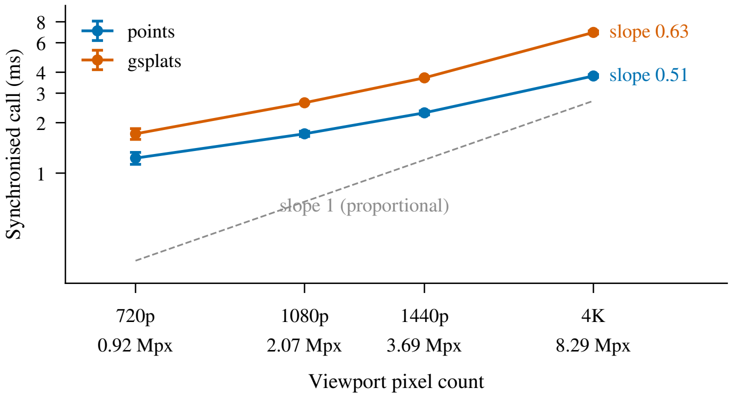


Figure 7: Latency against viewport pixel count, $N = 10^5$, `post_on`; medians of three repeats, bars: sample SD across the repeats. The ticks name the viewports; the dashed guide is a cost proportional to pixel count (slope 1) and the printed slopes are the log-log endpoint slopes from 720p to 4K.

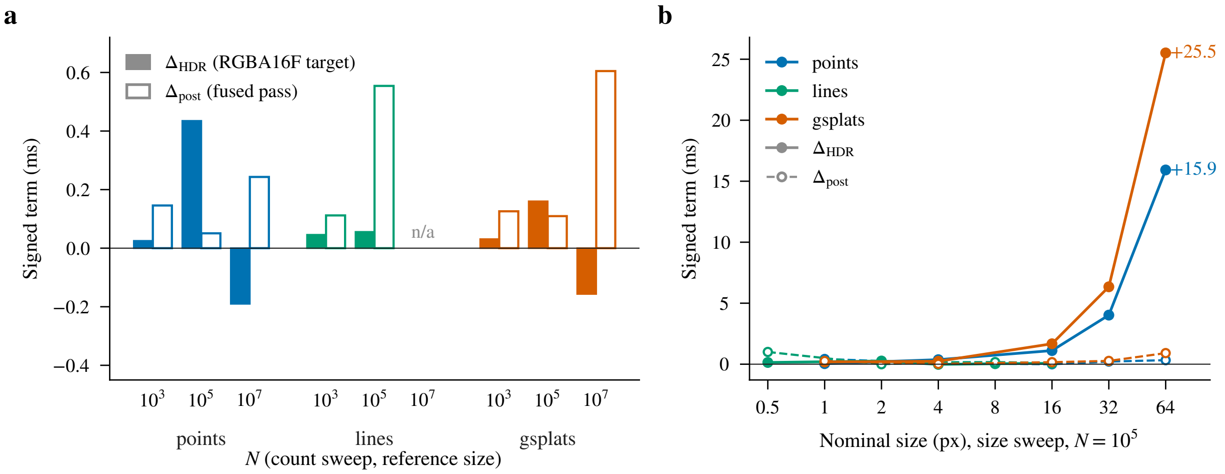


Figure 8: The post-processing overhead (`post_on` − `post_off`) split into its two signed terms: filled, the HDR-target term $\Delta_{HDR}$; hollow, the fused-pass term $\Delta_{post}$ (Fig. 2); colour names the geometry. (a) Count sweep at the reference size, $N = 10^3$, $10^5$ and $10^7$ per geometry (1280×720, medians of three repeats); lines have no $10^7$ cell. (b) Size sweep at $N = 10^5$ against nominal size, from the same cell medians: the HDR term grows with the covered pixels to +15.9 and +25.5 ms on the 64 px points and gsplats, while the fused-pass term (dashed, hollow markers) stays within about a millisecond on every cell.

### 4 Interpretation

#### 4.1 What the two numbers mean

The impulse to report an uncapped frame rate ("more than 2,000 renders per second") misreads the instrument. The browser presents at most one frame per tick of the begin-frame cadence, and the tested runtime's callbacks were capped at about 60 Hz, so the rAF rate saturates at 60 and the count-sweep windows are cap ticks (Fig. 4b). It answers one question, whether the cap was kept, and on the count sweep it was (§3.1). The synchronised call has no cap and scales with the scene (Fig. 4a); at $10^7$ elements it exceeds the budget, the regime in which dropped frames become possible, while the cell-median rate stays at the cap under either reading of §3.1.

Once frames overrun the cadence the two numbers part ways. On the heavy size cells the callback intervals of the rendering phase were 1.6 to 1.9 times the synchronised call, whether as one long frame or two frames of the call's length the rows do not say, and the rAF rate averaged that phase with idle ticks (§3.2), so neither was a delivered frame rate there. The call remains the number to quote for scaling, because it is defined the same way on every cell. The rate is the number to quote for whether the cap was kept, and only on cells where the viewer's loop was demonstrably drawing.

#### 4.2 Cost grows with covered area above about 16 px nominal size

Across both points and gsplats, Fig. 5 shows a close to quadratic increase in latency from 16 to 64 px at fixed $N = 10^5$, the signature of a per-pixel (fragment-fill) cost: each element's cost grows with the number of pixels it covers. Below 4 px the cost varies little with size, as a per-element cost that is paid regardless of how many fragments fall inside would. These are the natural readings of the curves; the sweep did not time GPU stages separately, so they are hypotheses about the bottleneck, not measurements of it.

#### 4.3 Post-processing splits into two terms

The decomposition of §3.4 shows that the overhead is not a single constant offset. The HDR-target term tracks coverage and reaches tens of milliseconds on the 64 px cells, while the fused pass costs at most a millisecond at 720p on every cell measured, independent of content with bloom off. On this hardware the post-processing pass is therefore never the limiting factor at 720p. When a scene is over budget, the cost lives in the scene render itself, and above all in the pixels the elements cover.

#### 4.4 Adaptive DPR

The production-default controller of §2.4 reacts to the rAF rate. On the $10^7$ count cells the rate reads 60 FPS, so it would not engage; on the heavy size cells (28–44 FPS) it would. What rate it reaches there was not measured, since every sweep ran with it pinned, and a quartering of the fill work at its floor would still leave the 64 px gsplat cell near $157/4 \approx 39$ ms per synchronised call if its cost scaled with pixel count. The numbers reported here are the raw cost at the documented pixel count; user-facing performance with the controller on depends on the scale it settles at.

##### 4.5 Geometry-type comparison

At $N = 10^5$ and $10^6$ the synchronised call orders lines < points < gsplats: 1.21, 1.23 and 1.72 ms at $10^5$ (lines and points within 2%), 3.31, 4.98 and 8.64 ms at $10^6$, with gsplats at 1.40–1.73× points. The ordering does not hold at the small counts. At $10^3$ all three are within 1.5% of each other (0.36–0.37 ms); at $10^4$ points (0.52 ms) are cheaper than gsplats (0.66 ms) and lines (0.72 ms), and the gsplat-to-point ratio is 1.00 and 1.27 there; at $10^7$ the two converge again (18.8 against 19.4 ms).

Points and gsplats share the same instanced-billboard pathway: each element is one screen-facing quad drawn by GPU instancing from a shared vertex template. The gsplat shader does more work in both stages (its vertex stage projects the 3D covariance to the screen and its fragment stage evaluates the resulting anisotropic Gaussian, where the point shader evaluates a soft disc), which is a plausible reading of the $10^5$–$10^6$ ratio, not one the timing alone establishes. Lines are cheapest at $10^5$–$10^6$ and their size curve rises more slowly than the other two (5.6× from 2 to 16 px against 4.6× and 6.7× for points and gsplats from 4 to 16 px), as a fragment count that grows linearly with width, and not with the square of a radius, would give.

Two caveats apply to the line numbers. The viewer draws a line segment with one of two primitives, the *capsule* (rounded per-segment geometry) or the *screen-space quad* (a flat rectangle of the requested on-screen width), and its automatic policy selects between them per scene from an effective segment load. The load is the authored segment count times a width factor. The factor starts from the authored `max_width` divided by the node's bounding-box diagonal and multiplied by 1024, a nominal viewport size in pixels, which the viewer's source describes as an order-of-magnitude proxy for the width in pixels at which the node opens (the line shaders draw about four times that, a bias that keeps the capsule longer). That opening width is divided by 1.5 px, the minimum width at which the viewer renders a line, and the factor is the larger of 1 and the quotient. The capsule is used below $2.000 \times 10^6$ effective segments and the quad at and above it. The primitive in force during the run is not recorded: the harness does not pin `?linePrimitive=`. Evaluating the rule on the sweep's cells (Table 3), the count cells at $10^3$–$10^5$ and the size cells at 0.5–8 px resolve to the capsule for any polyline extent the generator can produce. The 16 px cell ($2.011 \times 10^6$ effective segments for the generated cell on disk, $1.988 \times 10^6$ regenerated) and the $10^6$ count cell ($1.682 \times 10^6$ regenerated; $1.58$–$2.74 \times 10^6$ over the possible extents) sit at the switch, so the top endpoint of the line size curve and the top of the line count curve may have been drawn with the quad.

The two primitives differ in cost by a GPU-dependent factor, and the factors we can cite are not measurements of this sweep. The viewer's line-material documentation (`rendering/materials/line/README.md`) reports its own benchmark: the capsule at 1.04–1.11× the quad on an Apple GPU and, on a discrete NVIDIA GPU, about 1.5× on thin lines and 3.16–3.38× on wide lines. Neither GPU is the RTX 3070, this sweep timed no capsule-versus-quad pair, and it does not record which primitive drew any cell. If the 16 px cell was drawn with the quad, the

Table 3: Effective segment load of every line cell under the viewer's automatic line-primitive rule (§4.5): capsule below $2.000 \times 10^6$, screen-space quad at and above. On disk: from the attributes of the generated cells present on disk (no $10^6$ cell present); regenerated: a polyline regenerated with the cell's name-derived seed; any extent: the range over every bounding-box diagonal the generator can produce (0.9 to $0.9\sqrt{3}$ world units). The extent of the cells the run loaded is not recorded, so a cell at the switch cannot be resolved.

| Cell | Nominal px | Segments | Effective load ($\times 10^6$) | | | Auto primitive |
|---|---|---|---|---|---|---|
| | | | on disk | regenerated | any extent | |
| `countA_lines_n1000` | 1.5 | 999 | 0.002 | 0.002 | 0.002–0.003 | capsule |
| `countA_lines_n10000` | 1.5 | 9,999 | 0.020 | 0.021 | 0.016–0.027 | capsule |
| `countA_lines_n100000` | 1.5 | 99,999 | 0.187 | 0.220 | 0.158–0.274 | capsule |
| `countA_lines_n1000000` | 1.5 | 999,999 | – | 1.682 | 1.579–2.735 | at threshold |
| `sizeB_lines_px05` | 0.5 | 99,999 | 0.100 | 0.100 | 0.100–0.100 | capsule |
| `sizeB_lines_px2` | 2 | 99,999 | 0.214 | 0.235 | 0.211–0.365 | capsule |
| `sizeB_lines_px4` | 4 | 99,999 | 0.454 | 0.436 | 0.421–0.729 | capsule |
| `sizeB_lines_px8` | 8 | 99,999 | 0.976 | 1.224 | 0.842–1.459 | capsule |
| `sizeB_lines_px16` | 16 | 99,999 | 2.011 | 1.988 | 1.684–2.917 | at threshold |

5.6× rise understates the capsule's own width scaling, and the linear-in-width reading then rests on the 2–8 px span (3.0× for a 4× width increase), where every cell is a capsule cell; otherwise the line figures are capsule measurements.

### 5 Conclusion

For a user sizing a scene, the two sweeps give a rule of thumb that needs no rerun of the benchmark. Element count sets the cost through a sublinear empirical curve up to $10^6$ elements of any of the three geometries, where the viewer keeps every refresh at the reference size. A scene an order of magnitude larger crosses the budget even with small elements, and one whose elements exceed a nominal 16 px crosses it far earlier, because the cost then grows with the pixels each element covers. Element size, not count, is the knob to reach for first when a scene is slow.

For anyone quoting a rendering-cost figure, the synchronised call is the number to use, and it should be named as what it is: one render plus one readback, defined identically on every cell, which scales with the scene and predicts no frame rate. A rate in "renders per second" implies an interactive frame rate the user cannot observe, and the rAF rate says only whether the cadence was kept, and only where the viewer's loop was drawing.

When a scene is over budget, the post-processing pass is not where the time is. The fused pass costs about a millisecond at most at 720p, and the HDR target's cost is itself a coverage cost. The time is recovered by reducing the covered pixels: smaller elements, a lower render scale (which the adaptive DPR controller applies automatically in production) or fewer elements.
