## Supplementary material for "Luxar: Gaussian splatting for microscopy and scalable interactive web visualisation of multidimensional scientific data": supp_doc_07_compression_comparison

### Compression Comparison: Gaussian Splats vs. Voxel Codecs

Supplementary Document 7 – Luxar

**Abstract**

We ask how Gaussian-splat compression of microscopy volumes compares with a modern voxel codec. We benchmark Gaussian splats against H.265 video coding (`libx265`, Z planes as frames) and against bit-depth quantisation with `zstd` entropy coding on 17 microscopy volumes from four modalities. Fidelity is scored by blind-spot cross-validation applied identically to every method: a random 5% of voxels is hidden before encoding and each reconstruction is scored on those voxels against their original values, so a method that reproduces noise is penalised. Rate is the size of the stored representation in bits per voxel. Read at matched rate under the envelope rule, which credits H.265 with its best result at any rate up to the splat operating rate or its interpolated value at that rate, Gaussian splats recover more signal on 15 of 17 volumes and tie on 2 (median margin +0.98 dB). Read at exactly that rate the margin is +0.50 to +7.27 dB on 16 of 16 volumes. Each method's own held-out peak is higher for Gaussian splats on 12 of 17 by more than the replicate band. The advantage depends on rate: at the mostly lower rate of the H.265 peak the codec stays ahead on 9 of 12 comparable volumes, so the two held-out rate-distortion curves cross. Gaussian splats also reach rates below the lowest rate of the quantisation and lossless baselines, and they are rendered directly from their decoded parameters without reconstructing a voxel grid.

#### 1 Introduction

The main paper reports the compression ratio of every fitted splat set: the bytes of the source volume over the bytes of the store. A ratio alone says nothing about whether the representation is a good use of those bytes. The natural competitor for a Z-stack is a video codec. A stack is a sequence of correlated planes, and H.265 [6] is the most widely deployed codec built to exploit exactly that correlation, with block-based prediction between and within frames followed by transform coding. Two simpler voxel baselines bracket it: bit-depth quantisation followed by `zstd` [4] entropy coding is the plainest lossy voxel scheme, and lossless blosc [1] is the floor of the source data as stored.

Two words need fixing before any curve is drawn. *Rate* is the size of the stored representation in *bits per voxel (bpv)*: the MP4 container for H.265, the `.gsplats.zarr` store for the splats, the compressed bytes for the quantiser. *Fidelity* is held-out prediction *PSNR* under *blind-spot cross-validation* [3], applied identically to every method. Fidelity to the noisy measurement would be the wrong score: a codec that reproduces the noise exactly is perfect against the measurement while recovering nothing of the signal. Hiding a random 5% of voxels before encoding and scoring each reconstruction on those voxels rewards prediction of the signal instead.

The two methods reach their best fidelity at different rates, so one number cannot settle the comparison. We use four readings of the same curves. (i) *Peak to peak*: each method at its own best held-out PSNR, whatever the rate. (ii) *Exact rate*: H.265 interpolated at the bit rate of the splat operating point. (iii) The *envelope* reading: H.265 credited with the better of its best measured result at any rate up to that budget and its interpolated value at that rate. (iv) *At the H.265 peak rate*: the splats interpolated at the codec's own best rate. Sec. 2.6 defines the four precisely. The envelope reading is the compression claim; the exact-rate margin is its upper bound, peak to peak is a model-selection statement, and the reading at the H.265 peak rate is the symmetric check.

The document is organised as follows. Sec. 2 gives the protocol, the two encoders, the datasets and the four readings. Sec. 3 reports the peak comparison, the matched-rate comparison, the very-low-rate regime and the compression ratio at the cross-validation optimum. Sec. 4 interprets the results and Sec. 5 collects them.

#### Terms used in this document

**blind-spot cross-validation** A model-selection protocol in which a deterministic random 5% of voxels is hidden from the fit and the reconstruction is scored on those voxels afterwards, so that capacity spent on memorising noise shows up as a loss instead of a gain.

**held-out voxels** The masked voxels of a blind-spot split, whose original values the optimiser never sees and against which the held-out PSNR of each candidate splat count is scored.

**donut-median fill** The replacement of each masked voxel by the median of its $3^D - 1$ neighbours (the donut, 26 voxels in 3D); the shipped calibration draws the median over the unmasked neighbours only, and a document whose sweeps used all 26 neighbours, masked ones included, states that variant where it applies.

**PSNR** The peak signal-to-noise ratio, $10\log_{10}(\text{peak}^2/\text{MSE})$ in decibels, where peak is the reference range of the values compared; unless a document states another range, the supplements take peak 1 on volumes normalised to $[0, 1]$, so that $\text{PSNR} = -10\log_{10}\text{MSE}$.

***K****** The splat count recommended by a blind-spot sweep: the interior maximum of the held-out PSNR curve when there is a clear peak, the smallest count within 0.3 dB of the maximum on a plateau, and the largest count tested when the curve is still rising there (signal-limited). The shared operating-point table of these documents takes the knee instead, the smallest count within 0.3 dB of that maximum, which is the largest count itself on the two signal-limited benchmark volumes.

**plateau curve** A held-out PSNR curve of a splat-count sweep with a flat top and no clear interior peak, for which the smallest splat count within 0.3 dB of the maximum is reported as the operating point; the word is used in this sense only, not for a plateau of a training loss or a learning-rate schedule.

**signal-limited curve** A held-out PSNR curve whose maximum is the largest splat count tested and which is still climbing there (a total rise of at least 0.3 dB across the sweep, a trailing mean rise of at least 0.1 dB per step and a final step of at least 0.05 dB), so the peak lies beyond the sampled range.

**bits per voxel (bpv)** The compressed size of a volume divided by its voxel count, $8 \times$ store bytes/voxels, a measure of rate that does not depend on the source bit depth.

**CRF** The constant rate factor, the quality-targeting rate-control mode of the x264 and x265 encoders on a 0-51 scale, where a lower value means higher quality and a larger file.

**arm** One experimental condition of a comparison (for instance the blind-spot arm and the unmasked arm of a codec study), run with every other setting held equal.

**MAD** The median absolute deviation, a robust scale estimate that, divided by $0.6745 = 1/\Phi^{-1}(0.75)$, is a consistent estimator of the standard deviation of Gaussian noise.

**iSIM** Instant structured illumination microscopy, a super-resolution fluorescence technique that performs the structured-illumination reconstruction optically in a single exposure, giving roughly a twofold resolution gain at camera frame rates.

**relocation** The fixed-pool alternative to adding and removing splats during optimisation: the splat count is fixed at seeding, and the least informative splats are periodically moved to regions of high reconstruction residual, so the optimiser tensor shapes never change.

### 2 Methods

#### 2.1 Blind-spot cross-validation

We follow the blind-spot (Noise2Self, N2S) protocol of Batson and Royer [3], applied identically to both compression methods (Fig. 1). For each volume $V$ normalised to $[0, 1]$, a deterministic Bernoulli mask $M$ selects 5% of voxels as *held-out voxels* (`seed=42`). The selected voxels are replaced with the median of their $3\times3\times3$ donut neighbourhood (26-connected, centre excluded), the *donut-median fill*. The compression method sees only the masked, donut-filled volume $\tilde{V}$. Every row of this document, codec and splat alike, uses the all-donor fill of Supplementary Document 2's canonical sweeps, the median over all 26 donors with held-out donors included; the deposited runner implements the unmasked-donor variant, which on the 4 volumes Supplementary Document 2 re-swept with it moved held-out PSNR by at most 0.25 dB. We evaluate the held-out PSNR

$$\mathrm{PSNR_{ho}} = -10\log_{10}\mathrm{MSE_{ho}}, \qquad \mathrm{MSE_{ho}} = \frac{1}{|M|}\sum_{i\in M}(V_i - \hat{V}_i)^2, \tag{1}$$

where $|M|$ is the number of held-out voxels, $V_i$ the original (pre-replacement) value at held-out position $i \in M$, $\hat{V}$ the reconstruction produced by each method, and the peak is 1 on the normalised volume.

Under the blind-spot assumptions the held-out mean squared error is the clean-signal error plus the noise variance. The assumptions are that the noise is zero-mean and independent across voxels given the signal, and that the reconstruction at a held-out voxel does not depend on that voxel's own value. $\mathrm{PSNR_{ho}}$ then ranks methods by how well they predict the underlying signal, not the noise realisation. It is a held-out *prediction* score: its value is not the clean-signal PSNR. On volumes that were deconvolved or motion-aligned before the benchmark (Tribolium embryo nuclei, Zebrafish embryo nuclei, Neuromast membranes, Neuromast nuclei; marked * in the tables) the residual is spatially correlated, the independence assumption does not hold, and $\mathrm{PSNR_{ho}}$ measures prediction of the processed intensities (see Limitations).

The GSplat bits per voxel printed in this *arm* are those of the blind-spot fit's store (the fit of $\tilde{V}$). They differ from the main paper's Table 1, which measures the unmasked stores, by at most 0.0074 bpv at the same splat count, in the second decimal on three of the 17 volumes (Kidney nuclei, Tribolium embryo nuclei, Mouse heart nuclei; Kidney nuclei at 64K: 1.41 vs. 1.40 bpv).

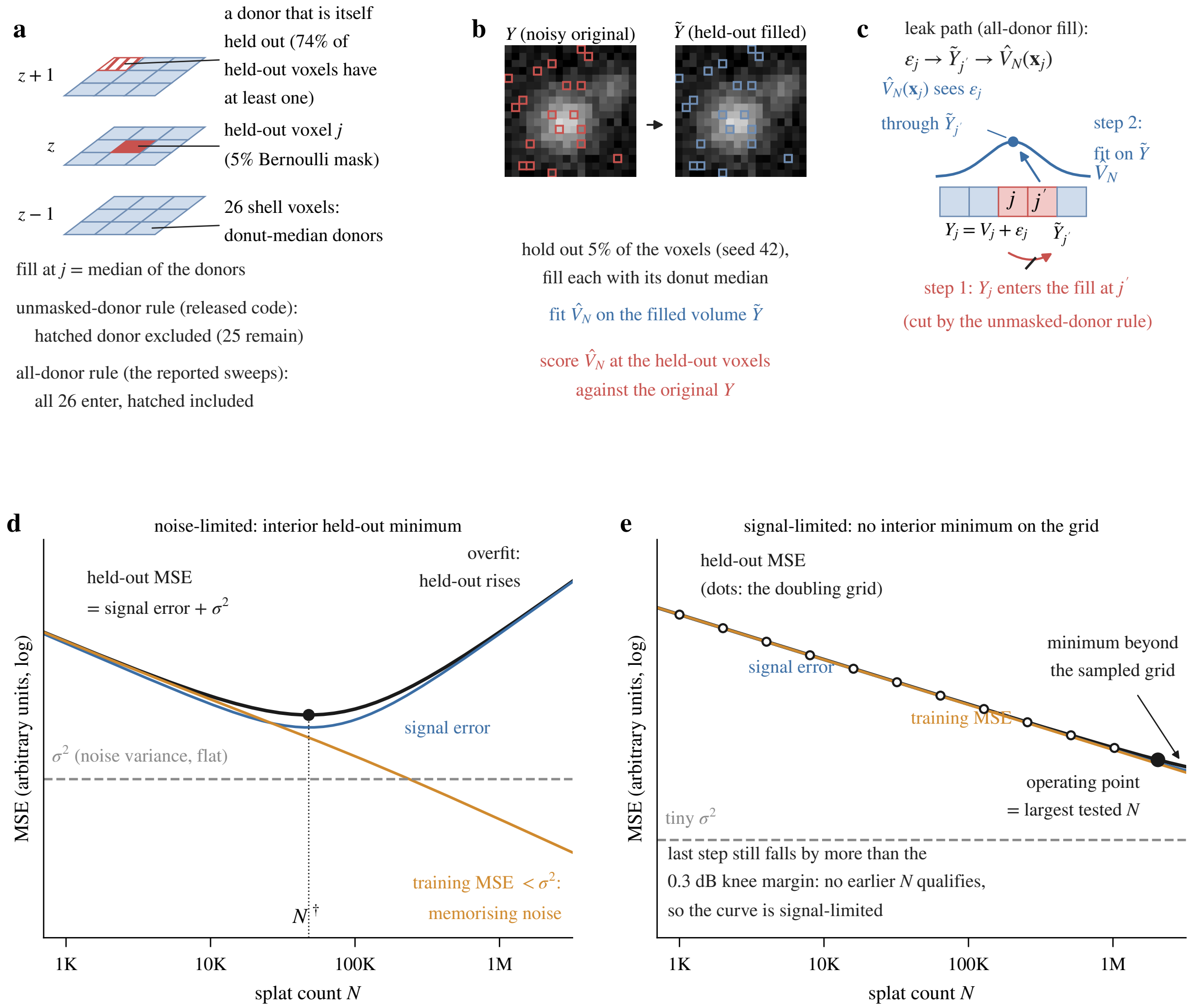


Figure 1: Schematic of the blind-spot protocol that scores every method in this document; shared with Supplementary Document 2, whose fitting sweep it also governs. (a) A held-out voxel $j$ (5% Bernoulli mask) and the 26 shell voxels of its $3 \times 3 \times 3$ donut, whose median replaces it; a donor can itself be held out, and the panel distinguishes the released code's unmasked-donor rule from the all-donor rule of the reported sweeps. (b) The pipeline: a codec or a fit sees only the filled volume $\tilde{V}$ and is scored at the held-out voxels against the original $V$. (c) The neighbour leak under the all-donor fill: a held-out value enters the fill of an adjacent held-out voxel, which the fit sees. (d, e) Idealised held-out error against capacity, no data plotted: the held-out MSE is the signal error plus the flat noise variance, so it has an interior minimum where the noise variance is resolvable (d) and none on the sampled grid where it is tiny (e, the signal-limited light-sheet case). Panels d and e are drawn for the splat count, labelled $N$ in the panel as in Supplementary Document 2 ($K$ in this document); the same decomposition governs a codec's rate knob, which is why the H.265 held-out curves of Fig. 3 decline past their peak.

#### 2.2 H.265 encoding

Each masked Z-stack is treated as a greyscale video whose frames are the Z planes, array axis 0 of the stored volume, in stored order. The physical z spacing is not passed to the encoder. It is coarser than the lateral sampling by more than 5% on 10 of the 17 volumes (within 5% of isotropic on 5, not recorded for 2), so inter-frame prediction treats anisotropic z neighbours as consecutive frames, and no resampling or axis choice was tuned for the codec.

Volumes are linearly mapped from $[0, 1]$ to 16-bit unsigned integers, written as raw frames, and encoded with `ffmpeg` using `libx265` [5], an H.265/HEVC [6] encoder, at `medium` preset and 12-bit internal precision (`gray12le` pixel format). We sweep the constant rate factor (*CRF*) over CRF $\in \{51, 45, 40, 35, 30, 25, 20, 15, 10, 5, 0\}$, where lower CRF means higher quality and larger files. At CRF 0 the encoder runs in x265's lossless mode (`lossless=1`), which is lossless for the 12-bit converted input (the 16-bit source is reduced to 12 bits before encoding), not for the 16-bit source. The encoder driver carries a fallback for `ffmpeg` builds without 12-bit `gray` pixel-format support (8-bit `yuv420p` at CRF $\geq 1$ without the lossless flag), which was not engaged on the runs reported here (main paper, Methods). Compressed file size is the size of the MP4 container. The video is decoded back to a uint16 volume, scaled to $[0, 1]$, and the held-out PSNR is computed against the original.

#### 2.3 GSplat encoding

For each masked volume we use the same fitting pipeline as in the main paper (Methods, "Optimisation"), sweeping the splat count $K$ over $\{1, 2, 4, 8, 16, 32, 64, 128, 256, 512\}$K on every volume (10 counts). The four light-sheet volumes (Tribolium embryo nuclei, Mouse heart nuclei, Zebrafish embryo nuclei, Drosophila embryo nuclei) carry two further counts, 1024K and 2048K, recorded in the same canonical sweep file as their base grid (Supplementary Document 2 documents the fitting protocol and the extension). The rendered volume is computed at full resolution and held-out PSNR is evaluated on the masked positions.

The cross-validation optimum $K^*$ is the count the held-out curve selects: the interior maximum when the curve has a clear peak, and otherwise the knee, the smallest count within 0.3 dB of the maximum. On a *plateau curve* the knee is the onset of the flat top; on a *signal-limited curve* it is the largest count tested whenever no earlier count comes that close to the maximum, as on both such volumes here (Sec. 3.4). It is the rule of the `luxar gsplat cal` command. The *operating point* of a volume is $K^*$ together with the bit rate of its store; the phrase is used in no other sense in this document. Table 1, Tables 2 and 3, $K^*$ and Fig. 3 all read the one merged curve per volume.

The compressed footprint is the measured on-disk size of each fit's `.gsplats.zarr` store as the default writer emits it: `uint16` fixed-point centres, `uint8`/`uint16` log-encoded Cholesky factors, amplitudes on a bounded-linear `uint8`/`uint16` or geometric-log `uint16` grid by dynamic range, Blosc-`zstd` chunks plus array and index metadata, fitting provenance excluded. It is measured on the cached store of every sweep cell (Supplementary Document 12 characterises the encoding). The byte accounting is the one the main paper's compression ratios use: the same writer, the same store measurement, and a numerator that is the source volume at its stored dtype (`uint16` for most volumes, `uint8` for the Mouse heart, `float32` for the deconvolved neuromast channels), never a float32 assumption. The stores differ, however: Tables 2 and 3 rate fits of the masked volume $\tilde{V}$, the main paper's Table 1 and Sec. 3.4 fits of the unmasked volume (Sec. 2.1).[1] The results we report under the "GSplats" label in Supplementary Fig. 7 of the main paper are precisely those generated by this pipeline.

##### 2.4 Datasets and metrics

The N2S sweep covers the main paper's benchmark volumes (Methods, "Datasets"; 17 volumes from 12 datasets, listed with modality, marker and source in the main-text dataset table). Table 1 gives one row per volume with a complete sweep, in the same order (spinning-disk, confocal, light-sheet, *iSIM*). Every volume passes through one loader, the benchmark's volume reader, which normalises it to $[0, 1]$ by an affine map of its intensity range with dataset-specific percentile clipping and, for the neuromast channels, background-floor subtraction (main paper, Methods). "The loader's normalised float32 array" below means that output, the payload every fit and every quantiser run consumes.

We report bits per voxel ($\text{bpv} = 8 \cdot \text{compressed\_bytes}/N_{\text{voxels}}$), held-out PSNR, and the per-dataset noise floor. The floor is the noise standard deviation $\hat{\sigma}$ estimated by an ensemble of robust median-absolute-deviation (*MAD*) estimators (main paper Methods, "Noise floor estimation"), reported as the PSNR ceiling $-20 \log_{10} \hat{\sigma}$.

##### 2.5 Arms and baselines

The *blind-spot arm* comprises every curve of Table 1, Tables 2 and 3 and Fig. 3. Each is an encoding of the masked, donut-filled volume $\tilde{V}$ (every codec is handed the same masked volume), not of the untouched original. The "raw" PSNR plotted alongside the held-out PSNR is that encoding of $\tilde{V}$ scored against the original noisy volume $V$ at all voxels, so it too carries the 5% of filled positions.

The *unmasked arm* (Sec. 3.3) encodes the original volume with no blind-spot mask and adds two voxel baselines. **Quantisation+zstd** is uniform quantisation of the loader's normalised float32 array to $\{4, 6, 8, 10, 12, 16\}$ bits followed by `zstd` level 9 with a 12-byte header; `zstd` (Zstandard) is a general-purpose lossless entropy coder. **Lossless blosc** [1, 2] is a chunked compression layer that reorders the bytes (`shuffle`) or bits (`bitshuffle`) of each element, grouped by the element size `typesize`, before a codec; we run `lz4+shuffle` level 5, `zstd+shuffle` level 5 and `zstd+bitshuffle` level 9.

Lossless blosc is measured on two bases. The *stored-dtype* baseline compresses exactly the benchmark's voxels (the same channel, timepoint and crop, verified by reproducing the loader's array bit for bit from the raw crop) at the dtype the source file stores them in: `uint16` for 14 volumes, `uint8` for the Mouse heart TIFF, `float32` for the two deconvolved neuromast channels. `typesize` equals that itemsize and the values are as stored: no percentile clip, no background-floor

[1] The encoding-agnostic float32 parameter footprint $F_D \cdot K \cdot 4$ bytes ($F_D = 10$ for 3D) is a different quantity. The shipped encoding stores a median of 11.3 B per splat over the 30 fresh-fit cells of Supplementary Document 12 (3.5× below the float32 footprint), and 9.0–12.9 B per splat (median 11.1 B, array and index metadata included) over the 17 stores at the operating points of Sec. 3.4; both are finite-sample summaries, not a large-$K$ limit.

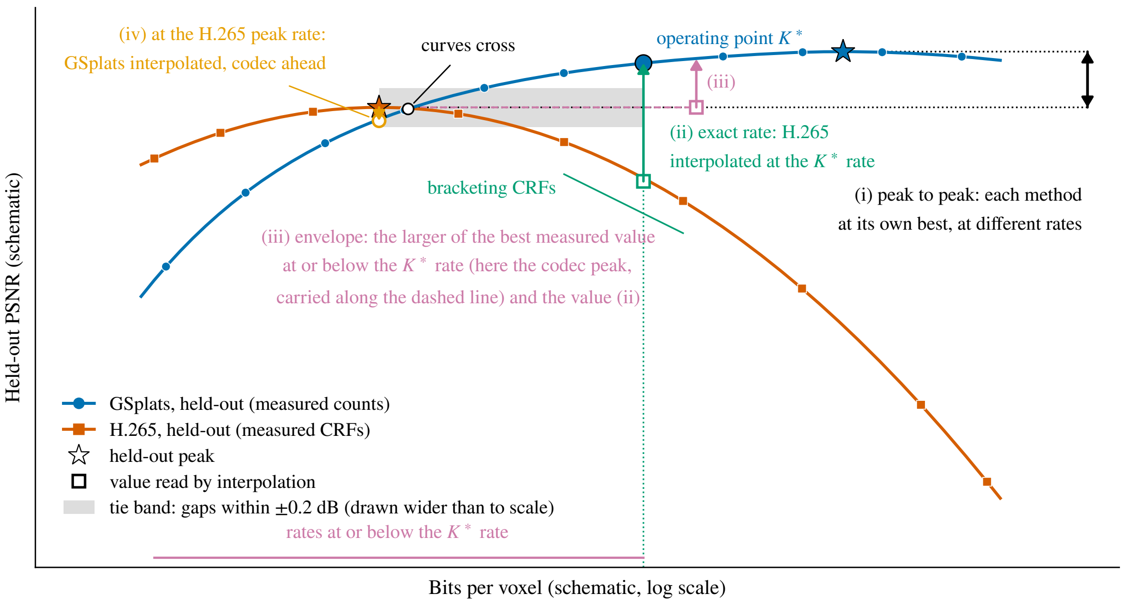


Figure 2: Schematic of the four readings on one idealised pair of held-out curves; no data is plotted. The GSplat curve (blue, circles at the measured counts) rises to a late peak; the H.265 curve (vermilion, squares at the measured CRFs) peaks early and then declines, so the two cross once. (i) Peak to peak: the two stars, reached at different rates. (ii) Exact rate: H.265 interpolated log-linearly between the two CRFs that bracket the operating rate of $K^*$ (open square) and read against the operating point. (iii) Envelope: H.265 credited with the larger of its best measured value at any rate at or below the operating rate (here the earlier peak, carried along the dashed line) and the interpolated value (ii); this gap can never exceed the exact-rate gap. (iv) At the H.265 peak rate: the GSplat curve interpolated at the codec's own peak rate (open circle), where the codec is ahead. The grey strip is the tie band, ±0.2 dB around the credited H.265 value and drawn wider than to scale: a gap inside it is read as a tie.

subtraction, no normalisation, because those steps condition the fit's input and a clip changes voxel values. It is the lossless floor of the source image data, and it is the basis of every "floor" comparison below. The *float32* baseline compresses the loader's normalised float32 array with `typesize` 4, is labelled as such wherever it appears, and is retained as the rate of the payload the quantiser and the fits consume. On either basis "lowest rate" means the smallest rate among the three configurations tested, not a codec-independent bound.

#### 2.6 Four readings and the tie band

The held-out curves of the two methods peak at different rates, so we read them four ways (Fig. 2).

**Peak to peak.** Each method at its own best held-out PSNR (Table 1). The gap is a model-selection statement, not a comparison at equal cost: the two peaks sit at unequal bit rates.

**Exact rate.** H.265 read at the bit rate of the splat operating point (Sec. 2.3). Its held-out PSNR at that rate is interpolated linearly in $\log_2(\text{bpv})$ between the two adjacent measured CRF settings that bracket the rate. We never extrapolate: when the rate falls outside the measured H.265 range the cell is left blank and flagged († below the range, ‡ above it). The interpolation is also *withheld* (¶) when the bracketing segment ends at the lossless CRF 0 and spans more than a factor of 4 in rate, two doublings: a log-linear segment across a lossy-to-lossless regime change that wide carries no information about the curve between its endpoints. The one such volume (Tribolium embryo nuclei) keeps its envelope reading, which uses measured points only, and is excluded from the exact-rate counts.

**Envelope.** A user with a byte budget would not encode H.265 to the splat rate if a smaller codec file scored better. The envelope rule therefore credits H.265 with the larger of two values: its best measured held-out PSNR at any rate at or below the splat operating rate, and its interpolated value at that rate (§ marks the volumes where the interpolated value is the larger). The envelope margin can never exceed the exact-rate one. We call it "the envelope rule" below, and it is the reading we treat as the compression claim.

**At the H.265 peak rate.** The symmetric check: the splat curve interpolated, by the same rule and with the same flags, at the bit rate of the H.265 held-out peak.

**Tie band.** Throughout, a gap within ±0.2 dB is read as a tie, a gap above +0.2 dB as a GSplat win and one below −0.2 dB as an H.265 win. The band is the maximum held-out spread between the two replicate seeds of the fit replicates in Supplementary Document 2 (0.20 dB), which that document gives as the level below which differences between table entries should not be read as dataset effects. A single replicate there differs from the canonical seed, the seed of every splat row here, by up to 0.41 dB. Read at ±0.41 dB instead, the peak, envelope and exact-rate splits below are unchanged, and the split at the H.265 peak rate (wins, ties and losses for GSplats) moves from 2/1/9 to 2/4/6. The band is symmetric and descriptive, not a significance statement: those replicates cover ten volumes (the spinning-disk and confocal volumes except the Fly brain neurons tile), and the codec arm has none. Sign-only counts (any positive gap a win) are given alongside for reference.

**Reproducibility.** The blind-spot results table deposited with the analysis records, per row, the volume, method, knob (CRF, bit depth or splat count), bits per voxel, compressed bytes, raw and held-out PSNR and a renderability flag, true for the splat rows, whose decoded parameters the viewer draws directly, and false for every voxel method, whose output is a voxel grid (Sec. 4); the unmasked arm and the two lossless-blosc baselines are deposited as companion tables. The codec rows carry the all-donor fill (Sec. 2.1); the deposited runner fills with unmasked donors only, so a rerun differs from the deposited rows at the held-out voxels that have a held-out donor. The `ffmpeg/libx265` build, the encoded pixel format and profile of each run and the encoder logs are not recorded, so the codec configuration cannot be verified from those tables alone: the invocation is the one printed in the main paper's Methods, and the fallback statement above rests on that record. No seed replicates were run for the codec arm and no uncertainty is propagated into the gaps.

### 3 Results

#### 3.1 Held-out prediction PSNR

The held-out and raw rate-distortion shapes of both methods (Fig. 3; equivalent to Supplementary Fig. 7 of the main paper) set up everything that follows. For voxel methods including H.265, the raw PSNR generally rises with bit rate, with small reversals at the highest rates because it is scored against the original noisy volume and not against the filled volume the codec encoded. The held-out PSNR rises, peaks, and then *declines*. At high bit rates H.265 starts to encode the masked-position fill values faithfully. Those values are donut medians of the surrounding noisy neighbourhood, so they drift away from the true signal at the masked positions, and the bits spent on them buy no held-out signal. GSplats show the same bias-variance trade-off with a smaller bias term: the global mixture model interpolates across masked regions with a continuous parametric representation instead of a codec's block-wise intra prediction, which extrapolates each block from its already coded neighbours and transform-codes the residual.

Table 1 reports the peak held-out PSNR of both methods on each of the 17 volumes, with the bit rate and knob value at which the peak is reached. Under the replicate band, GSplats lead on 12 of the 17 volumes, the two tie on 5 (OpenCell-MAP4 nuclei, OpenCell-MAP4 microtubules, OpenCell-LMNB1 nuclei, C. elegans embryo nuclei, Tribolium embryo nuclei) and H.265 leads on 0; by sign alone the split is 16 to 1. The two peaks are reached at different bit rates (the bpv columns), so the gap is peak to peak, not a comparison at equal cost; Sec. 3.2 makes that comparison.

Grouped by modality, the peak gaps read as follows. On the four spinning-disk OpenCell channels GSplats lead by a small margin, +0.14 dB (OpenCell-LMNB1 nuclei) to +0.52 dB (OpenCell-LMNB1 nuclear lamina), three of the four inside the band. On the seven confocal volumes the margin runs from +0.17 dB (C. elegans embryo nuclei; one of the seven inside the band) to +3.10 dB (Kidney actin), largest on the noisy kidney channels and the Cells3D membranes. On the two deconvolved iSIM neuromast channels the margins are the largest of the benchmark, +4.18 and +5.58 dB. On the light-sheet volumes GSplats win on the Mouse heart nuclei (+3.44 dB, at the 2048K end of its still-rising held-out curve), the Zebrafish embryo nuclei (+1.74 dB) and the *Drosophila* embryo nuclei (+1.24 dB), while on the *Tribolium* embryo nuclei the two tie (−0.09 dB). There H.265 peaks at 55.64 dB and the splat curve, still rising at the 2048K end of its extended sweep, reaches 55.55 dB at 4.5× the bit rate (1.47 vs. 0.33 bpv). The *Tribolium* outcome is specific to that volume and does not generalise to the other deconvolved light-sheet data.

We have no measured explanation for the *Tribolium* tie. Smoothness after deconvolution is our hypothesis, and one statistic argues against it. The held-out PSNR of the donut-median fill itself ("Fill" in Table 1: the 16-bit quantisation row, which stores the filled volume to within a $2^{-16}$ step) measures how predictable a volume's held-out voxels are from their neighbours, independent of either codec; read this way the fill is a *mask-fill predictor*. It is 53.83 dB on the *Tribolium* embryo nuclei and 56.34 dB on the Neuromast nuclei, so the second volume's held-out voxels are at least as predictable. H.265 clears its fill by +1.81 dB on the first and by +1.68 dB on the

Table 1: Peak held-out PSNR per volume across the 17 benchmark volumes with a complete sweep. "GSplats peak" and "H.265 peak" give the held-out PSNR at each method's own best point, with the splat count or CRF and the bit rate in brackets. Gap = GSplats − H.265; a gap within ±0.2 dB is read as a tie (Sec. 2.6). "NF" is the noise-floor ceiling $-20\log_{10}\hat{\sigma}$ in dB, with $\hat{\sigma}$ the median of the three MAD noise estimators (Laplacian, Haar, background) of Sec. 2.4; "n.e." (not estimable) marks the four volumes on which that ensemble estimate is 0 (only the median is deposited, so at least two of the three estimators returned 0 there). "Fill" is the held-out PSNR of the donut-median fill itself, read from the 16-bit quantisation row (Sec. 3.1). * marks volumes deconvolved before the benchmark (Sec. 2.1). GSplat rows are from the canonical sweep of Supplementary Document 2; H.265 rows are codec measurements independent of the fitting code.

| Dataset | GSplats peak<br>dB @ splats (bpv) | H.265 peak<br>dB @ CRF (bpv) | Gap (dB) | NF (dB) | Fill (dB) |
|---|---|---|---|---|---|
| OpenCell-MAP4 nuclei | 28.15 @ 16K (0.0877) | 28.00 @ 0 (11.2) | +0.15 | 32.61 | 28.00 |
| OpenCell-MAP4 microtubules | 24.89 @ 64K (0.301) | 24.70 @ 15 (4.66) | +0.19 | 30.45 | 24.69 |
| OpenCell-LMNB1 nuclei | 25.45 @ 32K (0.0735) | 25.31 @ 10 (5.68) | +0.14 | 29.96 | 25.31 |
| OpenCell-LMNB1 nuclear lamina | 34.00 @ 64K (0.136) | 33.48 @ 15 (2.78) | +0.52 | 42.56 | 33.47 |
| Kidney nuclei | 33.91 @ 64K (1.41) | 31.33 @ 35 (0.367) | +2.58 | 41.34 | 29.27 |
| Kidney actin | 30.65 @ 128K (2.65) | 27.55 @ 40 (0.322) | +3.10 | 43.47 | 25.67 |
| Cells3D nuclei | 35.45 @ 32K (0.773) | 35.02 @ 15 (3.51) | +0.43 | 37.57 | 34.99 |
| Cells3D membranes | 46.73 @ 32K (0.772) | 44.02 @ 25 (0.264) | +2.71 | 53.54 | 42.44 |
| Blastocyst nuclear lamina | 37.58 @ 32K (0.161) | 36.43 @ 35 (0.106) | +1.15 | 54.42 | 36.37 |
| C. elegans embryo nuclei | 39.23 @ 8K (0.0758) | 39.06 @ 40 (0.00750) | +0.17 | 40.73 | 39.02 |
| Fly brain neurons | 44.51 @ 512K (0.372) | 43.64 @ 35 (0.0121) | +0.87 | 52.08 | 43.45 |
| Tribolium embryo nuclei* | 55.55 @ 2048K (1.47) | 55.64 @ 15 (0.330) | −0.09 | 61.03 | 53.83 |
| Mouse heart nuclei | 45.30 @ 2048K (1.27) | 41.86 @ 30 (0.180) | +3.44 | n.e. | 39.21 |
| Zebrafish embryo nuclei* | 37.99 @ 1024K (0.483) | 36.25 @ 40 (0.0505) | +1.74 | n.e. | 35.45 |
| Drosophila embryo nuclei | 34.27 @ 2048K (1.67) | 33.03 @ 40 (0.0813) | +1.24 | 51.14 | 31.64 |
| Neuromast membranes* | 50.85 @ 128K (0.224) | 46.67 @ 35 (0.0304) | +4.18 | n.e. | 43.41 |
| Neuromast nuclei* | 63.60 @ 64K (0.0940) | 58.02 @ 25 (0.0151) | +5.58 | n.e. | 56.34 |

second, yet the peaks tie on the first and GSplats lead by +5.58 dB on the second. What differs is the splat side, +1.72 against +7.26 dB above the fill, so the fill statistic does not separate the two volumes and the tie is not explained by smoothness alone.

The other light-sheet volumes behave differently. On the Zebrafish embryo nuclei H.265 saturates at 36.25 dB and GSplats win; on the Mouse heart nuclei H.265 saturates at 41.86 dB while the splats reach 45.30 dB. Both the *Tribolium* and the Zebrafish volumes are multi-view-deconvolved, which introduces spatial correlations in the residual that partly violate the pixel-independence assumption of the blind-spot framework, as are the two iSIM neuromast channels (Richardson-Lucy deconvolved and motion-aligned). The deconvolution caveat applies to all four, the codec outcome nonetheless diverges, and the fill statistic above does not predict which way. A further caveat concerns cross-volume comparability. The light-sheet crops are not matched in style: the Zebrafish volume is a content-dense sub-region selected for signal, whereas the *Tribolium* and heart volumes are plain spatial crops that retain smooth background, which H.265 predicts near-losslessly. Part of the cross-volume difference therefore reflects content density, not intrinsic codec behaviour, and the content-dense Zebrafish volume arguably tests the codecs where reconstruction matters most. We therefore report the light-sheet outcomes individually and do not draw a general "H.265 wins on deconvolved light-sheet" conclusion.

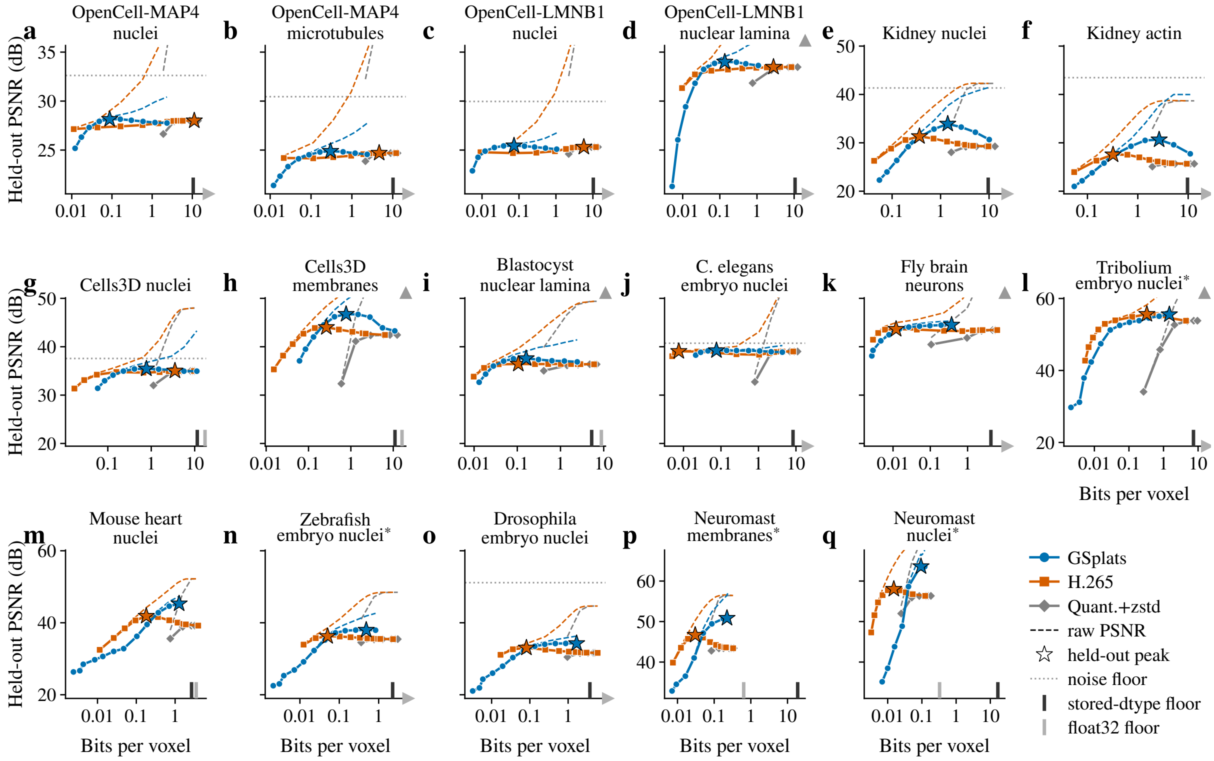


Figure 3: Held-out PSNR against bits per voxel for every volume of Table 1, one panel per volume in the table's order (spinning-disk a to d, confocal e to k, light-sheet l to o, iSIM p and q; the figure corresponds to Supplementary Fig. 7 of the main paper). Panels of one modality share a y range, so the height of a curve can be compared across the modality; the first panel of each modality and of each row carries the y tick labels. Solid lines with markers: held-out prediction PSNR of GSplats (blue circles), H.265 (vermilion squares) and quantisation+zstd under the same blind-spot protocol (grey diamonds). Thin dashed lines in the same colours: raw PSNR against the noisy measurement, which rises past the top of the panel at the highest rates because it rewards reproducing the noise. Stars: each method's peak held-out PSNR, the peak rows of Table 1; on a curve still rising at its largest count the star sits at the endpoint. Dotted horizontal line: the noise-floor ceiling where it is estimable and lies within the shared y range; a grey triangle at the top right marks a ceiling above that range, and the four "n.e." volumes carry neither. Ticks rising from the x axis: the lowest bit rate the three lossless blosc configurations reach on that volume, on the source at its stored dtype (dark) and on the normalised float32 array (light; Sec. 2.5); a light triangle at the right end of the axis marks a float32 floor beyond the plotted range.

#### 3.2 Matched-rate comparison

The peak-to-peak gaps of Table 1 compare points that spend very different numbers of bits. On 12 of the 17 volumes the splat operating point sits 1.5–10× *above* the H.265 peak in bits per voxel (median 4.4×; Kidney nuclei: 1.41 vs. 0.37 bpv), because the H.265 held-out curve peaks early and then declines while the splat curve keeps rising. On the other 5, the noisy spinning-disk and confocal volumes whose H.265 held-out peaks at CRF $\leq 15$, it sits 4.5–127× *below* it. The peaks themselves are further apart: the splat held-out argmax spends 1.5–31× more bits per voxel than the H.265 peak on 12 volumes and 4.5–127× fewer on the other 5. A higher peak is therefore not by itself evidence of better compression.

Table 2: Matched-rate comparison at the GSplat operating point. Exact-rate reading: the operating point ($K^*$ splats, bpv, held-out dB) against H.265 interpolated at the *same* bits per voxel; "CRFs" lists the two measured settings that bracket that rate, the segment the interpolation runs along; $\Delta$ = GSplats − H.265, positive meaning GSplats recover more signal for the same bits. Envelope reading: H.265 credited under the envelope rule of Sec. 2.6 (the credited point's bpv in brackets; § where the interpolated value is the larger) and $\Delta$ against it. †: rate below the measured H.265 range; ‡: above it; ¶: interpolation withheld (lossless bracket wider than 4×). *: deconvolved before the benchmark.

| | Exact-rate reading | | | | | Envelope reading | |
|---|---|---|---|---|---|---|---|
| Dataset | splats (bpv) | GSplats dB | H.265 dB | CRFs | $\Delta$ dB | H.265 (bpv) dB | $\Delta$ dB |
| OpenCell-MAP4 nuclei | 16K (0.0877) | 28.15 | 27.37 | 45–40 | +0.78 | 27.37 (0.0877)§ | +0.78 |
| OpenCell-MAP4 microtubules | 64K (0.301) | 24.89 | 24.26 | 45–40 | +0.63 | 24.26 (0.301)§ | +0.63 |
| OpenCell-LMNB1 nuclei | 32K (0.0735) | 25.45 | 24.69 | 45–40 | +0.76 | 24.78 (0.00910) | +0.67 |
| OpenCell-LMNB1 nuclear lamina | 64K (0.136) | 34.00 | 33.20 | 40–35 | +0.80 | 33.20 (0.136)§ | +0.80 |
| Kidney nuclei | 64K (1.41) | 33.91 | 30.23 | 25–20 | +3.68 | 31.33 (0.367) | +2.58 |
| Kidney actin | 128K (2.65) | 30.65 | 26.09 | 20–15 | +4.56 | 27.55 (0.322) | +3.10 |
| Cells3D nuclei | 32K (0.773) | 35.45 | 34.67 | 30–25 | +0.78 | 34.78 (0.157) | +0.67 |
| Cells3D membranes | 32K (0.772) | 46.73 | 43.31 | 20–15 | +3.42 | 44.02 (0.264) | +2.71 |
| Blastocyst nuclear lamina | 32K (0.161) | 37.58 | 36.42 | 35–30 | +1.16 | 36.43 (0.106) | +1.15 |
| C. elegans embryo nuclei | 8K (0.0758) | 39.23 | 38.73 | 35–30 | +0.50 | 39.06 (0.00750) | +0.17 |
| Fly brain neurons | 64K (0.0514) | 44.25 | 43.53 | 30–25 | +0.72 | 43.64 (0.0121) | +0.61 |
| Tribolium embryo nuclei* | 2048K (1.47) | 55.55 | –¶ | 5–0 | – | 55.64 (0.330) | −0.09 |
| Mouse heart nuclei | 2048K (1.27) | 45.30 | 39.95 | 15–10 | +5.35 | 41.86 (0.180) | +3.44 |
| Zebrafish embryo nuclei* | 256K (0.141) | 37.80 | 36.08 | 35–30 | +1.72 | 36.25 (0.0505) | +1.55 |
| Drosophila embryo nuclei | 256K (0.245) | 34.01 | 32.53 | 40–35 | +1.48 | 33.03 (0.0813) | +0.98 |
| Neuromast membranes* | 128K (0.224) | 50.85 | 43.58 | 5–0 | +7.27 | 46.67 (0.0304) | +4.18 |
| Neuromast nuclei* | 64K (0.0940) | 63.60 | 56.48 | 5–0 | +7.12 | 58.02 (0.0151) | +5.58 |

Tables 2 and 3 read each method at the rate of the other's reference point, H.265 at the splat operating rate and the splats at the H.265 held-out peak rate, under the rules of Sec. 2.6. The splat operating point is $K^*$, which coincides with the bare argmax of the held-out curve except on 3 plateau volumes (Fly brain neurons, Zebrafish embryo nuclei, Drosophila embryo nuclei), where $K^*$ is the first count within 0.3 dB of the maximum. The H.265 point is its held-out peak, as in Table 1.

Under the envelope rule (Table 2, right block) GSplats recover more signal on 15 of the 17 volumes, the two tie on 2 (C. elegans embryo nuclei, Tribolium embryo nuclei; +0.17 and −0.09 dB) and H.265 leads on 0. By sign alone GSplats are ahead on 16 of 17 (median +0.98 dB, range −0.09 to +5.58 dB). On three volumes (OpenCell-MAP4 nuclei, OpenCell-MAP4 microtubules, OpenCell-LMNB1 nuclear lamina, §) the interpolated value is the credited one, because the H.265 curve is still rising through the operating rate. On 12 of the 17 volumes the envelope value differs from the exact-rate one by more than 0.05 dB.

The exact-rate reading (left block) is the non-conservative one. GSplats recover more signal on 16 of the 16 volumes with an exact-rate value, by +0.50 dB (C. elegans embryo nuclei) to +7.27 dB (Neuromast membranes), median +1.32 dB, with 0 of them inside the replicate band; 0 of 17 are flagged as outside the measured H.265 range and one (Tribolium embryo nuclei) is withheld. These margins are larger than the peak-to-peak gaps (Kidney nuclei +3.68 vs. +2.58 dB; Mouse heart nuclei +5.35 vs. +3.44 dB) and exceed the envelope margin wherever H.265 has

Table 3: Matched-rate comparison at the H.265 held-out peak (CRF, bpv, dB) against GSplats interpolated at that bit rate; Δ has the sign convention of Table 2. †: rate below the measured GSplat range; ‡: above it. *: deconvolved before the benchmark.

| Dataset | CRF (bpv) | H.265 dB | GSplats dB | Δ dB |
|---|---|---|---|---|
| OpenCell-MAP4 nuclei | 0 (11.2) | 28.00 | –‡ | – |
| OpenCell-MAP4 microtubules | 15 (4.66) | 24.70 | –‡ | – |
| OpenCell-LMNB1 nuclei | 10 (5.68) | 25.31 | –‡ | – |
| OpenCell-LMNB1 nuclear lamina | 15 (2.78) | 33.48 | –‡ | – |
| Kidney nuclei | 35 (0.367) | 31.33 | 31.11 | −0.22 |
| Kidney actin | 40 (0.322) | 27.55 | 27.20 | −0.35 |
| Cells3D nuclei | 15 (3.51) | 35.02 | 35.13 | +0.11 |
| Cells3D membranes | 25 (0.264) | 44.02 | 44.93 | +0.91 |
| Blastocyst nuclear lamina | 35 (0.106) | 36.43 | 37.54 | +1.11 |
| C. elegans embryo nuclei | 40 (0.00750) | 39.06 | –† | – |
| Fly brain neurons | 35 (0.0121) | 43.64 | 43.29 | −0.35 |
| Tribolium embryo nuclei* | 15 (0.330) | 55.64 | 54.44 | −1.20 |
| Mouse heart nuclei | 30 (0.180) | 41.86 | 39.26 | −2.60 |
| Zebrafish embryo nuclei* | 40 (0.0505) | 36.25 | 35.72 | −0.53 |
| Drosophila embryo nuclei | 40 (0.0813) | 33.03 | 32.52 | −0.51 |
| Neuromast membranes* | 35 (0.0304) | 46.67 | 41.74 | −4.93 |
| Neuromast nuclei* | 25 (0.0151) | 58.02 | 43.20 | −14.82 |

passed its held-out peak before the splat rate.

The "CRFs" column shows how far apart the interpolation endpoints are. The bracket spans a factor of 2.4 in rate at the median, more than two on 12 volumes, widest on the C. elegans embryo nuclei (11.1×), and on three volumes (Tribolium embryo nuclei, Neuromast membranes, Neuromast nuclei) the upper setting is the lossless CRF 0. On the *Tribolium* embryo nuclei the operating rate (1.47 bpv) lies between CRF 5 (0.76 bpv) and the lossless CRF 0 (4.48 bpv), a single log-linear segment across a 5.9× span, so its exact-rate cell is withheld (¶); the envelope reading, which needs no interpolation there, is a tie.

Where the H.265 peak lies relative to the splat rate is a separate question, read off the bpv columns. The H.265 held-out peak sits below the splat operating rate on 12 of the 17 volumes and above it on the other 5 (OpenCell-MAP4 nuclei, OpenCell-MAP4 microtubules, OpenCell-LMNB1 nuclei, OpenCell-LMNB1 nuclear lamina, Cells3D nuclei), where the codec has not reached its held-out peak by the splat rate. The symmetric reading (Table 3) is the honest counterpart. At the H.265 peak rate, GSplats are ahead on only 3 of the 12 volumes where the splat sweep reaches that rate (2 by more than the replicate band; by up to +1.11 dB, Blastocyst nuclear lamina) and behind on 9. The median Δ over those 9 volumes is −0.53 dB and over all 12 comparable volumes −0.43 dB, down to −14.82 dB on the Neuromast nuclei, whose H.265 peak sits at 0.02 bpv where the splat sweep has only its first few thousand splats. On 5 volumes the H.265 peak lies outside the splat sweep's rate range and no comparison is possible. The two held-out rate-distortion curves therefore cross: H.265 is the better recovery at very low bit rates, and Gaussian splats are the better recovery at the rate the cross-validation selects, which is the rate a fitted representation is actually stored at.

Table 4: Lowest bit rates in the unmasked arm on the 17 volumes covered. "Rate" is the range of per-volume minima across volumes; "Factor" is that minimum divided by the same volume's 1K-splat rate, as a range across volumes with the median in brackets. Lossless blosc is the lowest of its three configurations on each volume, on two bases (Sec. 2.5); the stored-dtype rows are split by the dtype of the source and by modality. The H.265 row is its lowest measured rate, CRF 51, in the blind-spot arm against the 1K-splat rate of that arm.

| Method | Setting or subset | Rate (bpv) | Factor above 1K splats |
|---|---|---|---|
| GSplats | 1K splats | 0.002–0.06 | 1 (reference) |
| Quantisation+zstd | 4 bit (`quant4b-zstd`) | 0.02–2.2 | 3.5–407 (40) |
| Lossless blosc, stored dtype | all 17 volumes | 2.3–18.9 | 175–3506 (1079) |
| | `uint16` (14 volumes) | 2.3–11.5 | |
| | `uint8` (Mouse heart) | 2.7 | |
| | `float32` (two neuromast channels) | 16.6–18.9 | |
| | spinning-disk | | 839–2078 |
| | confocal | | 175–1565 |
| | light-sheet | | 1079–3506 |
| | iSIM | | 2371–2667 |
| Lossless blosc, float32 | loader's normalised array | 0.3–25.0 | 47–9762 (1455) |
| H.265 | CRF 51, blind-spot arm | | 0.24–5.9 (1.06) |

##### 3.3 The very-low-rate regime

Figure 3 also carries the two remaining voxel baselines under the same blind-spot protocol: uniform quantisation (4–16 bit) followed by zstd (grey), encoded from the mask-filled volume, and the lowest bit rate the lossless blosc configurations reach on that volume, on the source at its stored dtype (dark ticks rising from the x axis) and on the normalised float32 array (light ticks, with a light triangle at the right end of the axis where the floor lies beyond the plotted range; Sec. 2.5). Quantisation has no spatial model, so at high precision its held-out PSNR approaches that of the mask-fill predictor however many bits it spends (a finite quantisation step can land marginally closer to a target, so we compare against the best of its six measured settings, not a theoretical ceiling). Its raw PSNR beyond the noise floor measures fidelity to the sensor noise, not to the signal, which is why we do not report a raw rate-distortion comparison against it. GSplats exceed the quantiser's best measured held-out PSNR on all 17 volumes, by +0.14 dB (OpenCell-LMNB1 nuclei) to +7.44 dB (Neuromast membranes); on 6 volumes (OpenCell-MAP4 nuclei, OpenCell-MAP4 microtubules, OpenCell-LMNB1 nuclei, OpenCell-LMNB1 nuclear lamina, Cells3D nuclei, C. elegans embryo nuclei) the margin is below 0.6 dB.

The unmasked arm (fits of the original volume scored against it, alongside the quantisation and blosc rows of the same arm) shows the regime the other two voxel baselines do not reach in the configurations tested. Table 4 collects the lowest bit rate per method on the 17 volumes covered and its factor above the same volume's 1K-splat rate (per-volume values in Fig. 4).

The 1K-splat rate against the stored-dtype floor is the comparison that holds for the source image data. The two blosc bases are not interchangeable. On the 15 integer-stored volumes the float32 cast inflates the floor 1.3–2.9× (median 2.4×). On the two float32-stored neuromast channels the normalised array (floor-subtracted and clipped, hence mostly exact zeros) compresses 30–50× *smaller* than the stored frame. The float32 numbers are therefore the rate of the fits' input payload, never the floor of the source.

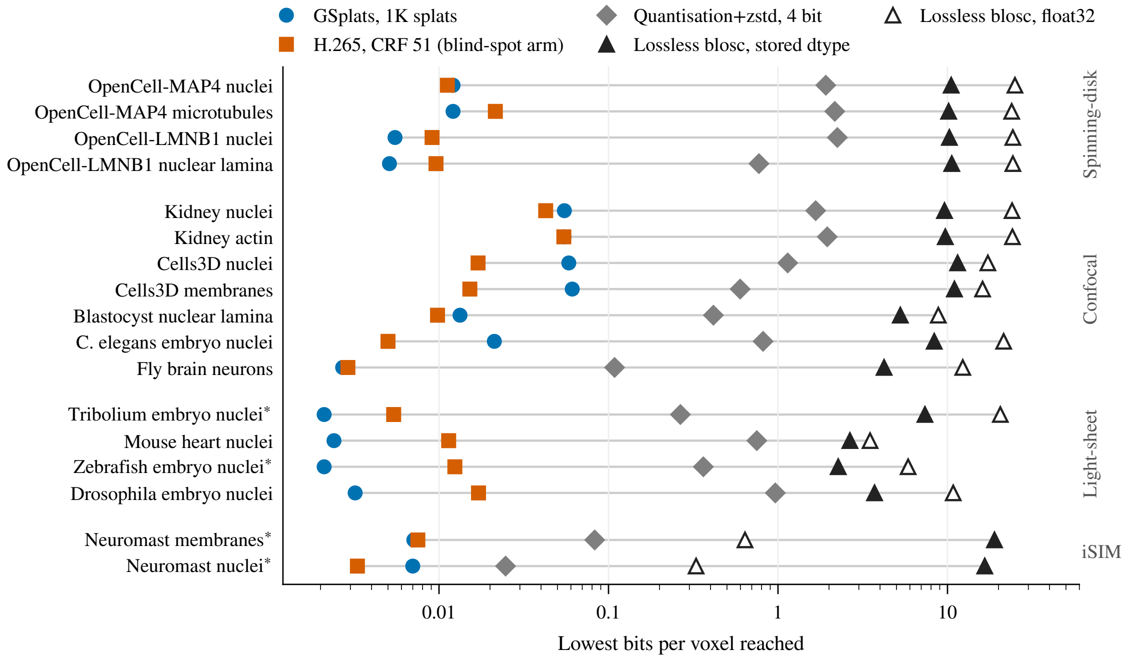


Figure 4: Lowest bit rate reached per volume by each baseline family against the 1K-splat rate: the per-volume values behind the ranges of Table 4. One row per volume in the order of Table 1, grouped by modality, on a log axis, so the horizontal distance from the blue circle to a marker is that row's factor above 1K splats (one decade is 10×). Blue circles: GSplats at 1K splats, unmasked arm. Grey diamonds: quantisation+zstd at 4 bit, its lowest setting. Triangles: the lowest of the three lossless blosc configurations on the source at its stored dtype (filled) and on the loader's normalised float32 array (open; Sec. 2.5). Vermilion squares: H.265 at CRF 51, its lowest measured rate, which exists only in the blind-spot arm, as in the last row of Table 4. A grey line joins each row's extremes. On the two float32-stored neuromast channels the open triangle lies left of the filled one: the normalised array, mostly exact zeros, compresses below the stored frame. *: deconvolved before the benchmark, as in Table 1.

At those very low bit rates GSplats deliver 21–38 dB PSNR against the original volume, depending on the volume: 22.4 dB on the Kidney nuclei at 0.055 bpv against 38.3 dB on the *C. elegans* embryo nuclei at 0.021 bpv. The low end is visibly lossy on the noisier confocal data and the high end is close to the source on the sparser, cleaner volumes. H.265 is a different case. Its lowest measured rate (CRF 51, blind-spot arm) is 0.24–5.9× the 1K-splat rate of the same arm (median 1.06×), lower on 8 of the 17 volumes, so it is the closest voxel-based competitor in the very-low-bpv regime. Its output is a coded voxel stream that is decoded to voxels before rendering, not a renderable primitive set (Sec. 4).

##### 3.4 Compression at the cross-validation optimum

Because the held-out PSNR provides a per-dataset stopping criterion (main paper, Fig. 3d), we can also report the compression ratio at the principled stopping point. Across the 17 benchmark volumes with a held-out sweep, the GSplat compression ratio at $K^*$ ranges from 6.0× (Kidney actin, optimum at 128K splats) to 340× (Neuromast nuclei, optimum at 64K splats), with a median of 99.3×. A finite optimum is reached on 15 of the 17 volumes. The exceptions are light-sheet volumes whose held-out PSNR is still rising at the largest tested capacity (signal-limited curves), so their ratio is reported at $K^*$, the smallest count within 0.3 dB of the maximum, which is the endpoint of the sweep: the Mouse heart nuclei at 2048K splats (6.3×; its noise floor sits below the precision of the blind-spot estimators) and the *Tribolium* embryo nuclei at 2048K (10.8×). The deconvolved Zebrafish embryo nuclei do reach a finite optimum: a held-out plateau from 256K (113.3×), with the maximum at 1024K and a decline at the 2048K endpoint of the extended sweep smaller than the replicate band.

### 4 Discussion

**What the two comparisons measure.** The matched-rate comparison (Sec. 3.2) is the compression claim: for the bits a fitted splat representation actually occupies, H.265 read under the envelope rule recovers less of the signal on 15 of 17 volumes and ties on 2 (median +0.98 dB); encoded to exactly that size it recovers less on 16 of 16 (median +1.32 dB). The peak comparison (Sec. 3.1) is a model-selection claim: the best held-out prediction each method reaches when its knob is chosen by cross-validation, irrespective of the bits spent. GSplats reach the higher optimum on 12 of 17 by more than the replicate band (16 by sign), but the optima occur at different rates, so that gap should not be read as a rate-distortion result. Neither claim extends to the very-low-rate regime, where H.265 leads (9 of 12 comparable volumes at the H.265 peak rate).

**Why GSplats win on noisy data.** The Gaussian-splat representation is a global mixture model with smooth, exponentially-decaying basis functions. As capacity grows, the representation can interpolate masked regions from the surrounding fitted splats but cannot easily reproduce the high-frequency, voxel-independent fluctuations of sensor noise: fitting such noise would require many low-amplitude, sub-voxel-scale splats. Two ingredients of the fitter plausibly reinforce this bias, though neither has been ablated here and neither is a guarantee. The L1 amplitude regulariser ($\lambda_a = 10^{-3}$ in the main paper Methods) penalises the *mean* absolute amplitude (it favours small amplitudes overall and does not penalise the number of splats). The fixed-pool dynamic *relocation* step (every 50 iterations) moves the least important splats to the strongest residual peaks found by non-maximum suppression, which concentrates capacity where the residual is largest without, by itself, telling an isolated noisy voxel from signal. The effect observed here is an implicit denoising bias: the model prefers low-frequency reconstructions that match the structural signal, and consequently predicts the held-out values better than methods that allow high-frequency residual to be encoded.

**A hypothesis for the *Tribolium* tie.** One reading of the *Tribolium* result is a smoothness effect: that volume's post-deconvolution statistics may match the prior built into block-DCT codecs (smooth, locally low-frequency patches with sharp edges), so that at its held-out peak (CRF 15) H.265 reconstructs the held-out voxels at 55.64 dB and the GSplat curve only draws level with it at 2048K splats (55.55 dB, a tie at 4.5× the bit rate). The Zebrafish embryo nuclei volume is also multi-view-deconvolved, and there H.265 saturates at 36.25 dB and GSplats win by +1.74 dB; on the Mouse heart nuclei H.265 saturates at 41.86 dB and GSplats win by +3.44 dB.

The fill statistic of Table 1 does not bear the hypothesis out (Sec. 3.1): the Neuromast nuclei's held-out voxels are at least as predictable from their neighbours (56.34 vs. 53.83 dB), H.265 clears its fill there by a similar margin (+1.68 vs. +1.81 dB), and GSplats lead by +5.58 dB. Whether the volume was deconvolved is not the determinant either, so we report the light-sheet outcomes individually and treat the *Tribolium* explanation as open.

**Direct renderability.** A property not captured by any rate-distortion plot: the GSplat representation is consumed directly by the GPU rasteriser. The store's compressed chunks are entropy-decoded like any other, but the decoded parameters go straight into vertex/instance buffers and are rendered without reconstructing a voxel grid. The voxel baselines here are decoded to a dense voxel buffer before rendering: at the volume sizes typical of light-sheet microscopy ($\sim 10^8$ voxels) that buffer is 100–400 MB at 1–4 bytes per voxel, before anything reaches the GPU. This is a property of our decode-to-file baselines, not of voxel formats as such (a chunked or multiscale voxel format decodes a region or level at a time, and a video decoder emits frames sequentially), but every voxel pipeline materialises the voxels it displays, whereas the splat representation never re-grids. For the Luxar use case of streaming multidimensional data into a web viewer, this is the property the viewer builds on.

**Limitations.** The blind-spot framework assumes pixel-independent noise; volumes that have already been denoised, deconvolved, or pre-processed in any other way that couples neighbouring voxels do not satisfy this assumption, and the held-out PSNR then measures prediction of the processed intensities, not recovery of an independent clean signal. The four deconvolved volumes of this benchmark (Tribolium embryo nuclei, Zebrafish embryo nuclei, Neuromast membranes, Neuromast nuclei; * in the tables) include the two iSIM channels that supply the largest matched-rate margins, so those margins are to be read in that light; *Tribolium* is the cleanest example of the effect on the codec side, but any preprocessing that introduces spatial correlation in the residual will move the optimum away from the unbiased estimator. We do not include "denoise then compress" pipelines (e.g., Noise2Self pretraining followed by H.265) in the present comparison. The H.265 arm is a single configuration, `libx265` at preset `medium` with its default tuning, and we tested no PSNR-tuned or slower preset, no other codec (HEVC range extensions, AV1 with 12-bit profiles) and no denoise-then-compress control, so every H.265 conclusion above is a statement about this configuration. Whether such alternatives would narrow the gap, and by how much, is not measured here.

#### 5 Summary

Four readings of one pair of held-out curves per volume give one consistent picture. Peak to peak, the splat optimum is the higher one on 12 of 17 volumes beyond the replicate band, with 5 ties inside it (OpenCell-MAP4 nuclei, OpenCell-MAP4 microtubules, OpenCell-LMNB1 nuclei, C. elegans embryo nuclei, Tribolium embryo nuclei), the *Tribolium* embryo nuclei the only one with a negative gap, and the peaks sit at different rates. At the splat operating rate, the envelope rule leaves GSplats ahead on 15 of 17 volumes and tied on 2; the exact-rate reading, non-conservative because the codec has often passed its peak by then, is ahead on every one of the 16 volumes with a value, with one cell withheld where the interpolation would span a lossy-to-lossless bracket. At the H.265 peak rate the codec is ahead on 9 of 12 comparable volumes. The curves cross, and the crossing lies below the rate at which a fitted representation is stored. Below the lowest rate of the quantisation and lossless baselines, 1K-splat fits still render, and every splat set is rendered from its decoded parameters without a voxel grid. The H.265 arm is one encoder at one preset, the codec arm has no replicates, and four volumes violate the independence assumption of the score; each of those limits the reach of the claim.

[2] Francesc Alted and The Blosc Development Team. Blosc: a blocking, shuffling and lossless compression library. https://www.blosc.org.

[3] Joshua Batson and Loïc Royer. Noise2Self: Blind denoising by self-supervision. In *International Conference on Machine Learning*, pages 524–533. PMLR, 2019.

[4] Yann Collet and Murray Kucherawy. Zstandard compression and the 'application/zstd' media type. RFC 8878, Internet Engineering Task Force, 2021.

[5] MulticoreWare, Inc. x265 HEVC encoder. https://www.videolan.org/developers/x265.html.

[6] Gary J. Sullivan, Jens-Rainer Ohm, Woo-Jin Han, and Thomas Wiegand. Overview of the High Efficiency Video Coding (HEVC) standard. *IEEE Transactions on Circuits and Systems for Video Technology*, 22(12):1649–1668, 2012. doi: 10.1109/TCSVT.2012.2221191.
