## Supplementary material for "Luxar: Gaussian splatting for microscopy and scalable interactive web visualisation of multidimensional scientific data": supp_doc_08_substitutive_lod

### Substitutive Levels of Detail for Gaussian Splats: Cost Metrics, Bin-wise Merging, and Hierarchical Construction

Supplementary Document 8 – Luxar

**Abstract**

A substitutive level of detail stands in for a Gaussian-splat dataset of $N$ splats with $\lceil N/K \rceil$ merged splats, for a compression factor $K > 1$, by grouping neighbouring splats and replacing each group with one Gaussian. Three results follow from experiments on synthetic mixtures and on a 7,740-splat fit of DAPI-stained nuclei. First, the sum of per-group errors is a faithful surrogate for the global $L^2$ error in the median once a spatial partition has separated the groups; single trials still deviate by tens of percent at moderate separation, and the separation this needs is not monotone in $K$: $s = 1.8$, 0.8 and 2.5 at $K = 2$, 4 and 8. Second, a $k$-means partition refined by cost-increment relocation matches greedy hierarchical merging to within 5.5 % of the median relative error at $K \geq 4$; greedy leads by up to 20.1 % at $K = 2$. Third, on the full fit that recipe reaches a relative $L^2$ error of 0.104 at $K = 4$, against 0.198 for culling with refitted amplitudes and 0.312 for plain culling; on subsamples whose neighbour spacing is 3.8× wider, both culling baselines win in the median. Neighbour spacing against splat width decides, not the merging algorithm. The theory behind these results surveys the candidate cost metrics, derives the $K$-wise moment-matched merge with $L^2$-optimal amplitude, decomposes the global error into per-group residuals plus an explicit cross-group interference term, and maps the algorithms onto the scalable variants that Luxar ships.

**10 Discussion and Open Questions** **29**

**11 Summary** **30**

### 1 Introduction and Problem Formulation

A scientific scene of Gaussian splats is often viewed from far away: the whole dataset occupies a few hundred pixels, yet every splat is still fetched, held in memory and drawn. The remedy is a coarser stand-in with fewer splats, shown whenever the fine one would be wasted. The question here is how such a stand-in should be built. Given $N$ splats and a compression factor $K$, we want the $\lceil N/K \rceil$ splats that best reproduce the scalar field the original ones define, a cost metric for that fidelity with a surrogate that can be optimised at scale, and a way to find them; how far they are from the best possible set is not known, and Section 6 shows why the natural spectral lower bound fails and claims no analytical one. Two families of answers exist: remove splats and keep the survivors verbatim, or merge groups of neighbouring splats into single new ones. The document develops the second family, substitution, and compares it with the first.

A Luxar scene of Gaussian splats represents a non-negative scalar field

$$f(\boldsymbol{x}) \;=\; \sum_{i=1}^{N} \phi_i(\boldsymbol{x}), \qquad \phi_i(\boldsymbol{x}) \;=\; a_i \exp\!\Big(-\tfrac{1}{2}\,(\boldsymbol{x}-\boldsymbol{\mu}_i)^{\top}\boldsymbol{\Sigma}_i^{-1}(\boldsymbol{x}-\boldsymbol{\mu}_i)\Big), \tag{1}$$

with peak amplitudes $a_i > 0$, centres $\boldsymbol{\mu}_i \in \mathbb{R}^D$, and positive definite covariances $\boldsymbol{\Sigma}_i \succ 0$. In the Luxar implementation each covariance is stored via its Cholesky factor $\boldsymbol{L}_i$ such that $\boldsymbol{\Sigma}_i = \boldsymbol{L}_i\boldsymbol{L}_i^{\top}$. Scientific scenes routinely reach $N \in [10^5, 10^7]$ splats. Count alone does not exhaust a commodity GPU: the viewer benchmark of Supplementary Document 6 holds its display rate through $10^6$ splats on a reference scene. But the work per frame, the bytes that must be fetched before anything is drawn, and the memory held resident all grow with $N$, whether or not the scene occupies more than a few pixels. A standard remedy is to provide multiple *levels of detail* (LOD): the viewer renders a coarser representation when fewer pixels are dedicated to the scene, and fetches the finest representation only on demand, so that the cost follows the screen footprint and the view, not the dataset size.

**Substitutive vs. removal LOD.** Two qualitatively different mechanisms produce a coarser representation from a finer one (Fig. 1):

- **Removal** (subset selection). Choose a subset $\mathcal{I} \subseteq \{1, \dots, N\}$ of size $M = \lceil N/K \rceil$ and retain $\{\phi_i\}_{i\in\mathcal{I}}$ verbatim. Removal is the strategy of Luxar's culling routines (*cull (retention)*); the contribution-based cull works from an $L^\infty$ error budget when the fitted volume is available and from a fractional-contribution threshold otherwise (see Section 8).
- **Substitution** (mixture reduction). Partition $\{1, \dots, N\}$ into $M$ *bins* $S_1, \dots, S_M$ (disjoint groups of splats that will be merged together) and replace each bin by a single representative splat $\bar{\phi}_j$, yielding the approximation

$$g(\boldsymbol{x}) \;=\; \sum_{j=1}^{M} \bar{\phi}_j(\boldsymbol{x}). \tag{2}$$

The two strategies live on opposite ends of a spectrum. Removal is faithful to the original splats

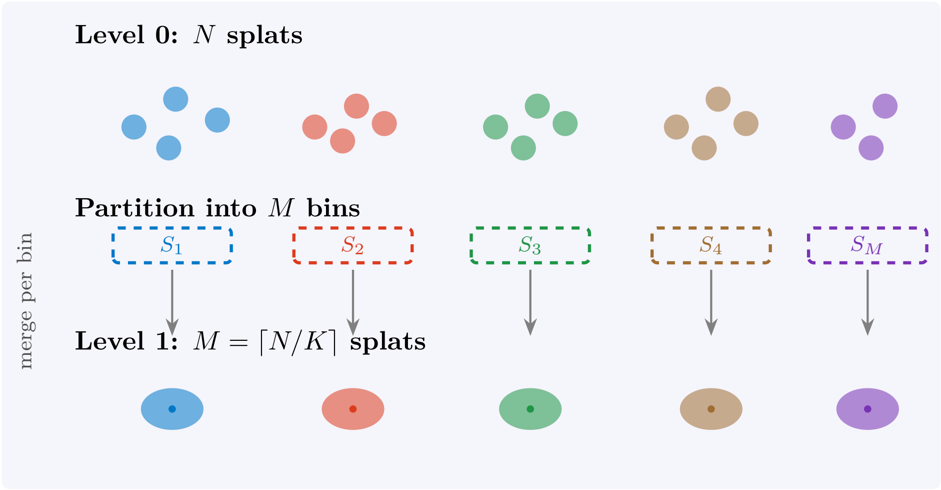

Figure 1: Substitutive LOD construction. The $N$ original splats are partitioned into $M = \lceil N/K \rceil$ bins; each bin is replaced by a single representative splat $\bar{\phi}_j$. The level-1 scene is the sum of $M$ representatives.

but cannot represent local *averages* of removed splats; substitution synthesises new splats that may lie outside the original set but can compactly summarise locally redundant clusters. This document develops the theory of substitution; Section 8 contrasts the two empirically.

**The two subproblems.** Substitutive LOD decomposes naturally into:

1. **Per-bin merge.** Given a bin $S_j$ of about $K$ splats, find the single splat $\bar{\phi}_j$ that best approximates $\sum_{i \in S_j} \phi_i$ under a chosen cost metric.
2. **Bin assignment.** Find the partition $\mathcal{P} : \{1, \ldots, N\} \to \{1, \ldots, M\}$ that minimises the total approximation error of $g$ to $f$.

The two problems interact: the optimal partition depends on what each bin can be approximated by, and the optimal representative depends on which splats fall in the bin. In analogy with $k$-means clustering [17], this motivates a cost-increment relocation between the two; we develop several variants in Section 5.

**Relation to mixture reduction.** The substitutive-LOD problem is closely related to *Gaussian mixture reduction*, a long-standing problem in target tracking and density estimation [6, 22, 23, 25, 26]. The classical formulation considers a *normalised* mixture $\sum_i w_i \mathcal{N}(\boldsymbol{x}; \boldsymbol{\mu}_i, \boldsymbol{\Sigma}_i)$ with $\sum_i w_i = 1$, and reduces it to a smaller normalised mixture under a divergence such as integrated squared error [24, 26] or KL [22]. Two aspects distinguish the substitutive-LOD problem in Luxar:

- Splats carry *amplitudes* (peak heights), not mixture weights; the underlying object is a non-negative scalar field, not a probability distribution. Generalised KL and Bregman divergences for non-negative measures [4, 11] (a Bregman divergence is the gap between a convex function and its tangent plane, of which squared distance and KL are the two familiar cases) are therefore more natural than ordinary KL.

- The approximation must be *embedded back into the renderer*, which evaluates $g(\boldsymbol{x})$ at sampled query points instead of comparing distributions. Pointwise integrated error metrics ($L^2$, $L^\infty$) are aligned with the rendering pipeline; the optimal-transport family is not.

**Companion document.** The pairwise ($K = 2$) merging problem under $L^2$ is treated in detail in the companion supplementary material (Supplementary Document 1) [18], which establishes the closed-form Gaussian inner product, the moment-matched single-Gaussian approximation, and the $\sin^2\alpha$ pairwise mergeability score. The present document generalises that machinery in three directions: from $K = 2$ to arbitrary $K$; from a fixed $L^2$ metric to a family of metrics; and from a single merge to the full LOD hierarchy.

**Outline.** Section 2 sets notation. Section 3 surveys cost metrics and argues for the recommended ones. Section 4 generalises moment matching to bins of size $K$ and gives closed-form bin merges in $L^2$ and Cauchy–Schwarz. Section 5 develops the bin-assignment subproblem, including the explicit cross-bin interference term, and states the three partition algorithms as pseudocode where each is introduced. Section 6 records what is, and is not, known about lower bounds. Section 7 discusses multi-level LOD. Section 8 relates the framework to the algorithms Luxar ships. Section 9 reports numerical experiments. Section 10 discusses limitations and open questions.

#### Terms used in this document

**cull (retention)** The removal of splats after a fit. The fit-time cull ranks splats by amplitude and keeps the top set accounting for a retention fraction of the total amplitude (0.95 by default); the contribution-based cull instead removes the splats whose contribution to the rendered field falls below an error budget or a fractional threshold, and is the baseline that removal-based level-of-detail methods use.

**leaf** A node that carries splat arrays directly (centres, amplitudes, Cholesky factors) together with an additive ladder of at least one sub-LOD, as opposed to a group node that only holds children.

**`kind=lod` group** A substitutive level-of-detail group whose children are alternative renderings of the same content at different splat counts, of which the viewer draws exactly one at a time: the coarsest level whose median projected splat footprint stays at or below 1.5 CSS pixels when the levels carry footprint stamps, otherwise the level selected by the size of its projected bounding box (its fraction of the viewport area for a screen-area ladder, its normalised projected diagonal for a coverage ladder).

**`kind=partition` group** A group whose `part_<i>` children each carry their own position bounds and are all drawn at once, each culled against the camera frustum; the parts are usually the regions of a recursive BSP split of one splat set, but any disjoint grouping is valid, such as one part per timepoint.

**recipe** A named LOD topology built from a fitted splat set: `flat` (one bare leaf), `stream` (one stream-laddered leaf), `levels` (coarse-to-fine replacement levels), `tiles` (a BSP partition with one ladder per tile), `overview` (one coarse level above a `tiles` branch) and `adaptive` (`tiles` where every tile carries its own `levels`).

**additive (streaming) ladder** One ordering of a leaf's splats cut into prefix-sum sub-LODs, so that every prefix of the download is a valid, progressively refining rendering of the same splat set.

**rung (sub-LOD)** One contiguous slab of an additive ladder, stored as its own group (`additive_<i>`, index 0 the coarsest) and appended to the committed prefix when it arrives; a document that uses "rung" for one step of a splat-count sweep says so where it first does.

**self-energy** The squared $L^2$ norm of one splat's Gaussian, $a_i^2\,\pi^{D/2}\,|\Sigma_i|^{1/2}$, the ordering weight of an additive ladder above the greedy size limit of 5,000 splats (below it a greedy residual ordering is used), the unit of the equal-energy breakpoints, and the quantity whose cumulative fraction stamps how much of a leaf a ladder prefix already carries.

**barrier dimension** A centre column (a time or channel axis) that the compiler and the LOD builders never mix across. The writer sorts the elements by that column first and computes the spatial-index curve over the spatial columns only, so every coordinate's elements are contiguous on disk; substitutive coarsening leaves the column out of `coarsen_dims`, so every coarse level stays separated per coordinate; and an additive ladder is built per coordinate, so no rung blends two of them.

**Morton and Hilbert ordering** Two space-filling curves that sort points so that neighbours in space become neighbours in memory or on disk: Morton (Z-order) interleaves the bits of the quantised coordinates, Hilbert follows a continuous curve whose consecutive cells are always adjacent, so it has no long jumps between neighbouring runs. The compiler orders the stored elements along one of them; the fitter re-sorts its splats by Morton code periodically for GPU locality, and the substitutive LOD builder uses a Morton sort as its warm-start partition.

#### 2 Notation and Setup

We work in $\mathbb{R}^D$ with $D \in \{2, 3\}$ for visualisation, but the formulas are dimension-agnostic. The notation extends the companion document [18].

**Splats and the Gaussian inner product.** A splat $\phi_i$ is the unnormalised Gaussian (1): unnormalised because its peak amplitude $a_i$ is free, so its integral is not one. Throughout, $|\mathbf{\Sigma}|$ denotes the determinant of a covariance matrix and $\pi$ the constant. The $L^2$ *self-energy* of a splat is $\|\phi_i\|^2_{L^2} = a_i^2\, \pi^{D/2}\, |\mathbf{\Sigma}_i|^{1/2}$. The $L^2$ inner product between two splats has the closed form

$$K_{ij} \;=\; \langle \phi_i, \phi_j \rangle_{L^2} \;=\; a_i a_j\, (2\pi)^{D/2}\, \frac{\sqrt{|\mathbf{\Sigma}_i||\mathbf{\Sigma}_j|}}{\sqrt{|\mathbf{\Sigma}_i + \mathbf{\Sigma}_j|}}\, \exp\!\Big(-\tfrac{1}{2}(\boldsymbol{\mu}_i-\boldsymbol{\mu}_j)^\top(\mathbf{\Sigma}_i+\mathbf{\Sigma}_j)^{-1}(\boldsymbol{\mu}_i-\boldsymbol{\mu}_j)\Big), \tag{3}$$

derived in detail in [18]. The Gram matrix $\boldsymbol{G} \in \mathbb{R}^{N\times N}$ has entries $\boldsymbol{G}_{ij} = K_{ij}$.

**Bins, partitions, representatives.** A partition $\mathcal{P}$ of $\{1, \dots, N\}$ is a collection of disjoint bins $S_1, \dots, S_M$ whose union is the whole index set; $\mathcal{P}(i)$ is the index of the bin holding splat $i$. We write $|S_j|$ for the bin size. The partition algorithms below do not force equal bins, so $|S_j|$ is only about $K$ on average, with $M = \lceil N/K \rceil$ bins in total. The bin function $f_{S_j}(\boldsymbol{x}) = \sum_{i\in S_j} \phi_i(\boldsymbol{x})$ collects the mass inside bin $j$. Each bin has an associated representative splat $\bar{\phi}_j$, parameterised by $(\bar{a}_j, \bar{\boldsymbol{\mu}}_j, \bar{\mathbf{\Sigma}}_j)$; its unit-amplitude template is $\tau_j(\boldsymbol{x}) = \exp(-\frac{1}{2}(\boldsymbol{x} - \bar{\boldsymbol{\mu}}_j)^\top \bar{\mathbf{\Sigma}}_j^{-1}(\boldsymbol{x} - \bar{\boldsymbol{\mu}}_j))$, so that $\bar{\phi}_j = \bar{a}_j\, \tau_j$. The approximation $g$ is the sum (2); a generic candidate function that has not been fixed yet is written $q$, and the levels of a multi-level hierarchy (Section 7) are $g_\ell$, with $g_0 = f$ and $g_1 = g$.

**Mass.** The integral mass of a splat is $m_i = a_i\, (2\pi)^{D/2}\, |\mathbf{\Sigma}_i|^{1/2}$, equal to its zeroth moment. Within bin $S_j$, the mass-weighted parameters are

$$m_j^{\text{tot}} = \sum_{i\in S_j} m_i, \qquad w_i^{(j)} = m_i / m_j^{\text{tot}}, \quad i \in S_j. \tag{4}$$

### 3 Cost-Metric Design Space

The choice of *which* discrepancy between $f$ and $g$ to minimise is the central design decision of substitutive LOD. Different choices yield different bin merges, different partition objectives, and different per-level error bounds. We deliberately survey the design space instead of committing prematurely to one metric, then argue from first principles for the recommended ones.

#### 3.1 Catalogue of candidate metrics

**Squared $L^2$ (integrated squared error, ISE).** The squared $L^2$ distance is $\|f - g\|_{L^2}^2 = \int (f - g)^2$. It is closed-form for Gaussian mixtures through the kernel (3) and was developed pairwise in the companion document. It is aligned with the voxel-domain rendering metrics (PSNR, MSE). It is *not* bin-additive: the residuals $r_j = f_{S_j} - \bar{\phi}_j$ of different bins are not $L^2$-orthogonal in general (Section 5).

**Cauchy–Schwarz divergence between mixtures.** For two non-zero, square-integrable functions $p, q \in L^2$ (a non-negative $L^1$ function need not be in $L^2$), the divergence is

$$D_{\mathrm{CS}}(p, q) \;=\; -\log \frac{\langle p, q\rangle^2}{\langle p, p\rangle \, \langle q, q\rangle} \;\geq\; 0, \tag{5}$$

with equality iff $p \propto q$. All three inner products are closed-form sums of pairwise Gaussian products (3), which yields a fully analytical divergence between Gaussian mixtures [13, 14, 19]. The divergence is scale-invariant in its second argument, $D_{\mathrm{CS}}(p, c\,q) = D_{\mathrm{CS}}(p, q)$ for any $c > 0$, so it does not constrain the total mass of $g$.

**Generalised Kullback–Leibler / I-divergence.** For non-negative measures $p, q$ the generalised KL divergence is $D_{\mathrm{gKL}}(p, q) = \int [p \log(p/q) - p + q]$, which extends the probability-measure KL to unnormalised functions [4, 11]. It is natural for splats because their total mass is meaningful. However, $\log(p/q)$ between two Gaussian mixtures has no closed form [12]; it must be estimated, by Monte Carlo sampling or by one of the deterministic approximations (Jensen-type variational bounds, matched-pair and unscented schemes) reviewed there.

**$W_2$ optimal transport.** The 2-Wasserstein distance between two Gaussians is $W_2^2(\mathcal{N}_1, \mathcal{N}_2) = \|\boldsymbol{\mu}_1 - \boldsymbol{\mu}_2\|^2 + \mathrm{tr}(\boldsymbol{\Sigma}_1 + \boldsymbol{\Sigma}_2 - 2(\boldsymbol{\Sigma}_1^{1/2}\boldsymbol{\Sigma}_2\boldsymbol{\Sigma}_1^{1/2})^{1/2})$ [1, 2]. For *mixtures*, $W_2$ has no closed form. The mixture-Wasserstein distance $\mathrm{MW}_2$ of Delon and Desolneux [7] (and the related construction of Chen et al. [5]), often called Gaussian-mixture optimal transport (GMM-OT), restricts the coupling to component-to-component transport. It is a genuine metric on Gaussian mixtures with $\mathrm{MW}_2 \geq W_2$. It is evaluated by a linear programme over a small cost matrix whose entries are the closed-form Gaussian $W_2^2$ above. The coupling, the matrix saying how much of each source component flows to each target component, is the solution of that linear programme (an LP coupling). The costs are therefore closed-form, the coupling is solved numerically, and the value is not the true mixture $W_2$. Balanced transport requires equal total mass. The metric is geometric and intuitive.

**Maximum mean discrepancy (MMD).** The MMD is the distance between two functions after each is embedded into a reproducing-kernel Hilbert space (RKHS), the space of functions spanned by a positive-definite kernel $\kappa$ in which evaluation at a point is an inner product with the kernel centred there. With kernel $\kappa$ and RKHS $\mathcal{H}_\kappa$, $\mathrm{MMD}^2_\kappa(p,q) = \int\int \kappa(\boldsymbol{x},\boldsymbol{x}')\,(p-q)(\boldsymbol{x})\,(p-q)(\boldsymbol{x}')\,d\boldsymbol{x}\,d\boldsymbol{x}'$. For Gaussian-mixture inputs and the unit-integral Gaussian kernel $\kappa_\sigma(\boldsymbol{x},\boldsymbol{x}') = (2\pi\sigma^2)^{-D/2}\exp(-\|\boldsymbol{x}-\boldsymbol{x}'\|^2/(2\sigma^2))$, the double integral is closed-form through the same kernel identity (3). Writing $h_\sigma = \mathcal{N}(0, \frac{1}{2}\sigma^2 I)$ for the convolution square root of $\kappa_\sigma$ ($h_\sigma \star h_\sigma = \kappa_\sigma$), one has $\mathrm{MMD}^2_{\kappa_\sigma}(p,q) = \|(p-q)\star h_\sigma\|^2_{L^2}$. The MMD is then an $L^2$ distance between the fields smoothed at bandwidth $\sigma/\sqrt{2}$, which downweights high-frequency discrepancies, and $\mathrm{MMD}^2_{\kappa_\sigma} \to \|p-q\|^2_{L^2}$ as $\sigma \to 0$. The normalisation matters: with the peak-one RBF kernel the same limit is zero.

**$L^1$ and total variation.** The $L^1$ distance is $\|f-g\|_{L^1} = \int |f-g|$. It is robust to local outliers and has no closed form for Gaussian mixtures. It is useful as a complementary *audit* metric.

**$L^\infty$ (max absolute error).** The $L^\infty$ distance is $\|f-g\|_\infty = \sup_{\boldsymbol{x}} |f(\boldsymbol{x}) - g(\boldsymbol{x})|$. It provides a worst-case voxel guarantee and is the metric that Luxar's contribution-based cull uses in its error-budget mode (Section 8). It is a peak measure, not an integrated one: it is attained wherever the residual field is largest (typically at a splat centre or where several residuals stack), and it is not bin-additive.

**Rendering / perceptual loss.** The metric the user actually cares about is the perceptual quality of the rendered image. Perceptual quality is intractable analytically and depends on viewpoint, lighting, and downstream image processing; it is not a candidate for the bin objective. It would be the natural endpoint sanity check; the experiments of Section 9 do not include one.

##### 3.2 First-principles criteria

A cost metric $d(f,g)$ for substitutive LOD should satisfy as many of the following as possible:

1. **Sampling fidelity.** $d(f,g)$ should be small whenever $g(\boldsymbol{x}) \approx f(\boldsymbol{x})$ at most $\boldsymbol{x}$.

2. **Mass preservation.** Splats represent positive scalar fields; the integral $\int g \approx \int f$ should be guaranteed per bin or by construction.

3. **Closed-form bin objective.** The per-bin minimum $\min_q d(f_{S_j}, q)$ over single Gaussians $q$ should be expressible in closed form (or at least via a small fixed-point) so that bin-wise updates are cheap.

4. **Approximate bin additivity.** The global error $d(f,g)$ should decompose, at least approximately, as $\sum_j d(f_{S_j}, \bar{\phi}_j)$. Additivity makes per-bin optimisation a rigorous surrogate for global optimisation.

5. **Compatibility with the Luxar parameterisation.** Splats are stored as $(\bar{a}, \bar{\boldsymbol{\mu}}, \boldsymbol{L})$ with $\bar{\boldsymbol{\Sigma}} = \boldsymbol{L}\boldsymbol{L}^\top$; the metric should not require explicit covariance inversion or whitening at every evaluation.

Table 1: Properties of candidate cost metrics for substitutive LOD, judged on the criteria of Section 3.2. The columns are defined in the text above the table and the two footnote marks below it.

| **Metric** | **Closed form** | **Bin-additive** | **Total mass** | **Used in code** | **Recommended role** |
|---|---|---|---|---|---|
| $L^2$ (ISE) | yes | no$^\dagger$ | sensitive; optimal amplitude $\neq$ mass | yes (merge cost, `lod/_kernels`) | primary objective |
| Cauchy–Schwarz | yes | no | invariant (scale-free) | no | secondary, scale-invariant |
| generalised KL | no | no$^\ddagger$ | sensitive; conserved at free-scale optimum | no | sanity check, audit |
| $W_2$ (mixtures) | no | no | required equal | no | geometric reasoning |
| $\mathrm{MW}_2$ (GMM-OT) | costs yes; LP coupling | no | required equal | no | geometric reasoning |
| $\mathrm{MMD}_\kappa$ (Gaussian $\kappa$) | yes | like $L^2$ | sensitive | no | smoothed $L^2$ variant |
| $L^1$ / TV | no | no$^\ddagger$ | sensitive | no | audit only |
| $L^\infty$ | no | no | sensitive | yes (contribution cull) | audit, certification |

$^\dagger$ The cross-bin interference term $2\sum_{j<k}\langle r_j, r_k\rangle$ admits an explicit closed form (Section 5) and vanishes when bins are well-separated. $^\ddagger$ Integrals of a pointwise integrand: exactly additive over bins with disjoint supports and approximately so when bins are well-separated, like $L^2$, but with no closed form for the cross-bin discrepancy.

Table 1 summarises which metrics satisfy which criteria. In its columns, "closed form" means that the metric admits an analytical formula between two Gaussian mixtures (or between a mixture and a single Gaussian); "bin-additive" that the global cost decomposes as a sum of per-bin costs without a cross-bin term; "total mass" records how the cost treats a difference between $\int g$ and $\int f$: *sensitive* (the cost registers it), *invariant* (it does not) or *required equal* (the metric is only defined for equal masses). None of the sensitive costs constrains an arbitrary candidate $q$ to equal mass. For generalised KL the unconstrained optimum over a freely scaled fixed shape does land on the target mass, while for $L^2$ it does not (Proposition 4.3). "Used in code" means implemented in the Luxar codebase.

##### 3.3 Recommended two-tier framework

Based on Table 1 and the criteria of Section 3.2, we adopt a two-tier framework:

- **Primary objective: $L^2$.** $L^2$ is closed-form, aligned with the Luxar pixel/voxel pipeline (rendering equivalent to MSE on query samples), and continuous with the companion document [18]. Its non-additivity across bins is not a fatal flaw: Section 5 shows that the cross-bin term is itself closed-form and vanishes for spatially separated bins; Section 9 finds numerically that it shrinks to a few percent of the global error in the median once the bins are spatially decoupled, while remaining noisy trial-to-trial.

- **Closed-form alternative: Cauchy–Schwarz divergence.** CS-divergence is closed-form between Gaussian mixtures (it expands into the same pairwise integrals as $L^2$, as does Gaussian-kernel MMD). Unlike MMD it is scale-invariant, which makes it useful for ranking

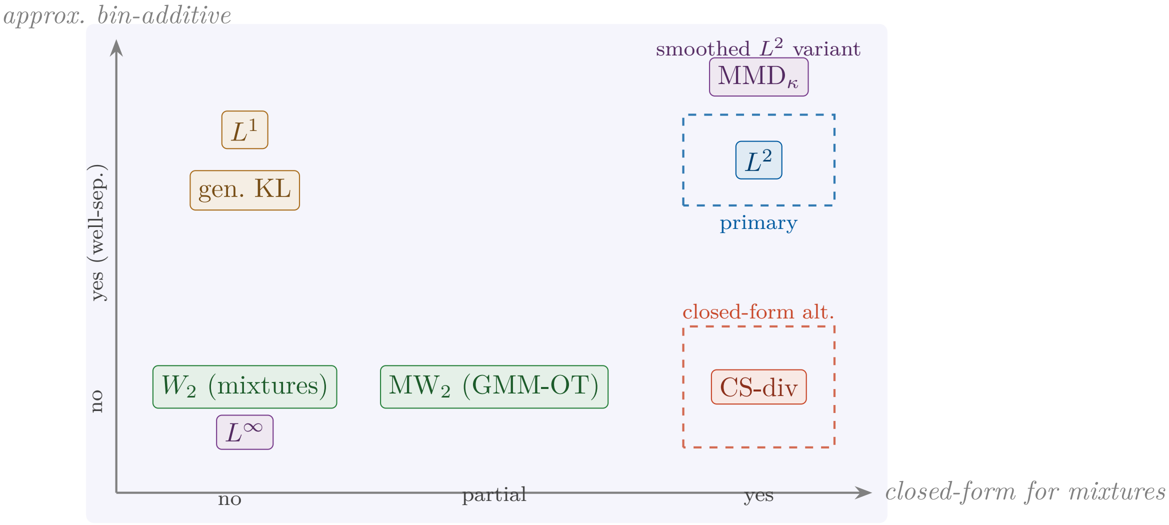

Figure 2: Cost-metric design space, placed as Table 1 classifies the metrics. Horizontal: closed-form between Gaussian mixtures ($L^\infty$ and the mixture $W_2$ have none; MW$_2$ has closed-form costs under a numerically solved coupling). Vertical: degree to which the global cost decomposes additively across bins under a spatial partition (generalised KL and $L^1$ are integrals of a pointwise integrand and so approximately additive for well-separated bins; the transport and $L^\infty$ costs are not). The recommended primary objective ($L^2$) sits in the upper right with Gaussian-kernel MMD, its smoothed variant, beside it; the recommended closed-form alternative (Cauchy–Schwarz divergence) is closed-form but not bin-additive.

partitions independently of an amplitude rescaling of $g$, and it sits naturally on a logarithmic scale that rewards proportional approximations.

- **Proposed audit metrics: $L^\infty$ and generalised KL.** $L^\infty$ would provide the same worst-case voxel guarantee that the contribution-based cull uses, enabling apples-to-apples comparison. Generalised KL would provide a complementary view that emphasises mass.

In the rest of the document, all bin-merge formulas and partition algorithms are derived in $L^2$, with side-by-side closed-form treatments in CS-divergence where they differ. The experiments of Section 9 report the $L^2$ quantities (the squared-error decomposition in Experiment A, `rel_l2` in Experiments B and C); $D_{\mathrm{CS}}$ is recorded in the deposited experiment records of Experiments B and C; the audit metrics above are proposed and are not measured in this document.

### 4 The $K$-wise Merge Subproblem

We now generalise the pairwise ($K = 2$) moment-matched merge of [18] to a bin of arbitrary size, then derive the $L^2$-optimal amplitude. The closed-form bin merge under Cauchy–Schwarz divergence is sketched.

#### 4.1 Moment-matched shape parameters

Treating $\sum_{i\in S_j} \phi_i$ as an unnormalised density and matching its zeroth, first, and second moments yields a single representative Gaussian.

**Proposition 4.1** ($K$-wise moment-matched parameters)**.** *For a bin $S_j$ of arbitrary size, the moment-matched single Gaussian $\bar{\phi}_j(\boldsymbol{x}) = \bar{a}_j \exp(-\frac{1}{2}(\boldsymbol{x}-\bar{\boldsymbol{\mu}}_j)^\top \bar{\boldsymbol{\Sigma}}_j^{-1}(\boldsymbol{x}-\bar{\boldsymbol{\mu}}_j))$ has shape parameters*

$$\bar{\boldsymbol{\mu}}_j = \sum_{i\in S_j} w_i^{(j)}\, \boldsymbol{\mu}_i, \tag{6}$$

$$\bar{\boldsymbol{\Sigma}}_j = \underbrace{\sum_{i\in S_j} w_i^{(j)}\, \boldsymbol{\Sigma}_i}_{\substack{\text{within-component}\\ \text{covariance}}} + \underbrace{\sum_{i\in S_j} w_i^{(j)}\, (\boldsymbol{\mu}_i - \bar{\boldsymbol{\mu}}_j)(\boldsymbol{\mu}_i - \bar{\boldsymbol{\mu}}_j)^\top}_{\substack{\text{between-component scatter}\\ \text{(within the bin)}}}, \tag{7}$$

*where the weights $w_i^{(j)}$ are defined by (4). Both sums run over the members of the single bin $S_j$.*

*Proof.* The first equality is the linearity of expectation under the mass-weighted distribution on $S_j$. The second is the law of total variance, which decomposes the second central moment of a mixture into the mass-weighted average of the component covariances plus the scatter of the component centres around the bin mean. □

*Remark* 4.2 (Recovery of the pairwise companion result). Setting $|S_j| = 2$ in (6)–(7) recovers the pairwise formulas of [18] verbatim; $w_1 + w_2 = 1$ and the between-component scatter becomes the rank-1 term $w_1 w_2(\boldsymbol{\mu}_1 - \boldsymbol{\mu}_2)(\boldsymbol{\mu}_1 - \boldsymbol{\mu}_2)^\top$.

#### 4.2 $L^2$-optimal amplitude

With shape parameters fixed, the residual is quadratic in the bin amplitude $\bar{a}_j$, so the $L^2$-optimal amplitude has a closed form. Recall from Section 2 that $\tau_j$ is the unit-amplitude template of bin $j$, the moment-matched Gaussian shape with amplitude one.

**Proposition 4.3** ($L^2$-optimal bin amplitude)**.** *The amplitude minimising $\|f_{S_j} - \bar{a}_j\,\tau_j\|_{L^2}^2$ is*

$$\bar{a}_j^\star \;=\; \frac{\langle f_{S_j}, \tau_j\rangle_{L^2}}{\|\tau_j\|_{L^2}^2} \;=\; \frac{\sum_{i\in S_j}\langle \phi_i, \tau_j\rangle_{L^2}}{\pi^{D/2}\,|\bar{\boldsymbol{\Sigma}}_j|^{1/2}}. \tag{8}$$

*The corresponding minimum bin error is the projection residual*

$$E_j^\star \;=\; \|f_{S_j}\|_{L^2}^2 - \frac{|\langle f_{S_j}, \tau_j\rangle_{L^2}|^2}{\|\tau_j\|_{L^2}^2}, \tag{9}$$

*expressible entirely in terms of the kernel (3).*

*Proof.* Identical to the pairwise derivation in [18]; quadratic-in-$\bar{a}_j$ minimisation extends unchanged from $|S_j| = 2$ to arbitrary $|S_j|$. □

**Corollary 4.4** (Mergeability score)**.** *The relative bin error $\varepsilon_j = E_j^\star/\|f_{S_j}\|^2$ equals $\sin^2\alpha_j$, where $\alpha_j$ is the angle in $L^2$ between the bin function $f_{S_j}$ and the template $\tau_j$.*

Figure 3 works the merge through on a one-dimensional bin of three members: the moment-matched template, its $L^2$-optimal and mass-conserving amplitudes, the residual each leaves, and the inflated representative that the shipped recipe writes (Section 8.2). Algorithm 1 collects the

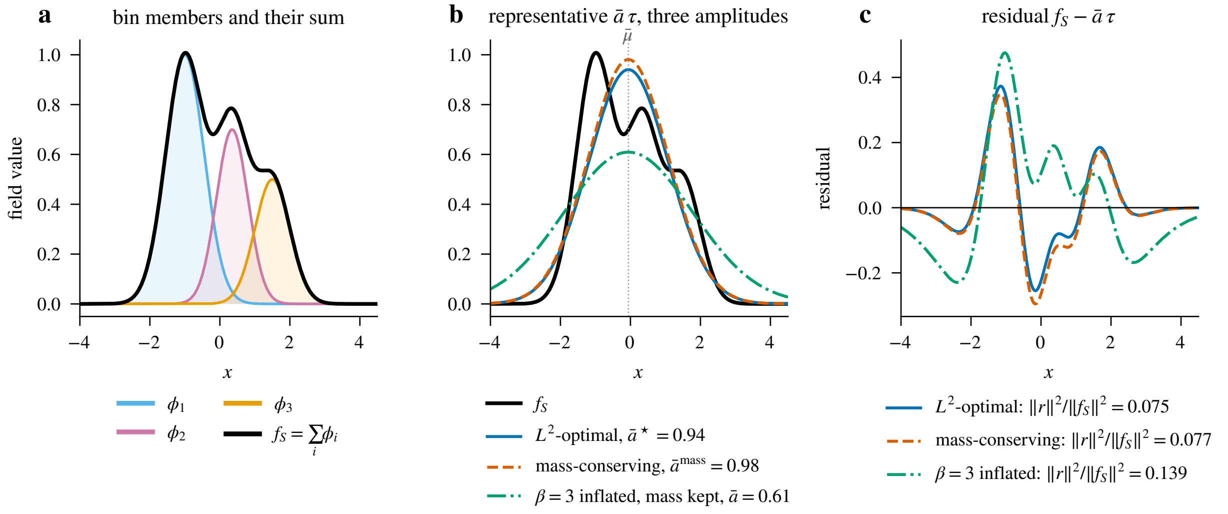

Figure 3: A one-dimensional bin merge, idealised (three members with chosen parameters, no data). (a) The members $\phi_i$ and their sum $f_S$. (b) The moment-matched template $\tau$ of Proposition 4.1 at the $L^2$-optimal amplitude $\bar{a}^\star$ of Proposition 4.3, at the mass-conserving amplitude $\bar{a}^{\text{mass}}$ of Section 4.3, and with the scatter term inflated by $\beta = 3$ at the amplitude that keeps the bin's mass, which is what the per-level mass conservation of Section 8.2 leaves for a level of one bin. The mass-conserving representative is the tallest of the three; the $L^2$-optimal one sits just below it and carries less mass than the bin; the inflated one is the lowest and the widest. (c) The residuals $f_S - \bar{a}\,\tau$ of the three choices; the $L^2$-optimal one has the smallest energy by construction, and the legend gives each energy as a fraction of $\|f_S\|^2$.

merge as pseudocode: masses and weights, the two moments, and the $L^2$-optimal amplitude, in that order.

**Algorithm 1:** $K$-wise bin merge under $L^2$.

**Input:** Bin of splats $S_j = \{\phi_i\}_{i \in S_j}$.
**Output:** Representative splat $\bar{\phi}_j$ with parameters $(\bar{a}_j, \bar{\boldsymbol{\mu}}_j, \bar{\boldsymbol{\Sigma}}_j)$.

Compute masses $m_i = a_i (2\pi)^{D/2} |\boldsymbol{\Sigma}_i|^{1/2}$ and weights $w_i^{(j)}$ via (4)
Compute $\bar{\boldsymbol{\mu}}_j$ via (6)
Compute $\bar{\boldsymbol{\Sigma}}_j$ via (7)
Compute $\bar{a}_j^\star$ via (8) using kernel (3)
**return** $\bar{\phi}_j = \bar{a}_j^\star \exp(-\frac{1}{2}(\boldsymbol{x} - \bar{\boldsymbol{\mu}}_j)^\top \bar{\boldsymbol{\Sigma}}_j^{-1} (\boldsymbol{x} - \bar{\boldsymbol{\mu}}_j))$

#### 4.3 Cauchy–Schwarz bin merge

With shape parameters fixed by moment matching, the CS-divergence $D_{\text{CS}}(f_{S_j}, \bar{a}_j \tau_j)$ is independent of $\bar{a}_j$ (since CS-divergence is scale-invariant in its second argument): the optimal bin amplitude under CS-divergence is undetermined and must be set externally (e.g. by mass conservation). The bin merge under CS-divergence is therefore the same as the moment-matched merge with mass-conservation amplitude $\bar{a}_j^{\text{mass}} = m_j^{\text{tot}} / [(2\pi)^{D/2} \, |\bar{\boldsymbol{\Sigma}}_j|^{1/2}]$, with residual measured by (5).

*Remark* 4.5 (Why CS-divergence still matters). Although the bin merge under CS coincides numerically with mass-conserved moment matching, the divergence value itself differs, and ranks candidate partitions on a log scale that is invariant to a global amplitude rescaling of $g$. That invariance is useful for partition selection in the presence of amplitude noise.

##### 4.4 $W_2$ component-barycentre surrogate (sketched)

For mass-normalised bin members $\mathcal{N}_i = \mathcal{N}(\boldsymbol{\mu}_i, \boldsymbol{\Sigma}_i)$ with weights $w_i^{(j)}$, the $W_2$-flavoured surrogate representative is the Gaussian whose mean is the weighted centroid and whose covariance is the Bures (Wasserstein) barycentre of the member covariances. The Bures barycentre is the covariance minimising the weighted sum of squared $W_2$ distances to the members; it is computed by the Álvarez-Esteban fixed-point iteration [1, 2]. The barycentre minimises the *component-wise* objective $\sum_i w_i^{(j)} W_2^2(\mathcal{N}_i, q)$ over single Gaussians $q$ (the same restriction to component-to-component transport as $\mathrm{MW}_2$). It is not the $W_2$ projection of the bin *mixture* $\sum_i w_i^{(j)} \mathcal{N}_i$ onto a single Gaussian.

That projection minimises $W_2^2(\sum_i w_i^{(j)} \mathcal{N}_i, q)$, has no closed form, and has a different minimiser as soon as the means differ. For two equally weighted, well-separated members the barycentre keeps the common member variance, whereas the mixture projection widens to cover both peaks.

As a merge rule the barycentre also behaves differently from moment matching. For identical covariances it returns that common covariance regardless of the means: for two equally weighted members $\mathcal{N}(\pm d, \sigma^2)$ the barycentre variance is $\sigma^2$, while the moment-matched variance of (7) is $\sigma^2 + d^2$. The between-component scatter makes the moment-matched matrix the larger one in the Loewner order (the partial order on symmetric matrices in which $A \succeq B$ means $A - B$ is positive semi-definite). When the covariances differ as well, the two are in general different matrices.

#### 5 The Partition / Assignment Subproblem

With a per-bin merge rule fixed (moment-matched shape and $L^2$-optimal amplitude, Section 4), the remaining problem is to find the partition $\mathcal{P}$ that minimises the global error.

##### 5.1 The cost of a partition

Let $r_j = f_{S_j} - \bar{\phi}_j$ be the residual of bin $j$. Then $f - g = \sum_j r_j$ and

$$\boxed{\|f - g\|_{L^2}^2 = \underbrace{\sum_j \|r_j\|_{L^2}^2}_{\text{per-bin residual energy}} + \underbrace{2\sum_{j<k} \langle r_j, r_k \rangle_{L^2}}_{\text{cross-bin interference}} .} \tag{10}$$

The cross-bin term *is fully closed-form* via the kernel (3): each $r_j$ is a sum of $|S_j| + 1$ Gaussians (the $|S_j|$ original splats plus the negative representative), so $\langle r_j, r_k \rangle$ is a finite sum of pairwise Gaussian inner products. The closed form makes the partition cost exactly evaluable, but (10) is *not* bin-additive in general: the cross-bin term may be of either sign.

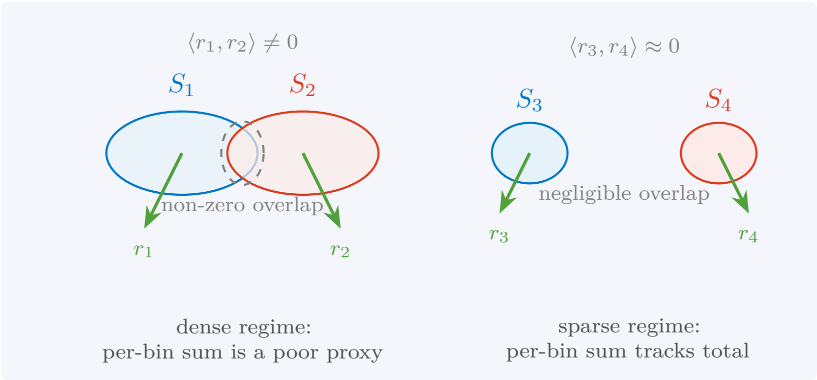

Figure 4: Cross-bin interference in the $L^2$ partition cost. When bins overlap spatially the residuals $r_j, r_k$ have non-zero $L^2$ inner product and the per-bin sum is not equal to the global error. When bins are well-separated the inner product vanishes and the per-bin sum becomes a faithful surrogate; this is the regime in which spatial clustering operates (Section 9, Experiment A).

##### 5.2 Greedy hierarchical partition (Runnalls)

A simple bottom-up algorithm builds the partition by repeated pairwise merges. Initialise each splat as its own cluster. Repeatedly merge the pair of clusters with smallest pairwise merge cost (using Proposition 4.3 on the union) until $M$ clusters remain. The procedure is the Luxar analogue of the Runnalls algorithm [22]; Algorithm 2 states it. We count cost in kernel evaluations (3). The naive implementation (rescan all pairs after every merge, as in Algorithm 2 and the analysis code) scores $O(N^3)$ candidate pairs; a heap with lazy recomputation reduces this to $O(N^2 \log N)$ candidates. Each candidate, however, is scored on the *union of original members* with Algorithm 1, which needs the bin's Gram sum and the overlap of every member with the moment-matched template, that is $O(n_b^2)$ kernel evaluations for a candidate bin of $n_b$ members. The cost therefore grows with the bins, and an unbalanced merge chain reaches $O(N^4)$ kernel evaluations in total. The $O(N^2 \log N)$ figure holds only under a bounded candidate-bin size, or when a candidate pair is scored by the constant-cost merge of its two already-merged *representatives* instead of the original members, which is the approximation the shipped Luxar greedy uses on a sparse neighbour graph (Section 8.3). The cosine ranking proxy of [18] (an $O(1)$ approximation to the pairwise merge cost using only $\langle \phi_i, \phi_j \rangle$) could accelerate the nearest-neighbour search; neither the reference implementation of Section 9 nor the shipped code uses it.

---

**Algorithm 2:** Greedy hierarchical partition.

**Input:** Splats $\{\phi_i\}_{i=1}^N$, target count $M$.
**Output:** Partition $S_1, \dots, S_M$.

Initialise clusters $\mathcal{C} \leftarrow \{\{i\} : i = 1, \dots, N\}$
**while** $|\mathcal{C}| > M$ **do**
  `// merge cost:  E* of the union, Algorithm 1`
  $(A, B) \leftarrow \arg\min_{A', B' \in \mathcal{C},\, A' \neq B'} \text{merge\_cost}(A' \cup B')$
  $\mathcal{C} \leftarrow (\mathcal{C} \setminus \{A, B\}) \cup \{A \cup B\}$
**return** $\mathcal{C}$

---

##### 5.3 Cost-increment relocation

The substitutive-LOD analogue of $k$-means clustering [17] is the sequential relocation of Algorithm 3. Fix the merge rule of Section 4: the shape of $\bar{\phi}_j$ is the moment-matched one, its amplitude the $L^2$-optimal one, so the bin cost $E^\star_j(S_j)$ of Proposition 4.3 is a deterministic function of the bin's membership alone. The algorithm then visits the splats one at a time and, for each, evaluates the exact change of the two affected bin costs, $\Delta_b(i) = [E^\star_a(S_a \setminus \{i\}) + E^\star_b(S_b \cup \{i\})] - [E^\star_a(S_a) + E^\star_b(S_b)]$, moving the splat only if the best candidate lowers the sum. The convention $E^\star(\emptyset) = 0$ covers a splat that leaves a singleton bin. In the spirit of Bregman clustering [4], the dissimilarity is the *cost increment*, not a pre-defined distance. Because every accepted move lowers $\sum_j E^\star_j(S_j)$ by construction, the per-bin sum surrogate is non-increasing; Section 5.1 relates it to the global $L^2$ error up to the cross-bin term, which Experiment A (Section 9.1) finds to be small in the median, though not uniformly, for spatially decoupled partitions.

**Algorithm 3:** Cost-increment relocation.

**Input:** Initial partition $S_1, \ldots, S_M$, splats $\{\phi_i\}$; convention $E^\star(\emptyset) = 0$.
**Output:** Refined partition with at most $M$ non-empty bins.

**repeat**
  **foreach** *splat* $\phi_i$ *with current bin* $a = \mathcal{P}(i)$ **do**
    **foreach** *candidate bin* $b \neq a$ **do**
      `// change of the per-bin sum from moving` $i$ `to` $b$ `(Algorithm 1)`
      $\Delta_b(i) \leftarrow [E^\star_a(S_a \setminus \{i\}) + E^\star_b(S_b \cup \{i\})] - [E^\star_a(S_a) + E^\star_b(S_b)]$
    $b^\star \leftarrow \arg\min_b \Delta_b(i)$
    **if** $\Delta_{b^\star}(i) < 0$ **then**
      Move $\phi_i$: $S_a \leftarrow S_a \setminus \{i\}$, $S_{b^\star} \leftarrow S_{b^\star} \cup \{i\}$
**until** *no splat moves (or a pass budget is exhausted)*

Termination follows from the finiteness of the set of partitions together with the strict decrease on every accepted move (a decreasing sequence bounded below would not suffice on its own); the endpoint is a one-move local optimum. A singleton bin may be emptied (its cost is 0 before and after), so the algorithm has at-most-$M$ semantics; the analysis code compacts the labels afterwards and evaluates whatever number of non-empty bins remains. Experiments B and C run five passes; on the full fit of Experiment C each splat's candidate bins are the 12 whose centroids are nearest other than its own, the shipped count.

The guarantee concerns the relocation on the moment-matched surrogate, not a Lloyd or Bregman alternation whose update step minimises each term: moment matching fixes the shape by its moments, not by minimising $L^2$ (or CS) over the shape, so the representative is $L^2$-optimal only in its amplitude (Proposition 4.3). For two unit peaks at $\pm 2$ with unit variance the moment-matched template (variance 5) has $L^2$-optimal-amplitude residual 0.897, whereas a template of variance 8.125 with re-optimised amplitude reaches 0.752; the shape-optimal single Gaussian is not the moment-matched one. A true alternation would need the $L^2$-optimal shape in its update step, which has no closed form. Figure 5a draws this example; panel b draws the Eckart–Young

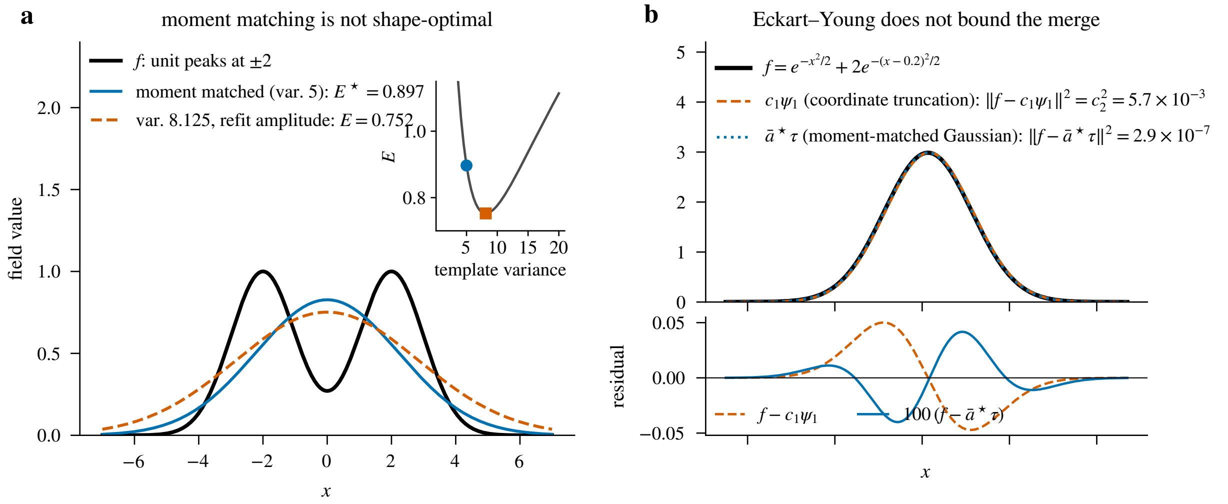

Figure 5: Two counterexamples, drawn from the numbers in the text. (a) Moment matching is not shape-optimal: for two unit peaks at $\pm 2$ with unit variance, the moment-matched template (variance 5) with $L^2$-optimal amplitude leaves a squared residual of 0.897, while a template of variance 8.125 with re-optimised amplitude leaves 0.752; the inset traces the residual against the template variance. (b) The Eckart–Young argument of Section 6 does not bound the merge: for $f = e^{-x^2/2} + 2e^{-(x-0.2)^2/2}$ the discarded Gram eigen-coordinate carries $c_2^2 \approx 5.7 \times 10^{-3}$, but that is the error of the best *coordinate truncation* $c_1\psi_1$, which is not a Gaussian; the single moment-matched Gaussian with $L^2$-optimal amplitude leaves $2.9 \times 10^{-7}$, four orders of magnitude less. Both residuals are drawn, the Gaussian one magnified $100\times$.

example of Section 6.

##### 5.4 Joint relaxation

Relaxing the partition constraint, treat the $M$ representatives $\{\bar{\phi}_j\}$ as $M$ free Gaussians and minimise $\|f - g\|_{L^2}^2$ directly by gradient descent over $\{(\bar{a}_j, \bar{\boldsymbol{\mu}}_j, \bar{\boldsymbol{\Sigma}}_j)\}$. The gradient is closed-form in the kernel (3). Relaxation loses the partition structure: each $\bar{\phi}_j$ no longer represents a specific subset of original splats. It would provide a reference value, though none of the experiments in Section 9 runs it. Any partition-and-merge result with $M$ bins can do no better than the joint-relaxation *global* optimum, since the latter solves the strictly less constrained problem "best $M$ Gaussians" over all of $\mathbb{R}_{>0}^M \times \mathbb{R}^{DM} \times (\mathrm{SPD}_D)^M$, where $\mathrm{SPD}_D$ is the cone of symmetric positive definite $D \times D$ matrices. That global optimum is not what a gradient-descent run returns, however: the objective is non-convex, so a numerical solution is a local minimum, i.e. an *upper* bound on the relaxed optimum, and a partition-and-merge result may in principle undercut it. We therefore treat the joint-relaxation error as an empirical reference value, not as a guaranteed lower bound on the partition-and-merge error; Section 6 explains why no analytical lower bound is claimed either.

##### 5.5 Recommended recipe

We recommend: **(i)** $k$-means on the splat centres (a $k$-means++ start [3]) to build a cheap warm-start partition; **(ii)** a few passes of the cost-increment relocation of Algorithm 3 to refine it; optionally **(iii)** a joint-relaxation polish. Steps (i) and (ii) are exactly the $k$-means + relocation arm of Experiments B and C (Section 9); greedy hierarchical (Algorithm 2) is run there as the quality reference at small $N$, not as the warm start (no experiment refines a greedy partition). Two optional refinement rounds of the Luxar implementation play the role of (iii). An $L^2$ *refit* optimises each merged level (by Adam gradient steps) against its fine-input mixture under a *sparse* form of the mixture $L^2$ objective: $\|g\|^2 - 2\langle f, g\rangle$ summed over neighbour lists of 16 fine-to-coarse and 64 coarse-to-coarse pairs within 3 and $5\sigma$, rebuilt every 20 steps. Total mass is pinned and any *barrier dimension* (a centre coordinate such as time or channel across which coarsening never merges splats) is frozen. Its acceptance guard re-evaluates that sparse objective on freshly rebuilt pair lists and keeps the seed unless a candidate is lower, so the result is never worse than the merge *in that sparse metric*, not in the full Gram sum. A *volume refit* instead warm-starts a re-fit of each coarse level against the source volume itself and keeps whichever of seed and candidate has the lower full-grid mean squared error. Neither refit is measured in this document; the experiments below evaluate the pure merge.

#### 6 On Lower Bounds

The distance between any substitutive-LOD method and the optimum is not known. We claim no analytical lower bound on the error of approximating $f$ by a sum of $M$ Gaussians. A tempting argument expands $f$ in the eigenbasis of the Gram matrix $\boldsymbol{G}$ and invokes Eckart–Young [8] to discard the $N - M$ smallest feature-space coordinates. It does not supply a bound: Eckart–Young bounds the error of the best *fixed* $M$-dimensional coordinate truncation of $f$, whereas the $M$ representatives are free Gaussians whose span is not a coordinate subspace of $\mathrm{span}\{\phi_1, \ldots, \phi_N\}$ and need not lie in that span at all. A one-dimensional example makes the failure explicit. Write $\boldsymbol{G} = \sum_k \lambda_k \boldsymbol{v}_k \boldsymbol{v}_k^\top$ for the eigen-decomposition of the Gram matrix; the functions $\psi_k = \lambda_k^{-1/2} \sum_i (\boldsymbol{v}_k)_i \, \phi_i$ are $L^2$-orthonormal and $f = \sum_k c_k \psi_k$ with coordinates $c_k = \sqrt{\lambda_k}\,(\boldsymbol{v}_k^\top \mathbf{1})$. For $f(x) = e^{-x^2/2} + 2\,e^{-(x-0.2)^2/2}$ ($N = 2$, $M = 1$) the discarded eigen-coordinate carries $c_2^2 \approx 5.7 \times 10^{-3}$, yet the single moment-matched Gaussian with $L^2$-optimal amplitude of Section 4 (mean 2/15, variance 1.0089) reaches $\|f - g\|_{L^2}^2 \approx 2.9 \times 10^{-7}$, four orders of magnitude below the purported bound (Fig. 5b). The only lower bound we rely on is the trivial one of Section 5.4: the global optimum of the joint relaxation (the best $M$ free Gaussians) is no larger than any partition-and-merge error. As discussed there, a numerical solution of that relaxation is a local minimum and hence an empirical reference value, not a certified bound, and the per-bin mergeability score of Corollary 4.4 remains a per-bin diagnostic, not a statement about the global optimum.

#### 7 Hierarchical Multi-level LOD

A single application of the partition-and-merge operator $\mathcal{R}_K$ produces a level with $M_1 = \lceil N/K \rceil$ splats. Iterating yields a hierarchy

$$N \xrightarrow{\mathcal{R}_K} M_1 \xrightarrow{\mathcal{R}_K} M_2 \xrightarrow{\mathcal{R}_K} \cdots \xrightarrow{\mathcal{R}_K} M_L, \qquad M_\ell = \lceil M_{\ell-1}/K \rceil = \lceil N/K^\ell \rceil, \tag{11}$$

where the collapse of the repeated ceilings holds for integer $K \geq 2$ (which the implementation enforces; for a non-integer $K$ the recurrence is the definition). The level-$\ell$ approximation $g_\ell$ therefore has a *splat-count* reduction $N/M_\ell$ of $K^\ell$ up to rounding; this is a count ratio, not the stored-byte compression ratio of the main text. Algorithm 4 states the construction: each level is built from the one before it by the partition step of Section 5 and the merge of Algorithm 1.

**Algorithm 4:** Build substitutive-LOD hierarchy.

**Input:** Splats $\{\phi_i\}_{i=1}^N$, branching factor $K$, depth $L$.
**Output:** Levels $g_0 = f, g_1, \ldots, g_L$ where $g_\ell$ has $M_\ell$ splats.

$g_0 \leftarrow f$, $M_0 \leftarrow N$
**for** $\ell = 1$ **to** $L$ **do**
    $M_\ell \leftarrow \lceil M_{\ell-1}/K \rceil$
    Partition the $M_{\ell-1}$ splats of $g_{\ell-1}$ into $M_\ell$ bins by $k$-means on the centres, then refine with a few passes of Algorithm 3 (Algorithm 2 instead when $M_{\ell-1}$ is small)
    $g_\ell \leftarrow$ sum of the $M_\ell$ bin representatives (each from Algorithm 1)
**return** $g_0, g_1, \ldots, g_L$

##### 7.1 Error compounding across levels

Because each level is built from the previous level and not from $f$, the error of a deep level is inherited across every step; this section bounds how it accumulates. In $L^2$, the cross-level error is sub-additive but not additive. Write $g_0 = f$ and let $\eta_{\ell'} = \|g_{\ell'-1} - g_{\ell'}\|_{L^2}/\|g_{\ell'-1}\|_{L^2}$ be the relative error of level $\ell'$ with respect to its *own input* $g_{\ell'-1}$. The quantity $\eta_{\ell'}$ is a norm ratio (the `rel_l2` of Section 9 for one level), not the squared per-bin score $\varepsilon_j$ of Corollary 4.4: for a level consisting of a single bin, $\eta^2 = \varepsilon$. Since $f - g_\ell = \sum_{\ell'=1}^{\ell}(g_{\ell'-1} - g_{\ell'})$, the triangle inequality on the $L^2$ norm gives

$$\|f - g_\ell\|_{L^2} \;\leq\; \sum_{\ell'=1}^{\ell} \eta_{\ell'}\,\|g_{\ell'-1}\|_{L^2}, \qquad \|f - g_\ell\|_{L^2}^2 \;\leq\; \Big(\sum_{\ell'=1}^{\ell} \eta_{\ell'}\,\|g_{\ell'-1}\|_{L^2}\Big)^2. \tag{12}$$

The intermediate norms cannot be dropped: a per-bin $L^2$ merge does not make the norm of the *sum* non-increasing, because the cross-bin terms of Section 5.1 can push $\|g_{\ell'}\|_{L^2}$ above $\|g_{\ell'-1}\|_{L^2}$ (in Experiment A, Section 9.1, 67 of the 120 trials have $\|g\|_{L^2} > \|f\|_{L^2}$). Bounding each intermediate norm by the same triangle inequality, $\|g_{\ell'}\|_{L^2} \leq (1 + \eta_{\ell'})\,\|g_{\ell'-1}\|_{L^2}$, turns (12) into the product bound

$$\frac{\|f - g_\ell\|_{L^2}}{\|f\|_{L^2}} \;\leq\; \prod_{\ell'=1}^{\ell}(1 + \eta_{\ell'}) - 1, \tag{13}$$

which is what controls a deep hierarchy. Per-level errors that are individually small do not keep the intermediate norms near $\|f\|$ unless their *sum* is small ($\eta_{\ell'} = \delta$ at every level gives $\|g_\ell\| = (1+\delta)^\ell \|f\|$ in the worst case). Only under that cumulative control, $\sum_{\ell'} \eta_{\ell'} \ll 1$ (or at fixed modest depth with each $\eta_{\ell'} \ll 1$), does (13) reduce to the familiar first-order form $\|f - g_\ell\|_{L^2}^2 \lesssim (\sum_{\ell'} \eta_{\ell'})^2\,\|f\|_{L^2}^2$. Equality in the triangle inequality is approached when consecutive level differences are aligned in $L^2$. Figure 6 draws the bound for the shipped defaults, three coarser

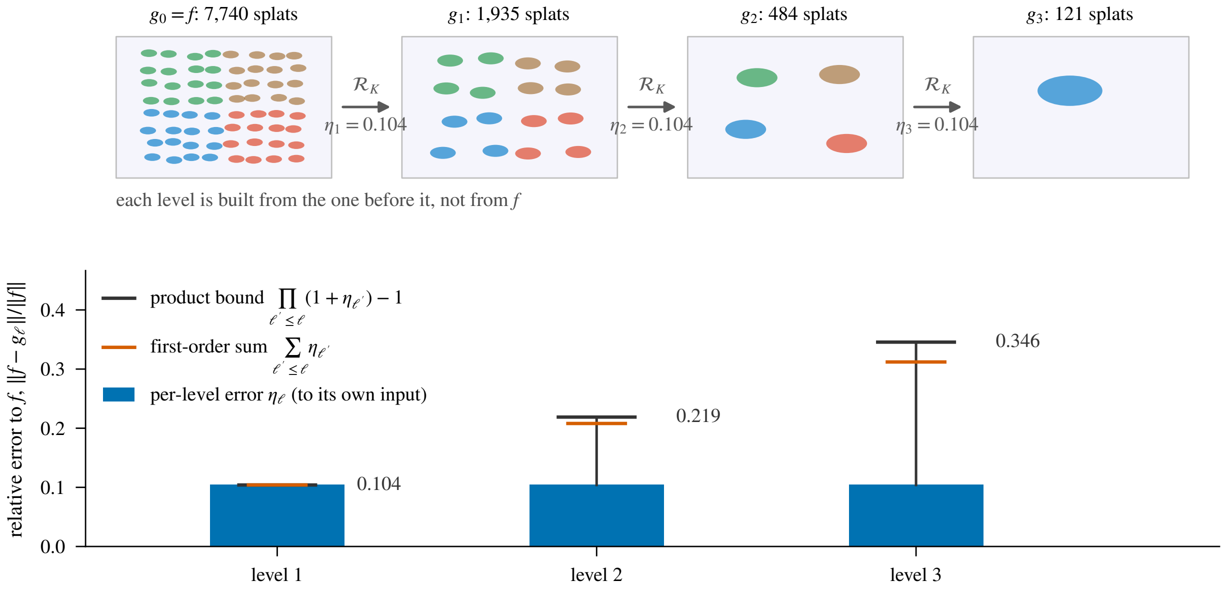

Figure 6: Error compounding across a three-level hierarchy, schematic. Top: the construction of Fig. 1 iterated, from $N = 7{,}740$ splats at the shipped $K = 4$ (Section 8.3), each level built from the one before it, so that its error to $f$ inherits every earlier step. Bottom: the relative error to $f$ that (13) allows at each level when every level has the same error $\eta = 0.104$ relative to its own input (the $k$-means + relocation error of Experiment C at $K = 4$, Table 3). The bar is the per-level error $\eta$, the whisker reaches the product bound $\prod(1+\eta) - 1$ and the tick marks the first-order sum $\ell\,\eta$; the gap between the two grows with depth.

levels at $K = 4$, with the Experiment C error of $k$-means + relocation at $K = 4$ standing in for every $\eta_{\ell'}$.

##### 7.2 Connection to existing 3D Gaussian-Splatting LOD methods

Several 3D Gaussian-Splatting LOD methods have appeared since the introduction of 3DGS [15]. The closest precedent is Hierarchical 3DGS [16]: it builds a bounding volume hierarchy by top-down spatial partitioning and then *merges* bottom-up, each interior node carrying a single Gaussian whose mean and covariance are the weighted first and second moments of its children (with weights derived from opacity and projected area). The interior nodes are then optimised against rendered images. The moment merge is the one of Proposition 4.1; the differences are the weights (integral mass here, opacity-times-area there), the amplitude (the fixed-shape $L^2$-optimal amplitude of Proposition 4.3 here, opacity heuristics followed by image-space optimisation there) and the objective (the scalar field $f$ itself, not rendered views). Octree-GS [21] attaches anchors to octree levels and selects a level per view. Mini-Splatting [10] contains an importance-sampled simplification stage inside a larger densification/re-initialisation pipeline; LightGaussian [9] and Reduced 3DGS [20] prune by importance scores (with distillation and quantisation). The substitutive-LOD framework of this document is the merging-based analogue with an analytic objective, complementary to the pruning-based methods, and may be combined with them in a hybrid pipeline (prune away low-importance splats, then substitutively merge the survivors into a coarse level).

### 8 Connection to Luxar Machinery

#### 8.1 How a hierarchy is stored and displayed

Luxar stores a fitted splat set as a tree of nodes. A *leaf* holds the splat arrays themselves (centres, amplitudes and Cholesky factors); a group holds children. A substitutive hierarchy is a ***kind=lod** group* group whose children are the levels $g_L, \ldots, g_1, g_0$, stored coarsest first, and the viewer draws exactly one of them at a time. Each level is one leaf: its centres are the $\bar{\boldsymbol{\mu}}_j$, its Cholesky factors those of $\bar{\boldsymbol{\Sigma}}_j$ (stored as a diagonal and an off-diagonal array) and its amplitudes the $\bar{a}_j$. The *recipe* that builds this tree is called `levels`. By default every level also carries an additive streaming *additive (streaming) ladder* (Supplementary Document 9): the level's splats are ordered and cut into *rung (sub-LOD)*s, each stored as its own sub-group, so that a level paints progressively while it downloads. The tree is then a replacement group above per-level leaves above rungs. The ladder can be switched off, in which case each level is one flat leaf. No storage code specific to substitutive LOD is needed beyond the standard leaf writer.

The viewer chooses the level from the screen footprint of the content, so the choice is independent of the splat count and of the display's pixel density. Two rules exist. The primary rule uses a *median footprint* that the builder stamps on every level: the median, over the level's splats, of the geometric mean of the marginal standard deviations $\sigma$ along the non-degenerate axes, together with the list of those axes. When every level of a group carries such a stamp and the stamped axes are the displayed dimensions, the viewer projects the stamped $\sigma$ (scaled by the node's world transform) to logical pixels at the depth of the group's centre. Logical (CSS) pixels are the page's pixel unit, which does not change with the display's pixel density. The viewer then displays the *coarsest* level whose projected median $\sigma$ is at most 1.5 logical pixels. Splats draw to about $3\sigma$, so a typical blob of the chosen level is about 9 pixels across; equivalently, the finest level is shown once even its own splats project larger than that. Refinements are immediate. A step to a coarser level requires the coarser level's footprint to drop below 1.35 pixels, a 10 % hysteresis, and a session-wide detail bias divides the limit by the square root of the bias. Figure 7 draws the rule.

The second rule is a fallback for a group whose stamps are absent or were measured on axes other than the displayed ones. The builder also writes a *screen-area* threshold per level: the fraction of the viewport that the node's projected bounding box must occupy for that level to be eligible. For a whole-object ladder the thresholds are $[0, \ldots, \frac{1}{8}, \frac{1}{4}, \frac{1}{2}]$, so the finest level is eligible while the object occupies at least half the screen and each halving of the occupied area admits one level coarser. A ladder built inside one part of a ***kind=partition** group* is anchored at 1 instead, because that part alone rarely fills the screen. The finest eligible level is chosen, with a hysteresis band of 10 % of the gap to the adjacent threshold. A store whose thresholds were written against the diagonal coverage metric of a coverage-selector ladder takes the same threshold path with that metric.

#### 8.2 Shipped merge defaults

The as-shipped coarse level deviates from the pure moment-matched representative and $L^2$-optimal amplitude derived above in two ways that are on by default. First, a coverage-inflation factor $\beta = 3$

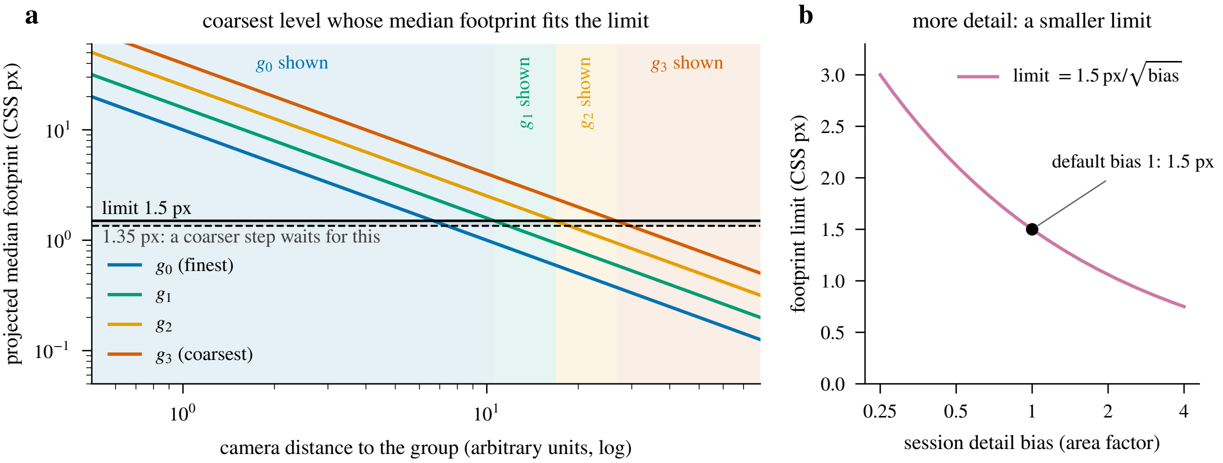

Figure 7: The viewer's level-selection rule, schematic. (a) Projected median footprint of each level of a group with three coarser levels against camera distance, drawn with the idealised ratio $K^{1/D} = 4^{1/3}$ between consecutive levels. The viewer shows the coarsest level whose footprint is at most 1.5 logical pixels (solid line); the shaded bands name the level shown at each distance. Refinement is immediate, while a step to a coarser level waits until that level's footprint drops below 1.35 pixels (dashed line, a 10 % hysteresis). (b) The limit against the session detail bias: the bias is an area factor, so the limit divides by its square root.

widens the between-component scatter term of (7), $\bar{\boldsymbol{\Sigma}}_j^{\text{out}} = \sum_i w_i \boldsymbol{\Sigma}_i + \beta \sum_i w_i(\boldsymbol{\mu}_i - \bar{\boldsymbol{\mu}}_j)(\boldsymbol{\mu}_i - \bar{\boldsymbol{\mu}}_j)^\top$, and rescales the amplitude by $|\bar{\boldsymbol{\Sigma}}_j|^{1/2}/|\bar{\boldsymbol{\Sigma}}_j^{\text{out}}|^{1/2}$ so that the representative's integral mass is unchanged. The value $\beta = 3$ is a calibrated smoothing heuristic: it turns the $d^2/12$ scatter of members spread uniformly over a bin of width $d$, the pitch of the bin tiling, into $(d/2)^2$ (two point members spaced $d$ apart would instead scatter by $d^2/4$), and it reduces the axis-aligned grid ripple that pure moment matching shows at coarse levels. It does not make neighbouring coarse splats sum exactly flat (a regular lattice of Gaussians at $\sigma = d/2$ retains a relative ripple of $2\exp(-\pi^2/2) \approx 1.4\,\%$), and $\beta = 1$ recovers the derivation here.

Second, per-level mass conservation applies one global amplitude factor per barrier group so that the level's integrated mass over the coarsened dimensions, $\sum_j \bar{a}_j |\bar{\boldsymbol{\Sigma}}_j|^{1/2}$ (the $(2\pi)^{D/2}$ factor cancels), equals that of its fine input. The factor is applied only when it lies in $[0.1, 10]$; otherwise the per-bin amplitudes are kept and a warning is printed. What this pins is the integrated mass, the DC component of the field (its zero-frequency term), not the sum of peak amplitudes and not pointwise brightness: two unit-mass Gaussians of variance 1 and 4 differ two-fold in peak height, so a level switch can still change local intensity and contrast. Conservation removes the DC brightness jump, which is what is visible as a pop in additive rendering, and the mass loss of the $L^2$-optimal amplitude. That loss is typical though not universal: on the Experiment C $k$-means + relocation bins at $K = 4$ (Section 9.3) the $L^2$-optimal amplitude is below the mass-conserving one in 79.8 % of the bins with at least two members (in a single-member bin the two amplitudes coincide), with a median ratio of 0.986 and a range of 0.925 to 1.048; a dominant member with a faint distant companion gives $\bar{a}^\star > \bar{a}^{\text{mass}}$. The Experiment C numbers in Section 9 were computed at $\beta = 1$ with the $L^2$-optimal amplitude (the pure derivation), not with these shipped defaults; the defaults trade a small increase in `rel_l2` for visibly smoother coarse levels.

#### 8.3 Theory, experiments and the shipped algorithms

The partition algorithms of Section 5 and the experiments of Section 9 use the exact per-bin cost of Proposition 4.3 on small problems. The shipped values quoted in this section were read at luxar commit `677c596f5`. The shipped `levels` recipe replaces each step by a scalable variant, and the experimental conclusions transfer to it only up to the differences that Table 2 lists. The rest of this section explains the three differences that matter most.

The first difference is the warm start. The shipped $k$-means-style methods do not run $k$-means: the splats are sorted along a Morton curve, a space-filling curve that interleaves the bits of the quantised coordinates so that neighbours in space become neighbours in the order, and the order is cut into $M$ contiguous chunks of equal size. The experiments use $k$-means++ [3] on the centres.

The second difference is the acceptance rule of the refinement. The shipped refinement reassigns every splat *synchronously* to the candidate bin whose unit template it projects onto best, $\langle \phi_i, \tau_b \rangle / \|\tau_b\|$, where the candidates are the current bins of its Morton-order neighbours. It then rebuilds the templates and commits the pass only if the total projection energy

$$\Psi \;=\; \sum_b \frac{\langle f_{S_b}, \tau_b \rangle^2}{\|\tau_b\|^2} \tag{14}$$

strictly increases; it stops at the first pass that does not. $\Psi$ is the term of (9) that depends on the templates, but the omitted term $\sum_b \|f_{S_b}\|^2$ also changes with the partition. An increase of $\Psi$ is therefore not the same as a decrease of the per-bin sum $\sum_b E_b^\star$, nor of the global residual; a pass can raise $\Psi$ while raising both. The monotone quantity is $\Psi$, and the guarantee of Algorithm 3 (an exact two-bin cost delta per sequential move) is not the shipped one.

The third difference is the greedy cost model. The shipped greedy keeps one representative per cluster and scores a candidate pair by the moment-matched merge of the two *representatives*, a constant-cost application of Proposition 4.3 to two Gaussians, not by the merge of the union of original members. It considers only pairs on a sparse spatial-hash neighbour graph with at most 8 neighbours per splat, managed by a lazy-deletion heap. Algorithm 2 and the experiments rescore the full union over all pairs.

The quality and timing conclusions of Experiments B and C therefore describe the algorithms as specified in Algorithms 1 to 3; a benchmark of the shipped variants against them is not part of this document.

#### 8.4 Substitutive merging vs. removal-based culling

Luxar implements removal-based decimation as a contribution-based cull. It selects the subset through an $L^\infty$ error budget when the fitted volume is supplied, and through a threshold on each splat's maximum fractional contribution to the predicted volume when it is not. In both cases a binary search then keeps the compounded removal within budget. Experiment C in Section 9 compares substitutive merging against two removal baselines that retain the $M$ splats (the budget of the merging arms) with the largest integral mass $m_i$, one keeping their amplitudes verbatim, one refitting the retained amplitudes jointly by least squares (the $M$-template analogue

Table 2: What this document analyses against what the shipped `levels` recipe runs. Each row names one quantity or step, states the shipped counterpart, and says where the two differ.

| **In this document** | **What ships** | **Where they differ** |
|---|---|---|
| Per-bin merge: moment-matched shape (Proposition 4.1) with $L^2$-optimal amplitude (Proposition 4.3); $\beta = 1$. | Moment-matched shape with the scatter term inflated by $\beta = 3$ and the amplitude rescaled to keep the representative's mass; then one mass-conservation factor per level and barrier group (Section 8.2). | Shape and amplitude both differ; the shipped level trades a small increase in relative $L^2$ error for smoother coarse levels. Experiment C uses the pure derivation. |
| Warm start: $k$-means on the centres with a $k$-means++ start. | The splats are sorted along a Morton space-filling curve (*Morton and Hilbert ordering*) and cut into $M$ contiguous chunks of equal size. | Geometric clustering with data-driven bin sizes against space-filling-curve chunking with equal bin sizes; no $k$-means is run. |
| Refinement (Algorithm 3): sequential single-splat moves, each accepted on the exact change of two bin costs; monotone in the per-bin sum $\sum_b E_b^\star$. | Every splat is reassigned at once to the candidate bin (the current bins of its Morton neighbours) whose template it projects onto best; the pass is kept only if the projection energy $\Psi$ increases, and the first pass that does not increase it stops the refinement. | Sequential against synchronous moves; the monotone quantity is $\Psi$, not the per-bin sum (see below). The candidate set in Experiment C (12 nearest centroids) matches the shipped count. |
| Greedy (Algorithm 2): every pair rescanned after each merge; a pair is scored on the union of its original members. | One representative per cluster; a pair is scored by merging the two representatives (Proposition 4.3 on two Gaussians); only pairs on a sparse spatial-hash neighbour graph (at most 8 neighbours) are considered, through a lazy-deletion heap. | Cost model (representatives against original members) and candidate set (neighbour graph against all pairs). |
| Method choice: every arm is run and compared (Section 9). | A method is chosen per level: greedy when the level's input has at most 5,000 splats, $k$-means-style chunking with refinement above. Defaults are $K = 4$ (an integer of at least 2), three coarser levels, five refinement passes and 12 candidate neighbours. | The experiments do not exercise the automatic choice. |
| Post-processing: none. | Coverage inflation and mass conservation follow the merge; the optional $L^2$ or volume refit (Section 5.5) follows those. | None of these steps is part of the experiments. |

of Proposition 4.3: with the unit templates $u_n$ of the kept shapes, solve $\sum_{n'} \langle u_n, u_{n'} \rangle a_{n'} = \langle u_n, f \rangle$ for all $n$). Both are strict simplifications of the contribution-based cull: they use the same "rank by per-splat scalar importance and keep the top $M$" template, but with an analytically clean importance score in place of a numerical $L^\infty$ budget search. The refit variant is an optimistic bound on what a removal of this kind can reach, because a renderable store needs positive amplitudes and the unconstrained solve may return negative ones. The qualitative comparison between substitutive merging and removal-based culling is expected to carry over to the full contribution-based cull; a fair head-to-head comparison is left for future work.

##### 8.5 Audit metric: `rel_l2`

The relative $L^2$ error $\|f - g\|_{L^2} / \|f\|_{L^2}$, written `rel_l2` in Luxar's quality metrics and in the experiment records below, is the natural endpoint quality measure for any LOD level. Experiments B and C below report `rel_l2` (and record $D_{\mathrm{CS}}$); Experiment A reports the squared-error decomposition normalised by $\|f\|^2$.

#### 9 Numerical Experiments

We run three experiments that test the framework empirically. Experiment A asks when the per-bin sum is a faithful surrogate for the global error; Experiment B compares the partition algorithms on synthetic mixtures; Experiment C repeats the comparison on a real fit and adds two removal baselines. The analysis code accompanying this document generates the figures deterministically from a fixed random seed. Its 45 unit tests guard the mathematical correctness of the building blocks: the Gaussian inner product against numerical 2D quadrature, moment matching against the pairwise formulas of the companion document, the $L^2$ merge cost, the partition decomposition $\|f - g\|^2 = \sum_j \|r_j\|^2 + 2 \sum_{j<k} \langle r_j, r_k \rangle$, the monotonicity of the cost-increment relocation, the vectorised kernel against the scalar one, and the two mass-ranked culling baselines.

**Provenance.** The synthetic mixtures are drawn as follows. Centres are uniform in $[0, s]^D$, where the side length $s$ sets the bin separation. Each covariance is $\boldsymbol{\Sigma}_i = Q \,\mathrm{diag}(\sigma_1^2, \ldots, \sigma_D^2)\, Q^\top$ with a random rotation $Q$ and axis standard deviations $\sigma_d \sim U[0.05, 0.4]$, and each amplitude is $a_i \sim U[0.5, 2]$. The draws come from NumPy's default generator, seeded with 20260425 for Experiment A, 20260426 for Experiment B and 20260427 for Experiment C; subsample $i$ of Experiment C draws its indices from a generator seeded with $20260427 + 1000\, i$. $k$-means uses a $k$-means++ start and at most 30 sweeps, and the relocation of Algorithm 3 runs five passes. Timings are wall-clock seconds of the partition step alone (the merge evaluation is excluded; for Experiment C the tabulated times are those of the relocation passes alone), measured on a single-process pure-NumPy implementation that recomputes the full Gram matrix per candidate. They characterise this reference code, not the shipped algorithms (Section 8.3).

**Reproducibility.** Three artefact types accompany this document: the analysis code, its unit tests, and one deposited experiment record per experiment. Each record holds the metric rows and the sweep parameters. The Experiment C record alone also carries a provenance block: the host platform and library versions of the run, the identity of the input store, the SHA-256 of the store file and of the decoded arrays, its stored and post-filter splat counts and the index set of every subsample.

##### 9.1 Experiment A: cross-bin interference

**Setup.** We draw $N = 64$ random anisotropic 2D Gaussians with means uniformly distributed in a box $[0, s]^2$, for varying side length $s$ (the "bin separation"). They are partitioned by $k$-means on the means into $M = 16$ bins ($K = 4$, the case the figure shows) and, in two further sweeps from the same seed, into $M = 32$ and $M = 8$ bins ($K = 2$ and $K = 8$), each bin merged by the $L^2$-optimal merge. Each setting is summarised by the median over 12 trials. The generator's median geometric-mean splat width is $\sigma = 0.21$, so $s/\sigma$ runs from 1.4 to 48.0; the quantity the result turns on is the spacing between neighbouring bin centroids in units of $\sigma$, which at $K = 4$ runs from 0.27 to 9.18 over the same $s$.

**Quantities.** We measure the per-bin sum $\sum_j \|r_j\|^2$, the global error $\|f - g\|^2$, and their difference (the cross-bin term $2\sum_{j<k}\langle r_j, r_k\rangle$), all normalised by $\|f\|^2$.

**Result.** Figure 8 shows that as bin separation grows, the cross-bin term shrinks and the per-bin sum becomes a faithful surrogate for the global error. In the densest regime ($s = 0.3$), the cross-bin term accounts for $\approx$+73.1 % of the global error (median over the 12 trials) and is *positive* (residuals reinforce: the per-bin sum *underestimates* the true error). For $s \geq 0.8$ the *median* per-bin sum lies within 3.0 % of the global error, but the cross-bin term is small only in aggregate, not uniformly. At $s = 0.8$ the per-trial cross-bin term still ranges from −33.3 % to +38.8 % of the global error with a sign that fluctuates trial-to-trial, and the range narrows only gradually with separation (−31.1 % to +19.0 % at $s = 1.8$; −1.9 % to +1.8 % at $s = 10$; the per-separation values are read from the deposited experiment record). The result supports the approximate-bin-additivity assumption of $L^2$-based partition algorithms in the median whenever the partition has spatially decoupled bins, quantifies the per-trial spread that a single-realisation user should expect, and marks the regime ($s \lesssim 0.5$) where the assumption breaks down outright.

That threshold belongs to one compression factor. At $K = 2$ ($M = 32$) the median per-bin sum is 82.1 % of the global error at $s = 0.8$ and 92.5 % at $s = 1.2$, reaching 98.6 % at $s = 1.8$. At $K = 8$ ($M = 8$) it *over*estimates, 119.4 % at $s = 0.8$ and 115.7 % at $s = 1.2$, where the per-trial cross-bin term runs from −69.9 % to +17.8 %. Taking as the criterion a median within 5 % of the global error at every larger separation, the reference separation is $s = 0.8$ at $K = 4$, 1.8 at $K = 2$ and 2.5 at $K = 8$; the median nearest-neighbour distance between bin centroids there is 0.73, 1.08 and 3.54 splat widths. The spacing that decouples the bins is therefore not one number, and it is not monotone in $K$: the reference separation is largest at $K = 8$ and smallest at $K = 4$, so the $K = 4$ figure should be read with that in mind.

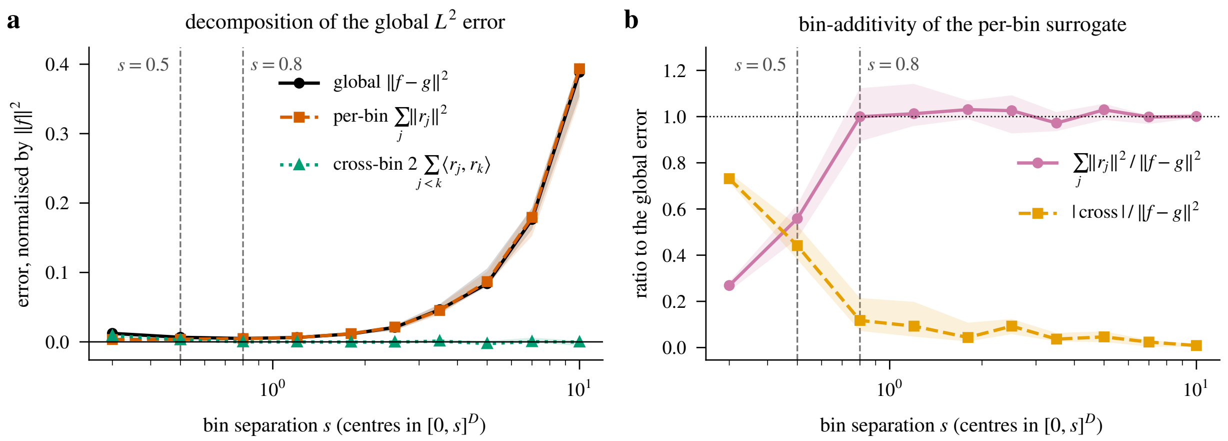

Figure 8: Experiment A: decomposition of the global $L^2$ approximation error. (a) The per-bin sum $\sum_j \|r_j\|^2$, the cross-bin term $2\sum_{j<k}\langle r_j, r_k\rangle$, and the global error $\|f - g\|^2$, all normalised by $\|f\|^2$, against bin separation. (b) The ratios $\sum_j \|r_j\|^2/\|f - g\|^2$ and $|\text{cross}|/\|f - g\|^2$. Lines are medians over 12 trials per separation; shaded bands are the interquartile range; the dashed guides mark $s = 0.5$, below which the assumption breaks down, and the reference separation $s = 0.8$ from which the median is faithful. Shown for $M = 16$ ($K = 4$); in units of the median splat width $\sigma = 0.21$ the separation axis runs from $s/\sigma = 1.4$ to 48.0 and the bin-centroid spacing from 0.27 to 9.18 $\sigma$. When bins are well-separated the per-bin sum tracks the global error in the median; individual trials still deviate by tens of percent at moderate separation, and the separation needed differs between compression factors without a monotone trend (text).

#### 9.2 Experiment B: algorithm comparison on synthetic mixtures

**Setup.** Synthetic 2D mixtures with $N \in \{32, 64, 128\}$, compression $K \in \{2, 4, 8\}$, fixed bin separation $s = 1.5$. Four arms, named as in Experiment C: (i) *random*, a random partition (control); (ii) *k-means* on the centres; (iii) *k-means + relocation*, the $k$-means partition followed by 5 passes of the cost-increment relocation of Algorithm 3; (iv) *greedy*, greedy hierarchical merging (Algorithm 2). All use the $L^2$-optimal $K$-wise merge of Algorithm 1 for representatives.

**Quantities.** Relative $L^2$ error $\sqrt{\|f - g\|^2/\|f\|^2}$, summarised by the median over 4 trials; CS-divergence between $f$ and $g$ is recorded in the deposited experiment record but not plotted.

**Result.** Figure 9 shows a clear ordering at small $K$ that softens at large $K$: **greedy** $\lesssim$ **$k$-means + relocation** $<$ **$k$-means** $\ll$ **random**. At $N = 128, K = 2$, the median relative $L^2$ errors are $\approx 0.017$ (greedy), 0.021 ($k$-means + relocation), 0.053 ($k$-means), 0.176 (random); at $N = 128, K = 8$ they are $\approx 0.058$ (greedy), 0.059 ($k$-means + relocation), 0.082 ($k$-means), 0.256 (random). At large compression $k$-means + relocation therefore *essentially matches* greedy while being $\approx 8.0\times$ faster (median assignment time 8.1 s vs. 64.5 s per trial at $N = 128$, $K = 8$). The speed-up is specific to the count regime of this Python reference implementation. The ratio of the median greedy assignment time to the median $k$-means + relocation one is 2.7 to 3.1$\times$ at $N = 32$, 4.4 to 5.5$\times$ at $N = 64$ and 7.7 to 8.3$\times$ at $N = 128$ across $K \in \{2, 4, 8\}$, growing with $N$ as the $O(N^3)$ candidate rescans of Algorithm 2 take over. It says nothing about the shipped algorithms (Section 8.3). The median random error stays 1.8 to 3.7$\times$ the $k$-means one at every

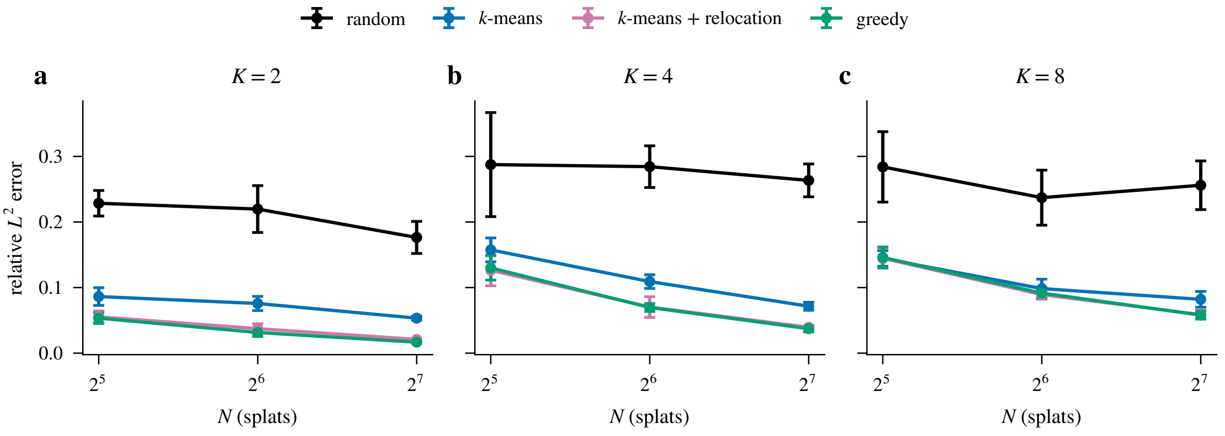

Figure 9: Experiment B: relative $L^2$ approximation error of the partition-and-merge LOD as a function of $N$, for compression factors $K = 2$, 4 and 8 (panels a to c). Markers are the median `rel_l2` over 4 trials per $(N, K)$; error bars are the population standard deviation over those trials. Spatial clustering ($k$-means, with and without relocation) and greedy both substantially outperform random partitioning at every compression factor.

$(N, K)$, with no monotone trend in $K$, while the gap between greedy and $k$-means + relocation shrinks with $K$ because the local greedy-pair choices become less informative when bins are large. The implication for practice: **$k$-means + relocation is the recommended workhorse**; greedy beats it only at $K = 2$, by 3.7 % ($N = 32$), 15.6 % ($N = 64$) and 20.1 % ($N = 128$) of the $k$-means + relocation median error (an advantage that grows with $N$), and is within 5.5 % of it either way at $K \geq 4$, at a several-fold higher assignment cost in this implementation.

##### 9.3 Experiment C: real DAPI nuclei fit

**Setup.** We load the hosted DAPI nuclei fit (3D), the checksum-pinned store from which Luxar's blastocyst DAPI nuclei demo compiles its scene (the deposited record names the store and its digests). We keep the splats that pass a validity filter (amplitude above $10^{-3}$ in the store's amplitude units and smallest covariance eigenvalue above $10^{-12}$ in squared scene units; every one of the 7,740 stored splats passes). Two blocks are run. The *full* block works on all 7,740 splats at $K \in \{2, 4, 8\}$ ($M =$ 3,870, 1,935 and 967; the analysis code rounds the budget down, $M = \lfloor N/K \rfloor$, which departs from the $\lceil N/K \rceil$ of Section 1 only at $K = 8$, where the ceiling would give one bin more, and Table 3 lists the budgets the code used) with four arms, named as in Tables 3 and 4. *$k$-means* runs $k$-means on the centres, then merges. *$k$-means + relocation* follows the $k$-means partition with 5 passes of the cost-increment relocation of Algorithm 3, each splat's candidates restricted to the 12 nearest bin centroids other than its own (the shipped count), then merges. *Culling* is mass-ranked culling with verbatim amplitudes, and *culling + refit* is the same culling with the joint least-squares amplitude refit of Section 8.4. *Greedy* (greedy hierarchical) is not run on the full fit (its all-pairs rescans are cubic in $N$). The *subsample* block draws 8 uniform subsamples of $N = 128$ splats without replacement (subsample $i$ from a generator seeded with $20260427 + 1000\,i$) and runs the four arms plus greedy, whose partitions at the three budgets come from one agglomeration per subsample. A subsample changes the geometry, not the splats:

Table 3: Experiment C, full fit: relative $L^2$ error of the four arms on all 7,740 splats. The relocation ran 5 passes in 90.0, 81.1 and 100.8 s at $K = 2, 4, 8$; the amplitude refit returned 11, 35 and 26 negative amplitudes out of $M$. $M$ is the requested budget; the relocation left 3,864, 1,933 and 967 non-empty bins (Algorithm 3 may empty a singleton bin).

| $K$ | $M$ | $k$-means | $k$-means + relocation | culling | culling + refit |
|---|---|---|---|---|---|
| 2 | 3,870 | 0.094 | 0.065 | 0.165 | 0.108 |
| 4 | 1,935 | 0.140 | 0.104 | 0.312 | 0.198 |
| 8 | 967 | 0.185 | 0.141 | 0.424 | 0.277 |

Table 4: Experiment C, subsamples: relative $L^2$ error of the five arms, median [minimum, maximum] over the 8 subsamples of 128 splats. The refit returned at most 0, 0 and 0 negative amplitudes at $K = 2, 4, 8$.

| $K$ | $k$-means | $k$-means + relocation | greedy | culling | culling + refit |
|---|---|---|---|---|---|
| 2 | 0.464 [0.410, 0.569] | 0.318 [0.271, 0.422] | 0.349 [0.305, 0.457] | 0.309 [0.266, 0.396] | 0.308 [0.264, 0.393] |
| 4 | 0.648 [0.551, 0.844] | 0.496 [0.436, 0.676] | 0.553 [0.512, 0.707] | 0.470 [0.429, 0.616] | 0.469 [0.428, 0.614] |
| 8 | 0.813 [0.673, 0.945] | 0.609 [0.546, 0.839] | 0.654 [0.598, 0.887] | 0.575 [0.519, 0.720] | 0.573 [0.518, 0.719] |

the median nearest-neighbour distance between centres is 1.69 scene units on the full fit and 6.43 on the subsamples (median over the 8), 3.8× wider, while the splats themselves, with a median geometric-mean width of 1.02 on the full fit, are unchanged.

**Quantities.** Relative $L^2$ error at matched splat budgets ($D_{\mathrm{CS}}$ is recorded in the deposited experiment record). For the full block the relocation's wall-clock time and the number of negative amplitudes returned by the refit are recorded; every subsample arm is summarised by its median, minimum and maximum over the 8 subsamples.

**Result.** Tables 3 and 4 hold the numbers and Fig. 10 draws them.

- **On the full fit, cost-aware merging beats both culling arms at every $K$.** $k$-means + relocation lowers the relative error by 61 to 67 % against culling and by 39 to 49 % against culling + refit; the refit closes 52 to 58 % of the gap between culling and $k$-means + relocation. Plain $k$-means is also below both culling arms at every budget, so on the full fit the geometric proxy is a usable warm start; what it leaves for the relocation to recover is 0.140 against 0.104 at $K = 4$.
- **On the subsamples the ordering inverts.** One of the two culling arms is the best arm in 8, 7 and 8 of the 8 subsamples at $K = 2, 4, 8$; $k$-means + relocation is the best merging arm in the median; greedy is below $k$-means + relocation in only 1, 0 and 0 subsamples. The refit changes little here (0.469 for culling + refit against 0.470 for culling at $K = 4$).

The two blocks are the same splats at two neighbour spacings, so the reversal is a regime statement, not a ranking of algorithms. Where neighbouring splats overlap (full fit: median centre spacing 1.69 against a median width of 1.02), a bin of $K$ neighbours is close to one blob and its moment-matched representative loses little, while removing a splat leaves a hole its neighbours do not fill.

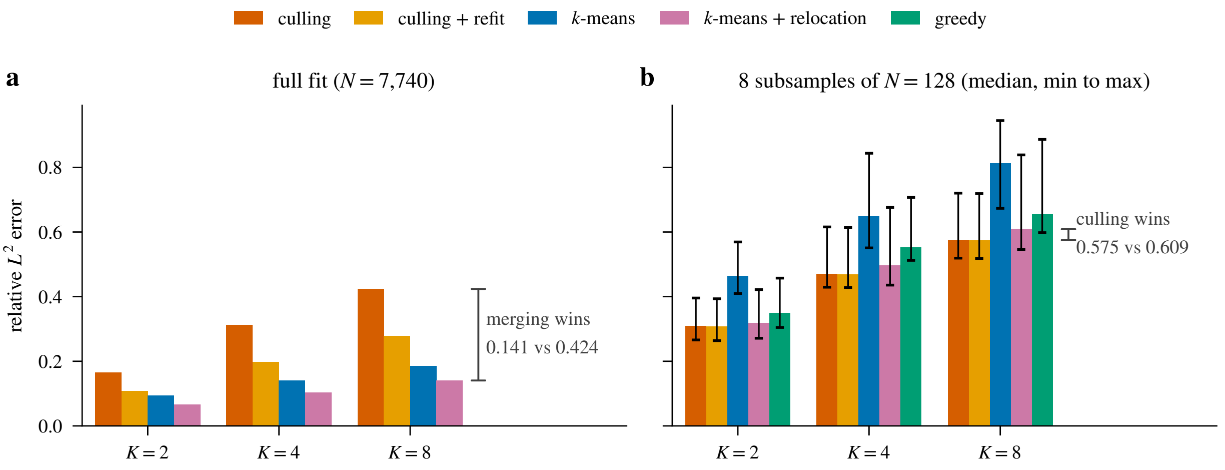

Figure 10: Experiment C: substitutive merging vs. mass-ranked culling on the DAPI nuclei fit (3D). (a) The full fit (7,740 splats; four arms, one run per budget). (b) 8 subsamples of 128 splats (five arms; bars are medians, whiskers span the minimum to the maximum over the subsamples). On the full fit every partition-and-merge arm beats both culling arms; on the sparse subsamples both culling arms win in the median and greedy does not improve on $k$-means + relocation. The annotation at $K = 8$ gives culling against $k$-means + relocation on each side of the inversion.

Where the spacing is 3.8× wider (the subsamples), the members of a bin are isolated peaks that a single Gaussian smears, and keeping the heaviest $M$ verbatim loses only the light ones. The $K = 2$ advantage of greedy on synthetic mixtures (Section 9.2) does not appear on the sparse subsamples. The $k$-means + relocation partition is where the mass-conservation discussion of Section 8.2 takes its numbers: at $K = 4$ the $L^2$-optimal amplitude is below the mass-conserving one in 79.8 % of the bins with at least two members (median ratio 0.986), at $K = 2$ and 8 in 67.4 % and 80.6 % of them. The conclusion is scoped to what was run: one fit, three budgets, two removal baselines; how the arms compare with the shipped contribution-based cull is not tested here.

#### 10 Discussion and Open Questions

**Choice of $K$.** Smaller $K$ (especially $K = 2$) gives the best per-level fidelity but builds a deeper tree; larger $K$ trades off detail for storage and rendering speed. The companion document's pairwise analysis covers $K = 2$ in full closed form; the present document generalises but does not optimise the choice of $K$, which depends on the deployment budget.

**Amplitude variation.** When bin members differ in amplitude by orders of magnitude (common in microscopy where small bright nuclei coexist with large dim background), the mass-weighted moment matching of Proposition 4.1 is dominated by the bright members. An amplitude-balanced variant that caps weight contributions or uses log-amplitude weights might be preferable for such data; we leave this as an open question.

**Generalised Gaussians (sharpness $\gamma \neq 2$).** The Luxar model uses the strict Gaussian falloff, exponent $\gamma = 2$ in $\exp(-\frac{1}{2}|\cdot|^{\gamma})$, which admits all closed-form inner-product expressions used here. With a per-splat super-Gaussian exponent $\gamma \neq 2$ the universal kernel (3) would no longer apply: the pairwise overlaps would need numerical integration (deterministic quadrature on a query grid, or Monte Carlo) or special-case formulas; equal-width Laplace kernels ($\gamma = 1$), for instance, have the exact one-dimensional overlap $(1 + |a - b|)\, e^{-|a-b|}$. The sample-based audits ($L^\infty$ on a query grid, perceptual loss) would remain available, and a perturbation analysis around $\gamma = 2$ is a plausible direction for future work.

**Per-channel and view-dependent splats.** Multi-channel splats with per-channel amplitudes can be handled by treating each channel as an independent scalar field and applying substitutive LOD per channel. View-dependent splats (with spherical-harmonic colour) require an angular extension of the kernel that is closed-form but more involved; we have not explored it.

**Re-fitting at each level.** Substitutive merging produces a coarse representation by *combining* existing splats; by itself it does not re-fit the underlying volume. The optional volume refit of Section 5.5 does exactly that: it re-runs the Luxar fitting pipeline on each level with the merged splats as initialisation and keeps the result only where it lowers the full-grid error. Whether its gain justifies its cost, and how large the gain is, is an empirical question that this document does not measure.

#### 11 Summary

The reader who wants to build a substitutive level of detail can take four things from this document. The cost metric to optimise is squared $L^2$: it is closed-form between Gaussian mixtures, and it is the metric the rendering pipeline measures. Its one defect, the cross-bin interference term, is itself closed-form and small in the median once bins are spatially separated; the separation this requires is $s = 1.8$, 0.8 and 2.5 at $K = 2$, 4 and 8, not a monotone function of $K$, and single trials spread by tens of percent at moderate separation. Cauchy–Schwarz divergence is the scale-invariant alternative for ranking partitions; $L^\infty$ and generalised KL are proposed as audit metrics and are not measured here.

The merge to use inside a bin is the moment-matched shape with the $L^2$-optimal amplitude of Proposition 4.3, the $K$-wise extension of the companion document's pairwise result. The partition to use is $k$-means on the centres followed by the cost-increment relocation of Algorithm 3. On synthetic mixtures it comes within 5.5 % of greedy hierarchical merging at $K \geq 4$, and greedy leads by up to 20.1 % only at $K = 2$. A numerical joint relaxation is a reference value, not a certified bound, and no analytical lower bound on the $M$-Gaussian approximation error is claimed.

Whether merging beats removal is a property of the data, not of the algorithm. On the full 7,740-splat DAPI nuclei fit, $k$-means + relocation reaches 0.104 at $K = 4$ against 0.198 for culling + refit and 0.312 for culling, and plain $k$-means also beats both culling arms at every budget. On 128-splat subsamples whose neighbour spacing is 3.8× wider, both culling baselines beat every merging arm in the median and greedy does not improve on $k$-means + relocation. Neighbour spacing against splat width decides.

The shipped `levels` recipe runs scalable variants of these steps (Morton-chunk warm start, projection-energy acceptance, representative-pair greedy, coverage inflation and mass conservation); Table 2 states where each departs from what was tested. The 45 unit tests that accompany the analysis guard the mathematical building blocks, including the sign handling of $\langle \phi_i, \phi_j \rangle$ for residuals with negative amplitude.

[9] Zhiwen Fan, Kevin Wang, Kairun Wen, Zehao Zhu, Dejia Xu, and Zhangyang Wang. Light-Gaussian: Unbounded 3D Gaussian compression with 15× reduction and 200+ FPS. In *Advances in Neural Information Processing Systems*, volume 37, 2024.

[10] Guangchi Fang and Bing Wang. Mini-Splatting: Representing scenes with a constrained number of Gaussians. In *European Conference on Computer Vision (ECCV)*, 2024.

[18] Luxar Project. Analytical merging error for Gaussian splats: An $L^2$ framework for optimal approximation, 2026. Supplementary material, this volume.

[19] Frank Nielsen. Closed-form information-theoretic divergences for statistical mixtures. In *Proceedings of the 21st International Conference on Pattern Recognition (ICPR)*, pages 1723–1726. IEEE, 2012.

[20] Panagiotis Papantonakis, Georgios Kopanas, Bernhard Kerbl, Alexandre Lanvin, and George Drettakis. Reducing the memory footprint of 3D Gaussian splatting. *Proceedings of the ACM on Computer Graphics and Interactive Techniques*, 7(1), 2024. doi: 10.1145/3651282.

[21] Kerui Ren, Lihan Jiang, Tao Lu, Mulin Yu, Linning Xu, Zhangkai Ni, and Bo Dai. Octree-GS: Towards consistent real-time rendering with LOD-structured 3D Gaussians. arXiv preprint arXiv:2403.17898, 2024.
