## Supplementary material for "Luxar: Gaussian splatting for microscopy and scalable interactive web visualisation of multidimensional scientific data": supp_doc_09_additive_lod

### Additive Levels of Detail for Gaussian Splats: Order Optimisation for Progressive Loading

Supplementary Document 9 – Luxar

**Abstract**

An additive level of detail for a Gaussian-splat scene is an ordering of its $N$ splats: the viewer draws the partial sum of the prefix that has arrived, so every prefix should approximate the full scene as closely as possible in the $L^2$ sense. The cumulative energy captured by a subset of splats is a monotone submodular set function whenever the splat Gram matrix is non-negative, as positive amplitudes guarantee, so the greedy ordering (always add the splat with the largest one-step drop in residual energy) captures at least $(1 - 1/e)$ of the optimal cumulative energy at every prefix length. The greedy rule is an $O(N^2)$ fixed-coefficient matching pursuit, made tractable at full scale by a support-pruned sparse Gram matrix. A Rayleigh–Ritz argument gives a lower bound on the prefix error; its tempting eigen-coordinate strengthening is false. Greedy has the lowest area under the loading curve (AUC, lower is better) on every synthetic mixture and comes within 0.08% of the optimum on exhaustively solved small instances. On four Luxar datasets subsampled to $N = 4\,000$ splats greedy again has the lowest AUC but lowers it by only 0.2–2.6% relative to the $O(N \log N)$ self-energy ordering, the gap growing with Gram density. The spectral ordering trails both on every dataset, and below a small leading cluster of splats its order is set at floating-point level. Self-energy is therefore a strong cheap default; greedy is the upgrade when its precompute is affordable.

#### Contents

**1 Introduction and Problem Formulation** **2**

**2 Notation and Setup** **5**

**3 Catalogue of Ordering Strategies** **7**

**4 Theory** **9**

**9 Discussion and Open Questions** **29**

**10 Summary** **30**

### 1 Introduction and Problem Formulation

A *level of detail* (LOD) is a cheaper stand-in for a scene that a viewer can show while the full scene is unavailable or not worth its cost. The companion supplementary documents [7, 8] treat the *substitutive* kind, which replaces $N$ splats by $M < N$ new representative splats. This document treats the *additive* kind, which keeps the original splats and only chooses the order in which they are sent. The question it answers is which order to choose, what can be proved about the best choice, and how much the provably good choice gains over cheap heuristics on real data.

A Luxar scene of Gaussian splats represents a non-negative scalar field

$$f(\boldsymbol{x}) = \sum_{i=1}^{N} \phi_i(\boldsymbol{x}), \qquad \phi_i(\boldsymbol{x}) = a_i \exp\Big(-\tfrac{1}{2}(\boldsymbol{x}-\boldsymbol{\mu}_i)^\top \boldsymbol{\Sigma}_i^{-1}(\boldsymbol{x}-\boldsymbol{\mu}_i)\Big), \tag{1}$$

with peak amplitudes $a_i > 0$, centres $\boldsymbol{\mu}_i \in \mathbb{R}^D$, and positive definite covariances $\boldsymbol{\Sigma}_i \succ 0$. The splats are *unnormalised*: no normalising constant divides the exponential, so $a_i$ is the height of the splat at its centre and its integral grows with the covariance volume. Two splats interact through their $L^2$ inner product, and the $N \times N$ matrix of these inner products, the *Gram matrix* $\boldsymbol{G}_{ij} = \langle \phi_i, \phi_j \rangle_{L^2}$, is the object on which every result below is stated; its closed form is given in Section 2.

**The progressive-loading question.** Suppose the full set of $N$ splats is already stored and must be transmitted or paged into the viewer over time. The viewer's display at time $t \propto k$ shows the partial sum of the first $k$ splats it has received. The problem is to choose *the order in which the $N$ splats are loaded so that the viewer's approximation is as faithful to the full scene as possible at every intermediate $k$.*

Formally, fix a permutation $\pi : \{1, \ldots, N\} \to \{1, \ldots, N\}$ and define the prefix approximation and residual at step $k$:

$$g_k(\boldsymbol{x}) \;=\; \sum_{j=1}^{k} \phi_{\pi(j)}(\boldsymbol{x}), \qquad \boldsymbol{r}_k(\boldsymbol{x}) \;=\; f(\boldsymbol{x}) - g_k(\boldsymbol{x}) \;=\; \sum_{j=k+1}^{N} \phi_{\pi(j)}(\boldsymbol{x}). \tag{2}$$

The set of the first $k$ splats is the *prefix* of length $k$. At the boundaries, $g_0 = 0$ and $g_N = f$, so the residual-energy curve $E_k = \|\boldsymbol{r}_k\|_{L^2}^2$ starts at $\|f\|^2$ and ends at 0. All permutations meet at the same endpoints; only the path between them depends on $\pi$.

**Anytime fidelity.** Because the user may interrupt loading at any $k$, the natural objective is *prefix-monotone*: it rewards a small $E_k$ at every prefix length $k$ at once, with no single target length. Two scalar objectives capture this:

- **Pointwise.** At a chosen $k$, minimise $E_k$, the size-$k$ subset-selection problem.
- **Anytime AUC.** Minimise the area under the loading curve $E_k/\|f\|^2$ plotted against $k/N$, evaluated by the trapezoidal rule on the $N+1$ prefix points,

$$\mathrm{AUC}(\pi) \;=\; \frac{1}{N\,\|f\|^2} \sum_{k=0}^{N-1} \frac{E_k(\pi) + E_{k+1}(\pi)}{2} \;=\; \frac{1}{N\,\|f\|^2} \sum_{k=0}^{N-1} E_k(\pi) \;-\; \frac{1}{2N} \;\in\; [0, 1), \tag{3}$$

  where the second form uses $E_N = 0$. The trapezoidal AUC differs from the plain prefix average $\frac{1}{N\|f\|^2} \sum_{k<N} E_k$ by the permutation-independent constant $1/(2N)$, so the two rank permutations identically at fixed $N$; every AUC reported in Section 8 is the trapezoidal value (3), and a lower AUC is better.

Proposition 4.1 below shows that the prefix sum $\sum_{k<N} E_k$, and hence the AUC, has an exact quadratic-assignment closed form in the Gram matrix, and Theorem 4.5 shows that a single greedy ordering captures at least $(1 - 1/e)$ of the optimal cumulative energy at *every* $k$ simultaneously; Corollary 4.7 turns this into an affine bound on the AUC.

**Scope and assumptions.** Three assumptions hold throughout and qualify every result. **(i) Scalar-field $L^2$ objective.** The fidelity metric is the analytical $L^2$ residual $\|f - g_k\|_{L^2}$, where $f$ is the non-negative scalar field (1). It matches the Luxar pixel/voxel pipeline (rendered MSE on query samples) and the companion $L^2$ framework of [7, 8], but it does not capture view-dependent shading, alpha compositing, or perceptual rendering metrics, which would require their own analyses. **(ii) Equal cost per splat.** Every splat is assumed to incur the same loading cost, so the prefix index $k$ doubles as both a splat budget and a "time" axis. Real progressive-streaming systems pay bytes (and splats can encode to different byte sizes), and an unequal-cost extension would require a knapsack constraint in place of a cardinality constraint. **(iii) Anytime AUC weights all prefixes equally.** The uniform AUC objective gives late prefixes the same weight as early ones; in practice progressive viewers usually weight the first prefixes more heavily. The pointwise objective in Theorem 4.5 is more flexible (any $k$-specific bound applies); the AUC bound in Corollary 4.7 is the uniform-weight specialisation.

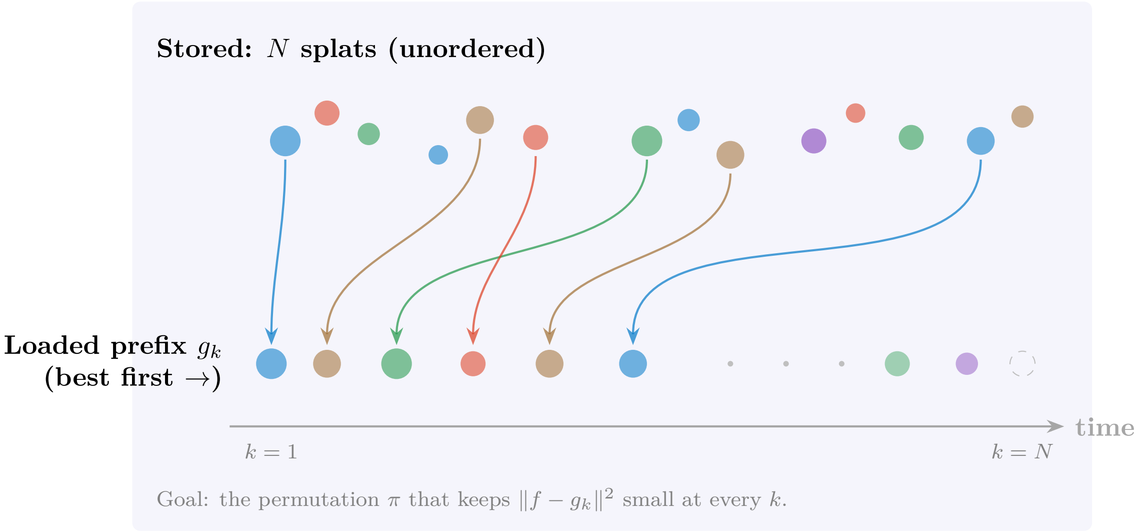

Figure 1: The progressive-loading question. All $N$ splats are already available in storage; the viewer receives them in some order and renders the partial sum of the prefix. The $L^2$ residual curve $E_k = \|f - g_k\|^2$ depends only on the loading order and not on which splats are eventually loaded (since $g_N = f$ in all cases). Picking the "best first" is the additive-LOD problem.

**What is proved and what is measured.** The $(1 - 1/e)$ guarantee of Theorem 4.5 is a statement about the greedy ordering. A shipped Luxar *additive (streaming) ladder* is a stored ordering cut into slabs, each a *rung (sub-LOD)*, and the default build runs the greedy ordering only for a *leaf* of at most 5 000 splats; above that size it switches to the $O(N \log N)$ $L^2$ *self-energy* ordering of Section 3, for which no approximation guarantee is proved here. The standing of that default rests on the measurements of Section 8.3, where greedy lowers the AUC relative to self-energy by a margin that is small on every dataset and grows with Gram density. The rung sizes, the per-rung *energy stamps* the viewer reads and the *brightness compensation* it applies while a ladder streams are engineering choices layered on top of the ordering, not consequences of the theory; Section 6.3 describes them.

**Substitutive vs. additive LOD.** The substitutive LOD problem [8] *synthesises* $M < N$ new splats that summarise the original $N$; the additive LOD problem retains the original splats and only *orders* them. Both collapse to identical endpoints: at $M = N$ (resp. $k = N$) the approximation is exact. The two are complementary in deployment: a substitutive hierarchy provides static coarse representations for fixed-budget decode targets (mip levels for cache-pinned LODs); an additive ordering provides anytime fidelity for streaming under variable bandwidth or memory. In a real viewer pipeline both are useful and orthogonal: a substitutive coarse level can be paged in first, then refined via the additive ordering on the next level.

**Outline.** Section 2 sets notation. Section 3 catalogues ordering strategies and gives their complexities. Section 4 develops the core theory: the Gram-matrix closed form for the AUC, submodularity of the cumulative-energy utility, the greedy $(1 - 1/e)$ guarantee, and the lazy-greedy implementation. Section 5 states a Rayleigh–Ritz lower bound on the prefix error and shows by counterexample that the natural stronger feature-space bound fails. Section 6 relates the framework to existing progressive-rendering methods (PRoGS, LapisGS) and describes the Luxar implementation as shipped. Section 7 gives pseudocode. Section 8 reports numerical experiments on synthetic and real datasets. Section 9 discusses limitations and open questions, and Section 10 states what to use in practice.

##### Terms used in this document

**additive (streaming) ladder** One ordering of a leaf's splats cut into prefix-sum sub-LODs, so that every prefix of the download is a valid, progressively refining rendering of the same splat set.

**rung (sub-LOD)** One contiguous slab of an additive ladder, stored as its own group (`additive_<i>`, index 0 the coarsest) and appended to the committed prefix when it arrives; a document that uses "rung" for one step of a splat-count sweep says so where it first does.

**leaf** A node that carries splat arrays directly (centres, amplitudes, Cholesky factors) together with an additive ladder of at least one sub-LOD, as opposed to a group node that only holds children.

**committed prefix (committed count)** The part of an additive ladder that has been processed, uploaded to the GPU and is on screen, as distinct from rungs whose download has merely completed.

**self-energy** The squared $L^2$ norm of one splat's Gaussian, $a_i^2\, \pi^{D/2}\, |\Sigma_i|^{1/2}$, the ordering weight of an additive ladder above the greedy size limit of 5,000 splats (below it a greedy residual ordering is used), the unit of the equal-energy breakpoints, and the quantity whose cumulative fraction stamps how much of a leaf a ladder prefix already carries.

**energy stamps** Two attributes written at build time: `energy_fraction_cum`, $e(k)$, on each sub-LOD is the cumulative self-energy fraction of the ladder prefix ending at rung $k$, and `reference_energy`, $w$, on each leaf is the leaf's absolute total self-energy used to weight leaves against one another.

**display gate** The viewer rule that keeps the previously displayed level on screen while a finer level is still streaming and releases the swap once the finer level's committed prefix carries at least 0.6 of its total self-energy ($e(k) \geq 0.6$), applied on upgrades only.

**brightness compensation** The factor $1/\max(e(k), 0.1)$ by which the viewer multiplies a leaf's opacity while its ladder is incomplete, holding the partial prefix near its final brightness so that arriving rungs do not pop, with the floor of 0.1 capping the boost at tenfold.

**`kind=lod` group** A substitutive level-of-detail group whose children are alternative renderings of the same content at different splat counts, of which the viewer draws exactly one at a time: the coarsest level whose median projected splat footprint stays at or below 1.5 CSS pixels when the levels carry footprint stamps, otherwise the level selected by the size of its projected bounding box (its fraction of the viewport area for a screen-area ladder, its normalised projected diagonal for a coverage ladder).

**`content_hash`** A post-order xxhash64 digest of a store (array bytes, storage identity, attributes and child digests) against which the viewer validates its cache, so any edit to the data invalidates stale chunks.

**PSNR** The peak signal-to-noise ratio, $10 \log_{10}(\text{peak}^2/\text{MSE})$ in decibels, where peak is the reference range of the values compared; unless a document states another range, the supplements take peak 1 on volumes normalised to $[0, 1]$, so that $\text{PSNR} = -10 \log_{10} \text{MSE}$.

#### 2 Notation and Setup

Notation extends the companion documents [7, 8]. We write $[N] = \{1, \ldots, N\}$ for the index set of the splats, $\mathbf{1}_S \in \{0, 1\}^N$ for the indicator vector of a subset $S \subseteq [N]$ (entry 1 on $S$, 0 elsewhere), $S^c = [N] \setminus S$ for its complement and $\mathbf{1} = \mathbf{1}_{[N]}$ for the all-ones vector.

**Splats and the Gaussian inner product.** A splat $\phi_i$ is the unnormalised Gaussian (1). The $L^2$ inner product between two splats has the closed form

$$K_{ij} \;=\; \langle \phi_i, \phi_j \rangle_{L^2} \;=\; a_i a_j \,(2\pi)^{D/2} \,\frac{\sqrt{|\boldsymbol{\Sigma}_i||\boldsymbol{\Sigma}_j|}}{\sqrt{|\boldsymbol{\Sigma}_i + \boldsymbol{\Sigma}_j|}}\, \exp\!\Big(-\tfrac{1}{2}(\boldsymbol{\mu}_i-\boldsymbol{\mu}_j)^\top(\boldsymbol{\Sigma}_i+\boldsymbol{\Sigma}_j)^{-1}(\boldsymbol{\mu}_i-\boldsymbol{\mu}_j)\Big). \tag{4}$$

For amplitudes $a_i > 0$ all entries are strictly positive: $K_{ij} > 0$ for all $i, j$. The Gram matrix $\boldsymbol{G} \in \mathbb{R}^{N\times N}_{\geq 0}$ has entries $\boldsymbol{G}_{ij} = K_{ij}$ and is symmetric positive semidefinite.

**Standard deviations and support.** The symbol $\sigma$ always denotes a standard deviation. A distance written $c\sigma$ is $c$ standard deviations along the widest axis of the splat concerned, $\sqrt{\lambda_{\max}(\boldsymbol{\Sigma}_i)}$, so a "$3\sigma$ support" is the ball of that radius around the centre. The Luxar renderer draws each splat only inside a finite Mahalanobis support of this kind, canonically $2.75\sigma$; the analytical field (1) is untruncated, and Section 7 explains how the two are reconciled.

**Splat scalar invariants.** We use three scalar invariants of a splat $\phi_i$:

- the *peak amplitude* $a_i$ (height of the splat at its centre $\boldsymbol{\mu}_i$);
- the *integral mass* $m_i \;=\; \int \phi_i(\boldsymbol{x})\, d\boldsymbol{x} \;=\; a_i\,(2\pi)^{D/2}\,|\boldsymbol{\Sigma}_i|^{1/2}$ (zeroth moment of the splat under the Lebesgue measure on $\mathbb{R}^D$);
- the $L^2$ *self-energy* $\|\phi_i\|^2_{L^2} \;=\; \int \phi_i(\boldsymbol{x})^2\, d\boldsymbol{x} \;=\; \boldsymbol{G}_{ii} \;=\; a_i^2\,\pi^{D/2}\,|\boldsymbol{\Sigma}_i|^{1/2}$ (squared $L^2$ norm; the diagonal of the Gram matrix obtained from (4) by setting $i = j$).

Peak amplitude depends on $a_i$ alone; integral mass is linear in $a_i$ and grows with covariance volume; $L^2$ self-energy is *quadratic* in $a_i$ but with the same covariance dependence as mass. The three invariants agree only in special cases. When all covariance volumes $|\boldsymbol{\Sigma}_i|$ are equal, all three rank by amplitude; when all amplitudes are equal, mass and self-energy both rank by covariance volume while the peak score ties every splat. On data heterogeneous in both amplitude and extent they rank splats differently and yield distinct ordering strategies (Section 3).

**Permutations, prefixes, residuals.** A permutation $\pi$ orders the splats; we write $\rho_i = \pi^{-1}(i) \in [N]$ for the *rank* of splat $i$ (rank 1 = first loaded). The prefix at step $k$ is the index set $S_k = \pi(\{1,\ldots,k\}) = \{i : \rho_i \leq k\}$. For any subset $S \subseteq [N]$ we write $g_S = \sum_{i\in S}\phi_i$ for its partial sum; the prefix function and residual function are $g_k = g_{S_k}$ and $\boldsymbol{r}_k = f - g_k = \sum_{i\notin S_k}\phi_i$, with squared $L^2$ norms

$$E_k(\pi) \;=\; \|\boldsymbol{r}_k\|^2_{L^2} \;=\; \sum_{i\notin S_k}\sum_{j\notin S_k} \boldsymbol{G}_{ij} \;=\; \mathbf{1}^\top_{S_k^c}\boldsymbol{G}\,\mathbf{1}_{S_k^c}. \tag{5}$$

The full-scene energy is $E_0 = \|f\|^2 = \mathbf{1}^\top\boldsymbol{G}\mathbf{1}$.

**Cumulative-energy utility.** For any subset $S \subseteq [N]$, define the cumulative-energy utility

$$U(S) \;=\; \|f\|^2 \;-\; \|f - g_S\|^2 \;=\; 2\sum_{i\in S}\sum_{j=1}^{N}\boldsymbol{G}_{ij} \;-\; \sum_{i,j\in S}\boldsymbol{G}_{ij}. \tag{6}$$

Then $U(\emptyset) = 0$, $U([N]) = \|f\|^2$, and the residual at prefix $S_k$ satisfies $E_k = \|f\|^2 - U(S_k)$. Maximising $U(S)$ over size-$k$ subsets is therefore equivalent to minimising $E_k$. Where a permutation must be named we write $E_k(\mathrm{g})$ for the residual of the greedy prefix of Section 3 and $E_k^\star = \min_{|S|=k} \|f - g_S\|^2$ for the size-$k$ optimum, the *lower envelope* that no single permutation need attain.

#### 3 Catalogue of Ordering Strategies

We consider six ordering strategies of increasing sophistication. All are evaluated empirically in Section 8; the design space is summarised in Table 1.

**(1) Random.** $\pi$ drawn uniformly at random; the trivial baseline. Combinatorial averaging over uniform $\pi$ gives the exact expectation

$$\mathbb{E}[E_k] \;=\; \tfrac{N-k}{N}\,\mathrm{tr}(\boldsymbol{G}) + \tfrac{(N-k)(N-k-1)}{N(N-1)}\,\big(\,\|f\|^2 - \mathrm{tr}(\boldsymbol{G})\,\big), \tag{7}$$

which, defining the *orthogonality fraction* $\eta = \mathrm{tr}(\boldsymbol{G})/\|f\|^2 \in (0,1]$, reads $\mathbb{E}[E_k]/\|f\|^2 = (1 - k/N)\,\eta + (1-k/N)^2(1-\eta) + O(1/N)$. The behaviour interpolates between *linear* in $k/N$ for nearly orthogonal dictionaries ($\eta \to 1$) and *quadratic* for highly redundant ones ($\eta \to 0$).

**(2) Peak amplitude.** Sort by $a_i$ descending. Implements the naive heuristic "brightest first". Ignores splat extent.

**(3) Integral mass.** Sort by $m_i = a_i(2\pi)^{D/2}|\boldsymbol{\Sigma}_i|^{1/2}$ descending. This is the splat's contribution to $\int f$ (zeroth moment) and weighs both peak amplitude and covariance volume.

**(4) $L^2$ self-energy.** Sort by $\boldsymbol{G}_{ii} = \|\phi_i\|^2 = a_i^2\pi^{D/2}|\boldsymbol{\Sigma}_i|^{1/2}$ descending. This is the squared $L^2$ norm of the splat in isolation. In the orthogonal limit ($\boldsymbol{G}_{ij} \approx 0$ for $i \neq j$) we have $\|f\|^2 \approx \sum_i \boldsymbol{G}_{ii}$, so the splat with largest $\boldsymbol{G}_{ii}$ contributes most to $\|f\|^2$ and is the correct first pick; greedy reduces to self-energy ordering in this limit (Remark 3.1).

**(5) Spectral.** Sort by $|\boldsymbol{u}_1[i]|$ descending, where $\boldsymbol{u}_1$ is the leading eigenvector of $\boldsymbol{G}$. The rule is a heuristic based on *dictionary coherence*, the degree to which the splats overlap one another in $L^2$: $\boldsymbol{u}_1$ is the unit coefficient vector whose splat combination $\sum_i \boldsymbol{u}_1[i]\phi_i$ has the largest squared $L^2$ norm, $\lambda_1$, and $|\boldsymbol{u}_1[i]|$ measures how much splat $i$ participates in that combination. It is a statement about the dictionary operator, not about the specific target $f = \sum_i \phi_i$: in the eigenbasis of $\boldsymbol{G}$ the energy of $f$ splits as $\|f\|^2 = \sum_\ell \lambda_\ell(\boldsymbol{u}_\ell^\top \mathbf{1})^2$, and the mode with the largest $\lambda_\ell$ need not carry the largest share $\lambda_\ell(\boldsymbol{u}_\ell^\top\mathbf{1})^2$ of $f$ (for $\boldsymbol{G} = \mathrm{diag}(3, [\begin{smallmatrix}1&1\\1&1\end{smallmatrix}])$ the eigenvalues are $3, 2, 0$ but the target shares are $3, 4, 0$). The Eckart–Young theorem [3] concerns low-rank approximation of the matrix $\boldsymbol{G}$ and does not supply a target-specific justification.

Table 1: Catalogue of ordering strategies. "Optimal at $k = 1$" indicates whether the strategy minimises $E_1$ (one-step greedy is by definition). "Submodular guarantee" refers to the $(1 - 1/e)$ bound from Theorem 4.5: only the matching-pursuit greedy attains it uniformly in $k$.

| **Strategy** | **Score / rule** | **Cost (precompute)** | **Optimal at $k = 1$** | **Submodular guarantee** |
|---|---|---|---|---|
| random | uniform($\pi$) | $O(N)$ | no | no |
| amplitude | sort by $a_i$ desc | $O(N \log N)$ | no | no |
| mass | sort by $a_i (2\pi)^{D/2} \lvert \boldsymbol{\Sigma}_i \rvert^{1/2}$ desc | $O(ND^3 + N \log N)$ | no | no |
| self-energy | sort by $\boldsymbol{G}_{ii} = \lVert \phi_i \rVert^2$ desc | $O(ND^3 + N \log N)$ | sometimes[a] | no |
| spectral | sort by $\lvert \boldsymbol{u}_1[i] \rvert$ desc, $\boldsymbol{u}_1 = \mathrm{argmax}_{\lVert \boldsymbol{u} \rVert = 1}\, \boldsymbol{u}^\top \boldsymbol{G} \boldsymbol{u}$ | Gram[b] + eigensolve[d] + $O(N \log N)$ | no | no |
| greedy / MP | $\arg\max_i\, 2\langle \phi_i, \boldsymbol{r}_S \rangle - \boldsymbol{G}_{ii}$ | Gram[b] + $O(N^2)$ ordering[c] | yes | yes $(1 - 1/e$ on $U)$ |

[a] Self-energy is optimal at $k = 1$ in the *orthogonal-dictionary* limit ($\boldsymbol{G}$ diagonal); for general overlapping splats it differs from greedy by the row-sum interaction correction (Remark 3.1).

[b] Dense Gram is $O(N^2)$ entries. The finite $(2.75\sigma)$ Mahalanobis support of Luxar splats, pruned here at the wider $3\sigma$, yields a support-pruned sparse surrogate with empirically $<1\%$ density on the $N = 4\,000$ subsamples of Section 8 (full-$N$ densities are not archived). It is computed in $O(N\bar{n})$, with $\bar{n}$ the mean number of candidate neighbours per splat, via neighbour-index pruning (Algorithm 4; a $k$-d tree in the experiments, a spatial hash in the shipped implementation). Its retained entries are the untruncated inner products (4), so it is an approximation to both the untruncated and the truncated objective (Section 8.3).

[c] The ordering loop is $O(N^2)$ for argmax scans on a dense Gram; with the sparse Gram, per-step updates touch only the non-zero entries of column $i^\star$.

[d] $O(t \cdot \mathrm{nnz}(\boldsymbol{G}))$ for $t$ Lanczos / power iterations to the requested tolerance on the sparse Gram; on a dense Gram a full eigendecomposition is $O(N^3)$. Spectral is therefore not an $O(N \log N)$ alternative: only amplitude, mass and self-energy are, for fixed $D$.

**(6) Greedy / matching pursuit.** At each step pick the splat $i \notin S$ that maximises the marginal energy gain

$$\Delta_i(S) \;=\; U(S \cup \{i\}) - U(S) \;=\; 2\,\langle \phi_i, \boldsymbol{r}_S \rangle_{L^2} \;-\; \lVert \phi_i \rVert_{L^2}^2, \tag{8}$$

where $\boldsymbol{r}_S = f - g_S$ is the residual after loading $S$. The rule is the natural analogue of *matching pursuit*, the greedy signal-approximation scheme that at each step adds the dictionary element most correlated with the residual [2, 9, 17], here in the fixed-coefficient setting where each splat enters with coefficient 1 and must eventually be loaded exactly once. Theorem 4.5 below shows that this rule captures at least $(1 - 1/e)$ of the optimal cumulative energy $U$ at every $k$; the corresponding statement for the residual $E_k$ is the affine inequality (12), not a multiplicative factor.

*Remark* 3.1 (Why $\boldsymbol{G}_{ii}$ is the correct first step). At $S = \emptyset$ the residual is $\boldsymbol{r}_S = f$, so $\langle \phi_i, \boldsymbol{r}_S \rangle = \sum_j \boldsymbol{G}_{ij}$ (the row-sum of $\boldsymbol{G}$). The greedy rule selects $\arg\max_i (2 \sum_j \boldsymbol{G}_{ij} - \boldsymbol{G}_{ii})$. In the *nearly-orthogonal* regime $\boldsymbol{G}_{ij} \approx 0$ for $i \neq j$, this reduces to $\arg\max_i \boldsymbol{G}_{ii}$, the self-energy ordering. For realistic, partially overlapping splats the greedy correction $2 \sum_{j \neq i} \boldsymbol{G}_{ij}$ favours splats located in dense regions, where adding one splat also reduces the residual amplitude near its neighbours.

### 4 Theory

#### 4.1 Closed-form quadratic-assignment AUC

A clean reformulation of the AUC objective expresses it as a quadratic form in the rank vector.

**Proposition 4.1** (AUC as a Gram-rank inner product)**.** *For any permutation $\pi$ with rank vector $\rho_i = \pi^{-1}(i)$,*

$$\sum_{k=0}^{N-1} E_k(\pi) \;=\; \sum_{i=1}^{N}\sum_{j=1}^{N} \boldsymbol{G}_{ij}\, \min(\rho_i,\, \rho_j). \tag{9}$$

*Proof.* By (5), $E_k = \sum_{i,j} \boldsymbol{G}_{ij}\, \mathbf{1}\{\rho_i > k\}\, \mathbf{1}\{\rho_j > k\}$. Summing over $k = 0, \ldots, N-1$ and exchanging summation order,

$$\sum_{k=0}^{N-1} E_k = \sum_{i,j} \boldsymbol{G}_{ij}\, \big|\{k : \rho_i > k \wedge \rho_j > k\}\big| = \sum_{i,j} \boldsymbol{G}_{ij}\, \min(\rho_i, \rho_j),$$

since the set of $k \in \{0, \ldots, N-1\}$ with $k < \min(\rho_i, \rho_j)$ has cardinality exactly $\min(\rho_i, \rho_j)$. □

*Remark* 4.2 (Quadratic-assignment hardness)*.* Equation (9) is an instance of the quadratic assignment problem (QAP), which is NP-hard in general [1, 15]. A direct combinatorial search is infeasible. The submodularity result of the next subsection provides a tractable approximation route.

#### 4.2 Submodularity and the greedy guarantee

**Theorem 4.3** (Monotone submodular utility)**.** *Let $\boldsymbol{G}$ be a symmetric Gram matrix with $\boldsymbol{G}_{ij} \geq 0$ for all $i, j$. Then the cumulative-energy utility $U : 2^{[N]} \to \mathbb{R}_{\geq 0}$ defined in (6) is monotone non-decreasing and submodular:*

$$\begin{aligned}
U(\emptyset) &= 0 && \textit{(normalisation)},\\
U(S) &\leq U(T) && \textit{for } S \subseteq T \textit{ (monotonicity)},\\
U(S \cup \{i\}) - U(S) &\geq U(T \cup \{i\}) - U(T) && \textit{for } S \subseteq T,\ i \notin T \textit{ (diminishing returns)}.
\end{aligned}$$

*Proof.* Direct expansion from (6) gives the marginal gain

$$\Delta_i(S) := U(S \cup \{i\}) - U(S) \;=\; \boldsymbol{G}_{ii} + 2 \sum_{j \notin S \cup \{i\}} \boldsymbol{G}_{ij}. \tag{10}$$

Monotonicity: every term on the right is non-negative since $\boldsymbol{G}_{ij} \geq 0$, so $\Delta_i(S) \geq 0$ for every $S$ and $i \notin S$. For $S \subseteq T$ write $T \setminus S = \{i_1, \ldots, i_m\}$ and $S_\ell = S \cup \{i_1, \ldots, i_\ell\}$; the difference telescopes as $U(T) - U(S) = \sum_{\ell=1}^{m} \Delta_{i_\ell}(S_{\ell-1}) \geq 0$. Diminishing returns: for $S \subseteq T$ with $i \notin T$,

$$\Delta_i(S) - \Delta_i(T) = 2 \sum_{j \in T \setminus S} \boldsymbol{G}_{ij} \;\geq\; 0,$$

which is non-negative entrywise. Hence $\Delta_i(S) \geq \Delta_i(T)$. □

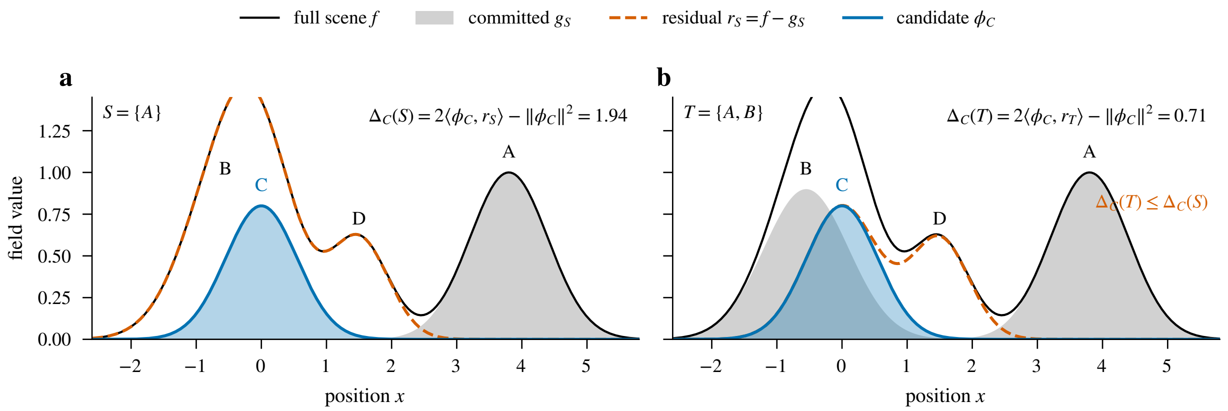

Figure 2: Schematic of diminishing returns (Theorem 4.3) as overlap, on a constructed one-dimensional mixture of four splats (no measured data). Both panels add the same candidate $\phi_C$ (blue); its gain is its correlation with the residual, $\Delta_C(S) = 2\langle \phi_C, \boldsymbol{r}_S \rangle - \|\phi_C\|^2$ (10), printed in each panel. (a) With $S = \{A\}$ committed (grey), the residual (dashed) still contains the neighbour $B$ under the candidate, and the gain is large. (b) With $T = \{A, B\}$ committed, the residual under the candidate has collapsed and the same splat gains less: $\Delta_C(T) \leq \Delta_C(S)$.

Figure 2 shows the inequality as overlap: the same candidate gains less once a committed splat covers part of its residual.

*Remark* 4.4 (Why non-negative Gram entries are essential). For Gaussian splats with positive amplitudes, all $\boldsymbol{G}_{ij}$ are strictly positive (equation (4)). If the dictionary were allowed signed amplitudes (residuals or signed splats), $\boldsymbol{G}$ could have negative entries and submodularity could fail. The proof of Theorem 4.3 pinpoints non-negativity of $\boldsymbol{G}$ as the structural property that makes greedy ordering tractable.

**Theorem 4.5** ($(1 - 1/e)$ greedy guarantee, uniform in $k$). *Let $S_k^\star = \arg\max_{|S|=k} U(S)$ be the size-$k$ optimum and let $S_k^{\mathrm{g}}$ be the prefix of size $k$ produced by greedy (8). Because $U$ is monotone, $S_k^\star$ also maximises $U$ over all $S$ with $|S| \leq k$: any smaller set can be extended to size $k$ without lowering $U$. Then for every $k = 1, \ldots, N$,*

$$U(S_k^{\mathrm{g}}) \;\geq\; (1 - 1/e)\, U(S_k^\star), \tag{11}$$

*or, equivalently, in residual energy*

$$E_k(\mathrm{g}) - E_k^\star \;\leq\; \frac{1}{e}\left(\|f\|^2 - E_k^\star\right). \tag{12}$$

*Proof.* Theorem 4.3 verifies the hypotheses of the Nemhauser–Wolsey–Fisher theorem [12], which states that the standard greedy algorithm on a monotone submodular function under the cardinality constraint $|S| \leq k$ achieves at least $(1 - 1/e)$ of the optimum over that constraint, which by the monotonicity noted in the statement equals $U(S_k^\star)$. The greedy ordering is exactly that algorithm at every $k$ simultaneously, because it grows the prefix by one element and never revisits decisions. □

*Remark* 4.6 (A single ordering, all $k$; utility, not residual). The strength of Theorem 4.5 is that one

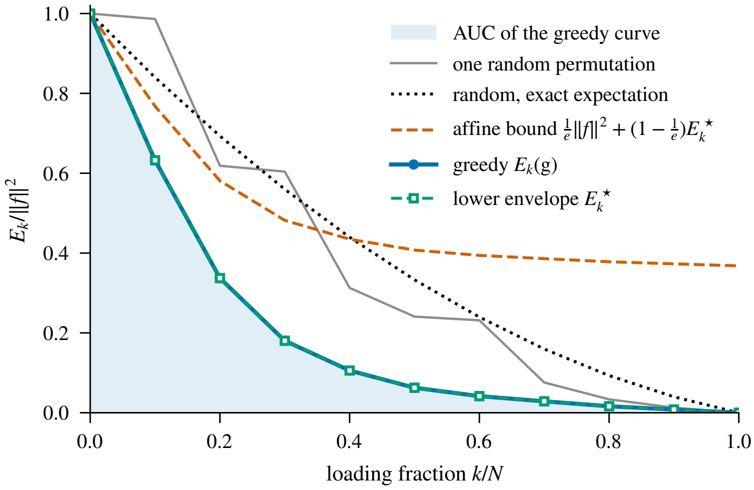

Figure 3: Schematic of the loading-curve objects of Theorem 4.5 and Corollary 4.7, on a constructed instance of $N = 10$ one-dimensional splats (no measured data). The greedy curve $E_k(\mathrm{g})/\|f\|^2$ (blue, its trapezoidal AUC shaded) coincides with the pointwise-optimal lower envelope $E_k^\star$ (green squares) at every $k$ on this instance; in general the envelope is only a floor that no single permutation need attain. The affine bound $\frac{1}{e}\|f\|^2 + (1-\frac{1}{e})E_k^\star$ of (12) (vermilion, dashed) holds at every $k$ and is loose: it ends at $\|f\|^2/e$ where every curve ends at zero. The exact expectation of a uniformly random permutation (7) (dotted) bends because the splats of this instance overlap heavily; one random draw (grey) wanders around it.

greedy permutation gives a $(1-1/e)$-approximation of the optimal *utility* for every prefix length $k$ at once. The factor does not transfer to the residual: an affine reparametrisation $E_k = \|f\|^2 - U(S_k)$ preserves exact minimisers but not multiplicative approximation ratios, and a multiplicative bound $E_k(\mathrm{g}) \le c\, E_k^\star$ with $c < 1$ would even contradict optimality. The residual statement is the additive inequality (12): the greedy excess over the size-$k$ optimum is at most $1/e$ of the energy that optimum has captured. The greedy curve is much closer to the exhaustive optimum (Section 8.1: ratio $\ge 0.9992$ on $N = 12$ densely-overlapping instances) than the worst-case $1 - 1/e \approx 0.632$ floor would suggest, which mirrors the empirical behaviour observed across submodular optimisation [6].

Figure 3 draws these objects for one constructed instance: the greedy curve and its AUC, the pointwise-optimal lower envelope it need not attain, the affine bound (12) and the exact expectation of a random ordering (7).

**Corollary 4.7** (Affine AUC bound from the lower envelope)**.** *Let* AUC$^\star$ *denote the trapezoidal AUC* (3) *evaluated on the* pointwise-optimal lower envelope $E_k^\star = \min_{|S|=k}(\|f\|^2 - U(S))$ *of size-$k$ optima. The envelope bounds every feasible AUC from below, but it need not be attained by any single permutation, because the size-$k$ optima need not be nested; it can lie strictly below the best feasible AUC. Then*

$$\mathrm{AUC}(\mathrm{g}) \;\le\; \tfrac{1}{e}\Big(1 - \tfrac{1}{2N}\Big) + \big(1 - \tfrac{1}{e}\big)\ \mathrm{AUC}^\star\,.$$

*Proof.* By (12), $E_k(\mathrm{g}) \le \frac{1}{e}\|f\|^2 + (1-\frac{1}{e})E_k^\star$ for every $k$ (at $k = 0$ both sides equal $\|f\|^2$; at $k = N$ the left side is 0). Averaging the trapezoids $(E_k + E_{k+1})/2$ over $k = 0, \dots, N-1$ and dividing by $\|f\|^2$ gives $\mathrm{AUC}(\mathrm{g}) \le \frac{1}{e}(1 - \frac{1}{2N}) + (1 - \frac{1}{e})\,\mathrm{AUC}^\star$, the constant being the trapezoidal AUC of the flat curve $E_k \equiv \|f\|^2$ for $k < N$, $E_N = 0$. In the plain-prefix-average convention the constant is $1/e$; the bound is affine in AUC$^\star$ in either convention. □

##### 4.3 Lazy greedy, complexity, and incremental updates

The naive greedy of (8) requires evaluating $\Delta_i(S_k)$ for all $N - k$ candidates at each step, giving an $O(N^2)$ work in the ordering loop on top of the $O(N^2)$ Gram matrix construction.

**Incremental residual inner products.** The marginal gain (10) can be rewritten using the running tail-row-sum $\beta_i^{(k)} = \sum_{j \notin S_k} \boldsymbol{G}_{ij}$:

$$\Delta_i(S_k) \;=\; 2\,\beta_i^{(k)} - \boldsymbol{G}_{ii} \quad \text{for } i \notin S_k. \tag{13}$$

After choosing $i^\star$, all $\beta_i^{(k+1)} = \beta_i^{(k)} - \boldsymbol{G}_{i,i^\star}$. The update is one column read from $\boldsymbol{G}$ per step, $O(N^2)$ total. See Algorithm 1 for pseudocode.

**Lazy greedy.** Since $U$ is submodular, marginal gains never increase as $S$ grows [10]: $\Delta_i(S_{k+1}) \leq \Delta_i(S_k)$ for any $i \notin S_{k+1}$. Maintaining a max-heap of upper-bound estimates and lazily re-evaluating only the top entry until it is up-to-date (Algorithm 2) replaces the $O(N)$ argmax scan of each step by a data-dependent number of heap operations. It does not change the asymptotic total: the tail-row-sum update $\beta_j \leftarrow \beta_j - \boldsymbol{G}_{j,i^\star}$ is still performed at every step, which is $\Theta(N^2)$ work on a dense Gram (or $O(\text{nnz}(\boldsymbol{G}))$ in total on the sparse surrogate), and in the worst case the number of stale re-evaluations is itself quadratic, giving $O(N^2 \log N)$ heap overhead, not a saving. Any advantage of the lazy variant is therefore empirical and instance-dependent; no timings for it are reported here. The shipped implementation uses the plain scan on a dense Gram below a size threshold and the lazy heap on the sparse Gram above it (Section 6.3); a vectorised C/CUDA implementation would change the constants. The stochastic-greedy variant of Mirzasoleiman et al. [11] evaluates $O(N \log(1/\epsilon))$ marginal gains in an oracle model and guarantees, for a *chosen* cardinality $k$, an *expected* utility of at least $(1 - 1/e - \epsilon)$ of the optimum. Its sample size depends on the target $k$, so the cited result does not by itself certify one streaming order at every prefix, and each oracle call still costs a Gram row or its sparse equivalent.

#### 5 Lower Bounds and Achievability

##### 5.1 Rayleigh–Ritz lower bound

**Proposition 5.1** (Rayleigh-quotient lower bound on prefix error)**.** *Let $\lambda_{\min}(\boldsymbol{G})$ denote the smallest eigenvalue of the splat Gram matrix. For any permutation $\pi$ and any $k \leq N$,*

$$E_k(\pi) \;\geq\; (N - k)\,\lambda_{\min}(\boldsymbol{G}). \tag{14}$$

*Proof.* By (5), $E_k(\pi) = \mathbf{1}_{S_k^c}^\top \boldsymbol{G}\,\mathbf{1}_{S_k^c}$ with $\|\mathbf{1}_{S_k^c}\|^2 = N - k$. The Rayleigh–Ritz inequality $\boldsymbol{v}^\top \boldsymbol{G}\,\boldsymbol{v} \geq \lambda_{\min}(\boldsymbol{G})\,\|\boldsymbol{v}\|^2$ then gives the bound. □

*Remark* 5.2 (The stronger feature-space bound is false). Define the feature-space coordinates of $f = \sum_i \phi_i$ as $\tilde{c}_\ell^2 := \lambda_\ell\,(\boldsymbol{u}_\ell^\top \mathbf{1})^2$ (so that $\|f\|_{L^2}^2 = \sum_\ell \tilde{c}_\ell^2$ in the eigenbasis of $\boldsymbol{G}$), and let $\tilde{c}_{(1)}^2 \geq \tilde{c}_{(2)}^2 \geq \cdots \geq \tilde{c}_{(N)}^2$ be their order statistics. A tempting strengthening of (14) would be

$$E_k(\pi) \;\geq\; \sum_{\ell=k+1}^{N} \tilde{c}_{(\ell)}^2 \qquad \text{(false in general)}, \tag{15}$$

which would be tighter whenever the small eigenvalues of $\boldsymbol{G}$ are not orthogonal to $\mathbf{1}$. It does not hold, even inside the model of this document (positive amplitudes, hence a strictly positive Gram). An explicit counterexample is $N = 7$ one-dimensional Gaussians with centres $\mu_i$, standard deviations $\sigma_i$ and amplitudes $a_i$,

$$\begin{aligned}
\mu &= (-0.4896,\ 1.0938,\ -2.2895,\ 2.4337,\ 2.5831,\ -3.8398,\ 3.4238),\\
\sigma &= (0.2885,\ 0.1402,\ 0.9568,\ 0.6459,\ 0.4574,\ 0.1769,\ 0.5311),\\
a &= (0.3948,\ 0.164,\ 0.5017,\ 4.711,\ 0.7183,\ 0.4635,\ 1.2077).
\end{aligned}$$

Every Gram entry is positive, $\lambda_{\min} = 6.16 \times 10^{-3}$, and the prefix that leaves only the second splat unloaded ($k = 6$) has $E_6 = \boldsymbol{G}_{22} = 6.68 \times 10^{-3}$. The right-hand side of (15), however, is $\tilde{c}_{(7)}^2 = 7.45 \times 10^{-3}$, a violation of 11.4% of the actual residual (Fig. 4). Multiplying every splat by the same transverse Gaussian factors embeds the example in 3D and rescales every Gram entry by one positive constant, so the violation persists in the dimension Luxar renders. The structural reason is that a prefix sum has coefficients fixed to $\{0,1\}^N$, not free real numbers, so neither an Eckart–Young argument (whose single-vector bound collapses to zero, since any 1-dimensional subspace containing $f$ gives zero residual) nor an eigen-coordinate truncation describes it. The Rayleigh–Ritz bound (14) remains the bound we rely on. A second valid bound follows from positivity alone: dropping the positive cross terms in (5) gives $E_k(\pi) \geq \sum_{i \notin S_k} \boldsymbol{G}_{ii} \geq$ the sum of the $N - k$ smallest diagonal entries, which is attained with equality in the counterexample above.

#### 5.2 Achievability gap

The Rayleigh–Ritz bound (14) is governed entirely by $\lambda_{\min}(\boldsymbol{G})$ and by the norm of the tail indicator $\mathbf{1}_{S_k^c}$, so it can be weak when the splat self-energies are unequal (the smallest eigenvalue then sits far below the energy of a typical remaining splat) or when the tail indicator has little projection on the eigenspace of $\lambda_{\min}$. The bound depends on the absolute size of $\lambda_{\min}$ and not on the conditioning of $\boldsymbol{G}$: rescaling every amplitude by a common factor rescales every eigenvalue and both sides of (14) alike while leaving the condition number unchanged. In the nearly-orthogonal, equal-energy limit $\boldsymbol{G} \approx c\,I$ the bound is tight ($E_k = (N-k)\,c$ for every permutation). The gap between the bound and the greedy / optimal prefix curve is the price paid for fixed-coefficient streaming: each loaded splat enters with coefficient 1, not an optimal real coefficient, so even the size-$k$ optimum can lie far above any spectrum-only bound that ignores how $\mathbf{1}_S$ aligns with the eigenvectors of $\boldsymbol{G}$. A substitutive coarse level [8] can improve on any fixed-coefficient subset of the same size by allowing free, rescaled representatives. Proposition 5.1 is a statement about prefix sums only: no analogous lower bound is known for substitutive levels, whose free Gaussians can

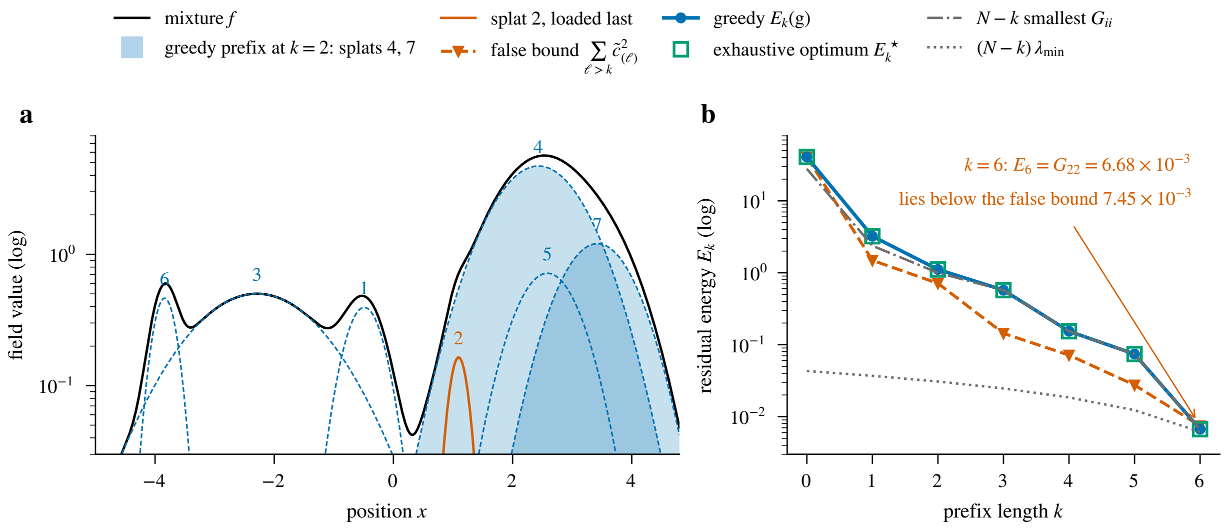

Figure 4: The $N = 7$ counterexample of Remark 5.2, recomputed from the centres, standard deviations and amplitudes listed there. (a) The mixture $f$ (black) and its components on a logarithmic scale, numbered as in the text; the two splats greedy loads first ($k = 2$, filled) are also the exhaustive size-2 optimum, and splat 2 (vermilion) is loaded last. (b) Residual energy $E_k$ for $k = 0, \ldots, 6$ ($E_7 = 0$ is off the logarithmic axis): the greedy curve coincides with the exhaustive optimum $E_k^\star$ at every $k$ on this instance; the Rayleigh–Ritz bound $(N-k)\lambda_{\min}$ (14) and the sum of the $N - k$ smallest diagonal entries stay at or below it; the eigen-coordinate tail $\sum_{\ell>k} \tilde{c}^2_{(\ell)}$ of (15) crosses above it at $k = 6$.

land four orders of magnitude below the corresponding eigen-coordinate tail (the section on lower bounds of Supplementary Document 8 [8], which compares those representatives numerically and claims no analytical lower bound for them).

### 6 Connections

#### 6.1 PRoGS, LapisGS, and rendering-based importance

PRoGS [19] orders the splats of a 3D Gaussian Splatting scene (3DGS, the photogrammetric radiance-field representation) for progressive rendering by computing for each Gaussian a contribution score $B_i = \sum_v \sum_p T_i(v,p)\alpha_i(v,p)$ summed over training views $v$ and pixels $p$, where $T_i$ is the transmittance and $\alpha_i$ the opacity from the alpha-compositing equation. The score is a viewpoint- and rendering-conditional analogue of our *integral mass* ordering: it weighs each splat by an effective contribution to the rendered image, not to the underlying scalar field. LapisGS [16] takes a different, optimisation-based route: it trains a layered representation in which each successive layer adds new Gaussians while the geometry of the earlier layers is frozen and only their opacities are re-optimised, prunes with a fixed opacity threshold, and interpolates opacities between adjacent layers at render time; the number of layers is a free choice, not two. Neither method orders a fixed dictionary by an analytical criterion, but both belong to the broad "importance-based" family in the sense that rendered contribution decides what is sent first. Our results show that for the analytical $L^2$ objective on the underlying scalar field, the residual-correlation greedy ordering captures at least $(1-1/e)$ of the optimal cumulative energy at every prefix (Theorem 4.5) and had the lowest AUC among the orderings we tested (Section 8); global optimality of greedy is not claimed. Rendering-conditional importance scores remain valuable when the rendering pipeline applies a viewpoint-dependent filter that $L^2$ does not capture; whether PRoGS- or LapisGS-style scores can outperform residual-correlation greedy on a rendering-aware metric is an open empirical question.

##### 6.2 Hierarchical 3DGS and other LOD methods

Two families of prior work reduce a splat set offline. Hierarchical 3DGS [5] and Octree-GS [14] build a *hierarchy* of merged representatives whose level is selected dynamically at render time from the view; LightGaussian [4], Reduced 3DGS [13] and the Luxar `cull_by_contribution` routine (`luxar.gsplats.culling`) perform *static* compression (pruning, quantisation, and distillation of the spherical-harmonic (SH) view-dependent colour coefficients) to a single reduced set. The additive ordering is complementary to both: a coarse level or a reduced set provides anchored fidelity at a fixed budget, and the additive ordering provides the streaming refinement *within* a level. In a deployed Luxar pipeline the natural composition is to (i) page in the coarsest LOD, (ii) within each level, stream splats in the selected additive order (Section 6.3).

##### 6.3 Luxar implementation

This subsection describes the shipped implementation (`luxar.gsplats.lod.additive` and the viewer's LOD registry), so that the theorems above can be connected to what is stored and rendered. Throughout, an *alpha-effective amplitude* is the stored amplitude multiplied by the splat's per-splat opacity (the alpha channel of an RGBA colour, 1 when absent), the amplitude the renderer actually draws.

**Ordering.** `--add-method auto` resolves to greedy for leaves of at most 5 000 splats and to self-energy above that (Section 1); `greedy`, `self_energy`, `mass`, `amplitude`, `spectral`, `random` and `radial` can be selected explicitly. Greedy and spectral build a sparse Gram by pruning pairs whose max-axis truncation balls (radius $R_i = \tau\sqrt{\lambda_{\max}(\mathbf{\Sigma}_i)}$, the criterion of Algorithm 4, with $\tau$ the dataset's own `truncation_radius`, canonically 2.75 standard deviations) do not overlap, using a spatial hash as the neighbour index of Algorithm 4 (the experiments use a $k$-d tree). Retained pairs carry the untruncated inner product (4) evaluated on alpha-effective amplitudes, so the production Gram is a support-pruned surrogate of the same construction as in the experiments (Section 8.3), pruned at $2.75\sigma$ instead of the $3\sigma$ used there and evaluated on alpha-effective, not stored, amplitudes. Greedy runs Algorithm 1 on a dense copy of that Gram up to a size threshold (default 2 000) and Algorithm 2 on the sparse matrix above it. `radial` is a *reveal* ordering by distance from a centre and makes no energy claim.

**Storage.** In the standalone gsplats format a ladder is stored as consecutive, disjoint `additive_`$i$ rungs of the chosen ordering, so paging in rungs $0,\ldots,i$ reconstructs the length-$k_i$ prefix; the rungs follow whichever ordering was selected, greedy or not. Each splat carries its centre, its

Cholesky factors and its amplitude. Rung sizes come from equal counts, a `stream:C` geometric schedule, equal self-energy shares (`equi-energy:n`), or explicit counts or energy fractions. On the command line the mechanism is exposed as `gsplat lod --recipe stream` (or `gsplat additive` to add a ladder to an existing tree); obtaining the greedy ordering of this document on a leaf of more than 5 000 splats requires `--add-method greedy` explicitly, because the default `auto` resolves to greedy only up to that size and to the cheaper `self_energy` ordering above it.

**Quality stamps.** Each rung $i$ (sub-LOD `additive_`$i$) carries in its `lod_stats` the attribute `energy_fraction_cum`, a cumulative *self-energy* fraction

$$e(k) \;=\; \frac{\sum_{j\le k} \boldsymbol{G}_{\pi(j)\pi(j)}}{\sum_{i=1}^{N} \boldsymbol{G}_{ii}},$$

evaluated at the rung's last splat from the $O(N)$ per-splat score $a_i^2\,|\boldsymbol{\Sigma}_i|^{1/2}$ on alpha-effective amplitudes (the shared $\pi^{D/2}$ of the true self-energy cancels in the fraction). It is the diagonal (orthogonal-limit) approximation to the utility fraction $U(S_k)/\|f\|^2$: it omits the cross terms of (6), and it is computed from the self-energy scores for every ordering, greedy and spectral included, even though those two build the sparse Gram. The absolute weight `reference_energy` $= \sum_i a_i^2\,\pi^{D/2}\,|\boldsymbol{\Sigma}_i|^{1/2}$, with the constant restored, lives one level up, in the `level_stats` of the leaf that owns the ladder (a substitutive level in the writer's terms), where it aggregates the fractions over a partition (the parts of a `kind=partition` group); it is written with `setdefault`, so a weight already stamped by a substitutive build (the group-consistent finest-content energy) wins over the ladder's own total. Rung boundaries and stamps can disagree: an `energy:` list of fractions is cut against the sparse-Gram residual-energy curve, i.e. the surrogate utility fraction, whenever greedy or spectral has built that Gram, and against the self-energy cumulative otherwise (`equi-energy:n` always uses the self-energy cumulative), while the $e(k)$ stamp is the self-energy fraction in every case. Reveal ladders (`radial`) carry neither attribute.

**Viewer behaviour.** The viewer uses the stamps in two ways, both outside the fixed-coefficient model of this document and both acting on the *committed prefix (committed count)* of a leaf, the rungs already on screen. (i) *Brightness compensation*: while a ladder inside a *`kind=lod` group* is incomplete, the registry multiplies the opacity of each of its leaves whose blend mode is additive, luminous or volumetric by $1/\max(e(k), 0.1)$, a boost of at most $10\times$, on by default and disabled by `?noLodEnergy`. Because the multiplier rescales the loaded prefix, the rendered field is no longer the plain prefix sum $g_k$; the $(1-1/e)$ guarantee and the AUC results concern $g_k$ with unit coefficients and say nothing about the compensated image. The multiplier is a heuristic: $e(k)$ is a fraction of *squared* amplitudes, whereas rendered brightness in additive compositing is linear in amplitude, so $1/e(k)$ conserves neither total brightness nor $L^2$ energy in general. Two disjoint splats of amplitudes 2 and 1 give $e = 4/5$ after the first, a multiplier of 1.25, and a compensated mass of 2.5 against a full mass of 3. A bare `stream` leaf outside any LOD group is rescaled by the density guard (`scene/density-guard.ts`), not by the registry. The guard thins any over-dense blendable node to a `keep` fraction of its elements and restores the aggregate brightness through the same

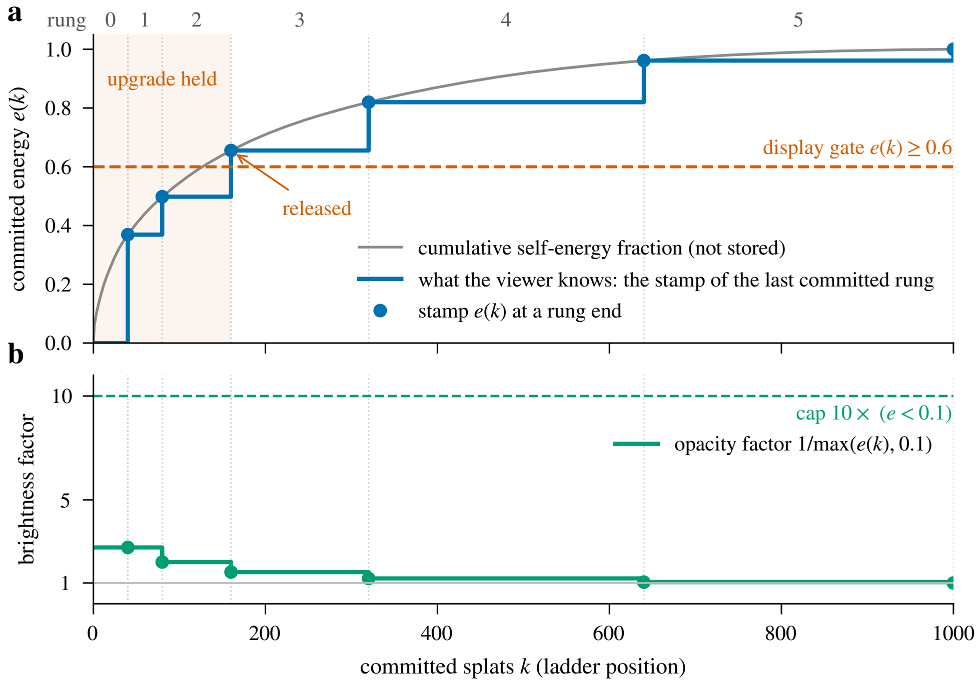

Figure 5: Schematic of a shipped ladder and what the viewer does with it, on a constructed ladder of $N = 1\,000$ splats in self-energy order cut by a `stream:C` geometric rung schedule with $C = 40$ (no measured data). (a) The cumulative self-energy fraction along the ordering (grey) and the stamps $e(k)$ written at each rung's last splat (dots); the viewer knows only the staircase (blue). The display gate holds an upgrade to this level until its committed prefix reaches $e(k) \geq 0.6$, here the end of rung 2. (b) The brightness-compensation factor $1/\max(e(k), 0.1)$ applied to the committed prefix, at most 10×; the cap acts only while $e(k) < 0.1$, which this ladder never reaches.

fade routine (`applyLodFade`), which multiplies in the $1/\max(e(k), 0.1)$ factor whenever it runs. The leaf's committed prefix is therefore compensated only at the moments the guard changes that node's keep step. The guard is on by default and disabled by `?noDensityGuard`; `?noLodEnergy` disables the energy factor in both paths. (ii) *display gate*: when the viewer upgrades to a finer substitutive level whose ladder is still streaming, it keeps the previously displayed level on screen until the finer level's committed prefix reaches $e(k) \geq 0.6$ (or its ladder completes, or its committed count passes the held level's). The threshold applies to upgrades only and to stamped datasets only. Figure 5 draws the stamps, the factor and the gate along one constructed ladder.

**Offline audit.** The metric `rel_l2` $= \|f - g_k\|/\|f\|$ in the gsplats metrics module is evaluated on a rasterised voxel grid, not analytically, and is an offline audit tool, not a readout the viewer computes.

### 7 Algorithms

We collect the four algorithms used in the experiments. Algorithm 1 is the principal $O(N^2)$ greedy / matching-pursuit ordering, with the marginal-gain update of (13). Algorithm 2 is the lazy-greedy variant that uses submodularity (Theorem 4.3) to skip stale heap entries while keeping the same per-step column update, so its total work is still $\Theta(N^2)$ on a dense Gram (Section 4). Algorithm 3 collects the three non-Gram-based scoring rules (peak amplitude, integral mass, $L^2$ self-energy), each computable in $O(N)$ from the raw splat parameters and sorted in $O(N \log N)$. Algorithm 4 is the $3\sigma$-truncation pruning routine that produces a sparse Gram matrix for use in Algorithm 1 at large $N$.

**Algorithm 1:** Greedy / matching-pursuit ordering ($O(N^2)$ post-Gram).

**Input:** Splats $\{\phi_i\}_{i=1}^N$ and Gram matrix $\boldsymbol{G}$.
**Output:** Permutation $\pi$ of $\{1, \dots, N\}$.

$\beta_i \leftarrow \sum_{j=1}^N \boldsymbol{G}_{ij}$ for all $i$ `// tail-row-sum init`
chosen$_i$ ← false for all $i$
**for** $k = 1$ **to** $N$ **do**
    $\Delta_i \leftarrow 2\beta_i - \boldsymbol{G}_{ii}$ for all $i$ with chosen$_i$ = false
    $i^\star \leftarrow \arg\max_{i:\, \neg\text{chosen}_i} \Delta_i$
    $\pi(k) \leftarrow i^\star$
    chosen$_{i^\star}$ ← true
    $\beta_j \leftarrow \beta_j - \boldsymbol{G}_{j,i^\star}$ for all $j$ `// O(N) column read`
**return** $\pi$

**Algorithm 2:** Lazy greedy with submodular upper bounds.

**Input:** Splats $\{\phi_i\}_{i=1}^N$ and Gram matrix $\boldsymbol{G}$.
**Output:** Permutation $\pi$.

$\beta_i \leftarrow \sum_{j=1}^N \boldsymbol{G}_{ij}$, chosen$_i$ ← false for all $i$ `// same init as Algorithm 1`
Build a max-heap $H$ of entries $(i, \Delta_i^{(0)}, \text{stale}_i{=}0)$ keyed on $\Delta_i^{(0)} = 2\beta_i - \boldsymbol{G}_{ii}$ for all $i$
  `// ties broken by smaller i`
**for** $k = 1$ **to** $N$ **do**
    **repeat**
        $(i, \Delta_i^{\text{ub}}, \text{stale}_i) \leftarrow$ HeapPopMax($H$)
        **if** chosen$_i$ **then**
            continue
        **else if** stale$_i < k$ **then**
            $\Delta_i^{\text{cur}} \leftarrow 2\beta_i - \boldsymbol{G}_{ii}$
            HeapPush($H, (i, \Delta_i^{\text{cur}}, k)$)
    **until** stale$_i = k$
    $\pi(k) \leftarrow i$; chosen$_i$ ← true; $\beta_j \leftarrow \beta_j - \boldsymbol{G}_{j,i}$ for all $j$
**return** $\pi$

**Algorithm 3:** Cheap orderings (no Gram).

**Input:** Splats $\{\phi_i\}_{i=1}^N$, type $\in$ {amplitude, mass, self-energy}.
**Output:** Permutation $\pi$.

**if** *type* = *amplitude* **then**
  $s_i \leftarrow a_i$
**else if** *type* = *mass* **then**
  $s_i \leftarrow a_i (2\pi)^{D/2} |\mathbf{\Sigma}_i|^{1/2}$
**else if** *type* = *self-energy* **then**
  $s_i \leftarrow a_i^2 \pi^{D/2} |\mathbf{\Sigma}_i|^{1/2}$
$\pi \leftarrow$ argsort of $s_i$ in decreasing order
**return** $\pi$

**Algorithm 4:** Support-pruned sparse Gram surrogate via $3\sigma$ truncation pruning.

**Input:** Splats $\{\phi_i\}_{i=1}^N$, truncation multiplier $\tau$ in standard deviations (default 3).
**Output:** Sparse symmetric matrix $\boldsymbol{G} \in \mathbb{R}^{N \times N}$ approximating the Gram matrix.

$R_i \leftarrow \tau \sqrt{\lambda_{\max}(\mathbf{\Sigma}_i)}$ for all $i$ `// max-axis truncation radius`
Build a neighbour index on $\{\boldsymbol{\mu}_i\}$ (a $k$-d tree in the experiment code of this document; a spatial hash in `luxar.gsplats.lod.additive`)
**for** $i = 1$ **to** $N$ **do**
  Query the index for $\{j : \|\boldsymbol{\mu}_i - \boldsymbol{\mu}_j\| \le R_i + R_{\max}\}$, with $R_{\max} = \max_j R_j$
  **for** *each candidate* $j$ *with* $j \ge i$ **do**
    **if** $\|\boldsymbol{\mu}_i - \boldsymbol{\mu}_j\| \le R_i + R_j$ **then**
      $\boldsymbol{G}_{ij} \leftarrow K_{ij}$ via (4); $\boldsymbol{G}_{ji} \leftarrow \boldsymbol{G}_{ij}$
**return** $\boldsymbol{G}$ *as a CSR sparse matrix*

**Sparse greedy.** Algorithm 4 keeps a pair only if the centre distance is at most the sum of the two support radii $R_i + R_j$, and stores for the kept pairs the *untruncated* inner product (4). The result is a surrogate of the Gram matrix of either field: relative to the untruncated Gaussians of (1) it drops small positive entries (every $K_{ij}$ is strictly positive). Relative to the finitely supported splats the renderer draws, it keeps untruncated values for pairs whose truncated supports may be disjoint (two unit-width splats at distance 5.8 have disjoint $2.75\sigma$ supports yet are retained by the $3\sigma$ prune with $K_{ij} = \sqrt{\pi}\, e^{-5.8^2/4} \approx 3.9 \times 10^{-4}$); Fig. 6 draws which pairs the prune keeps. The retained entries are non-negative, so Theorem 4.3 and Theorem 4.5 apply verbatim to the surrogate utility; they do not make it the exact $L^2$ utility of either field, and the surrogate's error against the dense analytical Gram on the real datasets is not quantified here. Algorithm 1 extends to the sparse surrogate verbatim: the only operation that touches $\boldsymbol{G}$ is the column read $\beta_j \leftarrow \beta_j - \boldsymbol{G}_{j,i^\star}$, which on a CSR/CSC representation costs only the $|\{j : \boldsymbol{G}_{j,i^\star} \neq 0\}|$ non-zero entries. The argmax over candidates is dominated by the $O(N)$ scan of $\beta$, not by Gram access, so per-step cost is $O(N)$ (argmax) plus $O(\text{nnz of column } i^\star)$ (update). Memory drops from $O(N^2)$ for the dense Gram to $O(\text{nnz}(\boldsymbol{G}))$ for the sparse one, which is a substantial saving on real datasets

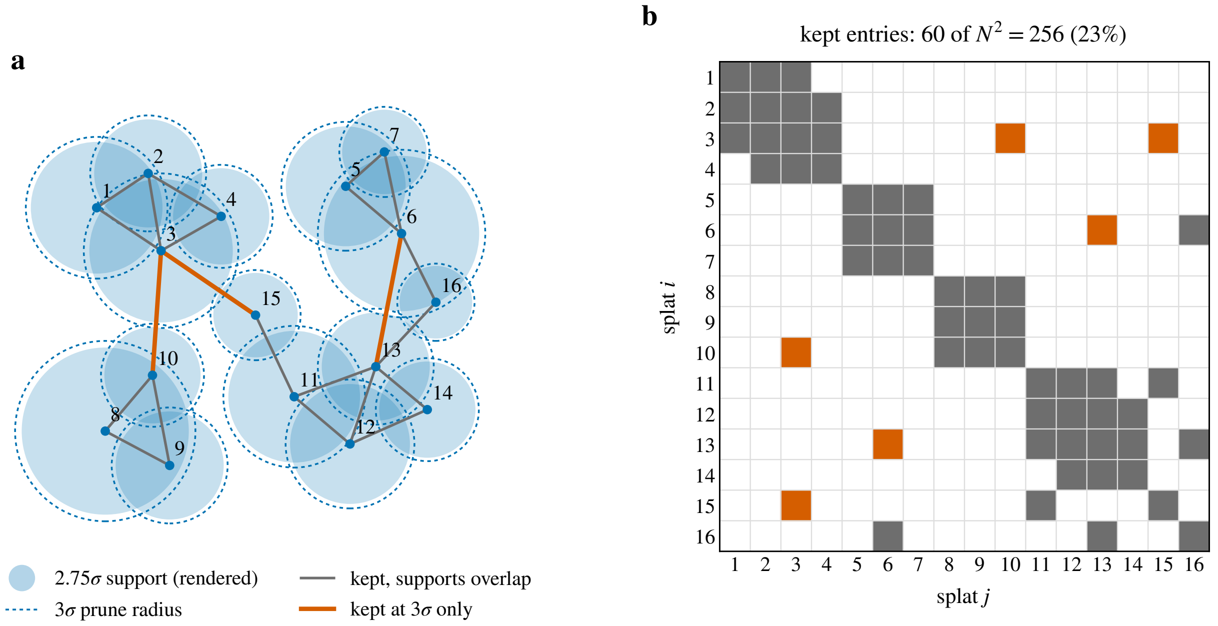

Figure 6: Schematic of the support-pruned surrogate (Algorithm 4) on a constructed layout of 16 isotropic two-dimensional splats (no measured data). (a) The filled disc is each splat's $2.75\sigma$ support, the finite Mahalanobis support the renderer draws; the dashed circle is its $3\sigma$ prune radius. A pair enters the surrogate when its $3\sigma$ discs overlap: a grey chord where the rendered supports overlap too, a vermilion chord for a pair kept at $3\sigma$ whose rendered supports are already disjoint and which therefore carries a small positive untruncated entry (4). (b) The resulting sparsity pattern of the $N \times N$ surrogate, same colours; the diagonal is always kept. On the $N = 4\,000$ subsamples of Section 8.3 the kept fraction is 0.04 to 0.84%.

(Section 8, Experiment C).

### 8 Numerical Experiments

We run three experiments: (A) verifies the theory on synthetic instances; (B) compares the six orderings on synthetic mixtures; (C) compares them on four real Luxar gsplat datasets. The experiment code accompanies this document; figures regenerate deterministically from a fixed RNG seed.

#### 8.1 Experiment A: theory verification

**Setup.** The experiment has two parts. (i) On $N = 64$ random anisotropic 2D splats with centres drawn uniformly in a box of side 1.5, run greedy and verify that the loading curve $E_k$ is monotone non-increasing and that the closed-form submodularity inequality holds on 24 random subset pairs; repeat for 16 trials. (ii) On small densely-overlapping instances, $N = 12$ in a box of side 0.4, enumerate all $\binom{N}{k}$ subsets to build the lower envelope $E_k^\star = \min_{|S|=k} \|f - g_S\|^2$; the small box forces heavy splat overlap, the regime in which greedy's locally-myopic choices are most likely to lag the global optimum. Then compare the greedy prefix utility $U_k^{\mathrm{g}} = \|f\|^2 - E_k(\mathrm{g})$ to $U_k^\star = \|f\|^2 - E_k^\star$ across 16 trials.

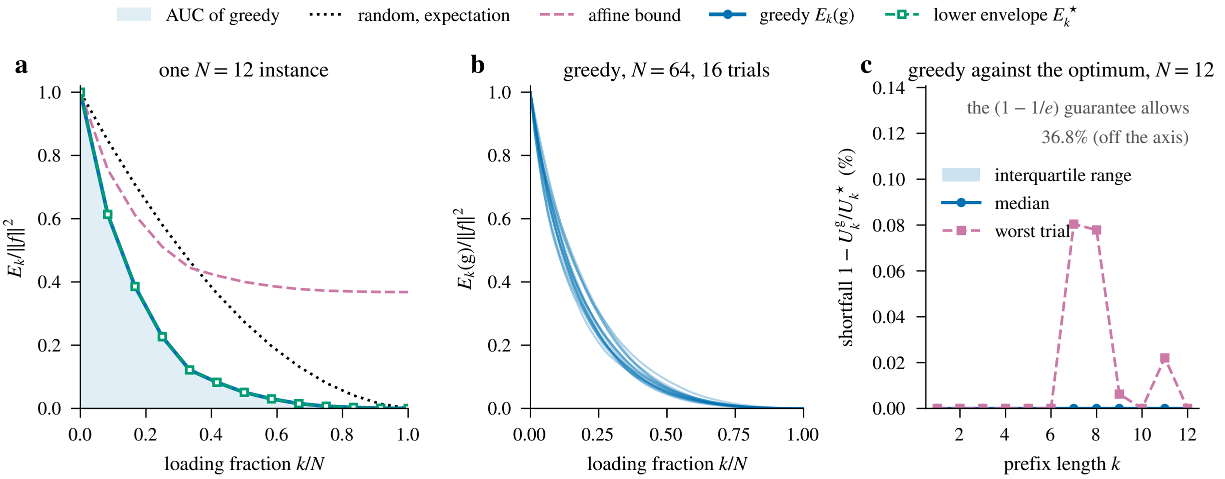

Figure 7: Experiment A: theory verification. (a) One of the $N = 12$ densely overlapping instances (box side 0.4): the greedy loading curve with its AUC shaded, the exhaustively enumerated lower envelope $E_k^\star$, the affine bound (12) and the exact expectation of a random permutation (7). (b) Greedy loading curves on 16 synthetic 2D mixtures ($N = 64$, centres uniform in a box of side 1.5), all non-increasing, as any ordering's must be under a non-negative Gram (a check of the code, not of the theory). (c) Shortfall of greedy against the exhaustive optimum, $1 - U_k^{\text{g}}/U_k^\star$, at every prefix length of the $N = 12$ instances: median, interquartile range and worst trial. The worst case is 0.08% (ratio 0.9992); the $1/e \approx 36.8\%$ that Theorem 4.5 allows is off the axis.

**Result.** Figure 7(b) shows $E_k(\text{g})/\|f\|^2$ for all 16 trials. All 16 curves are non-increasing, which is a sanity check of the loading-curve code and not a test of the theory: by (10) every step of every permutation removes $\boldsymbol{G}_{ii} + 2\sum_{j\notin S} \boldsymbol{G}_{ij} \geq 0$ from the residual, so a non-negative Gram makes $E_k$ non-increasing for any ordering. The real test is the diminishing-returns inequality of Theorem 4.3: it is checked on 384 random subset pairs (24 per trial) and violated 0 times. Figure 7(c) shows the shortfall $1 - U_k^{\text{g}}/U_k^\star$ over trials: the worst-case ratio across all $(k, \text{trial})$ pairs is 0.9992, so the empirical suboptimality gap $1 - U^{\text{g}}/U^\star = 0.08\%$ is more than two orders of magnitude (a factor of about 460) tighter than the worst-case $1/e \approx 0.368$ allowed by Theorem 4.5. We deliberately picked the densest-overlap regime (box side 0.4, splats with covariance length-scales up to 0.4) to stress greedy's myopia; even there, greedy is indistinguishable from the exhaustive combinatorial optimum, so on these small Gaussian-splat instances the $(1 - 1/e)$ bound is far from binding.

#### 8.2 Experiment B: synthetic ordering comparison

**Setup.** Synthetic 2D mixtures with $N \in \{64, 128, 256\}$, centres drawn uniformly in a fixed box of side 1.5, 6 trials each. All six orderings of Section 3 are evaluated; we report the loading curves $\|\boldsymbol{r}_k\|^2/\|f\|^2$ and the AUC.

**Result.** Figure 8 shows a consistent ordering at every $N$: **greedy** $<$ **{mass, self-energy, spectral}** $<$ **amplitude** $\ll$ **random** in AUC (lower is better). At $N = 256$ the median AUCs are 0.162 (greedy), 0.170 (mass), 0.169 (spectral), 0.172 (self-energy), 0.252 (amplitude), 0.339 (random). Greedy beats the best heuristic by 4% AUC and beats random by 52%, and has the lowest AUC in 18 of the 18 instances. Mass, self-energy and spectral cluster together: their

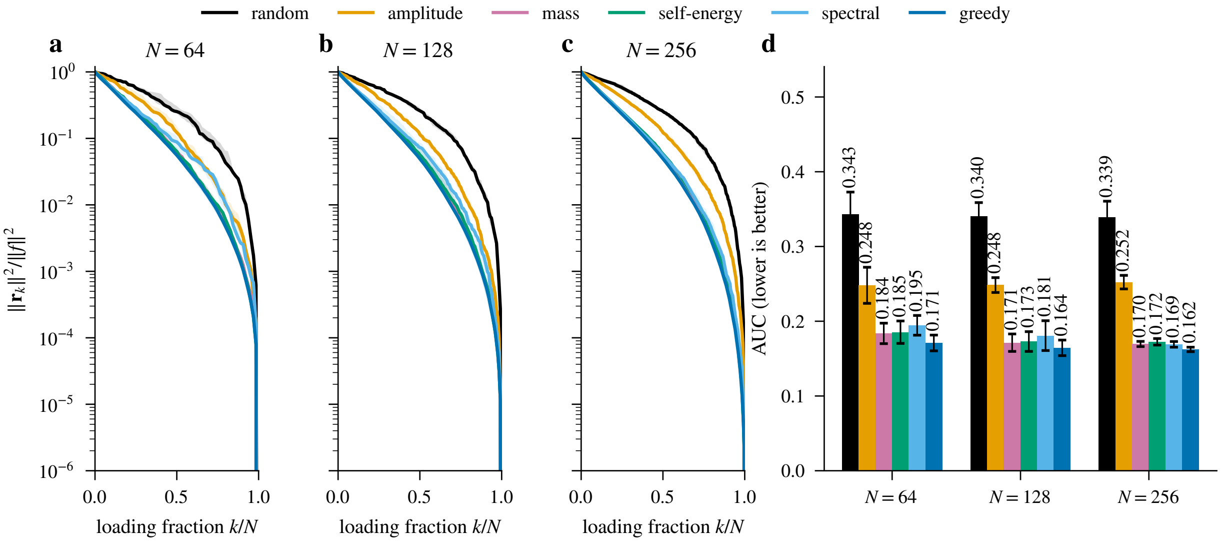

Figure 8: Experiment B: synthetic mixture loading curves. (a) to (c): median curves with interquartile-range bands (6 trials per $N$). (d): AUC bars are the median over the 6 trials, printed above each bar, with error bars of $\pm 1$ population standard deviation (not a confidence interval). Greedy has the lowest AUC in every instance; its median curve lies below every heuristic's except in the last few percent of the ladder ($k/N \geq 0.95$, relative residual below $10^{-3}$), where the self-energy curve crosses it (at $N = 64$ the mass curve too). Mass, self-energy and spectral cluster within 2.0% AUC of each other at $N = 256$ (5.5–5.9% at $N = 64$ and 128); mass and self-energy are the best $O(N \log N)$ alternatives. Amplitude trails the structured orderings substantially because splat extent is ignored.

median AUCs lie within 2.0% of each other at $N = 256$ and within 5.5–5.9% at $N = 64$ and 128. Of the three, mass and self-energy are $O(N \log N)$ alternatives; spectral needs the Gram matrix and an eigensolve (Table 1). Random and amplitude trail consistently: amplitude ignores splat extent (large covariance volume contributes substantially to $\|f\|^2$), so picking by peak height alone misses much of the energy. At late prefixes ($k/N > 0.7$) all non-random orderings concentrate near zero residual; differences shrink because most splats have been loaded and the residual is dominated by the few remaining tails. The greedy advantage is concentrated in the early prefixes ($k/N \in [0, 0.3]$), where the cross-splat correction terms in the marginal-gain formula (13) matter most.

##### 8.3 Experiment C: real Luxar gsplat datasets

**Datasets.** Four 3D Luxar gsplat demo scenes, one node each (Table 2), are read through Luxar's own scene reader, so that the encoded centres, amplitudes and Cholesky factors are decoded by the metadata stored with each array. Every scene is a public demo, built by `luxar demo run <key>` with the key of Table 2, and the analysis records the ***`content_hash`*** of each store it read, so a rebuilt store that differs from the one measured is detectable. A leaf that carries an additive ladder is read as the union of its rungs and checked against the leaf's declared count. The MAP4-GFP channel of the OpenCell scene is not used.

Table 2: The four real datasets of Experiment C. "Label" is the short name used in the figures, "Demo key" the public Luxar demo that builds the scene, "Node" the gsplat node read from it. "Stored" is the splat count of that node and "Rungs" the number of additive sub-LODs it ships with (1 = a single slab). The content hashes of the stores as measured are recorded with the analysis results.

| **Label** | `dapi_nuclei` | `acto3d_heart` | `tribolium_embryo` | `opencell_map4` |
|---|---|---|---|---|
| Demo key | `gsplats_3d_blastocyst_multichannel` | `gsplats_3d_acto3d_heart` | `gsplats_3d_tribolium_embryo` | `gsplats_3d_opencell_map4` |
| Node | `ch1_dapi` | `gsplats_sytox_green_nuclei` | `tribolium_embryo` | `gsplats_ch0` |
| Content | Blastocyst DAPI nuclei | Mouse heart nuclei (SYTOX Green, Acto3D) | *Tribolium* embryo nuclei | OpenCell-MAP4 nuclei (Hoechst channel) |
| Stored | 13 941 | 232 359 | 296 559 | 59 609 |
| Rungs | 4 | 1 | 1 | 4 |

**Filtering and subsampling.** Splats with amplitude at or below $10^{-3}$, or with a covariance whose smallest eigenvalue is at or below $10^{-12}$, are dropped before anything else. The amplitude criterion removes 38 684 splats (13.0%) from the *Tribolium* node and nothing from the other three, and the covariance criterion removes nothing, so the valid counts are 13 941, 232 359, 257 875 and 59 609. Each dataset is then reduced to a uniform-random subsample of $N = 4\,000$ splats (one draw, fixed across the six orderings) to keep the rasterised-PSNR loop bounded.

**Support-pruned surrogate.** Because Luxar splats carry a finite Mahalanobis support (canonically $2.75\sigma$), two splats whose support balls do not overlap have $\boldsymbol{G}_{ij} = 0$ exactly. We prune at $3\sigma$, deliberately wider than that support, so no pair of overlapping truncated splats is discarded. The retained pairs, however, carry the untruncated inner product (4), so the sparse matrix is a support-pruned surrogate of the Gram matrix, not the exact Gram of either the untruncated or the truncated field (Section 7, "Sparse greedy"). Every Experiment C AUC below is the AUC of that surrogate objective. At $N = 4\,000$ the surrogate is sparse: its density ranges from 0.04% on `tribolium_embryo` (6 324 non-zeros) through 0.15% on `acto3d_heart` and 0.24% on `opencell_map4` to 0.84% on `dapi_nuclei` (134 078 non-zeros) (Algorithm 4). Building it took 0.11–0.18 s per dataset and the sparse greedy ordering 0.09–0.10 s, against at most 0.3 ms for self-energy, 8 ms for mass and 6 ms for the sparse eigensolve of the spectral ordering (pure Python with NumPy/SciPy on the host of the run manifest). That eigensolve (ARPACK through SciPy) is started from the all-ones vector, the coefficient vector of $f$, so that the archived run is repeatable; its default random start is not. The two leading eigenvalues of the surrogate are separated by a relative gap of $6.2 \times 10^{-2}$ to $3.0 \times 10^{-1}$, and in a diagnostic run that is not deposited with the results the leading eigenvector was localised on a small cluster of splats: below that cluster its entries, and hence the order read off them, were at floating-point level, so the spectral AUC is fixed by the archived run and not by the data alone.

**Metrics.** For each dataset we report two metrics: (i) the analytical relative $L^2$ loading curve of the surrogate at all $k = 0, \ldots, N$; (ii) the rasterised *PSNR* against the fully-loaded ground truth, sampled at $k/N \in \{1\%, 2.5\%, 5\%, 10\%, 25\%, 50\%, 75\%, 90\%, 99\%\}$ (finer at the early prefixes where ordering quality matters most; the 100% checkpoint is omitted because rasterising all splats yields a numerical exact match and PSNR$\to \infty$). A single-window structural similarity index (SSIM) on the same support (global mean, variance and covariance of the two fields, with $C_1 = (0.01\,\text{peak})^2$ and $C_2 = (0.03\,\text{peak})^2$; not the local-window SSIM of Wang et al. [18]) is stored alongside the PSNR as an audit metric.

**Audit grid.** The grid behind the rasterised metrics is sized by the splats, not by a fixed voxel count. Its pitch is the subsample's median geometric-mean sigma $|\boldsymbol{\Sigma}_i|^{1/6}$ divided by 2: 0.67, 2.28, 0.52 and 0.53 in the datasets' own units for `dapi_nuclei`, `acto3d_heart`, `tribolium_embryo` and `opencell_map4`. The median splat therefore covers 7238–7247 voxel centres inside its $6\sigma$ support, and every splat of every subsample covers at least one (coverage 100.0%). The grid spans the axis-aligned box of the subsample's centres padded by $4\sigma_{\max}$, where $\sigma_{\max} = \max_i \sqrt{\lambda_{\max}(\boldsymbol{\Sigma}_i)}$ is the largest per-axis standard deviation in the subsample ($1951 \times 3549 \times 1900$ voxels on `tribolium_embryo`, whose splats are small against its extent). Only its *support voxels* are materialised and scored: the centres whose squared Mahalanobis distance $q_i = (\boldsymbol{x}-\boldsymbol{\mu}_i)^\top \boldsymbol{\Sigma}_i^{-1}(\boldsymbol{x}-\boldsymbol{\mu}_i)$ to at least one splat of the subsample satisfies $q_i < 36$, that is, the centres within $6\sigma$ of some splat ($2.4 \times 10^7$ to $6.7 \times 10^8$ per dataset). The rasteriser sums the untruncated Gaussians $a_i \exp(-q_i/2)$ there, and the MSE behind the PSNR is the mean over that fixed support, with the peak fixed to the maximum of the fully-loaded field. On a fixed support with a fixed peak every prefix of positive splats lies pointwise below the next, so the MSE cannot rise and the PSNR cannot fall along a ladder. The runner asserts this monotonicity on every curve before it writes the result table, and all 24 stored curves satisfy it.

**Resolution control.** A dense $32^3$ grid over the same box, used here as a resolution control, shows why the grid must be sized by the splats. Its pitch exceeds the splats' extent on every dataset; most of the `tribolium_embryo` subsample has no voxel centre inside $6\sigma$ at all (coverage 5.5%, 3 779 of 4 000 splats unseen, against 88.1% at best elsewhere), and the three energy orderings tie there to the third decimal over most of the ladder. On such a grid a PSNR curve saturates as soon as the few splats the grid can see have arrived: the grid is blind to the very splats the orderings disagree about. Visual progression on the *full* valid splat sets (no subsampling, $N$ up to 257 875) is shown separately in Fig. 10.

**Reproducibility.** Counts, thresholds, content hashes and the run manifest (Python 3.14.6, NumPy 2.4.6, luxar `429ddb065`) are archived with the results as a result table; the experiment code regenerates that table and every figure of this document deterministically from a fixed seed, and every number quoted here is read from the archived table.

**Result.** Figure 9 reports the analytical $L^2$ AUC and rasterised PSNR endpoints for all six orderings on every dataset. The AUCs are 0.01419/0.01422 (greedy/self-energy) for `tribolium_embryo`,

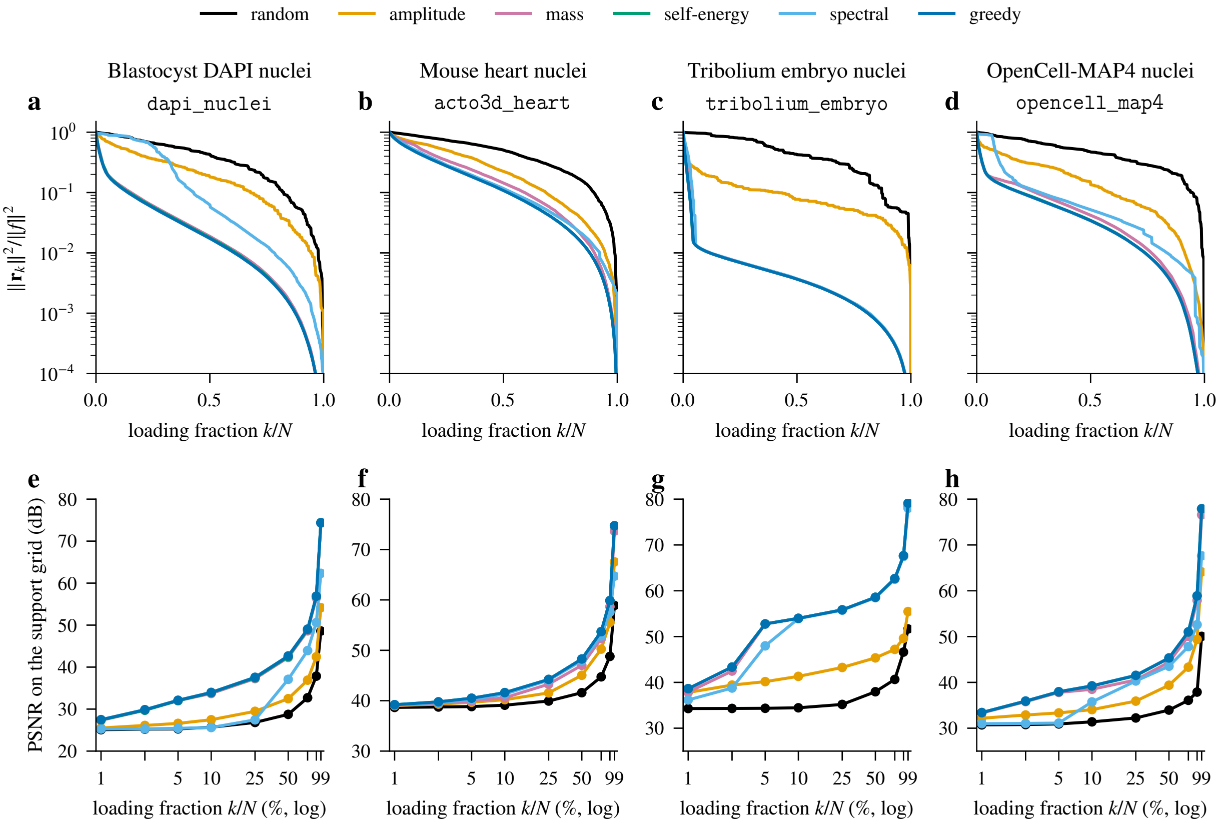

Figure 9: Experiment C: progressive loading on four real Luxar splat datasets, single trial, $N = 4\,000$ (one uniform-random subsample of the valid splats) on every dataset. Top row, (a) to (d): analytical relative $L^2$ residual energy $\|\boldsymbol{r}_k\|^2/\|f\|^2$ of the support-pruned surrogate vs. loading fraction (shared log-scale y-axis $10^{-4}$ to 1). Bottom row, (e) to (h): rasterised PSNR vs. the fully-loaded ground truth on the support voxels of a grid pitched at half the subsample's median sigma, with a fixed peak, sampled at $k/N \in \{1\%, 2.5\%, 5\%, 10\%, 25\%, 50\%, 75\%, 90\%, 99\%\}$ on a log $x$-axis to make the early-prefix regime readable (the tick marks are the checkpoints, labelled in percent where they fit), with $y$ capped at 80 dB (the 99% checkpoints reach up to 79 dB). Every PSNR curve is non-decreasing in $k$, as nested positive prefixes on a fixed support require; the runner asserts this before writing the results. Greedy / matching-pursuit ordering has the lowest AUC on every dataset, but it lowers the AUC relative to the cheap $O(N \log N)$ self-energy ordering by only a small margin (per-dataset values in the text), the gap increasing with Gram density across the four datasets. Self-energy beats mass by 2.7–17.8%. Spectral trails greedy and self-energy everywhere; below the small leading cluster of splats its order is at floating-point level, so its position relative to mass is not read from this figure. Random has the largest AUC on every dataset, and amplitude alone lies between them.

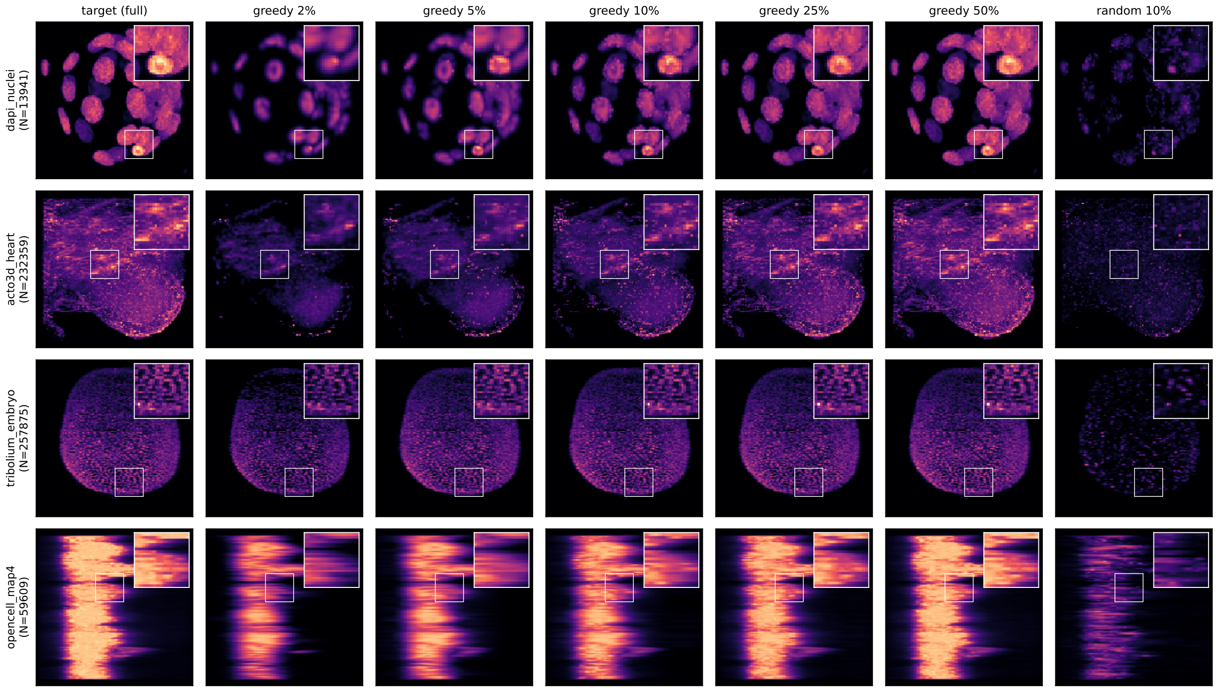

Figure 10: Experiment C: progressive refinement on the full real Luxar datasets (rows: `dapi_nuclei`, `acto3d_heart`, `tribolium_embryo`, `opencell_map4`). The per-row $N$ is the count of valid splats after the amplitude and covariance filters of Section 8.3: 13 941, 232 359, 257 875 and 59 609; on `tribolium_embryo` the amplitude filter removes 38 684 of 296 559 stored splats, on the other three it removes none. Each panel is rendered with Luxar's GPU-accelerated Python renderer `render_to_volume` (at the $2.75\sigma$ truncation the viewer also uses, a different field from the untruncated support-voxel rasteriser of Fig. 9) on a $128^3$ grid over one box per row (the full set's centre range padded by 5%) and shown as a Z-axis maximum-intensity projection so volumetric structure folds into a recognisable 2D image. Columns 2–6 show the residual-correlation greedy ordering at $\{2\%, 5\%, 10\%, 25\%, 50\%\}$ load. The right-most column shows one random ordering at 10% load as a same-budget baseline (a single draw, not a worst-case ordering); the random prefix is dominated by scattered low-energy splats and shows far less of the structure than the greedy prefix at the same budget. Greedy is computed via sparse Gram + sparse-greedy (Algorithm 4) directly at full $N$ (a dense float64 Gram of the largest row, `tribolium_embryo`, would occupy 532 GB). Each panel carries an upper-right inset (35% of the panel width) magnifying an 18%-side crop; vmin/vmax are shared within each row. The horizontal streaks of the `opencell_map4` row are its splats seen side on: the row is projected along the first centre axis and displays the second against the third, and the third axis spans about a sixth of the other two ($1236 \times 1236 \times 199$ audit voxels in Section 8.3), so the cubic $128^3$ render grid stretches that axis about six times relative to the others, and every splat with it. The figure is the archived render reproduced at reduced scale, so its printed row labels and column titles fall below print size; the rows and columns are identified in this caption, in the order listed.

0.05170/0.05310 for dapi_nuclei, 0.06233/0.06299 for opencell_map4, and 0.18789/0.18883 for acto3d_heart. Greedy has the lowest AUC on every dataset, but the reduction it achieves relative to self-energy, $(\mathrm{AUC}_{\mathrm{self}} - \mathrm{AUC}_{\mathrm{greedy}})/\,\mathrm{AUC}_{\mathrm{self}}$, is small and *increases with the Gram density* across the four datasets: 0.2% at 0.04% density (tribolium_embryo), 0.5% at 0.15% (acto3d_heart), 1.0% at 0.24% (opencell_map4), and 2.6% at 0.84% (dapi_nuclei). Expressed as self-energy's penalty relative to greedy, the same four gaps are 0.2–2.7%. The order of the four gaps is consistent with the mechanism behind greedy: its advantage comes from accounting for overlap between splats already sent and those still to come, so it has the most to add where splats overlap most and nothing to add on an orthogonal dictionary (Remark 3.1). The theory, however, predicts no monotone relation with the nonzero density, which counts support-pruned pairs without weighting them by amplitude or by the size of the overlap, and four single-trial points cannot establish one. On the rasterised metric the two are close everywhere: greedy's PSNR is at or above self-energy's at 36 of the 36 checkpoints, the largest difference in either direction is 0.17 dB, and at 50% load on dapi_nuclei, the dataset with the largest AUC gap, greedy reaches 42.66 dB against 42.49 dB for self-energy. Self-energy beats mass on every dataset, by 2.7% AUC on dapi_nuclei, 8.0% on tribolium_embryo, 13.2% on opencell_map4 and 17.8% on acto3d_heart (mass weights peak amplitude and covariance volume, self-energy weights amplitude *squared* and covariance volume, and squaring is what matches the $L^2$ objective). The spectral ordering trails greedy and self-energy on every dataset: the leading Gram eigenvector identifies splats that contribute most to a *single* global $L^2$ direction, but the $L^2$ energy of $f$ is spread across many directions and a single eigenvector ordering is suboptimal in early prefixes. The synthetic cluster of Section 8.2, where spectral sits within a few percent of mass and self-energy, is not reproduced. The mechanism is consistent with localisation. The leading eigenvector of a non-negative matrix is its Perron vector, and when the non-zeros of a support-pruned Gram form loosely coupled clusters that vector concentrates on the most strongly coupled cluster, so $|\boldsymbol{u}_1[i]|$ ranks that cluster first. In a diagnostic run on these Gram matrices the leading eigenvector was localised in this way: its entries outside a small cluster were at floating-point level relative to the largest, so the order of almost every splat was set by rounding, and a floating-point-level perturbation of the Gram moved the spectral AUC on acto3d_heart from below to above the mass AUC. That run is not deposited; the archived table records only the two leading eigenvalues and the start vector of the eigensolve. The recorded spectral AUCs are therefore quoted only where the comparison survives such perturbations: spectral trails greedy and self-energy everywhere, beats amplitude on acto3d_heart, tribolium_embryo and opencell_map4 and loses to it on dapi_nuclei; its position relative to mass is not reported. A dense Gram, as in Section 8.2, couples every splat to every other and has no such localisation. Random has the largest AUC on every dataset (0.4363–0.4944), and amplitude alone sits between the energy-based orderings and random: ahead of spectral on dapi_nuclei (0.2467 vs. 0.2692) but behind it on the other three datasets, since peak amplitude ignores how much of a splat's energy its extent carries.

**Tentative recipe (limited to these experiments).** Under the analytical-$L^2$ objective and the equal-cost-per-splat assumption stated in Section 1, the experiments suggest a working recipe: use the cheap $O(N \log N)$ self-energy ordering as a strong baseline, and upgrade to greedy (feasible at full $N$ via sparse Gram + sparse-greedy, Algorithm 4) when the AUC reduction measured above is worth the precompute. At $N = 4\,000$ that precompute took under a second in pure Python; the full-$N$ orderings behind Fig. 10 are not timed in the archived results. The recipe rests on single-trial experiments; repeated subsamples and rendering-aware metrics remain to be tested before it can be called a deployment default.

**Why self-energy is close to greedy.** Remark 3.1 predicts that greedy reduces to self-energy ordering in the nearly-orthogonal limit $\boldsymbol{G}_{ij} \approx 0$ for $i \neq j$. Empirically the gap is small (Fig. 9), which is *suggestive* that the Luxar fitting pipeline tends to produce splat dictionaries whose off-diagonal Gram energy is small relative to the diagonal: splats appear to settle on locally distinct features. A small gap is not by itself a proof of near-orthogonality, since a heuristic can also come close to greedy on a redundant dictionary. Nor is the nonzero density we report a measure of amplitude-weighted overlap: it counts support-pruned pairs, nearly touching broad splats can contribute negligible energy, and a change of amplitudes moves the couplings while leaving the density unchanged. Across our four datasets the gap increases with that density, but four single-trial points cannot establish a functional form, the theory does not predict one, and the gap plausibly also depends on amplitude heterogeneity and on how the overlap structure distributes mass across eigendirections. A direct measurement (normalised Gram cosines $\boldsymbol{G}_{ij}/\sqrt{\boldsymbol{G}_{ii}\boldsymbol{G}_{jj}}$, or the off-diagonal energy fraction $1 - \eta = 1 - \mathrm{tr}(\boldsymbol{G})/\|f\|^2$ with $\eta$ the orthogonality fraction of Section 3) would be the right diagnostic and is not recorded here. We treat the greedy/self-energy AUC gap as a *suggestive diagnostic* of the overlap structure, not as a clean quantitative measure of any single property.

**Visual progression.** Figure 10 shows the rasterised progressive refinement for each dataset using the *full* valid splat set (no subsampling; 13 941, 232 359, 257 875 and 59 609 splats) and the residual-correlation greedy ordering itself. Sparse Gram + sparse-greedy (Algorithm 4) is what brings full-$N$ greedy into reach: a dense float64 Gram of the largest of these, `tribolium_embryo`, would occupy 532 GB ($8N^2$ bytes), and 432 GB for `acto3d_heart`. The full-$N$ Gram and ordering times of this figure are not part of the archived results. Each panel is a Z-axis maximum-intensity projection (MIP, the brightest value along each line of sight) of the rasterised volume, so all volumetric structure folds into a single recognisable 2D image. Checkpoints are concentrated in the early-prefix regime ($\{2\%, 5\%, 10\%, 25\%, 50\%\}$): once the brightest splats are loaded, the MIP saturates quickly and later prefixes look almost identical, but at 2–5% load the structural skeleton is already visible. The right-most column shows a single random ordering at 10% load as a baseline at the same budget (one draw, not a worst-case ordering): the random prefix is dominated by scattered low-energy splats and shows markedly less of the structure the greedy prefix has already assembled, although some structure remains visible. The AUC analysis in Fig. 9 is the more sensitive metric for distinguishing greedy from the cheap heuristics; the visual figure is the easiest way to confirm that the greedy ordering loads structure in the right order.

### 9 Discussion and Open Questions

**Limits of the empirical evidence.** The real-data study uses a single uniform subsample of $N = 4\,000$ per dataset and, for PSNR, one support-voxel grid per dataset pitched at half the subsample's median sigma (the visual-progression Fig. 10 uses the full datasets at $128^3$, rendered with a different kernel). That is enough for a credible ranking under the analytical-$L^2$ objective, but not enough to support strong deployment claims. Four pieces are still missing. (a) Variability across repeated random subsamples would put error bars on the AUC and PSNR numbers. (b) Wall-clock scaling plots for greedy vs. self-energy on $N \gtrsim 10^5$ would replace our worst-case complexity arguments with measured runtime. (c) A head-to-head comparison against rendering-aware methods such as PRoGS [19] and LapisGS [16] on a rendered-image metric is absent. (d) View-conditioned variants of the greedy ordering for interactive viewing are untested. We see all four as natural follow-ups but treat the present results as a strong analytical-$L^2$ baseline, not a finished practical recommendation.

**Cost of the greedy precompute.** Constructing the dense $N \times N$ Gram matrix is $O(N^2)$ in entries and in memory and is therefore prohibitive at $N \sim 10^7$. Luxar splats carry a finite Mahalanobis support (`truncation_radius`, canonically $2.75\sigma$; we prune at the wider $3\sigma$, so no overlapping pair is discarded), so two splats with non-overlapping support balls have $\boldsymbol{G}_{ij} = 0$ exactly and the Gram matrix of the rendered field is naturally *sparse*. The support-pruned matrix we build is a surrogate of it (Section 7). At the $N = 4\,000$ subsamples of Section 8.3 its density is at most 0.84% (per-dataset values and timings there); the full-$N$ densities behind Fig. 10 are not archived. The same code orders the full valid sets of up to 257 875 splats for Fig. 10, where a dense float64 Gram would need 532 GB. Two further mitigations remain useful at scale ($N \gtrsim 10^7$): (i) lazy column reads; (ii) process the splats by spatial tiles or octree cells, running the greedy ordering *within* each tile. Tiling is a heuristic, not a decomposition of the objective: assigning two centres to different cells does not separate their supports, and a pair straddling a cell boundary can overlap almost as much as two coincident splats. The within-tile ordering therefore silently drops every cross-boundary $\boldsymbol{G}_{ij}$ unless each tile is built with a halo of at least one support radius (the companion document [8] likewise retains its cross-bin terms instead of assuming them away). The omitted cross-boundary energy should be measured before relying on this; tile-based processing is not benchmarked here.

**View-dependent ordering.** For an interactive viewer, the user's current viewpoint biases which splats are visible. A natural extension is to combine the greedy order with a view-frustum filter: load the greedy order *restricted to the visible cell set*, then continue with the global greedy as the user navigates. PRoGS does this with a frustum-aware ranking; the same extension applies here. An open question is whether view-conditioning preserves the $(1 - 1/e)$ guarantee; we conjecture yes if the conditional visibility is itself a monotone non-decreasing predicate as new tiles enter the frustum.

**Choice of metric beyond $L^2$.** We have optimised against the analytical $L^2$ residual of the support-pruned surrogate. Experiment C measures three different fields: that surrogate (untruncated pair products, pruned at $3\sigma$); the untruncated support-voxel rasteriser behind the PSNR and SSIM (Gaussians summed at the voxel centres within $6\sigma$ of some splat, pitch half the median sigma, fixed box and peak); and, for Fig. 10 only, Luxar's own renderer with its $2.75\sigma$ truncation at $128^3$. The rendered metrics agree with the analytical ranking to a quantifiable degree, short of confirming it. Over the 540 ordering pairs (four datasets, 9 checkpoints, 15 pairs of the six orderings) the rendered metric separates the two orderings in all but 5 cases for PSNR and 5 for SSIM (exact ties, where two prefixes coincide on every support voxel; a tie is neither agreement nor disagreement and is left out of the denominator). Among the 535 decided PSNR pairs the sign of the PSNR difference matches the sign of the AUC difference in 97.4% of cases, and the single-window SSIM in 95.0% of its 535. The disagreements sit almost entirely in pairs that involve the spectral ordering: 99.0% concordance inside the trio of energy-based orderings (108 pairs, 5 of them ties) against 97.0% over the remaining 432. Inside the trio the PSNR values are separated by fractions of a decibel over most of the ladder (greedy and self-energy lie within 0.17 dB at every checkpoint). The SSIM is a global-moment approximation, not the local-window statistic, and the pitch was set by the splats' own scale without a convergence sweep, so neither metric is offered here as a validated rendering-quality score. For radiance-field rendering with view-dependent shading, the situation is less clear: a view-dependent rendering metric does not reduce to $L^2$ on the parameters, and the corresponding utility may not be submodular. Quantifying this gap is a useful follow-up.

**Combining substitutive and additive LOD.** The natural pipeline is (i) build a substitutive hierarchy $g_0 = f, g_1, \ldots, g_L$ via the algorithms of [8]; (ii) at each level, additionally store an additive ordering $\pi_\ell$ via the greedy of this document. At runtime, the viewer pages in $g_{\ell^\star}$ for the level $\ell^\star$ matching the current bandwidth/memory budget, then refines within that level by streaming splats in the order $\pi_{\ell^\star}$. The interaction between levels is benign because each level is independently ordered; whether further joint optimisation across levels yields non-trivial gains is an open question.

**Multi-channel and time-varying scenes.** For multi-channel splats with per-channel amplitudes, applying the greedy ordering per channel and merging by interleaving is the simplest strategy. For time-varying scenes (4D / nD splats), the greedy can be applied either per timepoint or jointly with a Gram matrix that aggregates inner products across time (a temporally-stationary scene clusters splats that persist across timepoints to load earlier).

#### 10 Summary

The additive LOD problem has a clean structure and a modest practical payoff. The structure is that the cumulative energy captured by a subset of splats is monotone submodular whenever the Gram matrix is non-negative (Theorem 4.3), which every positive-amplitude splat set satisfies, so the greedy ordering carries a uniform $(1 - 1/e)$ guarantee on the captured energy at every prefix

(Theorem 4.5) and an affine bound on the AUC (Corollary 4.7). On exhaustively solved small instances greedy sits within 0.08% of the optimum, so the guarantee is a floor and not a forecast. The lower bound of Proposition 5.1 is the one that holds; its eigen-coordinate strengthening fails on a seven-splat example.

The payoff is modest because the cheap orderings are already good. Ordering by $L^2$ self-energy costs $O(N \log N)$, needs no Gram matrix, and on the four real datasets of Section 8.3 trails greedy by a margin that is small everywhere and largest on the dataset whose support-pruned Gram is densest. Integral mass trails self-energy on every dataset; the spectral ordering trails greedy and self-energy, with an order that is set at floating-point level below a small leading cluster of splats; and peak amplitude and random orderings are clearly worse than the energy-based ones.

The recommendation for building ladders follows. Use the self-energy ordering by default. Switch to greedy through the support-pruned sparse Gram (Algorithm 4) when a dataset's splats overlap densely and the precompute, a fraction of a second at $N = 4\,000$ in pure Python, is acceptable. The spectral ordering has no case here: it costs a Gram matrix and an eigensolve and loses to the free self-energy ordering on every real dataset. These recommendations hold for the analytical $L^2$ objective with equal cost per splat and rest on single-trial experiments. Rendering-aware and view-dependent metrics, unequal-cost streaming and a head-to-head comparison with PRoGS [19] and LapisGS [16] remain open.

[4] Zhiwen Fan, Kevin Wang, Kairun Wen, Zehao Zhu, Dejia Xu, and Zhangyang Wang. Light-Gaussian: Unbounded 3D Gaussian compression with 15× reduction and 200+ FPS. In *Advances in Neural Information Processing Systems*, volume 37, 2024.

[5] Bernhard Kerbl, Andreas Meuleman, Georgios Kopanas, Michael Wimmer, Alexandre Lanvin, and George Drettakis. A hierarchical 3D Gaussian representation for real-time rendering of very large datasets. *ACM Transactions on Graphics*, 43(4):1–15, 2024. doi: 10.1145/3658160.

[6] Andreas Krause and Daniel Golovin. Submodular function maximization. In Lucas Bordeaux, Youssef Hamadi, and Pushmeet Kohli, editors, *Tractability: Practical Approaches to Hard Problems*, pages 71–104. Cambridge University Press, 2014.

[7] Luxar Project. Analytical merging error for Gaussian splats: An $L^2$ framework for optimal approximation, 2026. Supplementary material, this volume.

[8] Luxar Project. Substitutive levels of detail for Gaussian splats: Cost metrics, bin-wise merging, and hierarchical construction, 2026. Supplementary material, this volume.

[14] Kerui Ren, Lihan Jiang, Tao Lu, Mulin Yu, Linning Xu, Zhangkai Ni, and Bo Dai. Octree-GS: Towards consistent real-time rendering with LOD-structured 3D Gaussians, 2024. arXiv:2403.17898.

[15] Sartaj Sahni and Teofilo Gonzalez. P-complete approximation problems. *Journal of the ACM*, 23(3):555–565, 1976. doi: 10.1145/321958.321975.

[16] Yuang Shi, Géraldine Morin, Simone Gasparini, and Wei Tsang Ooi. LapisGS: Layered progressive 3D Gaussian splatting for adaptive streaming. In *Proceedings of the International Conference on 3D Vision (3DV)*, 2025. arXiv:2408.14823.

[17] Vladimir N. Temlyakov. Greedy approximation. *Acta Numerica*, 17:235–409, 2008. doi: 10.1017/S0962492906380014.

[18] Zhou Wang, Alan C. Bovik, Hamid R. Sheikh, and Eero P. Simoncelli. Image quality assessment: From error visibility to structural similarity. *IEEE Transactions on Image Processing*, 13(4):600–612, 2004. doi: 10.1109/TIP.2003.819861.

[19] Brent Zoomers, Maarten Wijnants, Ivan Molenaers, Joni Vanherck, Jeroen Put, Lode Jorissen, and Nick Michiels. PRoGS: Progressive rendering of Gaussian splats. In *Proceedings of the IEEE/CVF Winter Conference on Applications of Computer Vision (WACV)*, pages 3118–3127, 2025. doi: 10.1109/WACV61041.2025.00308.
