## Supplementary material for "Luxar: Gaussian splatting for microscopy and scalable interactive web visualisation of multidimensional scientific data": supp_doc_10_scene_graph_pipeline

### Scene graph, transforms, overlays, and the viewer pipeline: a reference for the Luxar runtime

Supplementary Document 10 – Luxar

**Abstract**

The main text compresses Luxar's runtime into a few Methods paragraphs. This document is the reference for that runtime: the data structures, wire formats and rendering algorithms behind the viewer. A Luxar scene is a typed graph of groups and geometry leaves mirrored one-to-one onto Zarr groups, each node persisting a small attribute set of which only the type discriminator is mandatory; level-of-detail groups and spatial partitions are group kinds of the same tree, and a standalone splat file is one such subtree. Spatial transforms compose along the parent chain as column-major $4 \times 4$ matrices, and per-dimension transforms on non-displayed axes are applied in inverse to the slice query, so the spatial index is built once and never invalidated. Appearance attributes compose root-to-leaf by fixed rules and map to shader uniforms. Screen-space overlays carry dimension-aware visibility. GPU picking renders identities into a half-resolution float buffer and resolves the cursor by brightness-weighted majority voting; per-element labels and thumbnails use a compressed-sparse-row layout. The HDR pipeline runs one fused full-screen pass in a fixed stage order under an adaptive device-pixel-ratio controller, and the per-dimension animation system gates its advance on data arrival and drives a recording panel that writes stills, video and HDR EXR sequences. Chunk encoding, level-of-detail construction and the chunk cache are documented elsewhere.

**10 Conclusion** **27**

### 1 Introduction

A Luxar dataset is two things simultaneously. On disk it is a chunked Zarr archive with a typed group hierarchy and per-group attributes. In the browser it is a TypeScript scene graph driving a WebGL/WebGPU renderer with a post-processing composer, a screen-space overlay layer, a GPU picking subsystem, and a per-dimension animation controller. The shape of the on-disk format is dictated entirely by what the runtime needs to consume, and the runtime is structured to load and decode that format lazily, holding the chunks the current view needs and not the whole dataset, with minimal latency. The question this document answers is what, exactly, the two halves promise each other: which attributes and arrays a compiled archive must carry, and what the viewer does with each of them.

This document walks through that runtime layer by layer. Section 2 describes the typed node hierarchy and the attributes each node persists. Section 3 covers spatial and non-displayed-dimension transforms, including the inverse-query algorithm that keeps slice queries proportional

Table 1: Source map: where each section's contracts are defined at the documented commit.

| Section | Defining sources (relative to `packages/luxar/src/luxar/`, `packages/luxar-viewer/src/`, `docs/`) |
|---|---|
| §2 | `core/node/node.py`, `core/datanode.py`, `core/group/`, `io/_compiler/geometry_writers/`; the viewer's `scene/lod-group-registry.ts` and `scene/lod-selector-math.ts` (level selection); `guides/user/LUXAR_ZARR_FORMAT.md` (scene format 0.2), `specs/GSPLATS_ZARR_FORMAT.md` (gsplats format 3.4) |
| §3 | `core/transforms.py` and `validation/nd_transforms.py`; the viewer's `data/transforms/nd-transform.ts`; the specification `guides/specs/ND_TRANSFORMS_SPEC.md` |
| §4 | `data/attrs-composer.ts`, `rendering/display-range.ts`, `ui/layers/` |
| §5 | `validation/overlays.py`, `core/scene/overlays/`; `ui/overlay-manager.ts` |
| §6 | `rendering/picking/`: `picking-system.ts`, `picking-system/pick-render.ts`, `picking-system/settle-loop.ts`, and the per-geometry `shaders.ts` |
| §7 | `data/loaders/picking/label-loader.ts`, `image-label-loader.ts`; `utils/hover-template.ts`; `core/scene/overlays/hover_inject.py` |
| §8 | the fused pass `rendering/post-processing/mega/shader.glsl.ts`; the stage sequence and capture modes in `rendering/post-processing/post-processing-manager/` (`pipeline.ts`, `capture.ts`); the defaults in `config/sections/rendering-controls/data.ts`; `rendering/adaptive-dpr-manager.ts` and `rendering/adaptive-dpr/` |
| §9 | `config/sections/dimension-animation/data.ts`, `ui/recording-panel/` (`gui-builder.ts`, `ffmpeg-script.ts`, `screenshot-strategy.ts`, `offline-capture-strategy.ts`) |

to the number of dimensions instead of $O(N)$ per point, where $N$ is the number of elements in a node. Section 4 documents the layer model and how display-range/gamma/blending controls map to shader uniforms. Section 5 specifies the overlay system. Section 6 documents GPU picking and the brightness-weighted majority voting that determines which element a hover event lands on. Section 7 covers the compressed-sparse-row layout for per-element labels and image labels and the auto-injection of default hover overlays. Section 8 documents the HDR pipeline and adaptive resolution scaling. Section 9 covers the per-dimension animation system and the recording panel. Chunk layout and encoding (Supplementary Document 12), the level-of-detail recipes (Supplementary Documents 8 and 9) and the chunk cache are not covered here.

**Software state.** This document describes the contracts as implemented at the pinned luxar commit `5ec6c9852`: the Python package `luxar` at version `2026.06.05` (its `__version__` string) and the viewer package `@luxar/viewer` at version `2026.6.5`, built on Three.js `0.185` and `zarrita` `0.7`. Three format versions are independent of one another and of the software version: the compiled-scene format is `0.2` (root attributes `format_version` and `format_type = "luxar_zarr"`), the standalone `.gsplats.zarr` format is `3.4` (root `format_type = "gsplats_zarr"`), and the Zarr container format is `3` for every newly written store, with format-2 stores read unchanged (`luxar._zarr_compat.DEFAULT_ZARR_FORMAT`; the environment variable `LUXAR_ZARR_FORMAT=2` selects the older container). The text refers to source files as `packages/luxar/...` for the Python authoring layer and `packages/luxar-viewer/src/...` for the TypeScript runtime, but does not depend on specific line numbers; its claims are about the contracts between the two layers. Table 1 maps each section to the files and specifications that define those contracts, and Table 2 collects the defaults and constants those sections quote, with the module that defines each.

Table 2: Defaults and constants quoted in Sections 2 to 8, with the module that defines each value at the documented commit. Python modules are relative to `packages/luxar/src/luxar/`, viewer modules to `packages/luxar-viewer/src/`; the configuration modules of the last two blocks live under `config/sections/`.

| Constant | Value | Module |
|---|---|---|
| *Format and transforms* | | |
| Zarr container format of a new store | 3 | `_zarr_compat.py` |
| Translation slots of the on-disk transform | 12, 13, 14 (row-major 3, 7, 11 refused) | `rendering/node-factory/validation.ts` |
| Discrete grid anchor | range[0] + $k \cdot$ step | `data/transforms/nd-transform.ts` |
| Half-step membership window | 0.5 step | `workers/data-worker/projection/gsplats.ts`, `wasm/typescript/effective-radii.ts` |
| Footprint LOD limit; downgrade hysteresis | 1.5 CSS px $/\sqrt{\texttt{lodBias}}$ (default bias 1); 10 % | `scene/lod-selector-math.ts`, `scene/lod-group-registry.ts` |
| *Overlays* | | |
| `visible_range` exact-value window | $\pm 0.5$ | `ui/overlay-manager.ts` |
| *Picking* (under `rendering/picking/`) | | |
| Pick render target | `RGBA32F` | `picking-system.ts` |
| Node-id bound; background rule | $2^{24} - 1$; $R < 0.5$ | `picking-system/pick-render.ts` |
| Element index reconstruction | $\text{round}(A) \cdot 65536 + \text{round}(G)$ | `picking-system/pick-render.ts` |
| Pick-buffer scale; longer-side cap | 0.5× drawing buffer; 1024 px | `picking-system.ts` |
| Readback window | $5 \times 5$ texels | `picking-system.ts` |
| `HOVER_SETTLE_MS` | 120 ms | `picking-system/settle-loop.ts` |
| *Post-processing* (defaults in `config/sections/rendering-controls/data.ts`) | | |
| `msaaSamples` | 4 (MSAA off) | `rendering-controls/data.ts` |
| `bloomLevels` | 8 (bloom off) | `rendering-controls/data.ts` |
| `toneMapping` | ACES | `rendering-controls/data.ts` |
| FXAA, SSAA, vignette, noise, distortion | all off | `rendering-controls/data.ts` |
| EXR capture mode; pixel type | `hdr-effects-pre-tone`; HalfFloat | `post-processing/post-processing-manager/capture.ts` |
| *Adaptive DPR* (defaults in `config/sections/adaptive-dpr/data.ts`) | | |
| `minDPR`; `scaleDownFactor`; `scaleUpFactor` | 0.5; 0.9; 1.05 | `adaptive-dpr/data.ts` |
| `scaleDownFpsRatio`; `scaleUpFpsRatio` | 0.75; 0.9 of the refresh cap | `adaptive-dpr/data.ts` |
| `evaluationIntervalMs`; `hysteresisSeconds`; FPS window | 500 ms; 3 s; 1000 ms | `adaptive-dpr/data.ts`, `rendering/adaptive-dpr-manager.ts` |
| `probeWindowMs`; `probeImprovement` | 1500 ms; 1.05× | `adaptive-dpr/data.ts` |
| `floorTtlMs`; `backoffMultiplier`; `backoffMaxTtlMs` | 30 000 ms; 2; 300 000 ms | `adaptive-dpr/data.ts` |
| `ceilingTtlMs`; `punishedAscentWindowMs`; `punishedAscentThreshold` | 60 000 ms; 3000 ms; 2 | `adaptive-dpr/data.ts` |

#### Terms used in this document

**additive (streaming) ladder** One ordering of a leaf's splats cut into prefix-sum sub-LODs, so that every prefix of the download is a valid, progressively refining rendering of the same splat set.

**rung (sub-LOD)** One contiguous slab of an additive ladder, stored as its own group (`additive_<i>`, index 0 the coarsest) and appended to the committed prefix when it arrives; a document that uses "rung" for one step of a splat-count sweep says so where it first does.

**`kind=lod` group** A substitutive level-of-detail group whose children are alternative renderings of the same content at different splat counts, of which the viewer draws exactly one at a time: the coarsest level whose median projected splat footprint stays at or below 1.5 CSS pixels when the levels carry footprint stamps, otherwise the level selected by the size of its projected bounding box (its fraction of the viewport area for a screen-area ladder, its normalised projected diagonal for a coverage ladder).

**`kind=partition` group** A group whose `part_<i>` children each carry their own position bounds and are all drawn at once, each culled against the camera frustum; the parts are usually the regions of a recursive BSP split of one splat set, but any disjoint grouping is valid, such as one part per timepoint.

**BSP** Binary space partitioning, the recursive splitting of a set of elements by an axis-aligned plane (at the median by default) into two halves until every part falls below a size cap.

**recipe** A named LOD topology built from a fitted splat set: `flat` (one bare leaf), `stream` (one stream-laddered leaf), `levels` (coarse-to-fine replacement levels), `tiles` (a BSP partition with one ladder per tile), `overview` (one coarse level above a `tiles` branch) and `adaptive` (`tiles` where every tile carries its own `levels`).

**spatial index** The chunk-based index of a node: elements are reordered along a space-filling curve and grouped into fixed-size chunks, each with a stored axis-aligned bounding box (`chunk_bounds`), so a slice query reads only the chunks whose boxes intersect it.

**committed prefix (committed count)** The part of an additive ladder that has been processed, uploaded to the GPU and is on screen, as distinct from rungs whose download has merely completed.

**energy stamps** Two attributes written at build time: `energy_fraction_cum`, $e(k)$, on each sub-LOD is the cumulative self-energy fraction of the ladder prefix ending at rung $k$, and `reference_energy`, $w$, on each leaf is the leaf's absolute total self-energy used to weight leaves against one another.

**display gate** The viewer rule that keeps the previously displayed level on screen while a finer level is still streaming and releases the swap once the finer level's committed prefix carries at least 0.6 of its total self-energy ($e(k) \geq 0.6$), applied on upgrades only.

**HDR and LDR** High versus low dynamic range: HDR colours are unbounded floating-point intensities (supplied as float32; the AUTO encoding stores them as per-channel logarithmic `uint16` and decodes back to float32, PRECISION keeps float32), rendered into a half-float target, while LDR colours are clamped to $[0, 1]$ as a display expects; tone mapping converts the first into the second.

**anti-aliasing (MSAA, SSAA, FXAA, SMAA)** Techniques that soften the stair-stepping of rasterised edges: SSAA shades several samples per pixel and averages them, MSAA shades once per pixel and keeps per-sample coverage, while FXAA and SMAA are post-process filters that smooth edges detected in the finished image.

**rAF** `requestAnimationFrame`, the browser callback that fires once per begin-frame tick (the display refresh when there is one); a page's rAF rate is capped at that cadence and counts callbacks, not frames the viewer drew, so on its own it is not a frame rate.

**LUT** A lookup table, a small array indexed by a quantised value; a colormap LUT maps a scalar in $[0, 1]$ to a stored colour.

**AABB** An axis-aligned bounding box, the smallest box with faces parallel to the coordinate axes that contains a set of elements.

**EBML** The Extensible Binary Meta Language, the binary container syntax on which Matroska and WebM video files are built.

#### 2 Scene graph and node attributes

##### 2.1 Node hierarchy

A Luxar scene is rooted at a `Scene` object that owns the dimension definitions and the writer responsible for streaming metadata to disk. Interior `Group` nodes bundle children and may carry transforms; leaf `Points`, `Lines`, `Mesh` and `GSplats` nodes carry the actual geometry. A fifth leaf type, `sound`, is a node of the graph but not a geometry: it holds an audio clip and positions, carries no appearance, no level of detail and no picking, and is dispatched to the audio engine by its `type` alone. The hierarchy is exactly what is persisted to Zarr: each node corresponds to one Zarr group, and each parent-child relationship corresponds to one nested group. Group names are constrained to exclude the path separator (“`/`”) and must be unique among siblings; both constraints are enforced at construction time so that the disk layout is always well-formed. Figure 1a draws that mirror for a small scene: one Zarr group and one `zarr.json` per node, with the storage sub-groups of a laddered leaf and the two group kinds introduced below.

Groups can nest arbitrarily deep; data nodes are leaves by convention: the authoring API offers no way to add geometry beneath a `Points` node and the viewer never looks for one, although the base-class `add_group` is inherited by data nodes and is not refused, so the restriction is a design rule, not a validated invariant. Storage sub-groups do occur beneath a data node: an *additive (streaming) ladder* stores each *rung (sub-LOD)*, a prefix chunk holding one slab of a single ordering of the leaf's elements, so that sub-groups $0..i$ together form a prefix of that ordering, as an `additive_<i>/` sub-group of the leaf that owns it, and the loaders treat those sub-groups as part of the leaf, not as children of the scene graph. Every node knows its parent (or `None` for the scene root) and exposes a fully-qualified path that the viewer uses both for layer identification and for cache-key generation.

##### 2.2 Persisted attributes

Each node persists a small set of attributes to its Zarr group's user attributes (in the node's `zarr.json` for the default Zarr format 3, in `.zattrs` for a format-2 store; the attributes are identical in both). Table 3 enumerates the attributes common to every node type; Table 4 lists the fields the root and the specialised group kinds add. Most are optional; the only mandatory attribute is `type`, which is the discriminator the viewer uses to dispatch to the right loader and material. Type-specific data (e.g. `n_points`, `ndim`, `has_colors`, `position_bounds`) is written by the geometry writer as further top-level attributes of the same group; the Python node object keeps an in-memory copy of those values (its `_metadata` member), which is a cache and is not itself serialised.

The level-of-detail recipes of the command line (*recipe*) map onto these kinds directly (Fig. 1b). `stream` is one leaf with an additive ladder; `levels` is a ***`kind=lod`*** *group* whose children are (laddered) leaves; `tiles` is a ***`kind=partition`*** *group* of laddered leaves split by a spatial *BSP*;

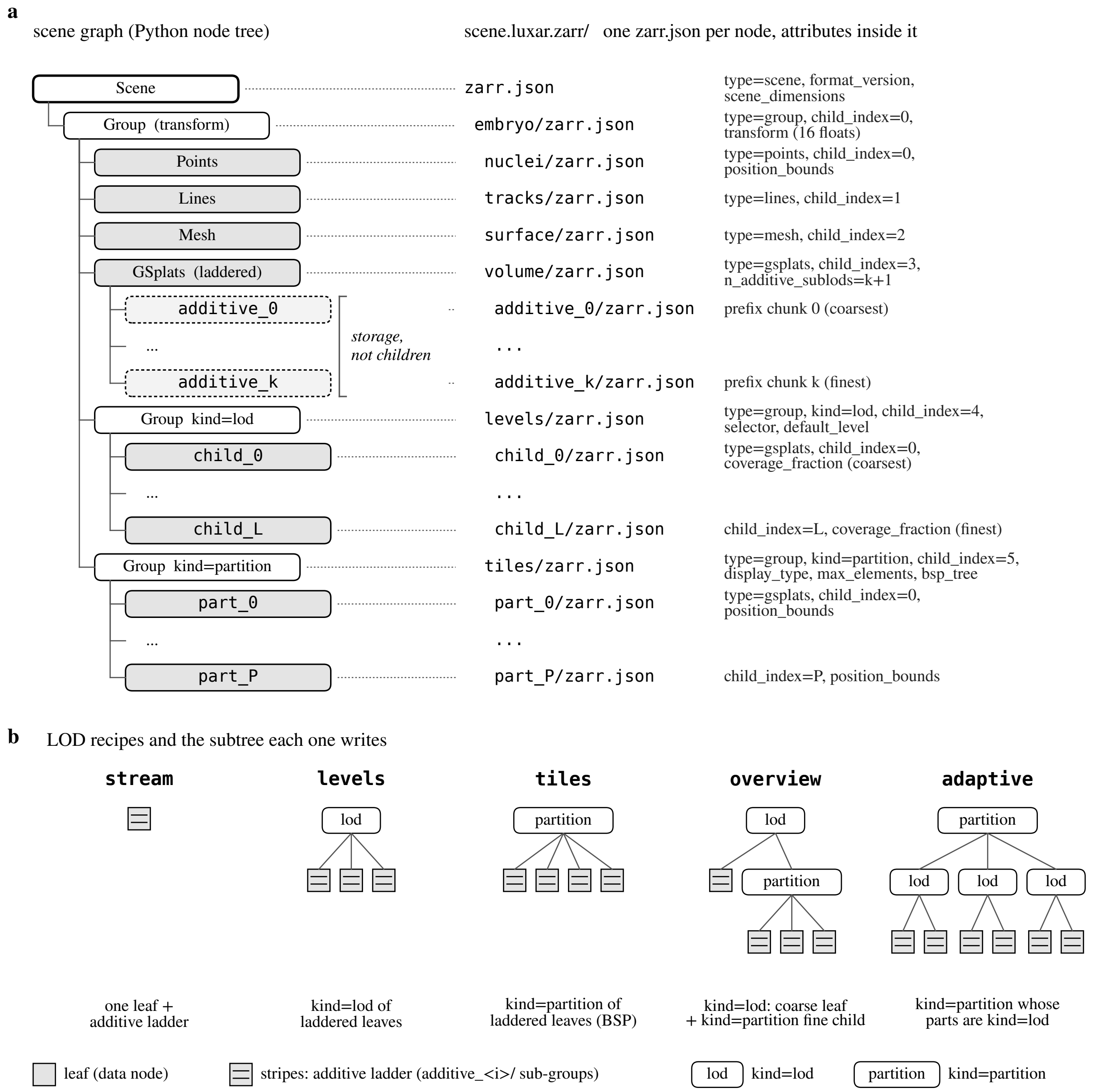

Figure 1: Schematic of the scene graph and its Zarr mirror. (a) The Python node tree of a small scene (left) and the group path plus the attributes each node persists (right); every node is one Zarr group with one `zarr.json`. The dashed `additive_<i>/` sub-groups of the laddered GSplats leaf are storage of that leaf and not children of the graph; `child_<i>/` under a `kind=lod` group and `part_<i>/` under a `kind=partition` group are children, ordered by `child_index`. (b) The subtree each level-of-detail recipe of the command line writes; a striped leaf carries an additive ladder.

Table 3: Common node attributes persisted to Zarr. Only `type` is mandatory. The rendering uniforms (`opacity`, `absorption`, `gamma`, `intensity`, `offset`, `blending_mode`, `colormap`) double as both authoring defaults (set in Python) and runtime state (modified by Layers-panel controls in the viewer). They compose along the parent chain as described in §4.3.

| Attribute | Type | Purpose |
| --- | --- | --- |
| `type` | string | node discriminator: `scene`, `group`, `points`, `lines`, `mesh`, `gsplats`, `sound` (a node but not a geometry) |
| `child_index` | int | insertion order among siblings, stamped by the node constructor on every non-root node. The viewer sorts siblings by it (a node without it sorts last, in enumeration order), so the Layers panel and `part_<i>` / `child_<i>` subgroups follow authoring order, not the store's alphabetical enumeration |
| `transform` | 16-float list | 4×4 spatial transform, column-major (THREE.js convention) |
| `nd_transform` | dict | per-dimension affine or permutation; see §3 |
| `visible` | bool | initial visibility (authoring-time default) |
| `extend_to_all` | list of strings | names of the non-displayed dimensions along which the node is not sliced (visible at every value); the Python `add_*` shorthand `"all"` is expanded to the scene's non-displayed dimension names before writing, so the stored value is always a resolved list |
| `layer` | bool | expose this node in the viewer's Layers panel |
| `opacity` | float [0,1] | uniform multiplier on alpha |
| `absorption` | float $\geq 0$ | absorption coefficient $\kappa$ of the `volumetric` blend mode (default 1) |
| `gamma` | float [0.1,10] | display gamma exponent |
| `intensity` | float [0,100] | display-range gain (see §4); this bound and the one on `offset` are checked by the Python validator when the attribute is authored, and the viewer clamps only the composed value, as Table 5 states |
| `offset` | float [−10,10] | display-range offset (see §4) |
| `blending_mode` | string | one of `additive`, `normal`, `max`, `opaque`, `luminous`, `volumetric` (emission–absorption; refused by meshes) |
| `colormap` | string | the name of a viewer built-in colormap, resolved against the viewer's registry; any other name (a matplotlib or colorcet colormap) and any $m \times 3$ LUT array are baked by the writer into a $256 \times 3$ LUT stored as `"custom"` with a sibling `colormap_lut` array |
| `layer_order` | int | cross-layer draw order; higher draws nearer the camera |

`overview` is a `kind=lod` group whose coarse child is one merged leaf and whose fine child is a `kind=partition`; `adaptive` is a `kind=partition` whose parts are themselves `kind=lod` groups. A partition's parts need not be spatial: the compiled zebrafish time-lapse scene of Supplementary Document 14 is a `kind=partition` with one part per timepoint, each part a laddered leaf whose `position_bounds` span a single time value, so the same culling that skips off-screen spatial tiles skips every timepoint but the current one (the *Drosophila* and neuromast scenes keep all frames in one node, whose chunks span several timepoints). A standalone `.gsplats.zarr` file is exactly one such subtree (a leaf, a `kind=lod` or a `kind=partition` root) with the gsplats-format header, which is why the viewer opens it directly and why a scene can graft it in without rewriting.

Two selectors read those attributes. When every child of a `selector="screen-area"` group carries `level_stats.median_footprint` and its `level_stats.footprint_dims` name exactly the displayed dimensions (as a set, in any order), the viewer projects each level's median footprint, scaled by the largest axis scale of the group's world matrix, at the group's bounding-box centre into CSS pixels and selects the *coarsest* level whose projected median footprint is at most the limit 1.5 px (`MAX_MEDIAN_FOOTPRINT_PX`) divided by the square root of the session's LOD bias (`lodBias`, an area factor, 1 by default, so the limit is 1.5 px unless the user sets it); the finest level is chosen only

Table 4: Attributes and children that specialise the root and the group kinds. The full field lists are in LUXAR_ZARR_FORMAT.md (scene format 0.2) and GSPLATS_ZARR_FORMAT.md (gsplats format 3.4).

| Node | Additional attributes and children |
|---|---|
| scene root | format_version ("0.2"), format_type ("luxar_zarr"), luxar_software_version (provenance, excluded from the digest), content_hash (post-order xxhash64 digest the viewer validates its cache against), scene_dimensions (the dimension table: name, unit, range, step, display, discrete, cyclic, scale, spatial, description, optional categories); optional citation, viewer_config; an overlays/ group (§5) |
| geometry leaf | element count (n_points, n_splats, ...), ndim, presence flags (has_colors, has_scalars, has_labels, has_keys, ...), position_bounds; the data arrays and the spatial-index arrays |
| additive ladder (on a leaf) | n_additive_sublods; sub-groups additive_<i>/ (coarsest first), each holding the next prefix chunk of the same splat set, so paging in sub-groups $0..i$ yields a prefix (Supplementary Document 9) |
| kind=lod group | type=group, kind=lod, display_type, selector ("screen-area" or "coverage"), default_level (0 = coarsest); children child_<i>/ coarsest to finest, each with a coverage_fraction switch threshold in the selector's units; a substitutive gsplat level also carries a level_stats block with median_footprint and footprint_dims (written by gsplats/lod/substitutive.py) (Supplementary Document 8) |
| kind=partition group | type=group, kind=partition, display_type, max_elements, optional bsp_tree (split axis is a position-column index); children part_<i>/, each a leaf or itself a kind=lod / kind=partition group, all resolving to the wrapper's display_type (a partition is homogeneous), frustum-culled by their own position_bounds |
| sound node | spatial, trigger, delay_ms, gain, bus, fades, and mandatory provenance (license, attribution, source_url); an optional positions array and the clip as a plain store key |

when no level qualifies, and the selector moves to a coarser level only once that level's footprint is 10% below the limit (scene/lod-selector-math.ts, pickChildByFootprintWithHysteresis, called from scene/lod-group-registry.ts). Where a stamp is missing on any child, the dimensions do not match, the centre has no finite projection (it lies at or behind the camera), or the group is a coverage-selector ladder, it falls back to the bounding-box selector, whose metric depends on the selector attribute. For a screen-area group the metric is the fraction of the viewport area that the group's projected bounding box covers. For a coverage ladder it is the projected length of the bounding box's diagonal, normalised by the axis the ladder was fitted against. Either metric is compared with the children's coverage_fraction thresholds, which are stored in that selector's units, with the same asymmetric hysteresis. An overview ladder, whose fine child is a partition and carries no footprint, always takes this path.

#### 2.3 Streaming write

The Python scene-graph object never holds bulk data. When a user calls a method like add_points on a group, the positions array is handed to the writer immediately and serialised to Zarr; only metadata (attributes, shape, dtype) is retained on the in-memory node. As a consequence, datasets that exceed available RAM compile naturally on a laptop provided each node does: the working set during compilation is the arrays handed to one add_* call (which the writer reorders along the *spatial index* and chunks before it returns), plus the scene-graph metadata, which grows with the number of nodes, not with the number of elements. This is the same property that makes the `LuxarZarrCompiler` usable inside a `with` block: the writer's `__exit__` method performs final consolidation and overlay auto-injection (see §7); the scene-graph object can be discarded immediately afterwards.

### 3 Spatial and nD transforms

#### 3.1 Spatial $4 \times 4$ transforms

Every node carries an optional $4 \times 4$ `transform` that maps its local frame into its parent's frame. The world transform of a leaf is the product of all transforms along the parent chain, computed with right multiplication: $T_{\text{world}} = T_{\text{root}} \cdot T_{\text{group}} \cdot \ldots \cdot T_{\text{leaf}}$, so for a column vector the leaf transform acts on coordinates first and the root transform last (a leaf scale of 2 under a root translation of $+10$ maps $x = 1$ to 12, not 22). The Python `world_transform` property collects the chain leaf-to-root and composes it in exactly this order. The viewer computes this product once per node at scene-graph load time and again whenever a transform attribute changes; geometry data itself is never re-transformed.

The on-disk representation is a flat 16-element list in **column-major** order, matching the layout of THREE.js's `Matrix4` and *not* NumPy's row-major convention. A naive serialisation that flattens row-major would silently swap row and column and place the translation in the wrong slots. The Luxar Python authoring layer transposes at the boundary: input $4 \times 4$ NumPy matrices are converted via $M \mapsto M^\top$ before flattening, and the inverse transposition is applied on read. Translation lives at indices $\{12, 13, 14\}$ in the on-disk list (the THREE.js convention); in NumPy row-major it would be at $\{3, 7, 11\}$. The viewer loader refuses a matrix whose translation sits at $\{3, 7, 11\}$ with $\{12, 13, 14\}$ zero, and also one with non-zero values in both slot sets, which it reports as ambiguous (`rendering/node-factory/validation.ts`, `validateTransformFormat`). Figure 2 shows the slot layout and the composition order on a one-dimensional example.

#### 3.2 Per-dimension transforms

Non-displayed dimensions are handled by a separate `nd_transform` attribute, a dictionary keyed by dimension name. Two value shapes are supported, depending on the dimension's type as declared at the scene level:

- **Continuous and discrete (ordinal) dimensions:** an affine $\{\texttt{scale} : s, \texttt{offset} : o\}$ mapping local-space coordinate $x$ to world-space coordinate $sx + o$. For a discrete (ordinal) dimension the world value is additionally rounded onto the dimension's declared grid $\text{range}[0] + k \cdot \text{step}$, i.e. the rounding is in units of the step and anchored at range[0], not at zero (§3.3). At the pinned commit a scale of 0 is flagged as a warning during validation (it collapses every value of the dimension to the same point) and not rejected. Such a transform has no inverse, and the viewer's inverse query (§3.3) leaves that dimension's query untransformed, so a zero-scale node is sliced as if it carried no transform on that dimension, and is not collapsed onto $o$. Zero scale should therefore be treated as unsupported.

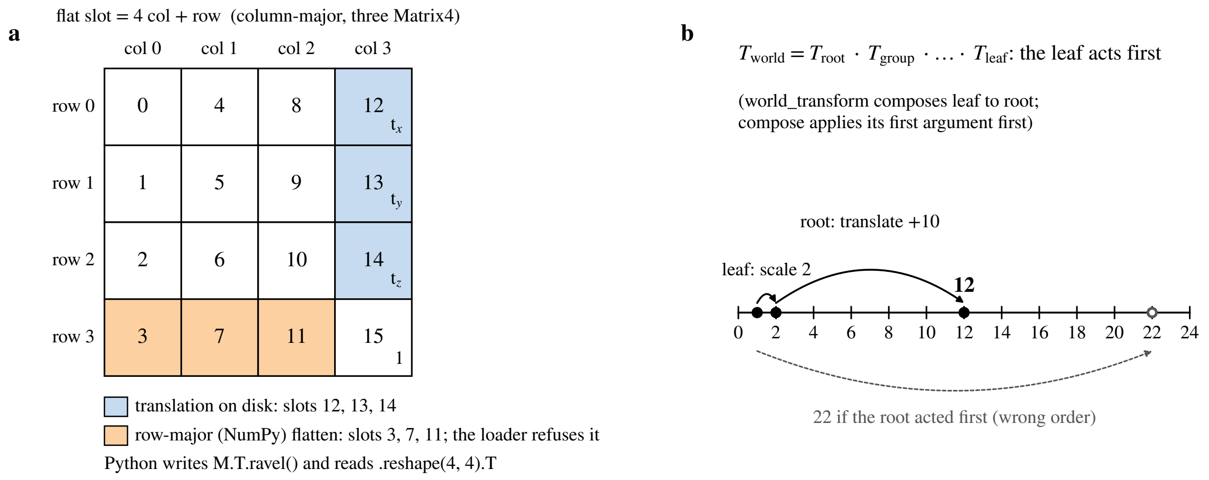

Figure 2: Schematic of the on-disk $4 \times 4$ transform. (a) The flat slot of entry (row, col) is $4\,\mathrm{col} + \mathrm{row}$; the translation occupies slots 12, 13 and 14 (blue), and a row-major flatten would put it in slots 3, 7 and 11 (orange), which the loader refuses. (b) Composition order along the parent chain: the leaf transform acts first, so a leaf scale of 2 under a root translation of +10 maps $x = 1$ to 12; 22 would be the result of the wrong order.

- **Categorical dimensions:** a permutation $\{\texttt{permutation} : [\pi_0, \pi_1, \ldots]\}$, where the list must be a valid permutation of $[0, n-1]$ and $n$ is the number of categories declared at the scene level. The permutation reorders category labels in the local-to-world direction: entry $i$ gives the world-space category index of local category $i$, world $= \pi[\mathrm{local}]$. For $\pi = [1, 2, 0]$, local category 0 is shown as world category 1, and world category 0 is local category 2.

Affine and permutation transforms cannot be mixed for the same dimension; the validator rejects the combination at write time.

Like spatial transforms, `nd_transforms` compose along the parent chain. For affines, $\{s_p, o_p\} \circ \{s_c, o_c\} = \{s_p s_c,\ s_p o_c + o_p\}$. For permutations, $\pi_p \circ \pi_c$ has $i$-th entry $\pi_p[\pi_c[i]]$. The composed transform is what the viewer uses when interpreting a slice position in world space.

##### 3.3 Inverse-query algorithm

The runtime never transforms point coordinates. Instead, the spatial index stores raw (untransformed) values and the slice query (slice position $\pm$ tolerance, per dimension) is transformed *into* the local frame at query time. This trades $O(N)$ per-point work at every camera change for work proportional to the number of non-displayed dimensions at every slice change (constant per affine dimension; one pass over the category list per categorical dimension), an overwhelming win when $N$ is in the millions and the dimensionality is small.

For an affine $\{s, o\}$ with $s \neq 0$ on a continuous dimension, the inverse query is

$$x_{\mathrm{raw}} = (x_{\mathrm{world}} - o)/s, \tag{1}$$

$$\Delta x_{\mathrm{raw}} = \Delta x_{\mathrm{world}}/|s|, \tag{2}$$

where $\Delta x$ is the half-width of the slice tolerance band along this dimension. The absolute value on the scale handles negative scales (axis flips) without changing the band width. A dimension named in the node's `extend_to_all` is not sliced at all; the viewer marks its tolerance with a reserved sentinel value standing for an unbounded band, and (2) passes that sentinel through unscaled.

On a discrete dimension the forward map rounds onto the declared grid ($\text{range}[0] + k \cdot \text{step}$), so the bare affine inverse (1) is not the answer. The viewer tests the two local grid points bracketing $(x_{\text{world}} - o)/s$ and keeps the one whose forward image rounds to the queried world value. Both candidates can qualify only when $|s| < 1$, since only then do two adjacent local grid points map to world values less than one step apart; in that case the nearer one is kept. The local query is then snapped onto the chosen grid point so that the downstream membership test, which accepts a local value lying within half a step of the query (the half-step window), selects exactly one category. When neither candidate qualifies, the queried world value is the image of no local value and the node renders nothing for that slice. Two examples: with $s = 2$, $o = 0$ and unit step, world value 7 has candidates 3 and 4 whose images are 6 and 8, so nothing is drawn (instead of querying local 3.5, which would admit two categories); with $s = 1$, $o = 0.4$, world value 7 resolves to local 7 (since $\text{round}(7.4) = 7$), not to 6.6. Figure 3 draws the affine inverse query and both discrete cases. Dimensions named in the node's `extend_to_all` are exempt, and a permutation always has exactly one preimage. For a permutation $\pi$, the inverse query builds the inverse permutation $\pi^{-1}[\pi[i]] = i$ when it derives the node's query (one pass over the category list per derivation, not a cached inverse; category counts are small, so the cost is negligible). It maps the queried world-space category index through $\pi^{-1}$ to recover the local-space index.

A subtle property of this design is that the spatial index can be built once at compile time and never invalidated: changing any `nd_transform` only changes the query, not the index. The same property holds for per-node opacity, blending mode, etc., so the entire visibility-and-styling state of the viewer is mutable in place without touching the underlying chunk catalogue.

### 4 Layers

#### 4.1 The `layer` flag

Not every node is interesting to expose to a user. Internal grouping nodes, helper layers, and metadata-only annotations are kept out of the Layers panel by default. The `layer` attribute (default `False`) is the explicit opt-in: nodes that authors mark with `layer=True` populate the panel; everything else is hidden. Groups can be flagged as layers, in which case the panel exposes them as composite controls that fan changes out to all descendants (see §4.3).

#### 4.2 Layer-panel uniforms

The Layers panel exposes per-layer controls for visibility, opacity, absorption (the `volumetric` mode's $\kappa$), display range, gamma, blending mode, and colormap. Of these, the display range needs special handling: it is a UI affordance (drag the two thumbs of a range slider, or type the bounds) but the GPU needs a pair of shader uniforms. For a non-degenerate window ($\max \neq \min$) the

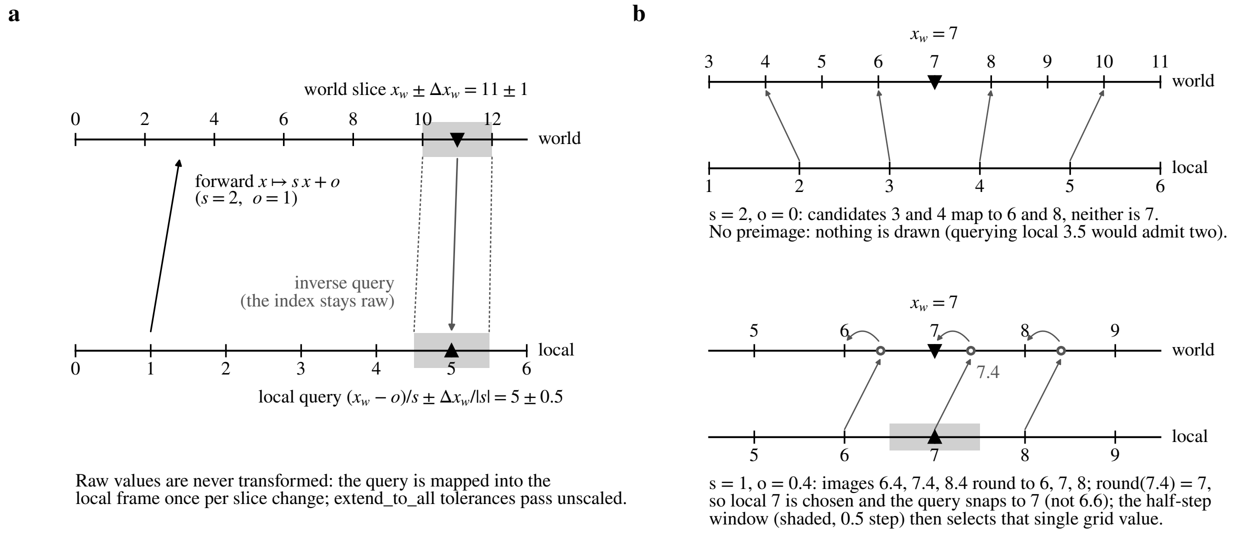

Figure 3: Schematic of the inverse slice query. (a) An affine `nd_transform` with $s = 2$, $o = 1$ on a continuous dimension: the world query $11 \pm 1$ is mapped once into the local frame as $(x_w - o)/s \pm \Delta x_w/|s| = 5 \pm 0.5$; the raw index is never transformed. (b) The discrete grid rule on the two examples of the text. Top: $s = 2$, $o = 0$ and unit step; the two local candidates bracketing 3.5 map to 6 and 8, neither rounds to the queried 7, so nothing is drawn. Bottom: $s = 1$, $o = 0.4$; the image 7.4 of local 7 rounds to 7, so the query snaps to local 7 and the half-step membership window (shaded) selects that single grid value.

viewer maps the $(\min, \max)$ pair to

$$\texttt{intensity} = 1/(\max - \min), \tag{3}$$

$$\texttt{offset} = -\min/(\max - \min), \tag{4}$$

so that the in-shader operation $\texttt{value} \cdot \texttt{intensity} + \texttt{offset}$ sends the two window endpoints to 0 and 1 regardless of how the data was originally normalised; values outside the window land outside $[0, 1]$. A degenerate window ($\max = \min$, e.g. a constant amplitude range) falls back to the identity pair $(1, 0)$ instead of dividing by zero. These per-layer controls are distinct from the global exposure, offset and gamma of the fused post-processing pass (§8.2), which act on the whole frame after every layer has been composited.

The pair of (3)–(4) is consumed differently on the two colour paths. A node rendering direct colours applies it to the colour itself, clamped below at zero but not above, so values brighter than the window remain HDR and are handled by the tone-mapping stage (§8); the layer's gamma is applied to the colour as $\texttt{value}^{1/\gamma}$ after windowing. A node rendering through a colormap receives the window instead as a scalar range: the per-element scalar is windowed and clamped to $[0, 1]$, gamma is applied to that scalar before the lookup-table fetch ($LUT$), and the post-lookup colour gain and offset are reset to 1 and 0 so the window is not applied twice. The `blending_mode` selects among additive, normal, max, opaque, luminous, and volumetric (emission–absorption compositing, not available on meshes) blend functions; the choice is per-layer and propagates immediately when changed.

Table 5: Inheritance of appearance attributes along the parent chain (`data/attrs-composer.ts`). The scene root (`type=scene`) is excluded from the chain.

| Attribute | Composition | Clamp / default when unset anywhere |
|---|---|---|
| `opacity` | product over the chain | clamped to $[0, 1]$; default 1 |
| `absorption` | product | clamped to $\geq 0$; default 1 |
| `gamma` | product | clamped to $[0.1, 10]$; default 1 |
| `intensity` | product | clamped to $\geq 0$, no upper clamp (the authored $[0, 100]$ of Table 3 is a write-time check); default 1 |
| `offset` | sum | not clamped (the authored $[-10, 10]$ of Table 3 is a write-time check); default 0 |
| `blending_mode` | nearest setter (leaf-most) wins | `additive` for points, lines and gsplats; `opaque` for meshes |
| `colormap` | nearest setter wins (its LUT travels with it) | direct colours |
| `layer_order` | nearest setter wins | unset (band 0; inferred containment order) |

##### 4.3 Multi-select and group-as-composite layers

The panel supports three click modes: a single click selects one layer, ctrl-click toggles a layer in or out of the selection, and shift-click range-selects from the last clicked anchor. Once a multi-selection is active, every uniform change applies to all selected layers simultaneously, a useful idiom for matched display ranges across channels of a multi-channel volume.

When a group is exposed as a layer (with `layer=True` on the group itself), changes to its uniforms are fanned out to all data descendants. The group does not have its own GPU material; it is purely a control affordance. This is the natural way to expose, e.g., all four channels of a single time point as a single "time point" layer with shared opacity.

A group's values do not overwrite what its descendants author; the two compose along the root-to-leaf chain by the rules of Table 5, which apply equally to authored attributes and to Layers-panel edits (a panel edit on a group layer recomputes every descendant's effective value through the same rule). Unset values are the identity (a gain of 1, an offset of 0), the scene root contributes nothing, and the composed multiplicative values are clamped once at the end. Two examples: a group with `opacity=0.5` over a leaf with `opacity=0.5` renders at 0.25; a group `offset` of 0.1 over a leaf `offset` of 0.2 gives 0.3. Writing $I$ and $O$ for the composed intensity and offset uniforms, the additive offset rule is deliberately not the chaining of the shader's $\texttt{value} \cdot I + O$ model through successive nodes. Chaining it, with the parent's map applied before the child's, would give a composed offset of $O_{\text{parent}} I_{\text{child}} + O_{\text{child}}$; the simpler rule is what the Python API documents and the viewer implements.

#### 5 Overlays

##### 5.1 Four overlay types

Beyond the rendered geometry, the viewer carries a screen-space annotation layer rendered as a stack of HTML `<div>` elements positioned absolutely above the WebGL canvas. Four types are supported, distinguished by their type attribute under the Zarr archive's overlays/ group:

- overlay_text: rendered via textContent, which is safe against cross-site scripting (XSS) by construction, since the browser performs no HTML parsing on it. Supports font presets (sans / serif / mono), font size, colour, alignment, line height, optional background and padding, and a stroke effected via a four-directional CSS text-shadow.

- overlay_image: a static image fetched from overlays/<name>/<image_file> inside the Zarr archive. Sizing uses viewport-relative units so that the overlay scales with the canvas.

- overlay_video: a WebM or MP4 clip stored verbatim under the overlay's group. The container signature (an *EBML* header for WebM, an ftyp box for MP4) is checked at authoring, so any other container fails in Python and not on the display; whether the clip's codec and profile decode remains the browser's decision. The clip is rendered as a <video> that is muted and looping by default (muted and loop are authoring parameters; autoplay=True requires muted=True) and that is played or paused with its own visible_range, so clips whose visibility ranges are mutually exclusive decode one at a time (clips visible together each decode; nothing arbitrates between them); autoplay=False shows native controls instead. Transparency is authored as a stacked alpha matte (alpha_matte="stacked": colour on top, the alpha channel as a grey matte below, one opaque frame twice as tall) that the viewer recombines in a small shader, because a codec's alpha plane plays transparent only in some browsers and is silently dropped by others.

- overlay_html: rendered via innerHTML after a client-side allowlist sanitisation. The allowed tag set is roughly that of a static blog post (<b>, <i>, <a>, <table>, <img>, <ul>, <p>, <h1 ... h6>, <sub>, <sup>, <code>, <pre>, ...); event-handler attributes (on*) and javascript: URLs are stripped. The allowed set is conservative; users who need richer dynamics should embed an <iframe> in their hosting page instead of trying to extend the overlay sanitiser.

#### 5.2 Anchoring, opacity, blend, transitions

All four overlay types share a common positioning and styling block:

- **Position:** normalised screen coordinates $(x, y) \in [0, 1]^2$ with origin at the top-left, mapped to CSS percentages on the overlay <div>.

- **Anchor:** a 9-point grid (top-left, top-center, ..., bottom-right); the anchor controls which point of the overlay coincides with the position (Fig. 4a). The six left and centre anchors are realised with a CSS left percentage and a transform: translate(...) that shifts the box by 0, $-50\%$ or $-100\%$ of its own width and height. The three right-hand anchors are instead pinned with right: $(1 - x) \cdot 100\%$ and a transform whose horizontal component is zero, so that a wrapping overlay measures its available width from the container's edge and not from its pre-transform position.

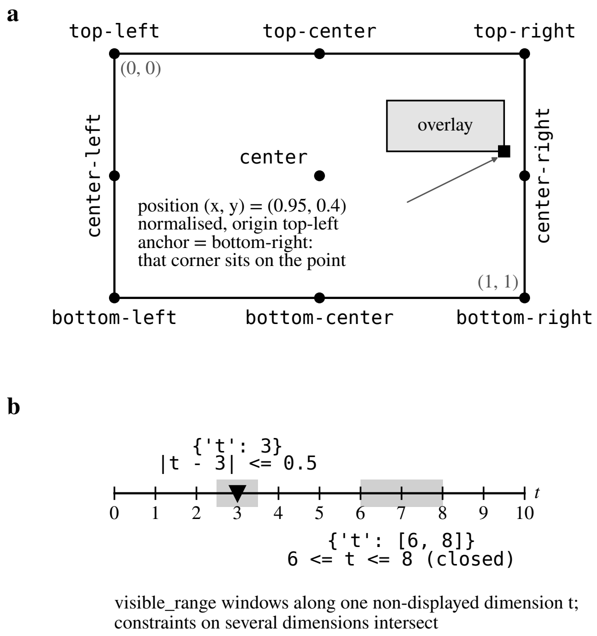

Figure 4: Schematic of overlay placement. (a) The nine anchors on the normalised canvas, origin top-left; an overlay at position $(0.95, 0.4)$ with anchor `bottom-right` puts that corner on the point. (b) Two `visible_range` windows along one non-displayed dimension $t$: the exact value `3` keeps the overlay visible while $|t - 3| \leq 0.5$, the list `[6, 8]` over the closed interval; constraints on several dimensions intersect.

- **Opacity:** a $[0, 1]$ value applied via CSS `opacity`.
- **Blend mode:** the Luxar overlay blend names map to CSS `mix-blend-mode` values: `normal`, `multiply`, `screen`, `overlay`, `additive` (`plus-lighter`), and `difference`.
- **Transitions:** either `none` or `fade` with a configurable duration, implemented via a CSS opacity transition.
- **Interactivity:** a non-interactive overlay sets `inert=true` and `pointer-events: none` so it cannot intercept mouse events meant for the canvas.

##### 5.3 Dimension-aware visibility

Overlays often pertain to a particular slice or range along a non-displayed dimension. The `visible_range` attribute is a dictionary keyed by dimension name; its values are either an exact target value (in which case the overlay is visible whenever the slider is within $\pm 0.5$ of that value) or a $[\min, \max]$ two-element list (visible when the slider falls in the closed interval; Fig. 4b). Multiple dimensions are combined by intersection: the overlay is visible only when every constraint is satisfied. A key naming a dimension the scene does not declare is skipped. This rule governs ordinary overlays. The hover overlays of §7 (`hover=True`) ignore `visible_range`: their visibility follows the pick result, subject only to the global hide toggle. A scene may author a guided tour as a sequence of story waypoints, camera poses (with their slice positions) that the viewer flies between. A waypoint authored with `reveal="on_arrival"` holds back, while its camera flight is in transit, an overlay whose range has only just become satisfied, until the flight lands; under the default `reveal="immediate"` the overlay appears during the flight.

The `visible_range` rule makes overlays useful for genuinely multidimensional data: an annotation tied to time point 17 of a 4D acquisition can be authored once and will appear and disappear automatically as the user sweeps the time slider, without the author having to write a controller.

### 6 GPU picking and hover annotations

#### 6.1 Pick-buffer encoding

The cursor needs to find which element it is over without iterating over the scene on the CPU. Luxar implements this via a hardware picking pass: each registered pickable node is rendered to an off-screen `RGBA32F` render target with materials that encode identity instead of colour (a float target that both the WebGL and the WebGPU backend read back through one asynchronous path). The encoding is (Fig. 5a)

- `R` channel: per-node ID, allocated centrally by the picking system and bounded by $2^{24} - 1$ so that consecutive IDs stay exactly representable in float32; a value below 0.5 marks background.
- `G` and `A` channels: the per-element index within the node (point index, line segment slot, splat index, mesh vertex index) split into its low and high 16 bits, so that indices above the $2^{24}$ float32 integer limit survive; the reader reconstructs $\text{round}(A) \cdot 65536 + \text{round}(G)$.
- `B` channel: per-fragment brightness in $[0, 1]$, derived from the same falloff function the rendering material uses (Gaussian weighting for splats, radial falloff for points, taper for lines, coverage alpha for meshes).
- `gl_FragDepth`: $1 -$ brightness for points, lines, and Gaussian splats and meshes in the commutative blend modes (`additive`, `max`, `luminous`, and `volumetric`); the projected surface depth for Gaussian splats and meshes in the `normal` and `opaque` modes, where the user sees an occluding surface and the front-most fragment must win.

The depth-from-brightness trick makes the buffer composable for emissive geometry: when two pickable elements overlap at a pixel, depth testing keeps the brighter one. Without this, picking would either pick whatever the draw order put on top last (unstable under camera changes) or require a separate sort pass. For a depth-sorted surface the same mechanism, fed the true depth, keeps the visible fragment instead.

To bound its cost, the buffer is rendered at *half* the drawing-buffer resolution, further capped at 1024 pixels on its longer side with the aspect ratio preserved: a $3840 \times 2160$ drawing buffer picks through a $1024 \times 576$ target, one pick pixel per $3.75 \times 3.75$ canvas pixels (Fig. 5d). The

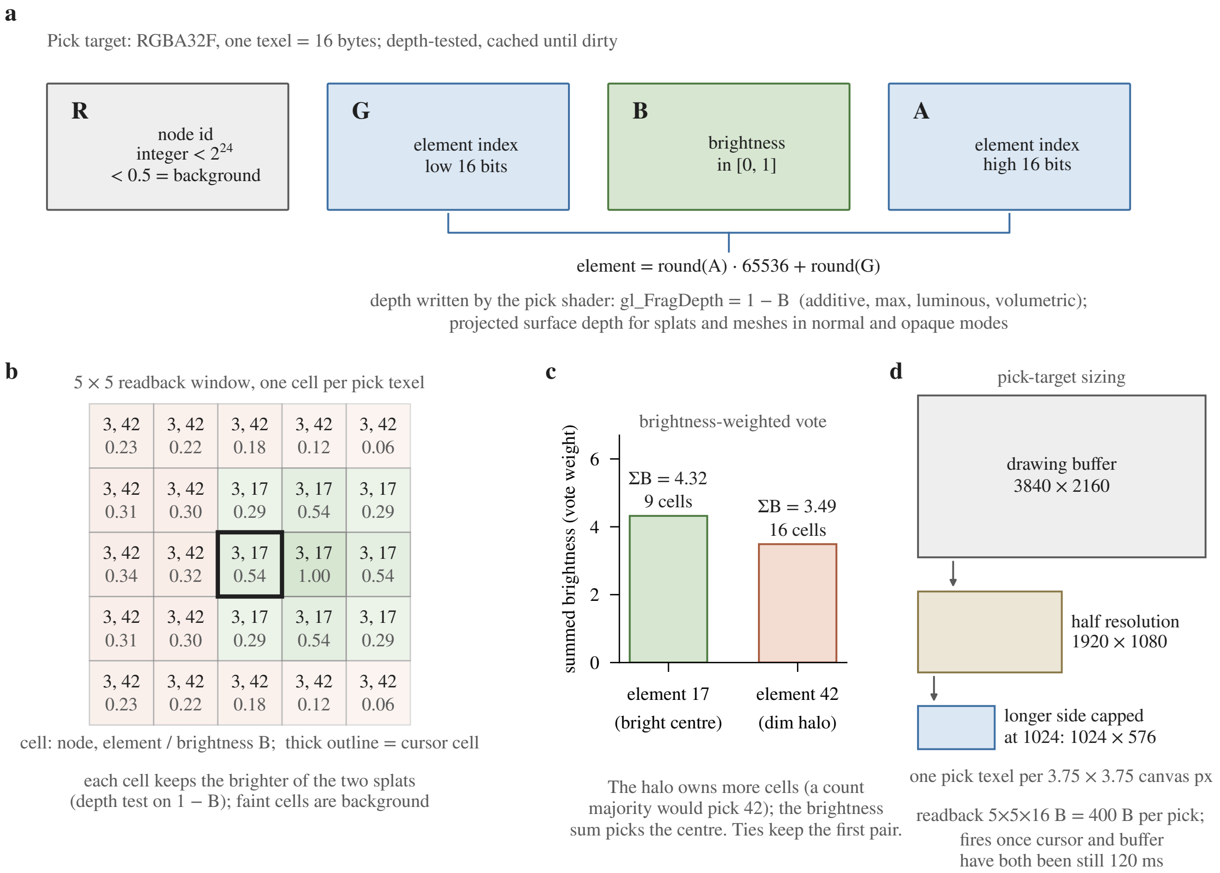

Figure 5: Schematic of GPU picking. (a) Channel layout of one `RGBA32F` pick texel: node id in R (below 0.5 is background), element index split across G (low 16 bits) and A (high 16 bits), brightness in B; the pick shader writes $1 - B$ as depth for the commutative blend modes and the projected surface depth for `normal` and `opaque`. (b) A $5 \times 5$ readback window over two overlapping splats of node 3; each cell shows the (node, element) pair that won the depth test and its brightness, and the thick outline is the cursor cell. (c) The brightness-weighted tally: element 42 owns 16 cells, but element 17 wins with the larger summed brightness. (d) Pick-target sizing for a $3840 \times 2160$ drawing buffer: half resolution, then the longer side capped at 1024 px.

IDs are integers and do not need sub-pixel precision; the coarseness is largely masked by the voting step described next, though on a dense scene at high resolution the cursor neighbourhood it samples is correspondingly wider.

#### 6.2 Brightness-weighted majority voting

When the cursor moves, the picker reads back a $5 \times 5$ pixel neighbourhood around the cursor's location in the half-resolution buffer. Each of the 25 pixels contributes a vote: the (node, element) pair encoded at that pixel, weighted by its brightness channel. The winner is the (node, element) with the highest total brightness; an exact tie keeps the pair encountered first in the readback order. Figure 5b,c works one such vote through.

The brightness weighting gives the picker intuitive behaviour on Gaussian splats: the centre of a splat is brighter than its halo, so a hover near the centre selects the splat even if the cursor is geometrically over the halo of an adjacent splat. Without brightness weighting, hovering over any pixel of a halo would select the halo's owner regardless of how dim it was at that pixel.

A ray–box pre-cull against each node's $AABB$ skips the readback entirely when the cursor's ray does not intersect any registered node's bounding box. This matters because the GPU readback is the most expensive part of the pipeline: even asynchronous, it is a GPU-to-CPU transfer per pick, and issuing one on every mouse move would compete with rendering.

##### 6.3 Cache and settle scheduling

The pick buffer is cached and re-rendered only when it is marked dirty: a camera pose or projection change, a resize, a geometry change, or an appearance or visibility change from the Layers panel (the picker also re-renders when the set of effectively visible nodes changes without a dirty event). On a static scene the buffer is dirty exactly once, after the initial render, and every subsequent hover event is just a $5 \times 5$ readback (400 bytes of `RGBA32F`). In place of a debounce, picks are scheduled by a two-axis settle rule evaluated once per animation frame: a pick fires only after *both* the cursor and the pick buffer (its last dirty event) have been still for 120 ms (`HOVER_SETTLE_MS`, the conventional tooltip delay) and something has changed since the last pick. When nothing has changed the loop idles until the next movement or dirty event re-arms it. The readback itself is asynchronous, and a result that a newer movement, dirty event or pointer-leave has superseded while it was in flight is discarded, not shown as a stale tooltip.

#### 7 Per-element labels and image labels

##### 7.1 Compressed-sparse-row encoding

For atlases with millions of elements, storing one fixed-width string per element is wasteful (most labels are short, a few are long) and storing one variable-length record per element fragments the I/O pattern. Luxar uses a compressed-sparse-row (CSR) layout: two parallel arrays, an offsets array of shape $(N+1)$ and a bytes array of shape $(\text{total\_bytes},)$, where the label of element $i$ is the byte range $[\text{offsets}[i], \text{offsets}[i+1])$ decoded as UTF-8.

Concretely the Zarr archive contains

- `label_offsets` (`uint64`, shape $N+1$): byte offsets into `label_bytes`.
- `label_bytes` (`uint8`, shape total): UTF-8 encoded label payload, concatenated.

A label is empty when $\text{offsets}[i] = \text{offsets}[i+1]$; the loader returns `null` in that case so that a UI bound to the loader naturally skips elements without labels. Figure 6 shows the layout on a four-element example.

The viewer's `LabelLoader` is lazy per node, not per element: on the first hover that lands on a node it reads both arrays in full, decodes all $N$ labels into a string array that is cached for the session (consecutive identical labels share one string), and coalesces concurrent requests for the same node. The retained cost is therefore the decoded labels of every hovered node, plus the $8(N+1)$-byte offsets array while decoding; for an atlas of millions of elements that is tens of megabytes per node. A parallel pair `key_offsets`/`key_bytes` (a machine-readable key per element, flagged by `has_keys`) goes through the same loader.

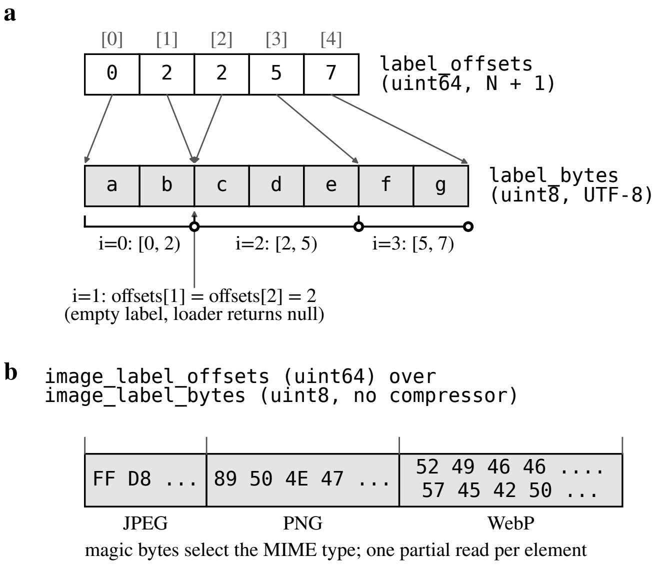

Figure 6: Schematic of the compressed-sparse-row label layout. (a) `label_offsets` (five entries for four elements) indexes into the concatenated UTF-8 `label_bytes`; element 1 has equal consecutive offsets, an empty label that the loader returns as `null`. (b) Image labels use the same pair over an uncompressed byte array of concatenated encoded images; the magic bytes select the MIME type and each image is fetched with one partial read.

#### 7.2 Image labels

Image labels (per-element thumbnails, e.g. a microscopy crop showing each cell of an atlas) use the same CSR pattern with a different payload type:

- `image_label_offsets` (`uint64`, shape $N + 1$).
- `image_label_bytes` (`uint8`): concatenated pre-compressed image blobs (JPEG, PNG, or WebP).

Image bytes are not Blosc-compressed because they are already in a content-aware codec; layering a generic compressor on top wastes CPU. The decoder identifies the format by magic bytes (`0xFF 0xD8` for JPEG, `0x89 0x50 0x4E 0x47` for PNG, `0x52 0x49 0x46 0x46` ... `0x57 0x45 0x42 0x50` for WebP; Fig. 6b) and creates a blob URL that the overlay's `<img>` tag consumes directly. Unlike text labels, image bytes are read per element: the offsets array is loaded once per node and each image is fetched with a partial-array read of its byte range. A least-recently-used (LRU) cache holds up to 50 MiB of *encoded* image payloads (the JPEG/PNG/WebP bytes, referenced through their blob URLs) at once; the browser's decoded raster and GPU memory are not counted against that budget. On eviction, the corresponding `URL.revokeObjectURL` is called so that the browser can reclaim the underlying memory.

#### 7.3 Auto-injection of default hover overlays

When at least one node carries label or image-label arrays, the scene finalisation step (called by the compiler's `__exit__` method) injects the matching default hover overlays, the text overlay when any node carries text labels, the image overlay when any node carries image labels, and both when both are present:

- `__hover_text`: an `overlay_text` positioned top-right with a 0.15 s fade transition; its template body is "`{hover_label}`".
- `__hover_image`: an `overlay_html` positioned top-right with a fade transition and a semi-opaque background; its template body is "`{hover_image_label}`", wrapped in an `<img>` tag with viewport-relative sizing.

The two are separate overlays because image loading is asynchronous: a single combined overlay would cause layout shift while the image fetched. Authors who want a different look can opt out by setting the scene's `suppress_hover_overlay` flag, or by providing their own overlay with `hover=True`. The auto-injection step detects any existing hover overlay and skips its own injection in that case.

The overlay manager substitutes five template variables on every pick result; a node's optional `link` and `copy` templates take four of them (`{hover_key}`, `{hover_label}`, `{hover_node}`, `{hover_index}`; not `{hover_image_label}`), with URL escaping in the former:

- `{hover_label}`: the text label for the picked element (HTML-escaped if the overlay is HTML, raw if it is text).
- `{hover_key}`: the element's machine-readable key from the `key_offsets`/`key_bytes` pair, kept separate from the label because a tooltip wants prose and a link wants a bare identifier.
- `{hover_image_label}`: only meaningful in HTML overlays; expands to an `<img>` tag wrapping the cached blob URL.
- `{hover_node}`: the scene-graph path of the layer the pick is reported against (for a `kind=partition` layer this is the outermost partition wrapper, the layer the user sees in the panel), useful for layered atlases where multiple node types are pickable.
- `{hover_index}`: the integer element index inside the leaf that was hit. Under a partition that leaf is a `part_<i>` child, so `{hover_node}` and `{hover_index}` refer to different nodes and do not together identify an element; an identifier meant to be stable outside the viewer should be authored as a per-element key and read through `{hover_key}`.

### 8 Post-processing pipeline

#### 8.1 HDR pipeline overview

The viewer renders into a `HalfFloatType` (16-bit float) render target, multisampled when MSAA is enabled (MSAA sets the target's sample count, 4 by default; MSAA, SSAA and FXAA, three of the *anti-aliasing (MSAA, SSAA, FXAA, SMAA)* techniques, are all *off* by default). This is the HDR domain: per-channel values can exceed 1.0, which is what bloom and tone-mapping operate on. The pipeline is a small custom stage sequence, not a library effect composer (Fig. 7):

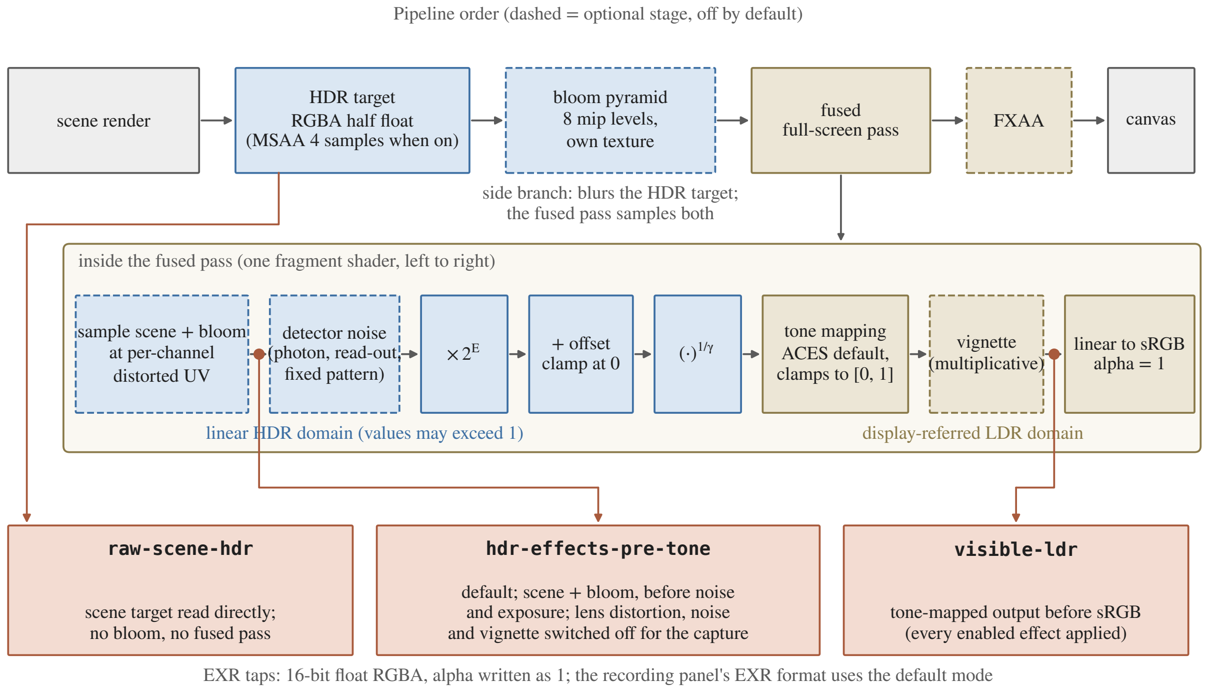

Figure 7: Schematic of the post-processing pipeline. Top row: the stage order from the scene render to the canvas; dashed stages are optional and off by default. Middle: the fused full-screen pass unfolded left to right, from the linear HDR domain (blue) through tone mapping into the display-referred LDR domain (tan). Bottom: the three EXR capture taps of §9.4 and the point of the chain each one reads; the default `hdr-effects-pre-tone` tap returns the scene plus bloom before noise and exposure.

1. **Scene render**: the scene is rendered into the HDR HalfFloat target.
2. **Bloom pyramid** (when enabled): a mipmap-based blur of the HDR target with configurable level count and threshold, rendered into its own texture.
3. **Fused full-screen pass**: one fragment shader samples the HDR scene plus bloom, applies the HDR-domain effects when enabled (chromatic distortion, a wavelength-dependent UV warp; simulated detector noise, a combined photon / read-out / fixed-pattern model), then exposure, offset and gamma, the tone-mapping operator (§8.2), the LDR-domain vignette, and finally the linear-to-sRGB display encoding.
4. **Anti-aliasing** (when enabled): FXAA as a separate final pass over the encoded LDR image; SSAA renders the whole chain at a higher resolution instead.

Depth of field, screen-space ambient occlusion, and SMAA are deliberately unsupported: the first two need depth-aware multi-pass blur or surface normals that point, splat, and line geometry do not provide, and SMAA's three-pass blend does not fit the single fused pass described next.

#### 8.2 Tone mapping with fused exposure, offset, and gamma

Chromatic distortion, detector noise, exposure/offset/gamma, tone mapping, and vignette are fused into one custom full-screen fragment pass. Three adjustments precede the operator-specific tone curve: exposure (in log-2 stops), an additive offset, and a gamma exponent, applied in that order. They are global, frame-wide controls, so their symbols carry the subscript g to keep them apart from the per-layer window and gamma of §4:

$$\mathbf{c}_1 = \mathbf{c}_0 \cdot 2^{E_\mathrm{g}}, \tag{5}$$

$$\mathbf{c}_2 = \max(\mathbf{c}_1 + o_\mathrm{g}\,\mathbf{1},\ \mathbf{0}), \tag{6}$$

$$\mathbf{c}_3 = \mathbf{c}_2^{1/\gamma_\mathrm{g}}, \tag{7}$$

$$\mathbf{c}_\mathrm{out} = \mathcal{T}(\mathbf{c}_3), \tag{8}$$

$$\mathbf{c}_\mathrm{disp} = \mathrm{sRGB}(\mathbf{c}_\mathrm{out}), \tag{9}$$

where $o_\mathrm{g}$ is a scalar offset replicated to the three channels, the clamp at zero keeps the fractional power defined, and $\mathcal{T}$ is one of LINEAR, REINHARD, CINEON, ACES filmic (the default), AgX, and NEUTRAL; "None" is an alias of LINEAR, and both clamp their output to $[0, 1]$, so neither is a raw-HDR passthrough. Raw HDR export does not run this stage at all: it uses a separate capture define that returns the linear scene sample before it (§9.4). Fusing the stages (5)–(9) into the same shader avoids one extra full-screen pass compared with a naive layered approach.

##### 8.3 Bloom and chromatic distortion ordering

Bloom is built on a mipmap-blur convolution; chromatic distortion is a per-channel UV warp. The two do not commute, so the fused pass fixes their order: the bloom pyramid is rendered first into its own texture, and the fused shader then samples both the HDR scene and the bloom texture at the same chromatically aberrated per-channel coordinates, so bloom is effectively added into the HDR buffer *before* the distortion samples from it. Detector noise, the stages (5)–(8), and the vignette follow in that order within the same pass, then the display encoding (9); FXAA, which needs the finished LDR edges, runs as a separate pass afterwards.

##### 8.4 Adaptive device-pixel-ratio scaling

A high-DPI display can render at more pixels than the GPU can sustain 60 FPS for. The adaptive DPR (device-pixel-ratio) controller observes the per-frame timestamps over a 1 s sliding window and re-evaluates every 500 ms. Instead of fixed frame-rate constants, its thresholds are set *relative to the display's estimated achievable refresh rate*, a live estimate of the fastest *rAF* cadence the monitor sustains. The render target is scaled down when the frame rate falls below a fraction of that cap (≈0.75×, i.e. 45 FPS on a 60 Hz display) and scaled back up above a higher fraction (≈0.9×, i.e. 54 FPS) once the configured `hysteresisSeconds` (default 3 s) has elapsed. Steps are multiplicative (down ×0.9, up ×1.05), not a single jump. Figure 8 lays the controller out as a state machine.

Upscaling can itself cost frames and trigger an oscillating DPR "U-shape," so each scale-down is *provisional*: the controller arms a short probe and, if throughput does not actually improve, reverts and records a *learned floor* that expires after 30 s, doubling on each re-learning up to a

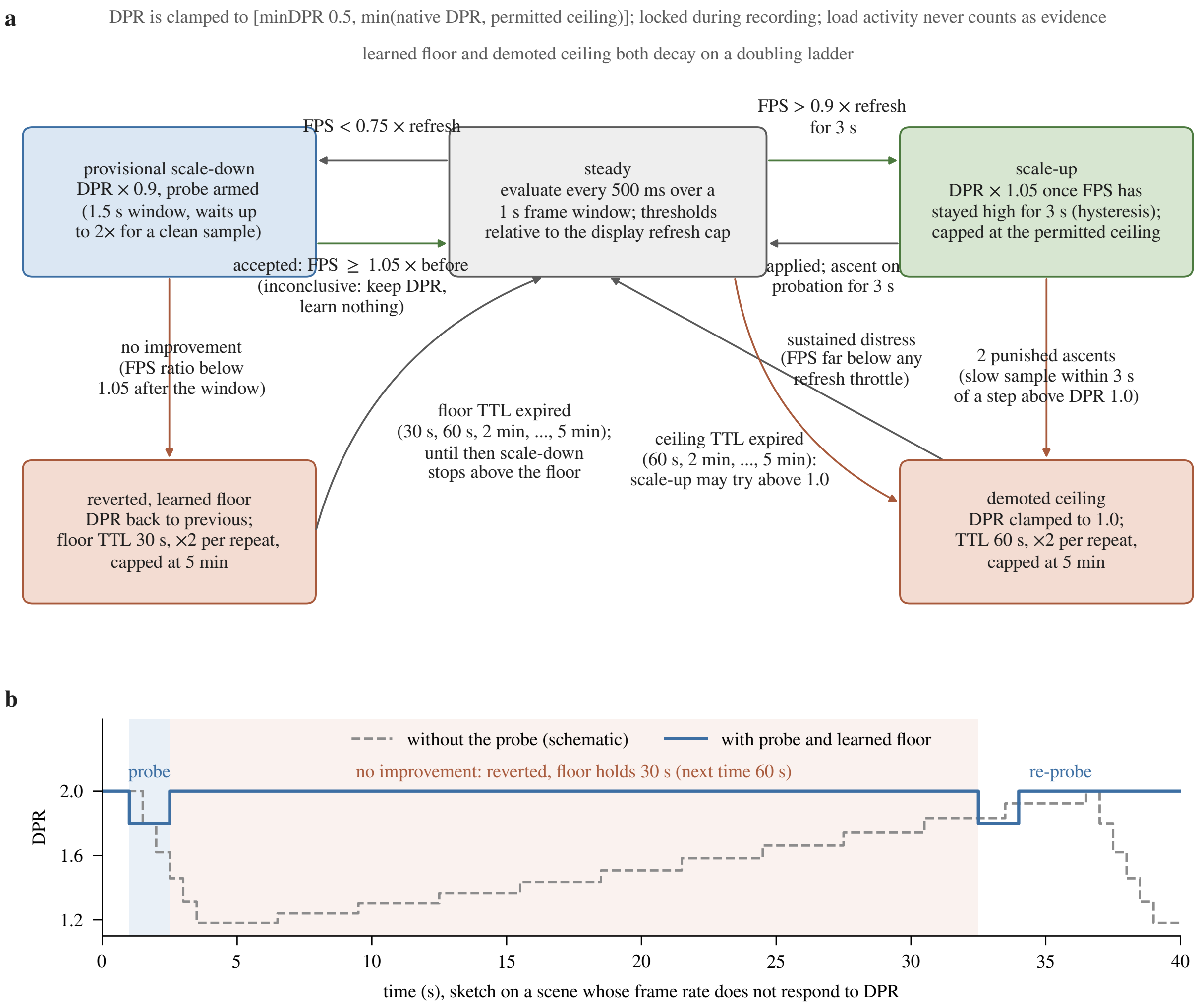

Figure 8: Schematic of the adaptive device-pixel-ratio controller. (a) States and transitions: the steady state evaluates every 500 ms over a 1 s frame window against thresholds set relative to the display refresh cap; a scale-down is provisional and arms a probe whose verdict is accepted (frame rate at least 1.05× the previous one), rejected (revert and learn a floor) or inconclusive (keep the DPR, learn nothing); two punished ascents above DPR 1.0, or sustained distress, demote the ceiling to 1.0. Floor and ceiling both decay on a doubling ladder capped at 5 min. (b) Sketch of the oscillation the probe prevents on a scene whose frame rate does not respond to DPR: without it the DPR walks down and back up repeatedly; with it one probe fails, the DPR reverts and the learned floor holds it there for 30 s before a re-probe.

5 min cap; a probe that sees no clean frame-rate sample inside its window is voided as inconclusive, so the DPR is kept and nothing is learned. Symmetrically, repeatedly punished attempts to exceed DPR 1.0 demote the scale-up ceiling to 1.0 on the same expiring ladder (60 s base, doubling, capped at 5 min), after which the ceiling is tried again; the same single demotion to 1.0 also fires when the refresh-rate estimator reports sustained distress, a frame rate too low to be any display throttle, while the DPR is above 1.0. Load activity is excluded from the FPS evidence, so transient loading jank never becomes a learned floor or ceiling.

The DPR is clamped to [`minDPR` (0.5), min(native DPR, permitted ceiling)], where the permitted ceiling follows the user's high-DPR policy (the native DPR when high DPR is allowed on a desktop, CSS resolution 1.0 otherwise, further lowered by a live demotion while interacting). On idle the controller rests at the policy ceiling for a sharp resting image (not necessarily at the native DPR) and re-probes after a content change. During recording the DPR is **locked**, so that the captured video has no mid-clip resolution changes.

### 9 Animation and recording

#### 9.1 Per-dimension animation

Each non-displayed dimension supports an independent animation state consisting of `isPlaying`, a `targetFPS` drawn from the preset $\{0.5, 1, 2, 5, 10, 15, 30, 60, 120\}$ Hz or a custom rate, a `loopMode` from `{once, loop, bounce}`, and a `direction` of `forward` or `backward`. The animation controller maintains a separate state per dimension and advances them in parallel; there is no global play-head.

For a discrete dimension, each step advances by the dimension's declared step size (typically 1). For a continuous dimension, the controller computes a per-frame increment as $\delta = (\max - \min)/(T_{\text{traverse}} \cdot \text{FPS})$, where $T_{\text{traverse}}$ is the configured traverse time, 10 s by default (the full range in ten seconds). A per-dimension step override (an explicit step value set from the animation menu, offered as multiples $\{0.1, 0.25, 0.5, 1, 2, 5\}$ of the base step) replaces both rules when set; on a discrete dimension it is quantised to the authored grid, one cell at minimum, and it never exceeds the dimension's range width. The default FPS is 10, and custom rates span 0.1–120 Hz.

The three loop modes have natural semantics. `once` stops at the boundary and emits a `complete` event. `loop` wraps around. `bounce` reverses direction at the boundary and emits a `directionChange` event so that any UI bound to the animation knows about the reversal.

#### 9.2 Frame-data synchronisation

A naive animation loop would advance the slider on every frame regardless of whether the data for the new slice has finished loading. On a slow link, this produces a jittery experience where some frames are blank because their chunks have not arrived yet. Luxar's animation loop explicitly gates the advance: a frame is marked `pending` in a Set when its data load starts, and the loop skips any further advance on this dimension until the pending frame clears. If the load is consistently slower than the target FPS, the loop's measured FPS drifts below target. The controller measures the achieved FPS once per second and, whenever that measurement is below 80% of the target (or the frame on screen is still filling in: the committed additive-ladder energy of that frame, the $e(k)$ of its committed prefix as stamped by the ladder's *energy stamps*, is still below the release threshold of the *display gate*), emits a `fpsWarning` event carrying the target FPS, the actual FPS and the committed energy fraction, which the UI can display as a warning indicator. A measurement at or above that fraction with a fully committed frame emits nothing.

#### 9.3 Recording panel

The recording panel composes the per-dimension animation system with a capture pipeline. Three top-level modes are supported:

- **Image mode**: a still snapshot, encoded as PNG, WebP, JPEG, or EXR (HDR; see §9.4).
- **Video mode**: real-time capture during interactive playback, encoded by `MediaRecorder` as a WebM container with VP8 or VP9. The bitrate is chosen as width $\cdot$ height $\cdot$ FPS $\cdot$ bpp, with bpp presets $\{0.04, 0.08, 0.15, 0.30\}$ for low / medium / high / max quality (yielding roughly $\{2.5, 5, 9, 19\}$ Mbps at 1080p 30).
- **Turntable mode**: a 360-degree camera orbit at a fixed slice, captured offline frame-by-frame through the `mediabunny` WebCodecs library from 8-bit canvas frames. The selectable codecs are H.264, H.265, VP9 and VP8; when the platform cannot encode the chosen codec at the requested size and bitrate, the encoder walks a fixed fallback list (VP9, AV1, VP8, H.264; H.264 first after a failed H.265) and can therefore deliver AV1 without the user having selected it. A WebM container restricts the choice to VP9, AV1 and VP8.

In Video mode, the controller can pair the capture with a slider sync: as the video records, a chosen dimension's slider advances in real time, producing a movie of the dataset, not a movie of a static view (Turntable mode records at a fixed slice, and Image mode has no slider sequence). Frame-sequence ZIPs are also supported (a directory of PNG / WebP / JPEG / EXR frames bundled by `fflate`). EXR sequences ship with a generated `ffmpeg` encode script. Its active command encodes an SDR H.265 master from the frames; its commented-out block, once the user enables it and chooses the nominal peak luminance (`npl`) that the scene-linear values should map to, converts the frames to 10-bit BT.2020/PQ HDR10 video. That is the only path to 10-bit output, since the in-browser encoders work from 8-bit canvas frames.

#### 9.4 HDR EXR capture

A standard PNG screenshot captures the display output: tone-mapped, sRGB-encoded and quantised to 8 bits. For quantitative HDR work this is lossy: the linear radiometry cannot be recovered from it, because the tone curve compresses the HDR range (Reinhard, for instance, sends 2 to 2/3 and 4 to 4/5) and whether values clip depends on the operator and the exposure chosen. The EXR exporter instead samples the linear HalfFloat pipeline *before* exposure, offset, gamma, tone mapping and display encoding and writes 16-bit float RGBA to a single-frame EXR. Its default mode (`hdr-effects-pre-tone`) runs the pipeline up to the fused pass and returns the HDR scene sample with bloom added when bloom is enabled, with lens distortion and detector noise switched off for the capture; a second mode (`raw-scene-hdr`) reads the scene render target alone, without bloom; a third (`visible-ldr`) returns the tone-mapped output before sRGB encoding (Fig. 7, bottom row). In every mode the alpha channel is written as 1 (additive blending leaves the target's alpha as a meaningless overdraw count). The dynamic range of the RGB channels is that of the renderer, so tone mapping becomes a post-processing decision the user can make in their image-editing tool of choice, not a destructive step at capture time; for quantitative use the capture mode and the enabled effects should be stated, and bloom disabled or the raw-scene mode chosen.

During a capture, adaptive DPR is pinned and the canvas resize handler is locked (so the capture resolution matches the user's intent); offline sequences (turntables and frame sequences) additionally register a `continuous: true` keep-alive callback with the animation controller so that the loop never idles between frames. After capture, all of these are restored.

#### 10 Conclusion

Every subsystem in this document rests on one decision taken in the scene graph: the on-disk tree is the runtime tree, and everything the viewer mutates (transforms, appearance, overlay visibility) lives in attributes or in the query, beside chunk data that is never rewritten. That decision is why a spatial index never needs rebuilding, why a Layers-panel edit or a group-level transform costs no data traffic, why a hover overlay can be injected at compile time without touching the geometry, and why a standalone splat file can be grafted into a scene without rewriting either.

The subsystems are also decoupled from one another so that each is cheap when idle: the picking buffer is reduced-resolution and re-rendered only on dirty events, the overlay layer is HTML and costs nothing per frame when nothing changes, and the animation controller is idle when no dimension is playing. Rendering-performance measurements are in Supplementary Document 6, which benchmarks the render pipeline itself; the combined cost of picking, overlays and animation on top of it has not been measured separately.
