## Supplementary material for "Luxar: Gaussian splatting for microscopy and scalable interactive web visualisation of multidimensional scientific data": supp_doc_11_tiled_fitting

### Tiled Fitting for Large Volumes

Supplementary Document 11 – Luxar

**Abstract**

A volume larger than GPU memory is fitted as overlapping tiles. Luxar lays a regular grid over the volume, multiplies each tile by a half-Hann taper, fits Gaussian splats to every tile independently and unions the splat sets. Adjacent tapers sum to one and splat rendering is linear, so in the ideal case the union reconstructs the volume without a seam. Whether real fits leave seams is measured on two volumes small enough to fit whole, a confocal blastocyst and a light-sheet *Tribolium* embryo. With the taper and an overlap of 5 to 20% of the tile size, the tiled union stays within −0.98 to +0.19 dB of the monolithic fit at equal seed budget, over grids of 4 to 100 tiles. The residual next to a seam is at most 13% above the residual deep inside a tile, relative to the fit without seams. A hard cut without overlap leaves a visible seam, and overlap without the taper is worse than a hard cut, so the partition of unity is what removes the seam. More overlap buys nothing at this budget. Tiling is a memory device: the working set is proportional to the voxels fitted at once, 43 to 46 MB per million, and tiles run in parallel with no communication. The document derives the taper, proves the partition of unity, records the grid and seed conventions and closes with rules: overlap near 10% of the tile size, seeds budgeted by content, memory sized for the widest folded tile.

**8 Summary and practical rules** **23**

### 1 Introduction

Luxar fits a volume with a Gaussian mixture. Splat $j$ has a centre $\boldsymbol{\mu}_j \in \mathbb{R}^D$, an amplitude $a_j$ and a lower-triangular Cholesky factor $\boldsymbol{\Lambda}_j$ of its covariance $\boldsymbol{\Sigma}_j = \boldsymbol{\Lambda}_j \boldsymbol{\Lambda}_j^\top$ (written $\boldsymbol{\Lambda}$ because $L$ is the overlap width throughout this document), and the reconstructed volume at voxel $\mathbf{x}$ is the sum

$$\hat{V}(\mathbf{x}) \;=\; \sum_j a_j \exp\Big(-\tfrac{1}{2}\|\boldsymbol{\Lambda}_j^{-1}(\mathbf{x}-\boldsymbol{\mu}_j)\|^2\Big)\,. \tag{1}$$

A $D$-dimensional splat therefore carries $F_D = D + D(D{+}1)/2 + 1$ parameters, $F_D = 10$ in 3D. The question of this document is how to fit that model to a volume that does not fit on the GPU, and what the answer costs in reconstruction quality at the places where the volume is cut.

The fitting cost of $K$ splats on a volume of $N$ voxels is dominated by two memory terms. The first is the volume itself, $4N$ bytes at float32. The second is the per-splat parameter, gradient and Adam optimiser state, about $4 \cdot K \cdot F_D \cdot 4$ bytes (parameter, gradient and the first and second moments, each a float32 per parameter). On an NVIDIA RTX 3090 (24 GB), a $2048^3$ float32 volume already requires 34 GB, exceeding working memory at the input alone, before any splat parameter is allocated. The input is not the binding term at smaller sizes. The fitting working set measured in Sec. 7.6 is 43–46 MB per million voxels, so a light-sheet stack of 2–4 giga-voxels, whose float32 input of 8–16 GB would still fit a 24 GB card, extrapolates to 86–185 GB to fit whole (from probes of at most a tenth of a giga-voxel, Sec. 7.6). A partitioning strategy is mandatory there. For smaller volumes, partitioning is also useful as a way to expose embarrassingly parallel work for cluster and multi-GPU scaling.

Three properties make the strategy a credible replacement for monolithic fitting. First, reconstruction quality at tile boundaries should not degrade visibly relative to the interior of a tile. Second, the tile geometry should be deterministic and resumable, so that batch systems such as Slurm can plan ahead and resume. Third, the post-fit composition of tile outputs should be a pure data operation (concatenation), not a second optimisation pass.

Luxar's answer is a *uniform* tiling with a fixed grid independent of content, which satisfies the second property by construction. Each tile's voxel data is multiplied by a raised-cosine (half-Hann) taper before fitting, the tiles are fitted independently and their splat sets are unioned. Two facts make the union a valid composition: adjacent tapers sum to one in the overlap (the partition of unity, Sec. 3) and Eq. 1 is linear in the splat set (Sec. 4). Both hold on the data the tiles actually see, so any background floor must be removed *before* windowing. Luxar resolves the floor once against the whole volume and subtracts it from each raw tile; a per-tile floor would differ between neighbours and reintroduce exactly the boundary intensity step the taper exists to prevent.

Luxar also ships a *content-adaptive* tiling (`gsplat fit --tiling content`). It partitions the volume with a content-balanced *BSP* over a coarse feature-density field and assigns each box a splat budget from the calibrated density relation $K \propto n_{\text{features}}^{\alpha}$, where $\alpha$ is the saturation exponent of Sec. 5.2 ($\alpha = 0.44$ by default). The second property still holds: the plan is computed once and persisted as a `FitPlan` JSON that every worker re-reads, so the geometry remains deterministic and resumable even though it is content-dependent. The content-adaptive fitter does not use the apodisation below, however. It fits each box on a halo-padded, unwindowed crop of the raw volume and keeps only the splats whose centres fall inside the box's half-open core (`gsplats/planner/fit_planned.py`), so adjacent boxes neither share retained splats nor blend. The window derivation, the partition-of-unity proof and the validation of Sec. 7 therefore concern uniform tiling only; the seam behaviour of content tiling has not been measured here.

The document proceeds from theory to measurement. Secs. 2 to 4 derive the tile grid, the taper, the partition of unity and the linearity argument. Secs. 5 and 6 record the conventions of the shipped implementation, including the rule that divides a seed budget over the tiles. Sec. 7 measures what the theory leaves open: whether the residual fitting error, which the taper cannot cancel, concentrates at the seams, and what tiling costs in quality, time and memory. Sec. 8 collects the practical rules.

#### Terms used in this document

**PSNR** The peak signal-to-noise ratio, $10\log_{10}(\text{peak}^2/\text{MSE})$ in decibels, where peak is the reference range of the values compared; unless a document states another range, the supplements take peak 1 on volumes normalised to $[0, 1]$, so that $\text{PSNR} = -10\log_{10}\text{MSE}$.

**arm** One experimental condition of a comparison (for instance the blind-spot arm and the unmasked arm of a codec study), run with every other setting held equal.

**BSP** Binary space partitioning, the recursive splitting of a set of elements by an axis-aligned plane (at the median by default) into two halves until every part falls below a size cap.

**cull (retention)** The removal of splats after a fit. The fit-time cull ranks splats by amplitude and keeps the top set accounting for a retention fraction of the total amplitude (0.95 by default); the contribution-based cull instead removes the splats whose contribution to the rendered field falls below an error budget or a fractional threshold, and is the baseline that removal-based level-of-detail methods use.

**relocation** The fixed-pool alternative to adding and removing splats during optimisation: the splat count is fixed at seeding, and the least informative splats are periodically moved to regions of high reconstruction residual, so the optimiser tensor shapes never change.

**blind-spot cross-validation** A model-selection protocol in which a deterministic random 5% of voxels is hidden from the fit and the reconstruction is scored on those voxels afterwards, so that capacity spent on memorising noise shows up as a loss instead of a gain.

**leaf** A node that carries splat arrays directly (centres, amplitudes, Cholesky factors) together with an additive ladder of at least one sub-LOD, as opposed to a group node that only holds children.

**`kind=partition` group** A group whose `part_<i>` children each carry their own position bounds and are all drawn at once, each culled against the camera frustum; the parts are usually the regions of a recursive

BSP split of one splat set, but any disjoint grouping is valid, such as one part per timepoint.

**part** One child of a `kind=partition` group (`part_<i>`), holding one disjoint subset of the splats (a BSP region, or a timepoint) together with its own position bounds for frustum culling.

**additive (streaming) ladder** One ordering of a leaf's splats cut into prefix-sum sub-LODs, so that every prefix of the download is a valid, progressively refining rendering of the same splat set.

**recipe** A named LOD topology built from a fitted splat set: `flat` (one bare leaf), `stream` (one stream-laddered leaf), `levels` (coarse-to-fine replacement levels), `tiles` (a BSP partition with one ladder per tile), `overview` (one coarse level above a `tiles` branch) and `adaptive` (`tiles` where every tile carries its own `levels`).

#### 2 Tile geometry

The volume has shape $\mathbf{S} = (S_1, \ldots, S_D)$, so $S_d$ is the length of axis $d$ in voxels. Each axis is partitioned into overlapping segments of nominal size $T_d$ with stride $T_d - L_d$, where $L_d$ is the overlap width. The enumeration (`compute_tile_specs` in `gsplats/tiling.py`) proceeds in two steps. First, candidate $i$ on axis $d$ covers the half-open index interval $[\, i(T_d - L_d),\ \min(i(T_d - L_d) + T_d,\ S_d)\,)$ for $i = 0, 1, \ldots$ as long as the start lies inside the axis. This gives $c_d = \lceil S_d/(T_d - L_d) \rceil$ candidates, the last of them clamped to the volume boundary; write $\mathrm{end}_i$ for the end of candidate $i$. Second, the *fold rule*. The last candidate's unique coverage is the part of the axis that only it covers, $S_d - \mathrm{end}_{c_d - 2}$. When that is *smaller than the overlap* $L_d$, the candidate is dropped and its predecessor is extended to $S_d$. That predecessor then spans up to $T_d + L_d - 1$ voxels, the one case in which a tile is larger than $T_d$. The number of tiles per axis is therefore $n_d = c_d$, or $c_d - 1$ when the fold applies. With $L_d = 0$ nothing folds, because no unique coverage is smaller than zero.

Two examples show the rule (Fig. 1a). For an axis of $S_d = 10$ voxels with $T_d = 8$ and $L_d = 4$ the candidates are $[0, 8)$, $[4, 10)$ and $[8, 10)$; the third has no voxel of its own ($10 - 10 = 0 < 4$), so it is folded and the axis holds the two tiles $[0, 8)$ and $[4, 10)$. The widest tile arises when the dropped candidate had $L_d - 1$ voxels of its own. For $S_d = 15$, $T_d = 8$, $L_d = 2$ (stride 6) the candidates are $[0, 8)$, $[6, 14)$ and $[12, 15)$; the third has one unique voxel ($15 - 14 = 1 < 2$), so the fold gives $[0, 8)$ and $[6, 15)$, a tile of $9 = T_d + L_d - 1$ voxels. Sec. 7.2 lists the grids this rule produces for the volumes of the sweep.

For each tile we record the *actual* per-face overlap sizes $(\ell_d^-, \ell_d^+)$ with the previous and next tile along axis $d$: the previous tile's end minus this tile's start, and this tile's end minus the next tile's start. Two boolean flags $\beta_d^-, \beta_d^+$ mark the *first and last tile on the grid axis* ($i = 0$ and $i = n_d - 1$). After the fold, the last tile on an axis is the only one that ends at $S_d$: a predecessor ending at $S_d$ would leave its successor no unique voxel, and the fold removes that successor. First and last tiles have zero overlap on their outer face and carry no ramp there; every interior face carries a ramp over its actual overlap. These per-tile descriptors are the input to the window construction in Sec. 3. Collecting them into a `TileSpec` object makes each tile fully self-describing, which matters for Slurm array jobs that must run independently with no shared state.

We require $L_d \leq T_d/2$ on every axis. On this grid the constraint guarantees that any voxel is contained in at most two tiles along each axis, hence in at most $2^D$ tiles overall: the tensor-product neighbours meeting at a face, edge or corner. With $L_d > T_d/2$ and at least three tiles on the axis,

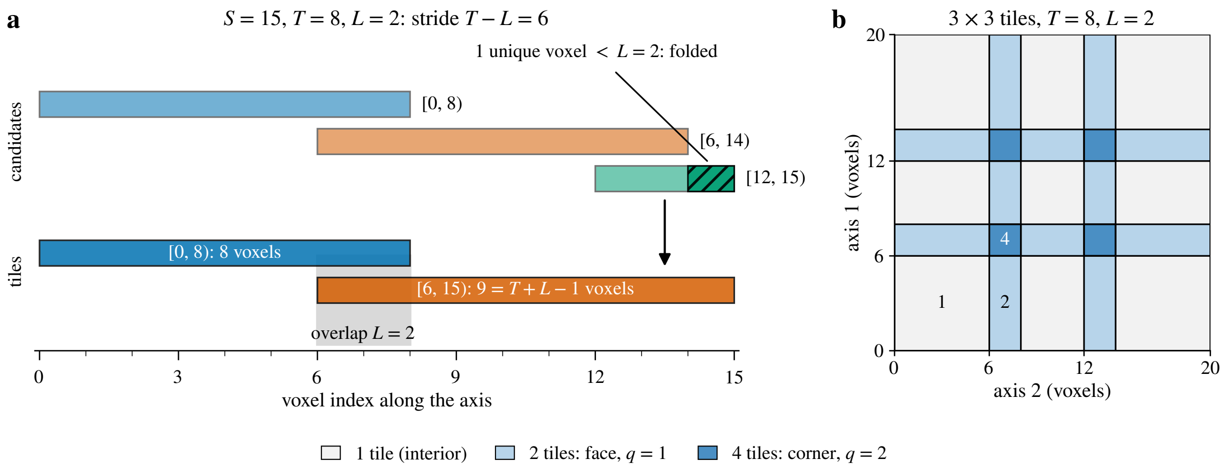

Figure 1: **Tile enumeration and the fold rule (schematic).** (a) The second worked example, an axis of $S = 15$ voxels with $T = 8$ and $L = 2$. The three candidate tiles start at multiples of the stride $T - L = 6$; the last one owns a single voxel (hatched), fewer than the overlap, so it is folded into its predecessor, which then spans $9 = T + L - 1$ voxels. The grey band is the overlap the two remaining tiles share. (b) A grid of $3 \times 3$ tiles with the same $T$ and $L$, coloured by the number of tiles that contain each voxel: one in a tile interior, two across a face and four at a corner crossing, $2^q$ for $q$ axes in overlap (Eq. 4); in three dimensions an edge holds four tiles and a corner eight.

consecutive tiles share samples three at a time and the two-ramp identity derived below no longer applies; the implementation rejects such configurations with a `ValueError`.

### 3 Cosine apodisation and partition-of-unity

#### 3.1 The 1D ramp

For an overlap of width $L$ samples, the rising half-cosine ramp is

$$r_L(k) \;=\; \frac{1}{2}\left(1 - \cos\frac{\pi k}{L+1}\right), \qquad k = 1, \ldots, L, \tag{2}$$

where $k$ is the overlap-local sample index. The denominator $L + 1$ (not $L$) ensures $r_L(k) \in (0, 1)$ strictly, avoiding the exact zeros that would erase boundary voxels and the exact ones that would defeat the apodisation at the opposite end of the ramp. The falling ramp is the time-reverse, $\bar{r}_L(k) = r_L(L + 1 - k)$.

#### 3.2 Two-tile partition of unity

The partition-of-unity claim is that two overlapping windows from adjacent tiles sum to 1 at every sample within the overlap zone. The two tiles meeting at a face record the same actual overlap for it: the left tile's $\ell_d^+$ and the right tile's $\ell_d^-$ are both the left tile's end minus the right tile's start (Sec. 2). Both ramps therefore have the same length $L = \ell_d^+ = \ell_d^-$ and are indexed by the same overlap-local coordinate $k \in \{1, \ldots, L\}$. For a rising ramp $r_L$ on the right tile and a falling

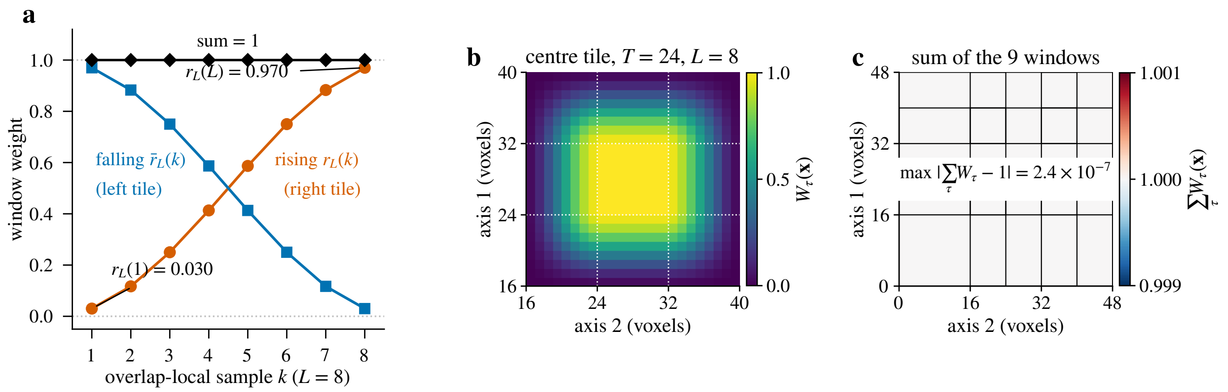

Figure 2: **The raised-cosine ramp and the partition of unity (schematic).** (a) The rising ramp $r_L(k)$ of the right tile, the falling ramp $\bar{r}_L(k)$ of the left tile and their sum over an overlap of $L = 8$ samples (Eq. 2); the denominator $L + 1$ keeps the end values at 0.030 and 0.970, never 0 or 1, and the sum is 1 at every sample (Eq. 3). (b) The separable window $W_\tau$ of the centre tile of a $3 \times 3$ grid with $T = 24$ and $L = 8$; the dotted lines mark where its ramps begin. (c) The sum of the nine windows of that grid over the whole domain, computed in single precision as the fitter does: it is 1 everywhere, faces, edges and corners included, to within $2.4 \times 10^{-7}$, two float32 units in the last place at 1 (Eq. 4).

ramp $\bar{r}_L$ on the left tile,

$$
\begin{aligned}
r_L(k) + \bar{r}_L(k) &= \tfrac{1}{2}\Big(1 - \cos\tfrac{\pi k}{L+1}\Big) + \tfrac{1}{2}\Big(1 - \cos\tfrac{\pi(L+1-k)}{L+1}\Big) \\
&= 1 - \tfrac{1}{2}\Big(\cos\tfrac{\pi k}{L+1} + \cos\Big(\pi - \tfrac{\pi k}{L+1}\Big)\Big) \\
&= 1 - \tfrac{1}{2}\Big(\cos\tfrac{\pi k}{L+1} - \cos\tfrac{\pi k}{L+1}\Big) \;=\; 1.
\end{aligned}
\tag{3}
$$

The identity $\cos(\pi - \theta) = -\cos\theta$ provides the cancellation. Consequently, in the overlap zone, the two adjacent windows sum exactly to 1 at every voxel; outside the overlap, one window is identically 1 and the other is undefined (the tile does not extend there).

The full window per tile is the separable product $W(\mathbf{x}) = \prod_{d=1}^{D} w_d(x_d)$, where each axial window $w_d$ is 1 in the interior, rising over $\ell_d^-$ samples on its low face if $\beta_d^- =$ False and falling over $\ell_d^+$ samples on its high face if $\beta_d^+ =$ False. Because the window is a product, the sum over the tiles that contain a voxel $\mathbf{x}$ factorises,

$$
\sum_{\mathbf{i}} \prod_{d=1}^{D} w_{d,i_d}(x_d) \;=\; \prod_{d=1}^{D} \sum_{i_d} w_{d,i_d}(x_d) \;=\; 1, \tag{4}
$$

where $\mathbf{i}$ runs over the $2^q$ tensor-product neighbours that meet at a $q$-fold face, edge or corner crossing ($q$ of the $D$ axes in overlap) and each axial sum has either one term (equal to 1) or the two terms of Eq. 3. Adjacent tiles therefore sum to 1 on the entire $D$-dimensional overlap manifold, edges and corners included, provided every one of those neighbours is present in the union (Fig. 2).

##### 3.3 Why the constraint $L \leq T/2$

If $L > T/2$ and the axis holds at least three tiles, consecutive tiles A, B, C share a non-empty intersection (the right overlap of A intersects the left overlap of C through B). In that triple-overlap region the window sum becomes $w_A + w_B + w_C$, which is generically not 1 with the two-ramp cosine windows. A partition of unity over triple coverage needs a different construction: a window family with wider support (quadratic B-spline bases, for instance, sum to one by construction), or simply any positive windows normalised by their pointwise sum. The Luxar implementation does not pursue either and instead enforces the simpler two-window guarantee.

#### 4 Why splat sets compose by union

The Gaussian-splat reconstruction operator of Eq. 1 is linear in the splat set. Write $\hat{V}_{\mathcal{G}}$ for the volume rendered from a set $\mathcal{G}$ of splats. If $\mathcal{G}_1$ and $\mathcal{G}_2$ are two collections of splats, the rendered volume from their disjoint union is the sum of the individually rendered volumes,

$$\hat{V}_{\mathcal{G}_1 \cup \mathcal{G}_2}(\mathbf{x}) \;=\; \hat{V}_{\mathcal{G}_1}(\mathbf{x}) + \hat{V}_{\mathcal{G}_2}(\mathbf{x}), \tag{5}$$

because $\hat{V}_{\mathcal{G}}(\mathbf{x}) = \sum_{j \in \mathcal{G}} a_j \exp(-\frac{1}{2}\|\mathbf{\Lambda}_j^{-1}(\mathbf{x} - \boldsymbol{\mu}_j)\|^2)$ is a sum and sums distribute over disjoint partitions of the index set. Combined with the partition of unity, this gives the per-tile windowing algorithm its correctness. Let $\tau$ label a tile of the grid and $W_\tau$ its window, extended by zero outside the tile.

1. Each tile $\tau$ multiplies the volume by its window, $V_\tau(\mathbf{x}) = V(\mathbf{x}) \cdot W_\tau(\mathbf{x})$.
2. Tile $\tau$ fits a splat set $\mathcal{G}_\tau$ to $V_\tau$ on the tile's own domain, so $\hat{V}_{\mathcal{G}_\tau}(\mathbf{x}) \approx V(\mathbf{x}) \cdot W_\tau(\mathbf{x})$ for $\mathbf{x}$ inside the tile.
3. The merged splat set $\mathcal{G} = \bigcup_\tau \mathcal{G}_\tau$ renders to

   $$\hat{V}_{\mathcal{G}}(\mathbf{x}) = \sum_\tau \hat{V}_{\mathcal{G}_\tau}(\mathbf{x}) \;\approx\; V(\mathbf{x}) \sum_\tau W_\tau(\mathbf{x}) \;=\; V(\mathbf{x}) \cdot 1 \;=\; V(\mathbf{x}),$$

   using Eq. 4 for the unit sum.

The error of step 3 has two parts. Inside tile $\tau$ it is the fit's residual against $V \cdot W_\tau$, the quantity the optimiser minimised. Outside the tile the same splats still render against a target that is zero there. A Gaussian is not clipped at the tile face: the renderer truncates it at $2.75\sigma$, with $\sigma$ the splat's standard deviation along the direction in question, and `fit_tile` merely translates the centres into global coordinates. A tile fit can therefore match its windowed target exactly on its own domain and still leak into its neighbours. Two unit-amplitude isotropic Gaussians of $\sigma = 1$ voxel, fitted to two unit samples one voxel apart on disjoint tiles, each reproduce their own sample exactly, yet their union renders $1 + e^{-1/2} = 1.61$ at both (Fig. 3a).

Exact reconstruction requires each $\mathcal{G}_\tau$ to render to $V \cdot W_\tau$ over the whole volume, zero extension included. The taper acts on exactly this term: near a face the target already falls to zero, so the

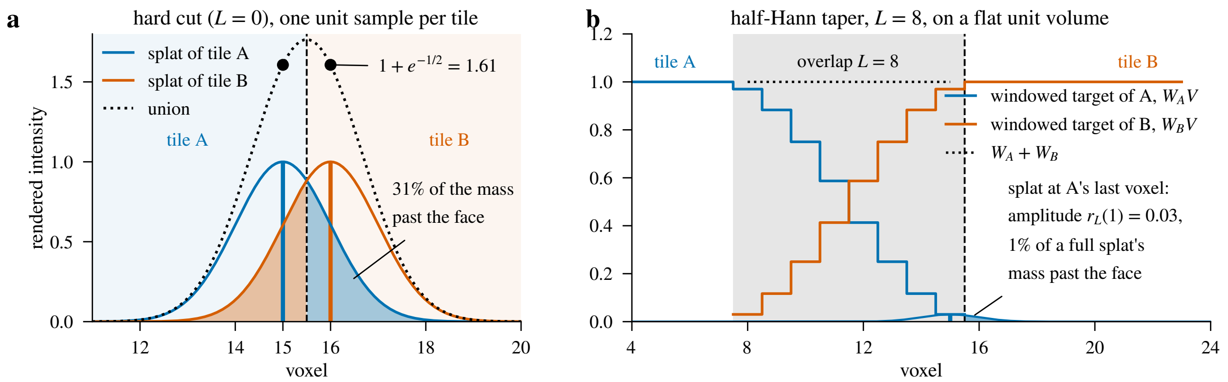

Figure 3: **Leakage across a tile face (schematic).** (a) A hard cut. Tile A holds a unit sample at its last voxel and tile B one at its first; each tile fits a unit-amplitude Gaussian of $\sigma = 1$ voxel that reproduces its own sample exactly. Neither Gaussian stops at the face (dashed): each carries 31% of its mass across it (shaded), and the union (dotted) renders $1 + e^{-1/2} = 1.61$ at both samples. (b) The same face under the half-Hann taper with $L = 8$, on a flat unit volume. The windowed targets of the two tiles fall and rise across the overlap (grey) and sum to 1 (dotted); the splat fitted at the last voxel of tile A now carries the window value $r_L(1) = 0.03$ as its amplitude, so what it leaks past the face is 1% of a full splat's mass.

splats fitted there carry little mass and leak little (Fig. 3b). Whether the two error terms are small in practice, whether they are uncorrelated across tiles, and how they scale with capacity are empirical questions, answered for the uniform tiling in Sec. 7; the hard-cut *arm* there ($L = 0$) shows what a Gaussian fitted to a target truncated at a plane does on the other side of it. There is no second optimisation pass after merging. Composition is a pure data union, with an optional cumulative-amplitude *cull (retention)* whose scope depends on the output shape (Sec. 5).

### 5 Implementation conventions

The Luxar implementation (`packages/luxar/src/luxar/gsplats/fit_tiled_gsplats.py`) exposes `fit_tile` (atomic, Slurm-ready) and `fit_tiled` (orchestrator). Every number in this document, the sweep of Sec. 7, its seed-allocation probe and the per-tile memory probe, was measured with one luxar build, recorded in every results record. The commit this document pins for the code it cites, `a5a18ca06`, is not that build. Between the two, `gsplats/tiling.py`, `gsplats/seeds/` and `gsplats/fitting/` are identical. `gsplats/fit_tiled_gsplats.py` and the CLI allocator differ by docstrings, the extraction of a helper that computes the same windowed mass, and the ceiling argument the allocator passes when a floor is applied (none was applied here); the diff of those two files between the two commits is deposited with the analysis. A re-run of the allocator on the volumes reproduces every recorded per-tile seed count at the build the sweep ran on; it was not repeated at the pinned commit. The batch runs behind Sec. 7.4 are described in Sec. 7.1.

Defaults are tile size 256 and overlap 32 on each axis, which yields a stride of 224 voxels and a tile-local volume of $256^3 = 16.8$ M voxels per tile, well within consumer-GPU working memory together with $\sim 10^4$ splats, their gradients and Adam state. For a $2048^3$ volume these parameters enumerate $\lceil 2048/224 \rceil = 10$ candidates per axis, the last starting at 2016. Since $2048 \bmod 224 = 32 = L$, the ninth candidate already ends at 2048 and the tenth has no voxel of its own, so the fold rule of Sec. 2 removes it. The grid is $9^3 = 729$ tiles, every one with voxels of its own, the last on each axis $[1792, 2048)$ and 256 voxels wide. All are fittable in parallel.

##### 5.1 Merge paths

Every tile fit disables its own post-fit cull (`fit_tile` keeps every fitted splat); what happens at the merge depends on the output shape, and three paths exist:

- **Flat union** (Python `fit_tiled(partition=False)`, or `--flat` on the CLI): the tiles are concatenated into one *leaf* and a single cumulative-amplitude cull is applied to the merged result at `cull_retention` (0.95 when nothing sets it, 0.999 under every `--preset`; 0 keeps all), so the cut is consistent across tile boundaries. A progressive fit (the multi-level fit of Supplementary Document 3) merges level by level: the coarsest level of detail (LOD 0) of every tile forms the global coarsest level, and so on. This is the path the validation of Sec. 7 uses.
- **Partition** (the CLI default for a tiled fit, sequential or `-j`): the tiles are kept as a ***`kind=partition`*** *group* with one *part* per tile, for viewer frustum culling. Each part is culled independently at the same retention (a global cut has no meaning once the parts stay separate) and each part keeps its own *additive (streaming) ladder*.
- **Batch** (`batch-fit`, Sec. 6): the workers keep every splat and the streaming merge applies no cull, so batch output retains every fitted splat.

The seed budget reaches the tiles through the same orchestrator. `fit_tiled` takes either one `seeds` value applied to every tile or an exact per-tile list, `tile_seed_counts`; the CLI resolves the latter from an integer `--seeds` budget (Sec. 5.2) and hands it down, and a tile whose count is 0 is skipped without fitting.

##### 5.2 Seed allocation

The CLI resolves an integer budget $K$ into one seed count per tile from the tile's windowed content. Let $\tau$ be a tile with window $W_\tau$, let $f$ be the background floor resolved once for the whole volume ($f = 0$ in the sweep, where no floor was applied) and let $v_{\max}$ be the shared intensity ceiling, the normalisation top the orchestrator resolves once for every tile of a volume. Two sums describe the tile,

$$n_{\mathrm{Hann},\tau} \;=\; \sum_{\mathbf{x}\in\tau} W_\tau(\mathbf{x}), \qquad m_\tau \;=\; \sum_{\mathbf{x}\in\tau} W_\tau(\mathbf{x})\,\max\big(V(\mathbf{x}) - f,\, 0\big), \tag{6}$$

its Hann-weighted voxel count and its windowed intensity mass above the floor. The tile's weight is

$$w_\tau \;=\; n_{\mathrm{Hann},\tau}\left(\frac{m_\tau}{n_{\mathrm{Hann},\tau}\, v_{\max}}\right)^{\alpha}, \tag{7}$$

the Hann-weighted size times the mean windowed intensity as a fraction of the ceiling, raised to the *saturation exponent* $\alpha = 0.44$ (`--saturation-exponent`). Equal content gets equal splat density in tiles of different sizes; between equal tiles the budgets stand in the ratio of their masses to the power $\alpha$. A tile with $m_\tau = 0$ has weight 0, is budgeted 0 and is not fitted. Let $M_+$ be the number of tiles with mass and $\bar{m} = \sum_\tau m_\tau / M_+$ the equal mass share. The provisional count of tile $\tau$ is $K_\tau = K\, w_\tau / \sum_{\tau'} w_{\tau'}$, subject to the floor

$$m_\tau \;\geq\; 0.25\,\bar{m} \quad\Longrightarrow\quad K_\tau \;\geq\; 0.25\,\frac{K}{M_+}, \tag{8}$$

so a tile holding at least a quarter of the equal mass share is never budgeted below a quarter of the equal seed share. Such a tile is pinned at that share and the rest of the budget is split proportionally among the others, repeated until no eligible tile falls below its floor. The weights are turned into integers summing to $K$ by deterministic largest-remainder rounding, every tile of positive weight receiving at least 1; a single-tile grid takes the whole budget. The CLI prints the resolved counts. The same exponent drives the content planner's $K \propto n_{\text{features}}^{\alpha}$ (Sec. 1).

##### 5.3 CLI reference

The uniform tiling of this document is driven by `luxar gsplat fit`. Table 1 lists the flags that matter for reproducing the workflow and the points where the CLI and the Python API differ.

#### 6 Slurm fan-in

Tile-level independence makes Slurm array jobs (one job script launched as many numbered tasks) a natural fit. `compute_tile_specs` produces a deterministic enumeration of tiles (row-major over the grid index), and the Luxar batch CLI (`luxar gsplat batch-fit submit`) plans one fitting task per $(t, c,$tile$)$ combination for OME-Zarr 5D inputs and submits them as a Slurm array: one array element per task by default, or `--tasks-per-job N` consecutive tasks per element when single tasks would not saturate a GPU. Its local sibling, `batch-fit run`, fans the same plan out over the GPUs of one machine. Each task is a `luxar gsplat fit --tile k/M` worker whose output is a small `.gsplats.zarr` archive; a single `luxar gsplat batch-fit merge` pass streams the per-tile splat sets into the final output.

With more than one spatial tile the merge writes a `kind=partition` tree with one part per spatial tile, each part holding that tile across all selected timepoints and channels. With a single spatial tile, a recording whose frames are smaller than the tile size, it writes one stacked leaf (or one LOD group under `--recipe`) with no partition wrapper. The merge holds one spatial region at a time, so its peak memory is one tile's splats over all timepoints, and it applies no cull (Sec. 5.1).

The input side is tile-local only on the batch path. A `batch-fit` plan whose floor and normalisation range are resolved at plan time, with no downscaling or denoising, hands every worker its exact tile region, which `load_volume` reads through the lazy zarr slice, so the workers of such a uniform plan request about $(T/(T-L))^D \approx 1.5$ frames of input in aggregate for the default $T$=256, $L$=32 in 3D. A worker spawned by the `-j` parent, or run by hand as `--tile k/M`, instead calls `load_volume` without a region: it selects the requested timepoint and channel lazily

Table 1: **Tiling flags of `luxar gsplat fit`.** Defaults in parentheses. The Python orchestrator `fit_tiled` takes the same tile size and overlap, and either a *per-tile* `seeds` value or the exact per-tile list `tile_seed_counts` the CLI resolves.

| Flag | Meaning |
|---|---|
| `--tiling` (`auto`) | `none` \| `uniform` \| `content` \| `auto`. `auto` fits the whole volume unless some dimension exceeds the tile size *and* the volume exceeds 64 M voxels; when it does tile, it picks `content` if a density (`--cal`, `--k-star-ref`, `--plan`) is given, else `uniform`. The 17.6 M-voxel blastocyst of Sec. 7 is therefore not tiled under `auto` although two of its axes exceed 256; pass `--tiling uniform` explicitly. |
| `--tile-size` (256), `--overlap` (32) | Per-axis nominal tile size and overlap in voxels; the default overlap fraction is $32/256 = 12.5\%$. Overlaps above half the tile size are rejected. The last candidate tile of an axis is folded into its predecessor when its unique coverage is thinner than the overlap (Sec. 2); that predecessor then spans up to $T + L - 1$ voxels, 287 at the defaults, so the memory budget is sized for it. |
| `--seeds K` | An integer is a *whole-volume* budget, divided over the tiles by the mass-weighted rule of Sec. 5.2, Eqs. 6 to 8. A ratio in $(0, 1]$ is applied per tile unchanged. `fit_tiled(seeds=)` in Python is per tile. |
| `--saturation-exponent` (0.44) | The exponent $\alpha$ of Eq. 7; it also drives the content planner's $K \propto n_{\text{features}}^{\alpha}$. |
| `-j`, `--jobs` (1) | Number of tiles fitted concurrently as subprocesses on *one* GPU (`auto` sizes it from GPU memory, host RAM and CPU threads); ignored with `--tiling none` or `--tile`. Multi-GPU and cluster fan-out is `batch-fit` (Sec. 6). |
| `--tile k/M` | Fit one tile of the grid and exit, the worker form `batch-fit` and `-j` spawn; the parent passes the tile's resolved count as `--tile-seed-count`. A hand-run worker without it divides an integer budget equally over the non-empty tiles. |
| `--flat` | Merge to a single leaf instead of the default `kind=partition` (one part per tile). |
| `--recipe stream`\|`levels` | Per-part LOD *recipe* built at fit time (`stream`: additive ladder per part, the `tiles` topology; `levels`: coarse-to-fine levels per part, `adaptive`). Requires a tiled partition: rejected with `--flat`, `--tiling none`, `--tile` and the plan-only forms. |
| `--cull-retention` | Cumulative-amplitude retention of the merge (scope as in Sec. 5.1). |

but materialises every spatial axis of that frame as float32 in host memory, and only then extracts its tile with `spec.slices`; when its seed budget is an integer and no parent has handed it an exact count, it also scans every tile of the grid to count the non-empty ones. Each such worker holds one whole spatial frame, so a fit with $M$ tiles requests $M$ frames in aggregate; only the GPU footprint is set by the tile.

### 7 Empirical validation

The partition-of-unity argument of Sec. 3 says what happens when every tile fit reproduces its apodised target exactly. Fits are not exact, so the question the theory leaves open is whether the residual fitting error, which the taper cannot cancel, concentrates at the seams. This section measures that directly.

**Reproducibility.** Every (dataset, window, $T$, $L$) cell is deposited as one per-cell metrics record, with the run manifest beside them; the tables and figures below are aggregated from those records alone, and the deposited module diff (Sec. 5) and the per-tile memory records complete the set.

#### 7.1 Protocol

**Volumes.** Tiling is for volumes that do not fit, but the comparison needs a monolithic reference, so the sweep uses two volumes that do: the confocal blastocyst, a nuclear-lamina label (channel 0, $236 \times 275 \times 271$ voxels, 17.6 M), and the light-sheet *Tribolium* embryo ($991 \times 104 \times 965$ voxels, 99.5 M), both min–max normalised to $[0, 1]$ as in Supplementary Document 2. The two differ in what a seam can damage. The blastocyst is sparse and mostly background; its signal, the lamina (the labelled nuclear envelopes), occupies a small part of the grid. The embryo is a dense sheet of bright nuclei in which almost every tile boundary cuts through structure. They also differ in how the intensity mass is spread over a grid (Sec. 7.2). On the blastocyst the most-seeded tile of the 4-tile grid at $T = 256$, $L = 13$ holds 95% of the summed intensity and receives 88% of the seeds. On the embryo the most-seeded tile of the 16-tile grid at the same $T$ and $L$ holds 9% of the intensity and receives 7% of the seeds.

**Budget and fitting settings.** Every fit, monolithic or tiled, spends the same whole-volume budget of $K = 32{,}000$ seeds, the 32,000-seed point of the rate–distortion sweep of Supplementary Document 2, with that document's settings: Adam at learning rate 0.01, $L_1$ loss, dynamic *relocation*, culling to 99.9% of cumulative amplitude on the final (merged) splat set, no background floor. The tiled arms call the Python orchestrator `fit_tiled` directly, with a flat merged output. The tile grid and the seed split are the CLI's own: the grid is `compute_tile_specs` with its fold rule (Sec. 2), and the runner obtains the per-tile counts from the CLI's allocator (`_weighted_uniform_seed_counts`, Sec. 5.2) and passes them to `fit_tiled` as `tile_seed_counts`, exactly as `luxar gsplat fit --tiling uniform --seeds 32000` would. The counts sum to 32,000 in every cell; a single-tile grid receives the whole budget. The no-taper control reuses the counts of its tapered sibling, allocated with the real window before the window is replaced, so the two arms differ in the window only.

Both arms are capped at 2,000 iterations with an early-stopping patience of 500, one fit per cell (torch seed 32,000). Supplementary Document 4 shows 99% of the 20,000-iteration PSNR by 2,000 iterations on 75 of 85 cells, all of them whole-volume fits of 1,000 seeds or more. The sweep applies the same cap to tile fits of 25 to 29,465 seeds as a cost control and records no per-tile stopping iteration, so whether a sparsely seeded tile would improve under a longer cap is not tested here. Every cell records the luxar commit it ran at, the torch build (2.14.0, CUDA 13.0), the GPU, that the compiled CUDA splatting extension is in use, the tile grid the fitter builds and the seed count handed to every tile; the grid is re-derived from the volume shapes and asserted against every cell.

**Arms.** The taper arm runs the shipped half-Hann window at $T = 128$ and $T = 256$ for $L/T \in \{0, 0.05, 0.1, 0.2, 0.4\}$ ($L$ rounded to whole voxels, always $\leq T/2$), plus $T = 64$ at the sweep's nominal $L/T = 0.1$ ($L = 6$) on the blastocyst; the shipped default is $32/256 = 0.125$. On the embryo, whose smallest axis is 104 voxels, $T = 128$ is sampled at $L/T = 0.1$ and 0.2 and $T = 64$ is not run: it would shred the volume into $17 \times 2 \times 17 = 578$ tiles, $\lceil K/M \rceil = 56$ seeds each on average. At $L = 0$ the window is identically 1, so that cell is a hard cut with no overlap.

The *no-taper control* repeats the $T = 128$ overlaps with the window replaced by a unit window: the tiles still overlap, but the overlap band is fitted twice at full amplitude and double-counted in the union. This is the degenerate arm that shows what the partition of unity buys, as distinct from what overlap alone buys. Three blastocyst cells ($T = 256$, $L = 26$, 51, 102) fold to a single tile (Sec. 7.2): they have no seam, and they enter the table as a same-budget repeat of the monolithic fit through the tiled code path.

**Metrics.** Reconstruction quality is full-volume *PSNR* against the original,

$$\text{PSNR} \;=\; 10\log_{10}\frac{1}{\text{MSE}}\;\text{dB}, \tag{9}$$

with MSE the mean squared error over the volume normalised to $[0, 1]$, reported as the difference to the monolithic fit. The seam question is answered by a *residual profile*. Every voxel is assigned its distance $\delta$ (in voxels) to the nearest interior seam, the centre plane of an overlap band, or the shared boundary plane at $L = 0$, and the mean absolute residual $|\hat{V} - V|$ is binned by $b = \lfloor\delta\rfloor$ from 0 to 64. The monolithic residual is binned on the same distance map, which gives a profile with no tile structure on exactly the same voxels; we call it the *null* profile and normalise every tiled profile by it.

Two scalars summarise a profile pair, both computed on the binned profiles and count-weighted over the bins. *Near/mono* is the tiled residual over the near band $0 \leq b \leq \max(1, \lfloor L/2\rfloor)$ divided by the null over the same bins. The *seam excess* is the ratio of near-band to deep-band residual for the tiled fit divided by the same ratio for the null, with the deep band $\min(\max(L, 8), 56) \leq b \leq 64$. The near band is the taper half-width; these are analysis bands on the distance map, not per-face masks. The deep band starts a full overlap width from the seam (at least 8 voxels) and is clamped at 56 so that at $L = 102$ it still holds nine bins; since the taper occupies only $\delta \leq L/2$, the deep band lies where the window is identically 1 in every cell (Fig. 4).

The deep band does not always lie inside the tile interiors. The distance is to the nearest seam, so an interior tile holds no voxel farther than $(T - L)/2$ from one, and when $\min(\max(L, 8), 56) \geq (T - L)/2$ the whole deep band samples the outer margins of edge tiles. That is the case for the blastocyst at $T = 128$, $L = 51$ (stride 77, half-stride 38.5), whose excess is therefore not a seam-to-interior figure and is not interpreted below. At $T = 128$ with $L \leq 26$ and at $T = 64$ the bins beyond the half-stride likewise come from edge tiles, while every $T = 256$ band lies inside the interiors. (The runner also stores a seam scalar with a deep band starting at $\min(2\max(L, 8), 56)$, which at $L = 26$–51 starts 52–56 voxels out and leaves the band few voxels; at $L = 102$ the two bands coincide. The table reports the definition above.)

A seam excess of 1.0 means the tiled fit is no worse at the seams than it is in the middle of its tiles, relative to a fit that has no seams. Near/mono alone also carries the tiled fit's overall deficit, which is why the excess normalises by the deep band; *deep/mono*, the same ratio on the deep band, is reported where that deficit needs locating.

**Hardware and provenance.** The sweep runs as one sequential process on one NVIDIA RTX PRO 6000 Blackwell (Workstation Edition, 96 GB, sm_120), all 26 cells in a fixed order, with the

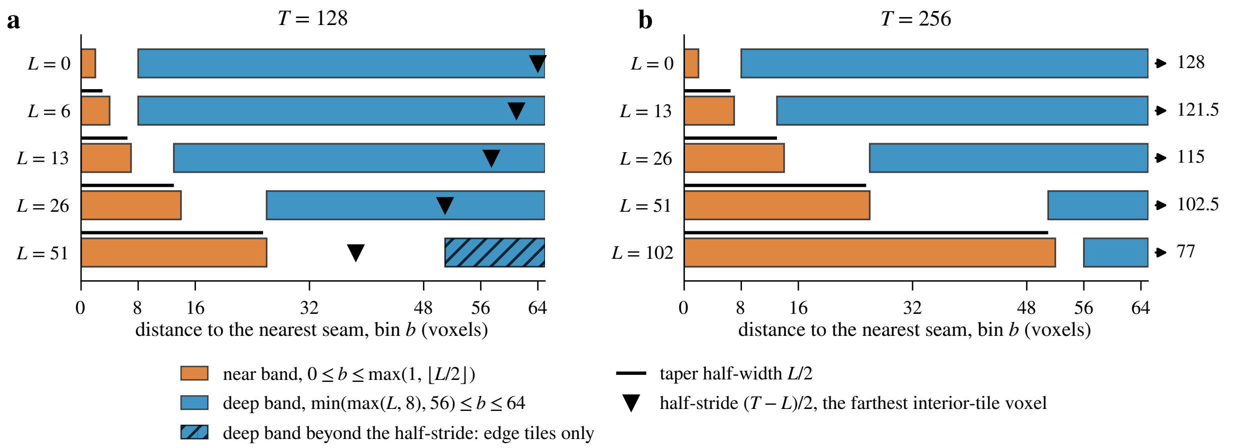

Figure 4: **The analysis bands on the distance-to-seam axis (schematic).** One row per overlap $L$ of the sweep at (a) $T = 128$ and (b) $T = 256$. The near band $0 \leq b \leq \max(1, \lfloor L/2 \rfloor)$ (orange) and the deep band $\min(\max(L, 8), 56) \leq b \leq 64$ (blue) are the bins of the residual profile that near/mono and the seam excess average over; the black bar is the taper half-width $L/2$, beyond which the window is 1. The triangle marks the half-stride $(T - L)/2$, the farthest any voxel of an interior tile lies from a seam; bins beyond it hold only the outer margins of edge tiles. At $T = 128$, $L = 51$ the deep band begins at or past the half-stride (hatched), so that cell's excess is not a seam-to-interior figure; at $T = 256$ every half-stride lies beyond the profile (arrows), so every band there is interior.

compiled CUDA splatting extension and no other fit on the card. Wall times are those of that process, one fit per cell. The sweep carries its own repeatability figure, since the two $T = 128$, $L = 0$ cells fit the identical target (the taper window is 1 everywhere at $L = 0$): they differ by 0.008 dB in PSNR and 6 splats, and take 379 and 405 s.

The peak memory of a whole `fit_tiled` call is not the per-tile footprint: the closing merged-quality pass renders the union over the full volume and sets the peak at essentially the monolithic value. The per-tile footprint is therefore measured separately, by a probe whose records (5 tile rows and the two whole-volume fits) are deposited with the analysis. The probe fits one tile per tile size: the most-seeded tile of the grid the fitter builds at the sweep's nominal overlap $L = \lfloor 0.1T \rceil$ ($L = 13$ for the blastocyst at $T = 256$, whose $L = 26$ grid is a single tile). The tile is fitted alone with the seed count the CLI allocator assigns it, through the same per-tile call `fit_tiled` makes, after one small warm-up fit in the same process. It records `torch.cuda.max_memory_allocated` for that fit, the peak of allocated tensor memory, which excludes the CUDA context and library workspaces; every row carries the commit, GPU and torch build.

The batch runs behind Sec. 7.4 used a uniform grid that keeps the last candidate tile of an axis unfolded and divides the budget equally over the non-empty tiles; this is the grid and the seed split that section reports.

##### 7.2 Tile grids and seed allocation

Table 2 lists the 18 distinct grids of the sweep, re-enumerated from the volume shapes and checked against the tile count, grid shape and tile slices every cell records. Its columns are the following.

Table 2: **Grids and seed splits of the sweep.** Rows grouped by volume; columns as defined in Sec. 7.2; $T$: tile size, $L$: overlap (voxels), $M$: tile count after the fold rule.

| $T$ | $L$ | grid | tiles | widest | seeds per tile min / median / max | busiest | tiles below $\lceil K/M \rceil/4$ tiles | seeds | intensity | pinned |
|---|---|---|---|---|---|---|---|---|---|---|
| *Blastocyst nuclear lamina* | | | | | | | | | | |
| 64 | 6 | $4 \times 5 \times 5$ | $125 \to 100$ | 64 | 71 / 278 / 925 | 3% | 3 | 0.7% | 0.2% | 0 |
| 128 | 0 | $2 \times 3 \times 3$ | 18 | 128 | 25 / 390 / 5,353 | 17% | 10 | 8.0% | 4.7% | 0 |
| 128 | 6 | $2 \times 3 \times 3$ | 18 | 128 | 60 / 693 / 4,869 | 15% | 3 | 1.7% | 1.0% | 0 |
| 128 | 13 | $2 \times 3 \times 3$ | $27 \to 18$ | 128 | 125 / 1,112 / 4,377 | 14% | 2 | 0.8% | 0.4% | 0 |
| 128 | 26 | $2 \times 3 \times 3$ | $27 \to 18$ | 134 | 292 / 1,760 / 3,505 | 11% | 2 | 2.2% | 2.9% | 0 |
| 128 | 51 | $3 \times 3 \times 3$ | $64 \to 27$ | 128 | 476 / 1,143 / 2,233 | 7% | 0 | – | – | 0 |
| 256 | 0 | $1 \times 2 \times 2$ | 4 | 256 | 56 / 1,240 / 29,465 | 92% | 3 | 7.9% | 4.7% | 0 |
| 256 | 13 | $1 \times 2 \times 2$ | 4 | 256 | 103 / 1,938 / 28,021 | 88% | 2 | 6.0% | 5.1% | 0 |
| 256 | 26 | $1 \times 1 \times 1$ | $8 \to 1$ | 275 | 32,000 (whole budget) | 100% | 0 | – | – | 0 |
| 256 | 51 | $1 \times 1 \times 1$ | $8 \to 1$ | 275 | 32,000 (whole budget) | 100% | 0 | – | – | 0 |
| 256 | 102 | $1 \times 1 \times 1$ | $8 \to 1$ | 275 | 32,000 (whole budget) | 100% | 0 | – | – | 0 |
| *Tribolium embryo nuclei* | | | | | | | | | | |
| 128 | 13 | $9 \times 1 \times 9$ | 81 | 128 | 46 / 434 / 636 | 2% | 1 | 0.1% | 0.1% | 0 |
| 128 | 26 | $10 \times 1 \times 9$ | $200 \to 90$ | 149 | 110 / 354 / 582 | 2% | 0 | – | – | 0 |
| 256 | 0 | $4 \times 1 \times 4$ | 16 | 256 | 1,203 / 2,100 / 2,477 | 8% | 0 | – | – | 0 |
| 256 | 13 | $4 \times 1 \times 4$ | $20 \to 16$ | 262 | 1,406 / 2,118 / 2,268 | 7% | 0 | – | – | 0 |
| 256 | 26 | $5 \times 1 \times 4$ | $25 \to 20$ | 275 | 261 / 1,857 / 2,423 | 8% | 4 | 4.1% | 4.1% | 0 |
| 256 | 51 | $5 \times 1 \times 5$ | 25 | 256 | 401 / 1,395 / 1,819 | 6% | 0 | – | – | 0 |
| 256 | 102 | $6 \times 1 \times 6$ | $49 \to 36$ | 256 | 590 / 876 / 1,199 | 4% | 0 | – | – | 0 |

- *grid*: the tiles per axis $(z, y, x)$ as `compute_tile_specs` enumerates them (the no-taper controls reuse the grids of their taper cells).
- *tiles*: the count, written *candidates* $\to$ *tiles* where the fold rule of Sec. 2 applies.
- *widest*: the largest tile extent on any axis (a folded tile reaches up to $T + L - 1$).
- *seeds/tile*: the counts the allocator of Sec. 5.2 resolves from $K = 32{,}000$.
- *busiest*: the share of $K$ in the most-seeded tile.
- *tiles below* $\lceil K/M \rceil/4$: the tiles seeded below a quarter of the equal share, with the share of $K$ they receive and the share of the volume's summed intensity they hold.
- *pinned*: the tiles the floor of Eq. 8 raised.

The intensity and pinned columns come from a re-run of the allocator on the volumes, whose counts are asserted equal to every cell's.

The fold rule acts on 11 of the grids: over the distinct grids it turns 727 candidate tiles into 494, and the widest tiles it produces are 134, 149, 262, 275 voxels. Two grids show what it does. On the embryo at $T = 128$, $L = 26$ the 104-voxel axis has a second candidate row with no voxel of its own and the 965-voxel axis a tenth with 21, both below 26, so the $10 \times 2 \times 10$ candidates become $10 \times 1 \times 9 = 90$ tiles, the last column 149 voxels wide. On the blastocyst at $T = 256$ the 236-voxel axis is one tile for any $L$, and at $L \geq 26$ the second rows on the other two axes (19 and 15 unique voxels) fold as well, so the three cells at $L = 26$, 51, 102 are single-tile fits of the whole volume.

The seed split follows the intensity. The ceiling $v_{\max}$ of Eq. 7 is 1.000 on the blastocyst and 0.969 on the embryo. On the sparse blastocyst the rule concentrates the budget where the lamina is: the 4-tile grid at $T = 256$, $L = 13$ puts 88% of $K$ into the tile holding 95% of the intensity, and on the 18-tile grids at $T = 128$ the most-seeded tile holds 22–24% of the intensity for 11–17% of the seeds. On the dense embryo the counts are close to even: at $T = 256$, $L = 13$ they run from 1,406 to 2,268 against an equal share of 2,000, and at $T = 128$, $L = 13$ from 46 to 636 against 396.

The quarter-share floor of Eq. 8 pins no tile on any of the 18 grids (0 pinned tiles), and no tile receives fewer than 25 seeds. The tiles seeded below a quarter of the equal share, 30 over 9 grids, hold at most 5.1% of a volume's intensity on any grid, and on 7 of those 9 grids they hold less intensity than seeds; on the blastocyst at $T = 128$, $L = 26$ they hold more, and on the embryo at $T = 256$, $L = 26$ the two shares are equal at the printed precision: they hold 2.9% and 4.1% of the intensity for 2.2% and 4.1% of the seeds. The largest gap between a tile's share of the seeds and its share of the intensity is 24 points, on the blastocyst at $T = 128$, $L = 51$, where the saturation exponent flattens the split of a grid whose busiest tile holds 31% of the intensity; on the embryo the gap is 2–7 points. Where a cell's residual is discussed below, this allocation is read alongside it.

#### 7.3 Seams

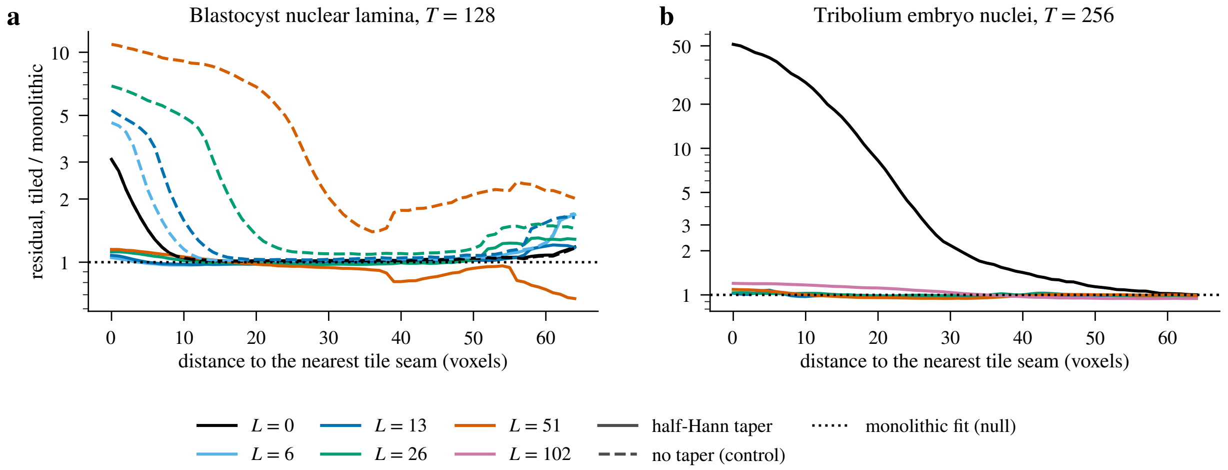

Figure 5: **Residual profiles against distance to the nearest tile seam.** For each cell, the mean absolute reconstruction error of the tiled union divided by that of the monolithic fit, binned by the distance $\delta$ (voxels) of each voxel from the nearest interior tile boundary of that cell's tile layout; the dotted line at 1 is the monolithic fit itself. One colour per overlap $L$ in both panels; tapered union solid, no-taper control dashed. (a) Blastocyst, $T = 128$; (b) *Tribolium*, $T = 256$. Bins holding fewer than $10^4$ voxels are dropped.

Figure 5 and Table 3 give the result. At zero overlap the seams are plainly visible in the residual: within two voxels of a boundary the blastocyst's residual is 2.19–2.84× the monolithic value and the embryo's is 50.40×, and the seam excess is 2.15–2.77 and 7.52 respectively. This is the failure mode a hard cut must produce: a Gaussian straddling the plane is fitted twice, once per tile, each time against a target truncated at the plane, and the two truncated fits neither

Table 3: **Every cell of the tiled-fitting sweep.** Rows grouped by volume. $T$: tile size; $L$: overlap (voxels); tiles: count after the fold rule (Table 2); seeds per tile: smallest and largest mass-weighted count on the grid; splats: count after the global 99.9% amplitude cull on the flat union; PSNR abs. and $\Delta$ to the monolithic fit of the same volume; seam near/mono and excess on the analysis bands of Sec. 7.1 (a single-tile grid has no seam; at $T = 128$, $L = 51$ the deep band lies outside the tile interiors); time: whole-call wall time divided by the tile count (monolithic: the whole fit); GPU: peak allocated tensor memory of one tile fit at that tile size, the most-seeded tile with the allocator's seed count, decimal GB (the probe of Sec. 7.1; monolithic and single-tile cells: the whole fit).

| window | $T$ | $L$ | tiles | seeds per tile | splats | PSNR (dB) abs. | $\Delta$ | seam near/mono | excess | time (s/tile) | GPU (GB) |
|---|---|---|---|---|---|---|---|---|---|---|---|
| *Blastocyst nuclear lamina* | | | | | | | | | | | |
| monolithic | – | – | 1 | 32,000 | 28,164 | 39.20 | – | – | – | 37 | 0.81 |
| taper | 64 | 6 | 100 | 71–925 | 27,764 | 39.04 | −0.16 | 1.03 | 1.07 | 20 | 0.01 |
| taper | 128 | 0 | 18 | 25–5,353 | 28,924 | 35.90 | −3.29 | 2.84 | 2.77 | 21 | 0.09 |
| taper | 128 | 6 | 18 | 60–4,869 | 29,477 | 39.25 | +0.06 | 1.03 | 1.03 | 14 | 0.09 |
| taper | 128 | 13 | 18 | 125–4,377 | 28,800 | 39.27 | +0.07 | 1.04 | 1.05 | 20 | 0.09 |
| taper | 128 | 26 | 18 | 292–3,505 | 28,819 | 39.14 | −0.06 | 1.07 | 1.08 | 26 | 0.09 |
| taper | 128 | 51 | 27 | 476–2,233 | 28,346 | 38.65 | −0.54 | 1.07 | 1.26 | 22 | 0.09 |
| taper | 256 | 0 | 4 | 56–29,465 | 27,740 | 38.61 | −0.59 | 2.19 | 2.15 | 22 | 0.71 |
| taper | 256 | 13 | 4 | 103–28,021 | 27,931 | 39.19 | −0.01 | 1.01 | 1.01 | 28 | 0.71 |
| taper | 256 | 26 | 1 | 32,000 | 28,171 | 39.20 | 0.00 | – | – | 40 | 0.81 |
| taper | 256 | 51 | 1 | 32,000 | 28,118 | 39.18 | −0.01 | – | – | 40 | 0.81 |
| taper | 256 | 102 | 1 | 32,000 | 28,189 | 39.21 | +0.01 | – | – | 42 | 0.81 |
| no taper | 128 | 0 | 18 | 25–5,353 | 28,918 | 35.91 | −3.28 | 2.84 | 2.76 | 23 | 0.09 |
| no taper | 128 | 6 | 18 | 60–4,869 | 29,461 | 31.18 | −8.02 | 4.17 | 3.91 | 14 | 0.09 |
| no taper | 128 | 13 | 18 | 125–4,377 | 28,797 | 27.94 | −11.26 | 4.55 | 4.31 | 17 | 0.09 |
| no taper | 128 | 26 | 18 | 292–3,505 | 29,132 | 23.57 | −15.62 | 5.65 | 5.10 | 24 | 0.09 |
| no taper | 128 | 51 | 27 | 476–2,233 | 29,172 | 17.45 | −21.75 | 9.03 | 4.08 | 29 | 0.09 |
| *Tribolium embryo nuclei* | | | | | | | | | | | |
| monolithic | – | – | 1 | 32,000 | 31,772 | 51.34 | – | – | – | 88 | 4.51 |
| taper | 128 | 13 | 81 | 46–636 | 27,170 | 50.36 | −0.98 | 1.19 | 1.13 | 28 | 0.07 |
| taper | 128 | 26 | 90 | 110–582 | 27,325 | 50.56 | −0.78 | 1.07 | 1.05 | 20 | 0.07 |
| taper | 256 | 0 | 16 | 1,203–2,477 | 30,641 | 29.50 | −21.84 | 50.40 | 7.52 | 26 | 0.33 |
| taper | 256 | 13 | 16 | 1,406–2,268 | 30,478 | 51.53 | +0.19 | 1.04 | 1.05 | 27 | 0.33 |
| taper | 256 | 26 | 20 | 261–2,423 | 30,127 | 51.42 | +0.09 | 1.02 | 1.03 | 30 | 0.33 |
| taper | 256 | 51 | 25 | 401–1,819 | 29,937 | 51.13 | −0.21 | 1.01 | 1.01 | 31 | 0.33 |
| taper | 256 | 102 | 36 | 590–1,199 | 29,342 | 49.66 | −1.68 | 1.09 | 1.15 | 30 | 0.33 |
| no taper | 128 | 13 | 81 | 46–636 | 27,698 | 20.52 | −30.81 | 69.92 | 4.38 | 23 | 0.07 |

match each other nor sum to the original. The light-sheet volume, in which boundary planes run through dense nuclei, loses 21.8 dB of full-volume PSNR to this alone.

With the half-Hann taper and an overlap of 5–20% of the tile size, the seams disappear from the blastocyst's residual. Over its 5 such cells, $T = 64$ (100 tiles), $T = 128$ (18 tiles) and $T = 256$ (4 tiles), the near-band residual is 1.01–1.07× the monolithic value and the deep-band residual 0.97–1.00× it, so the seam excess, their ratio, is 1.01–1.08. The tiled fit is at the same time within −0.16 to +0.07 dB of the monolithic fit in full-volume PSNR (Sec. 7.5). The profiles in Fig. 5 show no boundary spike on the blastocyst: over the first 30 voxels the ratio stays between 0.97 and 1.12, and on the *Tribolium* between 0.95 and 1.09. At $L = 0$ the same ratio is 3.1 (blastocyst) and 51.2 (*Tribolium*) in the $\delta = 0$ bin and falls below 1.2 by $\delta = 7$ and $\delta = 48$ voxels respectively, and the no-taper controls start at 5–11.

Each cell is a single fit and no seed replicate of a tapered cell was run (the only repeat, the two $L = 0$ cells, agrees to 0.008 dB), so a seam signal below a few percent of the residual would not be resolved here. The 100-tile cell, with 71 to 925 seeds per tile, is as clean as the 18-tile ones (excess 1.07 against 1.03–1.08), so on a sparse volume the tile count itself is not what matters over this range.

On the dense embryo the same configurations give the same answer. Over its 5 tapered cells at 5–20% overlap, $T = 256$ (16 to 25 tiles) and $T = 128$ (81 and 90 tiles), the near-band residual is 1.01–1.19× the null and the deep-band residual 0.99–1.05× it, so the excess is 1.01–1.13, and the union lands −0.98 to +0.19 dB from the monolithic fit. The tile count sets where in that band a cell falls. At $T = 256$ the three grids of 16 to 25 tiles, with 261 to 2,423 seeds per tile, have an excess of 1.01 to 1.05 and land −0.21 to +0.19 dB from the monolithic fit.

The two $T = 128$ grids, whose 81 and 90 tiles hold 46–636 and 110–582 seeds each, carry the largest near-band ratio (1.19 at $L = 13$) and the largest deficits, −0.98 and −0.78 dB, with the deep band at 1.02–1.05× the null. A tile of a few hundred seeds reproduces its nuclei a little less well throughout, and the excess of 1.05–1.13 on those two grids is the residual of a small tile fading out against the residual of its neighbour fading in. The taper does not fail on the dense volume; the small tiles cost a fraction of a decibel.

**The overlap of** 40%. On the blastocyst the 27-tile grid at $T = 128$, $L = 51$ loses 0.5 dB with near/mono 1.07; its deep band lies outside the tile interiors (Sec. 7.1), so its excess of 1.26 is not read as a seam figure. The voxels covered by two or more tiles are most of the volume (78%, Sec. 7.5). On the embryo the 36-tile grid at $T = 256$, $L = 102$, whose deep band is interior, loses 1.7 dB with excess 1.15, near/mono 1.09 and deep/mono 0.94, with 77% of its voxels in two or more tiles.

**The no-taper control.** Overlap alone is not what removes the seams; the partition of unity is. With the window replaced by 1, the same overlapping tile grids produce unions that are worse than a hard cut, and monotonically worse the more they overlap: $8.0, 11.3, 15.6, 21.7$ dB below the monolithic fit at $L/T = 0.05$, 0.1, 0.2 and 0.4 on the blastocyst (3.3 dB at $L = 0$). Their near-band residual is 4.17–9.03× the null and their seam excess 3.91–5.10. On the embryo at $T = 128$, $L/T = 0.1$, the control lands 30.8 dB below, with a near-band residual 69.92× the null.

The mechanism is the one Sec. 4 predicts. Two tiles (four at an edge, eight at a corner) each fit the overlap band at full amplitude, so the union carries twice the mass there or more, and the wider the band the larger the fraction of the volume rendered at multiple brightness. The tapered union of the same tiles, at the same overlaps and seed counts, is −0.54 to +0.07 dB from the monolithic fit on the blastocyst (Fig. 7, solid against dashed), with a seam excess of 1.03 to 1.08 at $L = 6$, 13 and 26 (the $L = 51$ sibling's excess is not a seam figure, Sec. 7.1). On the embryo it lands 1.0 dB below the monolithic fit against the control's 30.8 dB, with excess 1.13 against 4.38. That is the partition-of-unity argument of Sec. 3 measured: with the taper the windowed copies sum to one copy of the structure, without it they sum to two or more.

##### 7.4 Seams in amplitude, not in mass

The residual ratio above is a statement about reconstructed intensity, and reconstructed intensity is what the union of tile fits conserves. It is not what the viewer colours. A colour-mapped scene looks up each splat's colour from its own amplitude before compositing, so two fits with the same summed intensity can look different if they distribute that intensity over splats of different size, and a tile grid can be invisible in the residual and visible on screen. We met this on a fit of the kind shown in Supplementary Video 5 and measured it the following way, with the batch fitter (`luxar gsplat batch-fit`) of a build that is not the sweep's and whose run records carry no commit; it keeps the last candidate tile of an axis unfolded and splits the budget equally over the non-empty tiles (Sec. 7.1), without the fold rule and the mass-weighted split of Sec. 5.

One *Drosophila* frame ($108 \times 1352 \times 532$ voxels, floor 7.1 subtracted, 16-voxel slabs along the long axis) is fitted three times; the per-slab profiles, the run manifests and the per-tile seed counts are deposited with the analysis. First, uniform 512-voxel tiles with the 32-voxel taper and a 64,000-seed budget. The grid is $3 \times 2$, its second column a 52-voxel strip with 20 voxels of its own; one tile was found empty and each of the other 5 was seeded with 12,800. The frame holds 64,210 splats because the third-row full tile realises 13,010 from its 12,800 seeds, and the two strip tiles carry 25,600 of them. Second, content tiling from the frame's calibration (8 boxes, 683,899 splats). Third, a single tile seeded with 220,447 splats, the count the frame's *blind-spot cross-validation* retains after the cull at its top budget of 256,000 seeds (a `luxar gsplat cal` sweep of 6 budgets to 256,000 seeds at timepoint 150), the operating point of Table 1 of the main text; the batch workers apply no cull, so the seeded count is the splat count. For each slab we divide the summed splat *mass* (amplitude times the product of the three spatial standard deviations) and the summed *amplitude* by the slab's raw intensity, normalised to the interior median.

Mass tracks the raw intensity in all three fits, seam bands included (Fig. 6a): per-tile medians (over the three uniform tiles split at the band centres, for all three fits) of 0.98–1.01 for the uniform grid, 0.99–1.00 for the content boxes and 1.00 for the single tile, with 5th–95th percentiles over the interior of 0.86–1.14, 0.98–1.01 and 0.96–1.02. Amplitude does not (Fig. 6b). On the uniform grid it steps from 1.06 in the anterior tile to 0.79 in the middle tile and 1.84 in the posterior one, and falls to 0.65 and 0.49 in the two slabs of the 480–512 crossfade, which hold 1,549 and 1,288 splats against 1,069–1,138 in the two adjacent anterior slabs and 832–889 in the two adjacent middle ones. Equal seeds for unequal content give the busiest tile the coarsest, dimmest splats and the sparse tip the finest, brightest ones, and inside the taper each tile carries the shared structure at half amplitude. Content tiling removes the steps (per-tile medians 0.95–1.03, 0.97–1.01 in the former band positions) and the single tile has none (0.91–1.14, a smooth anterior-to-posterior gradient).

Viewed with a magma colour map whose top the nuclei reach, the uniform fit showed each tile in a different hue and a violet band at every seam, the content fit only thin lines along its box boundaries and the single tile nothing; no capture of these views is deposited. The metric of Sec. 7.3 would have passed all three. The remedy is to budget tiles by content, which is what the mass-weighted split of Sec. 5.2 does, and, for a colour-mapped scene, an intensity window that saturates the structure of interest, so that a halved amplitude still lands on the same colour.

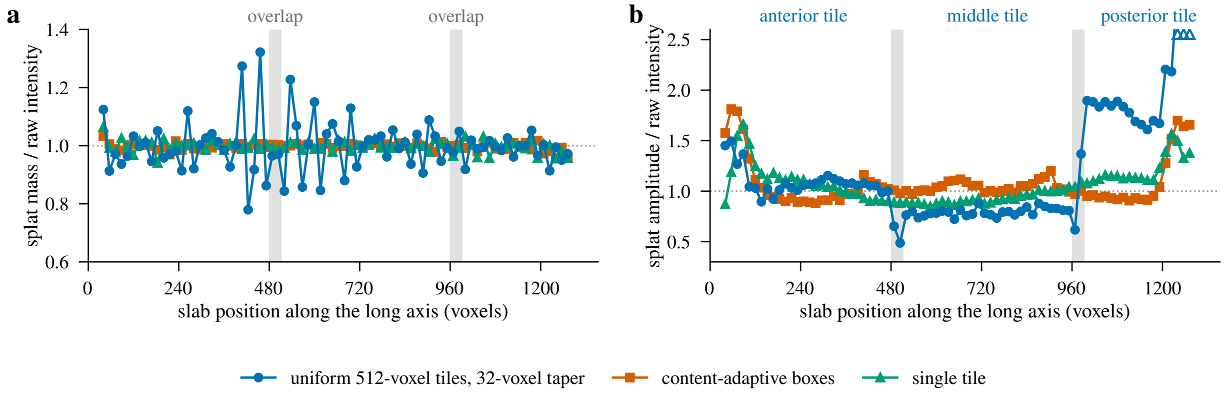

Figure 6: **Mass and amplitude per slab of the *Drosophila* frame.** For each 16-voxel slab along the long axis, (a) the summed splat mass and (b) the summed splat amplitude of the three fits, each divided by the slab's raw intensity and normalised to its interior median; the dotted line is 1. Grey bands are the two 32-voxel overlaps of the uniform grid, whose three tiles are named above (b); open triangles at the top edge of (b) mark three posterior slabs of that grid above the axis (peak 3.24). Slabs holding under 5% of the brightest slab's intensity are not drawn.

##### 7.5 Reconstruction quality

Tiling at a fixed budget need not cost quality, and on the blastocyst it does not (Fig. 7). In the 5 moderate-overlap cells the tapered union lands within −0.16 to +0.07 dB of the monolithic fit (39.0–39.3 against 39.20 dB) at essentially the same realised splat count (27,764–29,477 against 28,164). The three single-tile cells, the same budget through the tiled code path with its shared normalisation and merge-time cull, land −0.01 to +0.01 dB from the monolithic fit, so the path itself costs nothing.

Two effects pull in opposite directions, and one fit per cell does not separate them. The mass-weighted split gives the tiles holding the lamina the budget the background would have wasted, and each tile's optimiser then works on a smaller problem at the same iteration cap. Against it, the overlap bands are represented more than once: a structure straddling a face is modelled by two windowed halves (four copies at an edge, eight at a corner), each spending its own splats on what one Gaussian would cover. Table 4 quantifies the duplication. On an unbounded grid the fraction of the volume covered by two or more tiles is $1 - ((T-2L)/(T-L))^3$ and the share of tile samples that are duplicates is $1 - ((T-L)/T)^3$. The finite grids fitted here, with their clamped and folded edge tiles, sit below the infinite-grid covered fraction (enumerated from the tile specs).

The duplication grows with $L$ while the taper's benefit saturates at the smallest overlap, so more overlap does not help: on the blastocyst's $T = 128$ grid the delta moves from +0.06 through +0.07 and −0.06 to −0.54 dB at $L/T = 0.4$.

On the embryo the picture is the same to within a decibel. The 5 moderate-overlap cells land −0.98 to +0.19 dB from the monolithic fit (50.4–51.5 against 51.34 dB) at 27,170–30,478 splats against 31,772, and the 40% cell lands 1.7 dB below. The deficit follows the tile count, which sets the per-tile budget. At $T = 256$ the deltas from $L/T = 0.05$ to 0.4 are +0.19, +0.09, −0.21 and −1.68 dB over grids of 16, 20, 25 and 36 tiles. The two $T = 128$ grids of 81 and 90 tiles, with a

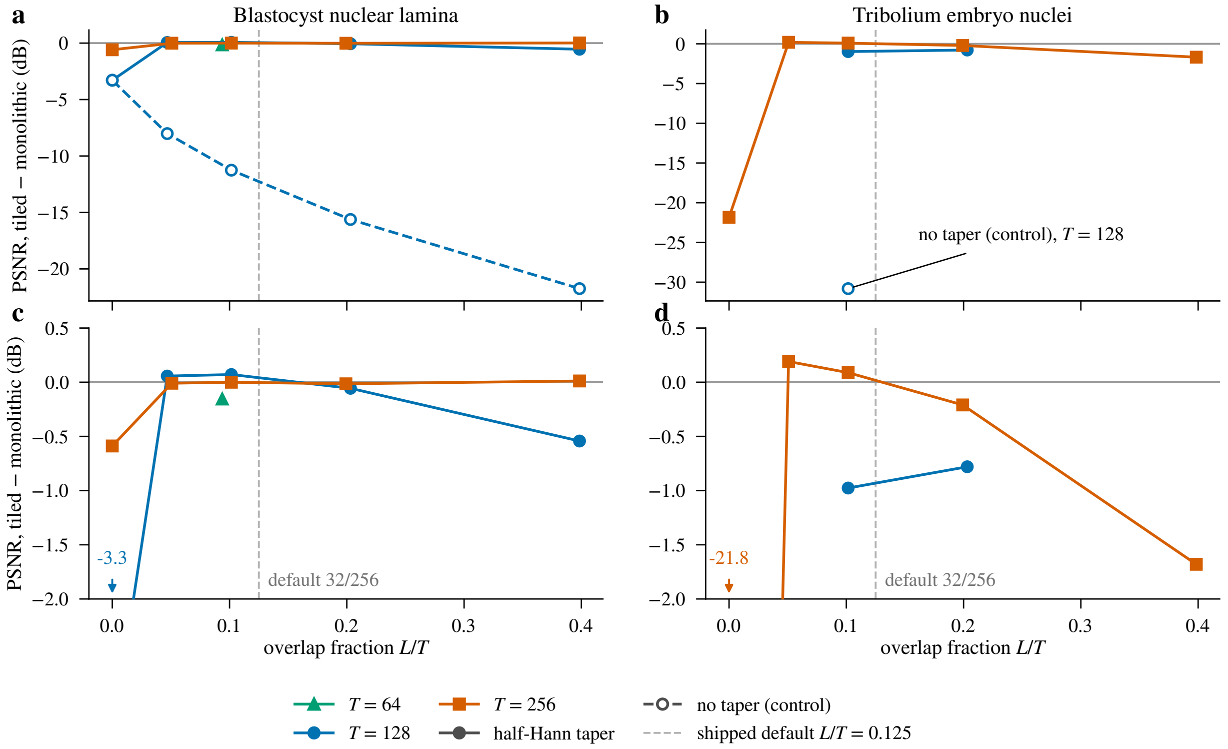

Figure 7: **Full-volume PSNR of the tiled union relative to the monolithic fit**, at the same total budget of 32,000 seeds, against overlap fraction $L/T$; one colour and marker per tile size, half-Hann taper solid, no-taper control dashed with open markers. (a, b) Every cell; the lone control on the *Tribolium* is labelled. (c, d) The tapered arms alone on a 2.5 dB window; a point below the window is marked at the lower edge with its value. Zero is the monolithic fit and the vertical dashed line is the shipped default $L/T = 32/256$. The blastocyst points at $T = 256$, $L/T \geq 0.2$ are single-tile grids (Sec. 7.2).

Table 4: **Voxel duplication against overlap fraction.** Covered: the fraction of the volume lying in two or more tiles; duplicates: the share of tile samples that are copies of a voxel already in another tile; both closed-form on an unbounded grid, and the covered fraction enumerated from the tile specs of the sweep's finite grids (range over cells at that $L/T$).

| $L/T$ | covered (unbounded) | duplicates (unbounded) | covered (sweep grids) |
|---|---|---|---|
| 0.05 | 15% | 14% | 8–11% |
| 0.1 | 30% | 27% | 18–23% |
| 0.2 | 58% | 49% | 37–42% |
| 0.4 | 96% | 78% | 77–78% |

median of 434 and 354 seeds each, lose 1.0 and 0.8 dB (Table 2). At the rate–distortion slope of this operating point (+0.80 and +0.84 dB per doubling of $K$ between the 16,000, 32,000 and 64,000 points on the blastocyst, from the rate–distortion table of Supplementary Document 2), a deficit of a decibel is what one halving of the budget would cost. The finest grids therefore behave like a fit at roughly half the budget: the duplicated overlap bands and the per-tile iteration cap take that much on a volume in which every tile is full of structure.

The practical guidance is therefore the shipped default and one allowance. Overlap near

$L/T = 0.1$ (the default is 32 voxels at $T = 256$, 12.5%): on both volumes every overlap from 5 to 20% is equivalent within the measurement, and 40% costs 0.54–1.68 dB. The mass-weighted split needs no correction on these volumes; a raised `--saturation-exponent` flattens the weighting, and a `fit_tiled(tile_seed_counts=)` call from Python takes any split. And spend a budget above what the monolithic rate–distortion curve would suggest when a volume must be tiled finely, by a factor a tiled budget sweep would have to establish.

#### 7.6 Cost

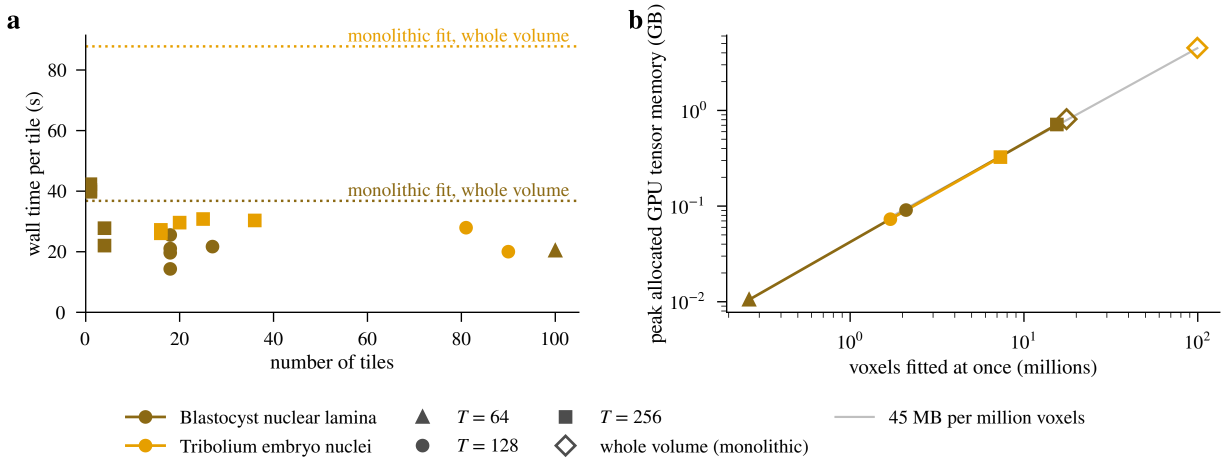

Figure 8: **Cost of tiling.** (a) Wall time per tile (the whole `fit_tiled` call, merge and quality pass included, divided by the tile count) against the number of tiles; one colour per volume, one marker per tile size, the monolithic fit's wall time dotted and labelled. Each tile has its own cap of 2,000 iterations, so the sequential total grows with the tile count and the saving is realised only through parallelism. (b) Peak allocated GPU tensor memory of a single tile fit against the number of voxels fitted at once (the probe of Sec. 7.1), the whole-volume fit as the open diamond and the 45 MB-per-million-voxel line in grey (decimal units).

Tiling is a memory device, not a speed-up. Each tile is an independent optimisation that runs its own iteration budget, so the sequential wall time of a tiled fit is roughly the tile count times the per-tile time. On these volumes, which fit whole, the elapsed time of the tapered `fit_tiled` calls with more than one tile is 2.4–55.3× the monolithic fit's 37 s on the blastocyst and 4.8–25.8× its 88 s on the embryo, i.e. 14–31 s per tile when the whole call is divided by the tile count. These figures are descriptive of one sequential process (Sec. 7.1). What tiling buys is that the $M$ tiles need no communication and can run on $M$ GPUs or Slurm array tasks at once.

The memory saving is what the tile size sets, and it is proportional. The per-tile probe (Fig. 8b; Table 3, last column; decimal MB and GB throughout) puts the peak allocated tensor memory of a single tile fit at 73 MB for a 128-voxel tile of the embryo (128 × 104 × 128, 1.7 M voxels, 636 seeds) and 325 MB for the folded 256-voxel tile (256 × 104 × 275, a folded-axis case, 7.3 M voxels, 2,423 seeds). The whole 99.5 M-voxel volume at 32,000 seeds takes 4.5 GB, a 62× and 14× reduction for the two tiles. On the blastocyst the 128-voxel tile takes 91 MB against 811 MB for the whole volume (9×), while the 256-voxel tile of the 4-tile grid, which covers 15.5 of the volume's 17.6 M voxels, saves almost nothing (713 MB).

Across every probe above a million voxels the peak is 43–46 MB per million voxels fitted at once, tile or whole volume alike, and the 64-voxel tile (0.3 M voxels, 10 MB) sits at 40.1 MB per million voxels. At these budgets the footprint is the voxel grid and its gradients, not the splats: the probed tiles carry 636 to 28,021 seeds, and the per-voxel figure does not follow them. The footprint of a fit is therefore predicted by its voxel count, whole volume or tile, whatever seed count the allocator hands it. Each probe follows one small warm-up fit in the same process, so one-time workspace allocation does not land in its row.

Because the footprint follows the voxel count, the memory rule must be stated for the widest tile the fold rule produces, up to $T + L - 1$ voxels on a folded axis ($134, 149, 262, 275$ here), not for $T$. The rule of thumb for the memory budget in Sec. 5 follows. A $256^3$ tile (16.8 M voxels) needs 0.7–0.8 GB of allocated tensor memory by this rule and a tile folded on all three axes ($287^3$, 23.6 M voxels) 1.0–1.1 GB, so a 24 GB card would hold some 22 such tiles concurrently (fewer with headroom) or one $\sim$520 M-voxel fit. These are extrapolations from single-process probes of 1.7–99.5 M voxels, with no concurrent-tile run measured and no allowance for the CUDA context and library allocations the metric excludes, so they need headroom.

##### 7.7 What this does and does not show

Within its scope the sweep answers the seam question. With the shipped taper, overlaps of 5–20% of the tile size keep the seam-to-interior residual ratio within 1.01 to 1.13 of the seam-free fit's on the confocal blastocyst and the light-sheet embryo, at most 13% above it and that on the 81-tile grid ($T = 128$, $L = 13$) of the dense embryo; the near-band residual itself is 1.01–1.19× the monolithic residual on the same voxels. This holds at tile sizes from 64 to 256 voxels and grids of 4 to 100 tiles, within $-0.98$ to $+0.19$ dB of the monolithic fit in full-volume PSNR; the no-taper control shows that this is the taper's doing, not the overlap's, more overlap buys nothing at this budget and 40% costs 0.54–1.68 dB.

The seed split is measured at one exponent ($\alpha = 0.44$) with the allocator's quarter-share floor, which binds on none of the 18 grids. Whether the finest grids, whose tiles of a few hundred seeds cost up to a decibel on the dense volume, would close that under a longer iteration cap or a larger budget is not tested.

What the sweep does not exercise is the regime tiling exists for. A volume larger than GPU memory has no monolithic reference, and the two volumes here are small enough that even $T = 256$ yields 4 to 36 tiles. Seam excess did not change with tile count on the blastocyst (1.03 at 18 tiles, 1.07 at 100). Tile count, geometry and per-tile seed allocation vary together in this sweep, so the 729-tile grid of Sec. 5 remains an untested extrapolation. Nor does the sweep test the CLI's content tiling (core-centre selection, no apodisation); its seams would need their own measurement. Sec. 7.4 adds that a grid over an unevenly filled volume can conserve intensity and still show its tiles, each tile's splats carrying a different amplitude per unit of mass.

#### 8 Summary and practical rules

The rules below are what the derivation and the sweep support for a uniform tiled fit.

**What to set.**

- **Tile size** is the memory knob. The working set of a fit is 43–46 MB of allocated tensor memory per million voxels fitted at once, whatever seed count the tile receives. Size it for the widest tile the fold rule can produce, $T + L - 1$ voxels per axis: a $256^3$ tile needs 0.7–0.8 GB and a $287^3$ folded tile 1.0–1.1 GB, plus headroom for the CUDA context, which the probe excludes.

- **Overlap** near 10% of the tile size (the shipped default is 32 voxels at $T = 256$, 12.5%). Every overlap from 5 to 20% was equivalent within the measurement; 40% cost 0.54–1.68 dB through duplicated overlap bands. Overlaps above $T/2$ are rejected (the two-ramp partition of unity fails there).

- **Taper** always on. Overlap without the taper is worse than a hard cut, 8.0–30.8 dB below the monolithic fit, and worse the wider the overlap.

- **Floor** resolved once against the whole volume and subtracted before windowing; a per-tile floor reintroduces the boundary step.

- **Seeds** as one integer whole-volume budget, left to the mass-weighted split of Sec. 5.2. On the 18 grids of the sweep the quarter-share floor bound nowhere and no tile received fewer than 25 seeds. Equal seeding of unequal tiles produces amplitude steps that are invisible in the residual and visible on screen (Sec. 7.4). A volume that must be tiled finely should be given more than the monolithic rate–distortion curve suggests: the finest grids of the dense embryo cost about a decibel, one halving of the budget.

- **Output shape**: the default `kind=partition` for viewing (frustum culling, one ladder per part), `--flat` for a single leaf with one global cull, `batch-fit` for several GPUs or a cluster.

**What it costs.** Tiling saves memory, not time. Run sequentially, the tapered calls took 2.4–55.3× the monolithic wall time on the blastocyst and 4.8–25.8× on the embryo; the saving is realised only when the tiles run in parallel (input reads are tile-local on the `batch-fit` path only, Sec. 6). At a fixed budget the tiled union lands $-0.98$ to $+0.19$ dB from the monolithic fit over the 10 moderate-overlap cells, with a seam excess of 1.01 to 1.13.

**What was not measured.** No volume larger than GPU memory was fitted, so the 729-tile default grid is an extrapolation from grids of at most 100 tiles. Every cell is one fit without a seed replicate. The iteration cap of 2,000 was not varied, so whether sparsely seeded tiles recover under a longer cap is open. No concurrent tiles were run on one card, and the memory metric excludes the CUDA context. The seams of content tiling, which merges by core-centre selection, were not measured. The amplitude seams of Sec. 7.4 were observed on one *Drosophila* frame with an equal seed split; the mass-weighted split was not tested on that frame.
