## Supplementary material for "Luxar: Gaussian splatting for microscopy and scalable interactive web visualisation of multidimensional scientific data": supp_doc_12_codec_selection

### Splat Parameter Encoding: Quantisation Fidelity and Codec Selection, Natively and in the Browser

Supplementary Document 12 – Luxar

**Abstract**

A Luxar store does not hold the fitted `float32` splat parameters. It quantises them to integer codes (16-bit fixed-point centres, 8-bit Cholesky factors that pass an encode-time error certificate, and amplitudes on an 8- or 16-bit grid chosen from the array's dynamic range) and then compresses the codes with a general-purpose codec. This document measures both steps. On fresh fits of 17 benchmark volumes at up to two operating points each (30 cells, the two coinciding on four volumes), quantisation changes the PSNR against the original volume by at most 0.679 dB (median 0.012 dB), an error 0.8 to 4.3 orders of magnitude below the fit's own error, while the store shrinks from 30–37 to 8.8–12.7 B/splat. A per-chunk sweep of 164 codec configurations over one light-sheet fit of 2,544,311 splats, decoded natively and in the browser, shows that the codec is the smaller lever: the quantisation tiers cut the raw payload by 2.9×, the codec adds tens of percent. Three findings follow. Blosc silently skips bit shuffle on any block whose element count is not a multiple of eight, so the filter a store pays for depends on its chunk size. Decode speed depends only weakly on the encoder level, but the browser reorders codecs: the `lz4` class loses 4.5–5.1× where `zstd` loses 3.2–3.4×. The shipped width-keyed `zstd` rule stores the fit at 9.63 B/splat; keying the filter on the code type as well saves a further 1.44%.

#### 1 Introduction

A fitted Gaussian splat in three dimensions is ten `float32` numbers: three centre coordinates, six Cholesky factors of its covariance and one amplitude. Stored as such, a scene costs 40 B per splat before any compression. Luxar's writer instead quantises each parameter array to integer codes on a grid fitted to that array (§2.1 lists the tiers), and it compands two of them first: Cholesky factors and wide-range amplitudes pass through a logarithm before uniform quantisation, so the grid is finer where values are small (log-companded codes). The codes are then compressed chunk by chunk with a general-purpose codec, preceded by an optional filter (byte shuffle, bit shuffle or a delta prefilter) that rearranges the bytes so that the codec sees longer repeats. Figure 1 sketches that path from the fitted parameters to the viewer and marks the two points at which this document measures it.

Quantisation and codec are separate levers with different currencies. Quantisation is lossy and changes the stored size by integer factors, and its cost is a reconstruction error. Codec selection is

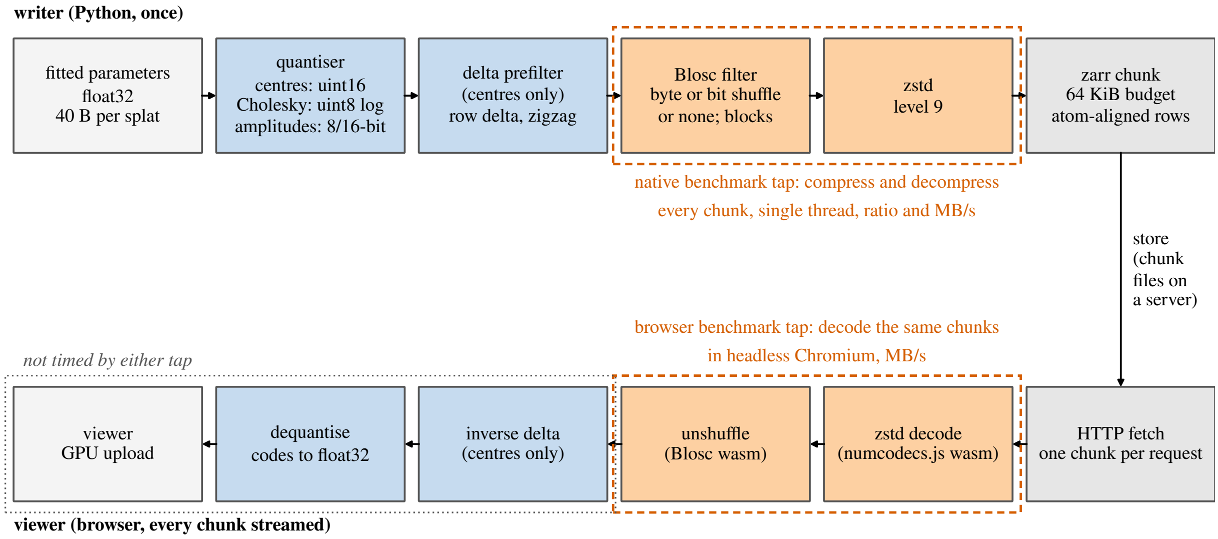


Figure 1: Schematic of the path a splat parameter takes from the writer to the viewer (no data plotted). The writer quantises the float32 parameters to integer codes, applies the delta prefilter to the centres, passes each chunk through a Blosc filter (byte shuffle, bit shuffle or none) and `zstd`, and writes it as a zarr chunk sized to a 64 KiB budget and aligned to the spatial-index atom. The viewer fetches one chunk per request, decodes it with the WebAssembly builds of the same codecs, inverts the delta filter and dequantises. The dashed boxes mark the two benchmark taps: the native sweep compresses and decompresses every chunk on one thread (§2.2); the browser run decodes the same compressed chunks in headless Chromium (§2.3). The dotted box marks the viewer stages neither tap times.

lossless and changes the size by tens of percent, and its cost is decode time, which a viewer pays on every chunk it streams. The two need different experiments, and this document runs both: a fidelity study of the quantised encoding on 17 benchmark volumes, and a codec sweep on the code arrays of one large fit.

The codec sweep is run natively and in the browser because the viewer is a browser. It reads zarr chunks through JavaScript and WebAssembly builds of the codec libraries, single-threaded and with an extra copy across the JavaScript/WebAssembly boundary per chunk. A codec that wins natively can lose there, so every deployable configuration is decoded both natively and in headless Chromium, and only configurations the viewer can decode are candidates for the shipped rule.

#### Terms used in this document

**Morton and Hilbert ordering** Two space-filling curves that sort points so that neighbours in space become neighbours in memory or on disk: Morton (Z-order) interleaves the bits of the quantised coordinates, Hilbert follows a continuous curve whose consecutive cells are always adjacent, so it has no long jumps between neighbouring runs. The compiler orders the stored elements along one of them; the fitter re-sorts its splats by Morton code periodically for GPU locality, and the substitutive LOD builder uses a Morton sort as its warm-start partition.

**atom** The spatial index's `chunk_size` in elements, of which every zarr chunk of a per-element array is a whole multiple, so an index range never straddles a zarr chunk.

**held-out voxels** The masked voxels of a blind-spot split, whose original values the optimiser never sees and against which the held-out PSNR of each candidate splat count is scored.

$\boldsymbol{K^*}$ The splat count recommended by a blind-spot sweep: the interior maximum of the held-out PSNR curve when there is a clear peak, the smallest count within 0.3 dB of the maximum on a plateau, and the largest count tested when the curve is still rising there (signal-limited). The shared operating-point table of these documents takes the knee instead, the smallest count within 0.3 dB of that maximum, which is the largest count itself on the two signal-limited benchmark volumes.

**signal-limited curve** A held-out PSNR curve whose maximum is the largest splat count tested and which is still climbing there (a total rise of at least 0.3 dB across the sweep, a trailing mean rise of at least 0.1 dB per step and a final step of at least 0.05 dB), so the peak lies beyond the sampled range.

**cull (retention)** The removal of splats after a fit. The fit-time cull ranks splats by amplitude and keeps the top set accounting for a retention fraction of the total amplitude (0.95 by default); the contribution-based cull instead removes the splats whose contribution to the rendered field falls below an error budget or a fractional threshold, and is the baseline that removal-based level-of-detail methods use.

**PSNR** The peak signal-to-noise ratio, $10 \log_{10}(\text{peak}^2/\text{MSE})$ in decibels, where peak is the reference range of the values compared; unless a document states another range, the supplements take peak 1 on volumes normalised to $[0, 1]$, so that $\text{PSNR} = -10 \log_{10} \text{MSE}$.

**SSIM** The structural similarity index, the mean over local windows of a product of luminance, contrast and structure comparisons between two images, lying in $[-1, 1]$ with 1 meaning identical.

### 2 Methods

#### 2.1 Dataset and arrays under test

All codec experiments use the last timepoint (`t252`) of the Zebrafish embryo nuclei light-sheet time-lapse (`h2afva`; the same series used in the main text), fitted with the content-adaptive tiled pipeline. That pipeline fits the volume in 43 content boxes, variable-size regions sized to the local splat density, whose fits were concatenated, giving $N = 2{,}544{,}311$ splats over a $404 \times 1976 \times 2039$ voxel extent. This is a scratch fit of one frame of the 10-timepoint batch on the benchmark workstation. The published time-lapse archive (Supplementary Document 14) retains the frames $t0, t5, \ldots, t250$ and does not include it. The fit was saved with the default writer, which sorts the splats along a Hilbert curve (*Morton and Hilbert ordering*) [Skilling, 2004]. The extent, the amplitude range and the covariance certificate quoted here are reproducible from the deposited writer-chunk manifest, which records the per-array encoding attributes of the same fit as re-written by the reference writer (writer-chunk stores, below). The quantisation tiers described in this document are those of the reference software state, and the stores' recorded dtypes and grid types are the ones that state writes. What the deposited record does not contain about these stores is listed in §4.5.

The benchmark operates on the *code arrays* as the reference writer encodes them (the stored integers, read back from zarr and re-chunked, §2.2) under three encoding scenarios, two of which the figures and tables report:

- **AUTO** (the default; scenario key `shipped` in the results files). The `centers` are `uint16` per-axis fixed-point codes ($N \times 3$, 15.3 MB). The `cholesky_factors_diag` and `cholesky_factors_offdiag` arrays are certified `uint8` per-column log codes ($N \times 3$ each, 7.6 MB).

Certified means that the encoder measured the covariance error the 8-bit grid causes before keeping the tier: the certificate is the 95th percentile of the covariance relative Frobenius error over an evenly spaced sample of 262,144 of the $N$ splats, and it measured $\mathrm{relF}_{p95} = 0.015 \leq 0.05$, so the uint8 tier was retained. The `amplitudes` are `uint16` geometric-log codes ($N$, 5.1 MB). That is the encoder's choice for this array because its dynamic range, $5\times10^{-4}$–$2\times10^{4}$, exceeds the 65,536:1 span a bounded-linear `uint16` grid covers (§2.4). Raw total: 35.6 MB.

- **PRECISION**: all four arrays as `float32` (30.5 MB, 30.5 MB, 30.5 MB, 10.2 MB; 101.8 MB), the archival scenario.

The third scenario (key `auto`) is a `float32`-amplitude variant of the same fit. Its centre and Cholesky codes compress to exactly the same sizes as AUTO's under every configuration, i.e. they are the same codes, so it serves only as a repeat measurement of those arrays (§2.2) and is excluded from every figure and table. Together the two reported scenarios cover the three payload classes a Luxar store contains: multi-byte integer codes, single-byte log codes, and raw IEEE-754 floats.

**Writer-chunk stores.** For the store-level measurement the reference writer writes the same fit: the float32 PRECISION store was loaded back and saved with `save_gsplats` (luxar at commit `5ec6c9852`, the software state this document describes; default Hilbert ordering; zarr format 3) under AUTO and under PRECISION. Their manifest, with the chunk geometry, on-disk bytes and versions, is deposited with this document (the writer-chunk manifest). The writer sizes each array's chunk to a 64 KiB budget computed on the float32 input and rounded down to a multiple of the spatial-index *atom* (1,170 rows here). The AUTO store therefore holds the centres in 4,680-row chunks (544 chunks of 14,040 `uint16` codes, 28,080 B of payload each), each Cholesky half in 4,680-row chunks (544 chunks of 14,040 B) and the amplitudes in 16,380-row chunks (156 chunks of 32,760 B), less than half the benchmark's 64 KiB. The PRECISION store holds the same row counts in float32. The writer's encode-time probe enabled the columnar delta+zigzag prefilter (`luxar_delta_v1`; numcodecs 0.16.5) on 1 of the four AUTO arrays (`centers`). That filter replaces each code by its difference from the previous row's code in the same column, then folds the signed difference onto the non-negative integers by interleaving positive and negative values (zigzag), so that small changes between neighbouring rows become small codes. It is defined for `uint8`/`uint16` codes only, so the float32 arrays are never candidates. The AUTO tiers and grids of this store are those listed above.

#### 2.2 Compression protocol

Compression is measured *per chunk*: Luxar stores target 64 KiB chunks (`TARGET_CHUNK_BYTES`) and zarr compresses each chunk independently, so that is the unit a store pays. Throughout this document the *compression ratio* of an array is its raw code bytes, at the stored dtype, divided by the compressed bytes of its chunks. This is a per-array codec ratio and not the compression ratio of Table 1 of the main text, which divides the bytes of the source volume at its stored dtype by the bytes of the whole archive; the two share the stored-dtype basis but have different numerators (parameter codes against source voxels) and denominators (one array's chunks against the whole store), so they are not comparable in magnitude. Each code array is split into row-aligned chunks of $\lfloor 65536/(\text{itemsize} \times \text{cols}) \rfloor$ rows (32,766 `uint16` elements per full centre chunk, 65,535 `uint8` elements per Cholesky chunk, 32,768 `uint16` amplitudes). Every configuration compresses and decompresses *all* chunks. Sizes are summed, and throughput is computed from the median wall time of 3 encode and 7 decode repetitions over the whole chunk sequence. Every configuration is verified by exact round-trip comparison on the first chunk. This 64 KiB run is a controlled re-chunked benchmark of the codes: the codec-selection experiment, on which the figures and the per-array comparison of §3.5 rest. The same 164 configurations were then run a second time on the writer-chunk stores (§2.1): every zarr chunk of each array is taken from disk exactly as the store's compressor saw it (the writer's chunk shape, the edge chunk padded to the full shape as zarr stores it, and the delta-filter residuals in place of the codes where the writer enabled the filter). Each such payload is verified before the sweep (re-encoding it with the store's own compressor reproduces the on-disk chunk size for every chunk, and inverting the filter chain reproduces the decoded codes). That size check is also the evidence that the recorded shuffle executed. Under zarr format 3 a Blosc filter whose element size is not forwarded records a shuffle it never applied, and the store's chunks would then match the unshuffled sweep. They match the byte-shuffled one instead. The cleanest case is the amplitude array, which carries no prefilter: its chunk files total 4,559,336 B, exactly the byte-shuffled level-9 sweep, where the unshuffled sweep gives 4,205,393 B. On the delta-filtered centre residuals the on-disk total likewise equals the byte-shuffled sweep (1.64× against 1.43× unshuffled, level 9). Every filter claim below is therefore a claim about bytes on disk and is independent of the recorded metadata. The shipped rule's compressed total on this chunk set therefore *is* the store's chunk-file total, and every claim below about the shipped rule and the recommended change is made on it (§3.5, Table 4). The prefiltered centres were additionally swept as plain codes at the same chunk geometry (scenario key `shipped_nodelta`) to separate the filter's contribution from the chunk geometry's. Neither run times the inverse delta filter, which the viewer applies after the codec.

The configuration matrix covers the Blosc meta-codec family [Alted, 2010] with internal codecs `zstd`, `lz4`, `lz4hc`, `zlib`, `blosclz`; compression levels $\{1, 3, 5, 7, 9\}$; the three filter settings (no shuffle, byte shuffle, bit shuffle [Masui et al., 2015]); and two block sizes (Blosc's automatic choice and a pinned 64 KiB). Outside Blosc it adds standalone codecs: raw `zstd` at levels $\{1, 3, 7, 12, 19\}$ (also, at levels $\{3, 12, 19\}$, with an explicit byte-shuffle prefilter), `zlib` $\{1, 6, 9\}$ (Python's `zlib.compress`, a zlib-wrapped stream, not gzip format; the results files label it `gzip`), `bz2` (9), and `lzma` $\{1, 6\}$: 164 configurations per array, 656 verified measurements per scenario, 1,968 over the three scenarios. A configuration is written throughout as *codec* l*level*/*filter*/*blocks*, for example `blosc-zstd` l3/bit/auto for Blosc with internal zstd at level 3, bit shuffle and automatic block size; the filter field is `byte`, `bit` or "–" for none, and the blocks field is `auto` or `64KiB` for the pinned 64 KiB block.

Blosc runs *single-threaded* throughout, for comparability with the single-threaded decoders in the browser scenario. Native multi-threaded decode is strictly faster than the numbers reported here. Hardware: the GPU workstation (Intel Core i9-9900K, 8 cores / 16 threads), Python 3.12.3 with `numcodecs` 0.16.5 and its bundled C-Blosc. The sweep ran with other background jobs paused, but a CI runner was active for part of it (the sweep pinned to one core; its load average was not recorded). The writer-chunk run (`numcodecs` 0.16.5, C-Blosc 1.21.7.dev, `zarr` 3.3.0) ran on the same host while other jobs were active at lower scheduling priority (1-minute load average 19–30 during the native sweep, 16–20 during the browser run, on 16 threads), so its throughputs are single-run medians under load and its byte counts are the deterministic part. Compression ratios are deterministic. Decode throughput is not. A repeat sweep re-measured the same centre and Cholesky codes as AUTO under every configuration (492 pairs with identical compressed sizes), and the paired throughputs differ by a median of 6.5%, by more than 41% in one pair in ten, and by up to 1.94×. The throughput figures below are therefore single-run medians on a shared host, and only the cross-family comparisons (factors of several) survive this variability. Close comparisons between configurations are not. The provenance the results files do not record is listed in §4.5.

#### 2.3 In-browser (wasm) decode protocol

The browser scenario decodes *the same compressed chunk blobs* through the codec classes the Luxar viewer's zarr reader (`zarrita`) dispatches to: the `numcodecs.js` registry (v0.3.2), in which `Blosc` and `Zstd` are WebAssembly builds of the C libraries while `Zlib` is a pure-JavaScript implementation (`fflate`). The browser stack is therefore mixed JS/wasm, and the standalone `zlib` family measures a JavaScript decoder, not a wasm one (`blosc-zlib`, inside the Blosc wasm build, is the wasm path for zlib). The benchmark page is bundled from the viewer's own pinned dependencies (esbuild), served locally, and driven in headless Chromium (Playwright's Chromium build, one core) on the same workstation as the native sweep; the writer-chunk run used Chromium 153.0.8010.12 under Playwright 1.63.0 (§4.5 for the 64 KiB run). In the 64 KiB run, for each configuration the first 48 chunks ($\approx 3$ MB raw; a contiguous prefix of the Hilbert-ordered array, whereas the native sweep times every chunk) are decoded; in the writer-chunk run *every* chunk of every configuration is decoded, the same chunk set the native sweep times. Timing is the median of 5 repetitions after a 4-chunk warm-up: 1,524 configurations over the three scenarios of the 64 KiB run (127 per array) and 1,016 over the two writer-chunk scenarios. The browser subset is Blosc levels $\{1, 3, 5, 9\}$ (level 7 omitted), standalone `zstd` at all five levels and `zlib` at $\{6, 9\}$; `bz2`, `lzma` and the external byte-shuffle prefilter are excluded because the viewer cannot decode them: deployability is a hard constraint. A missing browser cell in the tables below means the configuration is outside this subset. In the 64 KiB run, native and browser numbers come from the same machine and the same blobs but from different workloads: the browser decodes 48 of the 233 centre chunks (21%), of the 78 amplitude chunks (62%) and of the 117 chunks of each Cholesky array (41%), a spatial prefix whose compressibility need not match the whole array's. Its browser/native ratios are therefore ratios of two workloads and are not quoted as penalty factors. The writer-chunk run removes that mismatch: native and browser decode the identical chunk set (every chunk of 544 centre, 156 amplitude and 544 Cholesky chunks per array), with 7 native and 5 browser repetitions, and every browser penalty quoted in this document is that like-for-like ratio

(§3.4, Fig. 7). Both runs share the throughput caveat of §2.2: single-run medians on a shared host, so only family-level factors are claimed.

##### 2.4 Quantisation-fidelity protocol

For each of the 17 benchmark volumes of Supplementary Document 2 with a sweep scored on *held-out voxels* and two requested splat counts $K$ (the fixed 32K reference budget and the volume's cross-validation optimum $K^*$ as Supplementary Document 2 reports it; the two coincide on four volumes, giving 30 cells), a fit is produced in `float32` with the Supplementary Document 2 recipe (seed count $K$, random seed $K$, one fit per cell) and rendered. This in-memory render is the reference. For the two volumes with a *signal-limited curve* (Tribolium embryo nuclei, Mouse heart nuclei) the held-out curve is still rising at the largest tested count, so $K^*$ is the 2048K endpoint of the extended sweep, and that is the count fitted here. Every operating point Supplementary Document 2 reports is measured in this document. $K$ is the requested seed count; the retained count $N$ differs from $K$ (the seeder may return slightly more or fewer than $K$ seeds, and the post-fit *cull (retention)* at a retention of 0.999 removes a further fraction; Table 1 lists both) and is the denominator of every B/splat figure. The fit is then written as a plain `.gsplats.zarr` directory under the `PRECISION` mode (`float32` everywhere) and under the shipped `AUTO` mode, with `ordering="none"` so that the splat order is preserved and parameters can be compared splat by splat (a deployed store is Hilbert-ordered, so these are experimental order-preserved stores, not deployment stores). Each store is measured on disk (B/splat, including zarr metadata), loaded back and rendered. We report the *PSNR* of each render against the original volume,

$$\mathrm{PSNR} = 10 \log_{10} \frac{1}{\mathrm{MSE}} \ \mathrm{dB}, \tag{1}$$

with peak 1 because every volume is the $[0, 1]$-normalised copy held by the benchmark registry (the shared table of the 17 volumes, their sources and their normalisation that all supplementary documents draw on) and the mean squared error taken over all voxels with no held-out mask; a difference of 10 dB is one order of magnitude in MSE. We also report the PSNR of the quantised render against the float32 render (the quantisation error alone) and three per-splat distortions: the RMS centre displacement in voxels, the 95th percentile of the relative Frobenius error of the covariance, and the 95th percentile of the relative amplitude error (relative denominators floored at $10^{-12}$).

Under `AUTO` the amplitude array's grid is chosen from its dynamic range $\max / \min_{>0}$: bounded-linear `uint8` up to 256:1, bounded-linear `uint16` up to 65,536:1, and geometric-log `uint16` beyond that (the case of the `t252` store of §2.1); Cholesky factors are certified `uint8` (per-column log grids, escalating to `uint16` and then `float32` only if the encode-time covariance certificate fails); centres are per-axis fixed-point `uint16`.

The `MEMORY` mode, the writer's smallest tier set, was run on the same 30 cells as a control. It writes the Cholesky factors as `uint8` without the certificate and, above the 65,536:1 amplitude range, geometric-log amplitudes as `uint8`; below that range it selects the grids `AUTO` selects. On these cells it gives identical decoded parameters and metrics, not identical bytes: all 8 recorded reconstruction and decoded-parameter metrics (both PSNRs, the *SSIM*, the RMS centre displacement and the covariance and amplitude error percentiles) agree with `AUTO` to the recorded precision on every cell. Two differences remain. The recorded maximum absolute render difference differs on 17 cells, by at most $2 \times 10^{-5}$ relative, the extremum of a float32 render (the decoded parameters were not compared bit for bit). And every `MEMORY` store is 676–800 B smaller on disk, consistent with the two modes selecting the same tiers on these arrays and differing in metadata only. The `uint8` tier of `gsplat reencode -e memory` is a separate path outside this study.

**Deposited record.** The record comprises the native and in-browser benchmark tables (one record per scenario, array and configuration, for the 64 KiB run and for the writer-chunk run), the writer-chunk manifest (chunk geometry, on-disk bytes, per-array encoding attributes and library versions) and the per-cell quantisation-fidelity table. Every table and every numeric value quoted in this document is regenerated from those files when the document is built. The fidelity table holds 37 cells, of which the 30 at the two counts the study defines per volume are reported. The other seven (Cells3D membranes at 64K; C. elegans embryo nuclei at 16K; Tribolium embryo nuclei at 512K; Mouse heart nuclei at 512K and 1024K; Zebrafish embryo nuclei at 512K; Drosophila embryo nuclei at 512K) are counts that rule does not select; their largest loss is 0.196 dB (Tribolium embryo nuclei at 512K). What the record does not contain is collected in §4.5.

### 3 Results

#### 3.1 What quantisation costs in reconstruction

Table 1 gives the 30 cells and Fig. 3 plots them. The shipped encoding lowers the PSNR against the original volume by between 0.000 and 0.679 dB (median 0.012 dB); the largest loss, on Tribolium embryo nuclei at 2048K (56.7 dB in float32), is a 16.9% increase of the MSE against the original. The next largest loss among the 30 reported cells is 0.167 dB, so that cell stands apart from the rest. (The highest-fidelity fit in the table is Neuromast nuclei at 64K, 66.8 dB, so the size of the loss does not track the fit's PSNR.) Measured directly, the PSNR of the quantised render relative to the float32 render is 59–88 dB, which is 7.7–43.1 dB above each fit's own PSNR against the original, i.e. the quantisation error is 0.8–4.3 orders of magnitude (in MSE) below the approximation error the splat count sets. The smallest margin is on Tribolium embryo nuclei at 2048K, the cell with the largest loss: its fit error is low enough that the quantisation error sits only 7.7 dB below it. If the two errors add as uncorrelated terms, an error term 7.7 dB below the fit error raises the total MSE by 16.9%, which is exactly the 0.679 dB loss measured; the reconciliation rests on that independence assumption. The largest margin is on Fly brain neurons at 64K. Per splat, centres move by 0.002–0.006 voxels RMS (the `uint16` grid over each axis extent), the covariance changes by 0.7–2.3% in relative Frobenius norm at the 95th percentile (the certified `uint8` log tier), and the amplitude by 0.1–19.6% at the 95th percentile. The amplitude tail (*C. elegans* embryo nuclei, OpenCell-LMNB1 nuclear lamina, *Tribolium* at 2048K) is too large for a geometric-log grid, whose relative step is uniform across the range and is about 0.27% at 16 bits even over the full `float32` range (a maximum relative rounding error of 0.13%; Fig. 2). It is the signature of

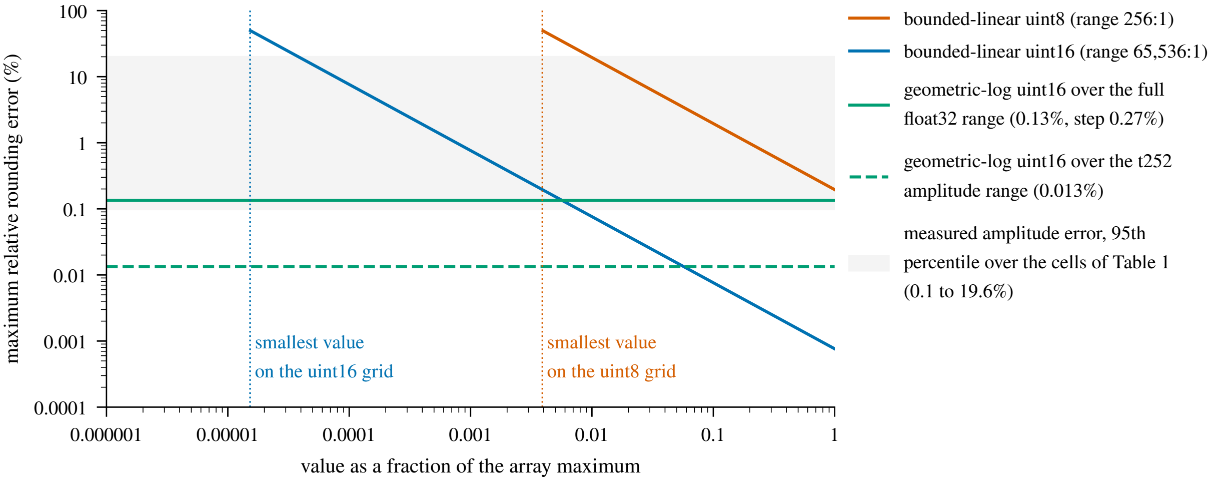


Figure 2: Maximum relative rounding error of the three amplitude grids, from their definitions in §4, against the value as a fraction of the array maximum. A bounded-linear grid has a uniform absolute step, so its relative error grows as the value shrinks: the `uint8` grid (drawn for the widest range it is selected for, 256:1) and the `uint16` grid (drawn for 65,536:1) both reach 50% at their smallest representable value. The geometric-log `uint16` grid has a uniform relative step, 0.27% over the full float32 range (maximum error 0.13%) and 0.027% over the range of the `t252` amplitude array (maximum error 0.013%). The grey band is the range of the measured 95th-percentile amplitude error over the cells of Table 1; the cells at its upper end are consistent with the bounded-linear `uint16` grid (§3.1).

the bounded-linear `uint16` grid the encoder selects when an array's dynamic range fits within 65,536:1 (§2.4), on which the smallest amplitudes carry the largest relative error. This attribution rests on the selection rule and on the error magnitude alone (§4.5). Whatever the mechanism, its effect on the image is bounded by the measured PSNR of the quantised render relative to the float32 render for those cells (Table 1, "vs. f32"). The float32 stores compress to 30–37 B/splat under the same codec policy (§3.5, §3.6). The quantised stores take 8.8–12.7 B/splat including metadata (median 11.3 B/splat over the 30 cells), 2.9–4.0× less per cell. These are order-preserved experimental stores of fresh fits: they are not Hilbert-ordered, they are not the deployed stores whose sizes the manuscript's compression ratios use, and their B/splat is not monotonic in $K$ (Table 1), so no large-$N$ limit is inferred from them. Within the tested cells the quantisation loss is at most 0.679 dB at every operating point Supplementary Document 2 reports, the two 2048K endpoints included. Quantisation is where the on-disk gain over `float32` comes from. The codec sweep that follows operates on the quantised codes.

##### 3.2 Rate versus decode speed

Figure 4 maps every configuration for the AUTO scenario. Three regimes structure all four panels. The `blosclz`/`lz4` family occupies the fast, shallow end (near-1.0 ratios at several GB/s single-threaded); `zstd` and `zlib` configurations form the knee of the Pareto front, the set of configurations that no other configuration beats on ratio and decode speed at once, at around a gigabyte per second; `lzma` and `bz2` extend ratios by only 0.02–0.14 beyond the best deployable configuration (and by nothing on the centres) while decoding at ≈24–31 MB/s (28–30 MB/s at

Table 1: Quantisation fidelity of the shipped splat encoding on 17 benchmark volumes at 32K and at the cross-validation optimum $K^*$ of Supplementary Document 2; for the two signal-limited volumes (Tribolium embryo nuclei, Mouse heart nuclei) $K^*$ is the 2048K endpoint of the extended sweep. $K$: requested seed count; $N$: splats retained after culling (the B/splat denominator). "float32": PSNR of the in-memory fit against the original (peak 1, all voxels, one fit per cell); "shipped": PSNR after writing under the shipped `AUTO` tiers and loading back; "vs. f32": PSNR of the quantised render relative to the float32 render. B/splat is measured on disk for a plain, order-preserved (`ordering="none"`) `.gsplats.zarr` directory under the shipped codec policy, metadata included. Distortions are the RMS centre displacement and the 95th-percentile relative covariance (Frobenius) and amplitude errors.

| | | | PSNR vs. original (dB) | | | vs. f32 | B/splat | | per-splat distortion | | |
|---|---|---|---|---|---|---|---|---|---|---|---|
| Dataset | $K$ | $N$ | float32 | shipped | $\Delta$ | (dB) | f32 | shipped | centre (vox) | cov. (%) | amp. (%) |
| OpenCell-MAP4 nuclei | 16K | 16,307 | 28.39 | 28.39 | -0.002 | 61.72 | 36.8 | 12.5 | 0.0037 | 1.0 | 5.6 |
| OpenCell-MAP4 nuclei | 32K | 32,314 | 28.61 | 28.61 | -0.002 | 61.65 | 36.0 | 11.5 | 0.0037 | 1.0 | 6.4 |
| OpenCell-MAP4 microtubules | 32K | 32,364 | 25.21 | 25.21 | -0.002 | 60.37 | 36.0 | 11.8 | 0.0037 | 1.0 | 5.9 |
| OpenCell-MAP4 microtubules | 64K | 62,156 | 25.68 | 25.68 | -0.002 | 59.02 | 35.9 | 11.2 | 0.0037 | 1.3 | 8.9 |
| OpenCell-LMNB1 nuclei | 32K | 31,969 | 25.60 | 25.60 | -0.001 | 61.98 | 36.3 | 11.9 | 0.0038 | 1.0 | 4.7 |
| OpenCell-LMNB1 nuclear lamina | 32K | 31,071 | 34.14 | 34.13 | -0.013 | 61.19 | 36.4 | 11.7 | 0.0038 | 1.2 | 17.6 |
| OpenCell-LMNB1 nuclear lamina | 64K | 62,078 | 34.58 | 34.56 | -0.014 | 61.28 | 36.1 | 11.2 | 0.0037 | 1.2 | 19.6 |
| Kidney nuclei | 32K | 31,658 | 34.79 | 34.78 | -0.013 | 59.95 | 36.4 | 12.2 | 0.0032 | 1.0 | 3.3 |
| Kidney nuclei | 64K | 63,038 | 38.16 | 38.13 | -0.026 | 60.36 | 36.2 | 11.7 | 0.0032 | 0.9 | 4.5 |
| Kidney actin | 32K | 31,341 | 31.13 | 31.12 | -0.007 | 59.18 | 36.2 | 12.2 | 0.0032 | 1.1 | 3.2 |
| Kidney actin | 128K | 124,298 | 39.12 | 39.07 | -0.049 | 58.80 | 35.7 | 11.3 | 0.0032 | 0.9 | 9.0 |
| Cells3D nuclei | 32K | 31,878 | 36.78 | 36.78 | -0.007 | 64.63 | 36.1 | 11.9 | 0.0016 | 1.0 | 2.2 |
| Cells3D membranes | 32K | 31,835 | 49.91 | 49.82 | -0.089 | 66.91 | 36.2 | 11.9 | 0.0016 | 1.0 | 5.0 |
| Blastocyst nuclear lamina | 32K | 28,154 | 39.18 | 39.18 | -0.002 | 75.28 | 36.2 | 12.6 | 0.0020 | 1.3 | 0.3 |
| C. elegans embryo nuclei | 8K | 7,947 | 39.29 | 39.27 | -0.017 | 63.94 | 37.1 | 12.7 | 0.0032 | 1.2 | 14.0 |
| C. elegans embryo nuclei | 32K | 31,816 | 39.36 | 39.33 | -0.033 | 60.77 | 35.8 | 10.9 | 0.0032 | 0.7 | 19.1 |
| Fly brain neurons | 32K | 29,622 | 44.59 | 44.59 | -0.001 | 83.84 | 35.9 | 11.6 | 0.0035 | 1.9 | 0.3 |
| Fly brain neurons | 64K | 60,739 | 44.87 | 44.87 | +0.000 | 88.00 | 35.6 | 10.6 | 0.0035 | 1.1 | 0.3 |
| Tribolium embryo nuclei | 32K | 31,767 | 51.45 | 51.28 | -0.167 | 65.59 | 36.0 | 11.3 | 0.0061 | 1.7 | 6.2 |
| Tribolium embryo nuclei | 2048K | 2,039,831 | 56.67 | 55.99 | -0.679 | 64.39 | 35.5 | 8.8 | 0.0061 | 1.1 | 11.0 |
| Mouse heart nuclei | 32K | 30,176 | 32.07 | 32.06 | -0.003 | 65.30 | 32.7 | 10.7 | 0.0049 | 2.1 | 4.4 |
| Mouse heart nuclei | 2048K | 1,452,304 | 47.70 | 47.69 | -0.008 | 75.30 | 35.7 | 11.1 | 0.0049 | 0.9 | 0.1 |
| Zebrafish embryo nuclei | 32K | 29,371 | 32.47 | 32.47 | -0.001 | 65.37 | 30.8 | 10.4 | 0.0035 | 1.8 | 3.5 |
| Zebrafish embryo nuclei | 256K | 190,172 | 39.70 | 39.69 | -0.013 | 66.25 | 33.9 | 10.0 | 0.0036 | 1.7 | 5.4 |
| Drosophila embryo nuclei | 32K | 30,887 | 30.47 | 30.46 | -0.004 | 61.86 | 33.5 | 11.3 | 0.0060 | 1.8 | 2.0 |
| Drosophila embryo nuclei | 256K | 228,929 | 34.84 | 34.83 | -0.011 | 61.04 | 34.6 | 10.6 | 0.0060 | 2.3 | 4.4 |
| Neuromast membranes | 32K | 31,706 | 52.67 | 52.65 | -0.018 | 75.39 | 30.6 | 9.5 | 0.0021 | 1.7 | 3.8 |
| Neuromast membranes | 128K | 74,732 | 57.17 | 57.13 | -0.040 | 77.65 | 31.1 | 10.5 | 0.0028 | 1.5 | 0.1 |
| Neuromast nuclei | 32K | 30,481 | 66.68 | 66.52 | -0.159 | 81.56 | 30.5 | 10.3 | 0.0022 | 1.3 | 0.1 |
| Neuromast nuclei | 64K | 32,783 | 66.79 | 66.66 | -0.133 | 81.34 | 29.9 | 10.0 | 0.0020 | 1.4 | 0.2 |

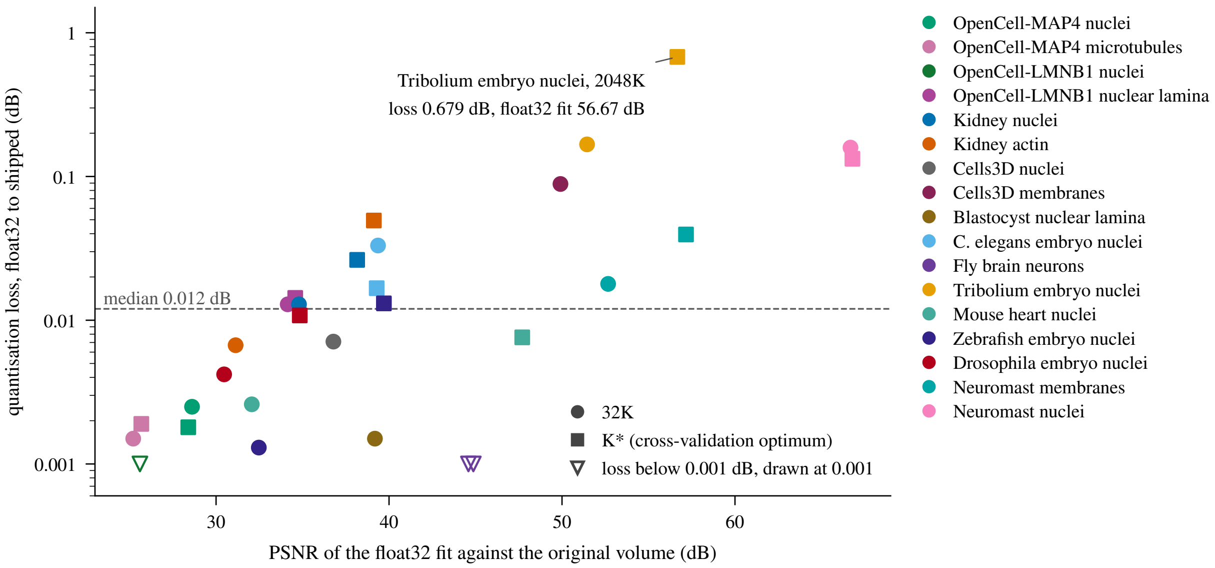


Figure 3: Table 1 as a scatter: the quantisation loss (the float32 PSNR minus the shipped PSNR against the original volume, log scale) against the PSNR of the float32 fit, one marker per cell, coloured by volume; circles are the 32K cells and squares the $K^*$ cells. The three cells with a loss below 0.001 dB are drawn at that floor as open triangles. The dashed line is the median loss (0.012 dB). The *Tribolium* 2048K cell stands apart at 0.679 dB; the size of the loss does not track the fit's PSNR.

the per-array optima), more than an order of magnitude slower, and not decodable by the viewer at all.

The star marks the uniform baseline (`blosc-zstd` level 3, bit shuffle, automatic blocks), one codec configuration applied to every array. The shipped writer uses the width-keyed level-9 rule of §3.5, whose configurations appear as ordinary points. The baseline's position differs sharply by array class. On the uint8 Cholesky arrays it sits on the front (1.39× and 1.42× at ∼1 GB/s), and the best deployable configurations improve the ratio by at most 3.0% on the unrounded ratios (the two-decimal entries of Table 2, 1.39× against 1.44×, give 3.6%). On the uint16 centres, however, it achieves only 1.06×, far below the 1.51× front. The uint16 geometric-log amplitudes are the least compressible array of all: the baseline reaches 1.10× against a 1.22× deployable front (1.34× with `lzma`). The next subsection explains the centres; the amplitudes are high-entropy codes (a log-companded scalar with little row-to-row structure for the codec's match finder, the encoder stage that searches earlier bytes for repeated strings, to use) on which both shuffle filters *cost* ratio (1.11–1.12× at level 9 against 1.22× unshuffled), so plain high-level `zstd` is the whole answer there.

##### 3.3 How filter, level and chunk size interact

Figure 6a isolates the uint16 anomaly. At level 1, bit shuffle behaves as theory suggests (1.28× vs. 1.07× unshuffled). From level 3 upward the bit-shuffled ratio equals the unshuffled value (1.06×), with equal compressed byte counts at every level. The mechanism is an alignment rule in C-Blosc's filter stage, not an entropy-coding effect: the bit-shuffle kernel transposes a block only when the block's element count is a multiple of eight and otherwise copies the block through

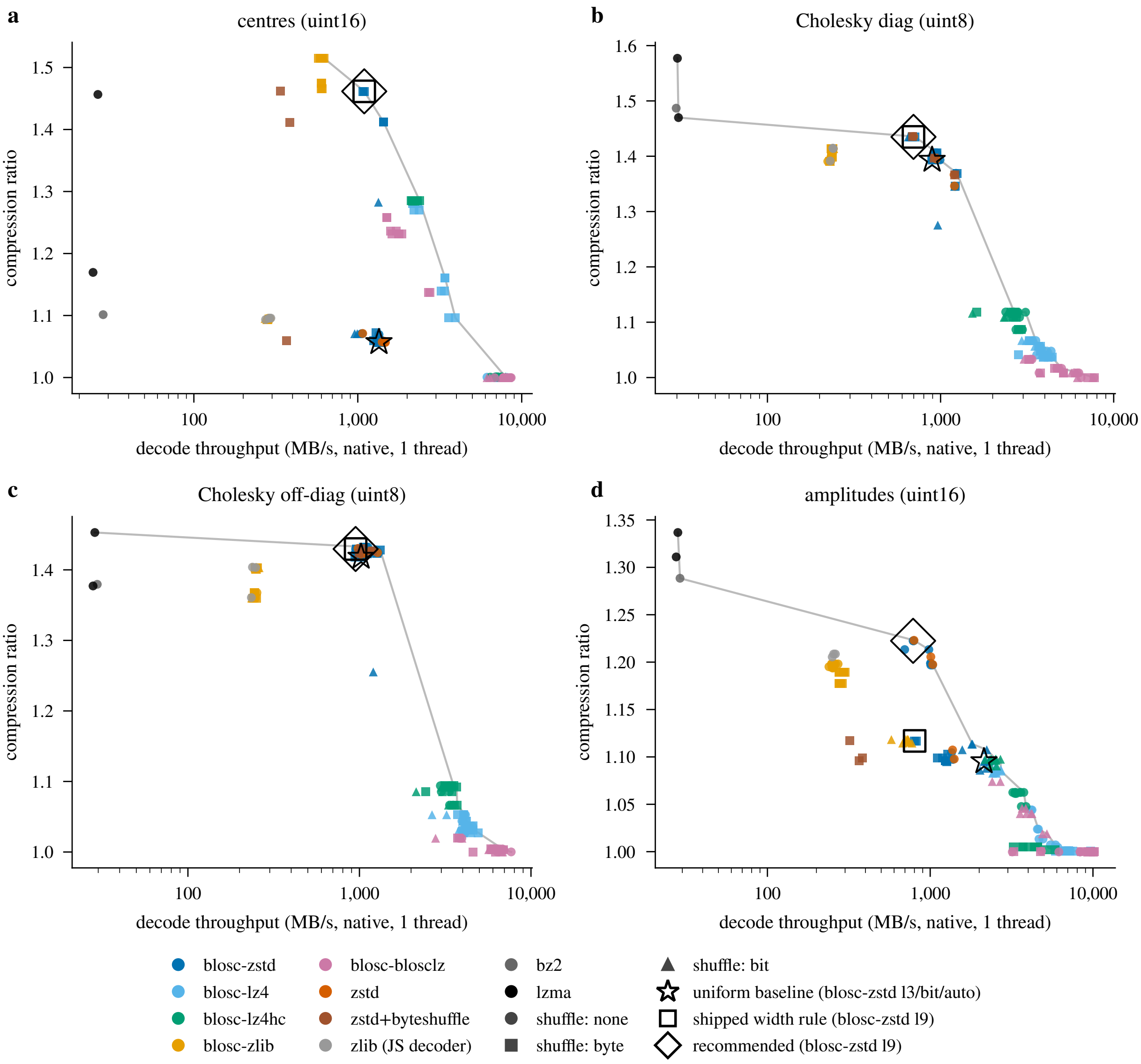


Figure 4: Compression ratio versus single-threaded native decode throughput for every configuration, AUTO scenario (the code arrays as the reference writer encodes them), re-chunked to 64 KiB: (a) centres, (b) Cholesky diagonal, (c) Cholesky off-diagonal, (d) amplitudes. Colour = codec family ("zlib (JS decoder)" is the standalone zlib stream; natively it is Python's `zlib`); marker = filter; grey line = Pareto front; star = the uniform baseline (`blosc-zstd` l3/bit/auto); square = the shipped width rule's configuration for the array (`blosc-zstd` l9, byte shuffle for the multi-byte integer codes, none otherwise); diamond = the recommended configuration (byte shuffle for the centres only, §3.5). On every array but the amplitudes the square and the diamond coincide.

unchanged (C-Blosc's block compressor; the chunk header still records the requested filter, so the frames are not identical even where the payloads are). A full benchmark centre chunk holds 32,766 `uint16` elements, not a multiple of eight, so whenever Blosc treats the chunk as a single block the filter is skipped (Fig. 5 draws the four chunk geometries measured here against that rule). Blosc's automatic block size grows with the level, and the data behave as that predicts: pinning the block size to the whole 64 KiB chunk (dashed) gives the unshuffled ratio already at level 1 (1.06×), whereas the automatic choice at level 1 splits the chunk into smaller blocks and the filter engages. The `uint16` amplitude chunks are the control: they hold 32,768 elements, a multiple of eight, and keep a distinct bit-shuffled size at every level (1.11× vs. 1.22× unshuffled at level 9). The writer's own chunks are the complementary control: a writer centre chunk holds 14,040 `uint16` codes, a multiple of eight, and there bit shuffle executes at every level and is the best filter for the plain codes (1.68× at level 9 vs. 1.46× byte-shuffled and 1.08× unshuffled; on the writer's delta-filtered residuals the two filters converge, 1.64× vs. 1.64×). A writer amplitude chunk, by contrast, holds 16,380 codes, not a multiple of eight, and its level-9 bit-shuffled size equals the unshuffled one (1.21× vs. 1.21×). The skip follows the element count, not the array. On the `uint8` Cholesky chunks of the writer (14,040 elements, so the filter executes) bit shuffle costs 14% of ratio at level 9 (1.23× vs. 1.44×), the level-1 behaviour of the 64 KiB run at every level. Byte shuffle has no such alignment rule and shows a different pattern on the centres: inert through level 5, then engaging at levels 7–9 to reach 1.41–1.46×, the best `zstd`-family results for this array. We did not isolate the cause of that level threshold. On the uint8 panel (Fig. 6b) byte shuffle is the identity and bit shuffle converges to it above level 1: all six filter/block variants coincide to within 0.1%, since a full Cholesky chunk holds 65,535 elements, again not a multiple of eight, so the same skip applies. Bit shuffle diverges downward at level 1, however (a 7–12% loss), where the filter does execute, so single-byte codes are only *near* filter-invariant, and only above level 1 (§4).

Two practical rules follow. First, the filter must be chosen *per element width and per code type* instead of globally: byte shuffle (with level $\geq 7$) for multi-byte fixed-point codes, no filter for single-byte codes and for the log-companded amplitude codes (the float32 arrays are a separate case, §3.6). Second, the chunk size, down to the element count per block, is silently part of the codec policy: conclusions from whole-array benchmarks do not transfer to chunked stores, and a writer that wants bit shuffle to execute must keep the element count per block a multiple of eight.

##### 3.4 Decode speed: levels, and the browser

Within every codec family the decode throughput depends less on the encoder level than the encode throughput does, but it is not level-independent. Going from `blosc-zstd` level 1 to level 9 costs 11–43% of native decode speed (median 27% over the twelve array/filter combinations; unshuffled centres 1347 → 1066 MB/s, Cholesky diagonal 1227 → 697 MB/s) and ≈ 24% in the browser (bit-shuffled centres: 335, 324, 254 MB/s at levels 1, 5, 9), while encode speed falls 41–69-fold over the same twelve combinations (811 → 16 MB/s on the unshuffled centres, 50×). The level is therefore predominantly, though not purely, a write-time budget decision: the read-time cost is about a quarter of the decode speed at the median, and it is part of the trade-off

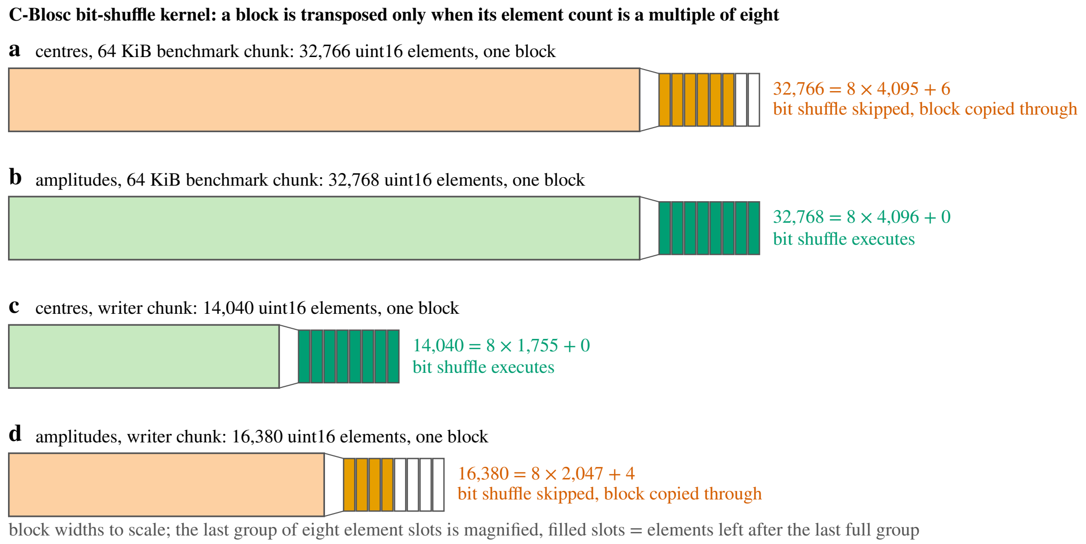


Figure 5: Schematic of the multiple-of-eight rule on the four `uint16` chunk geometries measured here (no data plotted). C-Blosc's bit-shuffle kernel transposes a block only when its element count is a multiple of eight and otherwise copies it through unchanged. Each chunk is drawn as a single block, the case from level 3 upward, where Blosc's automatic block size covers the whole chunk; the magnified group of eight slots at the end of each bar shows the remainder. The 64 KiB benchmark centre chunk (a, 32,766 elements) and the writer's amplitude chunk (d, 16,380) are copied through; the 64 KiB amplitude chunk (b, 32,768) and the writer's centre chunk (c, 14,040) are shuffled. At level 1 the automatic block size splits a 64 KiB chunk into smaller blocks and the filter engages (Fig. 6a).

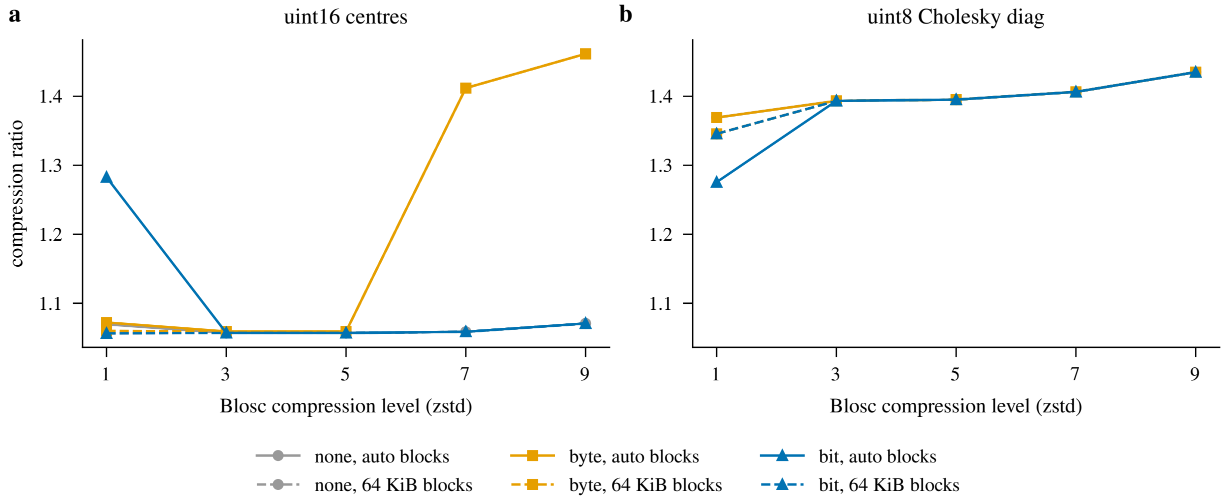


Figure 6: Compression ratio versus Blosc level (zstd) for all six filter/block-size variants (solid = Blosc's automatic block size; dashed = the block pinned to the whole 64 KiB chunk; both panels share the vertical axis). (a) uint16 centres: bit shuffle is skipped above level 1 (the single block's element count is not a multiple of eight); byte shuffle engages at level ≥ 7. (b) uint8 Cholesky diagonal: byte shuffle is the identity and bit shuffle converges to it above level 1 (it hurts at level 1, where it executes).

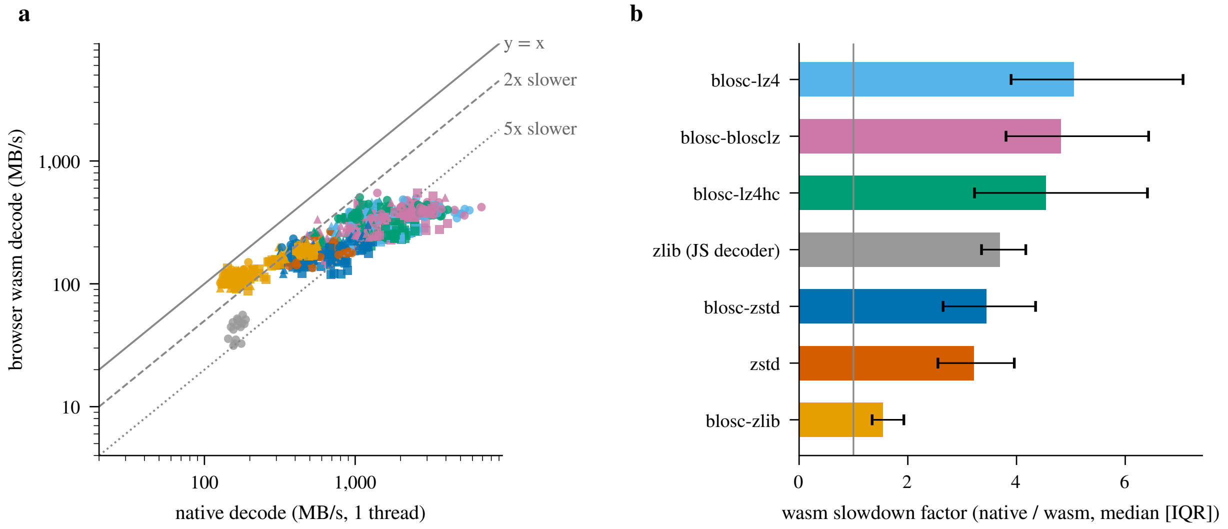


Figure 7: In-browser decode of the same compressed chunks as natively: the writer-chunk run, every chunk of every configuration on both sides (AUTO and PRECISION scenarios). (a) browser versus native throughput per configuration, both on the same workstation. (b) slowdown factor by codec family (median [IQR] over configurations): the `lz4` class loses 4.5–5.1×, `blosc-zstd` 3.4×, standalone `zstd` 3.2×, the JavaScript `zlib` decoder 3.7×, `blosc-zlib` (wasm) 1.5×.

the recommendation below accepts. Across families, the browser is *not* a scaled copy of native performance (Fig. 7). Measured on the same machine over the identical chunk set (the writer-chunk run, AUTO and PRECISION scenarios, every chunk on both sides, §2.3), the browser penalty is 1.5× for `blosc-zlib` (wasm), 3.7× for the standalone `zlib` stream (the JavaScript decoder), 3.4× for `blosc-zstd` and 3.2× for standalone `zstd`, but 4.5–5.1× for `lz4hc`, `blosclz` and `lz4` (medians over configurations; interquartile ranges in Fig. 7b). The 64 KiB run, whose browser side decoded only the 48-chunk prefix, orders the three families the same way at larger factors (AUTO and PRECISION scenarios pooled: `blosc-zlib` 1.7×, `zstd` ≈ 4.3×, the `lz4` class 8.3–10.4×). The two runs differ in chunk size, workload and host load, so only the like-for-like factors are quoted as penalties. The memory-bandwidth-bound codecs lose the most: in the browser the codecs run single-threaded with a copy across the JS/wasm heap boundary per chunk, which taxes precisely the codecs whose native decode approaches memcpy speed (whether the `numcodecs.js` 0.3.2 wasm builds use SIMD was not checked). In the browser the fastest family peaks at 0.55 GB/s regardless of configuration, so the native 2.6× median speed advantage of the `lz4` class over `zstd` (1.3 vs. 0.5 GB/s, AUTO writer chunks) shrinks to 2.0× (333 vs. 169 MB/s), while on the same chunks the best `zstd`-family ratio is 1.0 to 1.3 times the best `lz4`-class ratio, depending on the array. On the writer's chunks all recommended configurations decode at 138–215 MB/s of decoded bytes in the browser against 326–592 MB/s natively, in this warmed codec microbenchmark on the benchmark workstation. That is well above the compressed-byte rate of a typical residential connection, but the measurement times codec decode alone: it excludes the viewer's dequantisation, geometry preparation and GPU upload, and it covers no other browser or device. It bounds the codec's share of the read path, and it does not establish where the streaming bottleneck lies.

##### 3.5 Per-array policy and four-array payload

Table 2 lists, per array, the uniform baseline, the best viewer-deployable configuration, and the unconstrained best. Table 3 aggregates codec policies over the four-array payload (compressed chunk bytes of the four parameter arrays only, so unlike Table 1 it excludes metadata) at the 64 KiB geometry, and Table 4 the same policies on the writer's own chunks, the store-level numbers. Two comparisons have different denominators, and both are given. The *shipped* width rule (`zstd` level 9; byte shuffle for every multi-byte integer array, none for uint8; the compressor the reference writer resolves for these dtypes, plus the delta prefilter its probe enabled on the centres) stores the four arrays of the store as written in 24,513,722 B of chunk files: 24.5 MB, 9.63 B/splat (centres 9.31, amplitudes 4.56, Cholesky 5.30 and 5.34 MB). Compare 10.08 B/splat for the same compressor on the same chunks without the prefilter and 25.7 MB (10.09 B/splat) at the 64 KiB geometry. The writer's store is 4.5% smaller than the 64 KiB protocol's total for the same rule, and 98% of that difference is the delta prefilter on the centres (1.46 → 1.64×, 10.43 → 9.31 MB), the rest the smaller chunks. On those chunks the all-`zstd` policy recommended here (centres byte-shuffled `zstd` l9, the other three unshuffled `zstd` l9) saves 1.44% (0.14 B/splat, 354 kB on this store), all of it on the amplitude array, where byte shuffle costs ratio (1.12× against 1.21× unshuffled on the writer's 16,380-row chunks; 1.12 vs. 1.22× at 64 KiB). It also speeds that array's decode on the identical chunk set from 386 to 461 MB/s natively and from 136 to 141 MB/s in the browser (the browser difference is within the repeat variability of §2.2). Every array then decodes at $\geq$ 138 MB/s in the browser and $\geq$ 326 MB/s natively. That is the one actionable residual this measurement found: a one-line writer change (byte shuffle for *fixed-point* codes only, no filter for log-companded ones), whose size and decode effect are measured here on the writer's own chunks. The 64 KiB protocol gives 1.53% for the same change. Against the uniform baseline (`blosc-zstd` l3/bit/auto on every array; 29.9 MB, 11.77 B/splat; one codec configuration for every array), the shipped rule saves 14.3% and the recommendation 15.6%. Those figures measure the distance from a baseline the shipped rule does not use. The gain still available is the residual between the shipped rule and the recommendation. Taking the per-array optimum among all deployable codecs (which substitutes `blosc-zlib` l5/byte for the centres; natively it decodes at 598 against 1095 MB/s, in the browser, the environment this document argues from, at 242 against 244 MB/s, the same speed, and standalone `zstd` l19 for the amplitudes and the Cholesky diagonal) reaches 24.9 MB (9.78 B/splat), 16.9% below the baseline: the decision against it rests on keeping a single-codec pipeline, with decode speed playing no part, and 1.3 percentage points do not buy a second codec. Admitting non-deployable codecs adds only a further 3.0 points (24.0 MB, 9.42 B/splat; `lzma` at 28–30 MB/s decode): there is no hidden win outside the deployable set. The writer's chunks say the same (Table 4): the per-array deployable optimum reaches 9.44 B/splat (2.0% below the rule as written) and any codec 9.17 B/splat (4.8%).

##### 3.6 The PRECISION scenario, and where compression really comes from

Repeating the sweep on the float32 PRECISION store (Fig. 9) gives best deployable ratios of 1.54× (centres, zlib l9/byte/auto), 1.77× (amplitudes, zstd l19/–/auto; 1.97× with lzma l6/–/auto), 1.38× (Cholesky diagonal, zlib l9/byte/auto) and 1.16× (off-diagonal, zlib l7/byte/auto), while

Table 2: Best configurations per array (AUTO scenario, the code arrays as the reference writer encodes them). "Uniform baseline" = `blosc-zstd` l3/bit/auto, one codec configuration for every array (the shipped width rule is level 9, §3.5); "deployable" = decodable by the viewer's numcodecs.js registry; a "–" browser cell = configuration outside the browser subset (§2.3). Configuration syntax: codec level/filter/blocks; "zlib (JS)" is the standalone zlib stream decoded by the JavaScript codec. Array dtypes: "u16" = `uint16` codes, "u8" = `uint8` codes, "u16 geolog" = `uint16` geometric-log amplitude codes, the grid the encoder selects for an amplitude range beyond 65,536:1 (§2.1).

| array | role | configuration | ratio | native dec (MB/s) | wasm dec (MB/s) |
|---|---|---|---|---|---|
| centres (u16) | uniform baseline | blosc-zstd l3/bit/auto | 1.06 | 1346 | 328 |
| | best deployable | blosc-zlib l5/byte/auto | 1.51 | 598 | 242 |
| | best overall | blosc-zlib l5/byte/auto | 1.51 | 598 | 242 |
| amplitudes (u16 geolog) | uniform baseline | blosc-zstd l3/bit/auto | 1.10 | 2140 | 286 |
| | best deployable | zstd l19/–/auto | 1.22 | 795 | 204 |
| | best overall | lzma l6/–/auto | 1.34 | 28 | – |
| Chol. diag (u8) | uniform baseline | blosc-zstd l3/bit/auto | 1.39 | 893 | 194 |
| | best deployable | zstd l19/–/auto | 1.44 | 699 | 258 |
| | best overall | lzma l6/–/auto | 1.58 | 30 | – |
| Chol. off-diag (u8) | uniform baseline | blosc-zstd l3/bit/auto | 1.42 | 1022 | 300 |
| | best deployable | blosc-zstd l7/–/auto | 1.43 | 1100 | – |
| | best overall | lzma l6/–/auto | 1.45 | 29 | – |

Table 3: Four-array payload (compressed chunk bytes of the four parameter arrays, $N = 2{,}544{,}311$ splats; no metadata) under each codec policy at the 64 KiB benchmark geometry, with the size reduction relative to the uniform baseline and relative to the shipped width rule (a positive value is a smaller payload).

| | | | size reduction vs. | |
|---|---|---|---|---|
| policy (AUTO codes, four-array payload) | MB | B/splat | baseline | shipped rule |
| uniform baseline (zstd l3/bit on every array)<br>*zstd l3/bit/auto, zstd l3/bit/auto, zstd l3/bit/auto, zstd l3/bit/auto* | 29.9 | 11.77 | +0.0% | -16.7% |
| shipped width rule (zstd l9; byte shuffle for multi-byte codes)<br>*zstd l9/byte/auto, zstd l9/byte/auto, zstd l9/–/auto, zstd l9/–/auto* | 25.7 | 10.09 | +14.3% | +0.0% |
| recommended (zstd l9; byte shuffle for centres only)<br>*zstd l9/byte/auto, zstd l9/–/auto, zstd l9/–/auto, zstd l9/–/auto* | 25.3 | 9.93 | +15.6% | +1.5% |
| per-array best, viewer-deployable<br>*zlib l5/byte/auto, zstd l19/–/auto, zstd l19/–/auto, zstd l7/–/auto* | 24.9 | 9.78 | +16.9% | +3.0% |
| per-array best, any codec<br>*zlib l5/byte/auto, lzma l6/–/auto, lzma l6/–/auto, lzma l6/–/auto* | 24.0 | 9.42 | +19.9% | +6.6% |

Table 4: The same policies on the writer's own chunks (the AUTO store written by the reference writer, §2.1; every chunk as its compressor saw it, §2.2). The first row is the store as written: its total equals the store's chunk files byte for byte; a positive size reduction is a smaller payload. "Slowest array" = the lowest per-array decode throughput of the policy, native and in the browser, on the identical chunk set; "–" = a configuration outside the browser subset.

| policy (writer chunks, four-array payload) | MB | B/splat | size reduction vs. rule | slowest array (MB/s) native | slowest array (MB/s) browser |
|---|---|---|---|---|---|
| shipped width rule, as written (delta prefilter on centres)<br>*zstd l9/byte/auto, zstd l9/byte/auto, zstd l9/–/auto, zstd l9/–/auto* | 24.5 | 9.63 | +0.0% | 326 | 136 |
| recommended (byte shuffle for centres only)<br>*zstd l9/byte/auto, zstd l9/–/auto, zstd l9/–/auto, zstd l9/–/auto* | 24.2 | 9.50 | +1.4% | 326 | 138 |
| per-array best, viewer-deployable<br>*zlib l9/bit/auto, zstd l19/–/auto, zstd l19/–/auto, zstd l19/–/auto* | 24.0 | 9.44 | +2.0% | 327 | 135 |
| per-array best, any codec<br>*zlib l9/bit/auto, lzma l6/–/auto, lzma l6/–/auto, zstd l19/–/auto* | 23.3 | 9.17 | +4.8% | 21 | – |

the fast codecs sit at $\approx 1.00\times$. Three of the four float32 optima use byte shuffle, which groups the structured exponent bytes. The amplitude array is the exception: it compresses best unshuffled with a high-level standalone `zstd`, and we leave the cause of its larger ratio open (its values span more decades than the other arrays, so its exponent bytes may carry more structure, but this was not tested). The $32/23 \approx 1.4\times$ figure of §4 is a reference for independent random mantissas, not a bound: the centres and amplitudes exceed it, the off-diagonal falls well short. These float32 codec ratios are higher than the AUTO ratios of the same arrays, yet the compressed float32 stores are far larger, because the ratio applies to a $2.9\times$ larger raw payload: the PRECISION store holds the same four arrays in 101.8 MB raw where the quantised AUTO store holds them in 35.6 MB raw, and in 29.9 MB compressed even under the uniform baseline, the weakest codec policy tested. Quantisation moves the stored size by integer factors, codec selection by tens of percent (Fig. 8 lays the two levers end to end in bytes per splat). The codec is worth tuning (its read-time cost is the modest level penalty above), but it is the second of the two levers. The shipped rule's "no shuffle for floats" is not derived from this experiment (three of four float32 arrays prefer byte shuffle under `blosc-zlib`); it is an engineering simplification for the float32 fallbacks the default writer emits only when every integer tier fails its error certificate.

### 4 Theoretical Context

#### 4.1 What is left to compress after quantisation

The stored integer arrays are uniform-quantiser outputs in a companded domain. Centre coordinates use a per-axis linear grid: $u = \mathrm{round}\big((x-\ell)/(h-\ell)\cdot(2^{16}-1)\big)$ with $[\ell, h]$ the axis extent. The Cholesky columns and the geometric-log amplitudes use a grid with a *reserved zero*: exact zeros take code 0, and every nonzero entry is stored as $u = 1 + \mathrm{round}\Big((y-\ell)/(h-\ell)\cdot(2^b-2)\Big)$ of its companded value $y$, where $b$ is the code width in bits (8 or 16) and $[\ell, h]$ is the column's companded range over its nonzero entries. The companded value is $y = \ln(1+x)$ for the diagonal

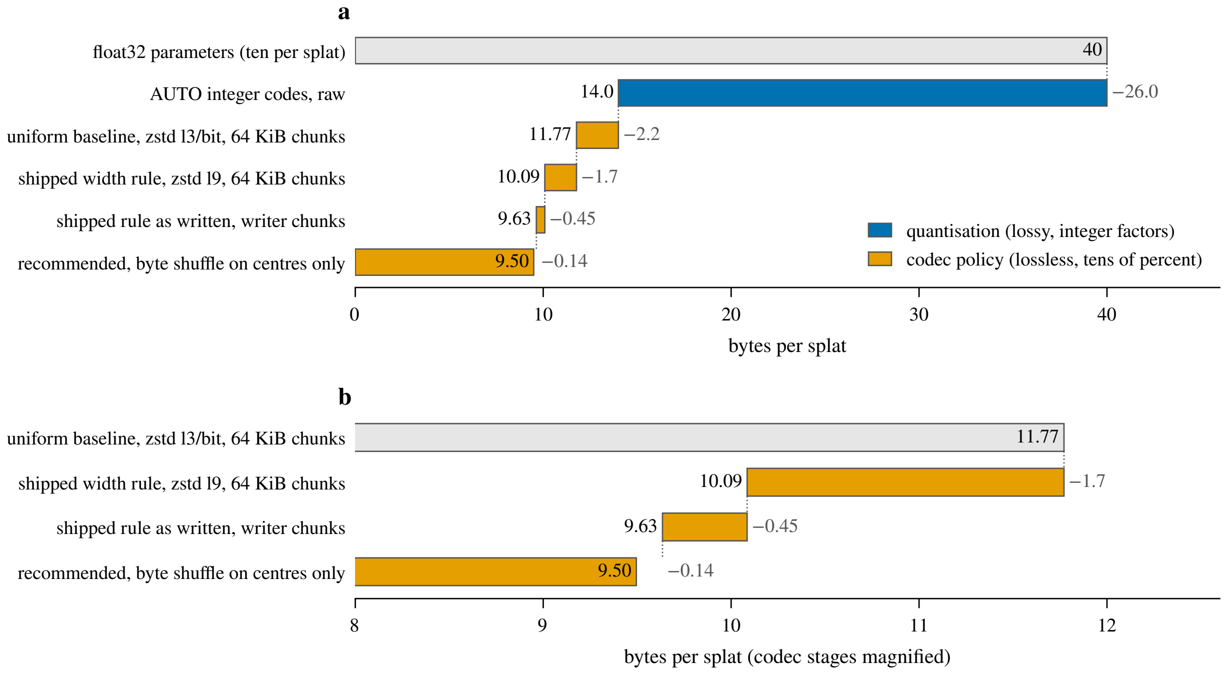


Figure 8: Bytes per splat of the `t252` fit at each stage, from the ten float32 parameters (40 B) to the recommended codec policy. (a) Blue: the quantisation tiers, which cut the raw payload from 40 to about 14 B/splat before any codec runs (35.6 MB over 2,544,311 splats). Orange: the codec policies in the order the document discusses them; the first two are measured at the 64 KiB benchmark geometry (Table 3), the last two on the writer's own chunks (Table 4), so the step between them measures the chunk geometry and the delta prefilter. (b) The codec stages on a magnified axis. Each codec bar is a row of Table 3 or Table 4; the two quantisation bars are the 40 B of ten float32 parameters and the 35.6 MB raw AUTO payload of §2.1, per splat. The step labels are differences of the unrounded byte totals.

columns, the signed $y = \operatorname{sign}(x)\ln(1+|x|)$ for the off-diagonal columns (which may be negative), and $y = \ln x$ for geometric-log amplitudes (bounded-linear amplitude grids use the centre formula with the array's $[\min, \max]$). A constant column is stored as a constant code. In the high-resolution regime the marginal entropy of such codes is $H(u) \approx h(Y) - \log_2 \Delta$ bits per symbol, where $h(Y)$ is the differential entropy of the companded source $Y$ (of which each stored $y$ is one sample) and $\Delta$ the quantiser step [Gray and Neuhoff, 1998], an approximation that assumes a fine grid and a regular density. Relative to $b$-bit storage the ideal saving is $b - H_{\text{rate}}$ bits per symbol, where the entropy rate $H_{\text{rate}}$ is at most the marginal entropy. The saving has two parts, the non-uniformity of the marginal code distribution and the gap between marginal and conditional entropy, i.e. the statistical dependence between neighbouring symbols. LZ-family codecs approach the conditional entropy of the stream given its recent past only asymptotically [Ziv and Lempel, 1977, Cover and Thomas, 2006]. A finite-window, finite-block codec such as those measured here need not attain it.

Luxar's writers enlarge the dependence part deliberately: splats are sorted along a Hilbert curve before writing [Skilling, 2004], so consecutive rows are spatial neighbours. Centre codes then change slowly between neighbours, and Cholesky codes, describing locally similar nuclei, cluster. We did not measure the neighbour-difference histograms, so this picture is illustrative. The codec's job is to convert this row-to-row coherence into bits saved. The filter's job (next

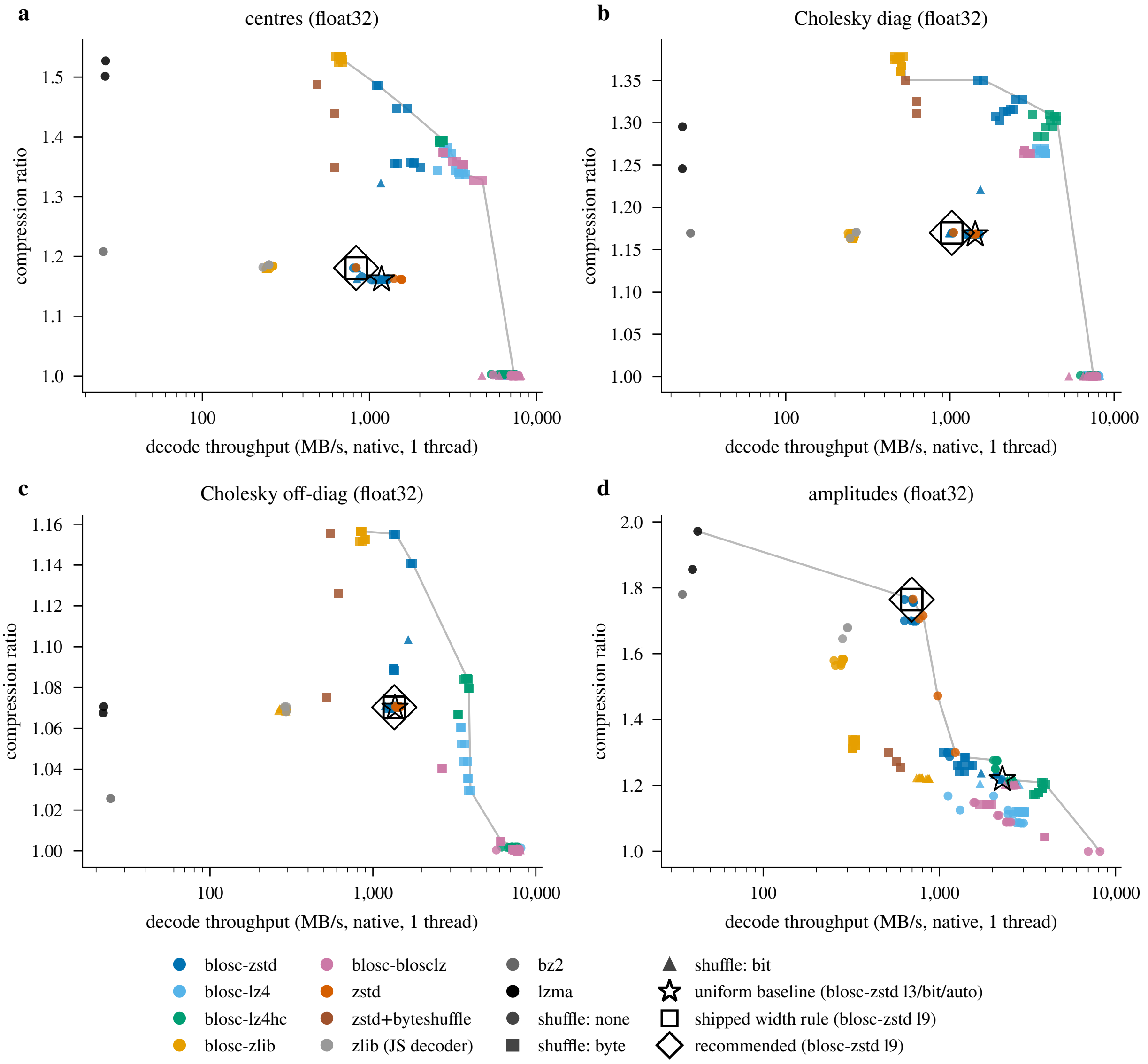


Figure 9: As Fig. 4, for the float32 PRECISION scenario: (a) centres, (b) Cholesky diagonal, (c) Cholesky off-diagonal, (d) amplitudes. Best deployable ratios per array: 1.54, 1.77, 1.38 and 1.16× (centres, amplitudes, Cholesky diagonal, off-diagonal); the fast codecs stay near 1.0×. The square (shipped width rule, `blosc-zstd` l9 unshuffled for floats) and the diamond (recommended) coincide on every float32 array, since the recommendation leaves the float32 rule unchanged.

section) is to present the coherence in a form the codec's match finder can see.

The `float32` arrays are the near-anti-case. If the mantissas of fitted continuous parameters were fully random, their $\approx 23$ bits would be irreducible while the structured sign+exponent ($1+8$ bits) compress away, giving a *reference* of $32/(23+\varepsilon) \approx 1.4\times$, where $\varepsilon$ stands for the few bits of the sign and exponent that do not compress away entirely. The reference is conditional on that independence assumption and is not a bound on actual fitted parameters. The measured PRECISION ratios scatter around it (§3.6): the centres reach 1.54× and the amplitudes 1.77× (fitted values do not fill 23 significant bits uniformly), while the wide-range off-diagonal, whose exponents vary too much to compress, reaches only 1.16×. Either way the compressed float32 stores stay far above the quantised AUTO stores in bytes: quantisation is where Luxar's on-disk compression actually comes from, and codec choice is secondary.

#### 4.2 Why bit shuffle, and when it does nothing

A `uint16` code whose neighbours differ by a few units has near-constant high bytes and volatile low bytes, but the byte stream interleaves them, so an LZ match finder sees no long runs. Byte shuffle transposes the two byte planes into contiguous blocks. Bit shuffle [Masui et al., 2015] goes further and transposes the 16 bit planes: the high planes become almost-constant runs (nearly free under LZ) and only the genuinely random low planes cost entropy (how many planes fall on each side depends on the neighbour-difference distribution, which we did not measure). Figure 10 shows the three byte orders on four example codes. Formally, shuffling is a permutation and cannot change the joint entropy, but it relocates redundancy from *across byte boundaries* (invisible to byte-granular LZ matching) to *along the stream* (exactly LZ's model). For single-byte codes *byte* shuffle has no boundary to transpose and is the identity permutation (our `uint8` byte- and no-shuffle configurations give equal compressed byte counts). *Bit* shuffle, however, still transposes the eight bit planes, a real permutation that, where it executes (level 1 in the 64 KiB sweep), *hurts* the `uint8` Cholesky arrays (a 7–12% ratio loss). On the 64 KiB chunks the filter is skipped above level 1 by the alignment rule of §3.3. On the writer's own chunks, where it executes at every level, it costs 14% of ratio at level 9. So single-byte codes want no filter: shuffling is formally a permutation, yet wherever bit-plane transposition executed, it cost ratio.

#### 4.3 Why decode speed depends only weakly on the level

zstd separates an asymmetric *search* problem from a symmetric *reconstruction* problem [Collet and Kucherawy, 2021]. The compression level controls encoder-side match-search effort (hash chains, binary trees, optimal parsing). The compressed representation keeps the same structure at every level, Huffman-coded literals (or raw/RLE literals) and sequence symbols coded with finite state entropy (FSE, a table-driven entropy coder of the asymmetric-numeral-systems family [Duda, 2013]), which the decoder replays in a single table-driven pass whose cost scales mainly with the *decoded* byte count. The same asymmetry holds for LZ4 and zlib. It is why encode speed falls 41–69-fold from level 1 to level 9 while decode speed falls 11–43% (§3.4). The residual decode cost is a measured quantity here, and it is real. For a write-once/read-many store the encoder level is therefore predominantly a write-time budget decision, *provided* the filter pipeline is not perturbed, which is exactly the failure mode quantified in §3.3 (Blosc's automatic block size at high levels leaving a single block whose element count is not a multiple of eight, on which bit shuffle is skipped).

#### 4.4 Blocking and the browser boundary

Blosc splits each chunk into cache-sized blocks compressed independently [Alted, 2010]: this bounds decoder working memory and enables (native) intra-chunk threading, at the cost of truncating the LZ window at block boundaries, a small cost when the correlation length is well below the block size, which we expect for Hilbert-ordered codes but did not measure. In the browser, Blosc

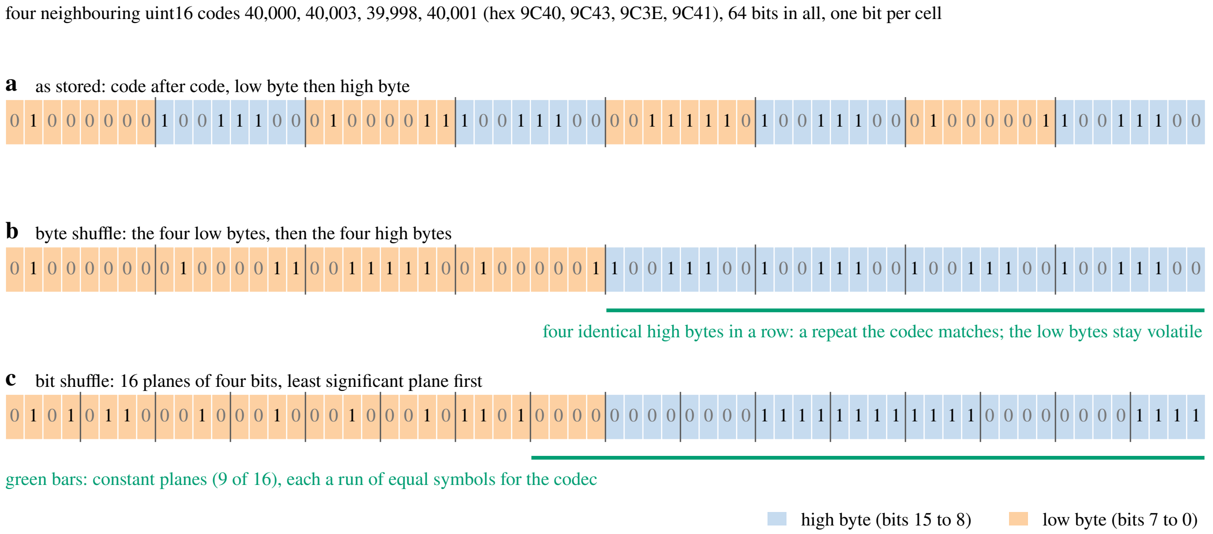


Figure 10: Schematic of the two shuffle filters on four example `uint16` codes chosen to differ by a few units, as Hilbert-sorted neighbours do (no data plotted). (a) As stored, each code contributes its low byte and then its high byte, so the near-constant high bytes are interleaved with the volatile low bytes. (b) Byte shuffle gathers the four low bytes and then the four high bytes: the four identical high bytes become one repeated string for the codec's match finder. (c) Bit shuffle gathers the 16 bit planes, least significant first (C-Blosc transposes the bytes of each code first and then the bits within each byte, so the low byte's eight planes precede the high byte's): every plane on which the four codes agree is a run of equal symbols (green), and only the low planes, now at the head of the stream, carry entropy. For single-byte codes byte shuffle is the identity and bit shuffle still transposes the eight planes (§3.3).

and zstd run as WebAssembly builds of the same C code (identical algorithms and compressed bytes, but single-threaded and with an extra copy across the JS/wasm heap boundary per chunk), while the standalone zlib codec is JavaScript. Whether the builds enable wasm SIMD was not checked. Theory therefore predicts a roughly constant-factor browser penalty, approximately uniform across levels within one codec, but *not* necessarily uniform across codecs, which is what makes the in-browser measurement worth doing instead of extrapolating.

##### 4.5 Limitations of the deposited record

The gaps in the deposited record bound what this document can claim, and none is closed by a rerun here. They fall into three groups.

**The fidelity study.** The 30 fidelity fits of §2.4 have no run manifest: their luxar commit, torch build and GPU are not recorded, so the fits are reproducible only as a recipe. The per-cell encoding attributes (which amplitude grid the encoder selected, whether the covariance certificate held) were not written into the deposited fidelity table, so the attribution of the amplitude tail to the bounded-linear grid rests on the selection rule and the error magnitude. The headline cell, Tribolium embryo nuclei at 2048K with a loss of 0.679 dB against 0.167 dB for the next reported cell, has no identified mechanism. The bounded-linear grid is consistent with it, and Table 1 gives the cell's three per-parameter distortions, but no decomposition of that cell's PSNR loss into the contributions of the centre, covariance and amplitude errors was made.

**The benchmark stores.** The three 64 KiB benchmark stores and the two writer-chunk stores are not deposited. The preparation step that concatenated the 43 content boxes and re-saved them, and the resulting stores' hashes, are recorded only from that session, so the concatenation is not reproducible from the deposited files. The luxar commit of the writer that produced the three 64 KiB stores was not recorded; the commit of the writer-chunk stores is recorded (§2.1).

**The codec sweeps.** The 64 KiB results file records the host name, the Python and `numcodecs` versions and the repeat counts, but not the C-Blosc, zstd or zlib library versions, the operating system, the core-pinning command or the host's load average during the run; the Chromium and Playwright versions of its browser run were likewise not recorded. Per-repeat times were not retained for either run, so no confidence interval accompanies any throughput, and the host carried other load during both (§2.2). The writer-chunk manifest does record its library, browser and driver versions.

### 5 Discussion

The recommended all-`zstd` compromise, under the constraint that the viewer must decode it, is not a single codec but a per-array policy within the `zstd` family. Multi-byte fixed-point codes get byte-shuffled high-level `blosc-zstd` (centres, at the 64 KiB geometry: 1.46× at 1.1 GB/s native, 244 MB/s in the browser). `blosc-zlib` l5/byte edges a further +3.6% ratio on the centres and decodes in the browser at 242 MB/s against `zstd`'s 244, the same speed (natively 598 against 1095 MB/s); it is declined in order to keep one codec in the pipeline. The uint8 codes and the uint16 amplitude codes get unshuffled high-level `zstd` (Cholesky: 1.43–1.44×; amplitudes: 1.22×). It is the policy selected for this workload (three encodings of one light-sheet fit; the 17-volume fidelity study tests quantisation, not codec choice). The amplitude exception already shows that element width alone does not determine the right filter, so transfer to other modalities should be measured, not assumed. Every recommended configuration is already decodable by the deployed viewer, and zarr arrays are self-describing, so adopting the policy is a writer-side change with no reader migration. The shipped policy is a width-aware per-dtype rule (multi-byte integer codes → `zstd` level 9 with byte shuffle; `uint8` codes and floats → `zstd` level 9 unshuffled), applied to `uint16` amplitude codes that are geometric-log for wide-range arrays such as this one and bounded-linear for narrow-range arrays (§2.4). The sweep reported here is on the codes that rule compresses. On the `t252` store as that writer writes it (its own atom-aligned chunks, the delta prefilter it enabled on the centres), the four arrays occupy 24.5 MB (9.63 B/splat) of chunk files, and the sweep on those chunks reproduces that total byte for byte (§2.2, Table 4). The 64 KiB protocol's 25.7 MB for the same rule is 4.5% larger, mostly because it omits the prefilter. The one refinement this measurement motivates is to key the filter on the code type as well as on the element width: the uint16 amplitude codes are log-companded scalars, a different code type from fixed-point coordinates, and compress 8% better unshuffled on the writer's own chunks (Table 4; 9% at 64 KiB, Table 3). That is the 1.44% residual saving of §3.5, with a native decode of 461 against 386 MB/s on that array (141 against 136 MB/s in the browser), measured on the identical chunk set. The encoder cost of the high levels (8–11 MB/s single-threaded at level 9) is a write-once cost, parallelises per chunk, and is irrelevant next to GPU fitting time. The read-time cost is the level penalty of §3.4.

The two cautionary results generalise beyond Luxar. First, a filter can silently fail to execute at deployment chunk sizes: C-Blosc skips bit shuffle on any block whose element count is not a multiple of eight, and its automatic block size grows with the level, so the same 64 KiB chunk is bit-shuffled at level 1 and not at level 9. This is a benchmarking trap: whole-array sweeps rank filters differently from 64 KiB-chunked sweeps, and only the latter reflect what a zarr store actually pays. Any pipeline that tunes codecs on whole arrays and deploys on chunks should re-measure on its real chunk geometry. Second, native decode rankings do not survive the browser: on the identical chunk set the `lz4`-class codecs lose 4.5–5.1× where `zstd` loses 3.2–3.4× and `blosc-zlib` 1.5×, so a native benchmark systematically overstates the case for speed-class codecs in browser-facing storage. Measured in the environment that matters, `zstd`'s ratio advantage costs little.

Finally, the PRECISION comparison keeps the codec question in proportion: lossless coding of the float32 splat parameters reached 1.16–1.77× with a deployable codec on this fit (around a reference of $\approx 1.4\times$ for independent random mantissas, which is not a bound), while the quantisation tiers (uint16 fixed-point coordinates, certified uint8 covariance, uint16 log-companded amplitudes) shrink the same arrays 2.9× before any codec runs. The codec policy recommended here is the cheap residual, worth taking after quantisation has done the real work.
