## Supplementary material for "Luxar: Gaussian splatting for microscopy and scalable interactive web visualisation of multidimensional scientific data": supp_doc_13_consumer_gpu_timing

### Fitting wall-time on a commodity GPU

Supplementary Document 13 – Luxar


**Abstract**

The main text measures its Gaussian-splat fitting times on a workstation-class card with 96 GB of memory. We asked whether a consumer card suffices. On an NVIDIA GeForce RTX 3070 with 8 GiB of memory, every one of the 17 benchmark volumes fits at the fixed reference budget of 32,000 requested splats in 39 to 379 s (median 142 s, about 2.4 min). The claim is scoped to that budget: twelve of the seventeen cross-validation operating points of Table 1 of the main text (16,000, 32,000, 64,000 and 256,000 splats) lie on the timed grid, the other five (8,000, 128,000 and 2,048,000 splats) were not timed here, and 256,000 splats already take up to 27 min, with 2 out-of-memory failures among the 85 cells of the sweep. Peak allocated CUDA memory did not track voxel count, so volume size alone does not predict where the 8 GiB limit binds. To cost the whole adoption path we also ran the pipeline once, end to end, on the same card and on public data: a 400-frame confocal recording of a *C. elegans* embryo was converted, calibrated, fitted frame by frame, merged, compiled, re-chunked and exported in 4.18 h of processing, 93% of it in the fits, with the card never the constraint. A modest laboratory GPU therefore carries both the single-volume fit at the reference budget and the full pipeline for a recording of this size.


#### 1 Introduction

The fitting times of the main text were measured on the workstation GPU that also produced its fidelity results. A laboratory adopting the method will more often have a consumer card, with a fraction of the memory and two hardware generations between it and the workstation part, so two questions decide whether the method is usable outside a well-equipped core facility. The first is whether such a card fits a volume at the reference splat budget in a time a user will wait for, and where its memory limit binds across a sweep of splat counts. The second is what the end-to-end path costs: a recording arrives as TIFF stacks or as an OME-Zarr, and turning it into a link that a collaborator opens in a browser means calibrating the splat count, fitting every frame, merging, compiling a scene, re-chunking it for hosting and serving it.

This document answers the first question with a splat-count sweep over the 17 benchmark volumes of the manuscript on an RTX 3070 (§2, §3), and the second by running the whole path once on the same card and on public data, with every stage timed and its memory sampled (§4). Practical rules for a laboratory close the document (§5).

##### Terms used in this document

***K**** The splat count recommended by a blind-spot sweep: the interior maximum of the held-out PSNR curve when there is a clear peak, the smallest count within 0.3 dB of the maximum on a plateau, and the largest count tested when the curve is still rising there (signal-limited). The shared operating-point table of these documents takes the knee instead, the smallest count within 0.3 dB of that maximum, which is the largest count itself on the two signal-limited benchmark volumes.

**blind-spot cross-validation** A model-selection protocol in which a deterministic random 5% of voxels is hidden from the fit and the reconstruction is scored on those voxels afterwards, so that capacity spent on memorising noise shows up as a loss instead of a gain.

**held-out voxels** The masked voxels of a blind-spot split, whose original values the optimiser never sees and against which the held-out PSNR of each candidate splat count is scored.

**donut-median fill** The replacement of each masked voxel by the median of its $3^D - 1$ neighbours (the donut, 26 voxels in 3D); the shipped calibration draws the median over the unmasked neighbours only, and a document whose sweeps used all 26 neighbours, masked ones included, states that variant where it applies.

**PSNR** The peak signal-to-noise ratio, $10\log_{10}(\text{peak}^2/\text{MSE})$ in decibels, where peak is the reference range of the values compared; unless a document states another range, the supplements take peak 1 on volumes normalised to $[0, 1]$, so that $\text{PSNR} = -10\log_{10}\text{MSE}$.

**cull (retention)** The removal of splats after a fit. The fit-time cull ranks splats by amplitude and keeps the top set accounting for a retention fraction of the total amplitude (0.95 by default); the contribution-based cull instead removes the splats whose contribution to the rendered field falls below an error budget or a fractional threshold, and is the baseline that removal-based level-of-detail methods use.

**relocation** The fixed-pool alternative to adding and removing splats during optimisation: the splat count is fixed at seeding, and the least informative splats are periodically moved to regions of high reconstruction residual, so the optimiser tensor shapes never change.

**leaf** A node that carries splat arrays directly (centres, amplitudes, Cholesky factors) together with an additive ladder of at least one sub-LOD, as opposed to a group node that only holds children.

**additive (streaming) ladder** One ordering of a leaf's splats cut into prefix-sum sub-LODs, so that every prefix of the download is a valid, progressively refining rendering of the same splat set.

**rung (sub-LOD)** One contiguous slab of an additive ladder, stored as its own group (additive_<i>, index 0 the coarsest) and appended to the committed prefix when it arrives; a document that uses "rung" for one step of a splat-count sweep says so where it first does.

**self-energy** The squared $L^2$ norm of one splat's Gaussian, $a_i^2\,\pi^{D/2}\,|\Sigma_i|^{1/2}$, the ordering weight of an additive ladder above the greedy size limit of 5,000 splats (below it a greedy residual ordering is used), the unit of the equal-energy breakpoints, and the quantity whose cumulative fraction stamps how much of a leaf a ladder prefix already carries.

**kind=partition group** A group whose part_<i> children each carry their own position bounds and are all drawn at once, each culled against the camera frustum; the parts are usually the regions of a recursive BSP split of one splat set, but any disjoint grouping is valid, such as one part per timepoint.

#### 2 Methods

**Units and notation.** $K$ is the *requested* splat count, the seed budget passed to the fitter; the post-fit cull retains fewer, and the retained count is recorded per cell. $K^*$ is the cross-validation operating point of Supplementary Document 2. Fidelity is quoted as *PSNR*, the peak signal-to-noise ratio,

$$\text{PSNR} = 10\log_{10}\frac{\text{peak}^2}{\text{MSE}}\ \text{dB}, \tag{1}$$

where MSE is the mean squared error between a reconstruction and its reference over the voxels scored and peak is the reference range of the values compared; this document meets two such ranges, named where they are used (§4). Memory is given in binary units as the GPU and process tools report it (MiB = $2^{20}$ bytes, GiB = $2^{30}$ bytes); the card's 8 GiB (8192 MiB) is its capacity as `nvidia-smi` reports it. Storage sizes are decimal (MB = $10^6$ bytes, GB = $10^9$ bytes), as file systems report them. Decibel values of this document's own calibration are quoted to two decimals, the precision at which its margins are resolved; values taken from Table 1 of the main text keep that table's one decimal. Table 1 of the main text is always named as such; Table 1 is this document's own.

**Hardware.** All fitting timings were measured on a single NVIDIA GeForce RTX 3070 (8 GiB GDDR6, compute capability sm_86) with PyTorch CUDA. The host has an Intel Core i9-9900K (8 cores, 16 threads) and 125 GiB of RAM, and its other GPU is the workstation card; the benchmark was pinned to the consumer card through `CUDA_VISIBLE_DEVICES`. The RTX 3070 is the GPU of the viewer-performance benchmark (Supplementary Document 6). It is deliberately two GPU generations and several price tiers below the RTX PRO 6000 Blackwell used for the fidelity analyses.

The 17 volumes were timed in two sessions on that host. The nine main-figure volumes, the volumes that the main-text figures draw on, were timed while the workstation GPU and the CPUs were occupied by unrelated work: the load average, the number of runnable processes averaged over the preceding minute, was 30–55 on 16 logical CPUs. The eight remaining volumes were timed with the host otherwise idle (load average 1–3). The host facts of the sweep are deposited with the analysis; the harness did not log its Luxar commit or library versions.

A replicate of the sweep on the nine main-figure volumes on the same host under a lighter load gives a contention control. Over the 25 of 45 paired cells whose fits succeed in both sessions and stop at the same iteration, the loaded-session wall time is 1.10 times the replicate's (interquartile range 1.06–1.13, linear-interpolation quartiles). The reported times therefore overstate what an idle host would take by roughly ten percent. The control is one fit per cell in each session, with the two sessions not otherwise matched (code version, cache state), so it indicates the load's effect and does not measure it.

**Fit configuration.** Each fit uses the configuration of the main rate-distortion analysis: L1 loss, learning rate 0.01, fixed-pool splat *relocation*, edge-plus-grid seeding, and a post-fit *cull (retention)* at 99.9% amplitude retention, with an Adam-iteration cap of 20,000 and early stopping at patience 500 (no loss improvement for 500 consecutive iterations). One setting differs, and it is documented here. These runs kept the shipped background-floor default (`--floor auto`) in place of the `--floor none` that the quality benchmarks pin. The background floor is a constant pedestal in the volume (camera offset, autofluorescence) that the fitter estimates from the intensity histogram and subtracts before normalisation; `auto` enables that estimate and `none` disables it. A different target changes the optimisation trajectory and where early stopping triggers, so these wall times describe the shipped floor-auto configuration, what a practitioner running the tool experiences. They are not a hardware-matched comparison with the floor-off fits of the quality benchmarks, with which they are also not bit-comparable.

The recorded wall time is the end-to-end time *to completion under the stopping rule*: the fit ends at early stop or at the 20,000-iteration cap (33 of the 83 successful cells reached the cap; the per-cell iteration count is in the deposited timing tables). It is measured with `time.perf_counter` around the entire `fit_gaussian_splats` call, seed generation, optimisation and culling included. We additionally log the fitter's internal optimisation time, the full-volume PSNR of a post-fit render against the input volume, the iteration count at early stop, and the peak allocated CUDA memory (`torch.cuda.max_memory_allocated`, that is tensor allocations, not the card's whole footprint), read after the fit and the post-fit quality render, so it covers both. The out-of-memory handler brackets the same two steps, so an OOM cell is one that failed in the fit or in that render; the harness does not separate the two. The outer timer is a different quantity from the optimisation time of Table 1 of the main text (the Adam loop alone): at $K = 32{,}000$ the outer time exceeds the fitter's internal optimisation time by a median factor of 1.08 (at most 1.19) across the 17 volumes. Per-cell RNG seeding is deterministic ($\texttt{seed} = K$), matching the main analysis.

**Sweep.** For each volume we fit at $K \in \{4{,}000,\ 16{,}000,\ 32{,}000,\ 64{,}000,\ 256{,}000\}$. $K = 32{,}000$ is the fixed reference budget at which the main text quotes the consumer-card median. $K = 16{,}000$ is the count of the fixed-capacity cell of the convergence analysis (Supplementary Document 4), so the two documents share a splat count. The cross-validation operating points of Table 1 of the main text range from 8,000 to 2,048,000; twelve of the seventeen lie on this grid (16,000, 32,000, 64,000 and 256,000 splats) and are timed here under the floor-auto configuration, while the five at 8,000, 128,000 and 2,048,000 splats are not; the remaining counts bracket the scaling.

### 3 Results

Table 1 reports the end-to-end fitting wall time on the RTX 3070 for every (dataset, $K$) cell. The 17 volumes are 12 physical stacks, 5 of which contribute two channels each. At the fixed reference budget ($K = 32{,}000$) all 17 volumes run to completion under the stopping rule in 39–379 s (median 142 s, about 2.4 min). Of the 17 reference cells, 4 stopped at the 20,000-iteration cap and the rest at patience 500. That range is the basis of the main text's "minutes per volume" figure at this budget; the five operating points of Table 1 of the main text that lie off the grid (8,000, 128,000 and 2,048,000 splats) were not timed on this card. At the top of the sweep ($K = 256{,}000$) the successful fits of the largest light-sheet volumes take longer, up to about 27 min, and two volumes (Fly brain neurons, Neuromast nuclei) exceed the card's memory at this count. Figure 1 plots every cell of Table 1 against $K$ and, beside it, the peak allocated memory of the top-of-sweep cells against volume size.

Two observations follow. First, wall time is not strictly monotonic in $K$. The cap is a fixed iteration budget with early stopping, so the runtime of a cell is governed by *when* the fit plateaus, which depends on the interaction of dataset and capacity and not on $K$ alone. A smaller model on a richly textured volume may run to the iteration cap while a larger model on the same volume plateaus early. Second, and the point that matters for a memory-limited card, the peak allocated CUDA memory stayed within 6.2 GiB (6346 MiB) for every successful cell. That peak did not

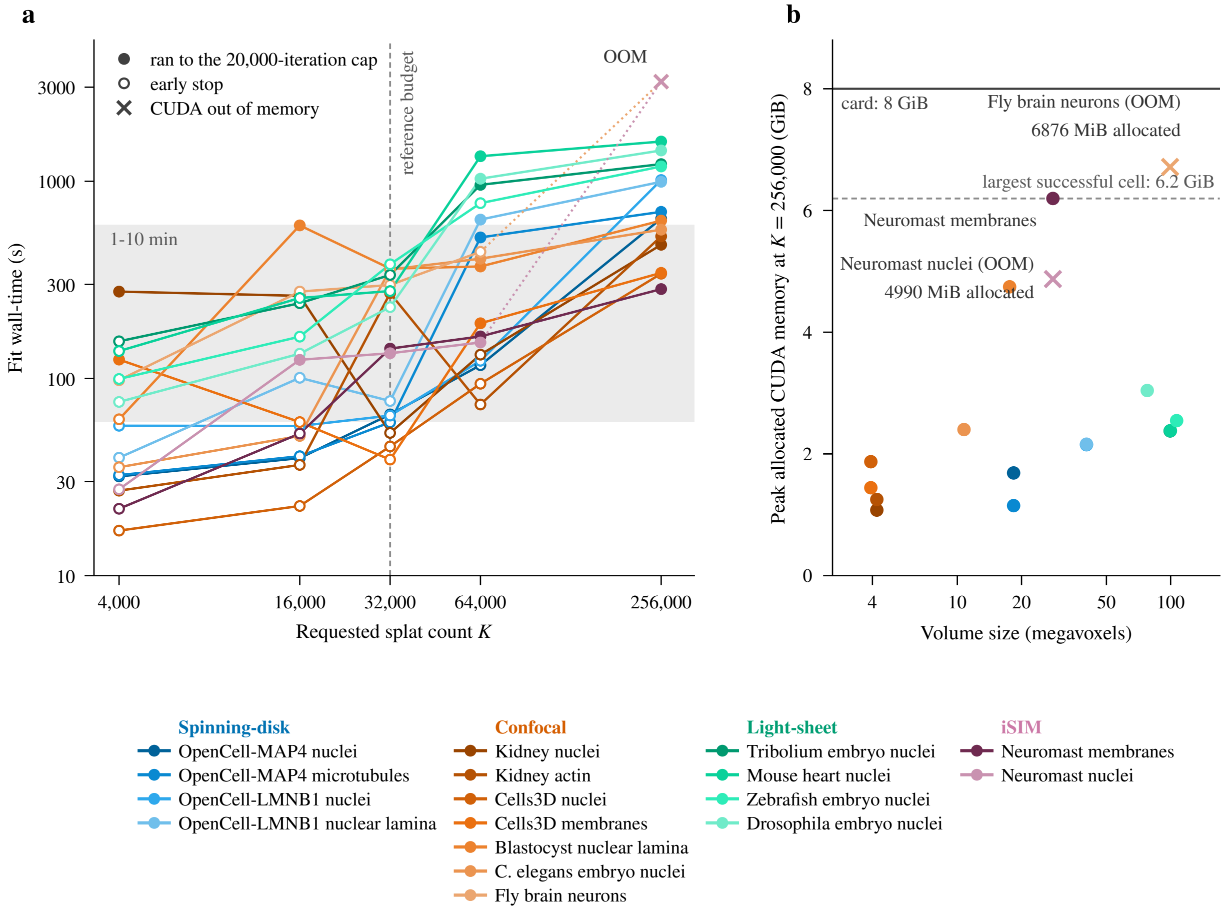


Figure 1: Fitting wall time and peak memory on the RTX 3070. (a) End-to-end fit wall time against the requested splat count $K$, one line per volume, coloured by imaging modality. A filled marker is a fit that ran to the 20,000-iteration cap, an open marker one that stopped early at patience 500, and a cross an out-of-memory cell, which has no wall time and is drawn above the data. The grey band spans 1 to 10 minutes and the dashed vertical line is the reference budget of 32,000. The lines cross and are not monotonic in $K$: the stopping rule, not the splat count, sets the run time. (b) Peak allocated CUDA memory at $K = 256{,}000$ against volume size, one point per volume, in the same colours. The solid line is the card's 8 GiB, the dashed line the 6.2 GiB (6346 MiB) of the largest successful cell; the two crosses are the out-of-memory cells at the bytes allocated when the allocator failed. Memory does not track voxel count, and one failure lies below the largest success.

track voxel count: at $K = 256{,}000$ the successful cells span 1.1–6.2 GiB across 4–107 megavoxels, the roughly 100-megavoxel light-sheet volumes among them. The two volumes that raised a CUDA out-of-memory error at that count, marked OOM in Table 1, are Fly brain neurons (99 megavoxels) and Neuromast nuclei (28 megavoxels).

The failures also show what the recorded peak does not measure. Fly brain neurons had 6876 MiB allocated when the allocator failed, and Neuromast nuclei only 4990 MiB, less than the 6346 MiB that a successful cell reached. A fit was therefore refused memory while its allocated tensors were below a level that another fit survived. The counter covers tensor allocations only; the CUDA context and the memory the caching allocator holds in reserve are outside it, so allocated bytes understate the card's footprint. The 6.2 GiB bound describes the successful cells and is

Table 1: Gaussian-splat fitting wall time (seconds, end-to-end) on a commodity NVIDIA RTX 3070 (8 GiB), across 17 benchmark volumes and a splat-count sweep. $K$ is the requested seed budget. A starred cell (*) ran to the 20,000-iteration cap (33 of the 83 successful cells); every other successful cell stopped early at patience 500. OOM marks the 2 cells that raised a CUDA out-of-memory error (in the fit or the post-fit quality render). Peak is the largest allocated CUDA memory over the row's successful cells (GiB = $2^{30}$ bytes); no successful cell exceeded 6.2 GiB. Voxel counts in millions.

| Dataset | Voxels (M) | Requested splat count $K$ 4,000 | 16,000 | 32,000 | 64,000 | 256,000 | Peak (GiB) |
|---|---|---|---|---|---|---|---|
| OpenCell-MAP4 nuclei | 18 | 32 | 39 | 66 | 117 | 646 | 1.7 |
| OpenCell-MAP4 microtubules | 18 | 32 | 40 | 60 | 519* | 700* | 1.2 |
| OpenCell-LMNB1 nuclei | 40 | 58 | 57 | 65 | 123 | 1014* | 2.2 |
| OpenCell-LMNB1 nuclear lamina | 40 | 40 | 101 | 77 | 639* | 994* | 2.2 |
| Kidney nuclei | 4 | 276* | 262* | 53 | 132 | 477* | 1.1 |
| Kidney actin | 4 | 27 | 36 | 271* | 74 | 526* | 1.3 |
| Cells3D nuclei | 4 | 17 | 23 | 45 | 94 | 340* | 1.9 |
| Cells3D membranes | 4 | 125* | 60 | 39 | 190* | 343* | 1.4 |
| Blastocyst nuclear lamina | 18 | 62 | 597* | 361* | 370* | 632* | 4.7 |
| *C. elegans* embryo nuclei | 11 | 35 | 51 | 357 | 405* | 568* | 2.4 |
| Fly brain neurons | 99 | 98 | 276 | 298 | 439 | OOM | 3.0 |
| *Tribolium* embryo nuclei | 99 | 154 | 241 | 335 | 959* | 1221* | 2.4 |
| Mouse heart nuclei | 100 | 138 | 256 | 277 | 1340* | 1590* | 2.4 |
| Zebrafish embryo nuclei | 107 | 99 | 163 | 379 | 774 | 1189* | 2.5 |
| *Drosophila* embryo nuclei | 78 | 76 | 134 | 230 | 1030* | 1433* | 3.0 |
| Neuromast membranes | 28 | 22 | 53 | 142* | 163* | 284* | 6.2 |
| Neuromast nuclei | 28 | 27 | 124* | 134* | 152* | OOM | 2.3 |

not a threshold below which a fit is safe. Volume size alone does not predict where the 8 GiB card binds, and the harness records nothing that would; the recorded peak also covers the post-fit quality render, whose two full-volume float32 tensors do grow with voxel count. Volumes whose *single-timepoint* size alone exceeds consumer VRAM, the regime that Supplementary Document 11 addresses with tiled fitting, are a separate case; none of the volumes timed here reaches it.

### 4 From an OME-Zarr recording to a shareable link

The timings above are single fits at a fixed budget. A laboratory adopting the method pays for a whole pipeline: calibrating the splat count, fitting every frame of a recording, merging, compiling a scene, re-chunking it for hosting and putting it behind a URL. To cost the processing stages of that path we ran it once, on the same 8 GiB RTX 3070 and on public data, with every stage timed and its host and GPU memory sampled every two seconds (Table 2, Figure 2).

**How the stages were measured.** Each stage command ran under a sampler wrapper. Its wall time is the wrapper's clock, from the child's start to the end of the wrapper's 2 s polling loop, so it is up to one polling interval plus a process-tree scan longer than the command itself. Host memory is the maximum sampled sum of resident set sizes (RSS) over the stage's process tree at 2 s intervals; shared pages are counted once per process, so the sum over-counts, and a peak between samples is missed, so it is not a strict bound. GPU memory is the maximum sampled

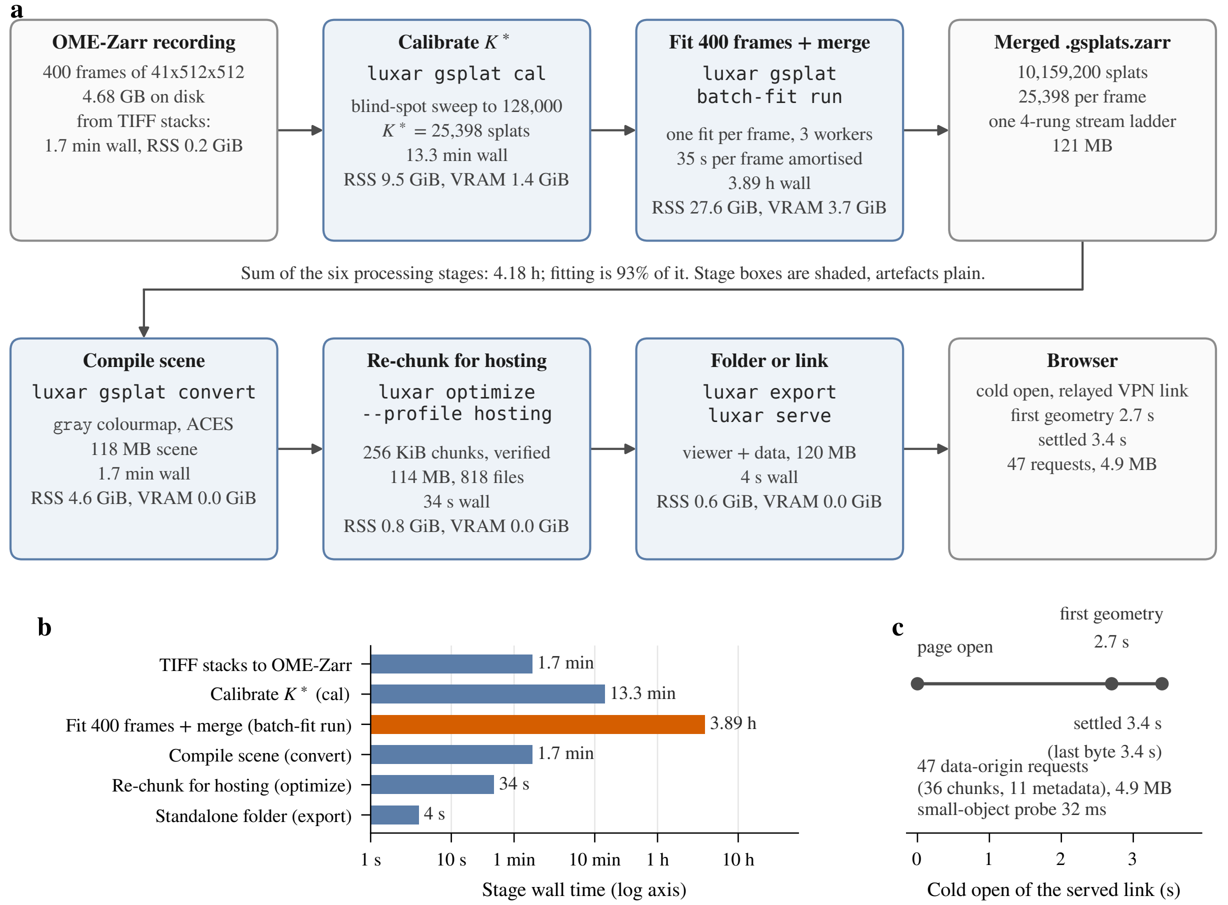


Figure 2: The OME-Zarr-to-link path on the RTX 3070 for the *C. elegans* recording. (a) The path from the TIFF stacks to the browser as eight boxes. Five of the six processing commands are the shaded boxes, each annotated with its wall time, peak host RSS and GPU memory above the idle baseline (export and serve share the last); the sixth, the TIFF-to-OME-Zarr conversion, is noted inside the recording's box with its wall time and RSS. Plain boxes are the artefacts that pass between them, with their sizes. (b) The six stage wall times on a logarithmic axis: the batch fit (orange) takes 93% of the 4.18 h sum, and every other stage is minutes or seconds. (c) The cold open of the served link from a laptop over a relayed VPN link: first geometry at 2.7 s, settled at 3.4 s after 47 data-origin requests (4.9 MB).

`nvidia-smi` usage above the card's idle baseline of 226 MiB. Table 2 sums the six successful processing stages; environment setup (608 s), data acquisition, the failed attempts described below and upload are excluded. The wall clock from the first stage's start to the export's end was 6.08 h.

**Input.** Sample 1 (`mskcc_confocal_s1`) of the confocal recording of *C. elegans* embryonic nuclei deposited in Zenodo record 6460303 (CC BY 4.0). The recording is CTC-style, that is organised as the Cell Tracking Challenge organises its data, one TIFF stack per timepoint: 400 timepoints of 41×512×512 uint16 at 75 s intervals, 8.67 GB of TIFF stacks. It is the recording behind the 4D embryo of Fig. 1b of the main text, taken here from its raw TIFFs and not from the demo's cached fits. A laboratory typically holds such data as one TIFF per timepoint or as an OME-Zarr; we start from the TIFFs and include the conversion as the first stage, because the fitting tools read the OME-Zarr directly.

Table 2: The OME-Zarr-to-link path on the RTX 3070, *C. elegans* sample 1 (400 frames of 41×512×512): wall time (minutes), peak host RSS and GPU memory above the idle baseline for each of the six processing stages, measured as described in the text. GiB = $2^{30}$ bytes.

| Stage | Settings | Wall time (min) | Host RSS (GiB) | VRAM (GiB) |
|---|---|---|---|---|
| TIFF stacks → OME-Zarr | 400 frames, 1×8×256×256 chunks, Blosc-zstd | 1.7 | 0.2 | 0.0 |
| Calibrate $K^*$ (`gsplat cal`) | one timepoint, blind-spot sweep to 128,000 | 13.3 | 9.5 | 1.4 |
| Fit 400 frames + merge (`batch-fit run`) | one tile per frame, one 4-rung stream ladder | 233.6 | 27.6 | 3.7 |
| Compile scene (`gsplat convert`) | `--colormap gray`, ACES | 1.7 | 4.6 | 0.0 |
| Re-chunk for hosting (`optimize`) | 256 KiB chunk target, read back and verified | 0.6 | 0.8 | 0.0 |
| Standalone folder (`export`) | viewer + data, serve anywhere | 0.1 | 0.6 | 0.0 |
| Sum of the six stages | | 251.0 | | |

**Environment.** Luxar at commit `f538cb4` (Python 3.12.3, PyTorch 2.14.0+cu130, CUDA 13.0 build, NVIDIA driver 580.173.02) on Ubuntu 24.04.4 LTS, on the same host as the sweep above (an Intel Core i9-9900K (8 cores, 16 threads) with 125 GiB of RAM). The RTX 3070 was selected for every GPU stage: `CUDA_VISIBLE_DEVICES=1`, scoped to the command, for the single-process stages, and `--gpus 1` (the card's physical index, with no `CUDA_VISIBLE_DEVICES` in the parent shell) for the batch fit. Setting the environment up from a clone took four one-off `make` steps not counted in the table: `make install-demo-deps` (246 s), `make build-cuda` (344 s) and, because `export` and `serve` ship the viewer, `make install-viewer-deps` and `make build-viewer` (18 s together, with a warm package cache and the Node and Rust toolchains assumed; a wheel install carries the built viewer and skips this step). The box was shared with an unrelated CPU load (load average 12–15) throughout, so the wall times are, if anything, pessimistic; this is an observation about the host, not a measured correction. The stage logs, the 2 s resource samples, the batch manifest and the merged store's root metadata are deposited with the analysis.

The batch manifest records the GPU name as "NVIDIA RTX PRO 6000 Blackwell Workstation Edition": at the pinned commit the planner probes CUDA device 0 for its name and per-task estimate (23 s) whichever card the workers are sent to, so that field does not say where the fits ran. The evidence is the resource sampler, which polls `nvidia-smi -i 1`, the RTX 3070, and records device 0 in a separate column: during the batch stage the RTX 3070 rose from its 226 MiB baseline to 4047 MiB while device 0 carried 4–3200 MiB of unrelated load.

**Stages.** The commands are the documented ones. They are written out in full, with every path, in a fail-fast script deposited with the analysis (its header lists the prerequisites), beside the command sequence as executed. The conversion script needs `numpy`, `tifffile`, `zarr` ≥ 3 and `numcodecs`; the resource sampler that wraps every stage needs GNU `time`, `bc` and `nvidia-smi` on Linux.

**Stage 1: conversion.** The manuscript's small converter, deposited with the analysis, turns the 400 TIFFs into one OME-Zarr 0.4 image (`t,z,y,x`; voxel steps 75 s and 0.75×0.15×0.15 µm;

1×8×256×256 chunks, Blosc-zstd), 4.68 GB on disk.

**Stage 2: calibration.** The command

```
luxar gsplat cal celegans_s1.ome.zarr cal.json --timepoint 100 --axes time,z,y,x
--k-max 128000 --device cuda --pdf cal.pdf
```

runs the *blind-spot cross-validation* sweep of Supplementary Document 2 on one timepoint under the tool's default calibration preset: 20,000 iterations, patience 500, 5% *held-out voxels*, mask seed 42, *donut-median fill* radius 1; the floor default subtracted a pedestal of 2132.0 grey levels first. The grid is 10 counts from 1,000 to 128,000, capped at 128,000 to bound the calibration time; the sweep, PDF report included, took 13.3 min (Table 2).

It returns $K^* = 25{,}398$ requested seeds (24,804 retained) with a held-out peak of 59.44 dB against a 63.71 dB noise ceiling, the held-out PSNR at which a reconstruction matches the volume up to its estimated noise and above which no further gain is real. Both figures are on the calibration's own scale (metric `psnr_minmax`): PSNR with the peak taken as the minimum-to-maximum range of the floor-subtracted raw grey levels of that timepoint. Table 1 of the main text scores the same timepoint on a $[0, 1]$-normalised, percentile-clipped basis with no floor, where the fit at its operating point of $K^* = 8{,}000$ scores 39.3 dB full-volume PSNR (40.5 dB at best over its sweep). The two dB scales are not comparable, and the offset between them is the change of reference range.

The operating point differs from the main text's for three reasons: the floor (auto here, 2132.0 grey levels subtracted; none there), the grid (10 log-spaced counts from 1,000 to 128,000 here, of which $K^*$ is the seventh; a doubling grid there), and the margin, which is small on a jagged curve. The peak lies 0.18 dB above the 128,000 endpoint, and the 8,640 point sits 2.88 dB below the peak on this scale. The tool's "confidence" of 0.18 dB is a heuristic score, the margin between the best and the second-best held-out PSNR (here the terminal 128,000 point at 59.26 dB), from a single sweep and not a statistical interval. The curve is jagged at this single-fit resolution. The tool's parametric rate-distortion model, which fits held-out error against $K$ as a floor plus a power law, reports a converged fraction of 0.67, the share of the achievable error drop that the sweep has realised by its largest $K$. Below 0.85 the tool flags the sweep as not converged, that is it extrapolates further gain beyond 128,000, even though the measured curve's last point lies 0.18 dB below its peak.

**Stage 3: batch fit and merge.** The command

```
luxar gsplat batch-fit run celegans_s1.ome.zarr fit/ --gpus 1 --axes time,z,y,x
--tile-size 1024 --seeds 25398 --jobs-per-gpu 3 --merge-recipe stream
--merge-n-lods 4
```

fits every frame as one tile (the tile size exceeds the frame) with three concurrent workers on the card, then runs the streaming merge.

With a single spatial tile for each frame the merge takes its single-tile branch: it emits a bare leaf instead of wrapping one part in a partition. The output (`fit/merged/final.gsplats.zarr`) is one stacked four-dimensional Gaussian-splat *leaf* of 10,159,200 splats (25,398 in each frame). It carries one *additive (streaming) ladder* of 4 rungs over the whole recording, built by the streaming merge (`--merge-recipe stream`): 2,539,800 splats per *rung (sub-LOD)*, cut at equal counts along one global *self-energy* ordering (`self_energy`), as the merged store's root metadata records; the per-timepoint split across the rungs is not recorded. There is no ***`kind=partition`*** *group* around it; the partition form is what a recording tiled into several spatial regions produces.

The batch fits use the batch command's default preset, `standard`, the one setting of the fit budget the batch manifest records; the preset is defined as a 5,000-iteration cap with patience 300, not the calibration's 20,000/500 budget. The planner resolved the floor once over four sampled frames (2136.4 grey levels subtracted by every task) and one shared normalisation range (1997–19110), and at the pinned commit the per-tile worker applied no post-fit cull, so every frame keeps all 25,398 requested seeds ($400 \times 25{,}398 = 10{,}159{,}200$). The realised per-frame iteration counts and the RNG policy of the 400 fits are not in the deposited logs, so the batch stage costs a `standard`-preset fit for each frame and is not evidence about the main text's 20,000-iteration protocol.

The batch stage is the one a laboratory budgets for: 3.89 h for 400 frames including the merge, an amortised 35 s for each frame at three workers, 93% of the stage sum. That amortised figure and the 357 s of the *C. elegans* cell at $K = 32{,}000$ in Table 1 are the same recording under two protocols, and they reconcile as follows. Each batch fit held a worker for about 105 s (the stage wall time times three workers over 400 frames, merge included), at $K = 25{,}398$ under the 5,000-iteration cap, with no cull and no quality render. The table's cell is a single fit at $K = 32{,}000$ under the 20,000-iteration cap that ran 19,625 iterations, with the cull (30,636 splats retained) and the full-volume quality render inside the timer. The two numbers differ mainly in the iteration budget and in what the timer encloses, and neither is the other's per-frame cost.

The worker count is pinned explicitly (`--jobs-per-gpu 3`) and not left to `auto`: an explicit count keeps the host's memory within budget on these small frames, and the script states it so that the trace is reproducible. The card is selected by its physical index (`--gpus 1`, no `CUDA_VISIBLE_DEVICES` in the parent shell, as above), so every worker runs on the RTX 3070. The resource samples of the batch stage (3,882 samples) show the RTX 3070's memory rising from its 226 MiB idle baseline within 19 s of launch to a 4047 MiB peak. The workstation GPU carried unrelated load throughout, so exclusivity rests on the pinning and on the RTX 3070's own memory trace, not on the manifest's GPU name (Environment).

**Stage 4: scene compilation.** The command

```
luxar gsplat convert fit/merged/final.gsplats.zarr celegans_s1.luxar.zarr
--center --colormap gray --tone-mapping ACES
```

writes the scene (118 MB). The OME-Zarr axis names are passed explicitly to the calibration and the batch fit (`--axes time,z,y,x`); the compiled scene names its time axis positionally, and both the stage's log and the deposited link trace record it as `dim3`.

**Stage 5: re-chunking for hosting.** The command

```
luxar optimize celegans_s1.luxar.zarr celegans_s1_hosting.luxar.zarr --profile
hosting --verify
```

re-chunks the scene to the compiler's 256 KiB hosting chunk profile with a full read-back check (114 MB, 818 files).

**Stage 6: export.** The command

```
luxar export celegans_s1_hosting.luxar.zarr -o export/ --overwrite
```

writes a standalone folder holding the viewer and the data (120 MB, 851 files), which any static file host or `luxar serve` turns into a link.

**Failed attempts.** The 1.89 h between the 6.08 h elapsed and the 4.18 h stage sum holds three discarded attempts at stage 3 and one at stage 4, kept as resource traces deposited with the analysis (no stdout of them is deposited). They are: a launch with an empty `--seeds` after a JSON-key error (2.3 s); a `--jobs-per-gpu auto` launch, recorded by the operator as having sized 90 workers for these small frames, killed after 25.5 s; and a run whose resource trace shows the RTX 3070 idle for 1.35 h while the fits ran (its RTX 3070 column never exceeds 388 MiB over its 1,384 samples); the operator's record attributes it to the workers landing on the workstation GPU, and it was stopped and discarded once noticed. The stage-4 retry with an empty merged-store path left no trace. The three traces sum to 1.36 h; the rest of the gap is diagnosis time.

**What it costs: time and memory.** The six processing stages sum to 4.18 h, of which fitting the 400 frames is 93%. Host memory, summed over the stage's process tree, peaked at 27.6 GiB during the batch fit (three fit workers plus the compile-worker pools that PyTorch's inductor spawns per process, shared pages counted once per process; the largest single process was 3124 MiB) on a host with 125 GiB (4.5 times the peak), and at 9.5 GiB during calibration. Sampled GPU memory never exceeded 3.7 GiB above baseline, so the 8 GiB card was not the constraint at this frame size and splat count. Summed RSS over-counts shared pages, so the true footprint is lower, but a 32 GB box may not run three workers at this frame size; `--jobs-per-gpu 1` or `2` is the setting to start from there.

**What it costs: storage.** Temporary storage is the batch output directory (0.25 GB, 22,188 files). Its per-frame tile stores (`fit/tiles`, 127 MB, 11,600 files) can be deleted once the merge has written its output, keeping the merged store (`fit/merged`, 121 MB) and the manifest. The deliverables are the hosting-profile scene (114 MB) or the standalone folder (120 MB).

**What it costs: opening the link.** The scene was served from the same machine with

```
luxar serve celegans_s1_hosting.luxar.zarr --viewer --host 0.0.0.0 --port 8421
  --viewer-port 5421 --cors-origin '*'
```

and opened cold at http://<host>:5421/?src=http://<host>:8421 from a laptop (Apple M4 Max, headless Chromium 151.0.7922.34 via ANGLE/Metal, 1280×800 viewport) over a relayed VPN link. The scene's first nonzero geometry count appeared after 2.7 s. The opening settled after 3.4 s (last data byte at 3.4 s), by which point 47 data-origin requests (36 chunks, 11 metadata; 4.9 MB; the viewer's own files excluded) had completed. 4 of 10 consecutive time steps needed any request at all (at most 0.3 MB, because one hosting chunk holds several of these 25,398-splat frames). A distant seek (median of 3) fetched 48 chunks (7.4 MB) with its last byte at 2.7 s, on a link whose small-object HTTP probe measured 32 ms (the playback-trace probe of Supplementary Document 14), and the browser's process tree stayed at 0.7 GiB.

**Publishing.** Publishing to a bucket adds an upload of the hosting store's bytes at the laboratory's uplink rate, which we do not time here. The viewer then reads the store from any reachable HTTP(S) host that serves its files and permits the viewer's origin through CORS: `luxar serve` does this itself (above, `--cors-origin '*'` for the remote laptop), a bucket needs an equivalent CORS rule, and co-hosting viewer and data from the exported folder on one origin avoids the cross-origin fetch altogether.

**What it does not cost.** No cluster and no workstation-class GPU were involved, and every successful stage is one documented command line (plus the conversion script). The two things a laboratory does have to decide (the splat count, which `cal` measures instead of guessing, and the appearance, which `convert` bakes in and the viewer's layer panel lets the recipient change) are the same two decisions that the manuscript's own recordings took (Supplementary Document 14).

### 5 Summary and practical rules

An 8 GiB consumer card runs the reference budget on every benchmark volume in minutes and the whole splat-count sweep to 256,000 on all but two of them. Of the 17 operating points of Table 1 of the main text, twelve (16,000, 32,000, 64,000 and 256,000 splats) lie on the timed grid and are covered by the sweep; the five at 8,000, 128,000 and 2,048,000 splats were not tested, and at 256,000 splats the largest light-sheet volumes already take up to 27 min. Memory, not time, is what fails first, and it fails in a way that voxel count does not predict, so a laboratory sizing a fit should treat the allocated-memory bound as descriptive and try the count it wants on its own volume before planning a batch.

For a recording, the fits dominate: 93% of the processing time of the end-to-end path went into fitting the frames, at three workers on one card, and conversion, calibration, compilation, re-chunking and export shared the rest. Three settings decided the outcome and are worth pinning in any script. Select the GPU by its physical index and check the card's own memory trace, because at the pinned commit the planner's manifest names the card it probed, not the one the workers used. Pin the worker count, since the automatic choice was recorded as sizing 90 workers for these small frames. Calibrate the splat count once with `cal` on a representative timepoint, read its operating point as a grid point on a jagged single-fit curve, and let the batch fit inherit it.

For the delivered link, neither host nor GPU memory bounded the batch fit on this host, though a 32 GB host may need fewer workers; storage stayed within a few hundred megabytes at every stage after conversion, and the viewer opened the recording cold in seconds over a relayed link, with most time steps needing no request at all.
