## Supplementary material for "Luxar: Gaussian splatting for microscopy and scalable interactive web visualisation of multidimensional scientific data": supp_doc_14_timelapse_splats

### Gaussian-splat time-lapses:
### three recordings, one timepoint resident at a time

Supplementary Document 14 – Luxar

**Abstract**

Three of the demo scenes are whole recordings fitted as Gaussian splats: a 51-frame zebrafish embryo (histone label, light-sheet), a 500-frame *Drosophila* embryo (30-second frames through gastrulation, SiMView light-sheet) and a 100-frame two-channel neuromast (iSIM, membranes and nuclei). Each is fitted one timepoint at a time and stacked into four-dimensional splat nodes whose fourth centre column is time. The archives are 157×, 67× and 157× smaller than the source voxels they represent. Time is a hard ordering barrier in every archive, so only one timepoint is ever resident on the GPU, and in the published chunk layout a time step costs 1 request on the *Drosophila* scene, 1 on the neuromast and one per-timepoint part (17.91–17.96 MB) on the zebrafish, whose frames are too large to share a chunk with their neighbours. That economy is paid at the opening, where a 1 MB chunk brings the neighbouring frames along. This document records how the three archives were built, what they cost to fit and to store, what a browser downloads when the time slider moves, and how faithful each shipped archive is to the raw frames on sampled timepoints.

#### 1 Introduction

The benchmark volumes of the main text are single stacks. A time-lapse poses three further questions: how to fit hundreds of frames within a fixed GPU and time budget, how to store them so that the whole recording is one archive, and how to stream them so that moving the time slider downloads only what the new frame needs. This document answers the three for the three recordings of Figure 4 of the main text.

The approach is the same for all three. Every timepoint is fitted on its own, with the single-volume fitter, and the per-frame splat sets are stacked along a fourth centre column that holds the time value. The result is a four-dimensional splat *node*, one scene-graph element that the viewer draws as a whole, of which the time axis is a *hidden dimension*: the viewer displays three axes and slices the fourth at the slider's position, so one frame is on screen at a time. The zebrafish recording keeps one *part* per timepoint inside its node, the *Drosophila* recording is one *leaf* and the two-channel neuromast is one leaf per channel. Within a leaf, an *additive (streaming) ladder* lets the viewer draw a frame progressively, one *rung (sub-LOD)* at a time, as its chunks arrive.

Section 2 gives the recordings and their fitting recipes. Section 3 explains why time is a hard ordering barrier and what that buys. Section 4 reports the archive sizes and Section 5 what a browser downloads at the opening, per time step and over a long stepped session. Section 6 scores the shipped archives against the raw frames, Sections 7 and 8 cover the two-channel scene and the capture of Figure 4, and Section 9 collects the practical rules.

#### Terms used in this document

**leaf** A node that carries splat arrays directly (centres, amplitudes, Cholesky factors) together with an additive ladder of at least one sub-LOD, as opposed to a group node that only holds children.

**part** One child of a `kind=partition` group (`part_<i>`), holding one disjoint subset of the splats (a BSP region, or a timepoint) together with its own position bounds for frustum culling.

**`kind=partition` group** A group whose `part_<i>` children each carry their own position bounds and are all drawn at once, each culled against the camera frustum; the parts are usually the regions of a recursive BSP split of one splat set, but any disjoint grouping is valid, such as one part per timepoint.

**`kind=lod` group** A substitutive level-of-detail group whose children are alternative renderings of the same content at different splat counts, of which the viewer draws exactly one at a time: the coarsest level whose median projected splat footprint stays at or below 1.5 CSS pixels when the levels carry footprint stamps, otherwise the level selected by the size of its projected bounding box (its fraction of the viewport area for a screen-area ladder, its normalised projected diagonal for a coverage ladder).

**additive (streaming) ladder** One ordering of a leaf's splats cut into prefix-sum sub-LODs, so that every prefix of the download is a valid, progressively refining rendering of the same splat set.

**rung (sub-LOD)** One contiguous slab of an additive ladder, stored as its own group (`additive_<i>`, index 0 the coarsest) and appended to the committed prefix when it arrives; a document that uses "rung" for one step of a splat-count sweep says so where it first does.

**self-energy** The squared $L^2$ norm of one splat's Gaussian, $a_i^2\,\pi^{D/2}\,|\Sigma_i|^{1/2}$, the ordering weight of an additive ladder above the greedy size limit of 5,000 splats (below it a greedy residual ordering is used), the unit of the equal-energy breakpoints, and the quantity whose cumulative fraction stamps how much of a leaf a ladder prefix already carries.

**committed prefix (committed count)** The part of an additive ladder that has been processed, uploaded to the GPU and is on screen, as distinct from rungs whose download has merely completed.

**spatial index** The chunk-based index of a node: elements are reordered along a space-filling curve and grouped into fixed-size chunks, each with a stored axis-aligned bounding box (`chunk_bounds`), so a slice query reads only the chunks whose boxes intersect it.

**atom** The spatial index's `chunk_size` in elements, of which every zarr chunk of a per-element array is a whole multiple, so an index range never straddles a zarr chunk.

**Morton and Hilbert ordering** Two space-filling curves that sort points so that neighbours in space become neighbours in memory or on disk: Morton (Z-order) interleaves the bits of the quantised coordinates, Hilbert follows a continuous curve whose consecutive cells are always adjacent, so it has no long jumps between neighbouring runs. The compiler orders the stored elements along one of them; the fitter re-sorts its splats by Morton code periodically for GPU locality, and the substitutive LOD builder uses a Morton sort as its warm-start partition.

**barrier dimension** A centre column (a time or channel axis) that the compiler and the LOD builders never mix across. The writer sorts the elements by that column first and computes the spatial-index curve over the spatial columns only, so every coordinate's elements are contiguous on disk; substitutive coarsening leaves the column out of `coarsen_dims`, so every coarse level stays separated per coordinate; and an additive ladder is built per coordinate, so no rung blends two of them.

**`content_hash`** A post-order xxhash64 digest of a store (array bytes, storage identity, attributes and child digests) against which the viewer validates its cache, so any edit to the data invalidates stale chunks.

**cull (retention)** The removal of splats after a fit. The fit-time cull ranks splats by amplitude and keeps the top set accounting for a retention fraction of the total amplitude (0.95 by default); the contribution-based cull instead removes the splats whose contribution to the rendered field falls below an error budget or a fractional threshold, and is the baseline that removal-based level-of-detail methods use.

**blind-spot cross-validation** A model-selection protocol in which a deterministic random 5% of voxels is hidden from the fit and the reconstruction is scored on those voxels afterwards, so that capacity spent on memorising noise shows up as a loss instead of a gain.

**held-out voxels** The masked voxels of a blind-spot split, whose original values the optimiser never sees and against which the held-out PSNR of each candidate splat count is scored.

$K^*$ The splat count recommended by a blind-spot sweep: the interior maximum of the held-out PSNR curve when there is a clear peak, the smallest count within 0.3 dB of the maximum on a plateau, and the largest count tested when the curve is still rising there (signal-limited). The shared operating-point table of these documents takes the knee instead, the smallest count within 0.3 dB of that maximum, which is the largest count itself on the two signal-limited benchmark volumes.

**SSIM** The structural similarity index, the mean over local windows of a product of luminance, contrast and structure comparisons between two images, lying in $[-1, 1]$ with 1 meaning identical.

**iSIM** Instant structured illumination microscopy, a super-resolution fluorescence technique that performs the structured-illumination reconstruction optically in a single exposure, giving roughly a twofold resolution gain at camera frame rates.

#### 2 Recordings and recipes

Table 1 collects the acquisition metadata of the three recordings; the paragraphs below give the fitting recipes, and Figure 1 lays the pipeline out stage by stage with the cost this document measures for each stage.

**Zebrafish embryo nuclei (*h2afva*).** A Zebrahub recording of a zebrafish embryo whose every nucleus carries a histone-H2A variant fusion, imaged on a light-sheet microscope: 253 timepoints of $407 \times 2048 \times 2048$ voxels, fused and deconvolved (0.40625 $\mu$m laterally, 1.625 $\mu$m axially)[2]. The full recording was fitted with the tiled batch fitter under *content* tiling: every timepoint its own fit, partitioned into content-balanced boxes (11,132 over the recording) that are fitted on halo-padded, unwindowed crops and merged by keeping each box's core-centre splats, one ***kind=partition** group* per frame; scaled isotropic and published as a 5.87 GB archive. This is not the apodised uniform tiling whose seams Supplementary Document 11 measures; that document notes that the seam behaviour of content tiling has not been measured. That fit occupied one RTX PRO 6000 for 112.5 hours (the box-completion timeline with one 11.7-h idle gap removed, since that fitter stamped no per-fit times).

The variant used here keeps every fifth frame (original indices $0, 5, \ldots, 250$; 51 frames), flattens the partition to one leaf, rebuilds a 4-rung equal-count progressive ladder and re-chunks the store at 1 MB. The partition and its substitutive levels (its ***kind=lod** groups*) are dropped because the viewer never selects them: the node is only ever seen one timepoint at a time, so they cost bytes and requests without changing the finest-level content. The demo re-authors that archive at build time, without refitting, into one laddered part per timepoint (51 parts of 2.1M–2.6M splats), so a time step loads exactly one part (17.91–17.96 MB, Section 5) and no rung spans two timepoints. Each part carries its own eight-rung equal-count progressive ladder (cumulative cuts at $k/8$ of the

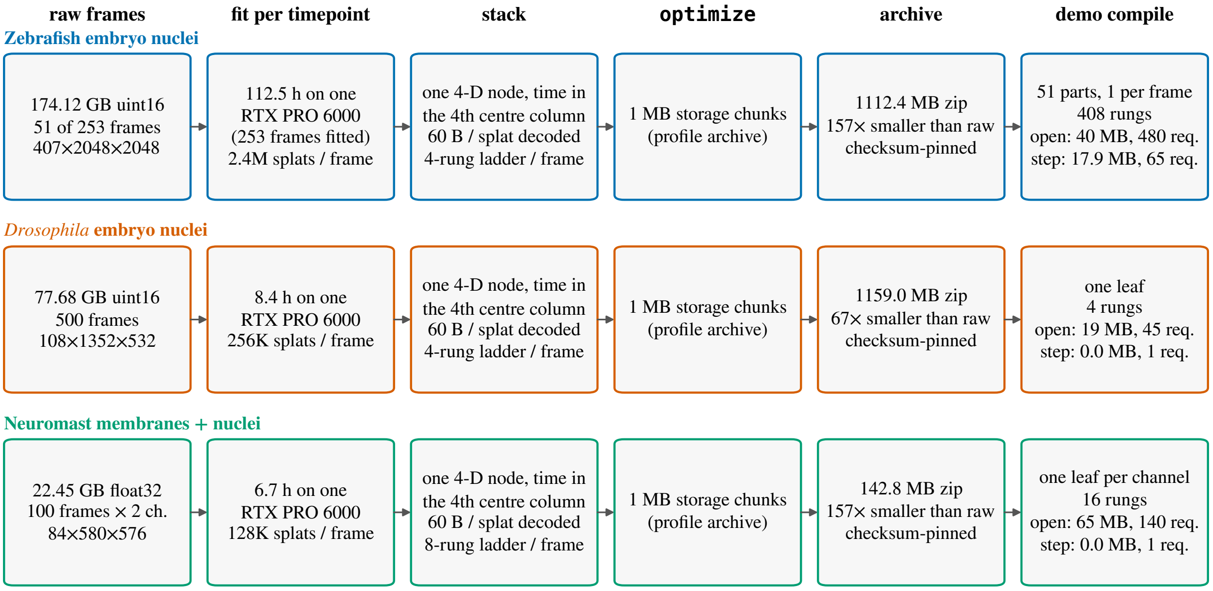


Figure 1: The pipeline of a splat time-lapse, one lane per recording: the raw frames, a fit per timepoint on one RTX PRO 6000, the stack into one four-dimensional node whose fourth centre column is time, the re-chunking pass, the published archive and the demo compile that grafts it into a scene. Each cell carries the cost this document measures for that stage: source bytes and frame grid, fit hours, the ladder and the 60-byte decoded record of the stacked node, the chunk profile, the archive size and ratio, and the compiled scene's structure with its opening and time-step cost (Sections 2, 4 and 5). Every number is one of this document's generated macros.

part's splats, $k = 1, \dots, 8$, ordered by *self-energy* so the brightest nuclei paint first; every per-part increment stays under the 900,000-splat commit cap, the largest rung the viewer processes in one step), 408 rungs over the 51 parts. The centres are in lateral-pixel units (the archive is published, so its bytes and checksum are fixed); the scene applies the calibration when it grafts the parts, that is, copies them into the scene store without refitting.

**Drosophila embryo nuclei.** A SiMView light-sheet recording of a *Drosophila melanogaster* embryo, `His2Av::mRFP1` nuclei, 500 timepoints at 30-second resolution ($500 \times 108 \times 1352 \times 532$ uint16; 19.3 GB compressed, 77.68 GB decoded). It was acquired in the Keller laboratory at HHMI Janelia Research Campus on the instrument of Royer *et al.*[6]. Its four views (two illumination arms, two detection cameras) were registered and fused into one volume per timepoint with the SiMView processing pipeline[1, 7], without deconvolution, and the recording was fitted at Biohub[5]. Every timepoint is fitted at full resolution with stock commands (names as registered in the luxar CLI of the build recorded with the analysis):

```
luxar gsplat batch-fit run DrosophilaHistone.zarr.zip out/ \
    --array-key data --axes time,z,y,x \
    --tiling uniform --tile-size 1400 --overlap 0 \
    --preset standard --iters 1500 --seeds 256000 --floor 8 \
    --gpus 0 --jobs-per-gpu 2 --merge-recipe stream --merge-n-lods 4
```

Table 1: The three recordings. Voxel pitches are those of the arrays that were fitted (the fused, deconvolved zebrafish stack; the fused *Drosophila* frames; the deconvolved neuromast frames, whose 0.25 / 0.1083 $\mu$m pitch is the 2.308 axial-to-lateral ratio the scene applies). The zebrafish interval is per represented frame (every fifth 220-s acquisition); the neuromast interval is read from the *iSIM* TIFF timestamps. Frame counts are per channel.

| Recording | Specimen, stage | Structure, marker | Microscope, processing | Voxel $z$ / $y$ / $x$ ($\mu$m) | Frame grid, dtype | Frames | Interval |
|---|---|---|---|---|---|---|---|
| Zebrafish embryo nuclei | Zebrafish embryo, gastrulation onward | nuclei, H2A.Z histone (*h2afva*) fusion | OpenSiMView light-sheet; fused, deconvolved | 1.625 / 0.40625 / 0.40625 | 407 × 2048 × 2048 `uint16` | 253 acquired, 51 represented | 18.3 min per represented frame (220 s acquired) |
| *Drosophila* embryo nuclei | *Drosophila melanogaster* embryo, cellular blastoderm to extended germband | nuclei, `His2Av::mRFP1` | SiMView light-sheet; four views registered and fused, not deconvolved | 1.93 / 0.40625 / 0.40625 | 108 × 1352 × 532 `uint16` | 500 | 30 s |
| Neuromast membranes + nuclei | Zebrafish lateral-line neuromast, 3 dpf | membranes, `cldnb:lyn-mScarlet`; nuclei, `she:GFP` | iSIM; Richardson–Lucy deconvolved, motion aligned | 0.25 / 0.1083 / 0.1083 | 84 × 580 × 576 `float32` | 100, 2 channels | 5.6 min |

```
luxar gsplat transform out/merged/final.gsplats.zarr um.gsplats.zarr \
    --scale 1.93,0.40625,0.40625,1
luxar optimize um.gsplats.zarr drosophila_embryogenesis_500tp.gsplats.zarr.zip \
    --profile archive
```

A tile size above every spatial extent makes the merge emit one stacked four-dimensional leaf, not a partition. The fan-in merge streams spatial tiles, so its peak host memory is one tile's splats over all timepoints and channels; for a recording fitted as one tile per frame, as here, that is the whole archive at float32, 60 bytes per splat before compression (Section 3 reconciles that four-dimensional record with the 40-byte three-dimensional payload a time step delivers). The archive keeps every fitted splat: 256K per frame, 128,000,000 in total, with no post-fit *cull (retention)*. The four equal-count ladder rungs hold 64,000 splats of every timepoint each, so each timepoint's first rung is a quarter of its frame.

Two ablations on frame 150 fix the recipe's two free choices. A cumulative amplitude cull keeping 95% of the amplitude mass removed 38% of the splats (256,000 to 157,651) and dropped the fit (a 1,000-iteration fit of that frame) from 41.7 / 37.1 dB to 32.0 / 27.1 dB (global / foreground PSNR against the floor-subtracted frame; the fit and score logs are deposited with the analysis), so the archive is not culled. Iterations are not the lever on this data: 5,000 iterations score 41.8 / 37.2 dB (global / foreground) against 41.8 / 37.2 dB at 1,500 (same logs). Seeding takes about a minute of the roughly two minutes a frame takes end to end. The 500 fits occupied one RTX PRO 6000 for 8.4 hours (two fits in flight). The per-frame times stamped in the tiles cover the optimisation loop only, not seeding or I/O, and sum to 8.56 h with a median of 0.8 min per frame. Each stamp was timed while two fits shared the card, and the loop is about half of each frame's two minutes, so the sum lands close to the elapsed time although it covers only the loop.

The time axis is expressed in minutes with a step of 0.5, which is exactly representable in binary. A discrete viewer axis snaps to its grid and fetches only within a quarter step of the snapped position, so a frame sitting off the grid would be a slider stop that renders nothing. The build asserts every frame lands on the grid.

**Neuromast membranes and nuclei.** A two-channel iSIM recording of a developing zebrafish lateral-line neuromast (`she:GFP; cldnb:lyn-mScarlet`, Jacobo laboratory, CZ Biohub), 100 timepoints of $84 \times 580 \times 576$ voxels per channel, Richardson–Lucy deconvolved and motion aligned, distributed on the permissively licensed demo record[4]. The splat budget per frame is a common 64,000 for both channels. That is the requested count at the $K^*$ of the *blind-spot cross-validation* sweep of Supplementary Document 2 for the nuclei channel (63.6 dB held-out at 64,000 requested); for the membranes the same sweep peaks at 128,000 requested (50.9 dB, against 50.7 dB at 64,000). Those are requested counts, not realised ones: the sweep's fits carry a post-fit cull that leaves the realised count at 32,715 (nuclei) and 74,825 (membranes) at those peaks, and Supplementary Document 2 classifies both channels as cull-limited, the realised count stopping its growth once the request passes the peak. The sweep also runs on the floor-subtracted, percentile-normalised benchmark frame, not on the raw frames fitted here, which are fitted without a cull and keep all 64,000 splats, so 64,000 is a chosen shared budget, not a per-channel calibration result.

Each channel was fitted with the fitter's noise-aware `n2s` preset, one tile per frame, on the raw frames with no floor subtraction and no post-fit cull, before the merge built one leaf per channel with an 8-rung equal-count progressive ladder. The two channel fits occupied one RTX PRO 6000 for 6.7 hours (twelve fits in flight; the per-fit times stamped in the tiles sum to about 80 h because of the sharing). The two channels are co-registered in one four-dimensional coordinate space and exposed as two layers, so they toggle independently but animate together.

Fitting the raw frames is deliberate. The recording carries a camera pedestal of about 106 counts, and a fit on the exactly floor-subtracted, zero-clipped frames loses the faint background (Table 2 and Figure 2; the fit logs are deposited with the analysis). The sweep fits frames 50 and 99 at 64,000 seeds on the frame minus a listed floor, clipped at 0, and scores every arm against the frame minus the recorded 105.99-count pedestal. Three intensity bands, given as fractions of the frame maximum, are scored separately: very low $[0.2\%, 1\%)$, dim $[1\%, 10\%)$ and foreground $[10\%, 100\%]$; each band lists the Pearson correlation of the render against the frame, and the very-low band also the ratio of rendered to reference mass. Dead seeds are the share of the 64,000 whose amplitude fell below $10^{-4}$ of the peak. Seeds that land on clipped zeros have nothing to fit and die (36% of them at frame 50, 65% at the bleached frame 99, against 1–2% on the raw frames). The very-low band, where the membrane-associated background structures live, correlates at only 0.16 with the data at frame 50, against 0.55 for the pedestal-inclusive fit (0.97–0.99 in the dim band and 1.00 in the foreground for both). Leaving a partial pedestal (subtracting 90–103 counts) lands in between (very-low-band correlation 0.26–0.45); relaxing the fit's amplitude sparsity changes nothing (0.13–0.16). The pedestal costs splats spent on a constant background, which the display window then hides.

Table 2: Neuromast membranes: what subtracting the camera pedestal before the fit does to the faint background. Frames 50 and 99 fitted at 64,000 seeds (`n2s` preset) on the frame minus the listed floor (clipped at 0), every arm scored against the frame minus the 105.99-count pedestal, so the raw-frame arm's global PSNR and very-low mass ratio measure the pedestal it reproduces, not its fidelity; its correlations are offset-invariant and are the numbers the text compares. Bands and dead seeds as defined in Section 2. The two relaxed-sparsity arms weight the fit's amplitude penalty by 0.1 and 0.

| Frame | Floor subtracted | Dead seeds (%) | Very-low corr. | Very-low mass ratio | Dim corr. | Foreground corr. | Global PSNR (dB) |
|---|---|---|---|---|---|---|---|
| 50 | none (raw frame) | 1 | 0.55 | 56.61 | 0.99 | 1.00 | 22.0 |
| | 90 | 25 | 0.45 | 9.16 | 0.98 | 1.00 | 42.4 |
| | 96 | 30 | 0.40 | 6.09 | 0.98 | 1.00 | 46.6 |
| | 100 | 32 | 0.33 | 4.16 | 0.98 | 1.00 | 50.9 |
| | 103 | 34 | 0.26 | 2.82 | 0.97 | 1.00 | 55.7 |
| | 105.99 (exact pedestal) | 36 | 0.16 | 1.77 | 0.97 | 1.00 | 60.7 |
| | 105.99, sparsity 0 | 36 | 0.16 | 1.74 | 0.97 | 1.00 | 60.5 |
| | 105.99, sparsity $\times 0.1$ | 36 | 0.16 | 1.74 | 0.97 | 1.00 | 60.6 |
| 99 | none (raw frame) | 2 | 0.43 | 108.56 | 0.98 | 1.00 | 18.7 |
| | 100 | 59 | 0.24 | 6.98 | 0.95 | 1.00 | 50.0 |
| | 105.99 (exact pedestal) | 65 | 0.12 | 2.39 | 0.94 | 1.00 | 61.3 |
| | 105.99, sparsity 0 | 65 | 0.13 | 2.34 | 0.94 | 1.00 | 61.6 |
| | 105.99, sparsity $\times 0.1$ | 64 | 0.13 | 2.33 | 0.94 | 1.00 | 61.4 |

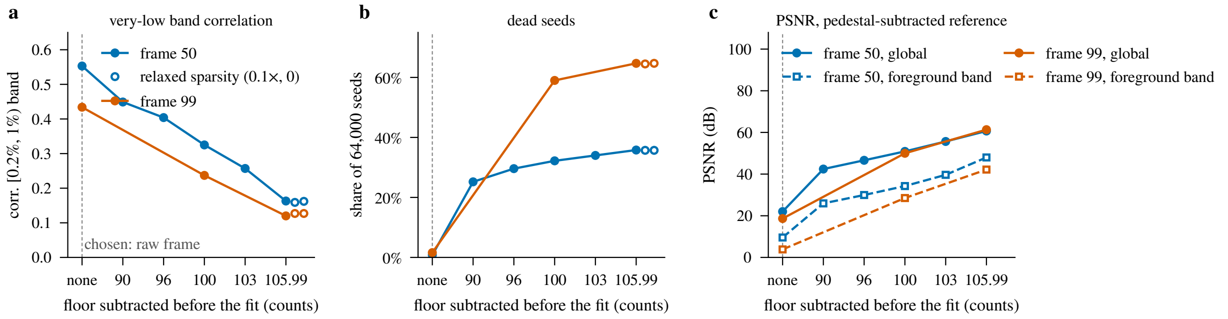


Figure 2: The floor sweep of Table 2 as curves: frames 50 and 99 of the membranes channel over the floor subtracted before the fit (none is the raw frame, the chosen recipe; 105.99 is the recorded pedestal). (a) Pearson correlation of the render with the frame in the very-low band, where the faint background lives: at frame 50 it falls from 0.55 on the raw frame to 0.16 at the exact pedestal. (b) The share of the 64,000 seeds whose amplitude died, which grows as more seeds land on clipped zeros. (c) Global PSNR (solid) and foreground-band PSNR (dashed) of every arm against the frame minus the recorded pedestal; on the raw-frame arm both measure the pedestal the fit reproduces, not its fidelity, which is why the text argues from (a) and (b). In (a) and (b) the hollow markers are the two relaxed-sparsity arms at the exact pedestal; in (c) the hollow squares are the foreground-band curve. The sweep left no images, so no crops are shown.

### 3 Time as a hard ordering barrier

In all three archives time is the fourth centre column, and a splat has no extent along it: $\sigma = 0$ along time, so a splat is instantaneous and cannot smear into the neighbouring frame. For the stored Cholesky factor this means that the $4 \times 4$ lower-triangular factor of a stacked splat is its $3 \times 3$ spatial factor padded with a fourth row of zeros, diagonal included, and no cross term between time and space. The decoded record of a splat is therefore 15 float32 values (four centres, one amplitude and the ten packed entries of the factor), 60 bytes; the compiled scenes store the same values quantised, the centres as four uint16 columns of 8 bytes per splat, which sets the pitch of the storage chunks in Figure 3. The payload a time step delivers to the renderer drops the time centre and the four zero entries of the padded row and keeps 10 values, the 40 bytes of Section 4.

The writer treats a stacked (categorical) centre column as an ordering *barrier dimension*: splats are sorted by time first and the Hilbert curve (*Morton and Hilbert ordering*) that orders them for the *spatial index* is computed over the three spatial columns only, so every timepoint's splats are contiguous. Two chunk grids lie over that order (Figure 3). The *index chunk* is the spatial index's *atom*, 1,024 splats with one stored bounding box; the *storage chunk* is the zarr chunk the file is compressed in, 1 MB in the published layout, a coarser grid of which every index chunk is a whole multiple. In the compiler's layout only the one index chunk at each timepoint boundary holds splats of two frames. In the compiled *Drosophila* scene, 1,995 of the 125,000 index chunks (over 4 rungs) hold two frames, at most one per boundary per rung, and 1,582 of 12,512 in the two neuromast nodes, counted over the compiled scenes' spatial-index chunks; the zebrafish scene, one part per timepoint, has none by construction. A 1 MB storage chunk can hold several neighbouring frames (Section 5). The additive ladder is built per timepoint for the same reason (the stacked axis is a hard coarsening barrier), so no rung blends two frames.

Two consequences follow. First, only one timepoint is ever resident: an nD node sliced on a hidden dimension commits only its current slice, so the *committed prefix (committed count)* on the GPU is one frame, about 2.4M, 256K and 128K splats for the three recordings (Table 3). The writer's warning that the *Drosophila* node holds more splats than one node can render on a 4096-class GPU (a GPU whose largest texture side is 4096 texels, the conservative floor the writer assumes) is about the whole archive and not about anything the viewer draws. Second, the cost of a time step does not grow with the recording's length. At a fixed per-frame content and chunk layout, stepping fetches the chunks whose bounds intersect the new slice (a 1 MB chunk that spans several timepoints brings its neighbours along), plus the first chunks of the adjacent slice that the viewer prefetches, and nothing from the rest of the recording, however many frames it holds.

### 4 Storage

Table 3 lists, per recording, the source bytes of the frames the archive represents, the published archive size, the resulting ratio, the splat counts per frame and the decoded parameter payload of one frame (float32 centres, amplitude and packed Cholesky factors: 40 bytes per splat, Section 3). That payload is what a time step delivers to the renderer, not the renderer's allocation: the viewer stores every splat as four RGBA32F texels plus two 32-bit ordering indices, 72 bytes per slot before padding (`rendering/gpu-buffer-pool/geometry-bytes.ts`), so the GPU footprint of a resident frame is 1.8× the payload. The archive sizes are the byte counts pinned in the software's dataset manifest for the published files.

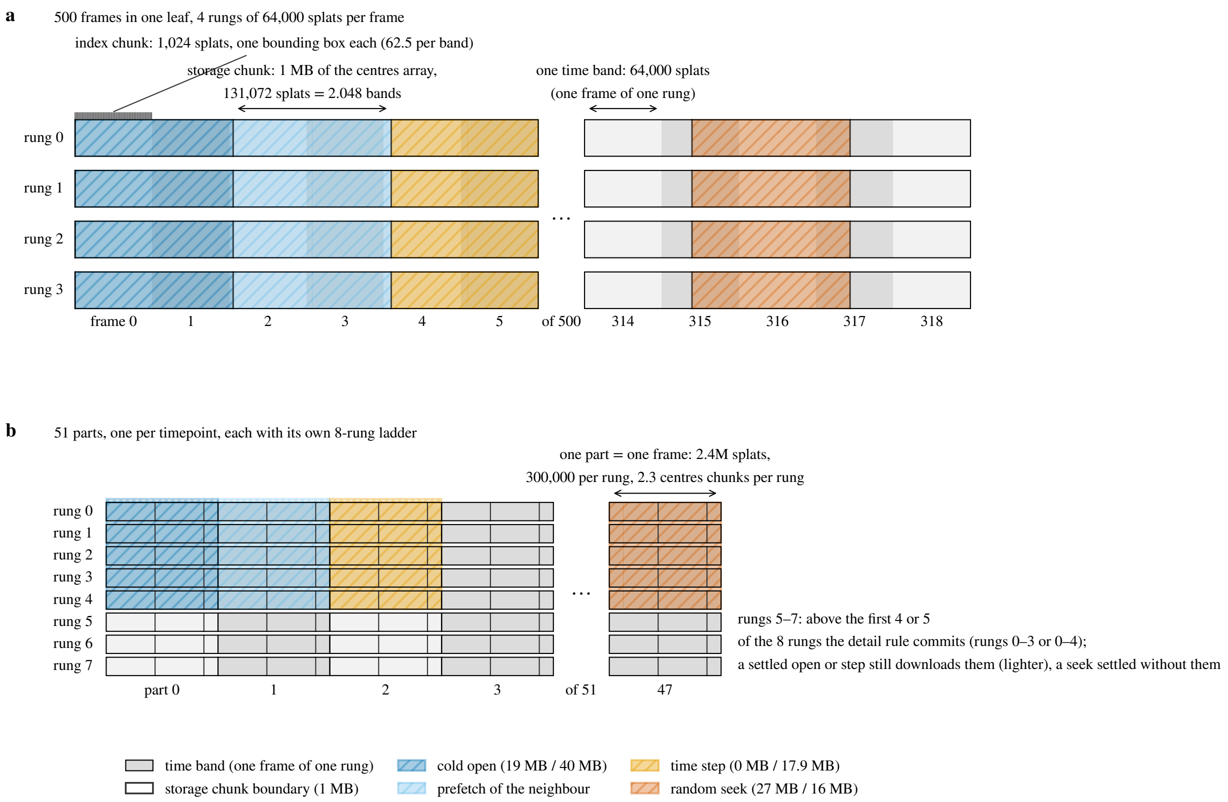


Figure 3: Schematic of the splat order and the two chunk grids, drawn with the counts of the compiled scenes; no measured curve is plotted. (a) The *Drosophila* node: one leaf whose splats are sorted by time first, one row per rung of its 4-rung ladder, so every frame is a contiguous band of 64,000 splats in each rung. Index chunks of 1,024 splats (the spatial index's atoms, ticks above rung 0) and 1 MB storage chunks of the centres array (131,072 splats of four uint16 columns, vertical lines) lie over that order. A storage chunk is 2.048 bands, so the cold open of frame 0 (blue) brings frame 1 and the head of frame 2 along and the viewer prefetches the next chunk (light blue); the first steps therefore move no chunk data, and about every second step needs one new centres chunk per rung (yellow, the step to frame 4), while the other arrays cross their own, coarser chunk boundaries. A random seek to a cold frame needs the one or two chunks that hold it in every rung (orange; the drawn target straddles a chunk boundary and needs two). (b) The zebrafish scene: one part per timepoint, each with its own eight-rung ladder of about 300,000 splats per rung, so a rung spans only about 2.3 chunks and nothing is shared between neighbours. A step loads one part: the detail rule commits its first 4 or 5 rungs (Section 5), and a settled open or step downloads the rungs above as well (lighter hatching), so its cost is the whole part; a random seek settled at the committed rungs. The legend gives the measured cost of each fetch for the two scenes: the cold open and the time step from the two-step measurement of Section 5, the random seek from Table 5.

### 5 Streaming

Two numbers describe what the viewer downloads: the cost of opening a frame cold and the cost of advancing one time step. We call their measurement the *two-step measurement*. Both are measured in a real browser (requests and bytes only, since timings depend on the network) on the scenes in the layout the demo site publishes: every store on the site goes through `luxar optimize --profile archive`, a 1 MB chunk target, before upload, so the measurement applies

Table 3: Three Gaussian-splat time-lapses. Raw bytes count the frames the archive represents, at the dtype each source is stored in (uint16 for the zebrafish and *Drosophila* stacks, float32 for the deconvolved neuromast), the same stored-byte basis as the single-volume tables of the main text. The zebrafish variant keeps every fifth frame of a 5.87 GB parent archive whose own ratio against all 253 source frames is 147×; all 253 frames were fitted. Splats per frame are the median over frames with the 5th–95th percentile range. Payload bytes are the decoded parameters of one frame at 40 bytes per splat, the median over the captured frames of Figure 4 of the main text. Requests per step are the measured range over single time steps in a real browser (the two-step measurement of Section 5).

| Recording | Raw (GB) | Archive (MB) | Ratio | Splats / frame median (p05–p95) | Payload (MB / frame) | Requests per step |
|---|---|---|---|---|---|---|
| Zebrafish embryo nuclei | 174.12 | 1112.4 | 157× | 2.4M (2.1M–2.6M) | 96.4 | 65 |
| *Drosophila* embryo nuclei | 77.68 | 1159.0 | 67× | 256K | 10.2 | 1 |
| Neuromast membranes + nuclei | 22.45 | 142.8 | 157× | 128K | 5.1 | 1 |

that pass to the compiled scene and serves the result from a local origin. The opening cost counts every request from a cold cache until the network has been idle for the settle window. The tracer waits a fixed interval (40 s) after the scene appears, then for network idle, then a second interval (15 s), and records requests and bytes only; it does not force the finest level of detail and does not record the drawn count. The opening cost is therefore the settled download of one timepoint at the viewer's default detail setting: not the first painted pixels, and not a guarantee that every ladder rung is resident.

The two recordings that keep all timepoints in one node pay at the opening and almost nothing per step. The *Drosophila* scene opens in 19.06 MB and 45 requests to show a timepoint whose own decoded parameters occupy about 10.2 MB, and the neuromast scene in 65.46 MB and 140 requests for both channels. A time step then costs 1 request on the former and 1 on the latter, in both cases a zero-byte conditional re-read of the scene's metadata document (the tracers record requests and bytes, not status codes), so the step moves no chunk data (0 and 0 MB). The amplification at the opening is the chunk size: with hundreds of frames in one node, a 1 MB chunk of any rung spans many timepoints, so drawing one frame downloads its neighbours as well, which is exactly why the next step is already resident.

The zebrafish scene is laid out the other way, one part per timepoint, and its opening load (39.87 MB, 480 requests) is the first part plus the parts of the neighbouring timepoints the viewer prefetches. A step then loads one per-timepoint part, 17.91–17.96 MB in 65 requests: at 2.4M splats a frame is far larger than any chunk and nothing of it can be shared with its neighbours.

The chunk size sets the trade (Figure 4). The same three scenes as the compiler writes them, at its default 64 KB target, open in 5.27, 40.58 and 3.71 MB (the *Drosophila* and neuromast openings shrink to roughly the bytes of one frame), but a time step then costs 266–268, 2247–2252 and 119–126 requests. A time-lapse is stepped far more often than it is opened, which is why the published layout accepts the larger opening.

**Reproducibility.** Every measurement in this document is taken on one build of each recording: the published archives, compiled into scenes by the demo code of one luxar build (recorded with

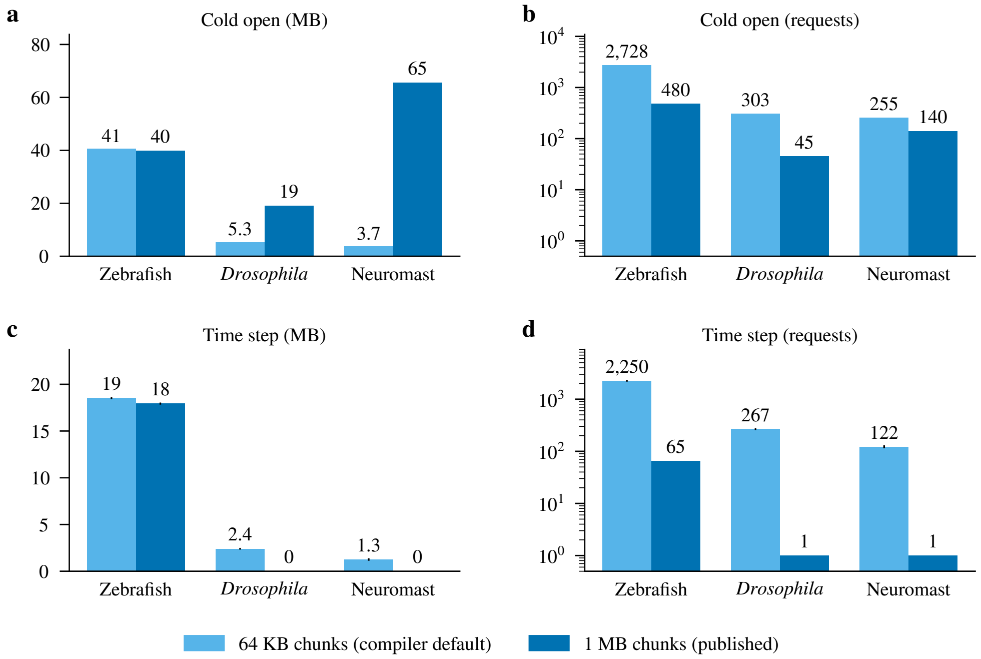


Figure 4: The chunk-size trade: the compiler's 64 KB target (light) against the published 1 MB layout (dark) for the three scenes, from the two-step measurement. (a, b) A cold open in MB and requests, the request axis logarithmic: the 1 MB layout opens the *Drosophila* and neuromast scenes at several times the bytes of one frame, because a chunk of any rung spans many timepoints. (c, d) One time step in MB and requests, the request axis again logarithmic: at 1 MB the same two scenes step in one request and no chunk data, at 64 KB in 266–268 and 119–126 requests; the zebrafish scene, one part per timepoint, moves the same bytes in either layout and only its request count changes. Thin black bars mark the range of the two measured steps.

the analysis), which fetches each checksum-pinned archive and grafts it without refitting. The archives are the Zenodo files `h2afva_51tp.gsplats.zarr.zip` (one leaf, 4 rungs)[2], `drosophila_embryogenesis_500tp.gsplats.zarr.zip` (one leaf, 4 rungs)[5] and the two neuromast zips (one leaf each, 8 rungs)[4]. The zebrafish node becomes 51 time parts with 408 rungs (Section 2); the *Drosophila* node keeps the archive's 4 rungs and the two neuromast nodes their 16. An archive and its scene hold the same splats; only the grouping, the ladders and the chunk layout differ.

The compile is reproducible at the level of chunk bytes. A second build of the neuromast scene from the same build on an Apple M4 Max laptop gives the same SHA-256 over the chunk bytes for 80 of 80 arrays of the compiled layout (13,328 chunks, 139.2 MB; 0 arrays differ). The root *`content_hash`* is not a portable build identity, because it also covers float64 encoder bounds in the array attributes whose reduction order is platform-dependent even when every quantised byte agrees. The per-array chunk digests of the three scenes in both layouts, the request traces and the per-frame scoring records are deposited with the analysis. The two-step measurement, Table 3, the chunk-span counts of Section 3 and the long trace below all describe those scenes (`gsplats_4d_h2afva_timelapse.luxar.zarr`, `gsplats_4d_drosophila_embryogenesis.luxar.zarr`, `gsplats_4d_neuromast_2ch.luxar.zarr`) in the published layout, served from the workstation that compiled them. The trace's sampled request URLs reach `additive_3` of the *Drosophila* node and `additive_7` of the neuromast nodes, the last rung of

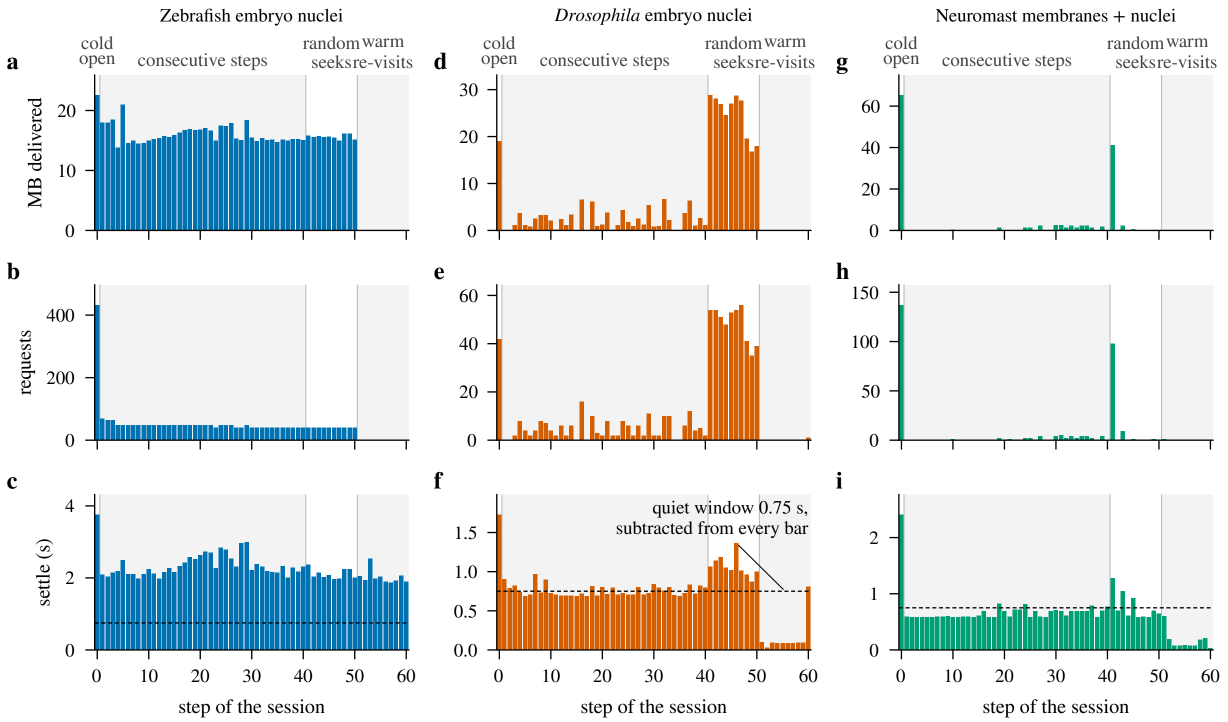


Figure 5: Every step of the long trace (Tables 4 and 5), one column per recording: MB delivered (top), requests (middle) and seconds to settle (bottom) for the cold open, the 40 consecutive steps, the 10 random seeks to unvisited frames and the 10 warm re-visits, the phases shaded. The zebrafish scene fetches one part on every step and every seek; the *Drosophila* scene fetches a few chunks on most steps and a cold frame on every seek; the neuromast opening download covers close to half of its archive, so little follows. The dashed line marks the tracer's 750 ms quiet window; the tracer subtracts it from every settle time it reports, so a bar below the line is a step whose data and drawn count stopped changing within that time of the step command.

each of those ladders; the trace's zebrafish steps stop short of a whole part (a median of 48 of a part's 64 data chunks, below).

**The long trace.** The long trace (Tables 4 and 5; every step of it in Figure 5) puts the two-step measurement in context and adds the timings the request counts leave out. From one cold open of each scene we stepped through 40 consecutive frames, jumped to 10 frames drawn at random (seed 42, without replacement) from those not yet visited and re-visited the first ten. The browser was Chromium headless, rendering on the workstation's GeForce RTX 3070 through ANGLE Vulkan (the compile and fitting box carries two cards; the browser took this one) and fetching from the same machine's `luxar serve` over its loopback interface (round trips of 2–4 ms and 320–1271 Mbit/s single-object throughput during the runs). The seconds below are those of a link with no wide-area latency and scale with the link.

Every statistic is recomputed from the per-step records: the median is the standard one (the mean of the two central values for an even count) and the 95th percentile interpolates linearly. A step is settled once no data response has arrived for 750 ms, the viewer reports no load in progress and the drawn count has held still for the same window.

A step here is a slice change while paused, which the viewer's automatic *detail rule* (Figure 6)

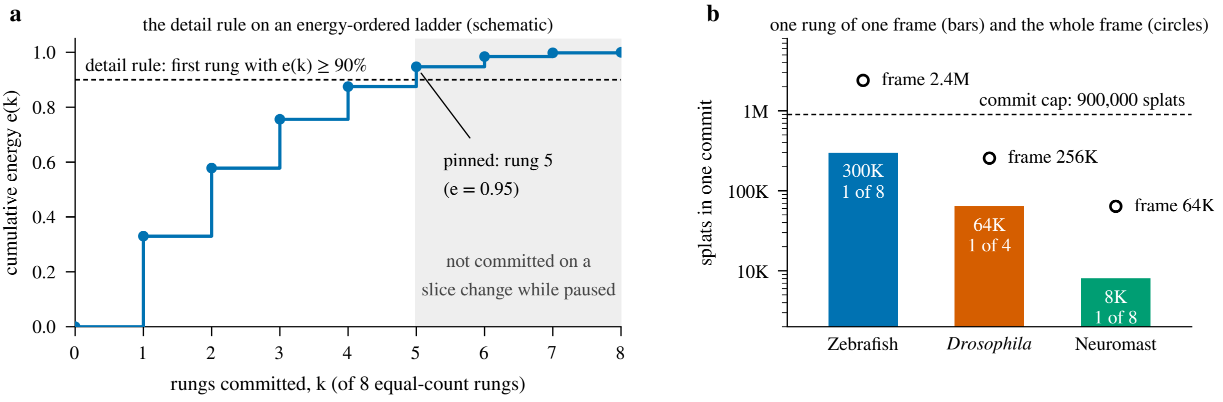


Figure 6: The detail rule and the commit cap, schematic. (a) Cumulative committed energy $e(k)$ after $k$ rungs of an eight-rung equal-count ladder ordered by self-energy, an idealised concave curve and not a measured stamp; the viewer's automatic detail setting pins the first rung whose $e(k)$ reaches 90% and commits nothing above it on a slice change while paused; the rungs above still download once the step settles (Section 5). On the zebrafish parts the trace shows that stop at rung 4–5 of 8. (b) One additive increment, a rung of one frame, for the three recordings (bars) against the 900,000-splat commit cap, the largest rung the viewer processes in one step, with the whole frame as a hollow marker: a zebrafish frame alone exceeds the cap, which is why a frame is laddered at all.

pins to the ladder rung that reaches 90% of the frame's energy, not to the full ladder. On the zebrafish parts that rule stops at 4–5 of the 8 equal-count rungs: the viewer reported a median 1.3M committed splats per frame, 50–63% of the 2.4M stored. The download does not stop at that rung. The two-step measurement above, which settles for longer, gives a zebrafish step of 17.91–17.96 MB in 65 requests, and those are the whole parts of frames 1 and 2: in the published layout every part holds 64 data chunks (8 per rung; the bounds arrays are fetched at the opening), and the chunk bytes of those two parts are 17.91 and 17.96 MB (the 51 parts span 17.9–24.6 MB). The trace's 750 ms quiet window closes earlier, after the committed rungs (a seek's 40 requests are the first five rungs at 8 chunks each), which is why its median step is 15.4 MB. The same settle rule makes the trace's zebrafish cold open (23 MB, 432 requests; Table 4) smaller than the two-step measurement's 39.87 MB, whose longer settle window also counts the neighbour-part prefetch. The four-rung *Drosophila* ladder is committed whole (256K of 256K per frame), as is the neuromast pair (128K).

Consecutive steps otherwise cost what the layout dictates. The zebrafish scene, one part per timepoint, fetched on every step: 48 requests (69 on the first, which also drew the neighbour prefetch) and 15.4 MB, taking 1.8 s to the last byte on this link (95th percentile 2.2 s, maximum 2.9 s). The *Drosophila* scene fetched on 34 of 40 steps, a median of 4 requests and 1.2 MB in 0.1 s (at most 6.7 MB), because a 1 MB chunk of any rung spans several of its frames and the remaining steps were already resident. The neuromast fetched on 13 of 40 steps, never more than 2.5 MB, because its opening download (65 MB) already covered close to half of its 142.8 MB archive.

A seek to an unvisited frame is a cold frame: 27.0 MB (0.8 s; up to 28.8 MB and 1.1 s) on the *Drosophila* recording, whose chunks then bring no cached neighbours, and 15.6 MB (1.7 s) on the zebrafish, whose parts are of similar size wherever the seek lands. On the neuromast only 4 of the

10 seeks needed any chunk, the first of them 41.2 MB for the frames the opening download had not covered.

Re-visiting a frame needed no chunk in any of the 30 warm steps (4 of them made one request, a zero-byte conditional re-read of the metadata document) and settled in a per-recording median of 80 (neuromast) to 1952 ms (zebrafish), at most 2528 ms. On the zebrafish that is the time to re-commit a 2.4M-splat frame from the client cache, with nothing downloaded. "One timepoint resident" therefore describes the GPU: on the CPU side the viewer keeps the frames it has seen in a budgeted cache pool. Where Chromium exposes its heap limit, the pool is 48% of `jsHeapSizeLimit` (the cache tiers take 60% of a 0.8 target heap usage; `cache/heap-budget.ts`); the fixed pools of 2 GiB, 1 GiB and 384 MiB are the fallback for a desktop-, laptop- or mobile-class device whose heap cannot be measured (WebKit), and `?cacheBudgetMB=` (MiB) overrides both. Chromium reported a limit of 4396 MB in these runs, a pool of about 2110 MB.

The client's memory grows to that budget plus the working set. The Chromium process tree peaked at 10.5, 2.8 and 2.1 GB of resident-set size during the zebrafish, *Drosophila* and neuromast traces (summed over Chromium's processes, so an upper bound on unique physical memory; 10.5, 2.7 and 2.0 GB at the last step), and `usedJSHeapSize`, which counts array-buffer backing stores, peaked at 8248, 1301 and 852 MB. The zebrafish run, whose decoded frames are 96.4 MB each, is the one whose working set outgrows the pool. The per-step request and byte counts are properties of the scenes' layout; the seconds are properties of this link and this card and scale with them.

#### 6 Fidelity

We measure the fidelity of the shipped archives to the raw frames. Five frames per recording are scored, spread over the whole time axis (Table 6); for the neuromast both channels are scored, so 20 frames are scored in all. For each frame, the scoring selects the timepoint in the shipped zip (the files named in Table 7), drops the time column, undoes the display scaling, renders it on the raw grid and scores it against the raw frame. It reads local copies whose bytes match the manifest pins, and every deposited score record carries the SHA-256 and byte count of the file it scored.

Two scores are reported, defined once here. With $R$ the range (maximum minus minimum) of the reference frame after the basis step described below and MSE the mean squared error over the whole frame,

$$\mathrm{PSNR} \;=\; 10\log_{10}\frac{R^2}{\mathrm{MSE}}, \qquad \mathrm{PSNR}_{\mathrm{fg}} \;=\; 10\log_{10}\frac{R^2\,N_{\mathrm{fg}}}{\mathrm{SSE}_{\mathrm{fg}}}, \tag{1}$$

where $N_{\mathrm{fg}}$ is the number of reference voxels above a 256-bin Otsu threshold of that frame and $\mathrm{SSE}_{\mathrm{fg}}$ their summed squared error. The foreground score thus replaces the whole-frame MSE by the mean squared error over those voxels while keeping the same whole-frame $R$ (`gsplats/metrics.py`, `compute_foreground_psnr`). The Otsu threshold is a contrast split of the frame, not the camera floor, and the share of voxels above it is listed beside each score. Global PSNR on a mostly empty light-sheet stack rewards predicting empty space, which is why both are given.

The basis step puts each frame on its recipe's basis before scoring. For the *Drosophila* the

Table 4: Stepped playback (slice changes while paused) of the three compiled scenes from one cold open: the cold open and 40 consecutive steps. Cold open: requests and MB until settled, seconds to the first drawn geometry and to settle. Per step: requests to the data host, MB delivered (chunks are stored compressed and served as-is, so this is the wire size) and seconds from the step command to the last data byte, the median on the recording's row and the (min–max) range beneath it; "fetched" is the number of steps that needed at least one chunk. Chromium headless rendering on the workstation's GeForce RTX 3070 (ANGLE Vulkan); the scenes are served by `luxar serve` on the same machine and fetched over its loopback interface (per-run RTT and throughput probes are stored with the trace).

| | Cold open | | | | Consecutive steps ($n = 40$) | | | |
|---|---|---|---|---|---|---|---|---|
| Recording | requests | MB | first geometry (s) | settled (s) | requests | MB | to last byte (s) | fetched |
| Zebrafish nuclei | 432 | 23 | 2.9 | 3.8 | 48 | 15.4 | 1.8 | 40 |
| | | | | | (40–69) | (13.8–20.9) | (1.6–2.9) | |
| *Drosophila* nuclei | 42 | 19 | 1.6 | 1.7 | 4 | 1.2 | 0.1 | 34 |
| | | | | | (0–16) | (0.0–6.7) | (0.0–0.8) | |
| Neuromast, 2 channels | 137 | 65 | 2.3 | 2.4 | 0 | 0.0 | 0.0 | 13 |
| | | | | | (0–5) | (0.0–2.5) | (0.0–0.7) | |

Table 5: The same runs, continued: 10 seeks to frames drawn at random (seed 42) from those not yet visited (drawn without replacement, so every seek is a cold frame), then a warm re-visit of the first 10 consecutive steps. Warm: requests per re-visit and seconds to settle. Peak memory: Chromium `usedJSHeapSize` (decimal MB; counts array-buffer backing stores) and the peak process-tree resident-set size sampled during the run (decimal GB, summed over Chromium's processes, an upper bound on unique physical memory).

| | Random seeks to unvisited frames ($n = 10$) | | | | Warm re-visits ($n = 10$) | | Peak memory | |
|---|---|---|---|---|---|---|---|---|
| Recording | requests | MB | to last byte (s) | fetched | requests | settle (s) | JS heap (MB) | RSS (GB) |
| Zebrafish nuclei | 40 | 15.6 | 1.7 | 10 | 0 | 1.95 | 8248 | 10.5 |
| | (40–41) | (15.0–16.1) | (1.6–1.9) | | (0–1) | (1.87–2.53) | | |
| *Drosophila* nuclei | 52 | 27.0 | 0.8 | 10 | 0 | 0.09 | 1301 | 2.8 |
| | (35–56) | (16.8–28.8) | (0.8–1.1) | | (0–1) | (0.03–0.81) | | |
| Neuromast, 2 channels | 0 | 0.0 | 0.0 | 4 | 0 | 0.08 | 852 | 2.1 |
| | (0–98) | (0.0–41.2) | (0.0–1.2) | | (0–1) | (0.03–0.60) | | |

Table 6: The frames scored per recording, spread over the whole time axis. The zebrafish variant stores every fifth source timepoint, so its frames are listed as variant frame (source timepoint).

| Recording | Frames scored |
|---|---|
| Zebrafish embryo nuclei | 0 (0), 12 (60), 25 (125), 37 (185), 50 (250) |
| *Drosophila* embryo nuclei | 0, 125, 250, 375, 499 |
| Neuromast membranes and nuclei | 0, 25, 50, 75, 99, both channels |

Table 7: Fidelity of the shipped archives (the zips the demo site serves: `h2afva_51tp.gsplats.zarr.zip`, `drosophila_embryogenesis_500tp.gsplats.zarr.zip`, `neuromast_membranes.gsplats.zarr.zip`, `neuromast_nuclei.gsplats.zarr.zip`) on sampled frames, scored against the raw frame as defined in Section 6 and Eq. 1. "fg PSNR" is the foreground score and "fg share" the fraction of voxels above the frame's Otsu threshold; the zebrafish "Frame" is the source timepoint. "Splats" is the archive's count at that frame.

| Recording | Frame | PSNR (dB) | fg PSNR (dB) | fg share (%) | Splats |
|---|---|---|---|---|---|
| Zebrafish embryo nuclei | 0 | 59.9 | 48.2 | 5.4 | 2,217,045 |
| | 60 | 55.2 | 36.9 | 1.2 | 2,183,179 |
| | 125 | 52.1 | 33.6 | 1.2 | 2,409,350 |
| | 185 | 44.8 | 28.0 | 1.8 | 2,448,113 |
| | 250 | 43.3 | 26.9 | 2.0 | 2,601,757 |
| *Drosophila* embryo nuclei | 0 | 39.5 | 34.0 | 20.7 | 256,000 |
| | 125 | 41.8 | 37.0 | 27.4 | 256,000 |
| | 250 | 42.1 | 37.6 | 29.2 | 256,000 |
| | 375 | 41.9 | 37.4 | 29.4 | 256,000 |
| | 499 | 48.7 | 44.2 | 31.3 | 256,000 |
| Neuromast membranes | 0 | 50.4 | 49.7 | 7.7 | 64,000 |
| | 25 | 49.4 | 49.9 | 6.9 | 64,000 |
| | 50 | 47.4 | 48.6 | 6.8 | 64,000 |
| | 75 | 47.0 | 47.9 | 7.3 | 64,000 |
| | 99 | 45.3 | 44.9 | 7.1 | 64,000 |
| Neuromast nuclei | 0 | 57.8 | 55.0 | 2.8 | 64,000 |
| | 25 | 56.4 | 53.8 | 2.7 | 64,000 |
| | 50 | 56.8 | 54.1 | 2.9 | 64,000 |
| | 75 | 53.3 | 50.1 | 2.9 | 64,000 |
| | 99 | 55.6 | 52.8 | 2.9 | 64,000 |

recipe's background floor of 8 is subtracted and clipped at 0, and the archive's frame index is its time value. The zebrafish frame is scored raw (its resolved floor is 0) against the source timepoint. The neuromast frame is scored raw with a least-squares gain, since the archive stores unit-normalised amplitudes. These are reconstruction scores of the shipped archive against its own raw frame. They are not the held-out blind-spot scores of Supplementary Document 2, and they are not on the normalised benchmark basis of the main text's single-volume tables. The references here are raw full frames of the stored dtype, uint16 for the zebrafish and *Drosophila* and float32 for the neuromast, with floors of 8, 0 and 0 counts and a per-frame gain for the neuromast, so their levels are not directly comparable with those tables. Every deposited per-frame score record carries the rounded PSNRs, *SSIM*, Otsu threshold and share, gain and floor, the mean squared error, the reference minimum, maximum and range, and the foreground error sum and voxel count. It also carries the SHA-256 and byte count of the zip it was scored on (the digest the demo manifest pins), so each dB of Table 7 is recomputable from its record alone through Eq. 1.

Scored this way (Table 7; the right column of Figure 7 plots the same scores), the *Drosophila* archive holds 39.5–48.7 dB (foreground 34.0–44.2 dB) over its five scored frames and the neuromast archives 45.3–57.8 dB across the two channels (foreground 44.9–55.0 dB). The zebrafish archive scores 43.3–59.9 dB (foreground 26.9–48.2 dB) against the raw uint16 frames, and the score falls along the time axis while the splat count per frame rises from 2.2M to 2.6M.

#### 7 Two channels, one coordinate space

The neuromast recording is the one case with two channels. Each channel is its own archive and its own scene node (`layer=true`), so the Layers panel gives per-channel visibility, display range, gamma and blending; but both nodes share the scene's four dimensions, so a time step moves both together and a cell's membrane and nucleus stay superimposed. The two archives were fitted with the same recipe from co-registered source stacks, so no cross-channel registration is applied at display time.

#### 8 Figure 4 of the main text

Figure 4 of the main text (Figure 7 here) shows the three recordings as three rows of six frames each, captured from the viewer on a GPU with the interface hidden, one time value per frame. The boxed region of every frame is magnified in a thin row beneath the frames (one box per recording, a quarter of the frame width, chosen as the window of highest local standard deviation averaged over the row's six frames, so it lands on structure and not on a saturated core; the *Drosophila* box is pinned to the midline). A side column per row holds three panels: a compression strip (raw, archive and decoded payload bytes as logarithmic bars); the splat count of every frame of the recording with the six shown frames marked (the deposited per-frame splat counts); and the PSNR of the shipped archive against the raw frame at the five scored frames of Table 7 (the neuromast channels in the layer colours).

Every frame is captured with bloom off (its glow is not data and blurs the frame edges and the magnified boxes) and with every level-of-detail group forced to its finest level. The shot is taken only once no load pass is in progress and the drawn splat count has held still for 8 s, so no frame shows a coarse substitutive level or a half-streamed additive ladder. The specimens drift in their recordings (the *Drosophila* embryo creeps up by about 2% of the frame over 250 min), so each row is composed through one window size, which keeps the six frames at the same pixel scale, placed per frame on the specimen's bounding box below the time label.

The time labels give elapsed time from the first frame shown. The frames are the stores' zero-based time values (Table 8), so the last labels read 15.3 h, 249.5 min and 9.2 h. The *Drosophila* store's time axis is in minutes (30-s frames). Neither the zebrafish nor the neuromast store records its interval, and the published description of the zebrafish recording[3] does not state it either, so both were read from the raw acquisition timestamps on the lab's cluster. The zebrafish recording (OpenSiMView multi-view microscope; its histone channel holds exactly the 253 timepoints of the fused store) was acquired every 220.0 s (ClearControl per-timepoint timestamps, 5th–95th percentile 219.2–220.8 s), so the 51-frame store, which keeps every fifth frame, advances 18.3 min per frame and spans 15.3 h. The neuromast iSIM TIFF headers give 5.6 min per frame (5 min 35 s; 9.2 h across the 100 frames).

Each capture's JSON sidecar records its camera and display state (`slice`, `pose`, `layerSet`, `exposure`, `bloom`, `lodFinest`). In all eighteen sidecars `layerSet` is empty and `exposure` unset: no per-capture display override was applied, each row shows the appearance baked into the demo store it was captured from, and the camera is the only per-row setting. Table 8 lists the frames,

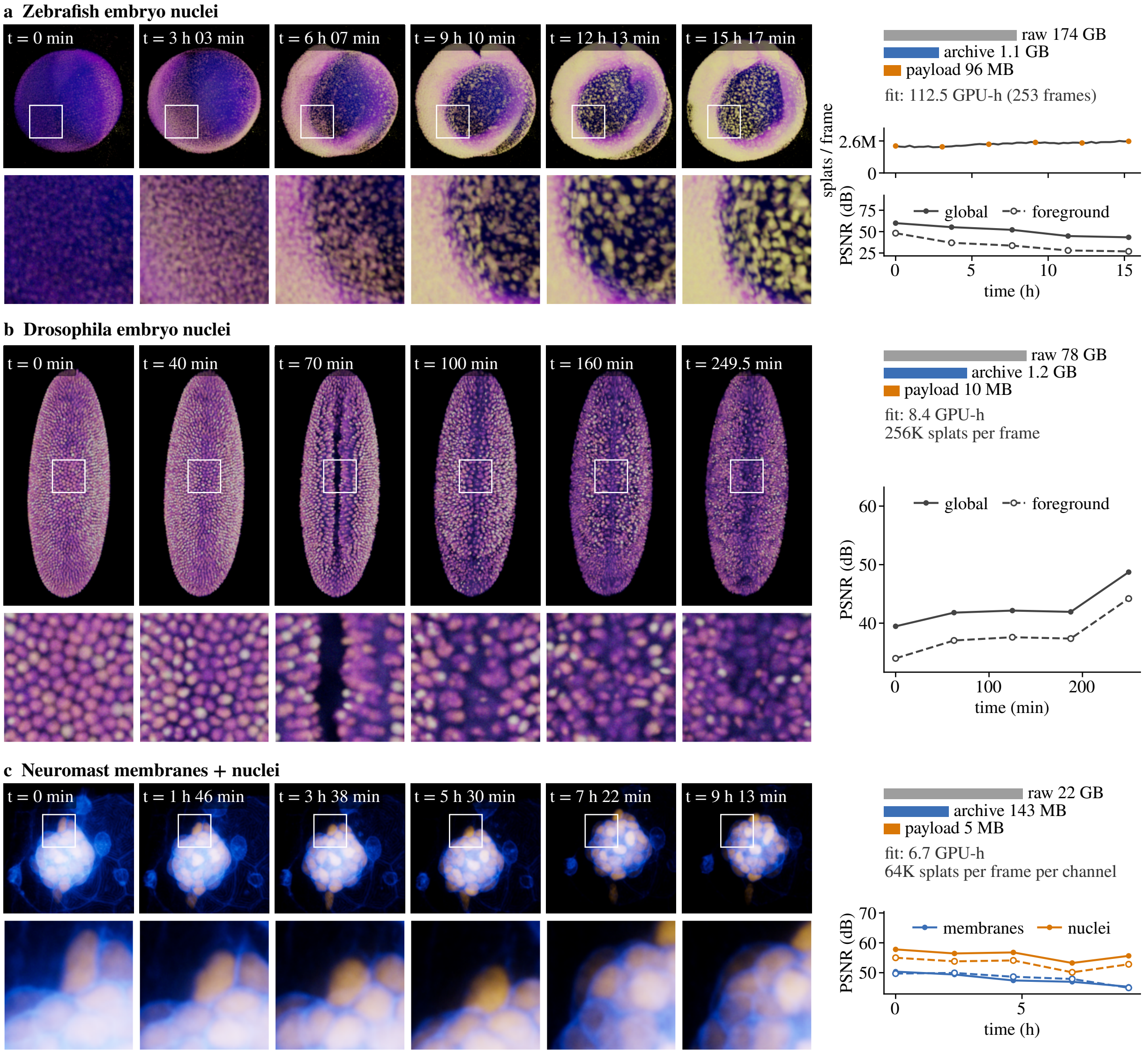


Figure 7: Figure 4 of the main text, set for this document: Gaussian-splat time-lapses. Rows: Zebrafish embryo nuclei (51 frames), *Drosophila* embryo nuclei (500 frames), Neuromast membranes and nuclei (100 frames, two channels). Columns: six frames per recording, each with its boxed region magnified in the thin row beneath (zero-based store frames 0–50, 0–249.5 min and 0–99; labels are elapsed time from the first frame shown). Right, per row: a strip of raw, archive and decoded payload bytes on a logarithmic scale, with the fit cost as elapsed hours of one RTX PRO 6000 (the zebrafish line counts all 253 fitted frames; Section 2 gives each row's basis) and, where every frame holds the same count, the splats per frame; for the zebrafish, whose count varies, the splat count of every frame of the recording with the six shown frames marked; and the fidelity of the shipped archive against the raw frame at the five scored frames of Table 7, its global PSNR (solid) and foreground PSNR (dashed, hollow markers), the neuromast channels in the layer colours. Requests per time step and per first paint are in Table 3 and Section 5.

Table 8: Frames, camera and baked appearance of the three rows of Figure 4. Frames are zero-based store time values (zebrafish: store frame, source timepoint in parentheses; *Drosophila*: minutes). The camera is the only per-row capture setting; window, gamma, opacity, absorption, blending and colour map are the appearance baked into each demo store.

| Row | Frames | Camera | Appearance |
|---|---|---|---|
| Zebrafish embryo nuclei | 0 (0), 10 (50), 20 (100), 30 (150), 40 (200), 50 (250) | side view: azimuth 300°, elevation −30° from the default pose, zoom 0.68, half-turn roll | window 0.001–0.025, gamma 2.2, opacity 0.55, absorption 1.09, volumetric blending, plasma colour map |
| *Drosophila* embryo nuclei | 0, 40, 70, 100, 160, 249.5 min | portrait 700 × 1400, zoom 0.7, azimuth 90° | window 0.059–0.868, gamma 1.0, opacity 0.20, absorption 1.19, volumetric blending |
| Neuromast membranes | 0, 19, 39, 59, 79, 99 | face-on along the imaging axis at the default distance | window 0–0.101, gamma 0.71, opacity 0.5; additive blending at exposure −3.4 stops |
| Neuromast nuclei | 0, 19, 39, 59, 79, 99 | as above | window 0–0.05, gamma 0.69, opacity 0.47; additive blending at exposure −3.4 stops |

the camera and the baked appearance per row. The neuromast is viewed face-on along its imaging axis, which the scene maps onto the viewer's $x$ axis, from an explicit camera at the viewer's default distance (the demo's own opening camera sits 37.5° off that axis); its axial step is the recording's measured 2.308 lateral pixels (0.25 / 0.1083 $\mu$m), so the row is not stretched along $z$. The zebrafish capture is turned by a half turn in the image plane, a camera roll, so the tail bud points up.

### 9 Summary and practical rules

**What to set.**

- **Fit per timepoint** with the single-volume fitter and stack the frames along a fourth centre column; the stacked column gets $\sigma = 0$, so a splat belongs to one frame and the decoded record grows from 40 to 60 bytes per splat without changing what a frame delivers.
- **Keep time a hard barrier**: sort by time first, order the spatial index within a frame, and build the additive ladder per timepoint, so that a time step touches nothing of the other frames.
- **Choose the chunk size for the step, not the opening.** At 1 MB chunks a step costs 1 request on the *Drosophila* scene and 1 on the neuromast; at the compiler's 64 KB it costs 266–268 and 119–126 requests. The price is an opening of 19.06 and 65.46 MB instead of 5.27 and 3.71 MB.
- **One part per timepoint** when a frame is far larger than a chunk, as for the 2.4M-splat zebrafish frames; a step then loads one part (17.91–17.96 MB) and nothing can be shared with the neighbours anyway.

- **A discrete time axis with a binary-exact step**, so every frame lands on the slider's grid.
- **Fit the raw frames** when a camera pedestal underlies faint structure; an exact floor subtraction kills the seeds that land on clipped zeros (36–65% of them on the neuromast) and loses the very-low band.
- **Do not cull the archive** at a per-frame budget of this size: a 95% cull cost 9.7 dB on the *Drosophila* frame. Take the budget from the blind-spot sweep where one exists, and state it as a shared choice where it does not.

**What it costs.** Fitting is the expensive step: 112.5, 8.4 and 6.7 hours of one RTX PRO 6000 for the three recordings. The archives are 1112.4, 1159.0 and 142.8 MB against 174.12, 77.68 and 22.45 GB of source voxels. A resident frame occupies 1.8× its payload on the GPU, and the client keeps the frames it has seen in a cache pool of about 2110 MB in these runs, so a long session on the zebrafish scene reached 10.5 GB of resident-set size. On a loopback link a zebrafish step took 1.8 s to the last byte and a *Drosophila* step 0.1 s; those seconds scale with the link.

**What was not measured.** The timings are those of one workstation and its loopback interface; no wide-area link was traced. Every step is a slice change while paused, so continuous playback, which the viewer's prefetcher serves, was not timed. One browser and one GPU were used. The neuromast budget of 64,000 splats per frame was not calibrated per channel on the raw frames. The shipped zebrafish archive's fitter configuration is not recorded in its metadata. The seams of the content tiling that fitted the zebrafish recording were not measured (Supplementary Document 11).

[3] Merlin Lange, Alejandro Granados, Shruthi VijayKumar, Jordao Bragantini, Sarah Ancheta, Yang-Joon Kim, Sreejith Santhosh, Michael Borja, Hirofumi Kobayashi, Erin McGeever, Ahmet Can Solak, Bin Yang, Xiang Zhao, Yang Liu, Angela M. Detweiler, Sheryl Paul, Ilan Theodoro, Honey Mekonen, Chris Charlton, Tiger Lao, Rachel Banks, Sheng Xiao, Adrian Jacobo, Keir Balla, Kyle Awayan, Samuel D'Souza, Robert Haase, Alexandre Dizeux, Olivier Pourquie, Rafael Gomez-Sjoberg, Greg Huber, Mattia Serra, Norma Neff, Angela Oliveira Pisco, and Loïc A. Royer. A multimodal zebrafish developmental atlas reveals the state-transition

dynamics of late-vertebrate pluripotent axial progenitors. *Cell*, 187(23):6742–6759, 2024. doi: 10.1016/j.cell.2024.09.047.

[4] Loïc A. Royer. Luxar demo datasets: permissively licensed (CC-BY, CC0, public domain), 2026. URL https://zenodo.org/records/22804363. Zenodo. Version DOI 10.5281/zenodo.22804363; concept DOI 10.5281/zenodo.21912279 resolves to the latest version.

[5] Loïc A. Royer. *Drosophila melanogaster* embryogenesis: a 500-timepoint light-sheet timelapse as Gaussian splats, 2026. URL https://zenodo.org/records/22804386. Zenodo. Version DOI 10.5281/zenodo.22804386; concept DOI 10.5281/zenodo.22118694.

[6] Loïc A. Royer, William C. Lemon, Raghav K. Chhetri, Yinan Wan, Michael Coleman, Eugene W. Myers, and Philipp J. Keller. Adaptive light-sheet microscopy for long-term, high-resolution imaging in living organisms. *Nature Biotechnology*, 34(12):1267–1278, 2016. doi: 10.1038/nbt.3708.

[7] Raju Tomer, Khaled Khairy, Fernando Amat, and Philipp J. Keller. Quantitative high-speed imaging of entire developing embryos with simultaneous multiview light-sheet microscopy. *Nature Methods*, 9(7):755–763, 2012. doi: 10.1038/nmeth.2062.
